# Development of selective pyrido[2,3-*d*]pyrimidin-7(8*H*)-one-based Mammalian STE20-like (MST3/4) kinase inhibitors

**DOI:** 10.1101/2023.11.24.568596

**Authors:** Marcel Rak, Amelie Menge, Roberta Tesch, Lena M. Berger, Dimitrios-Ilias Balourdas, Ekaterina Shevchenko, Andreas Krämer, Lewis Elson, Benedict-Tilman Berger, Ismahan Abdi, Laurenz M. Wahl, Antti Poso, Astrid Kaiser, Thomas Hanke, Thales Kronenberger, Andreas C. Joerger, Susanne Müller, Stefan Knapp

## Abstract

Mammalian STE20-like (MST) kinases 1-4 play key roles in regulating the Hippo and autophagy pathways, and their dysregulation has been implicated in cancer development. In contrast to the well-studied MST1/2, the roles of MST3/4 are less clear, in part due to the lack of potent and selective MST3/4 inhibitors. Here, we re-evaluated literature compounds, and used structure-guided design to optimize the p21-activated kinase (PAK) inhibitor G-5555 (**8**) to selectively target MST3/4. These efforts resulted in the development of MR24 (**24**) and MR30 (**27**) with good kinome-wide selectivity, high potency for MST3/4, and selectivity towards the closely related MST1/2. In combination with the MST1/2 inhibitor PF-06447475 (**2**) the two MST3/4 inhibitors can be used to elucidate the multiple roles of MST kinases in cells. We found that MST3/4-selective inhibition caused a cell cycle arrest in the G1 phase, while MST1/2 inhibition resulted in accumulation of cells in the G2/M phase. These data point to distinct functions of these closely related kinases, which can now be addressed with subfamily-selective chemical tool compounds.

## INTRODUCTION

Mammalian STE20-like (MST) kinases consist of five members: MST1 (STK4), MST2 (STK3), MST3 (STK24), MST4 (STK26), and YSK1 (STK25). MST1 and MST2 have been assigned to the germinal center kinase II (GCKII) subfamily, whereas MST3, MST4, and YSK1 (yeast Sps1/Ste20-related kinase 1) are members of the GCKIII subfamily.^1^ Although GCKs share a high sequence similarity, they may differ in their function and the mechanisms they regulate in cellular signaling. MST1/2 are well-studied regulators of the Hippo pathway, an important signaling pathway that controls organ size and, when dysregulated, causes the development of diverse diseases, including breast, liver, and lung cancers.^2-4^ In canonical Hippo signaling, activation of MST1/2 is initiated either by caspase cleavage,^1^ or by dimerization with other SARAH domain containing proteins.^5^ MST1/2 regulate apoptotic gene expression through phosphorylation of large tumor suppressor homolog 1/2 (LATS1/2) and MPS1 binder 1 A/B (MOB1A/B),^3,6^ which mediate activation of the downstream transcriptional coactivator YES-associated protein (YAP) and the transcriptional coactivator with PDZ-domain motif (TAZ).^6,7^ MST1/2 are also key regulators of the autophagy pathway by directly phosphorylating microtubule-associated proteins 1A/1B light chain 3B (LC3B) and beclin-1 (BECN1), which are important mediators of autophagy progression.^8^

In contrast, MST3 and MST4 have been studied less intensively. MST3 has also been suggested to regulate the Hippo pathway as well as cell-cycle progression by phosphorylating the LATS1/2 homologs nuclear Dbf2-related kinase 1/2 (NDR1/2),^9,10^ indicating potential applications in anti-cancer drug development.^11-15^ However, MST3 may also play a role in mediating autophagosome formation by controlling the intracellular translocation of BECN1 upon starvation and presumably by regulating NDR1/2 activity.^16,17^ MST4 has been linked to direct YAP phosphorylation, a post-translational modification associated with tumor progression,^7^ and has been shown to stimulate autophagy in glioblastoma through phosphorylation-dependent activation of cysteine protease and autophagy regulator ATG4B.^18^ Depletion of both MST3 and MST4 reduce YAP phosphorylation and phosphorylation of autophagosomal LC3-II.^19^ Notably, YSK1 has not yet been functionally characterized.^20^

However, especially the upstream regulators of each of the GCKIII-subfamily kinases in the Hippo and autophagy pathways and the dysregulation of these pathways in disease remain elusive, in part due to the lack of selective chemical probes that would allow to study them in cellular systems. Therefore, there is an urgent need for the development of highly potent and selective chemical tools that allow in-depth functional characterization of MST-family kinases. Our comprehensive analysis of published MST kinase inhibitors revealed that most of the available MST1/2 inhibitors lack selectivity, leaving only few suitable MST1/2 chemical tool compounds, and no inhibitors have been reported that selectively target MST3 or MST4. However, we have recently developed a highly selective MST3 inhibitor using a macrocyclization strategy.^21^ In the study presented here, we used a structure-based approach to shift the selectivity of the p21-activated kinase 1 (PAK1) inhibitor G-5555 (**8**) towards kinases of the GCKIII subfamily,^22^ eliminating the off-target activity for salt-inducible kinases 1-3 (SIK1-3) and other CAMK and STE family kinases. We studied the effects of the developed MST3/4-selective lead compounds on cell viability and cell-cycle progression and report their comprehensive characterization as chemogenomic compounds for these two interesting kinases.

## RESULTS AND DISCUSSION

The starting point of our study was a systematic review of all kinase inhibitors that have been reported to be potent inhibitors of MST kinases.^19,23,24^ We evaluated these compounds against defined criteria that we established for so-called chemogenomic compounds (CGCs). CGCs are typically less selective than chemical probes. However, they still have very narrow selectivity profiles, and several CGC compounds targeting the same kinase ideally have non-overlapping off-targets.^25,26^ This enables the assignment of phenotypic responses to a particular target or to closely related isoforms by comparing data measured using the entire compound set.^27^ The criteria we chose for our evaluation included selectivity criteria (selectivity screening data against at least 100 protein kinases with a selectivity score S(10) ≤ 0.025 or a Gini score ≥ 0.6 at a concentration of 1 µM and preferentially less than 10 off-targets with an IC50 < 1 µM outside the MST subfamily) and potency criteria (*in vitro* potency < 100 nM (IC50), and a cellular potency of less than 1 µM).^27,28^ The structures and on-target potencies as well as the available kinome-wide selectivity data of the published compounds are compiled in Figure 1A and B, respectively. The compounds were classified as MST1/2, MST3/4, and pan-MST inhibitors according to their on-target activity within the MST subfamily.

**Figure 1.**
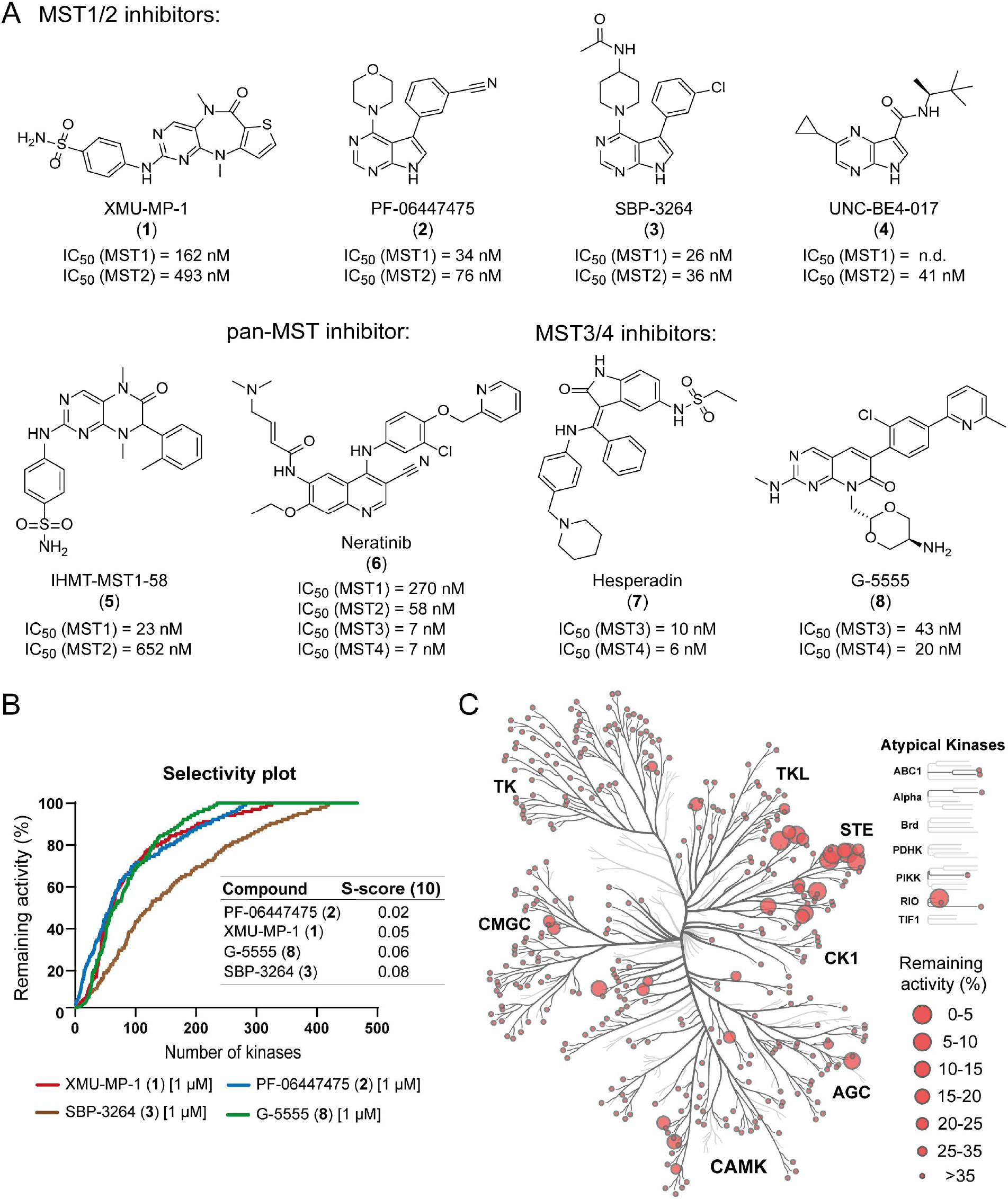
Selectivity profiles of published MST inhibitors. (A) Chemical structures of published MST inhibitors grouped by their isoform-specific inhibition. Published *in vitr*o potencies (IC50) against their main targets are shown. (B) Comparison of the published kinome-wide selectivity profiles (scanMAX, *Eurofins Discovery*) of compounds **1**, **2**, **3**, and **8**, shown as remaining kinase activity (%) in the presence of the corresponding inhibitor compared with the activity in its absence.^29,32,40,43^ The selectivity score of the compounds represents the number of kinases with a remaining activity lower than 10% divided by the total number of kinases tested. (C) Published selectivity profile of compound **2** which has been screened against 451 kinases (scanMAX, *Eurofins Discovery*), including kinase mutants, at a concentration of 1 µM.^32^ Selectivity data are mapped onto the phylogenetic tree, illustrated using the web-based tool Coral.^44^ Kinases tested are shown as dark gray branches and kinases inhibited by red cycles. The circle radius corresponds to the remaining activity in percent (%) compared with the kinase activity in the absence of the inhibitor.

The first inhibitor reported to selectively target MST1/2 was XMU-MP-1 (**1**).^43^ However, its *in vitro* activity on MST1/2 was not confirmed in a cellular context.^29^ In addition, XMU-MP-1 (**1**) exhibits prominent ULK1/2 and Aurora kinase activity, which is likely to cause toxicity.^29^ Due to the key role of ULK1/2 in autophagy and Aurora kinases in mitosis,^30,31^ respectively, these off-target activities would complicate the analysis of the phenotypic effect of MST kinases in these pathways.

PF-06447475 (**2**) was developed targeting the leucine-rich repeat kinase 2 (LRRK2) (IC50 = 3 nM).^32^ **2** was tested against 451 kinases at a concentration of 1 µM, revealing that it significantly inhibited only 8 kinases up to a remaining activity of 10%, resulting in a selectivity score of S(10) = 0.02 (Figure 1C). Activity on MST kinases has been reported with cellular EC50 values of 154 nM (MST1) and 197 nM (MST2).^29^ Further optimization of **2** with the goal to remove its LRRK2 activity, resulted in SBP-3264 (**3**) which has cellular EC50 values against MST1/2 of 139 nM and 122 nM, respectively.^29^ However, a lower kinome-wide selectivity has been described for **3** than for **2** (Figure S1).

The pyrrolopyrazine derivative UNC-BE4-017 (**4**) was developed to target JAK kinases,^33^ but this inhibitor also has high *in vitro* activity against MST2. The compound was resynthesized and further characterized by Schirmer *et al.* in 2022.^34^ Increased kinome-wide selectivity was observed with a selectivity score S(10) of 0.03 when tested against 468 kinases at a concentration of 1 µM. Unfortunately, the *in vitro* on-target potency was not confirmed in cells where UNC-BE4-017 had only an EC50 value of 7.6 µM (MST2).^34^

Recently, Wu *et al.* optimized XMU-MP-1 (**1**), resulting in the inhibitor IHMT-MST1-58 (**5**), which exhibited a 28-fold selectivity for MST1 over MST2.^35^ However, the reported selectivity data were inconclusive, and further evaluation of the selectivity of this inhibitor would be required in order to use it in mechanistic assays.

In summary, based on the data reported to date, PF-06447475 (**2**) appears to be the most promising compound among the MST1/2 kinase inhibitors evaluated as a candidate in a set of chemogenomic compounds. The potent LRRK2 activity of **2** could be controlled using highly selective LRRK2 chemical probes such as MLi2.^36^

Potential starting points for the development of selective MST inhibitors could be based on approved kinase inhibitors such as neratinib (**6**), a covalent ATP-competitive dual inhibitor of HER2/EGF approved for the treatment of breast cancer,^23,37^ but this compound has also been shown to be able to non-covalently inhibit several other kinases, including all four MST kinases.^24,37^

No selective inhibitor has yet been reported for the understudied kinases MST3 and MST4. The Aurora kinase inhibitor hesperadin (**7**) has been described to also inhibit MST3 and MST4.^19,38,39^ However, the selectivity of hesperadin (**7**) was only evaluated against 25 kinases at a concentration of 1 µM (Figure S2), and a more comprehensive assessment of its selectivity would be needed to evaluate the use of this inhibitor even as a chemical starting point. Intriguingly, the pyrido[2,3-*d*]pyrimidin-7(8*H*)-one based PAK1 inhibitor G-5555 (**8**), developed by Ndubaku *et al.*,^22^ is an inhibitor with favorable kinome-wide selectivity with a selectivity score S(10) of 0.06, mainly targeting kinases of the STE20 (PAK1, PAK2, MST3, and MST4) and of the CAMK group (SIK1–3 and NUAK2).^40^ Although the number of off-targets of G-5555 (**8**) is too large to consider it a tool compound for mechanistic studies, we chose this compound as the lead structure for the development of an MST3/4-selective inhibitor based on its strong potency for these two kinases and the significant SAR that we established developing this scaffold into a selective chemical probe targeting SIK kinases.^41,42^ Here we describe a new SAR, leading to the MST3/4-selective inhibitors MR24 (**24**) and MR30 (**27**).

### Cellular potency of G-5555 (8) on MST3/4

The *in vitro* activity of G-5555 (**8**) against recombinant MST3 and MST4 has been reported with IC50 values of 43 nM and 20 nM, respectively.^22^ However, the cellular binding may differ from enzyme kinetic inhibition data due to compound membrane penetration potentials, high concentration of cellular co-factors such as ATP or protein and domain interactions of the target kinase. Therefore, we evaluated the cellular on-target binding of G-5555 (**8**) on all four MST kinases using NanoBRET target engagement assays. In order to assess potential differences related to membrane penetration, we performed NanoBRET assays in intact cells as well as in cells that were permeabilized using digitonin. Since data analysis of measurements in intact cells was challenging for MST3, only data performed in permeabilized cells is shown for this target and G-5555 (**8**) as well as other weaker inhibitors. Consistent with the lack of *in vitro* activity of **8**, only weak binding was observed for MST1 and MST2, with EC50 values of 30.4 ± 8.3 µM and 4.6 ± µM, respectively. Surprisingly, the excellent potency of G-5555 (**8**) for MST3 and MST4 in enzyme kinetic assays was not confirmed in cell-based target engagement assays, with EC50 values of only 9.4 ± 6.7 µM against MST4 and 1.7 ± 0.1 µM against MST3 in permeabilized/lysed cells (Figure S3).

### Structure-based optimization of G-5555 (8)

To establish a rational design strategy for the development of selective MST3/4 ligands, the structurally best-characterized targets of G-5555 (**8**), MST3, PAK1, and SIK2, were chosen. As no crystal structure of any of the SIK kinases has been published, we used the previously established homology model of SIK2 based on the canonical binding mode of G-5555 (**8**) in MST4.^42^ We have previously shown that targeting the back pocket of the kinases is not beneficial for shifting the selectivity towards MSTs.^40^ With this in mind, we focused our analysis on the region around the activation loop, which contains the DFG motif and the catalytic loop of the kinases (Figure 2A), as well as the front-pocket region (Figure 2B).

**Figure 2.**
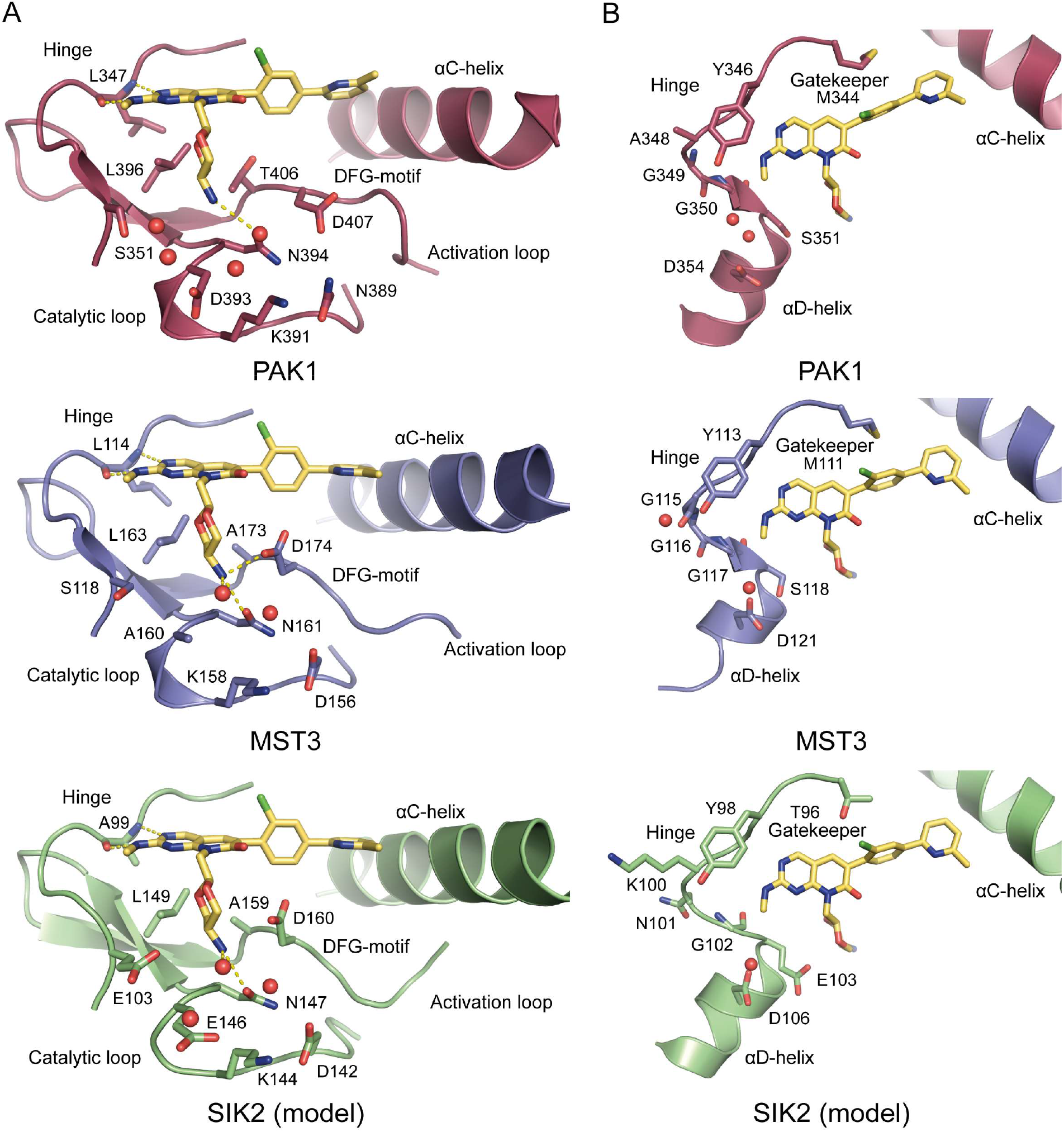
Structural comparison of PAK1 (PDB code 5DEY; red), MST3 (PDB code 7B30; blue), and SIK2 (homology model; green) in complex with G-5555 (**8**). Important amino acids and structural motifs are highlighted, with water molecules shown as red spheres and hydrogen bonds as dashed yellow lines. (A) Comparison of the amino acids located in the activation and catalytic loops of the three kinases (PAK1, MST3, and SIK2) around the (2*r*,5*r*)-2-methyl-1,3-dioxan-5-amine group of G-5555 (**8**). (B) Comparison of the amino acids lining the hinge and αD-helix regions of PAK1, MST3, and SIK2.

For all three kinases, the canonical binding mode of G-5555 (**8**) was characterized by hydrogen bonding to the hinge region amino acids Leu347 (PAK1), Leu114 (MST3), and Ala99 (SIK2). The regions around the A- and C-loops of the three kinases consist mainly of polar amino acids. Hydrogen-bond formation of G-5555 (**8**) was observed with a conserved asparagine in the C-loop of PAK1 (Asn394), MST3 (Asn161), and SIK2 (Asn147). While PAK1 (Asp393) and SIK2 (Glu146) share a negatively charged amino acid next to the conserved asparagine (−1 position), an alanine residue (Ala160) is found at the equivalent position in MST3. In addition, in the co-crystal structure of G-5555 (**8**) with MST3, the (2*r*,5*r*)-2-methyl-1,3-dioxan-5-amine group forms hydrogen bonds with the aspartate residue (Asp174) of the DFG motif and a water molecule. The adjacent −1 residue of the DFG motif differs in the three kinases studied. PAK1 harbors a threonine residue (Thr406), whereas MST3 and SIK2 share an alanine (Ala173 and Ala159, respectively).

The major difference in the hinge region of the three kinases is the gatekeeper amino acid: Met344 and Met111 in PAK1 and MST3, respectively, and a threonine (Thr96) in SIK2. However, targeting this difference appears not sufficient to shift the selectivity of the lead structure towards MST kinases, based on previous studies.^40^ The solvent-exposed front pocket was found to be more polar overall in SIK2 than in PAK1 and MST3. Adjacent to the amino acid of the hinge region forming hydrogen bonds with G-5555 (**8**), there is an alanine (Ala348) followed by two glycine residues (Gly349 and Gly350) in the structure of PAK1. MST3 harbors three glycine residues (Gly115, Gly116, and Gly117) at this position. In contrast, SIK2 has a lysine (Lys100) at position +1 of the hinge-binding amino acid, followed by an asparagine (Asn101) and a glycine (Gly102). In addition, the −1 residue of the αD-helix differs, with PAK1 and MST3 sharing a serine (Ser351 and Ser118, respectively) and SIK2 having a glutamate residue (Glu103) instead. All three kinases share an aspartate at the +3 position of the αD-helix.

In conclusion, optimizing the selectivity of the lead structure by targeting the region around the A-loop/C-loop appeared to be challenging as no key differences were identified that could be exploited. Targeting this region could, however, improve the binding affinity by extending the hydrogen-bond network with the MST kinases. In contrast, optimization of the front-pocket targeting group of G-5555 (**8**) was judged to be more promising for improving selectivity over SIK kinases as this would allow exploiting key amino acid differences in the overall more hydrophobic front pocket of MST3. Therefore, we first focused on modifying the (2*r*,5*r*)-2-methyl-1,3-dioxan-5-amine group of G-5555 (**8**), followed by derivatization of the methylamine group in the front-pocket region (Figure 3).

**Figure 3.**
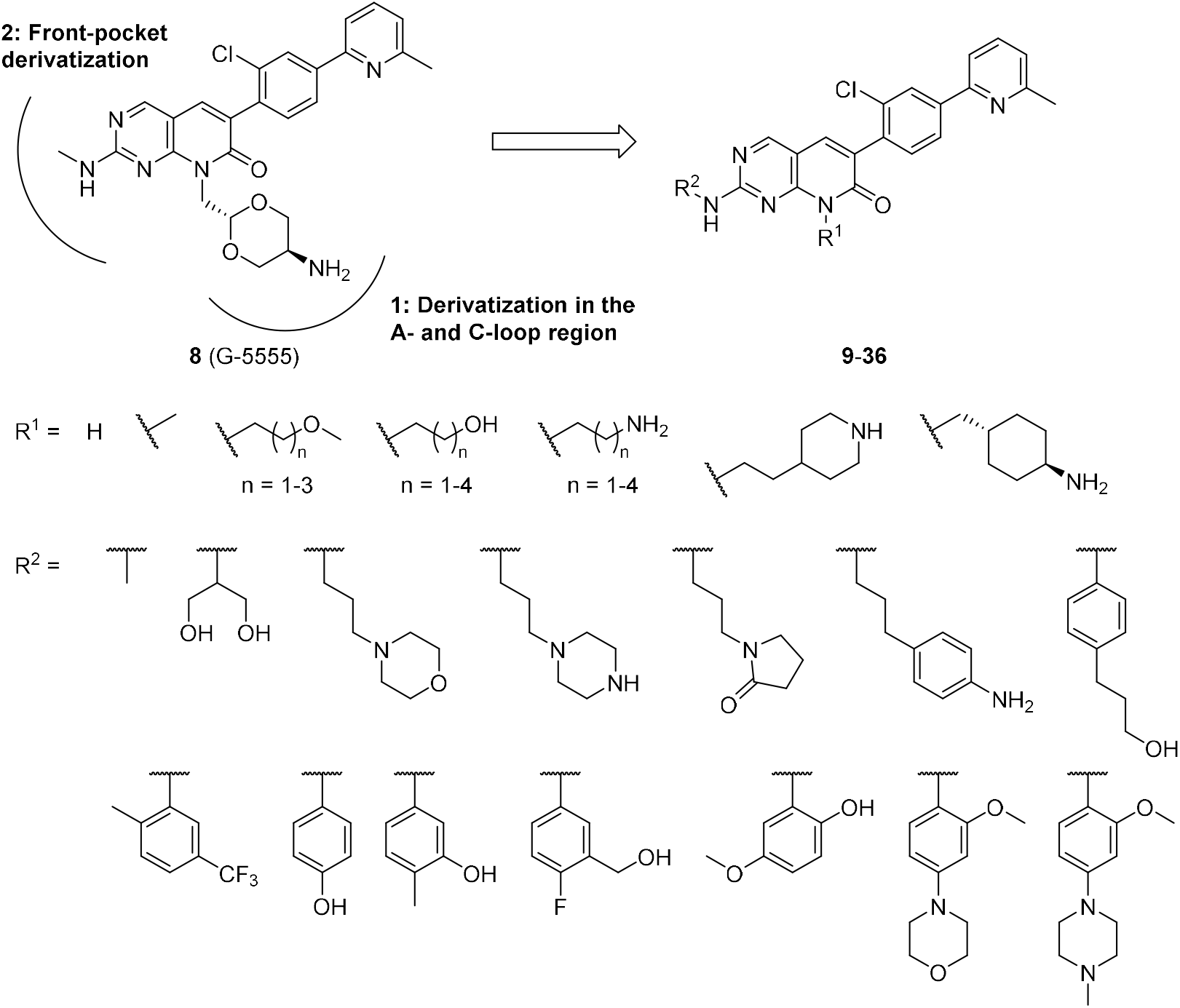
Design strategy for two series of pyrido[2,3-*d*]pyrimidin-7(8*H*)-one-based inhibitors modulating the selectivity of the lead structure towards MST kinases. In the first series, the (2*r*,5*r*)-2-methyl-1,3-dioxan-5-amine motif (R^1^) of G-5555 (**8**) was substituted with various aliphatic groups with potential hydrogen bond donors/acceptors. In the second series, the methylamine in the front-pocket binding region (R^2^) was modified by diverse aliphatic and aromatic amine moieties. For the corresponding combination of R^1^ and R^2^ see Table 1 and 2.

**Table 1.**
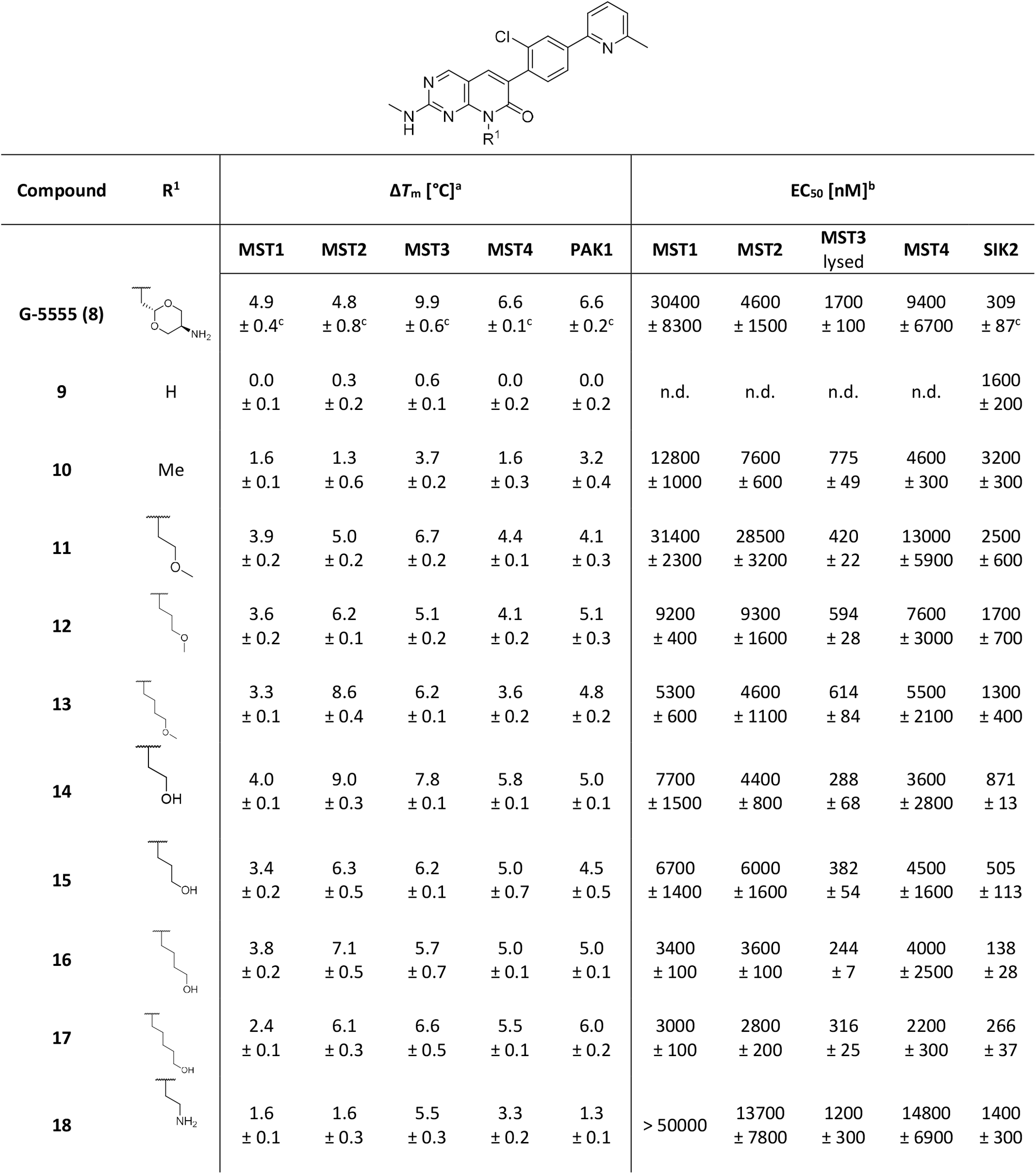
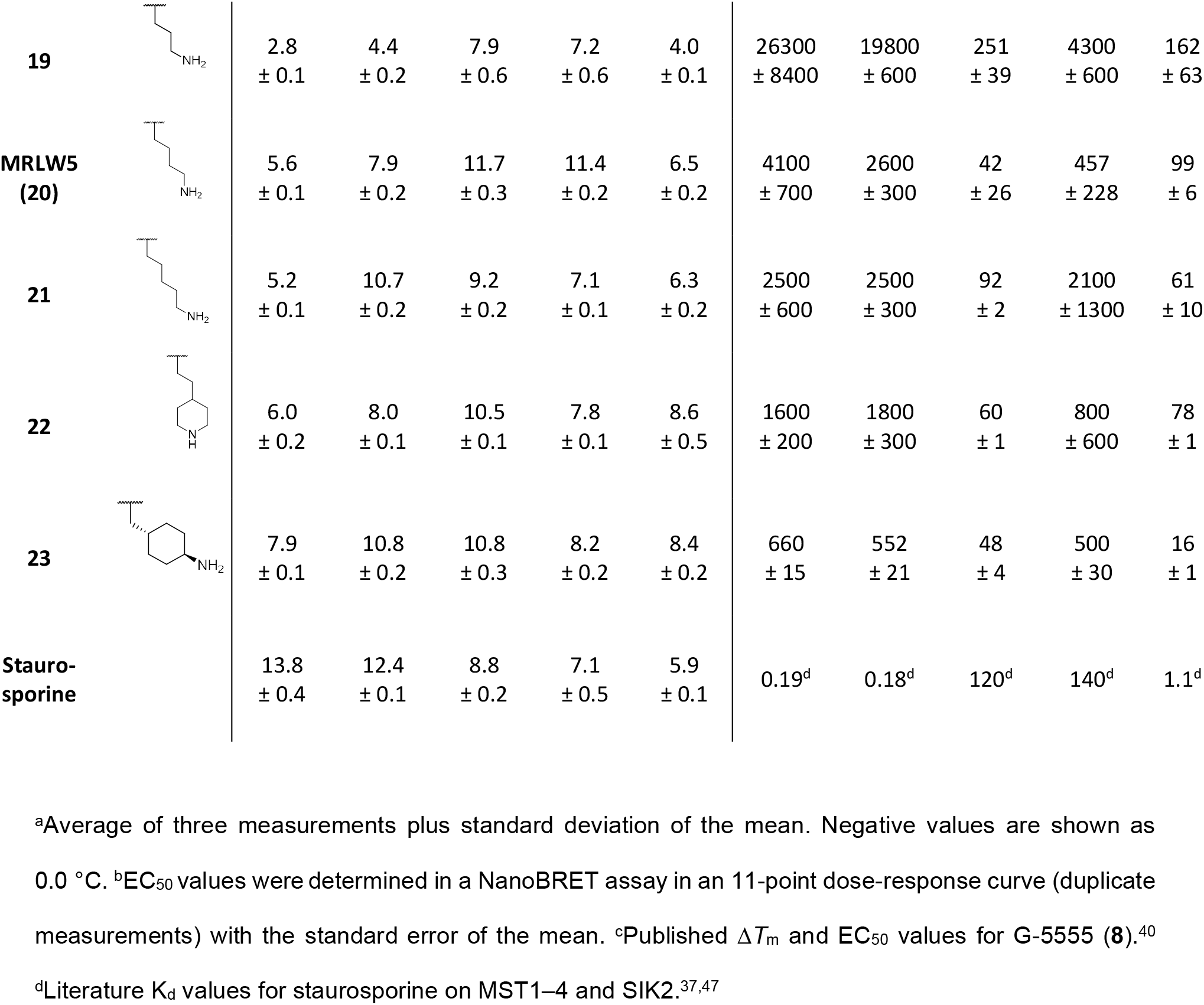
Selectivity profiles of pyrido[2,3-*d*]pyrimidin-7(8*H*)-one inhibitors 9–23 (A-loop/C-loop derivatives).

| Compound | R<sup>1</sup> | $\Delta T_m$ [°C]<sup>a</sup> | | | | | EC<sub>50</sub> [nM]<sup>b</sup> | | | | |
| --- | --- | --- | --- | --- | --- | --- | --- | --- | --- | --- | --- |
|  |  | MST1 | MST2 | MST3 | MST4 | PAK1 | MST1 | MST2 | MST3<br>lysed | MST4 | SIK2 |
| **G-5555 (8)** |  | 4.9<br>± 0.4<sup>c</sup> | 4.8<br>± 0.8<sup>c</sup> | 9.9<br>± 0.6<sup>c</sup> | 6.6<br>± 0.1<sup>c</sup> | 6.6<br>± 0.2<sup>c</sup> | 30400<br>± 8300 | 4600<br>± 1500 | 1700<br>± 100 | 9400<br>± 6700 | 309<br>± 87<sup>c</sup> |
| **9** | H | 0.0<br>± 0.1 | 0.3<br>± 0.2 | 0.6<br>± 0.1 | 0.0<br>± 0.2 | 0.0<br>± 0.2 | n.d. | n.d. | n.d. | n.d. | 1600<br>± 200 |
| **10** | Me | 1.6<br>± 0.1 | 1.3<br>± 0.6 | 3.7<br>± 0.2 | 1.6<br>± 0.3 | 3.2<br>± 0.4 | 12800<br>± 1000 | 7600<br>± 600 | 775<br>± 49 | 4600<br>± 300 | 3200<br>± 300 |
| **11** |  | 3.9<br>± 0.2 | 5.0<br>± 0.2 | 6.7<br>± 0.2 | 4.4<br>± 0.1 | 4.1<br>± 0.3 | 31400<br>± 2300 | 28500<br>± 3200 | 420<br>± 22 | 13000<br>± 5900 | 2500<br>± 600 |
| **12** |  | 3.6<br>± 0.2 | 6.2<br>± 0.1 | 5.1<br>± 0.2 | 4.1<br>± 0.2 | 5.1<br>± 0.3 | 9200<br>± 400 | 9300<br>± 1600 | 594<br>± 28 | 7600<br>± 3000 | 1700<br>± 700 |
| **13** |  | 3.3<br>± 0.1 | 8.6<br>± 0.4 | 6.2<br>± 0.1 | 3.6<br>± 0.2 | 4.8<br>± 0.2 | 5300<br>± 600 | 4600<br>± 1100 | 614<br>± 84 | 5500<br>± 2100 | 1300<br>± 400 |
| **14** |  | 4.0<br>± 0.1 | 9.0<br>± 0.3 | 7.8<br>± 0.1 | 5.8<br>± 0.1 | 5.0<br>± 0.1 | 7700<br>± 1500 | 4400<br>± 800 | 288<br>± 68 | 3600<br>± 2800 | 871<br>± 13 |
| **15** |  | 3.4<br>± 0.2 | 6.3<br>± 0.5 | 6.2<br>± 0.1 | 5.0<br>± 0.7 | 4.5<br>± 0.5 | 6700<br>± 1400 | 6000<br>± 1600 | 382<br>± 54 | 4500<br>± 1600 | 505<br>± 113 |
| **16** |  | 3.8<br>± 0.2 | 7.1<br>± 0.5 | 5.7<br>± 0.7 | 5.0<br>± 0.1 | 5.0<br>± 0.1 | 3400<br>± 100 | 3600<br>± 100 | 244<br>± 7 | 4000<br>± 2500 | 138<br>± 28 |
| **17** |  | 2.4<br>± 0.1 | 6.1<br>± 0.3 | 6.6<br>± 0.5 | 5.5<br>± 0.1 | 6.0<br>± 0.2 | 3000<br>± 100 | 2800<br>± 200 | 316<br>± 25 | 2200<br>± 300 | 266<br>± 37 |
| **18** |  | 1.6<br>± 0.1 | 1.6<br>± 0.3 | 5.5<br>± 0.3 | 3.3<br>± 0.2 | 1.3<br>± 0.1 | > 50000 | 13700<br>± 7800 | 1200<br>± 300 | 14800<br>± 6900 | 1400<br>± 300 |
| **19** |  | 2.8<br>± 0.1 | 4.4<br>± 0.2 | 7.9<br>± 0.6 | 7.2<br>± 0.6 | 4.0<br>± 0.1 | 26300<br>± 8400 | 19800<br>± 600 | 251<br>± 39 | 4300<br>± 600 | 162<br>± 63 |
| **MRLW5**<br>**(20)** |  | 5.6<br>± 0.1 | 7.9<br>± 0.2 | 11.7<br>± 0.3 | 11.4<br>± 0.2 | 6.5<br>± 0.2 | 4100<br>± 700 | 2600<br>± 300 | 42<br>± 26 | 457<br>± 228 | 99<br>± 6 |
| **21** |  | 5.2<br>± 0.1 | 10.7<br>± 0.2 | 9.2<br>± 0.2 | 7.1<br>± 0.1 | 6.3<br>± 0.2 | 2500<br>± 600 | 2500<br>± 300 | 92<br>± 2 | 2100<br>± 1300 | 61<br>± 10 |
| **22** |  | 6.0<br>± 0.2 | 8.0<br>± 0.1 | 10.5<br>± 0.1 | 7.8<br>± 0.1 | 8.6<br>± 0.5 | 1600<br>± 200 | 1800<br>± 300 | 60<br>± 1 | 800<br>± 600 | 78<br>± 1 |
| **23** |  | 7.9<br>± 0.1 | 10.8<br>± 0.2 | 10.8<br>± 0.3 | 8.2<br>± 0.2 | 8.4<br>± 0.2 | 660<br>± 15 | 552<br>± 21 | 48<br>± 4 | 500<br>± 30 | 16<br>± 1 |
| **Staurosporine** |  | 13.8<br>± 0.4 | 12.4<br>± 0.1 | 8.8<br>± 0.2 | 7.1<br>± 0.5 | 5.9<br>± 0.1 | 0.19<sup>d</sup> | 0.18<sup>d</sup> | 120<sup>d</sup> | 140<sup>d</sup> | 1.1<sup>d</sup> |

<sup>a</sup>Average of three measurements plus standard deviation of the mean. Negative values are shown as 0.0 °C. <sup>b</sup>EC<sub>50</sub> values were determined in a NanoBRET assay in an 11-point dose-response curve (duplicate measurements) with the standard error of the mean. <sup>c</sup>Published $\Delta T_m$ and EC<sub>50</sub> values for G-5555 (**8**).<sup>40</sup> <sup>d</sup>Literature K<sub>d</sub> values for staurosporine on MST1–4 and SIK2.<sup>37,47</sup>

**Table 2.**
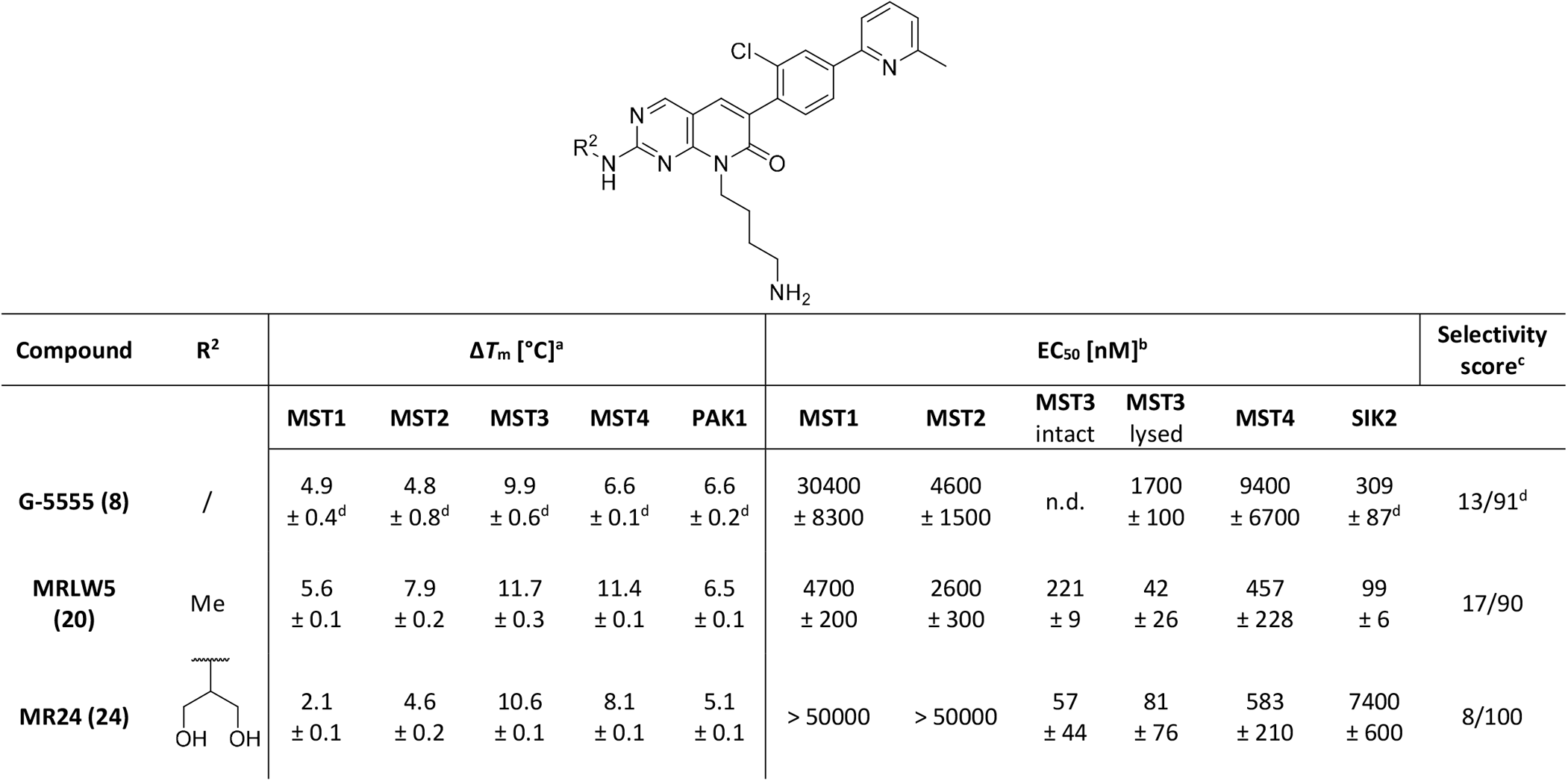
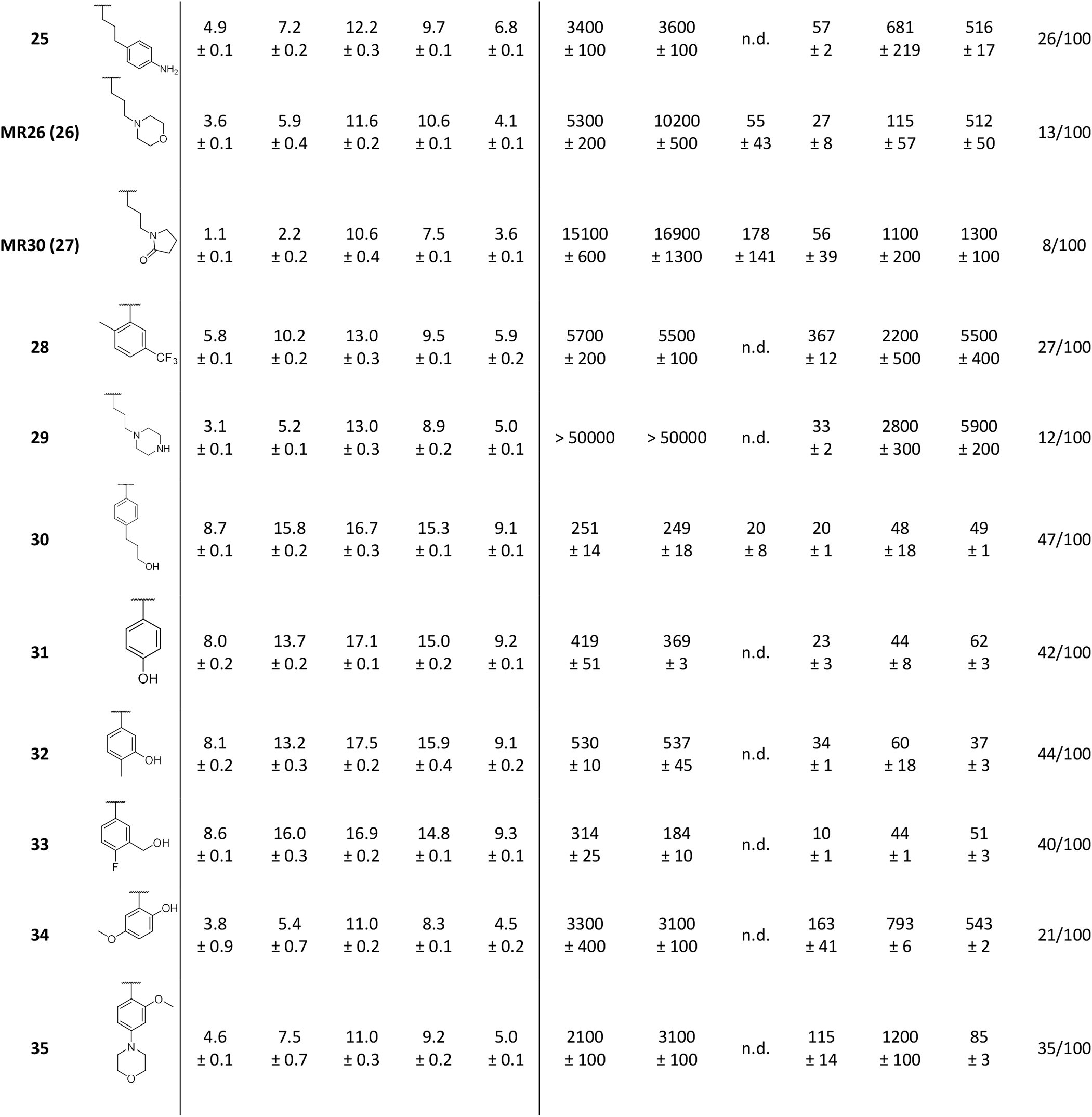
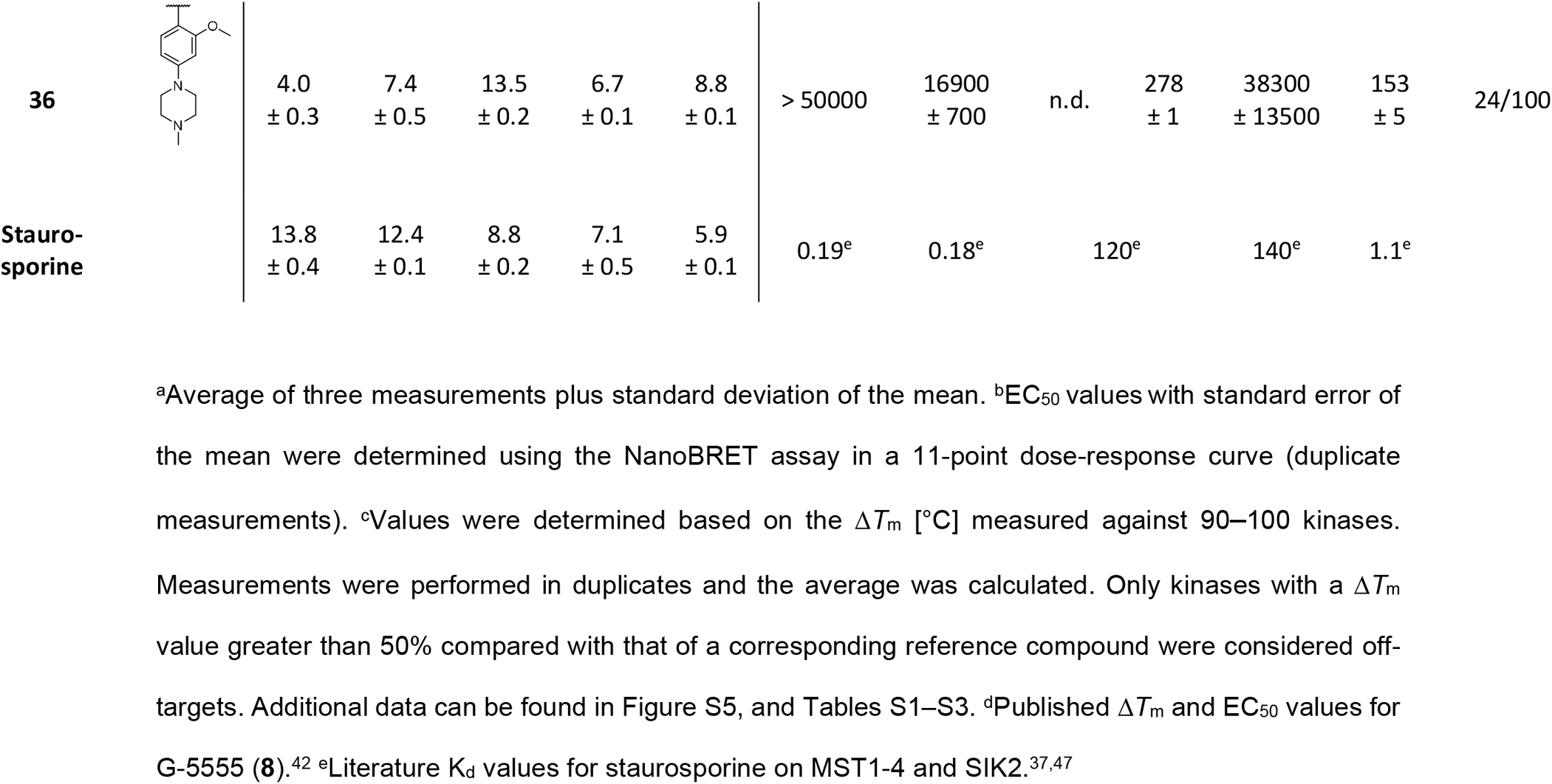
Selectivity profiles of pyrido[2,3-*d*]pyrimidin-7(8*H*)-one inhibitors 24–36 (front-pocket derivatives).

The (2*r*,5*r*)-2-methyl-1,3-dioxan-5-amine group of G-5555 (**8**) was substituted with various aliphatic methoxy, hydroxyl, or amino moieties of increasing length to investigate the effect on the binding affinity by mimicking the dioxane group of G-5555 (**8**). In addition, these moieties have a lower molecular weight than the lead structure. Following the improvement of the cellular binding affinity, key differences in the front-pocket region of MST3 were exploited by replacing the methylamine group with a diverse set of aromatic and aliphatic amine groups of varying size to optimize the ligand interactions in this region and thereby shift the selectivity towards MSTs.

### Synthesis of pyrido[2,3-*d*]pyrimidin-7(8*H*)-one-based MST inhibitors

The compounds in this work were synthesized in six to seven steps, depending on the derivative, as shown in Schemes 1–4. Main intermediates **41**, **52**, and **53** were prepared in larger scale, enabling the derivatization of the pyrido[2,3-*d*]pyrimidin-7(8*H*)-one scaffold at position 2 (front pocket) or 8 (A-loop/C-loop). The synthetic route was based on the one published by Rudolph *et al.*.^45^ Contrary, the route previously published by Tesch *et al.* was optimized to enable facile back-pocket derivatization.^40^ The synthesis of intermediate **41** (Scheme 1) started with a Fischer esterification of 2-(4-bromo-2-chlorophenyl)acetic acid (**37**). The resulting ethyl ester **38** was used in a subsequent Miyaura borylation reaction to give **39**. The transformation of **39** in a Suzuki cross-coupling with 2-bromo-6-methylpyridine resulted in the formation of **40**. The final step was a cyclocondensation of **40** with 4-amino-2-(methylthio)pyrimidine-5-carbaldehyde to obtain the pyrido[2,3-*d*]pyrimidin-7(8*H*)-one scaffold **41**. Starting from intermediate **41**, compounds **9**–**23** were synthesized in two to three steps (Scheme 2). Intermediates **42**–**56** were obtained in a nucleophilic substitution reaction, using the corresponding primary alkyl halides. This synthesis route included the key intermediates **52** and **53**, for which BOC- or phthalimide-protected 4-bromobutan-1-amine was used. In the case of intermediate **53**, a mixture of the isoindoline-1,3-dione and 2-carbamoylbenzoic acid derivative was obtained and then heated in acetic anhydride to fully convert the mixture to the isoindoline-1,3-dione containing intermediate **53**. The following steps led to compounds **9**–**17** and intermediates **57**–**62**. First, the methyl sulfide group at position 2 of the pyrido[2,3-*d*]pyrimidin-7(8*H*)-one scaffold was oxidized using 3-chloroperbenzoic acid (*m*-CPBA). This resulted in the formation of a mixture of the corresponding sulfoxide/sulfone derivatives, which was used in the next step without further separation. This mixture was then converted in a nucleophilic aromatic substitution (SNAr) under basic conditions using methylamine and *N*,*N*’-diisopropylethylamine (DIPEA). Subsequent deprotection of the BOC-protected amines **57**–**62** using trifluoroacetic acid (TFA) resulted in compounds **18**–**23**.

**Scheme 1.**
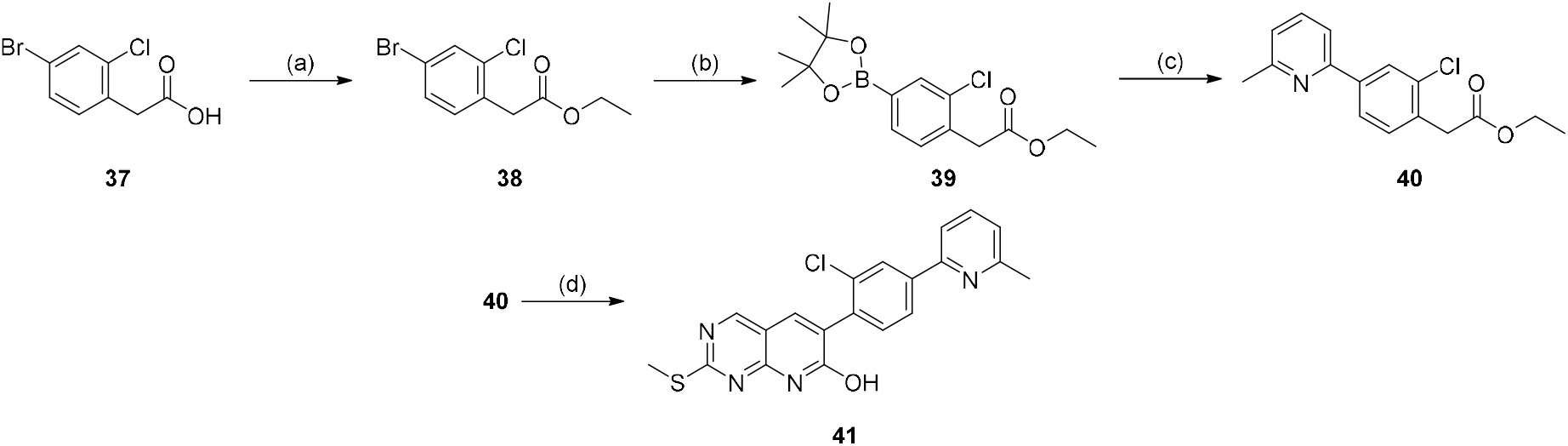
Reaction scheme of the synthesis of intermediate **41**. Reagents and conditions: (a) H2SO4, ethanol, 100 °C, 15 h; (b) bis(pinacolato)diboron, Pd(dppf)Cl2·DCM, KOAc, dioxane, 115 °C, 18 h; (c) 2-bromo-6-methylpyridine, Pd(dppf)Cl2, KOAc, dioxane/H2O (2:1), 107 °C, 18 h; (d) 4-amino-2-(methylthio)pyrimidine-5-carbaldehyde, K2CO3, DMF, 120 °C, 16 h.

**Scheme 2.**
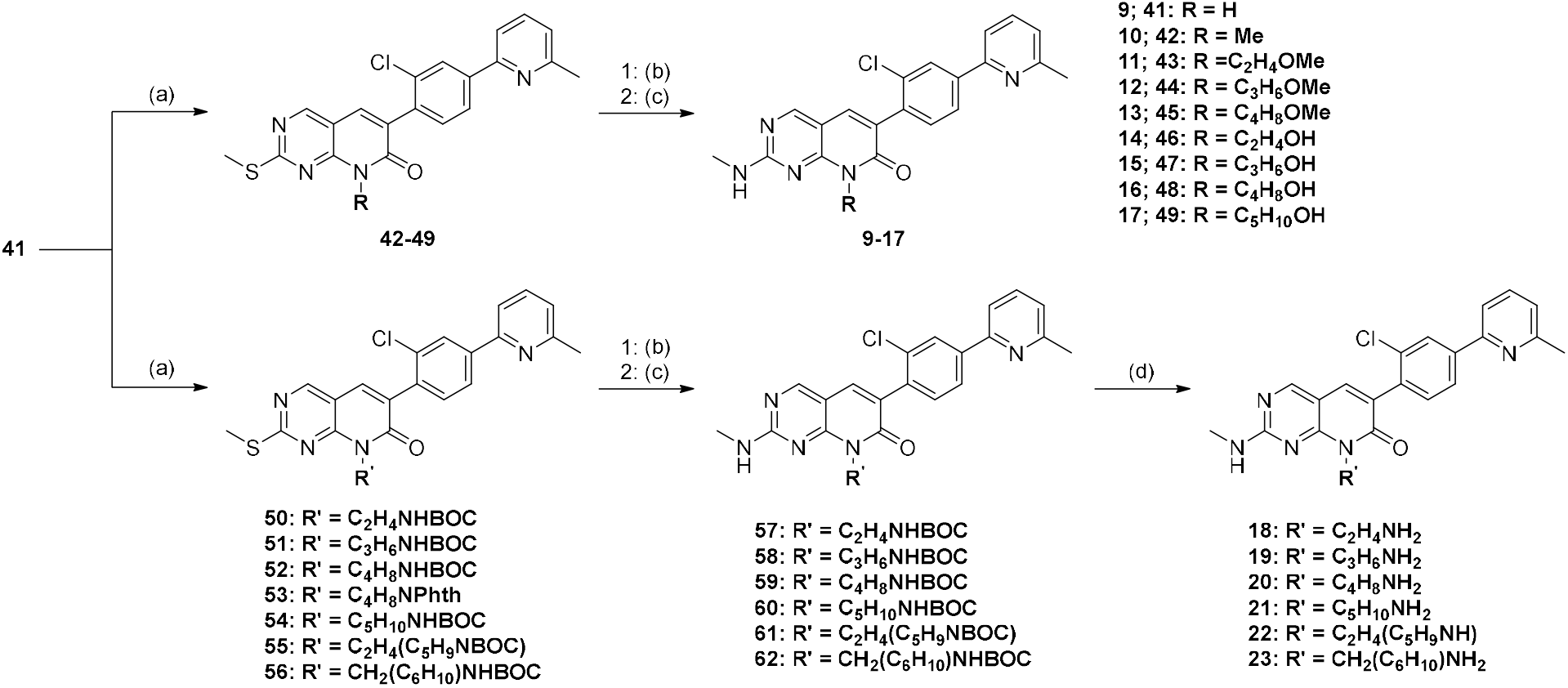
Synthesis of compounds **9**–**23** modified in the A-loop/C-loop region. Reagents and conditions: (a) primary alkyl halide, Cs2CO3, DMF, 110 °C, 18 h; except for **53**: 2-(4-bromobutyl)isoindoline-1,3-dione, K2CO3, DMSO, 110 °C, 18 h, then: Ac2O, 100 °C, 2 h; (b) 3-chloroperbenzoic acid (*m*-CPBA), DCM, RT, up to 3 h; (c) methylamine, *N,N’*-diisopropylethylamine (DIPEA), ethanol, 85 °C, 17 h; (d) TFA (20 vol%), DCM, RT, 0.5 h.

For the front-pocket derivatives **24**–**28**, main intermediate **53** was used as starting material (Scheme 3). As described above, the first step was the oxidation of the methyl sulfide group using *m*-CPBA. The resulting mixture of sulfoxide/sulfone product was then converted under different conditions. Compounds **24**–**27** were obtained by nucleophilic aromatic substitution using DIPEA and the corresponding primary aliphatic amine. In the case of compounds **24** and **25**, the reaction did not result in a complete deprotection of the phthalimide protected amine group. Therefore, the remaining residue was then converted in an additional nucleophilic substitution reaction using methylamine. In the case of compound **28**, the SNAr was performed based on a procedure described by Baiazitov *et al.*^46^ using lithium bis(trimethylsilyl)amide (LiHMDS) as the base. The reaction was performed at −70 °C, and 2-methyl-5-(trifluoromethyl)aniline was pre-activated with LiHMDS before the addition of the sulfoxide/sulfone mixture.

**Scheme 3.**
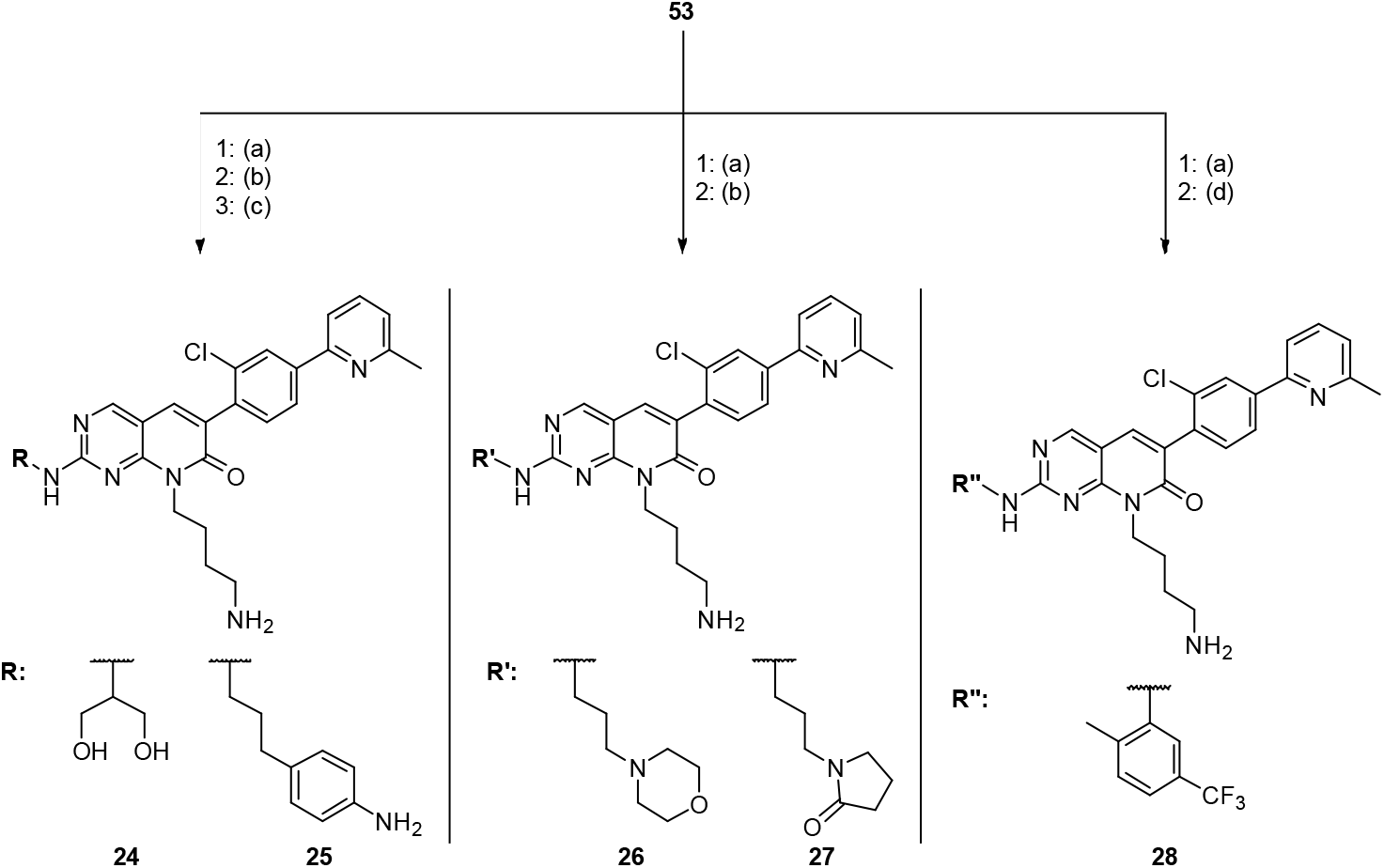
Synthesis of compounds **24**–**28** (front-pocket derivatives) using intermediate **53** as starting material. Reagents and conditions: (a) *m*-CPBA, DCM, RT, up to 3 h; (b) primary aliphatic amine, DIPEA, ethanol/DMF (5:1), 85 °C, 17 h; (c) methylamine, DIPEA, ethanol, 85 °C, 17 h; (d) 2-methyl-5-(trifluoromethyl)aniline, lithium bis(trimethylsilyl)amide (LiHMDS), THF, −70 °C, 1 h.

In addition, front-pocket derivatives **29**–**36** were synthesized using main intermediate **52** as starting material (Scheme 4). First, an oxidation of the methyl sulfide group was performed. In the case of compound **29**, this was followed by a SNAr using *tert*-butyl 4-(3-aminopropyl)piperazine-1-carboxylate to obtain **63**, and a subsequent deprotection of the BOC-protected amines using TFA. For compounds **30**–**36**, intermediates **64**–**70** were obtained under neutral conditions by stirring the sulfone/sulfoxide mixture with the corresponding aniline derivative at 85 °C for 17 hours. Finally, the BOC-protected amines **64**–**70** were deprotected using TFA to give compounds **30**–**36**.

**Scheme 4.**
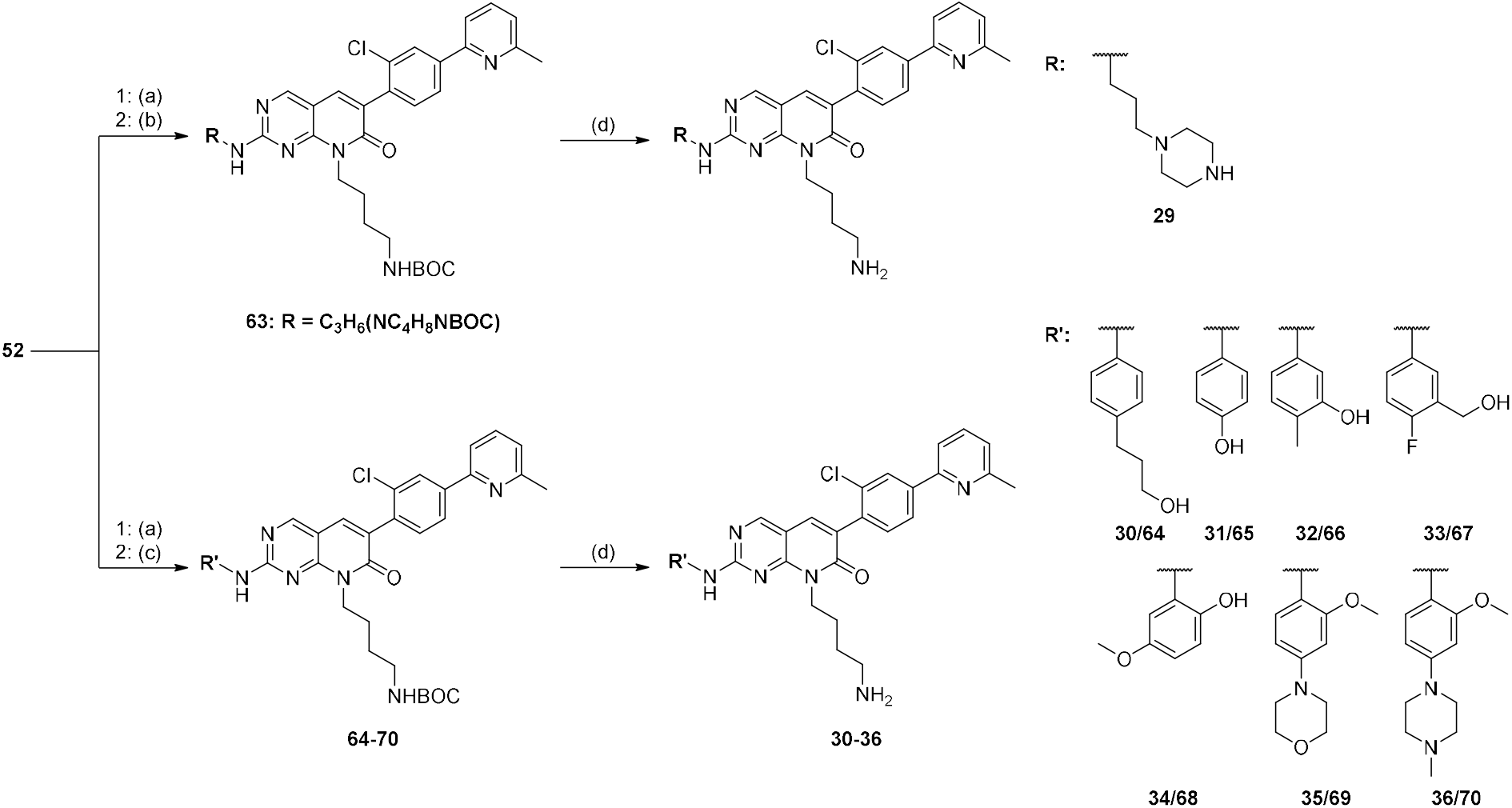
Synthesis of compounds **29**–**36** (front-pocket derivatives) using intermediate **52** as starting material. Reagents and conditions: (a) *m*-CPBA, DCM, RT, up to 3 h; (b) *tert*-butyl 4-(3-aminopropyl)piperazine-1-carboxylate, DIPEA, ethanol, 85 °C, 17 h; (c) primary aromatic amine, ethanol, 85 °C, 17 h; (d) TFA (20 vol%), DCM, RT, 0.5 h.

### Structure-activity relationship of pyrido[2,3-*d*]pyrimidin-7(8*H*)-one-derived MST inhibitors 9–23

On-target activity of the synthesized inhibitors against MST1, MST2, MST3, and MST4 was evaluated *in vitro* (Δ*T*m) by differential scanning fluorimetry (DSF) and *in cellulo* (EC50) using the NanoBRET target engagement assay (Table 1). For MST3, the EC50 values were determined in permeabilized cells in most cases due to technical challenges as outlined before. In addition, stabilization of the off-target kinase PAK1 was monitored *in vitro* (DSF assay). Selectivity towards SIK2 was assessed by determining the EC50 values in the NanoBRET assay. Compound **19** was previously published by Rudolph *et al.*^45^ and was resynthesized to evaluate for its potency and selectivity towards MST kinases. Published data of G-5555 (**8**) were used for comparison.^40^

Removal of the (2*r*,5*r*)-2-methyl-1,3-dioxan-5-amine group of G-5555 (**8**) in compounds **9** and **10** resulted in a complete loss of activity against all kinases tested. Compounds **11**–**13**, with an aliphatic methoxy group at position 8 of the pyrido[2,3-*d*]pyrimidin-7(8*H*)one scaffold, showed a comparable on- and off-target profile, and no strong binding of MST1, MST2, MST4, or SIK2 was observed in cells (EC50 values > 1 µM). Replacing the methoxy with a hydroxyl group (**14**–**17**) did generally also not result in a significant improvement for all MSTs, with one exception: an increased stabilization was observed for **14** on MST2, with a Δ*T*m value of 9.0 ± 0.3 °C compared with 4.8 ± 0.8 °C for G-5555 (**8**). In addition, all four compounds showed increased binding affinities against SIK2, with EC50 values below 1 µM. While **18**, with a 2-ethylamine group, showed no activity on either MST3 or SIK2, elongating the aliphatic chain by one carbon atom in **19** partially restored the stabilization of MST3 with Δ*T*m = 7.9 ± 0.6 °C. For MRLW5 (**20**), the highest Δ*T*m values were observed for MST3 and MST4 (11.7 ± 0.3 °C and 11.4 ± 0.2 °C, respectively). These values correlated well with strong binding affinities of 457 ± 228 nM (MST4) and 42 ± 26 nM (MST3; lysed). Unfortunately, SIK2 was also strongly inhibited with an EC50 = 99 ± 6 nM. A 5-pentylamine moiety (**21**) at position 8 of the main scaffold did not improve Δ*T*m values or potency compared with MRLW5 (**20**). **22** strongly bound MST3 and MST4 with EC50 values of 60 ± 1 nM (MST3, lysed) and 800 ± 600 nM (MST4). Interestingly, **23**, which has a (1*r*,4*r*)-4-methylcyclohexan-1-amine group, potently bound all four MST kinases (MST1-4) with EC50 values of 660 ± 15 nM, 552 ± 21 nM, 48 ± 4 nM (lysed), and 500 ± 30 nM, respectively. However, both compounds **22** and **23** were also potent inhibitors of SIK2 in the NanoBRET assay.

### Selectivity profiles of MRLW5 (20), 22, and 23 against the human kinome

Compounds MRLW5 (**20**), **22**, and **23**, which showed potent cellular on-target binding of MST4, were further evaluated in a subsequent DSF panel assay against up to 100 kinases to estimate the kinome-wide selectivity (Figure S4). In this assay, representative kinases from all branches of the phylogenetic kinome tree were included, as shown in Table S1. The obtained Δ*T*m values were compared with those of the corresponding reference compound, and only kinases with Δ*T*m values larger than 50% compared to the reference compound were considered potential off-targets. The suitability of the DSF panel for estimating kinome-wide selectivity was recently evaluated by comparing the results of G-5555 (**8**) in the DSF panel assay with the selectivity data from the scanMAX assay on 468 kinases (*Eurofins Discovery*).^42^ For G-5555 (**8**) this resulted in a selectivity score of 13/91 kinases and an excellent overlap between the two assays was observed.^42^

The increased potency of **23** on all four MST kinases correlated with lower overall selectivity, with 32 of 94 kinases stabilized by more than 50% compared with corresponding reference compounds. The selectivity score of **22** was also reduced compared with G-5555 (**8**), with 25/98 kinases. The best ratio between on-target potency and selectivity was observed for compound MRLW5 (**20**). Unlike the lead structure, MRLW5 (**20**) potently bound to MST3 and MST4 in cells, with EC50 values of 42 ± 26 nM and 457 ± 228 nM, respectively, as described above. In addition, a comparable selectivity score was obtained with 17 of 90 kinases.

To analyze the binding mode of MRLW5 (**20**), we determined the crystal structure of the complex with MST3 and compared it with the published structure of the MST3-G-5555 (**8**) complex.

Superimposition of the two structures revealed an essentially identical orientation of the central inhibitor scaffold (Figure 4). The newly introduced 4-aminobutyl group at position 8 of the pyrido[2,3-*d*]pyrimidin-7(8*H*)-one scaffold formed a salt-bridge with Asp174 of the DFG motif and hydrogen bonds with the side chain of Asn161 and the backbone oxygen of Ala160. Because of the improved affinity, MRLW5 (**20**) was selected as the lead structure for subsequent front-pocket optimization, resulting in the synthesis of compounds **24**–**36** with a 4-aminobutyl group interacting with the A-loop/C-loop region.

**Figure 4.**
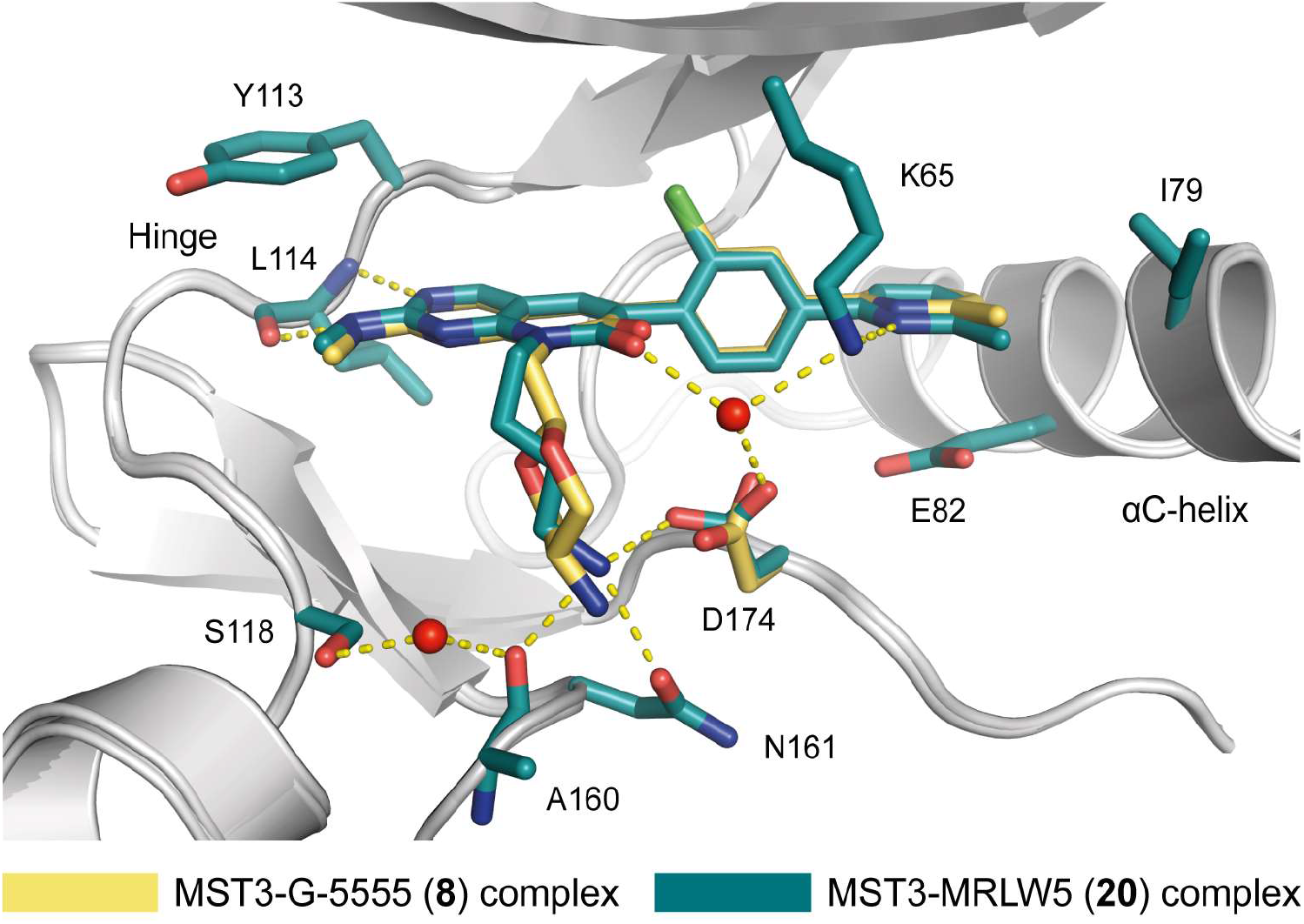
Comparison of the co-crystal structures of G-5555 (**8**) (PDB code 7B30) and MRLW5 (**20**) (PDB code 8BZJ) with MST3. G-5555 (**8**) is shown in yellow and MRLW5 (**20**) in cyan. MST3 is colored in gray. Water molecules are shown as red spheres and hydrogen bonds as dashed yellow lines. The water molecules, hydrogen-bond network, and amino acids are shown as in the structure of MRLW5 (**20**)-MST3.

### Modifications of the pyrido[2,3-*d*]pyrimidin-7(8*H*)-one scaffold in position R2

Similar to the lead structure G-5555 (**8**), compounds with aliphatic amine groups in the front-pocket region showed no activity on MST1 and MST2. Substitution of the methylamine group of MRLW5 (**20**) with a serinol (2-amino-1,3-propandiol) moiety in compound MR24 (**24**) resulted in slightly lower Δ*T*m values for MST3 and MST4 than for the parent molecule MRLW5 (10.6 ± 0.1 °C and 8.1 ± 0.1 °C, respectively). However, MR24 (**24**) showed a comparable binding of MST4 in cells with an EC50 value of 583 ± 210 nM *vs* 457 ± 228 nM for MRLW5 (**20**), and improved potency for MST3 with an EC50 = 57 ± 44 nM *vs* 221 ± 9 nM, respectively. In addition, MR24 (**24**) only weakly bound to SIK2, with an EC50 value in the micromolar range. The overall selectivity in the DSF panel assay against 100 kinases was significantly increased compared with G-5555 (**8**) and MRLW5 (**20**), with a selectivity score of only 8/100 kinases.

The largest aliphatic front-pocket motif (4-(3-aminopropyl)aniline) in compound **25** resulted in strong binding to MST3, MST4, and SIK2, with EC50 values below 1 µM. However, the overall selectivity was decreased with 26/100 kinases. A 3-morpholinopropan-1-amine group in the front-pocket region, as in MR26 (**26**), decreased the stabilization of PAK1, with a Δ*T*m shift of 4.1 ± 0.1 °C compared with a Δ*T*m shift of 6.6 ± 0.2 °C for G-5555 (**8**). Strong cellular on-target activity was observed, with EC50 values of 55 ± 43 nM (MST3) and 115 ± 57 nM (MST4), but MR26 (**26**) also strongly bound SIK2 in cells (EC50 = 512 ± 50 nM). The selectivity score was slightly decreased compared with MR24 (**24**), with 13/100 kinases stabilized.

In MR30 (**27**) the methylamine group of MRLW5 (**20**) was replaced by a 1-(3-aminopropyl)pyrrolidin-2-one group. This resulted in the lowest stabilization of PAK kinases observed for the different front-pocket modifications, with a Δ*T*m shift of 3.6 ± 0.1 °C. The cellular activity was comparable to MR24 (**24**), with an EC50 value of 178 ± 141 nM against MST3. Surprisingly, MR30 (**27**) only weakly bound to MST4 (EC50 > 1 µM), and no binding was detected for SIK2. MR30 (**27**) stabilized 8 out of 100 kinases by more than 50% compared with the stabilization by a corresponding reference compound.

The 2-methyl-5-(trifluoromethyl)aniline moiety of **28** induced a significant stabilization of MST2, MST3, and MST4 in the DSF assay with Δ*T*m values > 9 °C. Cellular binding affinity, however, was weak against all four kinases tested, as indicated by high EC50 values. Replacing the oxygen atom in the front-pocket motif of MR26 (**26**) with a nitrogen atom in **29** (3-(piperazin-1-yl)propan-1-amine moiety) resulted in a decreased cellular binding of MST4 and SIK2 and corresponding EC50 values higher than 1 µM.

Reduced selectivity was observed for most compounds with an aromatic amine group in the front-pocket region. **30** stabilized all kinases tested in the DSF assay significantly more than the lead structure G-5555 (**8**), and also more than the multi-kinase inhibitor staurosporine. It bound all four MSTs and SIK2 with EC50 values in the nanomolar range, but we observed only low overall selectivity (47/100 kinases).

Compounds **31** and **32**, containing a 4-aminophenol and a 5-amino-2-methylphenol group, showed the most pronounced Δ*T*m stabilization of MST3, twice as high as the one observed for staurosporine (Δ*T*m = 8.8 ± 0.2 °C), with Δ*T*m values of 17.1 ± 0.1 °C and 17.5 ± 0.2 °C, respectively. However, although this correlated with high cellular potency on MST1, MST2, MST4, and SIK2, the overall selectivity scores were again very low (42/100 and 44/100 kinases, respectively). Compound **33** also showed low EC50 values against all kinases tested and potently stabilized PAK1, with a Δ*T*m = 9.3 ± 0.1 °C.

The hydroxyl group at position 2 and the methoxy group at position 5 of the phenyl ring in **34** decreased the stabilization against all five kinases tested in the DSF assay (compared with **33**). In addition, the group was detrimental to the cellular potency on MST1/2, with EC50 values > 3 µM. The fact that substituents in position 2 (compounds **28** and **34**) led to lower stabilization on all four MST kinases suggests a suboptimal packing of the aromatic ring in the front pocket. Consistent with this hypothesis, compounds **35** and **36**, containing the largest aromatic groups occupying the MST front pocket, only weakly bound to MST1, MST2, or MST3. However, both compounds exhibited low EC50 values against SIK2 of 153 ± 5 nM and 85 ± 3 nM, respectively.

In summary, all compounds harboring an aliphatic moiety in the front-pocket region showed lower Δ*T*m values for MST kinases and PAK1 in the DSF assay compared with compounds with an aromatic group in this position. The latter, however, generally displayed a reduced selectivity compared with the lead structure G-5555 (**8**). The most potent cellular on-target activity was observed for compounds MR24 (**24**), MR26 (**26**), and MR30 (**27**), which was also associated with improved selectivity scores of 8/100, 13/100, and 8/100 kinases, respectively.

### Contribution of polar interactions and hydrophobic packing on the binding affinities of MR24 (24) and MR26 (26)

To provide a structural basis for understanding the potent on-target activity of compound MR24 (**24**), we determined high-resolution crystal structures of MST3 in complex with MR24 (**24**) and MR26 (**26**) (resolution of 1.9 and 1.6 Å, respectively). In addition, we determined the structure of the MST3 complex with the less potent but MST3-selective MR30 (**27**) (1.5 Å resolution). As already seen for the parent molecule (MRLW5 (**20**)), MR24 (**24**), MR26 (**26**), and MR30 (**27**) formed canonical hydrogen-bond interactions with the hinge-region backbone (Leu114), and the 4-aminobutyl moiety formed a salt bridge with the DFG motif (Asp174) and hydrogen bonds with C-loop residues (backbone of Ala160 and side chain of Asn161) of MST3 (Figure 5A, B, and C). The central scaffold perfectly superimposed in all three complexes (Figure 5D). Importantly, the two newly introduced hydroxyl groups in MR24 (**24**) formed a bidentate water-mediated hydrogen-bond network with Tyr113, Ser118, and Asp121, contributing to the potent binding of MR24 (**24**) to MST3. MR26 (**26**), however, only formed a water-mediated hydrogen bond with Tyr113 but did not engage in the water network extending to amino acids of the αD-helix or the hinge region (Ser118). Nevertheless, both compounds (**24** and **26**) exhibited similar on-target EC50 values of 57 ± 44 nM and 55 ± 43 nM on MST3, respectively, suggesting an increased contribution of van der Waals interactions to the binding affinity of MR26 (**26**) compared with MR24 (**24**).

**Figure 5.**
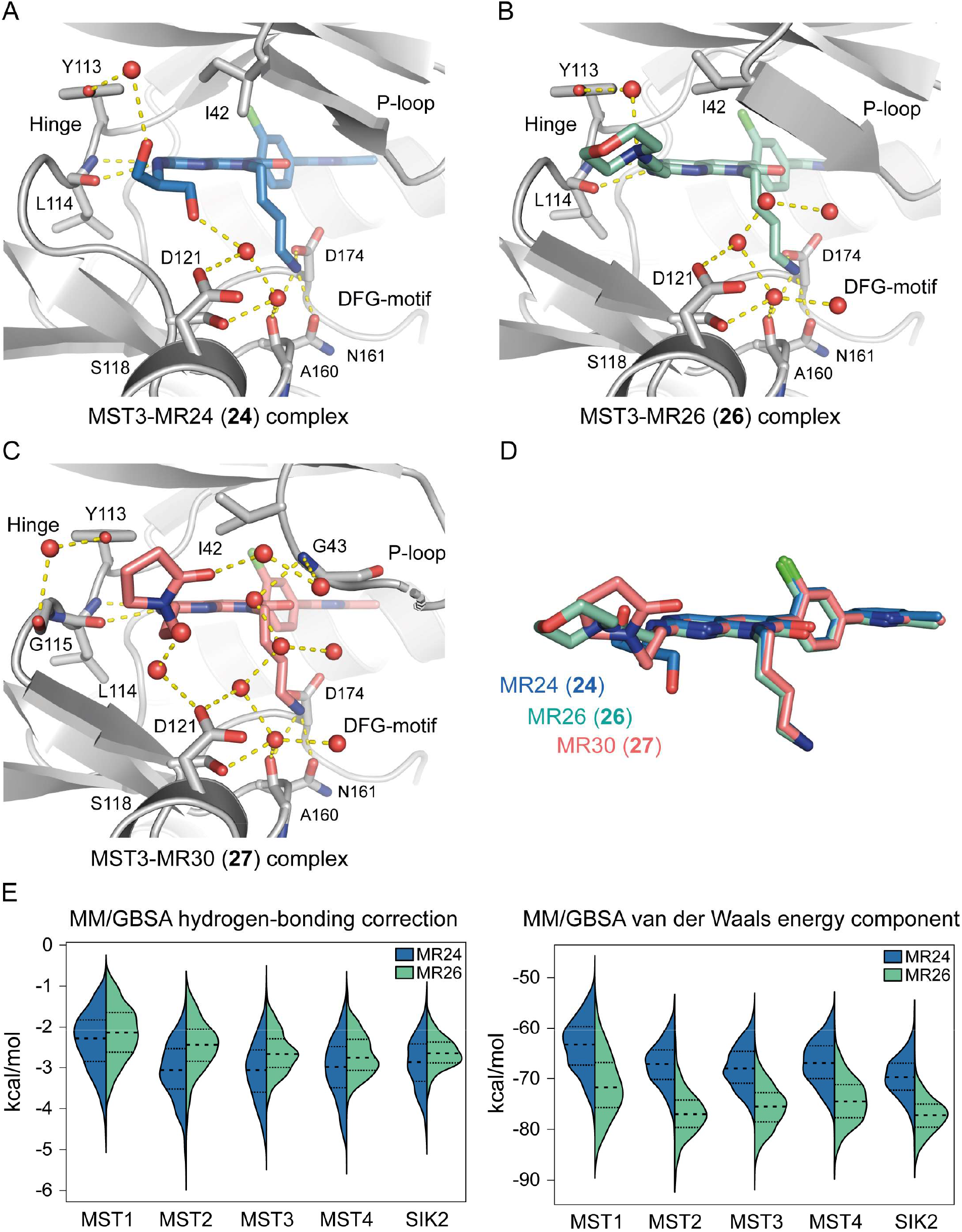
Analysis of the binding modes of MR24 (**24**), MR26 (**26**), and MR30 (**27**) in co-crystal structures with MST3 and molecular mechanics calculations. Hydrogen bonds are shown as yellow dashed lines and water molecules as red spheres. (A) Crystal structure of MST3 (gray) with bound MR24 (**24**; blue). An extensive water-mediated hydrogen-bond network can be found in the front-pocket region of the kinase. (B) Crystal structure of MST3 (gray) bound to MR26 (**26**; pale cyan). Fewer interactions between the kinase and the inhibitor in the front-pocket region were found. (C) Crystal structure of MST3 (gray) in complex with MR30 (**27**; salmon). The front-pocket motif interacts with the water network connecting the P-loop and the amino acids of the A-/C-loops. (D) Overlay of MR24 (**24**), MR26 (**26**), and MR30 (**27**) in the co-crystal structures with MST3. (E) Violin plot representation of MM/GBSA hydrogen-bonding correction component (*left*) and van der Waals energy component (*right*), derived from MD simulations. The bold dashed line represents the median. The thin dashed line represents the interquartile range (IQR). On each side of the vertical black line, a kernel density estimation shows the distribution shape of data. The wider sections represent a higher probability that members of the population will take on the given value.

In the structure of MR30 (**27**) in complex with MST3, the 1-(3-aminopropyl)pyrrolidin-2-one group in the front-pocket motif interacts with the water network connecting the P-loop with the amino acids of the A-/C-loops. However, the similar potency of MR30 (**27**) and the parent molecule MRLW5 (**20**) on MST3 (EC50 = 178 ± 141 nM *vs* 221 ± 9 nM, respectively) suggested that this additional hydrogen bonding did not contribute significantly to binding. In addition, no direct interaction of the front-pocket motif of **27** with Tyr113 was observed, in contrast to the structures of MR24 (**24**) and MR26 (**26**) in complex with MST3.

Binding of MR24 (**24**) and MR26 (**26**) was further investigated in molecular dynamic simulations (Figure S6 and Tables S6 and S7), and the free energy of both compounds in the binding pocket of MST3 was calculated by molecular mechanics with generalized Born and surface area solvation (MM/GBSA) and analyzed for its hydrogen-bonding correction component and van der Waals energy component (Figure 5E). The higher range of values and lower median observed for the hydrogen-bonding energy component of MR24 (**24**) compared with MR26 (**26**), as well as the lower values of the van der Waals energy component of MR26 (**26**), confirmed the hypothesis of an increased contribution of van der Waals interactions to the binding affinity of MR26 (**26**) compared with MR24 (**24**).

The potential off-targets of MR24 (**24**), MR26 (**26**), and MR30 (**27**) based on DSF data were further evaluated in a cellular context using the NanoBRET assay (Figures 6 and S7). As described above, MR26 (**26**) showed a selectivity score of 13/100 kinases and MR30 (**27**) of only 8/100 kinases in the DSF panel assay. The *in vitro* stabilization of NEK2, CK1e, and FES by both compounds was not confirmed in cells, as indicated by high EC50 values of more than 1 µM in NanoBRET assays. MR26 (**26**) and MR30 (**27**) are, therefore, suitable tool compounds for profiling GCKIII-subfamily kinases in a cellular context. In addition, MR30 (**27**) also showed selectivity within the MST subfamily, with only MST3 being potently inhibited in cells.

**Figure 6.**
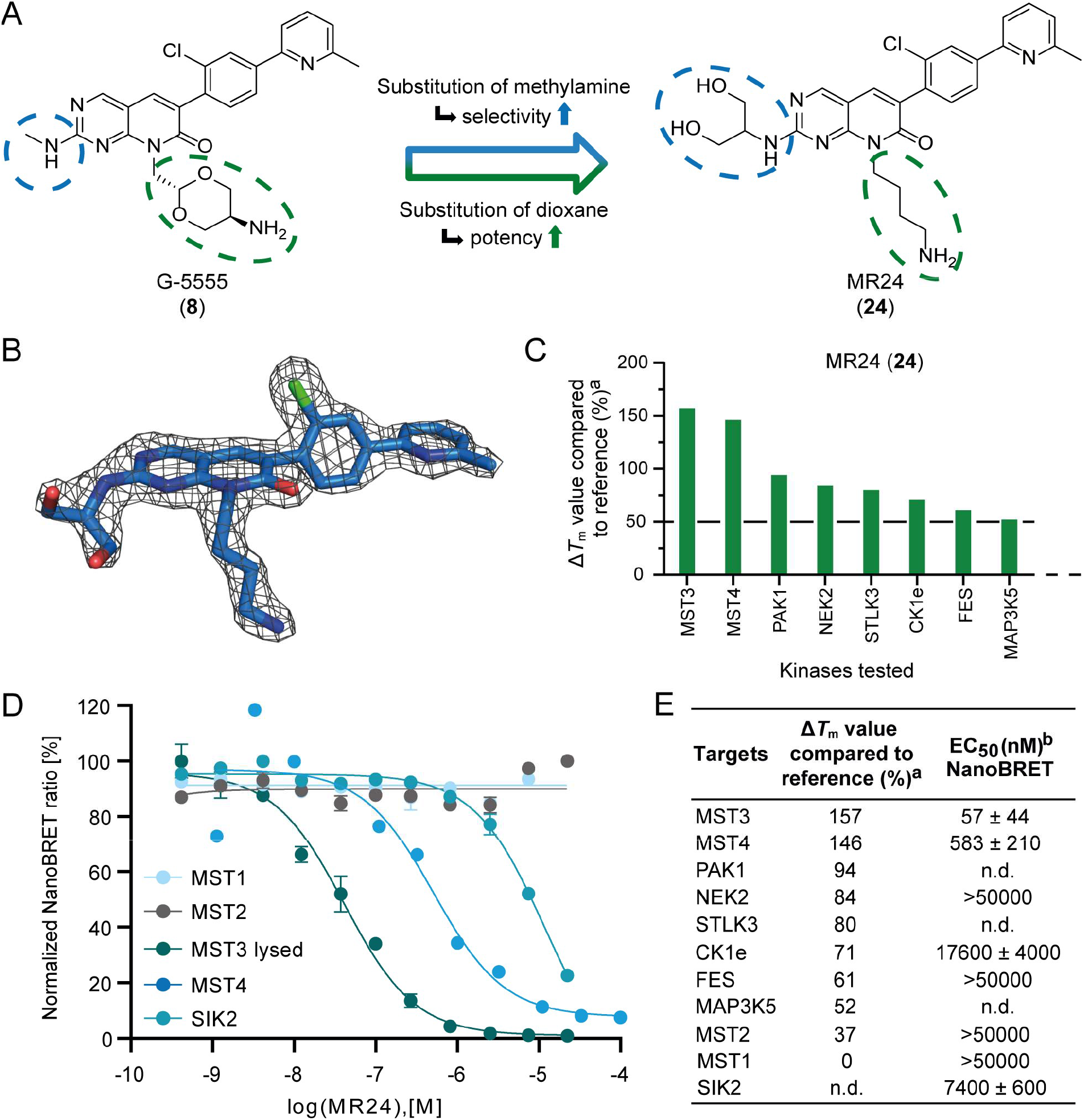
Selectivity profile of MST3/4 inhibitor MR24 (**24**). (A) Overview of the modifications of the lead structure G-5555 (**8**). Substitution of the (2*r*,5*r*)-2-methyl-1,3-dioxan-5-amine moiety of compound G-5555 (**8**) with a 4-aminobutyl group increased the cellular potency on MST3 and MST4. Replacement of the methylamine group with a serinol moiety improved the overall selectivity. (B) 2Fo–Fc electron density map around the ligand in the MST3-MR24 (**24**) complex, shown at a contour level of 2.0 σ. (C) Selectivity data of MR24 (**24**) in the DSF panel assay against 100 kinases, in percent (%). Obtained Δ*T*m values were compared with a corresponding reference compound, and only kinases with values higher than 50% were considered potential off-targets. (D) NanoBRET curves obtained on MST1-4 and SIK2 using compound MR24 (**24**). (E) Cellular on- and off-target evaluation of MR24 (**24**) using the NanoBRET assay. In addition to the kinases stabilized by more than 50% in the DSF panel assay, MST1, MST2, and SIK2 were tested. ^a^Data included in this figure are shown in Figure S5 and Table S1. ^b^EC50 values with standard error of the mean were determined using the NanoBRET assay in a 11-point dose-response curve (duplicate measurements).

In the case of MR24 (**24**), the lead structure G-5555 (**8**) was improved by replacing the (2*r*,5*r*)-2-methyl-1,3-dioxan-5-amine group by a 4-aminobutyl moiety and substituting the methylamine group in the front-pocket region with a serinol moiety (Figure 6A). In the DSF panel assay against 100 kinases, MR24 (**24**) stabilized only 8 kinases (MST3, MST4, PAK1, NEK2, STLK3, CK1e, FES, and MAP3K5) by more than 50% compared with a corresponding reference compound (Figure 6C). Similar to the lead structure, MR24 (**24**) showed no cellular binding for MST1/2 in NanoBRET assays (EC50 values > 50 µM) (Figure 6D). Importantly, the inhibitor also only weakly bound to NEK2, CK1e, and FES, with EC50 values in the micromolar range (Figure 6E). However, the other potential off-targets, PAK1, STK39, and MAP3K5, remain to be addressed in future studies as currently no NanoBRET assays are available for these targets. The strong Δ*T*m shift for PAK1 suggests, that MR24 (**24**) retains activity for this kinase. The modifications resulting in MR24 (**24**) further diminished the activity against SIK2 in cells (EC50 = 7.4 ± 0.6 µM) observed for the lead structure. In conclusion, MR24 (**24**) together with the chemogenomic compound PF-06447475 (**2**) can be used to investigate the entire MST subfamily *in cellulo* in mechanistic and phenotypic studies. The combination with the compound MR30 (**27**) allows target deconvolution between the understudied kinases MST3 and MST4 in cells.

### Cellular evaluation of the selective MST3/4 inhibitors

Comprehensive annotation of compounds, both in terms of selectivity and general properties in cell-based systems, is important to avoid false positive results in phenotypic assays.^48-51^ As a first assessment, we investigated effects on cell health using HCT116 cells in a high-content assay format.^52^ Most investigated compounds didn’t effect cell viability in comparison to the control, even at the highest concentration tested (10 µM) (Figure S8A and Table S8). However, MRLW5 (**20**), **30**, and MR26 (**26**) showed high intensity fluorescence after 24 h treatment (Figure S8B). Spectral analysis revealed that this effect was a result of intrinsic fluorescence of compound **30** (Figure S9). For compounds MRLW5 (**20**) and MR26 (**26**), no intrinsic fluorescence was detectable, suggesting that precipitation is responsible for the observed high fluorescence (Figure S8C). This behavior is most likely the cause of the significantly reduced cell health as compound precipitation can trigger apoptotic cell death.

Interestingly, PF-06447475 (**2**) increased the cell count after 48 h at both concentrations (1 µM and 10 µM) compared with cells treated with DMSO (0.1%), suggesting an increase in proliferation upon inhibition of MST1/2. MST1/2 are known to act as growth suppressors,^53^ and MST1 downregulation has been reported to increase cell proliferation in glioma cell lines by inhibition of AKT1 signalling.^54^

Compounds MR24 (**24**) and MR30 (**27**) showed no significant effect on the normalized cell count at a concentration of 1 µM. At 10 µM, a slight reduction in cell viability by about 30% was observed for both compounds. Treatment with MR24 (**24**) resulted in over 90% healthy nuclei after 24 h at both concentrations used (1 µM and 10 µM) (Figure S8D).

MST1, MST2, and MST3 regulate mitotic cell-cycle progression by phosphorylating the LATS1/2 homologs NDR1/2, which mediate gene transcription through phosphorylation of YAP/TAZ.^55^ Consistent with this finding, Cornils *et al.* showed that NDR1/2 are selectively activated in the G1 phase of the cell cycle and that inhibition of MST3 results in a cell-cycle arrest.^10^ We therefore performed a live-cell assay using the fluorescent ubiquitination-based cell cycle indicator (FUCCI) system.^56,57^ With this technology, it is possible to differentiate the cell-cycle phases G1, G2/M, or S phase at single-cell level. We tested the most promising compounds at three different concentrations (1 µM, 5 µM, and 10 µM) over 72 h (Figure 7A and Table S9). Like the CDK2 inhibitor milciclib,^58,59^ which was used as a positive control for G1-phase arrest, the MST3/4 inhibitors MR24 (**24**) and MR30 (**27**) increased the fraction of cells in G1 phase at 10 µM (Figure 7B and C). In contrast to the MST3/4-selective inhibitors, a dose-dependent G2/M phase arrest was observed for the MST1/2 inhibitor PF-06447475 (**2**).

**Figure 7:**
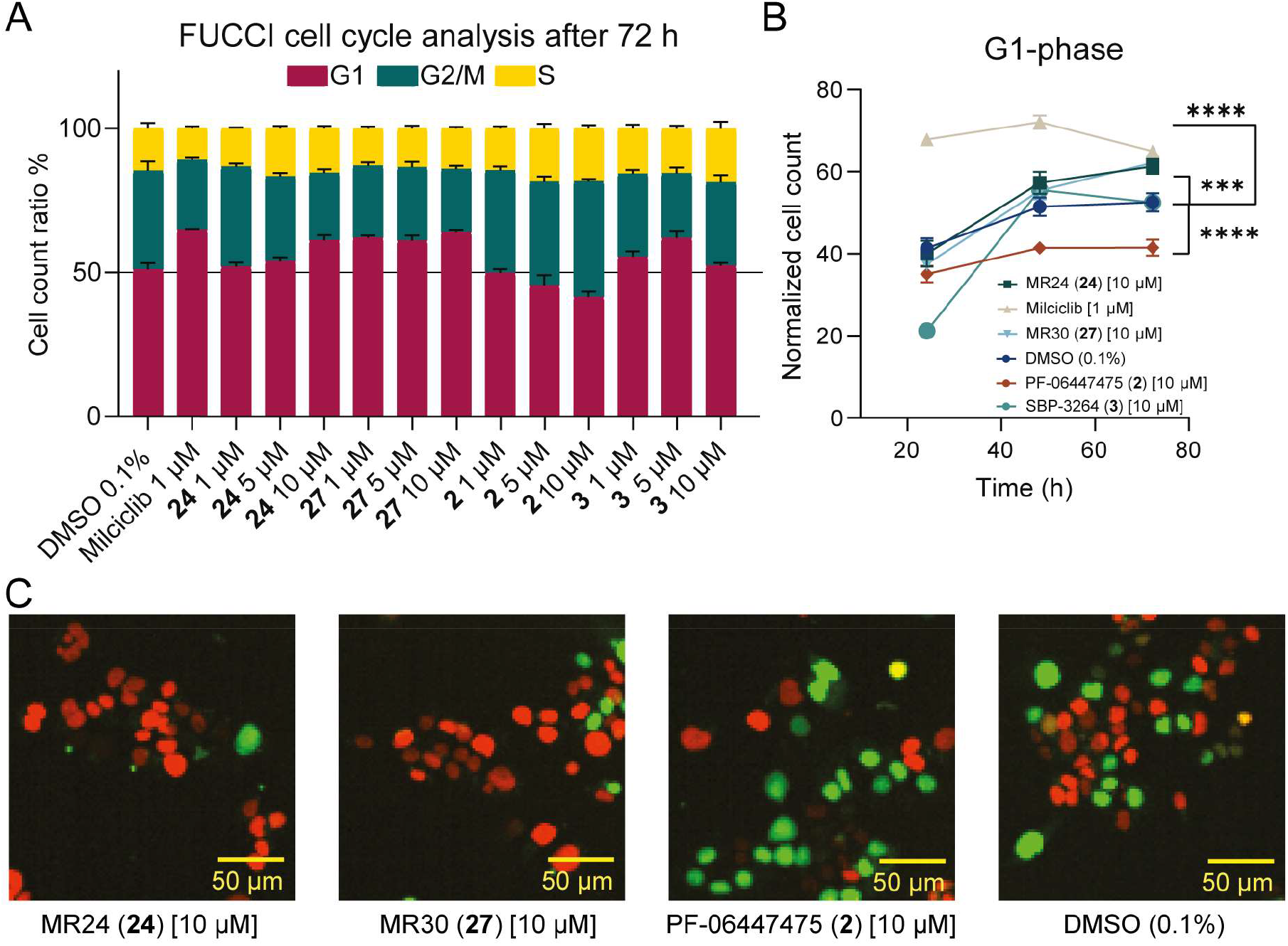
Cell-cycle analysis (FUCCI). (A) Fractions of red (G1), green (G2/M), or yellow (S) cells after 72 h of compound exposure (**2**, **3**, **24**, **27**, and milciclib) compared with cells exposed to DMSO. Error bars show SEM of two biological replicates. (B) Normalized cell count of cells in G1-phase after 24 h, 48 h, and 72 h of compound exposure (**2**, **3**, **24**, and **27**) at 10 µM compared with cells exposed to DMSO and cells exposed to milciclib (1 µM). Error bars show SEM of two biological replicates. Significance was calculated using a two-way ANOVA analysis for the 48 time-point. **** indicates a P value < 0.0001, *** indicates a P value < 0.002 (C) Fluorescence image of HCT116-FUCCI cells after 48 h of compound exposure (**2**, **24**, and **27**) in comparison with cells exposed to DMSO (0.1%). All data can be found in Table S9.

### Microsomal stability and prediction of pharmacokinetic properties

To assess the potential use of MR24 (**24**) *in vivo*, the compound was further profiled for its stability on activated liver microsomes of Sprague-Dawley rats. MR24 (**24**) exhibited an excellent microsomal stability of more than 90% over a time of 1 h, determined via high-performance liquid chromatography (HPLC) (Figure S10). In addition, the pharmacokinetic properties of MR24 (**24**) were calculated (QikProp; *Schrödinger*) and compared with the predicted values for MR30 (**27**) and MRLW5 (**20**) and the published values for the lead structure G-5555 (**8**).^40^ The prediction for MR24 (**24**) and MR30 (**27**) suggested a lower cell permeability in CACO-2 and HDCK cells compared with G-5555 (**8**). However, in our cellular NanoBRET binding assays for these compounds in HEK293T cells, similar EC50s for permeabilized and intact cells were determined indicating that cell permeability was adequate. The human oral bioavailability was predicted to be 52% (MR24 (**24**)) and 61% (MR30 (**27**)), respectively.

## SUMMARY AND CONCLUSION

The GCKII subfamily, consisting of MST1/2, and the structurally closely related GCKIII subfamily to which MST3/4 have been assigned, are involved in the activation of tumor suppressors and the assembly of transcription factors that, when dysregulated, can lead to malignant transformation. Although MST kinases share high sequence similarity, the associated biological pathways of these two subfamilies may differ considerably. It is, therefore, important to study the differences and specific functions of the two subfamilies to gain a comprehensive understanding of their cellular mechanisms of action. Mechanistic studies are best performed with selective inhibitors that are well profiled and annotated against a large number of potential targets. Several MST1/2 inhibitors such as XMU-MP-1 (**1**), PF-06447475 (**2**), and SBP-3264 (**3**) have been used for target validation, and the roles of both kinases are therefore well understood, which is, however, not the case for the closely related kinases MST3/4. One way to overcome the lack of inhibitors with kinome-wide selectivity is the use of chemogenomic compounds that inhibit several identical or structurally related targets and can be used in parallel for target deconvolution in phenotypic screens. A re-evaluation of published MST1/2 inhibitors identified the LRRK2 inhibitor PF-06447475 (**2**) as a good tool compound targeting MST1/2. PF-06447475 (**2**) showed activity against MST1/2 in the low nanomolar range in cells and good kinome-wide selectivity. To develop a tool compound for studying MST3/4, we optimized the selectivity of the pyrido[2,3-*d*]pyrimidin-7(8*H*)-one-based p21-activated kinase (PAK) inhibitor G-5555 (**8**) for those kinases and eliminated its residual activity on SIK kinases in the process. A systematic structural analysis of the key kinases targeted by the lead structure (MSTs, PAKs, and SIKs) suggested optimizing the pyrido[2,3-*d*]pyrimidin-7(8*H*)-one scaffold at position 8 for increased potency and modifying the front-pocket targeting moiety of the inhibitor (position 2) for improved selectivity. The SAR analysis was supported by high-resolution crystal structures, MD simulations, and molecular mechanics calculations (MM/GBSA). This resulted in compounds MR24 (**24**), MR26 (**26**), and MR30 (**27**), which exhibited the best balance between cellular on-target potency against MST3/4 and overall selectivity. Importantly, neither MR24 (**24**) nor MR30 (**27**) showed significant binding to SIK kinases. In cellular assays, the compounds had no adverse effect on cell health when treating HCT116 cells at a concentration of 1 µM.

The pivotal role of the Hippo signaling pathway on mitotic cell-cycle progression^55^ led us to perform a fluorescent ubiquitination-based cell-cycle indicator (FUCCI) assay^56^ to investigate the effect of the compounds on the cell cycle. Consistent with the key role of MST3/4 in the Hippo signaling pathway and mitotic cell-cycle progression, MR24 (**24**) and MR30 (**27**) induced cell-cycle arrest in G1 phase, supporting their use in phenotypic assays investigating the function of MST3/4. Based on this profiling, we propose the use of compounds MR24 (**24**) and MR30 (**27**) together with PF-06447475 (**2**) and the PAK1 allosteric chemical probe NVS-PAK1-1,^60^ to allow to account for off-target effects of class I PAKs in the study of the GCK subfamily, which in combination provides a valuable tool for elucidating the multifaceted roles of MST kinases.

## EXPERIMENTAL SECTION

### Chemistry

The synthetic procedure of compounds **9**–**70** will be explained in the following. Further information regarding the analytical data for **9**–**70** can be found in the Supporting Information. All commercial chemicals were purchased from common suppliers in reagent grade and used without further purification. Reactions were performed under argon atmosphere. For compound purification by column chromatography, silica gel with a particle size of 40-63 μm (*Machery-Nagel GmbH & Co.KG*) was used. Purification by flash chromatography was performed on a puriFlash XS 420 device with a UV-VIS multi-wave detector (200-400 nm) with prepacked normal phase PF-SIHP silica columns with a particle size of 30 µm (*Interchim*). All compounds synthesized were characterized by ^1^H/^13^CNMR and mass spectrometry (ESI +/-). In addition, final compounds were identified by HRMS, and their purity was evaluated by HPLC. All compounds used for further testing showed > 95% purity. ^1^H and ^13^CNMR spectra were measured on a DPX250, an AV400, an AV500 HD Avance III and an AV600 spectrometer from *Bruker Corporation*. Chemical shifts (*δ*) are reported in parts per million (ppm) and coupling constants (*J*) in hertz. DMSO-d6 was used as solvent, and the spectra were calibrated to the solvent signal: 2.50 ppm (^1^H NMR) or 39.52 ppm (^13^CNMR). Mass spectra (ESI +/-) were measured on a Surveyor MSQ device from *Thermo Fisher Scientific*. High-resolution mass spectra (HRMS) were obtained on a MALDI LTQ Orbitrap XL, also from *Thermo Fischer Scientific*. Preparative purification by HPLC was carried out on an *Agilent* 1260 Infinity II device using an Eclipse XDB-C18 (*Agilent*, 21.2 x 250mm, 7µm) reversed-phase column. A suitable gradient (flow rate 21 ml/min.) was used, with water (A; 0.1% TFA) and acetonitrile (B; 0.1% TFA) as mobile phase. All compounds purified by preparative HPLC chromatography were obtained as TFA salt. Determination of the compound purity by HPLC was carried out on the same *Agilent* 1260 Infinity II device together with HPLC-MS measurements using an LC/MSD device (G6125B, ESI pos. 100-1000). The purity of the final compounds was either analyzed on an Eclipse XDB-C18 (*Agilent*, 4.6 x 250mm, 5µm) reversed-phase column, with water (A; 0.1% TFA) and acetonitrile (B; 0.1% TFA) as a mobile phase, using the following gradient: 0 min. 2% B - 2 min. 2% B - 10 min. 98% B - 15 min. 98% B - 17 min. 2% B - 19 min. 2% B (flow rate of 1 mL/min.) (= Method A), or on a Poroshell 120 EC-C18 (*Agilent*, 3 x 150 mm, 2.7 µm) reversed phase column, with water (A; 0.1% formic acid) and acetonitrile (B; 0.1% formic acid) as a mobile phase, using the following gradient: 0 min. 5% B - 2 min. 5% B - 8 min. 98% B - 10 min. 98% B (flow rate of 0.5 mL/min.) (= Method B). UV-detection was performed at 254, or 280, and 310 nm.

### General procedure I for nucleophilic substitution reaction

6-(2-Chloro-4-(6-methylpyridin-2-yl)phenyl)-2-(methylthio)pyrido[2,3-*d*]pyrimidin-7-ol (**41**; 1 eq), cesium carbonate (3.0 eq) and the corresponding primary alkyl halide (1.0 or 1.5 eq) were dissolved in anhydrous DMF (20 mL) and stirred at 110 °C for 18 hours. The solvent was evaporated under reduced pressure, and the remaining residue was purified by column or flash chromatography on silica gel using DCM/MeOH as eluent.

### General procedure II for oxidation of the methyl sulfide to a sulfoxide/sulfone

The corresponding methyl sulfide derivative (1 eq.) was dissolved in anhydrous DCM (10 mL). *M*-chloroperoxybenzoic acid (*m*-CPBA) (≤ 77%, 2.1 eq) was added in one portion, and the reaction solution was stirred at room temperature for up to 3 h. The reaction was monitored by HPLC analysis and stopped after complete transformation by adding saturated aq. NaHCO3 solution (10 mL). The aqueous layer was extracted three times with a mixture of DCM/MeOH (4:1; 15 mL) and DCM (15 mL). The combined organic layers were dried over Na2SO4, and the solvent was evaporated under reduced pressure. The remaining residue, consisting of the corresponding sulfoxide and sulfone products, was used in the next step without further purification.

### General procedure III for nucleophilic aromatic substitution reaction

The remaining residue of the corresponding product obtained from general procedure II (1.0 eq) was dissolved in ethanol (10 mL), and a solution of methylamine in THF (2M, 4.0 eq) and *N*,*N’*-diisopropylethylamine (5.0 eq) was added. The reaction solution was stirred at 85 °C for 17 h. Afterwards the solvent was evaporated under reduced pressure. The remaining residue was purified either by flash chromatography on silica gel using DCM/MeOH as eluent, or by HPLC chromatography on silica gel (H2O/ACN; 0.1% TFA).

### General procedure IV for deprotection of BOC-protected amines

The corresponding BOC-protected amine was dissolved in anhydrous DCM (5 mL), and TFA (20 vol%) was added. The reaction solution was stirred at room temperature for 30 min. Then, the reaction solution was evaporated under reduced pressure. If necessary, the remaining residue was purified either by flash chromatography on silica gel using DCM/MeOH as eluent, or by HPLC chromatography on silica gel (H2O/ACN; 0.1% TFA). Compounds obtained from this reaction were isolated as TFA salt.

#### Ethyl 2-(4-bromo-2-chloro-phenyl)acetate (**38**)

The synthesis was performed as previously published.^40^

#### Ethyl 2-(2-chloro-4-(4,4,5,5-tetramethyl-1,3,2-dioxaborolan-2-yl)phenyl)acetate) (**39**)

Ethyl 2-(4-bromo-2-chlorophenyl)acetate (**38**; 200 mg, 0.72 mmol), bis(pinacolato)diboron (238 mg, 0.94 mmol), potassium acetate (212 mg, 2.16 mmol) and [1,1’-bis(diphenylphosphino)ferrocene]dichloropalladium(II) in complex with DCM (41 mg, 0.05 mmol) were dissolved in anhydrous dioxane (6 mL) and stirred at 115 °C for 18 h. The reaction solution was filtered over a pad of Celite^®^. Ethyl acetate (20 mL) was added, and the organic layer was washed with brine (20 mL). The aqueous layer was extracted twice with ethyl acetate (20 mL), and the combined organic layers were dried over Na2SO4. The solvent was evaporated under reduced pressure, and the remaining residue was purified by flash chromatography on silica gel using cyclohexane and ethyl acetate as eluent. **39** was obtained as a colorless oil in a yield of 90% (210 mg). ^1^H NMR (500 MHz, DMSO-d6, 300 K): *δ* = 7.62 (s, 1H), 7.57 (dd, *J* = 7.5, 0.9 Hz, 1H), 7.43 (d, *J* = 7.5 Hz, 1H), 4.09 (q, *J* = 7.1 Hz, 2H), 3.83 (s, 2H), 1.30 (s, 12H), 1.17 (t, *J* = 7.1 Hz, 3H) ppm. ^13^CNMR (126 MHz, DMSO-d6, 300 K): *δ* = 169.8, 136.0, 134.4, 133.7, 133.0, 132.0, 127.9, 84.1 (2C), 60.5, 38.7 (4C), 24.6, 14.1 ppm. MS (ESI+): m/z = 347.0 [M + Na]^+^; calculated: 347.1. *Ethyl 2-(2-chloro-4-(6-methylpyridin-2-yl)phenyl)acetate* (**40**). **39** (2.09 g, 6.44 mmol), potassium acetate (1.26 g, 12.88 mmol), [1,1′-Bis(diphenylphosphino)-ferrocene]dichloropalladium(II) (0.33 g, 0.45 mmol) and 2-bromo-6-methylpyridine (1.11 g, 6.44 mmol) were dissolved in a mixture of dioxane/water (2:1; 30 mL) and stirred at 107 °C for 18 hours. The solvent was evaporated under reduced pressure, and the remaining residue was purified by column chromatography on silica gel using cyclohexane/ethyl acetate as eluent. **40** was obtained as a pale yellow oil in a yield of 86% (1.60 g). ^1^H NMR (500 MHz, DMSO-d6, 300 K): *δ* = 8.15 (d, *J* = 1.5 Hz, 1H), 8.00 (dd, *J* = 8.0, 1.6 Hz, 1H), 7.82 - 7.75 (m, 2H), 7.51 (d, *J* = 8.0 Hz, 1H), 7.25 (d, *J* = 7.2 Hz, 1H), 4.12 (q, *J* = 7.1 Hz, 2H), 3.85 (s, 2H), 2.54 (s, 3H), 1.19 (t, *J* = 7.1 Hz, 3H) ppm. ^13^CNMR (126 MHz, DMSO-d6, 300 K): *δ* = 169.9, 157.9, 153.4, 139.6, 134.2, 133.2, 132.4, 126.7, 125.0, 122.5, 117.4, 60.5, 38.3, 24.3, 14.0 ppm. MS (ESI+): m/z = 290.0 [M + H]^+^; calculated: 290.1.

#### 6-(2-Chloro-4-(6-methylpyridin-2-yl)phenyl)-2-(methylthio)pyrido[2,3-d]pyrimidin-7-ol (**41**)

**40** (0.92 g, 3.19 mmol), 4-amino-2-(methylthio)-pyrimidine-5-carbaldehyde (0.54 g, 3.19 mmol) and potassium carbonate (1.32 g, 9.57 mmol) were dissolved in anhydrous DMF (70 mL) and stirred at 120 °C for 16 h. Water (300 mL) was added to the reaction solution, leading to the precipitation of **41**. After filtration, the aqueous layer was extracted three times with ethyl acetate (30 mL), and the combined organic layers were dried over MgSO4. The solvent was evaporated under reduced pressure, and the remaining residue was recrystallized from acetone. **41** was obtained as a white solid in a combined yield of 61% (0.77 g). ^1^H NMR (500 MHz, DMSO-d6, 300 K): *δ* = 12.67 (s, 1H), 8.92 (s, 1H), 8.24 (d, *J* = 1.7 Hz, 1H), 8.10 (dd, *J* = 8.0, 1.7 Hz, 1H), 8.01 (s, 1H), 7.88 (d, *J* = 7.8 Hz, 1H), 7.81 (t, *J* = 7.7 Hz, 1H), 7.53 (d, *J* = 8.0 Hz, 1H), 7.28 (d, *J* = 7.5 Hz, 1H), 2.59 (s, 3H), 2.57 (s, 3H) ppm. ^13^CNMR (126 MHz, DMSO-d6, 300 K): *δ* = 172.2, 161.3, 158.0, 156.9, 154.2, 153.3, 140.4, 137.7, 136.5, 135.0, 133.4, 132.1, 131.4, 126.9, 124.9, 122.7, 117.7, 108.8, 24.3, 13.7 ppm. MS (ESI+): m/z = 395.0 [M + H]^+^; calculated: 395.0.

#### 6-(2-Chloro-4-(6-methylpyridin-2-yl)phenyl)-8-methyl-2-(methylthio)pyrido[2,3-d]pyrimidin-7(8H)-one (**42**)

Compound **42** was prepared according to general procedure I, using **41** (150 mg, 0.38 mmol) and a solution of iodomethane in tert-butyl methyl ether (2M; 81 mg, 0.57 mmol) as starting materials. After purification by column chromatography on silica gel, a trituration from DCM/cyclohexane was performed to obtain **42** as a yellow solid in a yield of 67% (103 mg). ^1^H NMR (500 MHz, DMSO-d6, 300 K): *δ* = 8.95 (s, 1H), 8.25 (d, *J* = 1.6 Hz, 1H), 8.12 (dd, *J* = 8.0, 1.7 Hz, 1H), 8.06 (s, 1H), 7.89 (d, *J* = 7.8 Hz, 1H), 7.82 (t, *J* = 7.7 Hz, 1H), 7.53 (d, *J* = 8.0 Hz, 1H), 7.29 (d, *J* = 7.6 Hz, 1H), 3.68 (s, 3H), 2.64 (s, 3H), 2.57 (s, 3H) ppm. ^13^CNMR (126 MHz, DMSO-d6, 300 K): *δ* = 172.1, 160.8, 158.0, 157.3, 153.8, 153.2, 140.4, 137.7, 135.6, 135.3, 133.4, 132.1, 130.2, 126.9, 124.9, 122.7, 117.7, 109.2, 28.0, 24.3, 13.9 ppm. MS (ESI+): m/z = 409.1 [M + H]^+^; calculated: 409.1.

#### 6-(2-Chloro-4-(6-methylpyridin-2-yl)phenyl)-8-(2-methoxyethyl)-2-(methylthio)pyrido[2,3-d]pyrimidin-7(8H)-one (**43**)

Compound **43** was prepared according to general procedure I, using **41** (150 mg, 0.38 mmol) and bromo-2-methoxyethane (79 mg, 0.57 mmol) as starting materials. After purification by column chromatography on silica gel, a trituration from DCM/cyclohexane was performed to obtain **43** as a yellow solid in a yield of 63% (108 mg). ^1^H NMR (500 MHz, DMSO-d6, 300 K): *δ* = 8.95 (s, 1H), 8.25 (d, *J* = 1.7 Hz, 1H), 8.12 (dd, *J* = 8.0, 1.7 Hz, 1H), 8.07 (s, 1H), 7.88 (d, *J* = 7.8 Hz, 1H), 7.81 (t, *J* = 7.7 Hz, 1H), 7.54 (d, *J* = 8.0 Hz, 1H), 7.29 (d, *J* = 7.6 Hz, 1H), 4.58 (t, *J* = 6.2 Hz, 2H), 3.67 (t, *J* = 6.2 Hz, 2H), 3.28 (s, 3H), 2.63 (s, 3H), 2.57 (s, 3H) ppm. ^13^CNMR (126 MHz, DMSO-d6, 300 K): *δ* = 172.1, 160.4, 158.0, 157.5, 153.7, 153.2, 140.4, 137.7, 135.8, 135.2, 133.4, 132.1, 130.2, 126.9, 124.9, 122.7, 117.7, 109.1, 68.3, 58.1, 26.3, 24.3, 13.9 ppm. MS (ESI+): m/z = 453.2 [M + H]^+^; calculated: 453.1.

#### 6-(2-Chloro-4-(6-methylpyridin-2-yl)phenyl)-8-(3-methoxypropyl)-2-(methylthio)-pyrido[2,3-d]pyrimidin-7(8H)-one (**44**)

Compound **44** was prepared according to general procedure I, using **41** (150 mg, 0.38 mmol) and 1-bromo-3-methoxypropane (87 mg, 0.57 mmol) as starting materials. After purification by column chromatography on silica gel, a trituration from DCM/cyclohexane was performed to obtain **44** as a yellow solid in a yield of 57% (100 mg). ^1^H NMR (500 MHz, DMSO-d6, 300 K): *δ* = 8.94 (s, 1H), 8.25 (s, 1H), 8.15 - 8.09 (m, 1H), 8.05 (d, *J* = 0.7 Hz, 1H), 7.88 (d, *J* = 7.8 Hz, 1H), 7.81 (t, *J* = 7.7 Hz, 1H), 7.54 (d, *J* = 8.0 Hz, 1H), 7.28 (d, *J* = 7.6 Hz, 1H), 4.45 (t, *J* = 7.1 Hz, 2H), 3.41 (t, *J* = 6.0 Hz, 2H), 3.20 (d, *J* = 0.8 Hz, 3H), 2.63 (s, 3H), 2.57 (s, 3H), 1.96 - 1.90 (m, 2H) ppm. ^13^CNMR (126 MHz, DMSO-d6, 300 K): *δ* = 172.2, 160.4, 158.0, 157.4, 153.5, 153.2, 140.4, 137.7, 135.6, 135.3, 133.4, 132.1, 130.3, 126.9, 124.9, 122.7, 117.7, 109.1, 69.8, 57.9, 38.7, 27.4, 24.3, 13.9 ppm. MS (ESI+): m/z = 467.2 [M + H]^+^; calculated: 467.1.

#### 6-(2-Chloro-4-(6-methylpyridin-2-yl)phenyl)-8-(4-methoxybutyl)-2-(methylthio)pyrido[2,3-d]pyrimidin-7(8H)-one (**45**)

Compound **45** was prepared according to general procedure I, using **41** (150 mg, 0.38 mmol) and 1-bromo-4-methoxybutane (95 mg, 0.57 mmol) as starting materials. After purification by column chromatography on silica gel, a trituration from DCM/cyclohexane was performed to obtain **45** as a yellow solid in a yield of 73% (133 mg). ^1^H NMR (500 MHz, DMSO-d6, 300 K): *δ* = 8.94 (s, 1H), 8.25 (d, *J* = 1.6 Hz, 1H), 8.12 (dd, *J* = 8.0, 1.7 Hz, 1H), 8.05 (s, 1H), 7.88 (d, *J* = 7.8 Hz, 1H), 7.81 (t, *J* = 7.7 Hz, 1H), 7.54 (d, *J* = 8.0 Hz, 1H), 7.29 (d, *J* = 7.6 Hz, 1H), 4.41 - 4.36 (m, 2H), 3.35 (t, *J* = 6.3 Hz, 2H), 3.21 (s, 3H), 2.63 (s, 3H), 2.57 (s, 3H), 1.78 - 1.71 (m, 2H), 1.61 - 1.54 (m, 2H) ppm. ^13^CNMR (126 MHz, DMSO-d6, 300 K): *δ* = 172.1, 160.4, 158.0, 157.5, 153.4, 153.2, 140.4, 137.7, 135.6, 135.3, 133.4, 132.1, 130.3, 126.9, 124.9, 122.7, 117.7, 109.1, 71.5, 57.8, 40.7, 26.6, 24.3, 24.2, 13.9 ppm. MS (ESI+): m/z = 481.2 [M + H]^+^; calculated: 481.1.

#### 6-(2-Chloro-4-(6-methylpyridin-2-yl)phenyl)-8-(2-hydroxyethyl)-2-(methylthio)pyrido[2,3-d]pyrimidin-7(8H)-one (**46**)

Compound **46** was prepared according to general procedure I, using **41**(150 mg, 0.38 mmol) and 2-bromoethan-1-ol (71 mg, 0.57 mmol) as starting materials. After purification by column chromatography on silica gel, a trituration from DCM/cyclohexane was performed to obtain **46** as a yellow solid in a yield of 58% (97 mg). ^1^H NMR (500 MHz, DMSO-d6, 300 K): *δ* = 8.94 (s, 1H), 8.25 (d, *J* = 1.7 Hz, 1H), 8.12 (dd, *J* = 8.0, 1.7 Hz, 1H), 8.06 (s, 1H), 7.88 (d, *J* = 7.8 Hz, 1H), 7.81 (t, *J* = 7.7 Hz, 1H), 7.54 (d, *J* = 8.0 Hz, 1H), 7.29 (d, *J* = 7.6 Hz, 1H), 4.88 (t, *J* = 5.9 Hz, 1H), 4.49 (t, *J* = 6.6 Hz, 2H), 3.71 - 3.66 (m, 2H), 2.63 (s, 3H), 2.57 (s, 3H) ppm. ^13^CNMR (126 MHz, DMSO-d6, 300 K): *δ* = 172.0, 160.4, 157.9, 157.3, 153.7, 153.1, 140.3, 137.6, 135.6, 135.2, 133.3, 132.0, 130.1, 126.8, 124.8, 122.6, 117.6, 109.0, 57.5, 42.8, 24.2, 13.9 ppm. MS (ESI+): m/z = 439.1 [M + H]^+^; calculated: 439.1.

#### 6-(2-Chloro-4-(6-methylpyridin-2-yl)phenyl)-8-(3-hydroxypropyl)-2-(methylthio)pyrido[2,3-d]pyrimidin-7(8H)-one (**47**)

Compound **47** was prepared according to general procedure I, using **41** (150 mg, 0.38 mmol) and 3-bromopropan-1-ol (79 mg, 0.57 mmol) as starting materials. After purification by column chromatography on silica gel an additional purification by HPLC chromatography on silica gel (H2O/ACN; 0.1% TFA) was performed, to obtain **47** as a yellow solid in a yield of 25% (64 mg). ^1^H NMR (500 MHz, DMSO-d6, 300 K): *δ* = 8.95 (s, 1H), 8.24 (d, *J* = 1.6 Hz, 1H), 8.09 (d, *J* = 1.5 Hz, 1H), 8.06 (s, 1H), 7.97 - 7.96 (m, 2H), 7.58 (d, *J* = 8.0 Hz, 1H), 7.43 - 7.41 (m, 1H), 4.52 (t, *J* = 6.7 Hz, 1H), 4.48 - 4.44 (m, 2H), 3.51 (t, *J* = 6.4 Hz, 2H), 2.64 (s, 3H), 2.62 (s, 3H), 1.91 - 1.82 (m, 2H) ppm. ^13^CNMR (126 MHz, DMSO-d6, 300 K): *δ* = 172.3, 160.4, 157.5, 157.4, 153.5, 152.6, 139.4, 138.9, 135.9, 135.7, 133.5, 132.2, 130.2, 127.4, 125.4, 123.5, 118.9, 109.1, 58.8, 38.8, 30.8, 23.4, 13.9 ppm. MS (ESI+): m/z = 453.2 [M + H]^+^; calculated: 453.1.

#### 6-(2-Chloro-4-(6-methylpyridin-2-yl)phenyl)-8-(4-hydroxybutyl)-2-(methylthio)pyrido[2,3-d]pyrimidin-7(8H)-one (**48**)

Compound **48** was prepared according to general procedure I, using **41** (150 mg, 0.38 mmol) and 4-bromobutan-1-ol (87 mg, 0.57 mmol) as starting materials. After purification by column chromatography on silica gel, **48** was obtained as a white solid in a yield of 57% (100 mg). ^1^H NMR (500 MHz, DMSO-d6, 300 K): *δ* = 8.93 (s, 1H), 8.24 (d, *J* = 1.7 Hz, 1H), 8.10 (dd, *J* = 8.0, 1.8 Hz, 1H), 8.04 (s, 1H), 7.87 (d, *J* = 7.8 Hz, 1H), 7.80 (t, *J* = 7.7 Hz, 1H), 7.53 (d, *J* = 8.0 Hz, 1H), 7.28 (d, *J* = 7.6 Hz, 1H), 4.42 (t, *J* = 5.2 Hz, 1H), 4.40 - 4.35 (m, 2H), 3.46 - 3.42 (m, 2H), 2.62 (s, 3H), 2.56 (s, 3H), 1.77 - 1.70 (m, 2H), 1.52 - 1.47 (m, 2H) ppm. ^13^CNMR (126 MHz, DMSO-d6, 300 K): *δ* = 172.2, 160.3, 158.0, 157.4, 153.4, 153.2, 140.4, 137.7, 135.6, 135.3, 133.4, 132.0, 130.3, 126.9, 124.9, 122.7, 117.6, 109.1, 60.4, 40.8, 30.0, 24.3, 24.2, 13.9 ppm. MS (ESI+): m/z = 467.2 [M + H]^+^; calculated: 467.1.

#### 6-(2-Chloro-4-(6-methylpyridin-2-yl)phenyl)-8-(5-hydroxypentyl)-2-(methylthio)pyrido[2,3-d]pyrimidin-7(8H)-one (**49**)

Compound **49** was prepared according to general procedure I, using **41** (150 mg, 0.38 mmol) and 5-bromopentan-1-ol (95 mg, 0.57 mmol) as starting materials. After purification by column chromatography on silica gel, a trituration from DCM/cyclohexane was performed to obtain **49** as a yellow solid in a yield of 53% (97 mg). ^1^H NMR (500 MHz, DMSO-d6, 300 K): *δ* = 8.94 (s, 1H), 8.25 (d, *J* = 1.7 Hz, 1H), 8.12 (dd, *J* = 8.0, 1.8 Hz, 1H), 8.06 (s, 1H), 7.89 (d, *J* = 7.8 Hz, 1H), 7.82 (t, *J* = 7.7 Hz, 1H), 7.54 (d, *J* = 8.0 Hz, 1H), 7.29 (d, *J* = 7.6 Hz, 1H), 4.37 - 4.34 (m, 2H), 4.08 - 4.01 (m, 1H), 3.42 - 3.36 (m, 2H), 2.63 (s, 3H), 2.57 (s, 3H), 1.73 - 1.68 (m, 2H), 1.51 - 1.45 (m, 2H), 1.42 - 1.36 (m, 2H) ppm. ^13^CNMR (126 MHz, DMSO-d6, 300 K): *δ* = 172.2, 160.3, 158.3, 158.0, 157.8, 157.5, 153.4, 153.0, 138.4, 135.6, 133.5, 132.1, 130.3, 127.1, 125.1, 123.1, 118.1, 109.1, 60.5, 40.9, 32.1, 27.1, 24.0, 23.0, 13.9 ppm. MS (ESI+): m/z = 481.2 [M + H]^+^; calculated: 481.1.

#### 6-(2-Chloro-4-(6-methylpyridin-2-yl)phenyl)-2-(methylamino)pyrido[2,3-d]pyrimidin-7-ol (**9**)

Compound **9** was prepared according to general procedures II and III, with an addition of anhydrous DMF (5 mL) to improve the solubility during the reactions. **41** (100 mg, 0.25 mmol) was used as starting material. The crude product was purified by flash chromatography on silica gel using DCM/MeOH as eluent. A following recrystallization from acetone was performed to obtain **9** as a white solid in a yield of 89% (80 mg) over two steps. ^1^H NMR (500 MHz, DMSO-d6, 300 K): *δ* = 12.11 - 11.88 (m, 1H), 8.72 - 8.54 (m, 1H), 8.21 (d, *J* = 1.5 Hz, 1H), 8.07 (dd, *J* = 8.0, Hz, 1H), 7.86 (d, *J* = 7.8 Hz, 1H), 7.84 - 7.77 (m, 2H), 7.72 (s, 1H), 7.49 (d, *J* = 8.0 Hz, 1H), 7.28 (d, *J* = 7.5 Hz, 1H), 2.88 (d, *J* = 4.8 Hz, 3H), 2.57 (s, 3H) ppm. ^13^CNMR (126 MHz, DMSO-d6, 300 K): *δ* = 162.3, 161.8, 158.8, 158.0, 155.8, 153.4, 139.9, 137.6, 137.4, 135.8, 133.7, 132.3, 126.8, 125.4, 124.8, 122.6, 117.6, 103.7, 27.9, 24.3 ppm. HRMS (FTMS +p MALDI): m/z = 400.0945 [M + Na]^+^; calculated for C20H16ClN5ONa: 400.0936. HPLC: tR = 7.13 min; purity ≥ 95% (UV: 280/310 nm). LCMS (ESI+): m/z = 378.2 [M + H]^+^; calculated: 378.1.

#### 6-(2-Chloro-4-(6-methylpyridin-2-yl)phenyl)-8-methyl-2-(methylamino)pyrido[2,3-d]pyrimidin-7(8H)-one (**10**)

Compound **10** was prepared according to general procedures II and III, with an addition of anhydrous DMF (2 mL) to improve the solubility during the reactions. **42** (100 mg, 0.25 mmol) was used as starting material. After purification by column chromatography on silica gel using DCM/MeOH as eluent, an additional purification by HPLC chromatography on silica gel (H2O/ACN; 0.1% TFA) was performed to obtain **10** as a yellow solid (TFA salt) in a yield of 71% (90 mg) over two steps. ^1^H NMR (500 MHz, DMSO-d6, 300 K): *δ* = 8.75 - 8.64 (m, 1H), 8.22 (d, *J* = 1.7 Hz, 1H), 8.06 (dd, *J* = 8.0, 1.7 Hz, 1H), 7.98 - 7.88 (m, 3H), 7.85 (s, 1H), 7.53 (d, *J* = 8.0 Hz, 1H), 7.40 - 7.35 (m, 1H), 3.66 - 3.53 (m, 3H), 2.94 (s, 3H), 2.60 (s, 3H) ppm. ^13^CNMR (126 MHz, DMSO-d6, 300 K): *δ* = 161.7, 161.2, 159.1, 157.5, 155.4, 152.9, 138.8, 136.5, 136.3, 133.7, 132.4, 127.2, 125.1, 123.2, 122.6, 122.8, 118.6, 108.9, 27.9, 27.6, 23.7 ppm. HRMS (FTMS +p MALDI): m/z = 392.1273 [M + H]^+^; calculated for C21H19ClN5O: 392.1273. HPLC: tR = 11.34 min; purity ≥ 95% (UV: 254/280 nm). MS (ESI+): m/z = 392.1 [M + H]^+^; calculated: 392.1.

#### 6-(2-Chloro-4-(6-methylpyridin-2-yl)phenyl)-8-(2-methoxyethyl)-2-(methylamino)-pyrido[2,3-d]pyrimidin-7(8H)-one (**11**)

Compound **11** was prepared according to general procedures II and III. **43** (100 mg, 0.22 mmol) was used as starting material. The crude product was purified by flash chromatography on silica gel using DCM/MeOH as eluent to obtain **11** as a yellow solid in a yield of 36% (35 mg) over two steps. ^1^H NMR (500 MHz, DMSO-d6, 300 K): *δ* = 8.73 - 8.63 (m, 1H), 8.22 (d, *J* = 1.7 Hz, 1H), 8.07 (dd, *J* = 8.0, 1.8 Hz, 1H), 7.89 (d, *J* = 4.4 Hz, 1H), 7.87 - 7.82 (m, 2H), 7.80 (t, *J* = 7.7 Hz, 1H), 7.50 (d, *J* = 8.0 Hz, 1H), 7.27 (d, *J* = 7.5 Hz, 1H), 4.56 - 4.45 (m, 2H), 3.69 - 3.59 (m, 2H), 3.31 - 3.27 (m, 3H), 2.92 (d, *J* = 4.7 Hz, 3H), 2.56 (s, 3H) ppm. ^13^CNMR (126 MHz, DMSO-d6, 300 K): *δ* = 161.9, 160.9, 159.4, 158.0, 155.2, 153.4, 139.9, 137.6, 136.6, 136.1, 133.7, 132.3, 126.8, 124.8, 124.3, 122.6, 117.6, 103.9, 68.2, 57.9, 28.0, 27.9, 24.3 ppm. HRMS (FTMS +p MALDI): m/z = 436.1539 [M + H]^+^; calculated for C23H23ClN5O2: 436.1535. HPLC: tR = 11.36 min; purity ≥ 95% (UV: 254/280 nm). LCMS (ESI+): m/z = 436.1 [M + H]^+^; calculated: 436.2.

#### 6-(2-Chloro-4-(6-methylpyridin-2-yl)phenyl)-8-(3-methoxypropyl)-2-(methylamino)-pyrido[2,3-d]pyrimidin-7(8H)-one (**12**)

Compound **12** was prepared according to general procedures II and III. **44** (100 mg, 0.21 mmol) was used as starting material. After purification by flash chromatography on silica gel using DCM/MeOH as eluent, an additional purification by HPLC chromatography on silica gel (H2O/ACN; 0.1% TFA) was performed to obtain **12** as a yellow solid in a yield of 70% (67 mg) over two steps. ^1^H NMR (500 MHz, DMSO-d6, 300 K): *δ* = 8.75 - 8.63 (m, 1H), 8.21 (d, *J* = 1.7 Hz, 1H), 8.06 (dd, *J* = 8.0, 1.8 Hz, 1H), 7.97 - 7.86 (m, 3H), 7.84 (s, 1H), 7.54 (d, *J* = 8.0 Hz, 1H), 7.42 - 7.36 (m, 1H), 4.42 - 4.38 (m, 2H), 3.42 (t, *J* = 6.1 Hz, 2H), 3.22 (s, 3H), 2.93 (s, 3H), 2.61 (s, 3H), 1.97 - 1.86 (m, 2H) ppm. ^13^CNMR (126 MHz, DMSO-d6, 300 K): *δ* = 161.7, 160.9, 159.1, 157.5, 155.1, 152.8, 139.1, 138.6, 136.6, 136.4, 133.8, 132.4, 127.3, 125.2, 124.4, 123.3, 118.6, 104.0, 70.0, 57.8, 38.2, 27.8, 27.4, 23.5 ppm. HRMS (FTMS +p MALDI): m/z = 450.1690 [M + H]^+^; calculated for C24H24ClN5O2: 450.1691. HPLC: tR = 11.61 min; purity ≥ 95% (UV: 254/280 nm). MS (ESI+): m/z = 450.2 [M + H]^+^; calculated: 450.2.

#### 6-(2-Chloro-4-(6-methylpyridin-2-yl)phenyl)-8-(4-methoxybutyl)-2-(methylamino)-pyrido[2,3-d]pyrimidin-7(8H)-one (**13**)

Compound **13** was prepared according to general procedures II and III. **45** (100 mg, 0.21 mmol) was used as starting material. The crude product was purified by flash chromatography on silica gel using DCM/MeOH as eluent to obtain **13** as a yellow solid in a yield of 60% (57 mg) over two steps. ^1^H NMR (500 MHz, DMSO-d6, 300 K): *δ* = 8.73-8.61 (m, 1H), 8.21 (d, *J* = 1.7 Hz, 1H), 8.07 (dd, *J* = 8.0, 1.8 Hz, 1H), 7.89 - 7.83 (m, 2H), 7.83 - 7.77 (m, 2H), 7.50 (d, *J* = 8.0 Hz, 1H), 7.27 (d, *J* = 7.5 Hz, 1H), 4.39 - 4.24 (m, 2H), 3.36 (t, *J* = 6.4 Hz, 2H), 3.20 (s, 3H), 2.92 (d, *J* = 4.6 Hz, 3H), 2.56 (s, 3H), 1.79 - 1.65 (m, 2H), 1.62 - 1.52 (m, 2H) ppm. ^13^CNMR (126 MHz, DMSO-d6, 300 K): *δ* = 162.0, 160.8, 159.4, 158.0, 155.0, 153.4, 139.9, 137.6, 136.3, 136.2, 133.7, 132.3, 126.8, 124.8, 124.4, 122.6, 117.5, 103.9, 71.5, 57.8, 40.1, 27.8, 26.6, 24.3, 24.1 ppm. HRMS (FTMS +p MALDI): m/z = 464.1848 [M + H]^+^; calculated for C25H27ClN5O2: 464.1848. HPLC: tR = 11.67 min; purity ≥ 95% (UV: 254/280 nm). LCMS (ESI+): m/z = 464.1 [M + H]^+^; calculated: 464.2.

#### 6-(2-Chloro-4-(6-methylpyridin-2-yl)phenyl)-8-(2-hydroxyethyl)-2-(methylamino)-pyrido[2,3-d]pyrimidin-7(8H)-one (**14**)

Compound **14** was prepared according to general procedures II and III, with an addition of anhydrous DMF (2 mL) to improve the solubility during the reactions. **46** (64 mg, 0.15 mmol) was used as starting material. The crude product was purified by flash chromatography on silica gel using DCM/MeOH as eluent to obtain **14** as a white solid in a yield of 71% (44 mg) over two steps. ^1^H NMR (500 MHz, DMSO-d6, 300 K): *δ* = 8.76 - 8.58 (m, 1H), 8.25 - 8.20 (m, 1H), 8.07 (dd, *J* = 8.0, 1.7 Hz, 1H), 7.89 - 7.77 (m, 4H), 7.50 (d, *J* = 8.0 Hz, 1H), 7.27 (d, *J* = 7.5 Hz, 1H), 4.89 - 4.79 (m, 1H), 4.50 - 4.32 (m, 2H), 3.72 - 3.61 (m, 2H), 2.93 (d, *J* = 4.4 Hz, 3H), 2.56 (s, 3H) ppm. ^13^CNMR (126 MHz, DMSO-d6, 300 K): *δ* = 162.0, 161.0, 159.3, 158.0, 155.3, 153.4, 140.0, 137.7, 136.5, 136.2, 133.7, 132.3, 126.8, 124.8, 124.3, 122.6, 117.6, 104.0, 57.6, 42.3, 27.9, 24.3 ppm. HRMS (FTMS +p MALDI): m/z = 422.1381 [M + H]^+^; calculated for C22H21ClN5O2: 422.1378. HPLC: tR = 7.19 min; purity ≥ 95% (UV: 280/310 nm). LCMS (ESI+): m/z = 422.2 [M + H]^+^; calculated: 422.1.

#### 6-(2-Chloro-4-(6-methylpyridin-2-yl)phenyl)-8-(3-hydroxypropyl)-2-(methylamino)-pyrido[2,3-d]pyrimidin-7(8H)-one (**15**)

Compound **15** was prepared according to general procedures II and III, with an addition of anhydrous DMF (2 mL) to improve the solubility during the reactions. **47** (43 mg, 0.09 mmol) was used as starting material. The crude product was purified by flash chromatography on silica gel using DCM/MeOH as eluent to obtain **15** as a pale yellow solid in a yield of 56% (23 mg) over two steps. ^1^H NMR (500 MHz, DMSO-d6, 300 K): *δ* = 8.76 - 8.62 (m, 1H), 8.22 (d, *J* = 1.7 Hz, 1H), 8.08 (dd, *J* = 8.0, 1.8 Hz, 1H), 7.86 (d, *J* = 7.8 Hz, 2H), 7.83 - 7.78 (m, 2H), 7.50 (d, *J* = 8.0 Hz, 1H), 7.27 (d, *J* = 7.6 Hz, 1H), 4.51 - 4.43 (m, 1H), 4.42 - 4.27 (m, 2H), 3.54 - 3.45 (m, 2H), 2.92 (d, *J* = 4.6 Hz, 3H), 2.57 (s, 3H), 1.91 - 1.78 (m, 2H) ppm. ^13^CNMR (126 MHz, DMSO-d6, 300 K): *δ* = 162.0, 161.0, 159.4, 158.0, 155.0, 153.4, 139.9, • 137.6, 136.4, 136.2, 133.7, 132.3, 126.8, 124.8, 124.4, 122.6, 117.6, 104.0, 58.9, 38.1, 30.8, 27.8, 24.3 ppm. HRMS (FTMS +p MALDI): m/z = 436.1540 [M + H]^+^; calculated for C23H23ClN5O2: 436.1535. HPLC: tR = 7.35 min; purity ≥ 95% (UV: 280/310 nm). LCMS (ESI+): m/z = 436.1 [M + H]^+^; calculated: 436.2.

#### 6-(2-Chloro-4-(6-methylpyridin-2-yl)phenyl)-8-(4-hydroxybutyl)-2-(methylamino)-pyrido[2,3-d]pyrimidin-7(8H)-one (**16**)

Compound **16** was prepared according to general procedures II and III, with an addition of anhydrous DMF (2 mL) to improve the solubility during the reactions. **48** (72 mg, 0.15 mmol) was used as starting material. The crude product was purified by flash chromatography on silica gel using DCM/MeOH as eluent to obtain **16** as a pale yellow solid in a yield of 54% (37 mg) over two steps. ^1^H NMR (500 MHz, DMSO-d6, 300 K): *δ* = 8.73 - 8.64 (m, 1H), 8.21 (d, *J* = 1.7 Hz, 1H), 8.07 (dd, *J* = 8.0, 1.8 Hz, 1H), 7.97 - 7.73 (m, 5H), 7.51 (d, *J* = 8.0 Hz, 1H), 7.31 (d, *J* = 7.4 Hz, 1H), 4.36 - 4.27 (m, 2H), 3.44 (t, *J* = 6.3 Hz, 2H), 2.92 (d, *J* = 3.7 Hz, 3H), 2.58 (s, 3H), 1.76 - 1.68 (m, 2H), 1.52 - 1.47 (m, 2H) ppm. ^13^CNMR (126 MHz, DMSO-d6, 300 K): *δ* = 161.9, 160.9, 159.3, 157.8, 155.0, 153.2, 139.4, 138.1, 136.4, 136.3, 133.7, 132.3, 127.0, 124.9, 124.4, 122.8, 117.9, 103.9, 60.5, 40.3, 30.1, 27.8, 24.1 (2C) ppm. HRMS (FTMS +p MALDI): m/z = 450.1693 [M + H]^+^; calculated for C24H25ClN5O2: 450.1691. HPLC: tR = 7.43 min; purity ≥ 95% (UV: 280/310 nm). LCMS (ESI+): m/z = 450.0 [M + H]^+^; calculated: 450.2.

#### 6-(2-Chloro-4-(6-methylpyridin-2-yl)phenyl)-8-(5-hydroxypentyl)-2-(methylamino)-pyrido[2,3-d]pyrimidin-7(8H)-one (**17**)

Compound **17** was prepared according to general procedures II and III, with an addition of anhydrous DMF (2 mL) to improve the solubility during the reactions. **49** (83 mg, 0.15 mmol) was used as starting material. The crude product was purified by flash chromatography on silica gel using DCM/MeOH as eluent to obtain **17** as a pale yellow solid in a yield of 70% (56 mg) over two steps. ^1^H NMR (500 MHz, DMSO-d6, 300 K): *δ* = 8.73 - 8.63 (m, 1H), 8.22 (d, *J* = 1.7 Hz, 1H), 8.08 (dd, *J* = 8.0, 1.7 Hz, 1H), 7.89 - 7.79 (m, 4H), 7.50 (d, *J* = 8.0 Hz, 1H), 7.28 (d, *J* = 7.5 Hz, 1H), 4.35 - 4.24 (m, 3H), 3.41 - 3.37 (m, 2H), 2.92 (d, *J* = 4.7 Hz, 3H), 2.57 (s, 3H), 1.75 - 1.64 (m, 2H), 1.52 - 1.46 (m, 2H), 1.41 - 1.35 (m, 2H) ppm. ^13^CNMR (126 MHz, DMSO-d6, 300 K): *δ* = 162.0, 160.8, 159.4, 158.0, 155.0, 153.4, 139.9, 137.6, 136.3, 136.2, 133.7, 132.3, 126.8, 124.8, 124.4, 122.6, 117.6, 103.9, 60.6, 40.3, 32.1, 27.8, 27.0, 24.3, 23.1 ppm. HRMS (FTMS +p MALDI): m/z = 486.1667 [M + Na]^+^; calculated for C25H26ClN5O2Na: 486.1667. HPLC: tR = 7.79 min; purity ≥ 95% (UV: 280/310 nm). LCMS (ESI+): m/z = 464.2 [M + H]^+^; calculated: 464.2.

#### tert-Butyl (2-(6-(2-chloro-4-(6-methylpyridin-2-yl)phenyl)-2-(methylthio)-7-oxopyrido[2,3-d]pyrimidin-8(7H)-yl)ethyl)carbamate (**50**)

Compound **50** was prepared according to general procedure I, using **41** (150 mg, 0.38 mmol) and tert-butyl (2-bromoethyl)carbamate (128 mg, 0.57 mmol) as starting materials. The crude product was purified by flash chromatography on silica gel. **50** was obtained as a white solid in a yield of 74% (151 mg). ^1^H NMR (500 MHz, DMSO-d6, 300 K): *δ* = 8.93 (s, 1H), 8.25 (d, *J* = 1.7 Hz, 1H), 8.12 (dd, *J* = 8.1, 1.7 Hz, 1H), 8.06 (s, 1H), 7.89 (d, *J* = 7.8 Hz, 1H), 7.82 (t, *J* = 7.7 Hz, 1H), 7.53 (d, *J* = 8.0 Hz, 1H), 7.29 (d, *J* = 7.6 Hz, 1H), 6.90 (t, *J* = 6.1 Hz, 1H), 4.48 (t, *J* = 6.0 Hz, 2H), 3.37 - 3.32 (m, 2H), 2.65 (s, 3H), 2.57 (s, 3H), 1.29 (s, 9H) ppm. ^13^CNMR (126 MHz, DMSO-d6, 300 K): *δ* = 172.1, 160.6, 158.0, 157.3, 155.6, 153.9, 153.3, 140.4, 137.7, 135.7, 135.3, 133.4, 132.1, 130.1, 126.9, 124.8, 122.7, 117.7, 109.2, 77.5, 40.9, 37.5, 28.1 (3C), 24.3, 13.9 ppm. MS (ESI+): m/z = 438.2 [M-BOC+ H]^+^; calculated: 438.2.

#### tert-Butyl (3-(6-(2-chloro-4-(6-methylpyridin-2-yl)phenyl)-2-(methylthio)-7-oxopyrido[2,3-d]pyrimidin-8(7H)-yl)propyl)carbamate (**51**)

Compound **51** was prepared according to general procedure I, using **41** (150 mg, 0.38 mmol) and tert-butyl (3-bromopropyl)carbamate (136 mg, 0.57 mmol) as starting materials. The crude product was purified by flash chromatography on silica gel. **51** was obtained as a white solid in a yield of 80% (167 mg). ^1^H NMR (500 MHz, DMSO-d6, 300 K): *δ* = 8.94 (s, 1H), 8.25 (d, *J* = 1.7 Hz, 1H), 8.12 (dd, *J* = 8.0, 1.7 Hz, 1H), 8.06 (s, 1H), 7.89 (d, *J* = 7.8 Hz, 1H), 7.82 (t, *J* = 7.7 Hz, 1H), 7.55 (d, *J* = 8.0 Hz, 1H), 7.29 (d, *J* = 7.6 Hz, 1H), 6.83 (t, *J* = 5.7 Hz, 1H), 4.41 - 4.35 (m, 2H), 3.05 - 2.98 (m, 2H), 2.64 (s, 3H), 2.57 (s, 3H), 1.85 - 1.79 (m, 2H), 1.37 (s, 9H) ppm. ^13^CNMR (126 MHz, DMSO-d6, 300 K): *δ* = 172.2, 160.4, 158.0, 157.5, 155.5, 153.4, 153.2, 140.4, 137.7, 135.7, 135.3, 133.4, 132.1, 130.3, 126.9, 124.9, 122.7, 117.7, 109.2, 77.5, 39.0, 37.7, 28.2 (3C), 28.0, 24.3, 14.0 ppm. MS (ESI+): m/z = 452.2 [M-BOC + H]^+^; calculated: 452.1.

#### tert-Butyl (4-(6-(2-chloro-4-(6-methylpyridin-2-yl)phenyl)-2-(methylthio)-7-oxopyrido[2,3-d]pyrimidin-8(7H)-yl)butyl)carbamate (**52**)

Compound **52** was prepared according to general procedure I, using **41** (110 mg, 0.28 mmol) and tert-butyl (4-bromobutyl)carbamate (105 mg, 0.42 mmol) as starting materials. The crude product was purified by flash chromatography on silica gel. **52** was obtained as a white solid in a yield of 91% (144 mg). ^1^H NMR (500 MHz, DMSO-d6, 300 K): *δ* = 8.95 (s, 1H), 8.25 (d, *J* = 1.7 Hz, 1H), 8.12 (dd, *J* = 8.0, 1.8 Hz, 1H), 8.06 (s, 1H), 7.89 (d, *J* = 7.8 Hz, 1H), 7.82 (t, *J* = 7.7 Hz, 1H), 7.54 (d, *J* = 8.0 Hz, 1H), 7.29 (d, *J* = 7.6 Hz, 1H), 6.79 (t, *J* = 5.6 Hz, 1H), 4.37 (t, *J* = 7.1 Hz, 2H), 2.97 - 2.92 (m, 2H), 2.63 (s, 3H), 2.57 (s, 3H), 1.72 - 1.66 (m, 2H), 1.48 - 1.42 (m, 2H), 1.35 (s, 9H) ppm. ^13^CNMR (126 MHz, DMSO-d6, 300 K): *δ* = 172.2, 160.4, 158.0, 157.5, 155.6, 153.4, 153.3, 140.4, 137.7, 135.6, 135.3, 133.5, 132.1, 130.3, 126.9, 124.9, 122.7, 117.7, 109.1, 77.3, 40.7, 39.0, 28.2 (3C), 27.1, 24.9, 24.3, 13.9 ppm. MS (ESI+): m/z = 466.2 [M-BOC + H]^+^; calculated: 466.2.

#### 2-(4-(6-(2-Chloro-4-(6-methylpyridin-2-yl)phenyl)-2-(methylthio)-7-oxopyrido[2,3-d]pyrimidin-8(7H)-yl)butyl)isoindoline-1,3-dione (**53**)

Compound **53** was prepared in a two-step synthesis. **41** (450 mg, 1.14 mmol), 2-(4-bromobutyl)isoindoline-1,3-dione (**20**; 386 mg, 1.37 mmol) and potassium carbonate (472 mg, 3.42 mmol) were dissolved in anhydrous DMSO (25 mL) and stirred at 110 °C for 18 h. The reaction solution was cooled to room temperature and diluted with ethyl acetate (250 mL). The organic layer was washed three times with saturated aqueous ammonium chloride solution (100 mL) and brine (100 mL). It was dried over MgSO4, and the solvent was evaporated under reduced pressure. The remaining residue, containing a mixture of the corresponding isoindoline-1,3-dione and 2-carbamoylbenzoic acid derivative, was then dissolved in acetic anhydride (25 mL) and heated to 100 °C for 2 h. The solvent was evaporated under reduced pressure, and the crude product was purified by flash chromatography on silica gel using DCM/MeOH as eluent. **53** was obtained as a white solid in a yield of 49% (330 mg) over two steps. ^1^H NMR (500 MHz, DMSO-d6, 300 K): *δ* = 8.91 (s, 1H), 8.22 (d, *J* = 1.7 Hz, 1H), 8.11 (dd, *J* = 8.0, 1.8 Hz, 1H), 8.04 (s, 1H), 7.88 (d, *J* = 7.7 Hz, 1H), 7.86 - 7.81 (m, 5H), 7.53 (d, *J* = 8.0 Hz, 1H), 7.29 (d, *J* = 7.6 Hz, 1H), 4.40 (t, *J* = 6.8 Hz, 2H), 3.62 (t, *J* = 6.7 Hz, 2H), 2.57 (s, 3H), 2.57 (s, 3H), 1.78 - 1.72 (m, 2H), 1.70 - 1.64 (m, 2H) ppm. ^13^CNMR (126 MHz, DMSO-d6, 300 K): *δ* = 172.1, 167.9 (2C), 160.4, 158.0, 157.5, 153.4, 153.3, 140.4, 137.7, 135.6, 135.3, 134.4 (2C), 133.4, 132.0, 131.5 (2C), 130.3, 126.8, 124.9, 123.0 (2C), 122.7, 117.7, 109.1, 40.4, 37.2, 25.3, 24.6, 24.3, 13.8 ppm. MS (ESI+): m/z = 596.2 [M + H]^+^; calculated: 596.2.

#### tert-Butyl (5-(6-(2-chloro-4-(6-methylpyridin-2-yl)phenyl)-2-(methylthio)-7-oxopyrido[2,3-d]pyrimidin-8(7H)-yl)pentyl)carbamate (**54**)

Compound **54** was prepared according to general procedure I, using **41** (200 mg, 0.51 mmol) and tert-butyl (5-bromopentyl)carbamate (135 mg, 0.51 mmol) as starting materials. The crude product was purified by flash chromatography on silica gel. **54** was obtained as a white solid in a yield of 97% (285 mg). ^1^H NMR (500 MHz, DMSO-d6, 300 K): *δ* = 8.94 (s, 1H), 8.24 (d, *J* = 1.7 Hz, 1H), 8.12 (dd, *J* = 8.0, 1.8 Hz, 1H), 8.05 (s, 1H), 7.89 (d, *J* = 7.8 Hz, 1H), 7.82 (t, *J* = 7.7 Hz, 1H), 7.54 (d, *J* = 8.0 Hz, 1H), 7.29 (d, *J* = 7.6 Hz, 1H), 6.75 (t, *J* = 5.3 Hz, 1H), 4.39 - 4.32 (m, 2H), 2.90 (q, *J* = 6.7 Hz, 2H), 2.63 (s, 3H), 2.57 (s, 3H), 1.72 - 1.66 (m, 2H), 1.46 - 1.40 (m, 2H), 1.36 - 1.33 (m, 11H) ppm. ^13^CNMR (126 MHz, DMSO-d6, 300 K): *δ* = 172.2, 160.4, 158.1, 157.5, 155.6, 153.4, 153.3, 140.5, 137.7, 135.6, 135.4, 133.5, 132.1, 130.3, 126.9, 124.9, 122.8, 117.7, 109.1, 77.3, 40.8, 40.1, 29.1, 28.3 (3C), 27.0, 24.3, 23.8, 13.9 ppm. MS (ESI+): m/z = 580.2 [M + H]^+^; calculated: 580.2.

#### tert-Butyl 4-(2-(6-(2-chloro-4-(6-methylpyridin-2-yl)phenyl)-2-(methylthio)-7-oxopyrido[2,3-d]pyrimidin-8(7H)-yl)ethyl)piperidine-1-carboxylate (**55**)

Compound **55** was prepared according to general procedure I, using **41** (200 mg, 0.51 mmol) and tert-butyl 4-(2-bromoethyl)piperidine-1-carboxylate (148 mg, 0.51 mmol) as starting materials. The crude product was purified by flash chromatography on silica gel. **55** was obtained as a white solid in a yield of 61% (188 mg). ^1^H NMR (500 MHz, DMSO-d6, 300 K): *δ* = 8.94 (s, 1H), 8.25 (d, *J* = 1.7 Hz, 1H), 8.12 (dd, *J* = 8.0, 1.7 Hz, 1H), 8.06 (s, 1H), 7.88 (d, *J* = 7.8 Hz, 1H), 7.82 (t, *J* = 7.7 Hz, 1H), 7.54 (d, *J* = 8.0 Hz, 1H), 7.29 (d, *J* = 7.6 Hz, 1H), 4.45 - 4.38 (m, 2H), 3.92 (d, *J* = 10.3 Hz, 2H), 2.75 - 2.65 (m, 2H), 2.63 (s, 3H), 2.57 (s, 3H), 1.75 (d, *J* = 11.8 Hz, 2H), 1.65 - 1.59 (m, 2H), 1.54 - 1.45 (m, 1H), 1.39 (s, 9H), 1.09 - 1.04 (m, 2H) ppm. ^13^CNMR (126 MHz, DMSO-d6, 300 K): *δ* = 172.2, 160.3, 158.0, 157.5, 153.9, 153.3, 153.2, 140.4, 137.7, 135.6, 135.3, 133.4, 132.1, 130.3, 126.9, 124.9, 122.7, 117.7, 109.2, 78.4, 43.6, 40.1, 38.8, 33.8, 33.4, 31.6 (2C), 28.1 (3C), 24.3, 13.9 ppm. MS (ESI+): m/z = 506.1 [M-BOC + H]^+^; calculated: 506.2.

#### tert-Butyl ((1r,4r)-4-((6-(2-chloro-4-(6-methylpyridin-2-yl)phenyl)-2-(methylthio)-7-oxopyrido[2,3-d]pyrimidin-8(7H)-yl)methyl)cyclohexyl)carbamate (**56**)

Compound **56** was prepared according to general procedure I, using **41** (150 mg, 0.38 mmol) and tert-butyl ((1*r*,4*r*)-4-(bromomethyl)cyclohexyl)carbamate (167 mg, 0.57 mmol) as starting materials. After purification by column chromatography on silica gel, an additional purification by HPLC chromatography on silica gel (H2O/ACN; 0.1% TFA) was performed for a small fraction of the crude product. The solvent was evaporated under reduced pressure, leading to the deprotection of the BOC-protected amine. The larger fraction was used in the synthesis of **62** without further improvement of the purity. **56** was obtained as a yellow solid in a yield of 43% (100 mg). ^1^H NMR (500 MHz, DMSO-d6, 300 K): *δ* = 8.95 (s, 1H), 8.24 (d, *J* = 1.7 Hz, 1H), 8.11 (dd, *J* = 8.0, 1.7 Hz, 1H), 8.08 (s, 1H), 7.92 - 7.87 (m, 2H), 7.76 (d, *J* = 2.8 Hz, 3H), 7.56 (d, *J* = 8.0 Hz, 1H), 7.35 (d, *J* = 7.3 Hz, 1H), 4.28 (d, *J* = 7.0 Hz, 2H), 2.99 (s, 1H), 2.63 (s, 3H), 2.59 (s, 3H), 1.94 - 1.86 (m, 3H), 1.73 - 1.67 (m, 2H), 1.26 - 1.19 (m, 4H) ppm. ^13^CNMR (126 MHz, DMSO-d6, 300 K): *δ* = 172.5, 161.1, 158.3, 158.0, 154.3, 153.4, 140.4, 138.8, 136.2, 136.0, 133.9, 132.6, 130.8, 127.5, 125.6, 123.5, 118.6, 109.6, 49.6, 46.3, 35.7, 30.3 (2C), 28.6 (2C), 24.4, 14.4 ppm. MS (ESI+): m/z = 506. [M + H]^+^; calculated: 506.2.

#### tert-Butyl (3-(6-(2-chloro-4-(6-methylpyridin-2-yl)phenyl)-2-(methylamino)-7-oxopyrido[2,3-d]pyrimidin-8(7H)-yl)ethyl)carbamate (**57**)

Compound **57** was prepared according to general procedures II and III. **50** (100 mg, 0.19 mmol) was used as starting material. The crude product was purified by flash chromatography on silica gel using DCM/MeOH as eluent to obtain **57** as a white solid in a yield of 22% (21 mg) over two steps. ^1^H NMR (500 MHz, DMSO-d6, 300 K): *δ* = 8.74 - 8.60 (m, 1H), 8.22 (d, *J* = 1.7 Hz, 1H), 8.08 (dd, *J* = 8.0, 1.7 Hz, 1H), 7.90 - 7.80 (m, 4H), 7.50 (d, *J* = 8.0 Hz, 1H), 7.27 (d, *J* = 7.5 Hz, 1H), 6.84 (t, *J* = 5.6 Hz, 1H), 4.47 - 4.32 (m, 2H), 3.34 - 3.35 (m, 2H), 2.94 (d, *J* = 4.4 Hz, 3H), 2.57 (s, 3H), 1.31 (s, 9H) ppm. ^13^CNMR (126 MHz, DMSO-d6, 300 K): *δ* = 162.0, 161.1, 159.3, 158.0, 155.5, 155.3, 153.4, 139.8, 137.7, 136.5, 136.2, 133.7, 132.3, 130.6, 126.9, 124.7, 122.6, 117.6, 104.0, 77.5, 40.3, 37.7, 28.1 (3C), 27.9, 24.3 ppm. MS (ESI+): m/z = 404.1 [Mfr.]^+^; calculated: 404.1.

#### tert-Butyl (3-(6-(2-chloro-4-(6-methylpyridin-2-yl)phenyl)-2-(methylamino)-7-oxopyrido[2,3-d]pyrimidin-8(7H)-yl)propyl)carbamate (**58**)

Compound **58** was prepared according to general procedures II and III, using **51** (100 mg, 0.18 mmol) as starting material. The crude product was purified by flash chromatography on silica gel using DCM/MeOH as eluent. **58** was obtained as a white solid in a yield of 59% (57 mg) over two steps. ^1^H NMR (500 MHz, DMSO-d6, 300 K): *δ* = 8.76 - 8.62 (m, 1H), 8.22 (d, *J* = 1.7 Hz, 1H), 8.08 (dd, *J* = 8.0, 1.8 Hz, 1H), 7.92 - 7.84 (m, 2H), 7.83 (s, 1H), 7.80 (t, *J* = 7.7 Hz, 1H), 7.51 (d, *J* = 8.0 Hz, 1H), 7.27 (d, *J* = 7.5 Hz, 1H), 6.78 (t, *J* = 5.3 Hz, 1H), 4.40 - 4.23 (m, 2H), 3.06 - 2.97 (m, 2H), 2.93 (d, *J* = 4.5 Hz, 3H), 2.57 (s, 3H), 1.85 - 1.75 (m, 2H), 1.36 (s, 9H) ppm. ^13^CNMR (126 MHz, DMSO-d6, 300 K): *δ* = 162.0, 160.9, 159.4, 158.0, 155.5, 155.0, 153.4, 139.9, 137.7, 136.4, 136.2, 133.7, 132.3, 126.8, 124.8, 124.3, 122.6, 117.6, 104.0, 77.5, 38.4, 37.8, 28.2 (3C), 28.0, 27.9, 24.3 ppm. MS (ESI+): m/z = 435.1 [M-BOC + H]^+^; calculated: 435.2.

#### tert-Butyl (3-(6-(2-chloro-4-(6-methylpyridin-2-yl)phenyl)-2-(methylamino)-7-oxopyrido[2,3-d]pyrimidin-8(7H)-yl)butyl)carbamate (**59**)

Compound **59** was prepared according to general procedures II and III, using **52** (98 mg, 0.17 mmol) as starting material. The crude product was purified by flash chromatography on silica gel using DCM/MeOH as eluent. **59** was obtained as a white solid in a yield of 54% (51 mg) over two steps. ^1^H NMR (500 MHz, DMSO-d6, 300 K): *δ* = 8.75 - 8.61 (m, 1H), 8.21 (d, *J* = 1.7 Hz, 1H), 8.08 (dd, *J* = 8.0, 1.8 Hz, 1H), 7.90 - 7.83 (m, 2H), 7.83 - 7.78 (m, 2H), 7.50 (d, *J* = 8.0 Hz, 1H), 7.28 (d, *J* = 7.6 Hz, 1H), 6.79 (s, 1H), 4.36 - 4.21 (m, 2H), 2.97 - 2.90 (m, 5H), 2.57 (s, 3H), 1.71 - 1.60 (m, 2H), 1.49 - 1.41 (m, 2H), 1.35 (s, 9H) ppm. ^13^CNMR (126 MHz, DMSO-d6, 300 K): *δ* = 162.0, 160.8, 159.4, 158.0, 155.6, 155.0, 153.4, 139.9, 137.6, 136.3, 136.2, 133.7, 132.3, 126.8, 124.8, 124.4, 122.6, 117.6, 103.9, 77.3, 40.1, 39.8, 28.2 (3C), 27.8, 27.2, 24.8, 24.3 ppm. MS (ESI+): m/z = 449.1 [M-BOC + H]^+^; calculated: 449.2.

#### tert-Butyl (5-(6-(2-chloro-4-(6-methylpyridin-2-yl)phenyl)-2-(methylamino)-7-oxopyrido[2,3-d]pyrimidin-8(7H)-yl)pentyl)carbamate (**60**)

Compound **60** was prepared according to general procedures II and III, using **54** (306 mg, 0.53 mmol) as starting material. The crude product was purified by flash chromatography on silica gel using DCM/MeOH as eluent. **60** was obtained as a white solid in a yield of 21% (63 mg) over two steps. ^1^H NMR (500 MHz, DMSO-d6, 300 K): *δ* = 8.71 - 8.62 (m, 1H), 8.21 (d, *J* = 1.7 Hz, 1H), 8.07 (dd, *J* = 8.0, 1.7 Hz, 1H), 7.88 - 7.83 (m, 2H), 7.82 - 7.79 (m, 2H), 7.50 (d, *J* = 8.0 Hz, 1H), 7.27 (d, *J* = 7.6 Hz, 1H), 6.76 (s, 1H), 4.32 - 4.22 (m, 2H), 2.92 - 2.90 (m, 5H), 2.56 (s, 3H), 1.70 - 1.62 (m, 2H), 1.47 - 1.42 (m, 2H), 1.39 - 1.36 (m, 2H), 1.35 (s, 9H) ppm. ^13^CNMR (126 MHz, DMSO-d6, 300 K): *δ* = 162.0, 161.9, 160.9, 159.4, 158.0, 155.6, 155.0, 153.4, 139.9, 137.7, 136.3, 133.7, 132.3, 126.8, 124.8, 124.4, 122.6, 117.6, 104.0, 77.3, 40.2, 40.1, 29.1, 28.3 (3C), 27.9, 26.9, 24.3, 23.9 ppm. MS (ESI+): m/z = 563.3 [M + H]^+^; calculated: 563.3.

#### tert-Butyl 4-(2-(6-(2-chloro-4-(6-methylpyridin-2-yl)phenyl)-2-(methylamino)-7-oxopyrido[2,3-d]pyrimidin-8(7H)-yl)ethyl)piperidine-1-carboxylate (**61**)

Compound **61** was prepared according to general procedures II and III, using **55** (178 mg, 0.29 mmol) as starting material. The crude product was purified by flash chromatography on silica gel using DCM/MeOH as eluent. **61** was obtained as a white solid in a yield of 49% (84 mg) over two steps. ^1^H NMR (500 MHz, DMSO-d6, 300 K): *δ* = 8.72 - 8.64 (m, 1H), 8.22 (d, *J* = 1.7 Hz, 1H), 8.08 (dd, *J* = 8.0, 1.5 Hz, 1H), 7.90 - 7.85 (m, 2H), 7.83 - 7.81 (m, 2H), 7.50 (d, *J* = 8.0 Hz, 1H), 7.28 (d, *J* = 7.6 Hz, 1H), 4.39 - 4.28 (m, 2H), 3.94 - 3.88 (m, 2H), 2.92 (d, *J* = 4.8 Hz, 3H), 2.75 - 2.62 (m, 2H), 2.57 (s, 3H), 1.76 (d, *J* = 11.8 Hz, 2H), 1.64 - 1.58 (m, 2H), 1.52 - 1.47 (m, 1H), 1.39 (s, 9H), 1.09 - 1.04 (m, 2H) ppm. ^13^CNMR (126 MHz, DMSO-d6, 300 K): *δ* = 162.0, 160.8, 159.4, 158.0, 154.9, 153.9, 153.4, 139.9, 137.7, 136.3, 136.2, 133.7, 132.3, 126.8, 124.8, 124.4, 122.6, 117.6, 104.0, 78.4, 43.1, 40.0, 38.2, 33.7, 33.4, 31.6 (2C), 28.1 (3C), 27.9, 24.3 ppm. MS (ESI+): m/z = 489.2 [M-BOC + H]^+^; calculated: 489.2.

#### tert-Butyl ((1r,4r)-4-((6-(2-chloro-4-(6-methylpyridin-2-yl)phenyl)-2-(methylamino)-7-oxopyrido[2,3-d]pyrimidin-8(7H)-yl)methyl)cyclohexyl)carbamate (**62**)

Compound **62** was prepared according to general procedures II and III, using **56** (61 mg, 0.10 mmol) as starting material. The crude product was purified by flash chromatography on silica gel using DCM/MeOH as eluent. **62** was obtained as a yellow solid in a yield of 75% (45 mg) over two steps. ^1^H NMR (500 MHz, DMSO-d6, 300 K): *δ* = 8.72 - 8.62 (m, 1H), 8.21 (d, *J* = 1.7 Hz, 1H), 8.07 (dd, *J* = 8.0, 1.7 Hz, 1H), 7.88 - 7.78 (m, 4H), 7.50 (d, *J* = 8.0 Hz, 1H), 7.27 (d, *J* = 7.5 Hz, 1H), 6.64 (d, *J* = 8.0 Hz, 1H), 4.23 - 4.12 (m, 2H), 3.16 (s, 1H), 2.91 (d, *J* = 4.8 Hz, 3H), 2.57 (s, 3H), 1.87 - 1.80 (m, 1H), 1.78 - 1.72 (m, 2H), 1.66 - 1.59 (m, 2H), 1.36 (s, 9H), 1.16 - 1.03 (m, 4H) ppm. ^13^CNMR (126 MHz, DMSO-d6, 300 K): *δ* = 161.8, 161.1, 159.4, 158.0, 155.3, 154.8, 153.4, 139.9, 137.7, 136.3, 136.2, 133.7, 132.3, 126.8, 124.8, 124.5, 122.6, 117.6, 103.9, 77.3, 49.2, 45.6, 35.8, 32.2 (2C), 29.5 (2C), 28.3 (3C), 27.9, 24.3 ppm. MS (ESI+): m/z = 589.3 [M + H]^+^; calculated: 589.3.

#### 8-(2-Aminoethyl)-6-(2-chloro-4-(6-methylpyridin-2-yl)phenyl)-2-(methylamino)pyrido[2,3-d]pyrimidin-7(8H)-one (**18**)

Compound **18** was prepared according to general procedure IV, using **57** (12 mg, 0.02 mmol) as starting material. **18** was obtained as a pale yellow solid in a yield of 36% (5 mg). ^1^H NMR (500 MHz, DMSO-d6, 300 K): *δ* = 8.79 - 8.67 (m, 1H), 8.23 (d, *J* = 1.7 Hz, 1H), 8.09 (dd, *J* = 8.1, 1.8 Hz, 1H), 7.96 (d, *J* = 4.7 Hz, 1H), 7.91 - 7.80 (m, 6H), 7.54 (d, *J* = 8.1 Hz, 1H), 7.29 (d, *J* = 7.5 Hz, 1H), 4.64 - 4.43 (m, 2H), 3.24 - 3.19 (m, 2H), 2.95 (d, *J* = 4.8 Hz, 3H), 2.57 (s, 3H) ppm. ^13^CNMR (126 MHz, DMSO-d6, 300 K): *δ* = 166.1, 159.5, 158.0, 155.3, 153.3, 139.9, 137.7, 137.1, 133.6, 132.7, 132.5, 130.7, 128.8, 127.9, 126.9, 124.7, 122.7, 117.6, 38.2, 37.5, 29.0, 24.3 ppm. HRMS (FTMS +p MALDI): m/z = 421.1543 [M + H]^+^; calculated for C22H22ClN6O: 421.1538. HPLC: tR = 6.61 min; purity ≥ 95% (UV: 280/310 nm). LCMS (ESI+): m/z = 421.1 [M + H]^+^; calculated: 421.2.

#### 8-(3-Aminopropyl)-6-(2-chloro-4-(6-methylpyridin-2-yl)phenyl)-2-(methylamino)-pyrido[2,3-d]pyrimidin-7(8H)-one (**19**)

Compound **19** was prepared according to general procedure IV, using **58** (50 mg, 0.09 mmol) as starting material. **19** was obtained as a pale yellow solid in a yield of 13% (8 mg). ^1^H NMR (500 MHz, DMSO-d6, 300 K): *δ* = 8.78 - 8.67 (m, 1H), 8.22 (d, *J* = 1.8 Hz, 1H), 8.08 (dd, *J* = 8.0, 1.8 Hz, 1H), 7.97 (d, *J* = 4.1 Hz, 1H), 7.91 - 7.87 (m, 3H), 7.69 (s, 3H), 7.52 (d, *J* = 8.0 Hz, 1H), 7.34 (d, *J* = 7.1 Hz, 1H), 4.43 - 4.34 (m, 2H), 2.94 (d, *J* = 4.2 Hz, 3H), 2.90 - 2.79 (m, 2H), 2.59 (s, 3H), 2.07 - 1.98 (m, 2H) ppm. ^13^CNMR (126 MHz, DMSO-d6, 300 K): *δ* = 161.9, 161.3, 159.5, 157.8, 155.0, 153.1, 139.4, 138.3, 136.8, 136.3, 133.8, 132.3, 127.0, 125.0, 124.2, 123.0, 118.1, 104.1, 37.6, 37.2, 27.9, 25.8, 24.0 ppm. HRMS (FTMS +p MALDI): m/z = 435.1699 [M + H]^+^; calculated for C23H24ClN6O: 435.1695. HPLC: tR = 6.68 min; purity ≥ 95% (UV: 280/310 nm). LCMS (ESI+): m/z = 435.2 [M + H]^+^; calculated: 435.2.

#### 8-(4-Aminobutyl)-6-(2-chloro-4-(6-methylpyridin-2-yl)phenyl)-2-(methylamino)pyrido[2,3-d]pyrimidin-7(8H)-one (**20**)

Compound **20** was prepared according to general procedure IV, using **59** (46 mg, 0.08 mmol) as starting material. **20** was obtained as a pale yellow solid in a yield of 29% (16 mg). ^1^H NMR (500 MHz, DMSO-d6, 300 K): *δ* = 8.80 - 8.63 (m, 1H), 8.22 (d, *J* = 1.8 Hz, 1H), 8.07 (dd, *J* = 8.0, 1.8 Hz, 1H), 7.94 (s, 1H), 7.91 - 7.84 (m, 3H), 7.70 (s, 3H), 7.51 (d, *J* = 8.0 Hz, 1H), 7.33 (d, *J* = 7.7 Hz, 1H), 4.39 - 4.29 (m, 2H), 2.93 (d, *J* = 3.7 Hz, 3H), 2.89 - 2.81 (m, 2H), 2.58 (s, 3H), 1.83 - 1.70 (m, 2H), 1.67 - 1.56 (m, 2H) ppm. ^13^CNMR (126 MHz, DMSO-d6, 300 K): *δ* = 161.9, 161.0, 159.4, 157.8, 155.1, 153.1, 139.4, 138.4, 136.5, 136.3, 133.8, 132.3,127.1, 125.0, 124.4, 123.0, 118.1, 104.0, 40.1, 38.8, 27.9, 24.7, 24.5, 24.0 ppm. HRMS (FTMS +p MALDI): m/z = 449.1856 [M + H]^+^; calculated for C24H26ClN6O: 449.1851. HPLC: tR = 6.74 min; purity ≥ 95% (UV: 280/310 nm). LCMS (ESI+): m/z = 449.2 [M + H]^+^; calculated: 449.2.

#### 8-(5-Aminopentyl)-6-(2-chloro-4-(6-methylpyridin-2-yl)phenyl)-2-(methylamino)-pyrido[2,3-d]pyrimidin-7(8H)-one (**21**)

Compound **21** was prepared according to general procedure IV, using **60** (42 mg, 0.08 mmol) as starting material. The crude product was purified by flash chromatography on silica gel using DCM/MeOH as eluent to obtain **21** as a pale yellow solid in a yield of 95% (49 mg). ^1^H NMR (500 MHz, DMSO-d6, 300 K): *δ* = 8.76 - 8.62 (m, 1H), 8.22 (d, *J* = 1.7 Hz, 1H), 8.08 (dd, *J* = 8.0, 1.7 Hz, 1H), 7.92 - 7.79 (m, 4H), 7.68 (s, 3H), 7.50 (d, *J* = 8.0 Hz, 1H), 7.29 (d, *J* = 7.5 Hz, 1H), 4.40 - 4.20 (m, 2H), 2.92 (d, *J* = 4.8 Hz, 3H), 2.84 - 2.74 (m, 2H), 2.57 (s, 3H), 1.77 - 1.67 (m, 2H), 1.67 - 1.57 (m, 2H), 1.48 - 1.32 (m, 2H) ppm. ^13^CNMR (126 MHz, DMSO-d6, 300 K): *δ* = 162.0, 160.9, 159.4, 158.0, 155.0, 153.3, 139.8, 137.7, 136.4, 136.2, 133.7, 132.3, 126.8, 124.8, 124.4, 122.7, 117.6, 104.0, 40.1, 38.8, 27.9, 26.7, 26.6, 24.3, 23.4 ppm. HRMS (FTMS +p MALDI): m/z = 463.2001 [M + H]^+^; calculated for C25H28ClN6O: 463.2008. HPLC: tR = 7.07 min; purity ≥ 95% (UV: 280/310 nm). LCMS (ESI+): m/z = 463.1 [M + H]^+^; calculated: 463.2.

#### 6-(2-Chloro-4-(6-methylpyridin-2-yl)phenyl)-2-(methylamino)-8-(2-(piperidin-4-yl)ethyl)pyrido[2,3-d]pyrimidin-7(8H)-one (**22**)

Compound **22** was prepared according to general procedure IV, using **61** (50 mg, 0.09 mmol) as starting material. The crude product was purified by flash chromatography on silica gel using DCM/MeOH as eluent to obtain **22** as a pale yellow solid in a yield of 66% (40 mg). ^1^H NMR (500 MHz, DMSO-d6, 300 K): *δ* = 8.74 - 8.62 (m, 1H), 8.51 (s, 2H), 8.21 (d, *J* = 1.7 Hz, 1H), 8.07 (dd, *J* = 8.0, 1.7 Hz, 1H), 7.92 (d, *J* = 4.7 Hz, 1H), 7.87 - 7.78 (m, 3H), 7.50 (d, *J* = 8.0 Hz, 1H), 7.27 (d, *J* = 7.5 Hz, 1H), 4.39 - 4.25 (m, 2H), 3.27 (d, *J* = 12.3 Hz, 2H), 2.92 (d, *J* = 4.8 Hz, 3H), 2.85 (t, *J* = 11.7 Hz, 2H), 2.56 (s, 3H), 1.95 (d, *J* = 13.3 Hz, 2H), 1.63 (d, *J* = 15.6 Hz, 3H), 1.43 - 1.30 (m, 2H) ppm. ^13^CNMR (126 MHz, DMSO-d6, 300 K): *δ* = 162.0, 160.9, 159.5, 158.0, 154.9, 153.4, 139.9, 137.7, 136.4, 136.2, 133.8, 132.3, 126.8, 124.8, 124.4, 122.7, 117.6, 104.0, 43.2 (2C), 38.0, 33.3, 31.2, 28.4 (2C), 27.9, 24.3 ppm. HRMS (FTMS +p MALDI): m/z = 489.2161 [M + H]^+^; calculated for C27H30ClN6O: 489.2164. HPLC: tR = 7.06 min; purity ≥ 95% (UV: 280/310 nm). LCMS (ESI+): m/z = 489.3 [M + H]^+^; calculated: 489.2.

#### 8-(((1r,4r)-4-Aminocyclohexyl)methyl)-6-(2-chloro-4-(6-methylpyridin-2-yl)phenyl)-2-(methylamino)pyrido[2,3-d]pyrimidin-7(8H)-one (**23**)

Compound **23** was prepared according to general procedure IV, using **62** (19 mg, 0.03 mmol) as starting material. The crude product was purified by HPLC chromatography on silica gel (H2O/ACN; 0.1% TFA) to obtain **23** as a white solid in a yield of 87% (20 mg). ^1^H NMR (500 MHz, DMSO-d6, 300 K): *δ* = 8.74 - 8.64 (m, 1H), (d, *J* = 1.7 Hz, 1H), 8.07 (dd, *J* = 8.0, 1.7 Hz, 1H), 7.98 - 7.89 (m, 3H), 7.85 (s, 1H), 7.76 (s, 3H), 7.53 (d, *J* = 8.0 Hz, 1H), 7.41 - 7.34 (m, 1H), 4.29 - 4.11 (m, 2H), 2.98 (s, 1H), 2.92 (d, *J* = 2.3 Hz, 3H), 2.60 (s, 3H), 1.97 - 1.86 (m, 3H), 1.75 - 1.67 (m, 2H), 1.24 - 1.16 (m, 4H) ppm. ^13^CNMR (126 MHz, DMSO-d6, 300 K): *δ* = 161.6, 161.1, 159.2, 157.6, 155.4, 152.9, 139.0, 138.8, 136.6, 136.4, 133.8, 132.3, 127.1, 125.2, 124.5, 123.2, 118.4, 104.0, 49.3, 45.4, 35.3, 29.9 (2C), 28.4 (2C), 27.9, 23.7 ppm. HRMS (FTMS +p MALDI): m/z = 489.2160 [M + H]^+^; calculated for C27H30ClN6O: 489.2164. HPLC: tR = 5.49 min; purity ≥ 95% (UV: 280/320 nm). LCMS (ESI+): m/z = 489.3 [M + H]^+^; calculated: 489.2.

#### 8-(4-Aminobutyl)-6-(2-chloro-4-(6-methylpyridin-2-yl)phenyl)-2-((1,3-dihydroxypropan-2-yl)amino)pyrido[2,3-d]pyrimidin-7(8H)-one (**24**)

Compound **24** was prepared according to general procedure II, using **53** (50 mg, 0.08 mmol) as starting material. The remaining residue was dissolved in a mixture of anhydrous ethanol/DMF (5:1; 6 mL) and 2-aminopropane-1,3-diol (31 mg, 0.34 mmol), and DIPEA (54 mg, 0.42 mmol) were added. The reaction solution was heated to 85 °C for 17 h. Afterwards, the solvent was evaporated under reduced pressure. HPLC analysis showed complete conversion of the starting material by formation of the corresponding isoindoline-1,3-dione and 2-carbamoylbenzoic acid derivative. To enable the complete deprotection of the phthalimide protecting group, the remaining residue was converted according to general procedure III. The crude product was purified by HPLC chromatography on silica gel (H2O/ACN; 0.1% TFA) to obtain **24** as a pale yellow solid in a yield of 16% (10 mg) over three steps. ^1^H NMR (500 MHz, DMSO-d6, 300 K): (mixture of rotamers) *δ* = 8.74 - 8.66 (m, 1H), 8.22 (d, *J* = 1.7 Hz, 1H), 8.08 (dd, *J* = 8.0, 1.8 Hz, 1H), 7.90 - 7.83 (m, 3H), 7.66 (s, 3H), 7.51 (d, *J* = 8.0 Hz, 2H), 7.32 (d, *J* = 7.4 Hz, 1H), 4.35 - 4.29 (m, 2H), 4.11 - 4.10 (m, 0.3H), 4.02 - 4.01 (m, 0.7H), 3.74 (s, 2H), 3.61 (d, *J* = 5.5 Hz, 2.7H), 3.56 (d, *J* = 5.1 Hz, 1.3H), 2.88 - 2.80 (m, 2H), 2.58 (s, 3H), 1.79 - 1.71 (m, 2H), 1.66 - 1.55 (m, 2H) ppm. ^13^CNMR (126 MHz, DMSO-d6, 300 K): (mixture of rotamers) *δ* = 161.4, 161.0, 159.4, 157.9, 154.9, 153.2, 139.6, 138.1, 136.4, 136.2, 133.8, 132.3, 127.0, 124.9, 124.5, 122.9, 117.9, 104.3, 60.3, 60.1 (2C), 55.5, 54.9, 40.1, 38.8, 24.7, 24.6, 24.1 ppm. HRMS (FTMS +p MALDI): m/z = 509.2061 [M + H]^+^; calculated for C26H30ClN6O3: 509.2062. HPLC: tR = 6.32 min; purity ≥ 95% (UV: 280/310 nm). LCMS (ESI+): m/z = 509.2 [M + H]^+^; calculated: 509.2.

#### 8-(4-Aminobutyl)-2-((3-(4-aminophenyl)propyl)amino)-6-(2-chloro-4-(6-methylpyridin-2-yl)phenyl)pyrido[2,3-d]pyrimidin-7(8H)-one (**25**)

Compound **25** was prepared according to general procedure II, using **53** (50 mg, 0.08 mmol) as starting material. The remaining residue was dissolved in anhydrous ethanol (5 mL) and 4-(3-aminopropyl)aniline (126 mg, 0.84 mmol), and DIPEA (130 mg, 1.01 mmol) were added. The reaction solution was heated to 85 °C for 17 h. Afterwards, the solvent was evaporated under reduced pressure. HPLC analysis showed complete conversion of the starting material by formation of the corresponding isoindoline-1,3-dione and 2-carbamoylbenzoic acid derivative. To enable the complete deprotection of the phthalimide protecting group, the remaining residue was converted according to general procedure III. The crude product was purified by HPLC chromatography on silica gel (H2O/ACN; 0.1% TFA) to obtain **25** as a yellow solid in a yield of 34% (23 mg) over three steps. ^1^H NMR (500 MHz, DMSO-d6, 300 K): *δ* = 8.77 - 8.65 (m, 3H), 8.22 (d, *J* = 1.7 Hz, 2H), 8.06 (dd, *J* = 8.0, 1.7 Hz, 1H), 8.01 - 7.95 (m, 2H), 7.88 - 7.77 (m, 4H), 7.54 (d, *J* = 8.0 Hz, 1H), 7.44 (d, *J* = 7.5 Hz, 1H), 7.38 (d, *J* = 8.4 Hz, 2H), 7.31 (d, *J* = 8.3 Hz, 2H), 4.30 (t, *J* = 6.6 Hz, 2H), 3.47 - 3.38 (m, 2H), 2.91 - 2.78 (m, 2H), 2.75 - 2.66 (m, 2H), 2.63 (s, 3H), 1.99 - 1.88 (m, 2H), 1.78 - 1.72 (m, 2H), 1.63 - 1.56 (m, 2H) ppm. ^13^CNMR (126 MHz, DMSO-d6, 300 K): *δ* = 161.1, 161.1, 159.0, 157.3, 155.2, 152.6, 142.0, 139.7, 138.2, 136.8, 136.6, 133.9, 132.4, 129.7 (2C), 129.6, 127.5, 125.5, 124.5, 123.7, 123.1 (2C), 119.2, 104.2, 40.6, 40.1, 38.8, 32.2, 30.4, 24.8, 24.6, 23.2 ppm. HRMS (FTMS +p MALDI): m/z = 568.2579 [M + H]^+^; calculated for C32H35ClN7O: 568.2586. HPLC: tR = 6.79 min; purity ≥ 95% (UV: 280/310 nm). LCMS (ESI+): m/z = 568.2 [M + H]^+^; calculated: 568.3.

#### 8-(4-Aminobutyl)-6-(2-chloro-4-(6-methylpyridin-2-yl)phenyl)-2-((3-morpholinopropyl) amino)pyrido[2,3-d]pyrimidin-7(8H)-one (**26**)

Compound **26** was prepared according to general procedure II, using **53** (50 mg, 0.08 mmol) as starting material. The remaining residue was dissolved in a mixture of anhydrous ethanol/DMF (5:1; 6 mL) and 3-morpholinopropan-1-amine (121 mg, 0.84 mmol), and DIPEA (130 mg, 1.01 mmol) were added. The reaction solution was heated to 85 °C for 17 h. Afterwards, the solvent was evaporated under reduced pressure, and the crude product was purified by HPLC chromatography on silica gel (H2O/ACN; 0.1% TFA). **26** was obtained as a yellow solid in a yield of 35% (27 mg) over two steps. ^1^H NMR (500 MHz, DMSO-d6, 300 K): *δ* = 10.40 - 10.07 (m, 1H), 8.75 - 8.69 (m, 1H), 8.22 (d, *J* = 1.8 Hz, 1H), 8.11 (s, 1H), 8.07 (dd, *J* = 8.0, 1.8 Hz, 1H), 7.95 - 7.91 (m, 2H), 7.88 (s, 1H), 7.80 (s, 3H), 7.52 (d, *J* = 8.0 Hz, 1H), 7.42 - 7.35 (m, 1H), 4.44 - 4.24 (m, 2H), 3.99 (d, *J* = 12.0 Hz, 2H), 3.67 (t, *J* = 12.1 Hz, 2H), 3.51 - 3.42 (m, 4H), 3.23 - 3.18 (m, 2H), 3.13 - 3.02 (m, 2H), 2.91 - 2.79 (m, 2H), 2.61 (s, 3H), 2.07 - 1.94 (m, 2H), 1.86 - 1.72 (m, 2H), 1.65 - 1.61 (m, 2H) ppm. ^13^CNMR (126 MHz, DMSO-d6, 300 K): *δ* = 161.3, 161.0, 159.4, 157.5, 155.0, 152.8, 139.1, 138.7, 136.5, 136.5, 133.8, 132.3, 127.3, 125.2, 124.7, 123.4, 118.6, 104.4, 63.4 (2C), 54.3, 51.2 (2C), 40.1, 38.8, 38.2, 24.8, 24.6, 23.5, 22.9 ppm. HRMS (FTMS +p MALDI): m/z = 562.2672 [M + H]^+^; calculated for C30H37ClN7O2: 562.2692. HPLC: tR = 6.27 min; purity ≥ 95% (UV: 280/310 nm). LCMS (ESI+): m/z = 562.3 [M + H]^+^; calculated: 562.3.

#### 8-(4-Aminobutyl)-6-(2-chloro-4-(6-methylpyridin-2-yl)phenyl)-2-((3-(2-oxopyrrolidin-1-yl)propyl)amino)pyrido[2,3-d]pyrimidin-7(8H)-one (**27**)

Compound **27** was prepared according to general procedure II, using **53** (50 mg, 0.08 mmol) as starting material. The remaining residue was dissolved in anhydrous ethanol (5 mL) and 1-(3-aminopropyl)pyrrolidin-2-one (119 mg, 0.84 mmol), and DIPEA (130 mg, 1.01 mmol) were added. The reaction solution was heated to 85 °C for 17 h. Afterwards, the solvent was evaporated under reduced pressure, and the crude product was purified by HPLC chromatography on silica gel (H2O/ACN; 0.1% TFA). **27** was obtained as a yellow solid in a yield of 42% (28 mg) over two steps. ^1^H NMR (500 MHz, DMSO-d6, 300 K): *δ* = 8.73 - 8.67 (m, 1H), 8.22 (d, *J* = 1.7 Hz, 1H), 8.08 (dd, *J* = 8.0, 1.7 Hz, 1H), 7.99 (t, *J* = 5.5 Hz, 1H), 7.89 - 7.84 (m, 3H), 7.67 (s, 3H), 7.50 (d, *J* = 8.0 Hz, 1H), 7.31 (d, *J* = 7.4 Hz, 1H), 4.37 - 4.30 (m, 2H), 3.39 - 3.35 (m, 4H), 3.30 - 3.25 (m, 2H), 2.89 - 2.80 (m, 2H), 2.58 (s, 3H), 2.25 - 2.21 (m, 2H), 1.96 - 1.91 (m, 2H), 1.81 - 1.72 (m, 4H), 1.65 - 1.58 (m, 2H) ppm. ^13^CNMR (126 MHz, DMSO-d6, 300 K): *δ* = 174.1, 161.4, 160.9, 159.5, 157.9, 155.0, 153.2, 139.6, 138.0, 136.5, 136.2, 133.7, 132.3, 126.9, 124.9, 124.4, 122.8, 117.8, 104.2, 46.5, 40.1 (2C), 38.8 (2C), 30.5, 26.5, 24.7, 24.6, 24.1, 17.5 ppm. HRMS (FTMS +p MALDI): m/z = 560.2526 [M + H]^+^; calculated for C30H35ClN7O2: 560.2535. HPLC: tR = 6.89 min; purity ≥ 95% (UV: 280/310 nm). LCMS (ESI+): m/z = 560.3 [M + H]^+^; calculated: 560.3.

#### 8-(4-Aminobutyl)-6-(2-chloro-4-(6-methylpyridin-2-yl)phenyl)-2-((2-methyl-5-(trifluoromethyl)phenyl)amino)pyrido[2,3-d]pyrimidin-7(8H)-one (**28**)

Compound **28** was prepared according to general procedure II, using **53** (25 mg, 0.04 mmol) as starting material. The remaining residue obtained was dissolved in anhydrous THF (1.5 mL) and cooled to −70 °C using an ethyl acetate/liquid-nitrogen cooling bath. 2-Methyl-5-(trifluoromethyl)aniline (29 mg, 0.17 mmol) was dissolved in anhydrous THF (1.5 mL), cooled to −70 °C, and a solution of LiHMDS in THF (1 M, 42 mg, 0.25 mmol) was added. After 15 min, the first solution, containing the sulfone/sulfoxide intermediate, was added to the second one using a syringe. The combined solutions were stirred at −70 °C for 1 h. HPLC analysis showed complete conversion of the starting material via formation of the corresponding isoindoline-1,3-dione and 2-carbamoylbenzoic acid derivative. The reaction was stopped, and unreacted LiHMDS was quenched by adding water (5 mL) dropwise. The aquatic layer (pH 7) was first acidified using HCl (1M) and extracted three times with ethyl acetate (5 mL). Second, it was basified using NaOH (1M) and extracted three times using DCM (5 mL). The combined organic layers were dried over MgSO4, and the solvent was evaporated under reduced pressure. To enable the complete deprotection of the phthalimide protecting group, the remaining residue was converted according to general procedure III. The crude product was purified by HPLC chromatography on silica gel (H2O/ACN; 0.1% TFA) to obtain **28** as a yellow solid in a yield of 29% (10 mg) over three steps. ^1^H NMR (500 MHz, DMSO-d6, 300 K): *δ* = 9.63 (s, 1H), 8.86 (s, 1H), 8.23 (d, *J* = 1.7 Hz, 1H), 8.10 (dd, *J* = 8.1, 1.6 Hz, 1H), 8.07 (s, 1H), 7.96 (s, 1H), 7.89 (d, *J* = 7.7 Hz, 1H), 7.87 - 7.83 (m, 1H), 7.63 (s, 3H), 7.52 (d, *J* = 7.9 Hz, 2H), 7.47 (d, *J* = 8.1 Hz, 1H), 7.32 (d, *J* = 7.2 Hz, 1H), 4.23 - 4.21 (m, 2H), 2.79 - 2.71 (m, 2H), 2.58 (s, 3H), 2.39 (s, 3H), 1.71 - 1.61 (m, 2H), 1.50 - 1.44 (m, 2H) ppm. ^13^CNMR (126 MHz, DMSO-d6, 300 K): *δ* = 160.9, 159.9, 159.7, 157.9, 154.8, 153.2, 139.9, 138.0, 137.9, 136.9, 136.2, 135.9, 133.7, 132.2, 131.4, 127.2 - 126.4 (m, 2C), 126.3, 124.9, 128.1 - 120.6 (m, 1C), 122.8, 121.5 - 121.2 (m), 121.2 - 121.0 (m), 117.8, 106.1, 40.1, 38.6, 24.4, 24.3, 24.2, 18.1 ppm. HRMS (FTMS +p MALDI): m/z = 593.2036 [M + H]^+^; calculated for C31H29ClF3N6O: 593.2038. HPLC: tR = 8.29 min; purity ≥ 95% (UV: 280/310 nm). LCMS (ESI+): m/z = 593.3 [M + H]^+^; calculated: 593.2.

#### tert-Butyl 4-(3-((8-(4-((tert-butoxycarbonyl)amino)butyl)-6-(2-chloro-4-(6-methylpyridin-2-yl)phenyl)-7-oxo-7,8-dihydropyrido[2,3-d]pyrimidin-2-yl)amino)propyl)piperazine-1-carboxylate (**63**)

Compound **63** was prepared according to general procedure II, using **52** (47 mg, 0.08 mmol) as starting material. The obtained remaining residue was dissolved in ethanol (5 mL) and tert-butyl 4-(3-aminopropyl)piperazine-1-carboxylate (82 mg, 0.33 mmol), and DIPEA (54 mg, 0.42 mmol) were added. The reaction solution was heated to 85 °C for 17 h. Afterwards the solvent was evaporated under reduced pressure and the crude product was purified by flash chromatography on silica gel using DCM/MeOH as eluent. **63** was obtained as a yellow solid in a yield of 92% (59 mg) over two steps. ^1^H NMR (500 MHz, DMSO-d6, 300 K): *δ* = 8.70 - 8.63 (m, 1H), 8.21 (d, *J* = 1.7 Hz, 1H), 8.07 (dd, *J* = 8.0, 1.7 Hz, 1H), 8.00 (t, *J* = 5.7 Hz, 1H), 7.86 (d, *J* = 7.8 Hz, 1H), 7.83 - 7.78 (m, 2H), 7.49 (d, *J* = 8.0 Hz, 1H), 7.27 (d, *J* = 7.6 Hz, 1H), 6.83 - 6.77 (m, 1H), 4.32 - 4.24 (m, 2H), 3.43 - 3.38 (m, 2H), 3.31 - 3.24 (m, 4H), 2.96 - 2.92 (m, 2H), 2.57 (s, 3H), 2.39 - 2.36 (m, 2H), 2.33 - 2.30 (m, 4H), 1.83 - 1.73 (m, 2H), 1.68 - 1.61 (m, 2H), 1.49 - 1.42 (m, 2H), 1.39 (s, 9H), 1.35 (s, 9H) ppm. ^13^CNMR (126 MHz, DMSO-d6, 300 K): *δ* = 161.5, 160.8, 159.5, 158.0, 155.5, 154.9, 153.8, 153.4, 139.9, 137.7, 136.3, 136.2, 133.7, 132.3, 126.8, 124.8, 124.4, 122.6, 117.6, 104.1, 78.7, 77.3, 55.6, 52.6 (2C), 43.8, 42.9, 40.1 (3C), 28.2 (3C), 28.1 (3C), 27.3, 25.9, 25.0, 24.3 ppm. MS (ESI+): m/z = 761.4 [M + H]^+^; calculated: 761.4.

#### 8-(4-Aminobutyl)-6-(2-chloro-4-(6-methylpyridin-2-yl)phenyl)-2-((3-(piperazin-1-yl)propyl) amino)pyrido[2,3-d]pyrimidin-7(8H)-one (**29**)

Compound **29** was prepared according to general procedure IV, using **63** (16 mg, 0.02 mmol) as starting material. The crude product was purified by HPLC chromatography on silica gel (H2O/ACN; 0.1% TFA) to obtain **29** as a yellow solid in a yield of 85% (14 mg). ^1^H NMR (500 MHz, DMSO-d6, 300 K): *δ* = 9.26 (s, 2H), 8.77 - 8.66 (m, 1H), (d, *J* = 1.8 Hz, 1H), 8.08 (dd, *J* = 8.0, 1.8 Hz, 2H), 7.90 - 7.83 (m, 3H), 7.74 (s, 3H), 7.50 (d, *J* = 8.0 Hz, 1H), 7.32 (d, *J* = 7.5 Hz, 1H), 4.35 - 4.31 (m, 2H), 4.14 - 4.13 (m, 2H), 3.47 - 3.36 (m, 8H), 3.20 - 3.16 (m, 2H), 2.86 - 2.82 (m, 2H), 2.58 (s, 3H), 2.02 - 1.95 (m, 2H), 1.79 - 1.73 (m, 2H), 1.62 - 1.57 (m, 2H) ppm. ^13^CNMR (126 MHz, DMSO-d6, 300 K): *δ* = 161.4, 161.0, 159.6, 157.9, 155.0, 153.2, 139.7, 138.1, 136.5, 136.1, 133.7, 132.3, 127.0, 124.9, 124.7, 122.9, 117.9, 104.4, 67.5, 54.1, 40.6 (2C), 40.1, 38.8, 38.2 (2C), 24.6, 24.1, 23.9, 23.2 ppm. HRMS (FTMS +p MALDI): m/z = 561.2843 [M + H]^+^; calculated for C30H38ClN8O: 561.2852. HPLC: tR = 6.06 min; purity ≥ 95% (UV: 280/310 nm). LCMS (ESI+): m/z = 561.3 [M + H]^+^; calculated: 561.3.

#### tert-Butyl (4-(6-(2-chloro-4-(6-methylpyridin-2-yl)phenyl)-2-((4-(3-hydroxypropyl)phenyl) amino)-7-oxopyrido[2,3-d]pyrimidin-8(7H)-yl)butyl)carbamate (**64**)

Compound **64** was prepared according to general procedure II, using **52** (50 mg, 0.09 mmol) as starting material. The obtained remaining residue was dissolved in ethanol (5 mL), and 3-(4-aminophenyl)propan-1-ol (53 mg, 0.35 mmol) was added. The reaction solution was heated to 85 °C for 17 h. Afterwards, the solvent was evaporated under reduced pressure, and the crude product was purified by flash chromatography on silica gel using DCM/MeOH as eluent. **64** was obtained as a yellow solid in a yield of 63% (37 mg) over two steps. ^1^H NMR (500 MHz, DMSO-d6, 300 K): *δ* = 10.12 (s, 1H), 8.83 (s, 1H), 8.23 (d, *J* = 1.7 Hz, 1H), 8.09 (dd, *J* = 8.0, 1.8 Hz, 1H), 7.91 (s, 1H), 7.87 (d, *J* = 7.8 Hz, 1H), 7.81 (t, *J* = 7.7 Hz, 1H), 7.72 (d, *J* = 8.4 Hz, 2H), 7.52 (d, *J* = 8.0 Hz, 1H), 7.28 (d, *J* = 7.6 Hz, 1H), 7.21 (d, *J* = 8.3 Hz, 2H), 6.83 (t, *J* = 5.7 Hz, 1H), 4.44 (t, *J* = 5.1 Hz, 1H), 4.34 - 4.29 (m, 2H), 3.45 - 3.41 (m, 2H), 3.01 - 2.95 (m, 2H), 2.61 - 2.57 (m, 5H), 1.74 - 1.68 (m, 4H), 1.54 - 1.49 (m, 2H), 1.35 (s, 9H) ppm. ^13^CNMR (126 MHz, DMSO-d6, 300 K): *δ* = 160.8, 159.3, 159.0, 158.0, 155.6, 154.6, 153.4, 140.1, 137.7, 137.2, 136.5, 136.1, 135.9, 133.7, 132.2, 128.4 (2C), 126.8, 126.0, 124.8, 122.7, 119.6 (2C), 117.6, 105.6, 77.4, 60.2, 40.7, 40.1, 34.5, 31.1, 28.2 (3C), 27.5, 25.0, 24.3 ppm. MS (ESI+): m/z = 569.3 [M-BOC + H]^+^; calculated: 569.2.

#### tert-Butyl (4-(6-(2-chloro-4-(6-methylpyridin-2-yl)phenyl)-2-((4-hydroxyphenyl)amino)-7-oxopyrido[2,3-d]pyrimidin-8(7H)-yl)butyl)carbamate (**65**)

Compound **65** was prepared according to general procedure II, using **52** (50 mg, 0.09 mmol) as starting material. The obtained remaining residue was dissolved in ethanol (5 mL), and 4-aminophenol (77 mg, 0.71 mmol) was added. The reaction solution was heated to 85 °C for 17 h. Afterwards, the solvent was evaporated under reduced pressure, and the crude product was purified by flash chromatography on silica gel using DCM/MeOH as eluent. **65** was obtained as a yellow solid in a yield of 75% (41 mg) over two steps. ^1^H NMR (500 MHz, DMSO-d6, 300 K): *δ* = 9.94 (s, 1H), 9.18 (s, 1H), 8.77 (s, 1H), 8.23 (d, *J* = 1.6 Hz, 1H), 8.08 (dd, *J* = 8.0, 1.6 Hz, 1H), 7.89 - 7.85 (m, 2H), 7.80 (t, *J* = 7.7 Hz, 1H), 7.57 (d, *J* = 7.4 Hz, 2H), 7.51 (d, *J* = 8.0 Hz, 1H), 7.27 (d, *J* = 7.6 Hz, 1H), 6.81 (t, *J* = 5.5 Hz, 1H), 6.77 (d, *J* = 8.8 Hz, 2H), 4.36 - 4.20 (m, 2H), 3.00 - 2.91 (m, 2H), 2.57 (s, 3H), 1.73 - 1.64 (m, 2H), 1.51 - 1.45 (m, 2H), 1.35 (s, 9H) ppm. ^13^CNMR (126 MHz, DMSO-d6, 300 K): *δ* = 160.8, 159.3, 159.1, 158.0, 155.6, 154.7, 153.4, 153.1, 140.0, 137.7, 136.1, 136.0, 133.7, 132.2, 131.0, 126.8, 125.5, 124.8, 122.6, 121.5 (2C), 117.6, 115.0 (2C), 105.3, 77.4, 40.5, 40.1, 28.3 (3C), 27.4, 25.0, 24.3 ppm. MS (ESI+): m/z = 527.2 [M-BOC + H]^+^; calculated: 527.2.

#### tert-Butyl (4-(6-(2-chloro-4-(6-methylpyridin-2-yl)phenyl)-2-((3-hydroxy-4-methylphenyl) amino)-7-oxopyrido[2,3-d]pyrimidin-8(7H)-yl)butyl)carbamate (**66**)

Compound **66** was prepared according to general procedure II, using **52** (50 mg, 0.09 mmol) as starting material. The obtained remaining residue was dissolved in ethanol (5 mL), and 5-amino-2-methylphenol (87 mg, 0.71 mmol) was added. The reaction solution was heated to 85 °C for 17 h. Afterwards, the solvent was evaporated under reduced pressure, and the crude product was purified by flash chromatography on silica gel using DCM/MeOH as eluent. **66** was obtained as a yellow solid in a yield of 74% (42 mg) over two steps. ^1^H NMR (500 MHz, DMSO-d6, 300 K): *δ* = 9.99 (s, 1H), 9.22 (s, 1H), 8.81 (s, 1H), 8.23 (d, *J* = 1.8 Hz, 1H), 8.09 (dd, *J* = 8.0, 1.8 Hz, 1H), 7.90 (s, 1H), 7.88 (d, *J* = 7.8 Hz, 1H), 7.81 (t, *J* = 7.7 Hz, 1H), 7.52 (d, *J* = 8.0 Hz, 1H), 7.34 (s, 1H), 7.28 (d, *J* = 7.7 Hz, 1H), 7.15 (s, 1H), 7.04 (d, *J* = 8.2 Hz, 1H), 6.79 (t, *J* = 5.6 Hz, 1H), 4.33 (t, *J* = 7.2 Hz, 2H), 2.97 - 2.92 (m, 2H), 2.57 (s, 3H), 2.10 (s, 3H), 1.71 - 1.65 (m, 2H), 1.51 - 1.46 (m, 2H), 1.35 (s, 9H) ppm. ^13^CNMR (126 MHz, DMSO-d6, 300 K): *δ* = 160.8, 159.2, 159.0, 158.0, 155.6, 155.2, 154.7, 153.4, 140.0, 138.1, 137.7, 136.0, 136.0, 133.7, 132.2, 130.2, 126.8, 125.9, 124.8, 122.6, 118.4, 117.6, 110.5, 106.7, 105.5, 77.3, 40.7, 40.1, 28.2 (3C), 27.3, 25.0, 24.3, 15.5 ppm. MS (ESI+): m/z = 541.2 [M-BOC + H]^+^; calculated: 541.2.

#### tert-Butyl (4-(6-(2-chloro-4-(6-methylpyridin-2-yl)phenyl)-2-((4-fluoro-3-(hydroxymethyl) phenyl)amino)-7-oxopyrido[2,3-d]pyrimidin-8(7H)-yl)butyl)carbamate (**67**)

Compound **67** was prepared according to general procedure II, using **52** (50 mg, 0.09 mmol) as starting material. The obtained remaining residue was dissolved in ethanol (5 mL), and (5-amino-2-fluorophenyl)methanol (100 mg, 0.71 mmol) was added. The reaction solution was heated to 85 °C for 17 h. Afterwards, the solvent was evaporated under reduced pressure, and the crude product was purified by flash chromatography on silica gel using DCM/MeOH as eluent. **67** was obtained as a yellow solid in a yield of 79% (46 mg) over two steps. ^1^H NMR (500 MHz, DMSO-d6, 300 K): *δ* = 10.20 (s, 1H), 8.84 (s, 1H), 8.23 (d, *J* = 1.8 Hz, 1H), 8.09 (dd, *J* = 8.0, 1.8 Hz, 1H), 7.92 (s, 1H), 7.90 - 7.85 (m, 2H), 7.82 - 7.76 (m, 2H), 7.52 (d, *J* = 8.0 Hz, 1H), 7.28 (d, *J* = 7.6 Hz, 1H), 7.17 (t, *J* = 9.3 Hz, 1H), 6.80 (t, *J* = 5.7 Hz, 1H), 5.28 (t, *J* = 5.5 Hz, 1H), 4.57 (d, *J* = 5.4 Hz, 2H), 4.33 (t, *J* = 7.2 Hz, 2H), 2.96 - 2.89 (m, 2H), 2.57 (s, 3H), 1.72 - 1.65 (m, 2H), 1.51 - 1.45 (m, 2H), 1.34 (s, 9H) ppm. ^13^CNMR (126 MHz, DMSO-d6, 300 K): *δ* = 160.8, 159.3, 159.0, 158.0, 155.6, 155.2 (d, *J* = 240.2 Hz), 154.7, 153.4, 140.1, 137.7, 136.0, 135.9, 135.7 (d, *J* = 2.3 Hz), 133.7, 132.2, 129.2 (d, *J* = 16.0 Hz), 126.8, 126.2, 124.8, 122.6, 120.57 - 119.95 (m), 119.69 - 119.24 (m), 117.6, 114.7 (d, *J* = 22.4 Hz), 105.8, 77.4, 56.8 (d, *J* = 3.7 Hz), 40.7, 40.1, 28.2 (3C), 27.4, 25.0, 24.3 ppm. MS (ESI+): m/z = 559.2 [M-BOC + H]^+^; calculated: 559.2.

#### tert-Butyl (4-(6-(2-Chloro-4-(6-methylpyridin-2-yl)phenyl)-2-((2-hydroxy-5-methoxyphenyl)amino)-7-oxopyrido[2,3-d]pyrimidin-8(7H)-yl)butyl)carbamate (**68**)

Compound **68** was prepared according to general procedure II, using **52** (50 mg, 0.09 mmol) as starting material. The obtained remaining residue was dissolved in ethanol (5 mL), and 2-amino-4-methoxyphenol (98 mg, 0.71 mmol) was added. The reaction solution was heated to 85 °C for 17 h. Afterwards, the solvent was evaporated under reduced pressure, and the crude product was purified by flash chromatography on silica gel using DCM/MeOH as eluent. **68** was obtained as a dark orange solid in a yield of 64% (37 mg) over two steps. ^1^H NMR (500 MHz, DMSO-d6, 300 K): *δ* = 8.98 (s, 1H), 8.24 (d, *J* = 1.7 Hz, 1H), 8.11 (dd, *J* = 8.0, 1.7 Hz, 1H), 8.07 (s, 1H), 7.88 (d, *J* = 7.8 Hz, 1H), 7.81 (t, *J* = 7.7 Hz, 1H), 7.52 (d, *J* = 8.0 Hz, 1H), 7.31 - 7.27 (m, 1H), 6.90 (d, *J* = 8.7 Hz, 1H), 6.73 (t, *J* = 5.6 Hz, 1H), 6.40 (d, *J* = 2.9 Hz, 1H), 6.17 (dd, *J* = 8.8, 2.9 Hz, 1H), 5.01 - 4.89 (m, 2H), 4.11 (t, *J* = 7.2 Hz, 2H), 3.69 (s, 3H), 2.90 - 2.83 (m, 2H), 2.57 (s, 3H), 1.60 - 1.49 (m, 2H), 1.36 (s, 9H), 1.32 - 1.26 (m, 2H) ppm. ^13^CNMR (126 MHz, DMSO-d6, 300 K): *δ* = 164.5, 160.5, 160.5, 158.0, 157.5, 155.6, 155.5, 153.3, 141.6, 140.4, 137.7, 135.4, 135.4, 133.5, 133.3, 132.1, 129.3, 126.9, 124.9, 122.7, 122.5, 117.7, 108.9, 101.3, 100.9, 77.4, 55.0, 40.7, 40.1, 28.3 (3C), 27.0, 24.8, 24.3 ppm. MS (ESI+): m/z = 557.3 [M-BOC + H]^+^; calculated: 557.2.

#### tert-Butyl (4-(6-(2-chloro-4-(6-methylpyridin-2-yl)phenyl)-2-((2-methoxy-4-morpholinophenyl)amino)-7-oxopyrido[2,3-d]pyrimidin-8(7H)-yl)butyl)carbamate (**69**)

Compound **69** was prepared according to general procedure II, using **52** (50 mg, 0.09 mmol) as starting material. The obtained remaining residue was dissolved in ethanol (5 mL), and 2-methoxy-4-morpholinoaniline (147 mg, 0.71 mmol) was added. The reaction solution was heated to 85 °C for 17 h. Afterwards, the solvent was evaporated under reduced pressure, and the crude product was purified by flash chromatography on silica gel using DCM/MeOH as eluent. **69** was obtained as a purple solid in a yield of 64% (41 mg) over two steps. ^1^H NMR (500 MHz, DMSO-d6, 300 K): *δ* = 8.74 (s, 1H), 8.67 (s, 1H), 8.22 (d, *J* = 1.7 Hz, 1H), 8.08 (dd, *J* = 8.0, 1.7 Hz, 1H), 7.88 - 7.84 (m, 2H), 7.80 (t, *J* = 7.7 Hz, 1H), 7.72 (s, 1H), 7.50 (d, *J* = 8.0 Hz, 1H), 7.27 (d, *J* = 7.6 Hz, 1H), 6.78 (t, *J* = 5.5 Hz, 1H), 6.67 (d, *J* = 2.5 Hz, 1H), 6.58 (d, *J* = 8.0 Hz, 1H), 4.18 (s, 2H), 3.83 (s, 3H), 3.77 - 3.74 (m, 4H), 3.15 - 3.12 (m, 4H), 2.96 - 2.88 (m, 2H), 2.56 (s, 3H), 1.65 - 1.55 (m, 2H), 1.47 - 1.40 (m, 2H), 1.36 (s, 9H) ppm. ^13^CNMR (126 MHz, DMSO-d6, 300 K): *δ* = 160.7, 159.7, 159.3, 158.0, 155.6, 154.7, 153.4, 151.9, 149.1, 140.0, 137.7, 136.1, 136.0, 133.7, 132.2, 126.8, 125.6, 124.8, 123.5, 122.6, 119.6, 117.6, 106.4, 105.3, 99.6, 77.4, 66.2 (2C), 55.6, 48.9 (2C), 40.4, 40.1, 28.2 (3C), 27.3, 24.9, 24.3 ppm. MS (ESI+): m/z = 626.3 [M-BOC + H]^+^; calculated: 626.3.

#### tert-Butyl (4-(6-(2-chloro-4-(6-methylpyridin-2-yl)phenyl)-2-((2-methoxy-4-(4-methylpiperazin-1-yl)phenyl)amino)-7-oxopyrido[2,3-d]pyrimidin-8(7H)-yl)butyl)carbamate (**70**)

Compound **70** was prepared according to general procedure II, using **52** (85 mg, 0.15 mmol) as starting material. The obtained remaining residue was dissolved in ethanol (5 mL), and 2-methoxy-4-(4-methylpiperazin-1-yl)aniline (100 mg, 0.45 mmol) was added. The reaction solution was heated to 85 °C for 17 h. Afterwards, the solvent was evaporated under reduced pressure, and the crude product was purified by flash chromatography on silica gel using H2O/ACN as eluent. **70** was obtained as an orange solid in a yield of 23% (25 mg) over two steps. ^1^H NMR (500 MHz, DMSO-d6, 300 K): *δ* = 8.75 (s, 1H), 8.67 (s, 1H), 8.22 (d, *J* = 1.7 Hz, 1H), 8.08 (dd, *J* = 8.0, 1.8 Hz, 1H), 7.88 - 7.86 (m, 2H), 7.81 (t, *J* = 7.7 Hz, 1H), 7.70 (s, 1H), 7.51 (d, *J* = 8.0 Hz, 1H), 7.28 (d, *J* = 7.6 Hz, 1H), 6.78 (t, *J* = 5.5 Hz, 1H), 6.66 (d, *J* = 2.5 Hz, 1H), 6.56 (d, *J* = 8.4 Hz, 1H), 4.25 - 4.16 (m, 2H), 3.83 (s, 3H), 3.19 - 3.14 (m, 4H), 2.95 - 2.88 (m, 2H), 2.57 (s, 3H), 2.48 - 2.46 (m, 4H), 2.23 (s, 3H), 1.64 - 1.56 (m, 2H), 1.44 - 1.39 (m, 2H), 1.36 (s, 9H) ppm. ^13^CNMR (126 MHz, DMSO-d6, 300 K): *δ* = 160.7, 159.3, 158.8, 158.0, 155.6, 154.7, 153.4, 152.2, 149.1, 140.0, 137.7, 136.2, 136.1, 136.0, 133.7, 132.2, 126.8, 125.5, 124.8, 122.6, 119.2, 117.6, 106.6, 105.3, 99.8, 77.4, 55.6 (2C), 54.7, 48.5, 45.8 (2C), 40.1 (2C), 28.3 (3C), 27.3, 24.9, 24.3 ppm. MS (ESI+): m/z = 739.4 [M + H]^+^; calculated: 739.4.

#### 8-(4-Aminobutyl)-6-(2-chloro-4-(6-methylpyridin-2-yl)phenyl)-2-((4-(3-hydroxypropyl)phenyl)amino)pyrido[2,3-d]pyrimidin-7(8H)-one (**30**)

Compound **30** was prepared according to general procedure IV, using **64** (37 mg, 0.06 mmol) as starting material. The crude product was purified by HPLC chromatography on silica gel (H2O/ACN; 0.1% TFA) to obtain **30** as a yellow solid in a yield of 59% (26 mg). ^1^H NMR (500 MHz, DMSO-d6, 300 K): (mixture of rotamers) *δ* = 10.17 (d, *J* = 11.8 Hz, 1H), 8.85 (d, *J* = 3.5 Hz, 1H), 8.24 (d, *J* = 1.4 Hz, 1H), 8.10 (dd, *J* = 8.0, 1.3 Hz, 1H), 7.95 (d, *J* = 2.4 Hz, 1H), 7.90 - 7.83 (m, 2H), 7.80 - 7.70 (m, 5H), 7.53 (d, *J* = 8.0 Hz, 1H), 7.31 (d, *J* = 7.5 Hz, 1H), 7.22 (dd, *J* = 13.0, 8.5 Hz, 2H), 5.20 (s, 1H), 4.42 - 4.36 (m, 3H), 3.43 (t, *J* = 6.4 Hz, 1H), 2.90 - 2.83 (m, 2H), 2.67 (t, *J* = 7.6 Hz, 1H), 2.62 - 2.59 (m, 1H), 2.58 (s, 3H), 2.11 - 1.98 (m, 1H), 1.88 - 1.79 (m, 2H), 1.74 - 1.63 (m, 3H) ppm. ^13^CNMR (126 MHz, DMSO-d6, 300 K): (mixture of rotamers) *δ* = 160.9, 159.4, 157.9, 154.7, 153.2, 139.8, 138.0, 137.6, 137.2, 136.5, 136.0 (d, *J* = 1.9 Hz), 134.8, 133.7, 132.3, 128.4 (d, *J* = 2.6 Hz, 2C), 127.0, 126.0 (d, *J* = 10.9 Hz), 124.9, 122.8, 119.6 (d, *J* = 7.0 Hz, 2C), 117.9, 105.7, 67.9, 60.1, 40.3, 38.7, 34.4, 31.1, 30.4, 29.2, 24.7, 24.5, 24.1 ppm. HRMS (FTMS +p MALDI): m/z = 569.2419 [M + H]^+^; calculated for C32H34ClN6O2: 569.2426. HPLC: tR = 7.17 min; purity ≥ 95% (UV: 280/310 nm). LCMS (ESI+): m/z = 569.3 [M + H]^+^; calculated: 569.2.

#### 8-(4-Aminobutyl)-6-(2-chloro-4-(6-methylpyridin-2-yl)phenyl)-2-((4-hydroxyphenyl)amino) pyrido[2,3-d]pyrimidin-7(8H)-one (**31**)

Compound **31** was prepared according to general procedure IV, using **65** (16 mg, 0.03 mmol) as starting material. **31** was obtained as a yellow solid in a yield of 95% (18 mg). ^1^H NMR (500 MHz, DMSO-d6, 300 K): *δ* = 9.99 (s, 1H), 8.80 (s, 1H), 8.21 (d, *J* = 1.7 Hz, 1H), 8.07 (dd, *J* = 8.0, 1.8 Hz, 1H), 7.97 - 7.90 (m, 3H), 7.74 (s, 3H), 7.61 - 7.53 (m, 3H), 7.40 (dd, *J* = 6.1, 2.4 Hz, 1H), 6.99 (s, 1H), 6.80 (d, *J* = 8.7 Hz, 2H), 4.34 (t, *J* = 7.0 Hz, 2H), 2.90 - 2.80 (m, 2H), 2.61 (s, 3H), 1.84 - 1.73 (m, 2H), 1.67 - 1.59 (m, 2H) ppm. ^13^CNMR (126 MHz, DMSO-d6, 300 K): *δ* = 160.9, 159.4, 159.1, 157.5, 154.8, 153.3, 152.8, 139.2, 138.7, 136.5, 136.3, 133.8, 132.4, 131.0, 127.3, 125.4, 125.3, 123.4, 121.6 (2C), 118.7, 115.2 (2C), 105.3, 40.1, 38.8, 24.7, 24.5, 23.5 ppm. HRMS (FTMS +p MALDI): m/z = 527.1946 [M + H]^+^; calculated for C29H28ClN6O2: 527.1957. HPLC: tR = 6.94 min; purity ≥ 95% (UV: 280/310 nm). LCMS (ESI+): m/z = 527.2 [M + H]^+^; calculated: 527.2.

#### 8-(4-Aminobutyl)-6-(2-chloro-4-(6-methylpyridin-2-yl)phenyl)-2-((3-hydroxy-4-methylphenyl)amino)pyrido[2,3-d]pyrimidin-7(8H)-one (**32**)

Compound **32** was prepared according to general procedure IV, using **66** (38 mg, 0.06 mmol) as starting material. **32** was obtained as a yellow solid in a yield of 96% (43 mg). ^1^H NMR (500 MHz, DMSO-d6, 300 K): *δ* = 10.04 (s, 1H), 8.84 (s, 1H), 8.59 (s, 1H), 8.23 (d, *J* = 1.8 Hz, 1H), 8.08 (dd, *J* = 8.0, 1.8 Hz, 1H), 7.99 - 7.92 (m, 3H), 7.72 (s, 3H), 7.56 (d, *J* = 8.0 Hz, 1H), 7.42 (dd, *J* = 6.9, 1.4 Hz, 1H), 7.29 (s, 1H), 7.21 (s, 1H), 7.04 (d, *J* = 8.3 Hz, 1H), 4.40 (t, *J* = 7.1 Hz, 2H), 2.89 - 2.79 (m, 2H), 2.62 (s, 3H), 2.11 (s, 3H), 1.84 - 1.75 (m, 2H), 1.67 - 1.60 (m, 2H) ppm. ^13^CNMR (126 MHz, DMSO-d6, 300 K): *δ* = 160.9, 159.3, 159.0, 157.4, 155.3, 154.8, 152.7, 139.4, 138.6, 138.1, 136.5, 136.3, 133.8, 132.4, 130.2, 127.4, 125.8, 125.4, 123.5, 118.9, 118.5, 110.5, 106.7, 105.6, 40.3, 38.8, 24.7, 24.5, 23.4, 15.6 ppm. HRMS (FTMS +p MALDI): m/z = 541.2107 [M + H]^+^; calculated for C30H30ClN6O2: 541.2113. HPLC: tR = 7.17 min; purity ≥ 95% (UV: 280/310 nm). LCMS (ESI+): m/z = 541.2 [M + H]^+^; calculated: 541.2.

#### 8-(4-Aminobutyl)-6-(2-chloro-4-(6-methylpyridin-2-yl)phenyl)-2-((4-fluoro-3-(hydroxymethyl)phenyl)amino)pyrido[2,3-d]pyrimidin-7(8H)-one (**33**)

Compound **33** was prepared according to general procedure IV, using **67** (38 mg, 0.06 mmol) as starting material. The crude product was purified by HPLC chromatography on silica gel (H2O/ACN; 0.1% TFA) to obtain **33** as a dark yellow solid in a yield of 56% (25 mg). ^1^H NMR (500 MHz, DMSO-d6, 300 K): (mixture of rotamers) *δ* = 10.36 (s, 0.3H), 10.26 (s, 0.7H), 9.61 (s, 1H), 8.89 (s, 0.3H), 8.87 (s, 0.7H), 8.24 (s, 1H), 8.08 (d, *J* = 8.0 Hz, 1H), 7.99 - 7.91 (m, 4.3H), 7.78 - 7.66 (m, 3.7H), 7.56 (d, *J* = 8.0 Hz, 1H), 7.42 (d, *J* = 6.6 Hz, 1H), 7.34 (t, *J* = 9.2 Hz, 0.3H), 7.17 (t, *J* = 9.3 Hz, 0.7H), 5.53 (s, 0.6H), 4.59 (s, 1.4H), 4.39 (t, *J* = 6.7 Hz, 2H), 2.90 - 2.78 (m, 2H), 2.62 (s, 3H), 1.84 - 1.76 (m, 2H), 1.68 - 1.60 (m, 2H) ppm. ^13^CNMR (126 MHz, DMSO-d6, 300 K): (mixture of rotamers) *δ* = 160.9, 159.5, 159.0, 157.4, 155.6 (d, *J* = 210.8 Hz), 155.2 (d, *J* = 239.8 Hz), 154.8, 152.7, 139.4, 138.8 - 138.6 (m), 136.5 - 136.2 (m, 2C), 136.1 (d, *J* = 2.3 Hz), 135.8(d, *J* = 2.4 Hz), 133.8, 132.3, 129.3 (d, *J* = 15.8 Hz), 127.4, 126.0, 125.4, 123.5, 122.4 (d, *J* = 8.9 Hz), 122.2 (d, *J* = 2.2 Hz), 120.7 (d, *J* = 15.5 Hz), 120.1 (d, *J* = 2.1 Hz), 119.8 - 119.4 (m), 118.8, 115.8 (d, *J* = 22.0 Hz), 114.8 (d, *J* = 22.4 Hz), 105.8, 64.1 (d, *J* = 3.1 Hz), 56.8 (d, *J* = 3.7 Hz), 40.3, 38.8, 24.7, 24.6, 23.4 ppm. HRMS (FTMS +p MALDI): m/z = 559.2010 [M + H]^+^; calculated for C30H29ClFN6O2: 559.2019. HPLC: tR = 7.25 min; purity ≥ 95% (UV: 280/310 nm). LCMS (ESI+): m/z = 559.2 [M + H]^+^; calculated: 559.2.

#### 8-(4-Aminobutyl)-6-(2-chloro-4-(6-methylpyridin-2-yl)phenyl)-2-((2-hydroxy-5-methoxy-phenyl)amino)pyrido[2,3-d]pyrimidin-7(8H)-one (**34**)

Compound **34** was prepared according to general procedure IV, using **68** (30 mg, 0.05 mmol) as starting material. **34** was obtained as a dark red solid in a yield of 92% (33 mg). ^1^H NMR (500 MHz, DMSO-d6, 300 K): *δ* = 10.53 (s, 1H), 8.86 (s, 1H), 8.60 (s, 1H), 8.23 (d, *J* = 1.8 Hz, 1H), 8.07 (dd, *J* = 8.0, 1.8 Hz, 1H), 8.03 - 7.96 (m, 3H), 7.77 - 7.65 (m, 4H), 7.57 (d, *J* = 8.0 Hz, 1H), 7.45 (d, *J* = 7.5 Hz, 1H), 6.86 (d, *J* = 8.7 Hz, 1H), 6.57 (dd, *J* = 8.8, 3.0 Hz, 1H), 4.36 (t, *J* = 7.0 Hz, 2H), 3.73 (s, 3H), 2.86 - 2.80 (m, 2H), 2.63 (s, 3H), 1.82 - 1.75 (m, 2H), 1.65 - 1.58 (m, 2H) ppm. ^13^CNMR (126 MHz, DMSO-d6, 300 K): *δ* = 160.9, 159.4, 158.8, 157.3, 154.9, 152.5, 152.2, 141.5, 139.8, 138.2, 136.5, 136.3, 133.8, 132.3, 127.5, 127.2, 126.3, 125.5, 123.7, 119.2, 115.2, 108.1, 107.6, 106.0, 55.5, 40.2, 38.7, 24.5, 24.5, 23.1 ppm. HRMS (FTMS +p MALDI): m/z = 557.2049 [M + H]^+^; calculated for C30H30ClN6O3: 557.2062. HPLC: tR = 7.45 min; purity ≥ 95% (UV: 280/310 nm). LCMS (ESI+): m/z = 557.2 [M + H]^+^; calculated: 557.2.

#### 8-(4-Aminobutyl)-6-(2-chloro-4-(6-methylpyridin-2-yl)phenyl)-2-((2-methoxy-4-morpholino-phenyl)amino)pyrido[2,3-d]pyrimidin-7(8H)-one (**35**)

Compound **35** was prepared according to general procedure IV, using **69** (32 mg, 0.04 mmol) as starting material. **35** was obtained as a dark green solid in a yield of 85% (32 mg). ^1^H NMR (500 MHz, DMSO-d6, 300 K): *δ* = 8.79 (d, *J* = 8.2 Hz, 2H), 8.24 - 8.20 (m, 1H), 8.08 - 8.02 (m, 2H), 7.99 (d, *J* = 7.8 Hz, 1H), 7.92 (s, 1H), 7.83 - 7.65 (m, 4H), 7.57 (d, *J* = 8.0 Hz, 1H), 7.49 (d, *J* = 7.7 Hz, 1H), 6.78 (d, *J* = 2.4 Hz, 1H), 6.64 (dd, *J* = 8.8, 2.4 Hz, 1H), 4.28 (t, *J* = 6.6 Hz, 2H), 3.85 (s, 3H), 3.81 - 3.77 (m, 4H), 3.22 - 3.18 (m, 4H), 2.87 - 2.78 (m, 2H), 2.65 (s, 3H), 1.78 - 1.69 (m, 2H), 1.62 - 1.54 (m, 2H) ppm. ^13^CNMR (126 MHz, DMSO-d6, 300 K): *δ* = 160.9, 159.4, 159.1, 157.1, 155.0, 152.3, 152.0, 148.1, 140.4, 137.7, 136.8, 136.3, 133.8, 132.5, 132.4, 127.7, 125.6, 124.0, 123.7, 120.4, 119.5, 107.1, 105.6, 100.3, 66.0 (2C), 55.8, 49.5 (2C), 40.1, 38.7, 24.7, 24.5, 22.8 ppm. HRMS (FTMS +p MALDI): m/z = 626.2624 [M + H]^+^; calculated for C34H37ClN7O3: 626.2641. HPLC: tR = 7.80 min; purity ≥ 95% (UV: 280/310 nm). LCMS (ESI+): m/z = 626.3 [M + H]^+^; calculated: 626.3.

#### 8-(4-Aminobutyl)-6-(2-chloro-4-(6-methylpyridin-2-yl)phenyl)-2-((2-methoxy-4-(4-methylpiperazin-1-yl)phenyl)amino)pyrido[2,3-d]pyrimidin-7(8H)-one (**36**)

Compound **36** was prepared according to general procedure IV, using **70** (23 mg, 0.03 mmol) as starting material. The crude product was purified by HPLC chromatography on silica gel (H2O/ACN; 0.1% TFA) to obtain **36** as an orange solid in a yield of 11% (3 mg). ^1^H NMR (500 MHz, DMSO-d6, 300 K): *δ* = 9.85 (s, 1H), 8.77 (s, 1H), 8.74 (s, 1H), 8.23 (d, *J* = 1.8 Hz, 1H), 8.10 (dd, *J* = 8.1, 1.8 Hz, 1H), 7.92 (s, 1H), 7.88 (d, *J* = 7.8 Hz, 1H), 7.82 (t, *J* = 7.2 Hz, 1H), 7.78 - 7.61 (m, 3H), 7.51 (d, *J* = 8.0 Hz, 1H), 7.30 (d, *J* = 7.6 Hz, 1H), 6.75 (d, *J* = 2.4 Hz, 1H), 6.61 (d, *J* = 8.6 Hz, 1H), 4.30 (t, *J* = 5.8 Hz, 2H), 3.88 (d, *J* = 13.5 Hz, 2H), 3.85 (s, 3H), 3.55 (d, *J* = 12.1 Hz, 2H), 3.23 - 3.13 (m, 2H), 2.98 (t, *J* = 12.4 Hz, 2H), 2.89 (s, 3H), 2.86 - 2.80 (m, 2H), 2.57 (s, 3H), 1.82 - 1.69 (m, 2H), 1.61 - 1.53 (m, 2H) ppm. ^13^CNMR (126 MHz, DMSO-d6, 300 K): *δ* = 160.9, 159.4, 158.5, 158.0, 156.3, 154.8, 153.3, 152.2, 149.9, 140.0, 137.8, 136.3, 136.2, 135.9, 133.7, 132.2, 126.9, 125.7, 124.8, 122.7, 120.0, 117.7, 107.2, 100.7, 55.8, 52.4 (2C), 46.1 (2C), 42.1, 39.8, 38.7, 24.6, 24.5, 24.3 ppm. HRMS (FTMS +p MALDI): m/z = 638.2863 [M]^+^; calculated for C35H39ClN8O2: 638.2879. HPLC: tR = 6.87 min; purity ≥ 95% (UV: 280/310 nm). LCMS (ESI+): m/z = 639.4 [M + H]^+^; calculated: 639.3.

### Statistical analyses and figures

Structural images were generated using PyMol 2.4.3 (*Schrödinger*), and graphs were plotted using GraphPad Prism version 8.4 (*GraphPad Software*, San Diego, California, USA, www.graphpad.com).

### NanoBRET assay

The assay was performed as previously described.^61^ In brief: full-length kinases were obtained as plasmids cloned in frame with a terminal NanoLuc-fusion (*Promega*) as specified in Table S4. Plasmids were transfected into HEK293T cells using FuGENE HD (*Promega*, E2312), and proteins were allowed to express for 20 h. Serially diluted inhibitor and NanoBRET Kinase Tracer K8/K9/K10 (*Promega*), at a concentration determined previously as the Tracer KD,app (Table S4), were pipetted into white 384-well plates (*Greiner* 781207) using an Echo acoustic dispenser (*Labcyte*). The corresponding protein-transfected cells were added and reseeded at a density of 2 x 10^5^ cells/mL after trypsinization and resuspending in Opti-MEM without phenol red (*Life Technologies*). The system was allowed to equilibrate for 2 h at 37 °C/5% CO2 prior to BRET measurements. To measure BRET, NanoBRET NanoGlo Substrate + Extracellular NanoLuc Inhibitor (*Promega*, N2540) was added as per the manufacturer’s protocol, and filtered luminescence was measured on a PHERAstar plate reader (*BMG Labtech*) equipped with a luminescence filter pair (450 nm BP filter (donor) and 610 nm LP filter (acceptor)). For permeabilized-mode NanoBRET experiments, digitonin (Promega, #G9441) was added as per the manufacturer’s instructions to a final concentration of 50 ng/mL. Competitive displacement data were then graphed using GraphPad Prism 8 software using a normalized 3-parameter curve fit with the following equation: Y=100/(1+10^(X-LogEC50)).

### DSF-based selectivity screening against a curated kinase library

The assay was performed as previously described.^62,63^ Briefly, recombinant protein kinase domains at a concentration of 2 μM were mixed with 10 μM compound in a buffer containing 20 mM HEPES, pH 7.5, and 500 mM NaCl. SYPRO Orange (5000×, *Invitrogen*) was added as a fluorescence probe (1 µl per mL). Subsequently, temperature-dependent protein unfolding profiles were measured using the QuantStudio™ 5 realtime PCR machine (*Thermo Fisher Scientific*). Excitation and emission filters were set to 465 nm and 590 nm, respectively. The temperature was raised with a step rate of 3 °C per minute. Data points were analyzed with the internal software (Thermal Shift Software^TM^ Version 1.4, *Thermo Fisher Scientific*) using the Boltzmann equation to determine the inflection point of the transition curve. Differences in melting temperature are given as Δ*T*m values in °C.

### Protein expression and purification

MST3 (STK24) was expressed and purified as previously described by Tesch *et al.*^40^ MST3 (R4-D301) was subcloned in pNIC-CH vector (non-cleavable C-terminal His6-tag). The expression plasmids were transformed in *Escherichia coli* Rosetta BL21(D3)-R3-pRARE2 and BL21(D3)-R3-λPP competent cells, respectively. Initially, cells were cultured in Terrific Broth (TB) media at 37 °C to an optical density (OD) 2.8 prior to induction with 0.5 mM IPTG at 18 °C overnight. Cells were harvested and resuspended in a buffer containing 50 mM HEPES pH 7.5, 500 mM NaCl, 30 mM imidazole, 5% glycerol, and 0.5 mM TCEP. The recombinant protein was initially purified by Ni^2+^-affinity chromatography. The Ni-NTA fraction containing MST3 was concentrated using a 30 kDa cutoff ultrafiltration device and further purified by size exclusion chromatography (SEC; HiLoad 16/600 Superdex 200) pre-equilibrated with SEC buffer (25 mM HEPES pH 7.5, 200 mM NaCl, 0.5 mM TCEP, 5% glycerol). Quality control was performed by SDS-gel electrophoresis and ESI-MS (MST3: expected 34,507.7 Da, observed 34,506.9 Da).

### Crystallization and structure determination

Co-crystallization trials were performed using the sitting-drop vapor-diffusion method at 293 K with a mosquito crystallization robot (*TTP Labtech*, Royston UK). MST3 protein (11 mg/mL in 25 mM HEPES pH 7.5, 200 mM NaCl, 0.5 mM TCEP, 5% glycerol) was incubated with inhibitor at a final concentration of 1 mM prior to setting up crystallization trials. Crystals grew with the following reservoir buffers: MRLW5: 12% PEG 6000, 0.1 M HEPES, pH 7.4; MR24: 12% PEG 6000, 0.1 M HEPES, pH 7.4; MR26: 10% PEG 6000, 0.1 M HEPES, pH 7.2; MR30: 24% PEG 3350, 0.1 M citrate, pH 5.2. Crystals were cryo-protected with mother liquor supplemented with 20% ethylene glycol and flash-frozen in liquid nitrogen. X-ray diffraction data sets were collected at 100 K at beamline X06SA of the Swiss Light Source, Villigen, Switzerland. Diffraction data were integrated with the program XDS^64^ and scaled with AMLESS,^65^ which is part of the CCP4 package.^66^ The MST3 structures were solved by difference Fourier analysis using PHENIX^67^ with PDB entry 7B32 as a starting model. Structure refinement was performed using iterative cycles of manual model building in COOT^68^ and refinement in PHENIX. Dictionary files for the compounds were generated using the Grade Web Server (http://grade.globalphasing.org). X-ray data collection and refinement statistics are shown in Table S5.

### Homology modeling and molecular docking

System preparation and docking calculations were performed using the *Schrödinger* Drug Discovery suite for molecular modeling (version 2021.2). Protein−ligand complexes were prepared with the Protein Preparation Wizard^69^ to fix protonation states of amino acids, add hydrogens, and fix missing side-chain atoms. The MST1 model is based on MST1 crystal structure (PDB code 8A5J), the missing part of the G-rich loop (E137-G141) was filled with AlfaFold model derived from UniProt sequence Q13043. The MST2 model is based on MST2 crystal structure (PDB code 4LG4), the missing part of the G-rich loop (E35-Y38) was filled with AlfaFold model derived from UniProt sequence Q13043. The MST3 model is based on MST3 crystal structure (PDB code 7B30). The MST4 model is based on MST3 crystal structure (PDB code 7B36), the missing part of the G-rich loop (K32-S34) was filled with AlfaFold model derived from UniProt sequence Q9P289. The models were built as a multi-template composite/chimera type with Prime,^70^ followed by hydrogen bond assignment and energy minimization with Protein Preparation Wizard. All ligands for docking were drawn using Maestro and prepared using LigPrep^71^ to generate the 3D conformation, adjust the protonation state to physiological pH (7.4), and calculate the partial atomic charges with the OPLS4^72^ force field. Docking studies with the prepared ligands were performed using Glide (Glide V7.7)^73,74^ with the flexible modality of induced-fit docking with extra precision (XP), followed by a side-chain minimization step using Prime. Ligands were docked within a grid around 12 Å from the centroid of the co-crystallized ligand, generating ten poses per ligand.

### Molecular dynamics simulation

MD simulations were carried out using Desmond^75^ with the OPLS4 force-field.^72^ The simulated system encompassed the protein-ligand complexes, a predefined water model (TIP3P)^76^ as a solvent, and counter ions (5 Na^+^ adjusted to neutralize the overall system charge in MST4 bound simulations, 4 Na^+^ in MST3, and 3 Na^+^ in MST2 and MST1 simulations). The system was treated in a cubic box with periodic boundary conditions specifying the box’s shape and size as 10 Å distance from the box edges to any atom of the protein. In all simulations, we used a time step of 1 fs, the short-range coulombic interactions were treated using a cut-off value of 9.0 Å using the short-range method, while the Smooth Particle Mesh Ewald method (PME) handled long-range coulombic interactions.^77^ Initially, the system’s relaxation was performed using Steepest Descent and the limited-memory Broyden-Fletcher-Goldfarb-Shanno algorithms in a hybrid manner, according to the established protocol available in the Desmond standard settings. During the equilibration step, the simulation was performed under the NPT ensemble for 5 ns implementing the Berendsen thermostat and barostat methods.^78^ A constant temperature of 310 K was kept throughout the simulation using the Nosé-Hoover thermostat algorithm,^79^ and the Martyna-Tobias-Klein Barostat algorithm^80^ was used to maintain 1 atm of pressure. After minimization and relaxation of the system, we continued with the production step of at least 500 ns, with frames being recorded/saved every 1,000 ps. Five independent replicas with the length of 1 µs were produced for each ligand-protein system, resulting in a total of 10 µs simulation per protein (MR24 bound to MST1 (1 µs x 5 replicas) + MR26 bound to MST1 (1 µs x 5 replicas) = 10 µs). Trajectories and interaction data are available on the Zenodo repository (DOI: 10.5281/zenodo.7638701). The representative structures were selected by inspecting changes in the Root-mean-square deviation (RMSD). Figure S6 represents the fluctuation of RMSD values of the ligand-protein complexes along the MD trajectory, divided by replicas (1–5 accordingly). Finally, protein-ligand interactions were determined using the Simulation Event Analysis pipeline implemented in Maestro (Maestro v2021.3). The ligands’ binding energy was calculated using the Born and surface area continuum solvation (MM/GBSA) model, using Prime^70^ and the implemented thermal MM/GBSA script. Calculated free-binding energies are represented by the MM/GBSA.

### PK-prediction

QikProp (implemented in the Schrödinger Drug Discovery suite for molecular modeling, version 2021.2) was used to predict pharmacokinetic properties, and acceptable limits were derived from their manual references.

### Viability assessment

To assess cell viability, a live-cell assay based on nuclear morphology was performed as previously described.^52^ In brief, HCT116 cells were stained with 60 nM Hoechst33342 (*Thermo Fischer Scientific*) and seeded at a density of 1200 cells per well (Cell culture microplate, PS, f-bottom, µClear, 781091, *Greiner*) in culture medium (50 µL per well) in a 384 well plate. After 6 h, 12 h, 24 h, 48 h, and 72 h of compound treatment, fluorescence and cellular shape was measured using the CQ1 high-content confocal microscope (*Yokogawa*). Compounds were added directly to the cells at three different concentrations (1 µM, 5 µM, and 10 µM). The following parameters were used for image acquisition: Ex 405 nm/Em 447/60 nm, 500ms, 50%; Ex 561 nm/Em 617/73 nm, 100 ms, 40%; Ex 488/Em 525/50 nm, 50 ms, 40%; Ex 640 nm/Em 685/40, 50 ms, 20 %; bright field, 300ms, 100% transmission, one centered field per well, 7 z stacks per well with 55 µm spacing. Analysis of images was performed using the CellPathfinder software (*Yokogawa*) as previously described.^81^ The cell count was normalized against the cell count of cells treated with DMSO (0.1%). Cells showing Hoechst High Intensity Objects were detected. All normal gated cells were further classified into cells containing healthy, fragmented, or pyknosed nuclei. Error bars show SEM of biological duplicates. All data can be found in Supplementary Table S8.

### FUCCI cell-cycle assay

To test the influence of the compounds on cell cycle, we performed a fluorescent ubiquitination-based cell-cycle indicator (FUCCI) assay as previously described.^52^ In brief, HCT116-FUCCI cells, stably expressing the FUCCI system, introduced by the Sleeping Beauty transposon system,^82^ were seeded at a density of 1200 cells per well in a 384 well plate (Cell culture microplate, PS, f-bottom, µClear, 781091, *Greiner*) in culture medium (50 µL per well) and stained additionally with 60 nM Hoechst33342 (*Thermo Fischer Scientific*). Fluorescence and cellular shape were measured 6 h, 12 h, 24 h, 48 h, and 72 h after compound treatment using the CQ1 high-content confocal microscope (*Yokogawa*). Compounds were added directly to the cells at three different concentrations (1 µM, 5 µM, and 10 µM). The following parameters were used for image acquisition: Ex 405 nm/Em 447/60 nm, 500ms, 50%; Ex 561 nm/Em 617/73 nm, 100 ms, 40%; Ex 488/Em 525/50 nm, 50 ms, 40%; Ex 640 nm/Em 685/40, 50 ms, 20 %; bright field, 300ms, 100% transmission, one centered field per well, 7 z stacks per well with 55 µm spacing. Analysis of images was performed using the CellPathfinder software (*Yokogawa*) as previously described.^81^ All normal gated cells were further classified in cells containing healthy, fragmented, or pyknosed nuclei. Cells that showed a healthy nucleus were gated in red, green, or yellow. Error bars show SEM of biological duplicates. All data can be found in Table S9.

### Microsomal stability

The solubilized test compound (5 μL, final concentration 10 µM) was preincubated at 37 °C in 432 μL of phosphate buffer (0.1 M, pH 7.4) together with 50 μL NADPH regenerating system (30 mM glucose-6-phosphate, 4 U/mL glucose-6-phosphate dehydrogenase, 10 mM NADP, 30 mM MgCl2). After 5 min, the reaction was started by the addition of 13 μL of microsome mix from the liver of Sprague−Dawley rats (Invitrogen; 20 mg protein/mL in 0.1 M phosphate buffer) in a shaking water bath at 37 °C. The reaction was stopped by adding 500 μL of ice-cold methanol at 0, 15, 30, and 60 min. The samples were centrifuged at 5000 g for 5 min at 4 °C, then the supernatants were analyzed, and the test compound was quantified by HPLC. The composition of the mobile phase was adapted to the test compound in a range of MeOH 40-90% and water (0.1% formic acid) 10-60%; flow-rate: 1 mL/min; stationary phase: Purospher STAR, RP18, 5 μm, 125×4, precolumn: Purospher STAR, RP18, 5 μm, 4×4; detection wavelength: 254 and 280 nm; injection volume: 50 μL. Control samples were performed to check the test compound’s stability in the reaction mixture: the first control was without NADPH, which is needed for the enzymatic activity of the microsomes, the second control was with inactivated microsomes (incubated for 20 min at 90 °C), and the third control was without test compound (to determine the baseline). The amounts of the test compound were quantified via an external calibration curve. Data are expressed as the mean ± SEM remaining compound from three independent experiments.

## ASSOCIATED CONTENT

### Supporting Information

Published selectivity profile of SBP-3264 (**3**); Selectivity profile of aurora kinase inhibitor hesperadin (**7**); NanoBRET curves of G-5555 (**8**) against MST1-4; Selectivity data of compounds MRLW5 (**20**), **22**, and **23** tested in our in-house Δ*T*m panel; Bar charts of the selectivity data of compounds **24**–**36** obtained in our in-house DSF panel; RMSD of MR24 (**24**)- and MR26 (**26**)- protein complexes; Selectivity data of compounds MR26 (**26**) and MR30 (**27**); Viability assessment; Absorption spectra of compounds **2**, **3**, **20**, **24**, **26**, and **30**; Cell cycle analysis in living cells; Pharmacokinetic properties of MR24 (**24**); Results of the DSF panel assay of compounds **20**, **22**–**26** and **33**–**36** on 100 kinases; NanoBRET assay information; X-ray data collection and refinement statistics; MR24 (**24**) and MR26 (**26**) interaction patterns derived from MD simulations; Percentage of hydrophobic ligand–protein interactions derived from MD simulations for selected systems; Analytical data of all compounds presented. (PDF) Raw data of cellular viability assessment and FUCCI cell cycle analysis. (Excel) Molecular formula strings (CSV) with activity data are provided.

## AUTHOR INFORMATION

### Corresponding Author

**Stefan Knapp** — Institute of Pharmaceutical Chemistry, Goethe University Frankfurt, Frankfurt am Main 60438, Germany; Structural Genomics Consortium (SGC), Buchmann Institute for Life Sciences, Johann Wolfgang Goethe University, Frankfurt am Main 60438, Germany; German Translational Cancer Network (DKTK) and Frankfurt Cancer Institute (FCI), Frankfurt am Main 60438, Germany;

### Author Contributions

M. R., R. T. and S. K. designed the project; M. R., I. A., L. M. W. and T. H. synthesized the compounds; R. T. and D.-I. B. expressed, purified and co-crystallized MST3; R. T., D.-I. B. and A. C. J. refined the structures of MST3 inhibitor complexes; E.S., T.K. and A. P. performed, analyzed and interpreted the MD simulations and MM/GBSA calculations; A. Krämer, L. E. and M. R. performed DSF assays; L. B. and B.-T. B. performed the NanoBRET assays; A. M. performed the viability assessment and FUCCI cell cycle analysis; A. Kaiser performed microsomal stability assay; S. M. and S. K. supervised the research. The manuscript was written by M. R., A. M., A. C. J. and S. K. with contributions from all coauthors.

### Notes

The authors declare the following competing financial interest(s): L.B. is a co-founder and B.-T.B. is a co-founder and the CEO of CELLinib GmbH (Frankfurt am Main, Germany). The other authors declare that the research was conducted in the absence of any commercial or financial relationships that could be construed as a potential conflict of interest.

### Accession Codes

Coordinates and structure factors of the MST3-inhibitor complexes are available in the Protein Data Bank (PDB) under accession codes 8BZJ (MRLW5), 8QLR (MR24), 8QLS (MR26), and 8QLT (MR30).

Molecular dynamics simulation trajectories and additional data are freely available (DOI: 10.5281/zenodo.7638701). Namely, for each simulated system we attached an additional archive with the raw data on MSTnumber–MRnumber interactions in the *.dat format separated for interaction components (*i.e.* Hydrophobic interactions/Hydrogen Bonding) for concatenated trajectories with a total length of 5 μs each. Furthermore, the *.csv files provide the raw MM/GBSA output for concatenated trajectories of a given system.

## ACKNOWLEDGMENT

The authors are grateful to the Structural Genomics Consortium (SGC), a registered charity (No:1097737) that received funds from Bayer AG, Boehringer Ingelheim, Bristol Myers Squibb, Genentech, Genome Canada through Ontario Genomics Institute, Janssen, Merck KGaA, Pfizer, and Takeda. This project received funding from the Innovative Medicines Initiative 2 Joint Undertaking (JU) under grant agreement No. 875510. The JU receives support from the European Union’s Horizon 2020 research and innovation program, EFPIA, Ontario Institute for Cancer Research, Royal Institution for the Advancement of Learning McGill University, Kungliga Tekniska Hoegskolan, and Diamond Light Source Limited. Disclaimer: This communication reflects the views of the authors, and JU is not liable for any use that may be made of the information contained herein. M. R. is grateful for the support by the Dr. Hilmer-Stiftung. A. M. is supported by the German Research Foundation (DFG) – grant number 259130777 (SFB1177) and B.-T. B. and S.K. are grateful for support by the SFB1399 (grant number: 413326622). R. T. was supported by the German Research Foundation (DFG) grant 397659447. A. C. J. is supported by the German Research Foundation (DFG) grant JO 1473/1-3. The CQ1 microscope was funded by FUGG (INST 161/920-1 FUGG). We want to thank Julia Frischkorn for her passionate cell culture work. Data collection at the Swiss Light Source (SLS) was supported by funding from the European Union’s Horizon 2020 research and innovation program grant agreement number 730872, project CALIPSOplus. We also thank the staff at SLS beamline X06SA for assistance during data collection. T. K., A. P. and E. S. would like to thank the TϋCAD2, which is funded by the Federal Ministry of Education and Research (BMBF) and the Baden-Württemberg Ministry of Science as part of the Excellence Strategy of the German Federal and State Governments EXC 2180–390900677, T. K. is supported by the Fortüne initiative (No. 2613-0-0) and by the iFIT, which are both initiatives from the Excellence Strategy of the German Federal and State Governments. We thank the CSC-Finland for the generous computational resources provided.

## ABBREVIATIONS

AKT: protein kinase B
AMPKα: AMP-activated protein kinase
ATF: cAMP-dependent transcription factor
ATG: autophagy-related gene
BECN1: beclin-1
CGC: chemogenomic compound
CK1: casein kinase 1
DSF: differential scanning fluorimetry
EGFR: epidermal growth factor receptor
FES: feline sarcoma/Fujinami avian sarcoma oncogene homolog
FUCCI: fluorescent ubiquitination-based cell cycle indicator
GCK: germinal center kinase
GFP: green fluorescent protein
HER2: tyrosine kinase-type cell surface receptor
HPLC: high-performance liquid chromatography
IQR: interquartile range
JAK: Janus kinase
LATS: large tumor suppressor homolog
LC3B: microtubule-associated proteins 1A/1B light chain 3B
LRRK2: leucine-rich repeat kinase 2
MAP3K5: mitogen-activated protein kinase kinase kinase 5
MD: molecular dynamics
MM/GBSA: molecular mechanics with generalized Born and surface area solvation
Mob1A/B: Mps one binder 1 A/B
MST: mammalian STE20-like protein kinase
NDR: Dbf2-related kinase
NEK2: NimA-related protein kinase 2
NUAK: SNF1/AMP kinase-related kinase
PAK: p21-activated kinase
RFP: red fluorescent protein
Sav1: Salvador homolog 1
SIK: salt-inducible kinase
STE: homologous kinases of the yeast proteins STE20, STE11, and STE7
STK: serine/threonine kinase
TAZ: transcriptional coactivator with PDZ-domain motif
TEAD: TEA domain family member
ULK1: Unc-51-like kinase 1
YAP: Yes-associated protein
YSK1: Yeast Sps1/Ste20-related kinase 1

