## Supplemental material for "Development of selective pyrido[2,3-*d*]pyrimidin-7(8*H*)-one-based Mammalian STE20-like (MST3/4) kinase inhibitors"

**Table of Content**

Supplementary Figures S1-S10 .......... S3-S15

Table S1: DSF assay panel results of compounds **20** and **22**–**26** .......... S16-S19

Table S2: DSF assay panel results of compounds **27**–**32** .......... S20-S23

Table S3: DSF assay panel results of compounds **33**–**36** .......... S24-S26

Table S4: NanoBRET assay information .......... S26

Table S5: Data collection and refinement statistics .......... S27

Table S6: MR24 (**24**) and MR26 (**26**) interaction patterns derived from Molecular Dynamics simulations .......... S28-S29

Table S7: Percentage of hydrophobic ligand–protein interactions derived from Molecular Dynamics simulations .......... S30

Analytical data of compounds **9**–**36** and intermediates **38**–**70** .......... S31-S168

References .......... S169

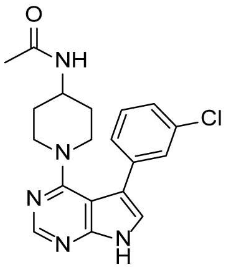

SBP-3264
(**3**)

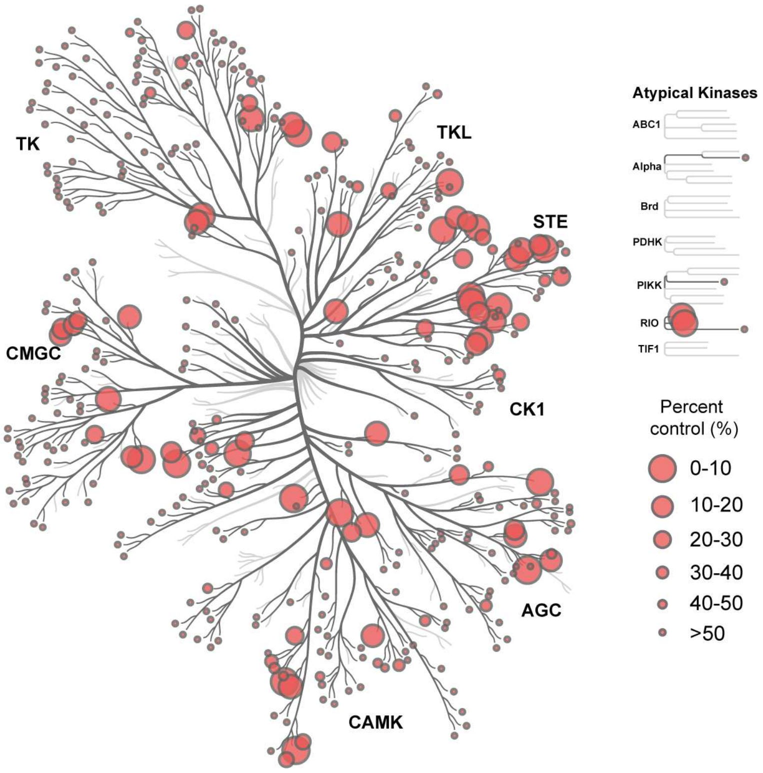


**Figure S1.** Published selectivity profile of SBP-3264 (**3**).[1] **3** was tested using the scanMAX kinase assay panel of 468 kinases (*Eurofins Discovery*) at a concentration of 1 μM. Results are shown as a phylogenetic tree using the web-based tool CORAL.[2] Kinases tested are shown as dark gray branches and kinases inhibited by red circles. The circle radius represents the remaining activity in the presence of the inhibitor (%) compared with the kinase activity in its absence.

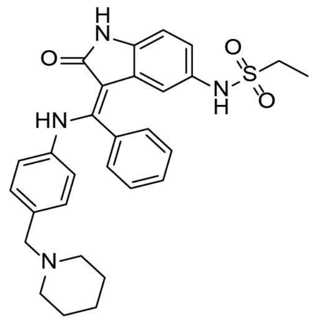

Hesperadin
(**7**)

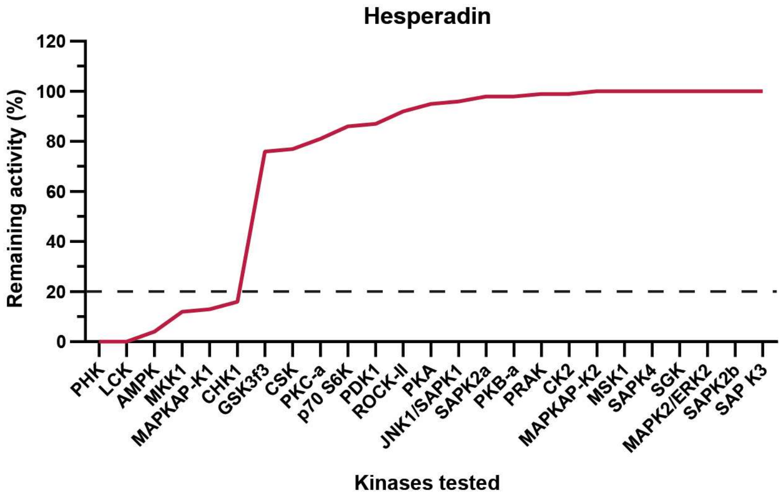


**Figure S2.** Selectivity profile of aurora kinase inhibitor hesperadin (**7**).[3] Hesperadin (**7**) was tested against a panel of 25 kinases, including kinase domains and active mutants, at a concentration of 1 μM. Data are shown as percent of remaining activity in the presence of the inhibitor compared with the activity in its absence.

NanoBRET *in vitro* measurement of **8**

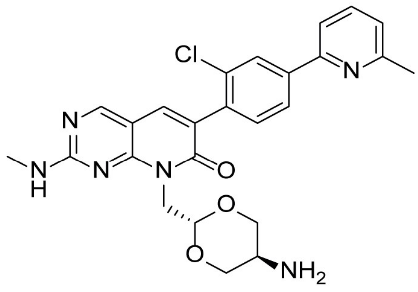


G-5555
(**8**)

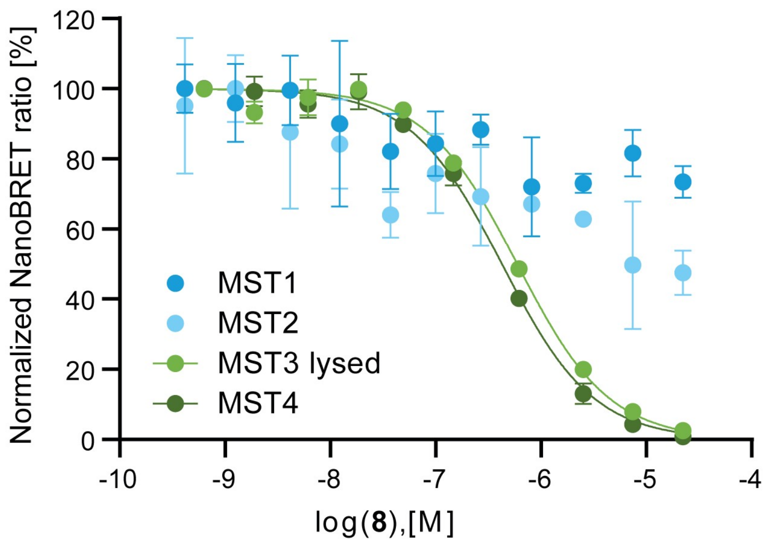


**Figure S3.** NanoBRET curves of G-5555 (**8**) against MST1-4. $EC_{50}$ values with standard error of the mean were determined in a 11-point dose-response curve (duplicate measurements).

In-house selectivity profiles of compounds MRLW5 (**20**), **22**, and **23**

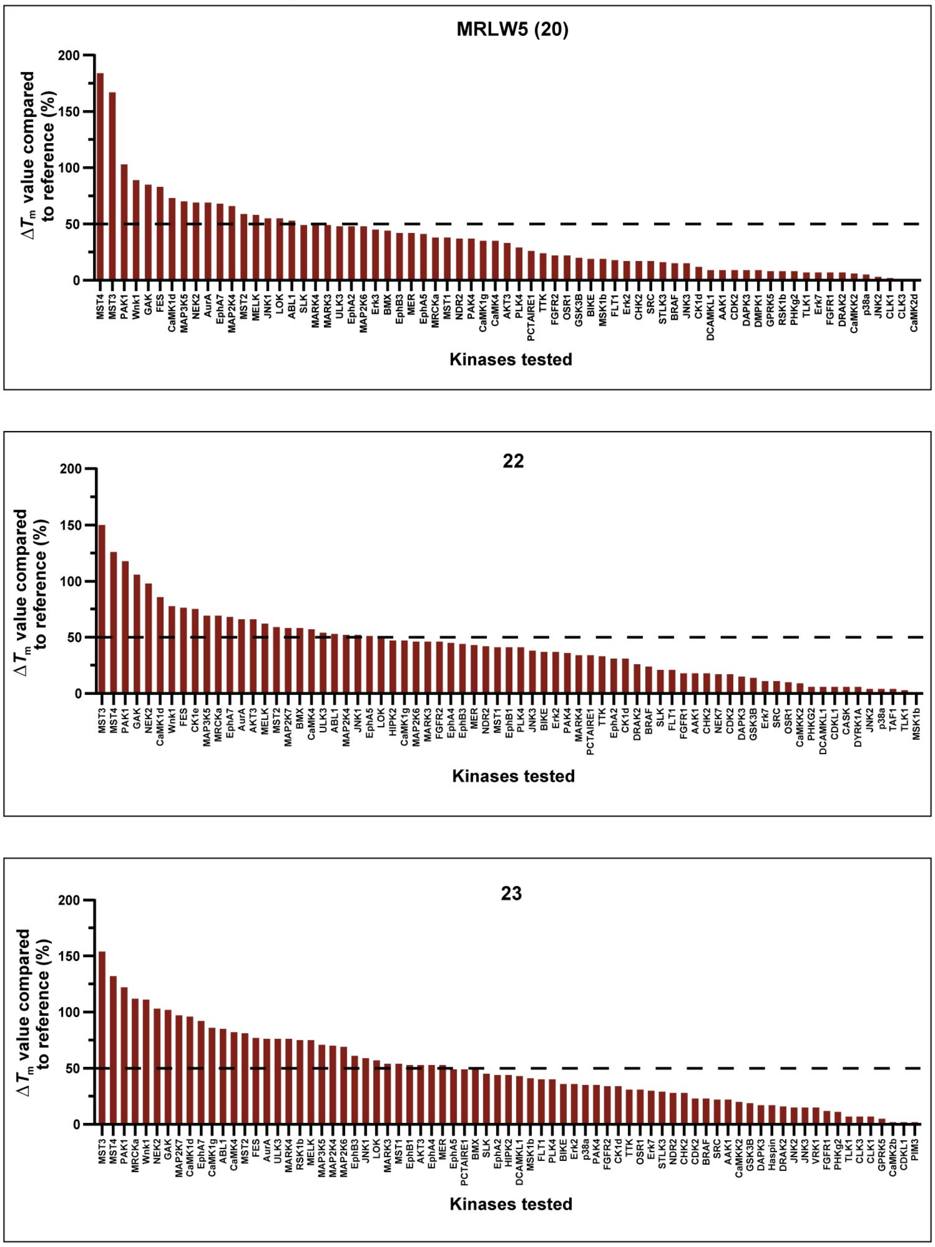


**Figure S4.** Selectivity data of compounds MRLW5 (**20**), **22**, and **23** tested in our in-house DSF panel assay against 90 and 94 kinases, respectively. Data are shown as bar charts. Determined $\Delta T_m$ values were compared to a corresponding reference compound and are shown in percent (%). Only kinases that were stabilized are shown. The raw data can be found in Table S1. Average of duplicate measurements. All kinases higher 50% were considered as off-targets.

In-house selectivity profile of front-pocket modified compounds **24**-**36**

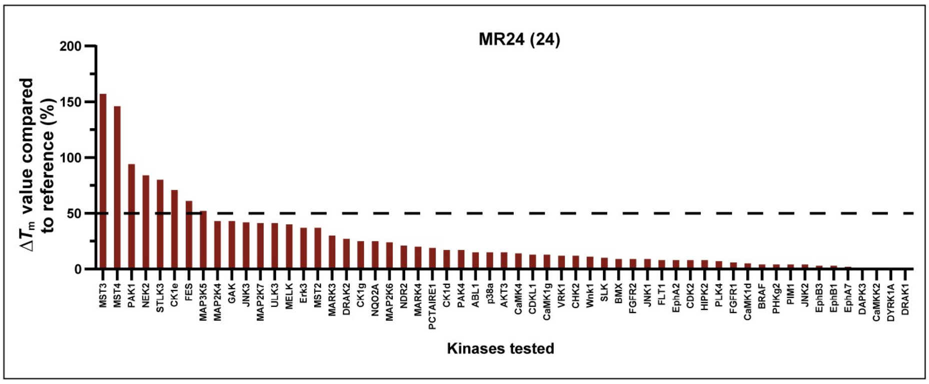


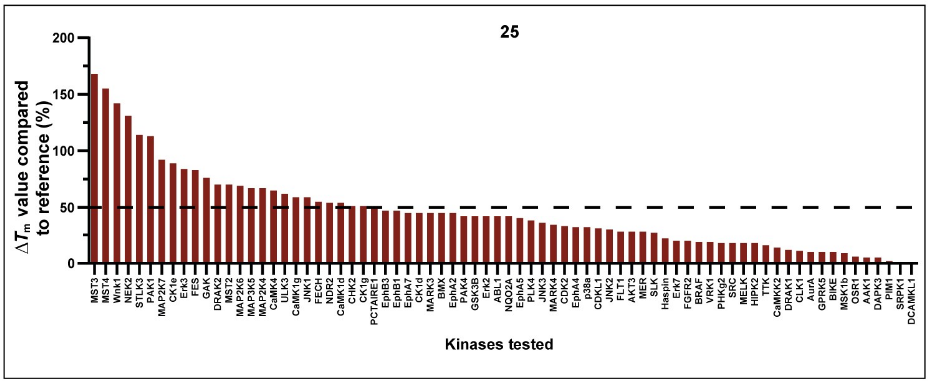


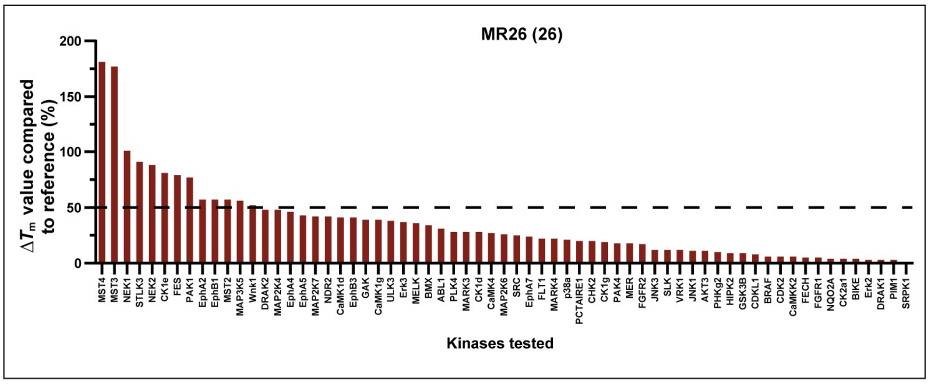


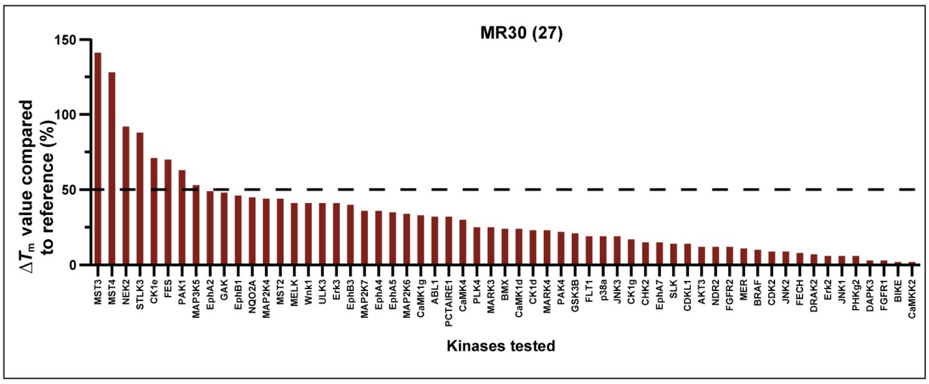

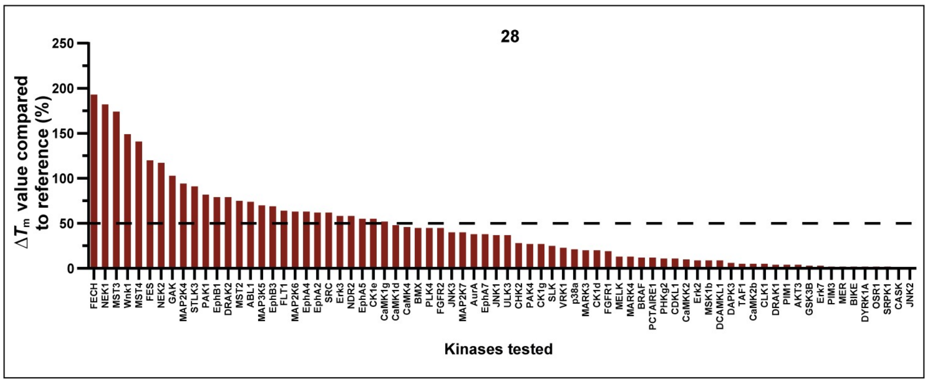
28
ΔTm value compared to reference (%)
250
200
150
100
50
0
Kinases tested

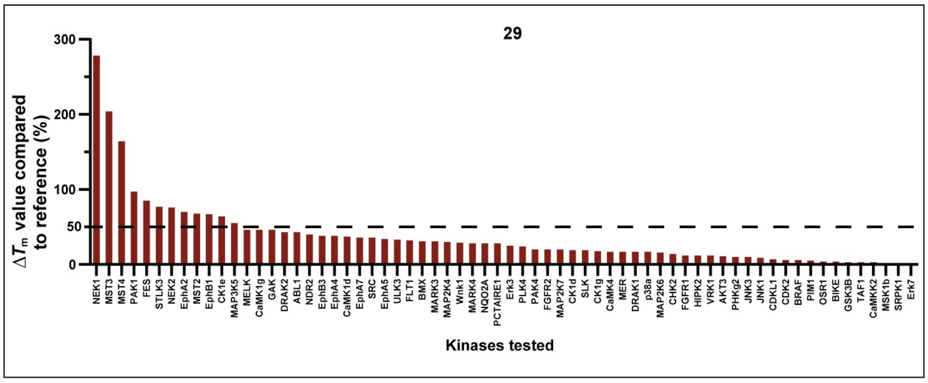
29
ΔTm value compared to reference (%)
300
200
100
50
0
Kinases tested

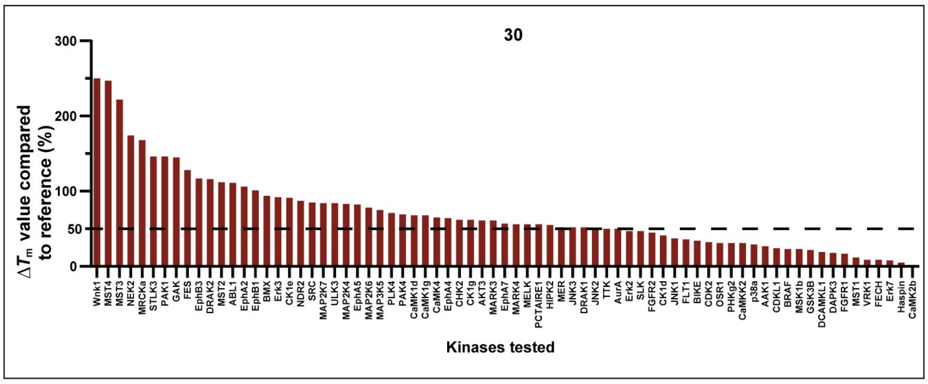
30
ΔTm value compared to reference (%)
300
200
100
50
0
Kinases tested

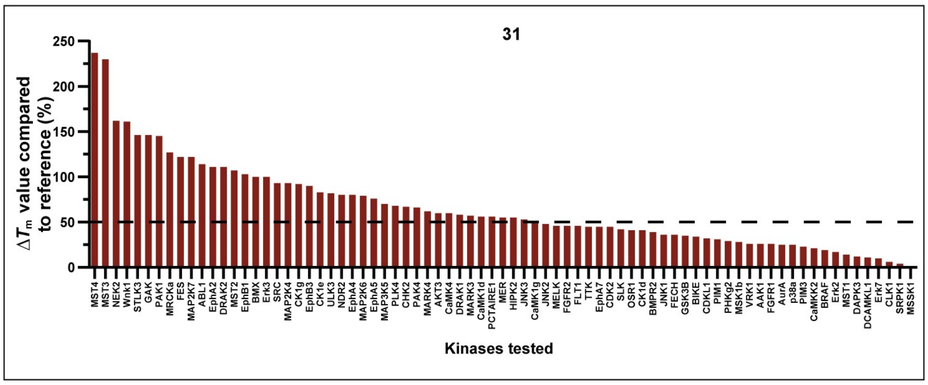
31
ΔTm value compared to reference (%)
250
200
150
100
50
0
Kinases tested

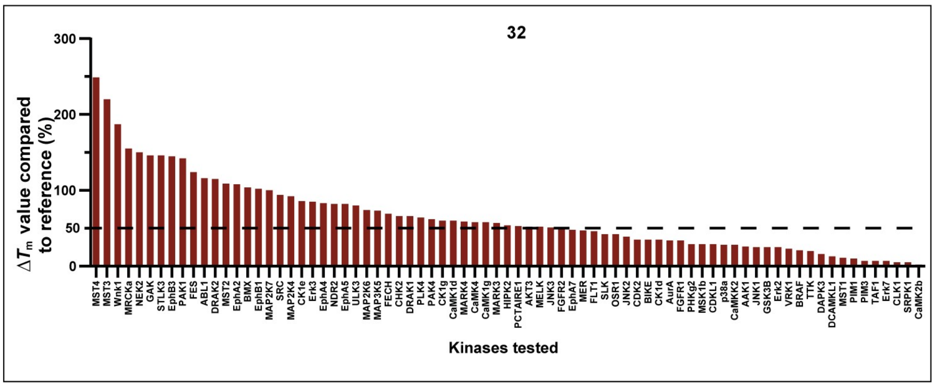
32
$\Delta T_m$ value compared to reference (%)
300
200
100
50
0
MST4 MST3 Wnk1 MRCKa NEK2 GAK STLK3 EphB3 PAK1 FES ABL1 DRAK2 MST2 EphA2 BMX EphB1 MAP2K7 SRC MAP2K4 CK1e Erk3 EphA4 NDR2 EphA5 ULK3 MAP2K6 MAP3K5 FECH CHK2 DRAK1 PLK4 PAK4 CK1g CaMK1d MARK4 CaMK4 CaMK1g MARK3 HIPK2 PCTAIRE1 AKT3 MELK JNK3 FGFR2 EphA7 MER FLT1 SLK OSR1 JNK2 CDK2 BIKE CK1d AurA FGFR1 PHKg2 MSK1b CDKL1 p38a CaMKK2 AAK1 JNK1 GSK3B Erk2 VRK1 BRAF TTK DAPK3 DCAMKL1 MST1 PIM1 PIM3 TAF1 Erk7 CLK1 SRPK1 CaMK2b
Kinases tested

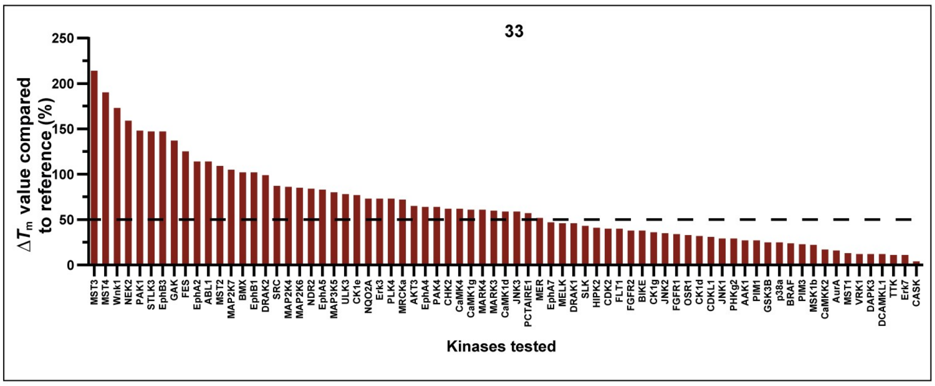
33
$\Delta T_m$ value compared to reference (%)
250
200
150
100
50
0
MST3 MST4 Wnk1 NEK2 PAK1 STLK3 EphB3 GAK FES EphA2 ABL1 MST2 MAP2K7 BMX EphB1 DRAK2 SRC MAP2K4 MAP2K6 NDR2 EphA5 MAP3K5 ULK3 CK1e NQO2A Erk3 PLK4 MRCKa AKT3 EphA4 PAK4 CHK2 CaMK4 CaMK1g MARK4 MARK3 CaMK1d JNK3 PCTAIRE1 MER EphA7 MELK DRAK1 SLK HIPK2 CDK2 FLT1 FGFR2 BIKE CK1g JNK2 FGFR1 OSR1 CK1d CDKL1 JNK1 PHKg2 AAK1 PIM1 GSK3B p38a BRAF PIM3 MSK1b CaMKK2 AurA MST1 VRK1 DAPK3 DCAMKL1 TTK Erk7 CASK
Kinases tested

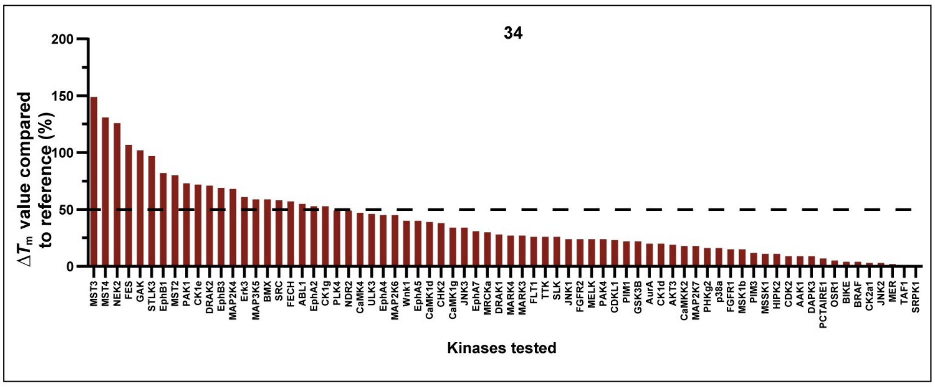
34
$\Delta T_m$ value compared to reference (%)
200
150
100
50
0
MST3 MST4 NEK2 FES GAK STLK3 EphB1 MST2 PAK1 CK1e DRAK2 EphB3 MAP2K4 Erk3 MAP3K5 BMX SRC FECH ABL1 EphA2 CK1g PLK4 NDR2 CaMK4 ULK3 EphA4 MAP2K6 Wnk1 EphA5 CaMK1d CHK2 CaMK1g JNK3 EphA7 MRCKa DRAK1 MARK4 MARK3 FLT1 TTK SLK JNK1 FGFR2 MELK PAK4 CDKL1 PIM1 GSK3B AurA CK1d AKT3 CaMKK2 MAP2K7 PHKg2 p38a FGFR1 MSK1b PIM3 MSSK1 HIPK2 CDK2 AAK1 DAPK3 PCTAIRE1 OSR1 BIKE BRAF CK2a1 JNK2 MER TAF1 SRPK1
Kinases tested

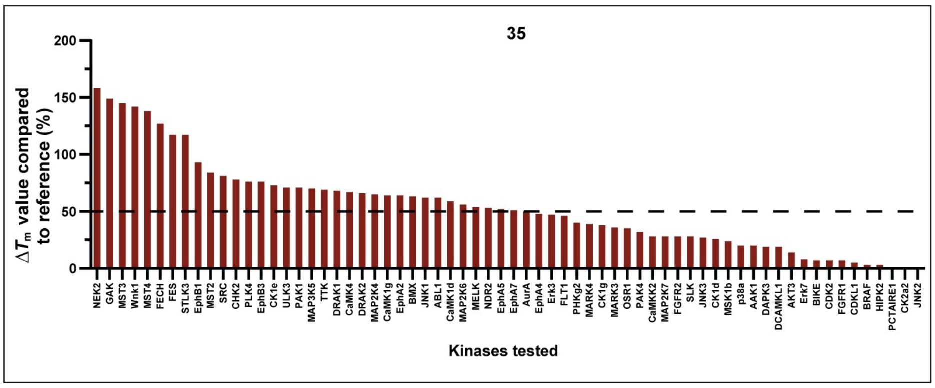
35
$\Delta T_m$ value compared to reference (%)
200
150
100
50
0
NEK2 GAK MST3 Wnk1 MST4 FECH FES STLK3 EphB1 MST2 SRC CHK2 PLK4 EphB3 CK1e ULK3 PAK1 MAP3K5 TTK DRAK1 CaMK4 DRAK2 MAP2K4 CaMK1g EphA2 BMX JNK1 ABL1 CaMK1d MAP2K6 MELK NDR2 EphA5 EphA7 AurA EphA4 Erk3 FLT1 PHKg2 MARK4 CK1g MARK3 OSR1 PAK4 CaMKK2 MAP2K7 FGFR2 SLK JNK3 CK1d MSK1b p38a AAK1 DAPK3 DCAMKL1 AKT3 Erk7 BIKE CDK2 FGFR1 CDKL1 BRAF HIPK2 PCTAIRE1 CK2a2 JNK2
Kinases tested

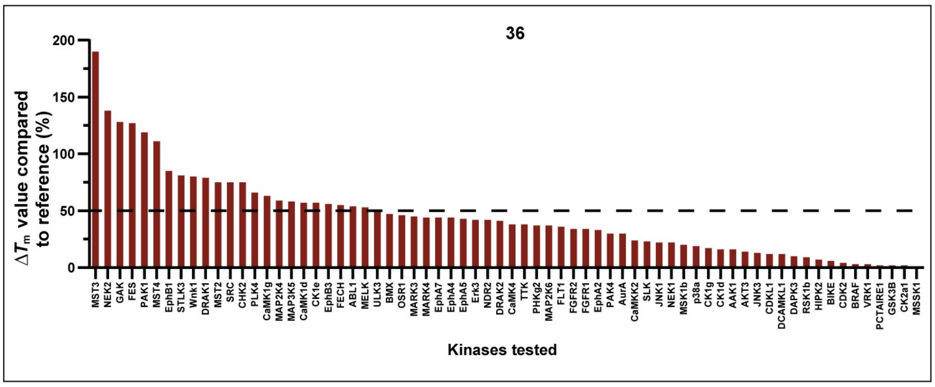


**Figure S5.** Bar charts of the selectivity data of compounds **24–36** tested in our in-house DSF panel assay on 100 kinases. Determined $\Delta T_m$ values were compared to a corresponding reference compound and are shown in percent (%). Only kinases that were stabilized are shown. The raw data can be found in the Tables S1, S2, and S3. Average of duplicate measurements. All kinases higher 50% were considered as off-targets.

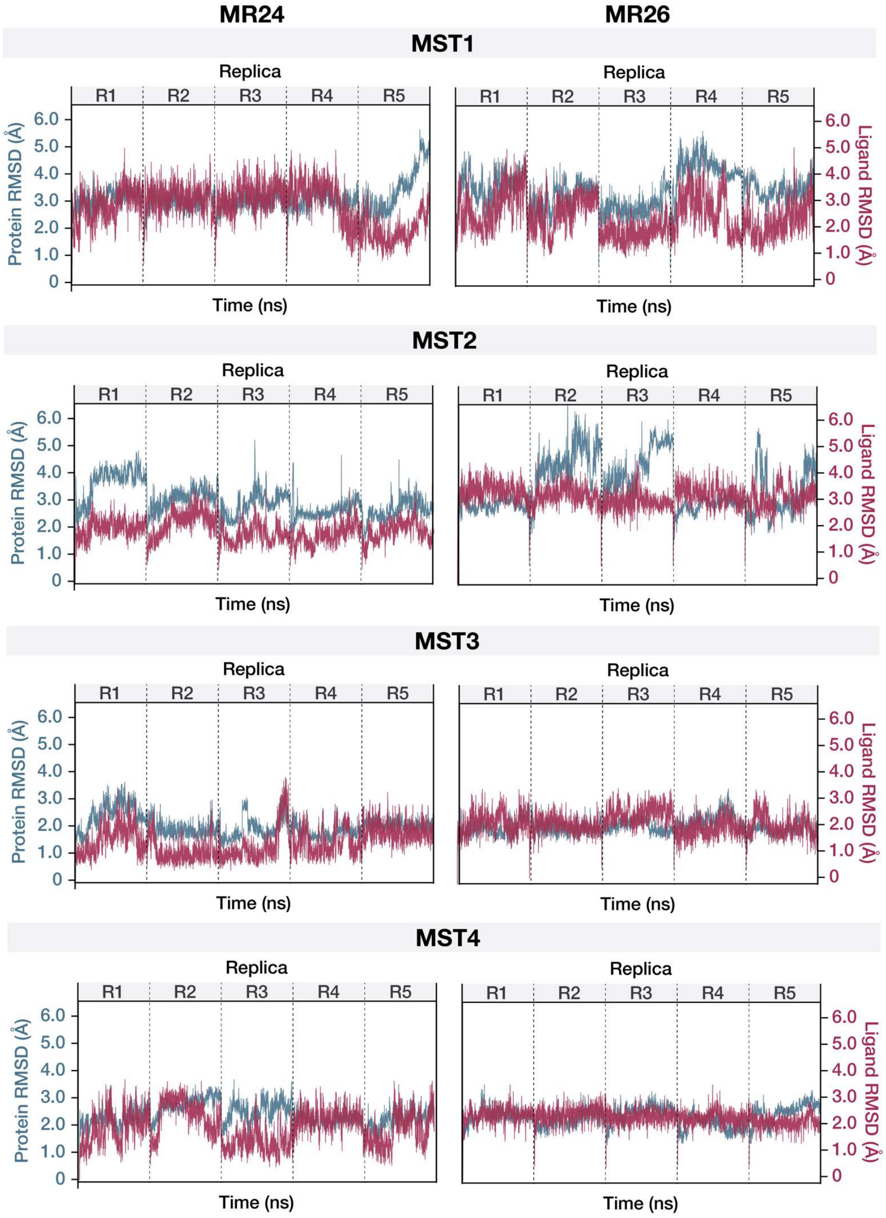


**Figure S6.** RMSD of MR24 (**24**)- and MR26 (**26**)-protein complexes. Protein RMSDs were calculated for $C_a$–atoms, shown in blue. Ligand RMSDs were calculated as aligned on protein (the RMSD measures the fluctuations of the ligand with respect to the protein), shown in red. The RMSD values are divided by replicas: R1–replica 1, R2–replica 2, etc. Each replica comprises 1000ns (1μs) simulation time.

A

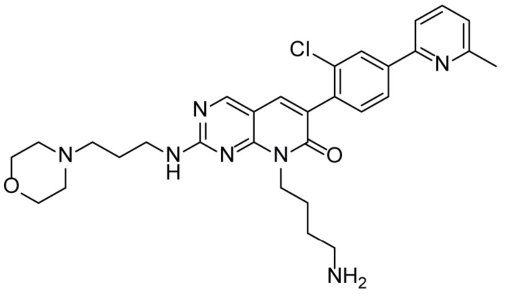

MR26
(**26**)

| Targets | $\Delta T_m$-value compared to reference (%)[a] | $EC_{50}$(nM)[b] NanoBRET |
|---|---|---|
| MST4 | 181 | 115 ± 57 |
| MST3 | 177 | 55 ± 43 |
| NEK1 | 101 | n.d. |
| STLK3 | 91 | n.d. |
| NEK2 | 88 | 6400 ± 100 |
| CK1e | 81 | 4000 ± 1100 |
| FES | 79 | 8600 ± 200 |
| PAK1 | 77 | n.d. |
| EpHA2 | 57 | n.d. |
| EpHB1 | 57 | n.d. |
| MST2 | 57 | 10200 ± 500 |
| MAP3K5 | 56 | n.d. |
| Wnk1 | 52 | n.d. |
| MST1 | 3.6 | 5300 ± 200 |
| SIK2 | n.d. | 512 ± 50 |

B

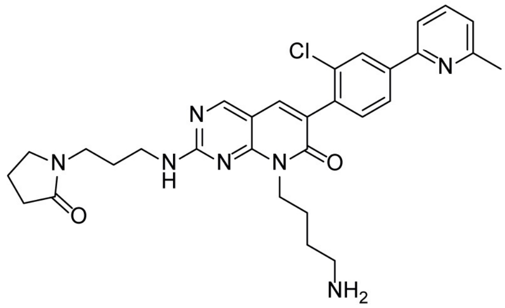

MR30
(**27**)

| Targets | $\Delta T_m$-value compared to reference (%)[a] | $EC_{50}$(nM)[b] NanoBRET |
|---|---|---|
| MST3 | 141 | 178 ± 141 |
| MST4 | 128 | 1100 ± 200 |
| NEK2 | 92 | 5600 ± 100 |
| STLK3 | 88 | n.d. |
| CK1e | 71 | 9900 ± 1200 |
| FES | 70 | 7800 ± 700 |
| PAK1 | 63 | n.d. |
| MAP3K5 | 53 | n.d. |
| MST2 | 44 | 16900 ± 1300 |
| MST1 | 1.1 | 15100 ± 600 |
| SIK2 | n.d. | 1300 ± 100 |

**Figure S7.** Selectivity data of compounds MR26 (**26**) and MR30 (**27**). Both compounds were tested in a DSF panel assay on 100 kinases. Determined $\Delta T_m$ values were compared with a corresponding reference compound and are shown in percent (%). Kinases with values higher than 50% are shown together with MST1, MST2, and SIK2. *In cellulo* $EC_{50}$ values of the corresponding kinases were measured in the NanoBRET assay. [a]Average of duplicate measurements. Data included in this figure can be found in Tables S1 and S2. [b]$EC_{50}$ values with standard error of the mean were determined in a 11-point dose-response curve (duplicate measurements).

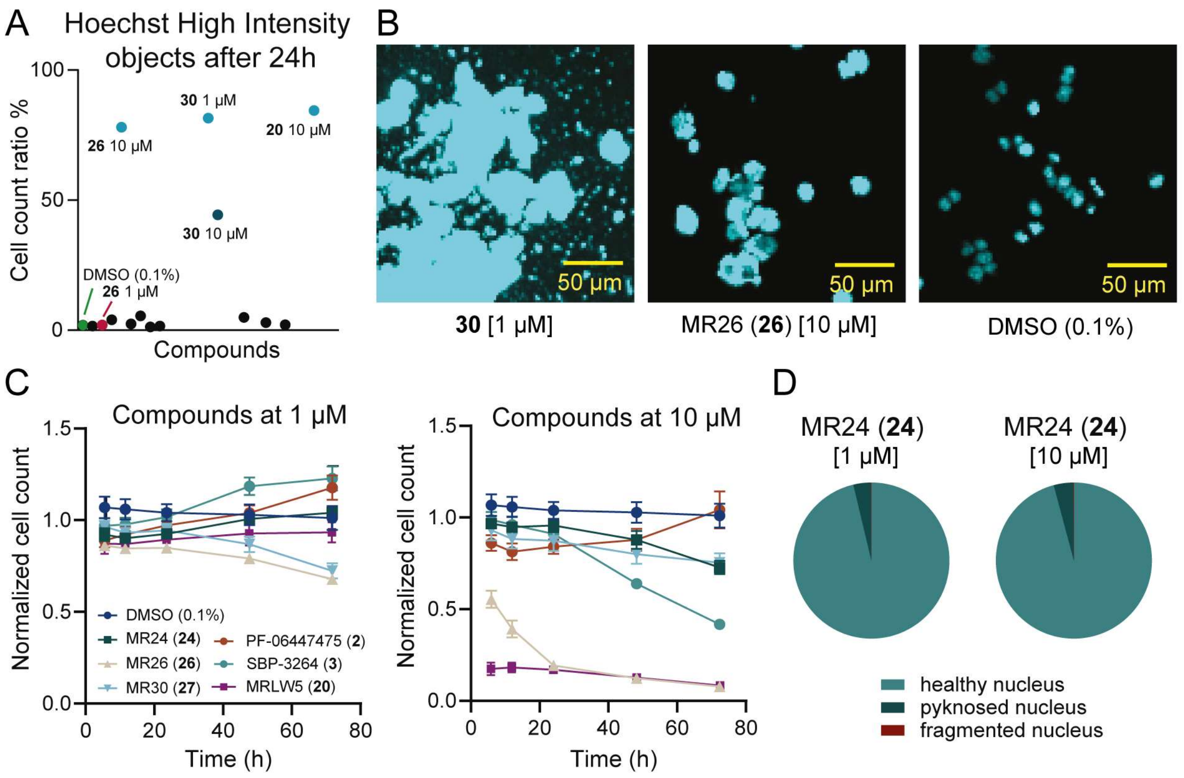


**Figure S8:** Viability assessment. (A) Ratio of cells showing Hoechst High Intensity Objects per well after 24 h of compound exposure (**2**, **3**, **20**, **24**, **26,** and **27**) at 1 µM and 10 µM. Data of two biological replicates are shown. Light blue color indicates Hoechst High Intensity Objects > 50%. (B) Fluorescent confocal image of stained (blue: DNA/nuclei) HCT116 cells after 48 h of compounds exposure (**30** [10 µM], **26** [1 µM]) compared with cells exposed to DMSO (0.1%). (C) Normalized cell count of HCT116 cells after 6 h, 12 h, 24 h, 48 h, and 72 h of compound exposure (**2**, **3**, **20**, **24**, **26**, and **27**) at 1 µM and 10 µM compared with cells exposed to DMSO (0.1%). Error bars show SEM of two biological replicates. (D) Fraction of healthy, fragmented, and pyknosed nuclei after 48 h of compound MR24 (**24**) exposure at 1 µM and 10 µM in HCT116 cells. Average data of two biological duplicates are shown.

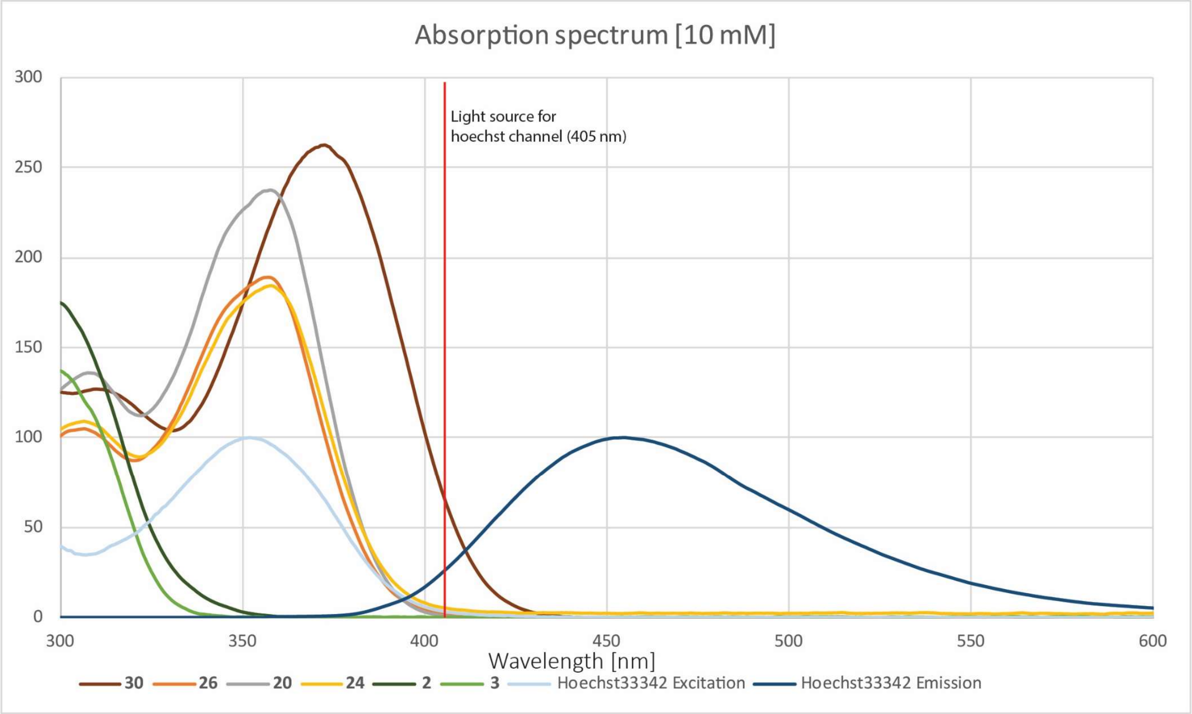


**Figure S9.** Absorption spectra of compounds **2**, **3**, **20**, **24**, **26**, and **30**. Absorbance was measured at a concentration of 10 mM using a NanoDrop UV-Vis Spectrophotometer.

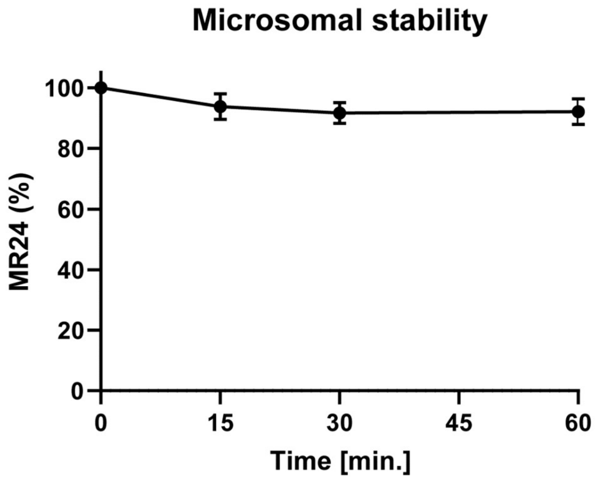


| Descriptor | Interpretation | Limit Values | G-5555[a] | MRLW5[+] | MR24[+] | MR30[+] |
|---|---|---|---|---|---|---|
| $P_{Caco-2}$ | Predicted apparent Caco-2 cell permeability in nm/sec[b]. Caco-2 cells are a model for the gut-blood barrier | <25 poor, >500 great | 140.1 | 96.5 | 22.1 | 23.9 |
| logB/B | Predicted brain/blood partition coefficient[c] | - 3.0 – 1.2[b] | -0.8 | -1.1 | -2.2 | -1.9 |
| $P_{MDCK}$ | Predicted apparent MDCK cell permeability in nm/sec[b]. MDCK cells are considered to be a good mimic for the blood-brain barrier | <25 poor, >500 great | 125.4 | 88.8 | 16.5 | 41.3 |
| $logK_p$ | Predicted skin permeability | - 8.0 – -1.0 | -4.4 | -4.5 | -5.3 | -4.7 |
| $logK_{hsa}$ | Prediction of binding to human serum albumin | -1.5 – 1.5 | 0.1 | 0.5 | 0.0 | 0.4 |
| Oral Abs. | Predicted human oral absorption on 0 to 100% scale. The prediction is based on a quantitative multiple linear regression model | >80% is high <25% is poor | 81 | 84 | 52 | 61 |

**Figure S10.** Pharmacokinetic properties of MR24 (**24**). Microsomal stability was determined by incubating MR24 (**24**) with activated liver microsomes over a time of 1 h and monitoring its stability via HPLC. Data is shown as ± SEM of three experiments. ADME prediction of compounds MR24 (**24**) and MR30 (**27**) were performed using the software QikProp (*Schrödinger, LLC, New York, NY, 2021*). [a]Predicted literature values for G-5555 (**8**).[4] [b]Predictions are for non-active transport. [c]Predictions are for orally delivered drugs so, for example, dopamine and serotonin are CNS negative because they are too polar to cross the blood-brain barrier.

**Table S1**. Results of the DSF panel assay of compounds **20** and **22–26** on 100 kinases. Kinases are shown with their common abbreviation. For each kinase, a reference compound is shown with its corresponding $\Delta T_m$ value. Average of duplicate measurements.

| | | | **$\Delta T_m$ [°C]** | | | | | |
|---|---|---|---|---|---|---|---|---|
| **Kinases** | **Reference compound** | **$\Delta T_m$ shift reference compound** | **MRLW5 (20)** | **22** | **23** | **MR24 (24)** | **25** | **MR26 (26)** |
| **AAK1** | Staurosporine | 15.3 | 1.4 | 2.8 | 3.3 | -0.4 | 0.8 | -0.2 |
| **ABL1** | Staurosporine | 9.3 | 4.9 | 4.9 | 7.9 | 1.4 | 3.9 | 2.9 |
| **AKT3** | Staurosporine | 7.0 | 2.3 | 4.6 | 3.7 | 1.0 | 2.0 | 0.8 |
| **AurA** | Staurosporine | 17.2 | 11.8 | 11.4 | 13.1 | -1.7 | 1.8 | -0.9 |
| **BIKE** | Staurosporine | 19.0 | 3.7 | 7.1 | 6.9 | -0.4 | 1.8 | 0.7 |
| **BMPR2** | Lestaurtinib | 2.6 | -0.1 | -0.3 | -0.7 | -0.1 | -0.2 | -0.2 |
| **BMX** | Staurosporine | 7.0 | 3.1 | 4.1 | 3.4 | 0.6 | 3.1 | 2.4 |
| **BRAF** | Dabrafenib | 26.6 | 4.1 | 6.3 | 6.2 | 1.2 | 5.2 | 1.7 |
| **BRD4** | JQ1 | 7.0 | -0.7 | 0.0 | -1.1 | -1.7 | -2.9 | -1.2 |
| **BRPF1** | GSK6853 | 14.0 | n.d. | -0.4 | -1.8 | -1.5 | -1.4 | -1.0 |
| **CaMK1d** | Staurosporine | 9.9 | 7.2 | 8.5 | 9.5 | 0.5 | 5.3 | 4.1 |
| **CaMK1g** | Staurosporine | 11.0 | 3.9 | 5.1 | 9.5 | 1.4 | 6.5 | 4.3 |
| **CaMK2b** | Staurosporine | 13.2 | -1.3 | -0.9 | 0.2 | -0.8 | -0.1 | -0.3 |
| **CaMK2d** | Staurosporine | 16.5 | 0.1 | 0.0 | -0.2 | -0.2 | -0.2 | -0.1 |
| **CaMK4** | Staurosporine | 8.4 | 2.9 | 4.8 | 6.9 | 1.2 | 5.4 | 2.2 |
| **CaMKK2** | Staurosporine | 24.5 | 1.4 | 2.3 | 4.9 | 0.2 | 3.3 | 1.5 |
| **CASK** | Staurosporine | 5.0 | -0.5 | 0.3 | -0.4 | -0.4 | -0.4 | -0.3 |
| **CDK2** | Staurosporine | 15.0 | 1.3 | 2.5 | 3.5 | 1.2 | 4.9 | 0.9 |
| **CDKL1** | CEP-32496 | 6.6 | -0.1 | 0.4 | 0.1 | 0.9 | 2.1 | 0.5 |
| **CHK2** | Staurosporine | 16.7 | 2.8 | 3.0 | 4.6 | 2.0 | 8.5 | 3.3 |
| **CK1d** | SR-3029 | 6.8 | 0.8 | 2.1 | 2.3 | 1.2 | 3.1 | 1.9 |
| **CK1e** | PF-670462 | 9.0 | n.d. | 6.8 | n.d. | 6.4 | 8.0 | 7.3 |
| **CK1g** | Staurosporine | 2.0 | n.d. | n.d. | n.d. | 0.5 | 1.0 | 0.4 |
| **CK2a1** | Similtasertib | 14.6 | -0.2 | -0.6 | -0.4 | 0.0 | -0.4 | 0.6 |
| **CK2a2** | Similtasertib | 15.7 | -0.6 | -1.4 | -0.9 | -2.0 | -1.9 | -1.0 |
| **CLK1** | Staurosporine | 12.3 | 0.3 | -1.6 | 0.8 | -0.4 | 1.4 | -0.1 |
| **CLK3** | T3 | 15.0 | 0.2 | -0.4 | 1.0 | -0.2 | -0.3 | -0.4 |

| | | | ΔTm [°C] | | | | | |
|---|---|---|---|---|---|---|---|---|
| **Kinases** | **Reference compound** | **$\Delta T_m$ shift reference compound** | **MRLW5 (20)** | **22** | **23** | **MR24 (24)** | **25** | **MR26 (26)** |
| **DAPK1** | Staurosporine | 9.4 | -0.8 | -0.8 | -0.4 | -0.8 | -1.0 | -0.7 |
| **DAPK3** | Staurosporine | 16.2 | 1.4 | 2.4 | 2.8 | 0.2 | 0.8 | 0.0 |
| **DCAMKL1** | Staurosporine | 10.7 | 1.0 | 0.7 | 4.6 | -2.0 | 0.1 | -2.0 |
| **DMPK1** | Staurosporine | 9.4 | 0.8 | -1.7 | -2.8 | -1.3 | -2.7 | -1.3 |
| **DRAK1** | Staurosporine | 7.7 | -0.5 | -1.3 | -1.0 | 0.0 | 0.9 | 0.2 |
| **DRAK2** | Staurosporine | 10.6 | 0.7 | 2.7 | 1.7 | 2.9 | 7.5 | 5.1 |
| **DYRK1A** | Staurosporine | 9.6 | -1.0 | 0.5 | -0.3 | 0.1 | -0.1 | -0.3 |
| **DYRK2** | Staurosporine | 7.3 | -0.1 | -0.4 | -0.2 | -0.3 | -0.6 | -0.2 |
| **EphA2** | Staurosporine | 7.3 | 3.5 | 2.3 | 3.2 | 0.6 | 3.3 | 4.2 |
| **EphA4** | Staurosporine | 5.5 | n.d. | 2.5 | 2.9 | -0.3 | 1.8 | 2.5 |
| **EphA5** | Staurosporine | 8.1 | 3.3 | 4.1 | 4.0 | -0.4 | 3.2 | 3.5 |
| **EphA7** | Staurosporine | 11.3 | 7.7 | 7.7 | 10.4 | 0.3 | 5.1 | 2.7 |
| **EphB1** | Staurosporine | 6.4 | n.d. | 2.6 | 3.4 | 0.2 | 3.0 | 3.7 |
| **EphB3** | Staurosporine | 5.9 | 2.5 | 2.6 | 3.6 | 0.2 | 2.8 | 2.4 |
| **Erk2** | GDC-0994 | 7.5 | 1.3 | 2.8 | 2.7 | -0.9 | 3.1 | 0.2 |
| **Erk3** | Staurosporine | 1.2 | 0.5 | n.d. | n.d. | 0.4 | 1.0 | 0.4 |
| **Erk7** | Staurosporine | 14.2 | 1.0 | 1.5 | 4.2 | -0.4 | 2.9 | -0.4 |
| **FECH** | Vemurafenib | 5.5 | n.d. | -0.3 | n.d. | -0.5 | 3.0 | 0.3 |
| **FES** | Staurosporine | 7.1 | 5.9 | 5.4 | 5.5 | 4.3 | 5.9 | 5.6 |
| **FGFR1** | Staurosporine | 6.0 | 0.4 | 1.1 | 0.7 | 0.4 | -0.4 | 0.3 |
| **FGFR2** | Staurosporine | 8.5 | 1.9 | 3.9 | 2.9 | 0.8 | 1.7 | 1.4 |
| **FLT1** | Staurosporine | 12.2 | 2.2 | 2.6 | 4.9 | 1.0 | 3.5 | 2.7 |
| **GAK** | Staurosporine | 9.1 | 7.7 | 9.7 | 9.3 | 3.9 | 6.9 | 3.6 |
| **GPRK5** | Staurosporine | 5.9 | 0.5 | -1.4 | 0.3 | -1.0 | 0.6 | -0.5 |
| **GSK3B** | Staurosporine | 9.0 | 1.8 | 1.3 | 1.7 | 0.0 | 3.8 | 0.8 |
| **Haspin** | Staurosporine | 8.9 | -0.4 | -1.2 | 1.5 | -1.4 | 1.9 | -1.5 |
| **HIPK2** | Staurosporine | 4.8 | n.d. | 2.3 | 2.1 | 0.4 | 0.8 | 0.4 |
| **JNK1** | Staurosporine | 7.6 | 4.2 | 4.0 | 4.5 | 0.7 | 4.5 | 0.8 |
| **JNK2** | SBI-0069279 | 8.6 | 0.3 | 0.3 | 1.3 | 0.4 | 2.6 | 0.0 |
| **JNK3** | CEP-32496 | 8.8 | 1.3 | 3.3 | 1.3 | 3.7 | 3.2 | 1.1 |
| **LOK** | Staurosporine | 22.8 | 12.5 | 11.4 | 13.0 | -0.6 | -0.7 | -0.4 |

| | | | $\Delta T_m$ [°C] | | | | | |
|---|---|---|---|---|---|---|---|---|
| **Kinases** | **Reference compound** | **$\Delta T_m$ shift reference compound** | **MRLW5 (20)** | **22** | **23** | **MR24 (24)** | **25** | **MR26 (26)** |
| **MAP2K4** | Staurosporine | 11.7 | 7.7 | 6.1 | 8.2 | 5.0 | 7.9 | 5.6 |
| **MAP2K6** | Staurosporine | 12.0 | 5.7 | 5.6 | 8.3 | 2.8 | 8.3 | 3.1 |
| **MAP2K7** | Staurosporine | 7.3 | n.d. | 4.3 | 7.1 | 3.0 | 6.7 | 3.1 |
| **MAP3K5** | Staurosporine | 17.5 | 12.3 | 12.0 | 12.4 | 9.0 | 11.8 | 9.8 |
| **MAPKAPK2** | Staurosporine | 3.4 | 0.0 | -0.1 | -0.9 | -0.1 | -0.6 | -0.2 |
| **MARK3** | Staurosporine | 19.1 | 9.3 | 8.8 | 10.4 | 5.8 | 8.6 | 5.4 |
| **MARK4** | Staurosporine | 16.8 | 8.2 | 5.8 | 12.7 | 3.3 | 5.8 | 3.7 |
| **MELK** | Staurosporine | 13.9 | 8.0 | 8.5 | 10.4 | 5.5 | 2.4 | 5.0 |
| **MER** | Staurosporine | 5.9 | 2.5 | 2.6 | 3.1 | -0.7 | 1.6 | 1.0 |
| **MRCKa** | Staurosporine | 2.6 | 1.0 | 1.8 | 2.9 | -0.7 | 0.0 | -0.2 |
| **MSK1b** | Staurosporine | 15.6 | 2.9 | 0.1 | 6.4 | -1.0 | 1.4 | -0.4 |
| **MSSK1** | Staurosporine | 3.7 | n.d. | -0.1 | n.d. | -0.2 | -0.4 | -0.3 |
| **MST1** | Staurosporine | 14.7 | 5.6 | 6.0 | 7.9 | -0.6 | -1.4 | -0.5 |
| **MST2** | Staurosporine | 13.4 | 7.9 | 8.0 | 10.8 | 4.9 | 9.4 | 7.7 |
| **MST3** | Staurosporine | 7.0 | 11.7 | 10.5 | 10.8 | 11.0 | 11.8 | 12.4 |
| **MST4** | Staurosporine | 6.2 | 11.4 | 7.8 | 8.2 | 9.1 | 9.6 | 11.2 |
| **MYT1** | Dasatinib | 4.5 | -1.0 | -4.0 | -0.9 | -2.0 | -2.3 | -1.1 |
| **NDR2** | Staurosporine | 12.3 | 4.6 | 5.1 | 3.5 | 2.6 | 6.7 | 5.2 |
| **NEK1** | Staurosporine | 0.1 | -1.9 | -2.0 | -2.7 | 0.0 | -1.2 | 0.1 |
| **NEK2** | Staurosporine | 6.7 | 4.6 | 6.6 | 6.9 | 5.6 | 8.8 | 5.9 |
| **NEK7** | Staurosporine | 1.3 | -0.1 | 0.2 | -0.3 | -0.9 | -0.5 | -0.7 |
| **NQO2A** | Staurosporine | 1.2 | n.d. | -0.1 | n.d. | 0.3 | 0.5 | 0.1 |
| **OSR1** | Staurosporine | 7.4 | 1.6 | 0.7 | 2.3 | -0.5 | 0.4 | -0.1 |
| **p38a** | Dorama-pimod | 18.9 | 1.0 | 0.7 | 6.7 | 2.8 | 6.1 | 4.0 |
| **PAK1** | Staurosporine | 7.3 | 7.5 | 8.6 | 8.9 | 6.9 | 8.3 | 5.7 |
| **PAK4** | Staurosporine | 12.9 | 4.8 | 4.7 | 4.5 | 2.2 | 5.5 | 2.4 |
| **PCTAIRE1** | Staurosporine | 9.0 | 2.3 | 3.0 | 4.4 | 1.7 | 4.5 | 1.8 |
| **PHKG2** | Staurosporine | 21.8 | 1.8 | 1.4 | 2.4 | 1.0 | 4.0 | 2.1 |
| **PIM1** | Staurosporine | 12.2 | -0.1 | -0.4 | -0.7 | 0.5 | 0.3 | 0.3 |
| **PIM3** | Staurosporine | 19.8 | -1.2 | -1.2 | 0.3 | -1.9 | -0.8 | -0.1 |
| **PLK4** | Staurosporine | 17.9 | 5.2 | 7.3 | 7.1 | 1.3 | 6.7 | 5.1 |

| | | | $\Delta T_m$ [°C] | | | | | |
|---|---|---|---|---|---|---|---|---|
| **Kinases** | **Reference compound** | **$\Delta T_m$ shift reference compound** | **MRLW5 (20)** | **22** | **23** | **MR24 (24)** | **25** | **MR26 (26)** |
| **RSK1b** | Staurosporine | 3.6 | 0.3 | -0.4 | 2.7 | -0.4 | -1.4 | -0.6 |
| **SLK** | Staurosporine | 18.4 | 9.1 | 3.9 | 8.3 | 1.9 | 5.0 | 2.3 |
| **SRC** | Staurosporine | 5.4 | 0.9 | 0.6 | 1.2 | 0.0 | 1.0 | 1.4 |
| **SRPK1** | Staurosporine | 6.7 | -0.2 | -0.2 | -0.5 | -0.1 | 0.1 | 0.1 |
| **STLK3** | Staurosporine | 11.4 | 1.8 | -3.0 | 3.3 | 9.1 | 13.0 | 10.4 |
| **TAF1** | Bromosporine | 7.4 | 0.0 | 0.3 | -0.5 | -0.1 | -0.1 | 0.0 |
| **TLK1** | Staurosporine | 8.5 | 0.6 | 0.2 | 0.6 | -1.8 | -0.9 | -1.6 |
| **TTK** | Staurosporine | 11.2 | 2.7 | 3.7 | 3.5 | -3.0 | 1.8 | -0.2 |
| **ULK3** | Staurosporine | 17.8 | 8.6 | 9.7 | 13.5 | 7.2 | 11.1 | 6.7 |
| **VRK1** | Staurosporine | 3.4 | 0.0 | -0.7 | 0.5 | 0.4 | 0.6 | 0.4 |
| **Wnk1** | Staurosporine | 1.9 | 1.7 | 1.5 | 2.1 | 0.2 | 2.7 | 1.0 |

**Table S2**. Results of the DSF panel assay of compounds **27–32** on 100 Kinases. Kinases are shown with their common abbreviation. For each kinase a reference compound is shown with its corresponding $\Delta T_m$ shift. Two measurements were performed, and the average is shown.

| | | | $\Delta T_m$ [°C] | | | | | |
|---|---|---|---|---|---|---|---|---|
| **Kinases** | **Reference compound** | **$\Delta T_m$ shift reference compound** | **MR30 (27)** | **28** | **29** | **30** | **31** | **32** |
| **AAK1** | Staurosporine | 15.3 | -0.5 | -0.1 | 0.0 | 4.1 | 4.0 | 4.0 |
| **ABL1** | Staurosporine | 9.3 | 3.0 | 6.9 | 4.0 | 10.3 | 10.6 | 10.8 |
| **AKT3** | Staurosporine | 7.0 | 0.9 | 0.3 | 0.8 | 4.3 | 4.2 | 3.6 |
| **AurA** | Staurosporine | 17.2 | -0.3 | 6.5 | -2.1 | 8.6 | 4.4 | 5.9 |
| **BIKE** | Staurosporine | 19.0 | 0.4 | 0.3 | 0.7 | 6.4 | 6.5 | 6.6 |
| **BMPR2** | Lestaurtinib | 2.6 | -0.1 | -0.1 | 0.0 | -0.2 | 1.0 | -0.4 |
| **BMX** | Staurosporine | 7.0 | 1.7 | 3.1 | 2.1 | 6.6 | 7.0 | 7.3 |
| **BRAF** | Dabrafenib | 26.6 | 2.6 | 3.2 | 1.5 | 6.2 | 5.0 | 5.7 |
| **BRD4** | JQ1 | 7.0 | -3.9 | -3.7 | -2.2 | -2.4 | -1.6 | -2.7 |
| **BRPF1** | GSK6853 | 14.0 | -1.8 | -1.6 | -1.2 | -4.4 | -3.4 | -1.4 |

| | | | $\Delta T_m$ [°C] | | | | | |
|---|---|---|---|---|---|---|---|---|
| **Kinases** | **Reference compound** | **$\Delta T_m$ shift reference compound** | **MR30 (27)** | **28** | **29** | **30** | **31** | **32** |
| **CaMK1d** | Staurosporine | 9.9 | 2.4 | 4.8 | 3.7 | 6.8 | 5.6 | 6.0 |
| **CaMK1g** | Staurosporine | 11.0 | 3.6 | 5.7 | 5.0 | 7.5 | 5.5 | 6.4 |
| **CaMK2b** | Staurosporine | 13.2 | -1.1 | 0.7 | -0.5 | 0.3 | -0.4 | 0.3 |
| **CaMK2d** | Staurosporine | 16.5 | -0.5 | -0.4 | -0.1 | 0.1 | -0.1 | 0.0 |
| **CaMK4** | Staurosporine | 8.4 | 2.5 | 3.9 | 1.5 | 5.5 | 5.0 | 4.9 |
| **CaMKK2** | Staurosporine | 24.5 | 0.5 | 2.4 | 0.7 | 7.5 | 5.2 | 6.8 |
| **CASK** | Staurosporine | 5.0 | -0.3 | 0.0 | -0.5 | -0.5 | -0.2 | 0.0 |
| **CDK2** | Staurosporine | 15.0 | 1.4 | -0.2 | 0.9 | 4.8 | 6.7 | 5.3 |
| **CDKL1** | CEP-32496 | 6.6 | 0.9 | 0.7 | 0.5 | 1.6 | 2.1 | 1.9 |
| **CHK2** | Staurosporine | 16.7 | 2.5 | 4.6 | 2.4 | 10.4 | 11.2 | 11.0 |
| **CK1d** | SR-3029 | 6.8 | 1.6 | 1.4 | 1.3 | 2.8 | 2.8 | 2.3 |
| **CK1e** | PF-670462 | 9.0 | 6.4 | 4.9 | 5.8 | 8.2 | 7.5 | 7.7 |
| **CK1g** | Staurosporine | 2.0 | 0.3 | 0.5 | 0.4 | 1.2 | 1.8 | 1.2 |
| **CK2a1** | Silmitasertib | 14.6 | -0.4 | -0.6 | -0.2 | -1.1 | 0.0 | -0.3 |
| **CK2a2** | Silmitasertib | 15.7 | -6.3 | -4.2 | -1.6 | -1.8 | -0.9 | -1.6 |
| **CLK1** | Staurosporine | 12.3 | -0.7 | 0.6 | -0.3 | -0.2 | 0.8 | 0.7 |
| **CLK3** | T3 | 15.0 | -0.3 | 0.0 | 0.0 | -0.2 | -0.5 | 0.1 |
| **DAPK1** | Staurosporine | 9.4 | -1.2 | -0.7 | -0.9 | -0.7 | -0.6 | -0.9 |
| **DAPK3** | Staurosporine | 16.2 | 0.6 | 1.0 | 0.0 | 2.9 | 2.0 | 2.6 |
| **DCAMKL1** | Staurosporine | 10.7 | -1.8 | 0.9 | -1.7 | 2.0 | 1.1 | 1.4 |
| **DMPK1** | Staurosporine | 9.4 | -1.4 | -2.4 | -1.3 | -5.0 | -2.0 | -4.5 |
| **DRAK1** | Staurosporine | 7.7 | -0.4 | 0.3 | 1.3 | 4.0 | 4.5 | 5.1 |
| **DRAK2** | Staurosporine | 10.6 | 0.8 | 8.4 | 4.6 | 12.3 | 11.7 | 12.2 |
| **DYRK1A** | Staurosporine | 9.6 | -1.2 | 0.2 | -0.3 | -1.8 | -1.0 | -0.4 |
| **DYRK2** | Staurosporine | 7.3 | -0.4 | -0.7 | -0.2 | -1.1 | -0.8 | -1.4 |
| **EphA2** | Staurosporine | 7.3 | 3.6 | 4.5 | 5.1 | 7.7 | 8.1 | 7.9 |
| **EphA4** | Staurosporine | 5.5 | 2.0 | 3.4 | 2.1 | 3.5 | 4.4 | 4.6 |
| **EphA5** | Staurosporine | 8.1 | 2.8 | 4.5 | 2.7 | 6.6 | 6.2 | 6.6 |
| **EphA7** | Staurosporine | 11.3 | 1.7 | 4.2 | 4.1 | 6.4 | 5.1 | 5.4 |
| **EphB1** | Staurosporine | 6.4 | 3.0 | 5.1 | 4.3 | 6.5 | 6.6 | 6.6 |
| **EphB3** | Staurosporine | 5.9 | 2.3 | 4.1 | 2.3 | 6.9 | 5.3 | 8.5 |

| | | | ΔTm [°C] | | | | | |
|---|---|---|---|---|---|---|---|---|
| **Kinases** | **Reference compound** | **ΔTm shift reference compound** | **MR30 (27)** | **28** | **29** | **30** | **31** | **32** |
| **Erk2** | GDC-0994 | 7.5 | 0.4 | 0.7 | -0.4 | 3.5 | 1.3 | 1.9 |
| **Erk3** | Staurosporine | 1.2 | 0.5 | 0.7 | 0.3 | 1.1 | 1.2 | 1.0 |
| **Erk7** | Staurosporine | 14.2 | -0.4 | 0.5 | 0.1 | 1.2 | 1.4 | 1.0 |
| **FECH** | Vemurafenib | 5.5 | 0.5 | 10.6 | -0.4 | 0.5 | 2.0 | 3.8 |
| **FES** | Staurosporine | 7.1 | 5.0 | 8.5 | 6.0 | 9.1 | 8.7 | 8.8 |
| **FGFR1** | Staurosporine | 6.0 | 0.2 | 1.1 | 0.7 | 1.0 | 1.5 | 2.0 |
| **FGFR2** | Staurosporine | 8.5 | 1.0 | 3.8 | 1.7 | 3.9 | 3.9 | 4.2 |
| **FLT1** | Staurosporine | 12.2 | 2.4 | 7.8 | 3.9 | 4.3 | 5.6 | 5.6 |
| **GAK** | Staurosporine | 9.1 | 4.3 | 9.4 | 4.2 | 13.2 | 13.3 | 13.3 |
| **GPRK5** | Staurosporine | 5.9 | -1.1 | -0.6 | -0.4 | -0.9 | -0.2 | -0.4 |
| **GSK3B** | Staurosporine | 9.0 | 1.9 | 0.3 | 0.3 | 1.9 | 3.2 | 2.2 |
| **Haspin** | Staurosporine | 8.9 | -0.7 | -1.3 | -1.2 | 0.4 | -0.4 | -1.5 |
| **HIPK2** | Staurosporine | 4.8 | 0.0 | -0.5 | 0.6 | 2.6 | 2.6 | 2.6 |
| **JNK1** | Staurosporine | 7.6 | 0.4 | 2.8 | 0.7 | 2.8 | 2.7 | 1.9 |
| **JNK2** | SBI-0069279 | 8.6 | 0.7 | 0.0 | -0.1 | 4.3 | 4.2 | 3.3 |
| **JNK3** | CEP-32496 | 8.8 | 1.7 | 3.5 | 0.8 | 4.6 | 4.7 | 4.5 |
| **LOK** | Staurosporine | 22.8 | -0.6 | -0.7 | -0.4 | -0.4 | -0.4 | -0.2 |
| **MAP2K4** | Staurosporine | 11.7 | 5.2 | 11.0 | 3.5 | 9.7 | 10.8 | 10.8 |
| **MAP2K6** | Staurosporine | 12.0 | 4.1 | 7.5 | 1.9 | 9.4 | 9.4 | 8.9 |
| **MAP2K7** | Staurosporine | 7.3 | 2.6 | 2.9 | 1.4 | 6.2 | 8.9 | 7.3 |
| **MAP3K5** | Staurosporine | 17.5 | 9.2 | 12.2 | 9.6 | 13.1 | 12.3 | 12.7 |
| **MAPKAPK2** | Staurosporine | 3.4 | -0.3 | -0.2 | -0.2 | -0.3 | -0.1 | -0.1 |
| **MARK3** | Staurosporine | 19.1 | 4.7 | 3.8 | 5.8 | 11.6 | 10.9 | 10.9 |
| **MARK4** | Staurosporine | 16.8 | 3.8 | 2.2 | 4.8 | 9.5 | 10.4 | 9.9 |
| **MELK** | Staurosporine | 13.9 | 5.8 | 1.8 | 6.5 | 7.8 | 6.5 | 7.2 |
| **MER** | Staurosporine | 5.9 | 0.7 | 0.1 | 1.0 | 3.1 | 3.3 | 2.8 |
| **MRCKa** | Staurosporine | 2.6 | -0.2 | -1.0 | -0.3 | 4.4 | 3.3 | 4.0 |
| **MSK1b** | Staurosporine | 15.6 | -1.4 | 1.4 | 0.2 | 3.6 | 4.3 | 4.6 |
| **MSSK1** | Staurosporine | 3.7 | -0.4 | -0.9 | -0.3 | -0.2 | 0.0 | -0.2 |
| **MST1** | Staurosporine | 14.7 | -1.0 | -1.6 | -0.5 | 1.7 | 2.0 | 1.6 |
| **MST2** | Staurosporine | 13.4 | 5.9 | 10.1 | 9.1 | 15.0 | 14.3 | 14.7 |

| | | | $\Delta T_m$ [°C] | | | | | |
|---|---|---|---|---|---|---|---|---|
| **Kinases** | **Reference compound** | **$\Delta T_m$ shift reference compound** | **MR30 (27)** | **28** | **29** | **30** | **31** | **32** |
| **MST3** | Staurosporine | 7.0 | 9.8 | 12.1 | 14.3 | 15.5 | 16.1 | 15.4 |
| **MST4** | Staurosporine | 6.2 | 7.9 | 8.8 | 10.2 | 15.3 | 14.7 | 15.5 |
| **MYT1** | Dasatinib | 4.5 | -2.8 | -4.3 | -0.9 | -2.8 | -2.3 | -2.6 |
| **NDR2** | Staurosporine | 12.3 | 1.5 | 7.1 | 4.9 | 10.7 | 9.9 | 10.1 |
| **NEK1** | Staurosporine | 0.1 | -0.3 | 0.2 | 0.3 | -3.1 | -3.3 | -3.0 |
| **NEK2** | Staurosporine | 6.7 | 6.2 | 7.8 | 5.1 | 11.6 | 10.8 | 10.1 |
| **NEK7** | Staurosporine | 1.3 | -0.4 | -0.7 | -0.2 | -0.9 | -1.6 | -0.6 |
| **NQO2A** | Staurosporine | 1.2 | 0.5 | -0.7 | 0.3 | -0.2 | -0.3 | -0.2 |
| **OSR1** | Staurosporine | 7.4 | -1.1 | 0.1 | 0.3 | 2.3 | 3.1 | 3.1 |
| **p38a** | Dorama-pimod | 18.9 | 3.6 | 3.9 | 3.2 | 5.5 | 4.7 | 5.3 |
| **PAK1** | Staurosporine | 7.3 | 4.6 | 6.0 | 7.1 | 10.7 | 10.6 | 10.4 |
| **PAK4** | Staurosporine | 12.9 | 2.8 | 3.5 | 2.6 | 9.0 | 8.5 | 7.9 |
| **PCTAIRE1** | Staurosporine | 9.0 | 2.9 | 1.1 | 2.5 | 5.0 | 5.0 | 4.8 |
| **PHKG2** | Staurosporine | 21.8 | 1.2 | 2.4 | 2.2 | 6.7 | 6.4 | 6.4 |
| **PIM1** | Staurosporine | 12.2 | -0.5 | 0.5 | 0.6 | -0.4 | 3.7 | 1.2 |
| **PIM3** | Staurosporine | 19.8 | -2.7 | 0.5 | 0.0 | -0.8 | 4.5 | 1.5 |
| **PLK4** | Staurosporine | 17.9 | 4.5 | 8.0 | 4.2 | 12.6 | 12.2 | 11.5 |
| **RSK1b** | Staurosporine | 3.6 | -0.8 | -1.4 | -0.3 | -2.0 | -1.6 | -1.5 |
| **SLK** | Staurosporine | 18.4 | 2.6 | 4.5 | 3.5 | 8.6 | 7.7 | 7.8 |
| **SRC** | Staurosporine | 5.4 | -0.5 | 3.4 | 1.9 | 4.6 | 5.0 | 5.1 |
| **SRPK1** | Staurosporine | 6.7 | -0.2 | 0.1 | 0.1 | 0.0 | 0.3 | 0.4 |
| **STLK3** | Staurosporine | 11.4 | 10.0 | 10.4 | 8.8 | 16.6 | 16.6 | 16.6 |
| **TAF1** | Bromosporine | 7.4 | 0.0 | 0.4 | 0.2 | -0.5 | -1.0 | 0.5 |
| **TLK1** | Staurosporine | 8.5 | -2.0 | -1.8 | -1.5 | -0.9 | -1.6 | -0.8 |
| **TTK** | Staurosporine | 11.2 | -0.3 | -11.0 | -0.4 | 5.6 | 5.1 | 2.2 |
| **ULK3** | Staurosporine | 17.8 | 7.3 | 6.5 | 5.8 | 14.9 | 14.5 | 14.3 |
| **VRK1** | Staurosporine | 3.4 | -0.1 | 0.8 | 0.4 | 0.3 | 0.9 | 0.8 |
| **Wnk1** | Staurosporine | 1.9 | 0.8 | 2.8 | 0.6 | 4.8 | 3.1 | 3.5 |

**Table S3**. Results of the DSF panel assay of compounds **33–36** on 100 Kinases. Kinases are shown with their common abbreviation. For each kinase a reference compound is shown with its corresponding $\Delta T_m$ shift. Average of duplicate measurements.

| | | | **$\Delta T_m$ [°C]** | | | |
|---|---|---|---|---|---|---|
| **Kinases** | **Reference compound** | **$\Delta T_m$ shift reference compound** | **33** | **34** | **35** | **36** |
| **AAK1** | Staurosporine | 15.3 | 4.1 | 1.4 | 3.0 | 2.4 |
| **ABL1** | Staurosporine | 9.3 | 10.6 | 5.2 | 5.8 | 5.0 |
| **AKT3** | Staurosporine | 7.0 | 4.6 | 1.3 | 1.0 | 1.0 |
| **AurA** | Staurosporine | 17.2 | 2.8 | 3.4 | 8.6 | 5.1 |
| **BIKE** | Staurosporine | 19.0 | 7.2 | 0.8 | 1.3 | 1.0 |
| **BMPR2** | Lestaurtinib | 2.6 | -0.1 | 0.0 | -1.0 | -0.2 |
| **BMX** | Staurosporine | 7.0 | 7.1 | 4.1 | 4.4 | 3.3 |
| **BRAF** | Dabrafenib | 26.6 | 6.3 | 1.1 | 0.8 | 0.8 |
| **BRD4** | JQ1 | 7.0 | -3.7 | -1.8 | -3.6 | -3.5 |
| **BRPF1** | GSK6853 | 14.0 | -2.7 | -1.6 | -2.8 | -1.2 |
| **CaMK1d** | Staurosporine | 9.9 | 5.9 | 3.9 | 5.8 | 5.6 |
| **CaMK1g** | Staurosporine | 11.0 | 6.7 | 3.8 | 7.1 | 7.0 |
| **CaMK2b** | Staurosporine | 13.2 | -0.2 | -0.8 | -1.4 | -0.6 |
| **CaMK2d** | Staurosporine | 16.5 | -0.2 | -0.2 | -0.9 | -0.2 |
| **CaMK4** | Staurosporine | 8.4 | 5.2 | 3.9 | 5.7 | 3.2 |
| **CaMKK2** | Staurosporine | 24.5 | 4.3 | 4.3 | 6.9 | 5.8 |
| **CASK** | Staurosporine | 5.0 | 0.2 | -0.2 | 0.0 | -0.4 |
| **CDK2** | Staurosporine | 15.0 | 6.0 | 1.4 | 1.0 | 0.6 |
| **CDKL1** | CEP-32496 | 6.6 | 2.0 | 1.5 | 0.3 | 0.8 |
| **CHK2** | Staurosporine | 16.7 | 10.4 | 6.4 | 13.0 | 12.6 |
| **CK1d** | SR-3029 | 6.8 | 2.2 | 1.3 | 1.8 | 1.1 |
| **CK1e** | PF-670462 | 9.0 | 6.9 | 6.5 | 6.6 | 5.1 |
| **CK1g** | Staurosporine | 2.0 | 0.7 | 1.1 | 0.8 | 0.3 |
| **CK2a1** | Similtasertib | 14.6 | -0.4 | 0.5 | -0.1 | 0.2 |
| **CK2a2** | Similtasertib | 15.7 | -1.2 | 0.1 | 0.1 | -0.5 |
| **CLK1** | Staurosporine | 12.3 | -0.2 | -0.3 | -1.6 | -0.9 |
| **CLK3** | T3 | 15.0 | -0.5 | -1.0 | -1.3 | -0.7 |

| | | | $\Delta T_m$ [°C] | | | |
|---|---|---|---|---|---|---|
| **Kinases** | **Reference compound** | **$\Delta T_m$ shift reference compound** | **33** | **34** | **35** | **36** |
| **DAPK1** | Staurosporine | 9.4 | -1.1 | -0.8 | -2.1 | -1.2 |
| **DAPK3** | Staurosporine | 16.2 | 1.9 | 1.5 | 3.2 | 1.6 |
| **DCAMKL1** | Staurosporine | 10.7 | 1.2 | -0.3 | 2.0 | 1.3 |
| **DMPK1** | Staurosporine | 9.4 | -4.7 | -0.8 | -2.1 | -1.5 |
| **DRAK1** | Staurosporine | 7.7 | 3.5 | 2.2 | 5.2 | 6.1 |
| **DRAK2** | Staurosporine | 10.6 | 10.5 | 7.5 | 7.0 | 4.3 |
| **DYRK1A** | Staurosporine | 9.6 | -0.6 | -1.0 | -2.3 | -0.8 |
| **DYRK2** | Staurosporine | 7.3 | -1.0 | 0.0 | -0.6 | -0.4 |
| **EphA2** | Staurosporine | 7.3 | 8.3 | 3.9 | 4.6 | 2.4 |
| **EphA4** | Staurosporine | 5.5 | 3.5 | 2.5 | 2.7 | 2.4 |
| **EphA5** | Staurosporine | 8.1 | 6.8 | 3.2 | 4.2 | 3.5 |
| **EphA7** | Staurosporine | 11.3 | 5.3 | 3.4 | 5.8 | 5.0 |
| **EphB1** | Staurosporine | 6.4 | 6.5 | 5.2 | 6.0 | 5.5 |
| **EphB3** | Staurosporine | 5.9 | 8.7 | 4.1 | 4.5 | 3.3 |
| **Erk2** | GDC-0994 | 7.5 | -0.7 | -2.7 | -3.0 | -2.7 |
| **Erk3** | Staurosporine | 1.2 | 0.9 | 0.7 | 0.6 | 0.5 |
| **Erk7** | Staurosporine | 14.2 | 1.6 | -0.4 | 1.1 | -0.5 |
| **FECH** | Vemurafenib | 5.5 | -2.9 | 3.2 | 7.0 | 3.0 |
| **FES** | Staurosporine | 7.1 | 8.9 | 7.6 | 8.3 | 9.0 |
| **FGFR1** | Staurosporine | 6.0 | 2.0 | 0.9 | 0.4 | 2.0 |
| **FGFR2** | Staurosporine | 8.5 | 3.3 | 2.0 | 2.4 | 2.9 |
| **FLT1** | Staurosporine | 12.2 | 4.9 | 3.2 | 5.6 | 4.4 |
| **GAK** | Staurosporine | 9.1 | 12.5 | 9.3 | 13.5 | 11.7 |
| **GPRK5** | Staurosporine | 5.9 | -0.8 | -0.7 | -2.1 | -1.3 |
| **GSK3B** | Staurosporine | 9.0 | 2.3 | 2.0 | -0.3 | 0.2 |
| **Haspin** | Staurosporine | 8.9 | -0.2 | -0.2 | -0.1 | -0.7 |
| **HIPK2** | Staurosporine | 4.8 | 2.0 | 0.5 | 0.1 | 0.3 |
| **JNK1** | Staurosporine | 7.6 | 2.2 | 1.8 | 4.7 | 1.7 |
| **JNK2** | SBI-0069279 | 8.6 | 3.0 | 0.3 | 0.1 | -0.3 |
| **JNK3** | CEP-32496 | 8.8 | 5.2 | 3.0 | 2.3 | 1.1 |
| **LOK** | Staurosporine | 22.8 | -0.3 | -0.3 | -0.8 | -0.4 |

| | | | $\Delta T_m$ [°C] | | | |
|---|---|---|---|---|---|---|
| **Kinases** | **Reference compound** | **$\Delta T_m$ shift reference compound** | **33** | **34** | **35** | **36** |
| **MAP2K4** | Staurosporine | 11.7 | 10.0 | 8.0 | 7.6 | 6.9 |
| **MAP2K6** | Staurosporine | 12.0 | 10.2 | 5.4 | 6.8 | 4.4 |
| **MAP2K7** | Staurosporine | 7.3 | 7.7 | 1.3 | 2.0 | 0.0 |
| **MAP3K5** | Staurosporine | 17.5 | 14.1 | 10.3 | 12.2 | 10.1 |
| **MAPKAPK2** | Staurosporine | 3.4 | -0.5 | -0.1 | -0.6 | -0.1 |
| **MARK3** | Staurosporine | 19.1 | 11.5 | 5.1 | 6.9 | 8.5 |
| **MARK4** | Staurosporine | 16.8 | 10.3 | 4.6 | 6.6 | 7.4 |
| **MELK** | Staurosporine | 13.9 | 6.4 | 3.3 | 7.5 | 7.4 |
| **MER** | Staurosporine | 5.9 | 3.1 | 0.1 | 0.0 | -0.2 |
| **MRCKa** | Staurosporine | 2.6 | 1.9 | 0.8 | -2.1 | -0.9 |
| **MSK1b** | Staurosporine | 15.6 | 3.5 | 2.4 | 3.8 | 3.0 |
| **MSSK1** | Staurosporine | 3.7 | -0.4 | 0.4 | -0.2 | 0.0 |
| **MST1** | Staurosporine | 14.7 | 2.0 | -0.9 | -1.0 | -0.4 |
| **MST2** | Staurosporine | 13.4 | 14.6 | 10.8 | 11.3 | 10.1 |
| **MST3** | Staurosporine | 7.0 | 15.0 | 10.4 | 10.1 | 13.3 |
| **MST4** | Staurosporine | 6.2 | 11.8 | 8.1 | 8.6 | 6.9 |
| **MYT1** | Dasatinib | 4.5 | -1.8 | -2.7 | -3.5 | -2.8 |
| **NDR2** | Staurosporine | 12.3 | 10.4 | 6.0 | 6.5 | 5.2 |
| **NEK1** | Staurosporine | 0.1 | -3.6 | -1.0 | -0.7 | 0.0 |
| **NEK2** | Staurosporine | 6.7 | 10.6 | 8.4 | 10.6 | 9.2 |
| **NEK7** | Staurosporine | 1.3 | -1.4 | -1.0 | -2.4 | -1.4 |
| **NQO2A** | Staurosporine | 1.2 | 0.9 | -1.3 | -2.4 | -1.0 |
| **OSR1** | Staurosporine | 7.4 | 2.4 | 0.4 | 2.6 | 3.4 |
| **p38a** | Doramapimod | 18.9 | 4.7 | 3.0 | 3.8 | 3.7 |
| **PAK1** | Staurosporine | 7.3 | 10.8 | 5.3 | 5.2 | 8.7 |
| **PAK4** | Staurosporine | 12.9 | 8.2 | 3.0 | 4.1 | 3.9 |
| **PCTAIRE1** | Staurosporine | 9.0 | 5.2 | 0.6 | 0.1 | 0.2 |
| **PHKG2** | Staurosporine | 21.8 | 6.2 | 3.5 | 8.8 | 8.0 |
| **PIM1** | Staurosporine | 12.2 | 3.3 | 2.7 | -0.9 | -0.1 |
| **PIM3** | Staurosporine | 19.8 | 4.6 | 2.4 | -2.4 | -2.1 |
| **PLK4** | Staurosporine | 17.9 | 13.1 | 8.8 | 13.6 | 11.8 |

| | | | $\Delta T_m$ [°C] | | | |
|---|---|---|---|---|---|---|
| **Kinases** | **Reference compound** | **$\Delta T_m$ shift reference compound** | **33** | **34** | **35** | **36** |
| **RSK1b** | Staurosporine | 3.6 | -1.8 | -0.3 | -0.6 | 0.3 |
| **SLK** | Staurosporine | 18.4 | 8.0 | 4.7 | 5.1 | 4.2 |
| **SRC** | Staurosporine | 5.4 | 4.7 | 3.1 | 4.4 | 4.1 |
| **SRPK1** | Staurosporine | 6.7 | -0.2 | 0.1 | -0.5 | 0.0 |
| **STLK3** | Staurosporine | 11.4 | 16.8 | 11.0 | 13.4 | 9.3 |
| **TAF1** | Bromosporine | 7.4 | -0.2 | 0.1 | -0.7 | -0.4 |
| **TLK1** | Staurosporine | 8.5 | -0.8 | -1.1 | -2.5 | -1.9 |
| **TTK** | Staurosporine | 11.2 | 1.3 | 2.9 | 7.8 | 4.3 |
| **ULK3** | Staurosporine | 17.8 | 13.9 | 8.3 | 12.7 | 8.9 |
| **VRK1** | Staurosporine | 3.4 | 0.4 | 0.0 | -0.2 | 0.1 |
| **Wnk1** | Staurosporine | 1.9 | 3.3 | 0.8 | 2.7 | 1.5 |

**Table S4**. NanoBRET assay information. Tracer K8, K9 and K10 have the following ordering numbers at Promega: N2622, N2632 and N2642, respectively)

| **Protein Kinase** | **Alias** | **Catalog #/ CAS #** | **NanoLuc orientation** | **Tracer, used** | **Tracer $K_{D,app}$ [nM]** | **Tracer, used [nM]** |
|---|---|---|---|---|---|---|
| STK4 | MST1 | NV4351 | N | K10 | 165 | 170 |
| STK3 | MST2 | NV4301 | N | K10 | 270 | 300 |
| STK24 | MST3 | NV4281 | C | K9 | 150 | 150 |
| STK26 | MST4 | NV4291 | C | K9 | 480 | 500 |
| SNF1LK2 | SIK2 | NV2061 | N | K10 | 30 | 50 |
| CSNK1E | CK1e | NV2971 | N | K8 | 1000 | 1000 |
| FES | | NV3091 | C | K10 | 985 | 900 |
| NEK2 | | NV1781 | N | K10 | 577 | 600 |

**Table S5.** X-ray data collection and refinement statistics of MST3 inhibitor complexes

| | MRLW5 (**20**) | MR24 (**24**) | MR26 (**26**) | MR30 (**27**) |
|---|---|---|---|---|
| **Data collection** | | | | |
| Space group | P1 | $P2_1$ | C2 | $P2_1$ |
| Molecules asymmetric unit | 2 | 2 | 1 | 1 |
| a, b, c (Å) | 55.0, 57.7, 61.8 | 55.5, 93.6, 61.8 | 99.1, 58.8, 61.5 | 43.6, 58.6, 61.2 |
| α, β, γ (°) | 88.9, 89.2, 62.1 | 90.0, 92.3, 90.0 | 90.0, 93.1, 90.0 | 90.0, 107.3, 90.0 |
| Resolution (Å)[a] | 48.7-2.52 (2.62-2.52) | 47.7-1.85 (1.89-1.85) | 39.6-1.61 (1.64-1.61) | 41.6-1.47 (1.50-1.47) |
| Unique reflections[a] | 22,207 (2,545) | 52,593 (3,231) | 45,283 (2,235) | 49,300 (2,453) |
| Completeness (%)[a] | 97.8 (97.8) | 98.0 (98.3) | 99.0 (98.0) | 98.6 (98.9) |
| Multiplicity[a] | 3.4 (3.5) | 5.9 (6.0) | 7.0 (6.5) | 5.1 (5.2) |
| $R_{merge}$ | 0.052 (0.628) | 0.037 (0.717) | 0.035 (0.725) | 0.030 (0.725) |
| CC1/2[a] | 0.988 (0.779) | 1.000 (0.890) | 1.000 (0.837) | 1.000 (0.811) |
| Mean $I/\sigma(I)$[a] | 10.0 (1.6) | 19.5 (2.1) | 24.2 (2.5) | 21.8 (2.2) |
| **Refinement** | | | | |
| $R_{work}$, (%)[b] | 18.2 | 18.0 | 15.2 | 15.2 |
| $R_{free}$, (%)[b] | 25.0 | 21.7 | 18.6 | 17.6 |
| No. of atoms | | | | |
| Protein[c] | 4017 | 4044 | 2111 | 2132 |
| Inhibitor | 64 | 72 | 40 | 40 |
| Ethylene glycol | 4 | 12 | 36 | 12 |
| Water | 78 | 200 | 245 | 260 |
| Overall B-factor ($Å^2$) | 57.9 | 49.6 | 35.7 | 29.7 |
| RMSD bond lengths (Å) | 0.008 | 0.008 | 0.007 | 0.006 |
| RMSD bond angles (°) | 0.96 | 0.87 | 0.89 | 0.85 |
| Ramachandran favored (%)[d] | 94.1 | 96.1 | 96.1 | 97.0 |
| Ramachandran outliers (%)[d] | 0.2 | 0.0 | 0.0 | 0.0 |
| **Protein Data Bank entry** | 8BZJ | 8QLR | 8QLS | 8QLT |

[a]Values in parentheses are for the highest-resolution shell.
[b]$R_{work}$ and $R_{free}$ = $\sum||F_{obs}| - |F_{calc}||/\sum|F_{obs}|$, where $R_{free}$ was calculated with 5% of the reflections chosen at random and not used in the refinement.
[c]Number includes alternative conformations.
[d]MolProbity statistics

**Table S6.** MR24 (**24**) and MR26 (**26**) interaction patterns derived from Molecular Dynamics simulations for selected systems. The percentages give the fraction of the simulation time that the ligand is in contact with protein residues. **HB** – hydrogen bond, **WB** – water bridge, **SB** – salt bridge, **BB** – backbone, **SC** – side chain. (i.e., HB SC – hydrogen bond from the ligand to side chain of a residue in the protein). The conserved residues within the kinase binding site are abbreviated as following: I–VIII – β-sheets I–VIII; **g.l.** – G-rich loop, **b.l.** – loop connecting αC-helix to IV; **GK** – gatekeeper; **linker** – loop connecting the hinge to αD-helix; **c.l.** – catalytic loop; DFG – DFG-motif. The residues in the rows are superimposed, with conserved residues highlighted in yellow. For example, catalytic lysine from β-V is present in MST1 (K32), MST2 (K41), MST3 (K65), MST4 (K36), and SIK2 (K49) at the same position, therefore the cell is colored in yellow. The hinge residues are different among the proteins. However, they are at the same position among superimposed structures. Molecular dynamics timescale for individual replica is 5 μs. An asterisk indicates interactions with multiply substructures of the protein. The displayed minimum contact strength is 20%.

| | MST1 | | | MST2 | | | MST3 | | | MST4 | | |
|---|---|---|---|---|---|---|---|---|---|---|---|---|
| **Residue location** | Res № | **MR24** | **MR26** | Res № | **MR24** | **MR26** | Res № | **MR24** | **MR26** | Res № | **MR24** | **MR26** |
| **β-V** | **K32** | 61% Pi-cat | 60% Pi-cat | **K41** | 80% Pi-cat | 70% Pi-cat | **K65** | 35% Pi-cat | 40% Pi-cat | **K36** | 40% Pi-cat | 36% Pi-cat |
| | | 37% HB SC | 49% HB SC | | | | | 94% HB SC | 87% HB SC | | 90% HB SC | 96% HB SC |
| | | 61% WB | 63% WB | | 61% WB | 55% WB | | 67% WB | 68% WB | | 68% WB | 65% WB |
| **Hinge** | **C78** | 97% HB BB | 99% HB BB | **C87** | 99% HB BB | 99% HB BB | **L114** | 99%* HB BB | 99% HB BB | **L85** | 99%* HB BB | 99% HB BB |
| **DFG** | D140 | | | **D149** | 41% WB | 49% WB | **D174** | 55%** WB | 43%** WB | **D145** | 43%** WB | 47%** WB |
| | | | | | | 25% HB SC | | | | | 40% Pi-cat | 36% Pi-cat |
| **DFG** | F141 | | | **F150** | 20% WB | | F175 | | | F146 | | |
| **c.l** | G126 | | | **G135** | | 25% HB BB | **A160** | 51% HB BB | | **A131** | 43% HB BB | 59% HB BB |
| **g.l** | E11 | | | **E20** | 21% HB SC | 29% HB SC | K44 | | | K15 | | |
| **αD** | D85 | | | **D94** | 40% HB SC | | D121 | | | D92 | | |
| | | | | | | | D174 ** | 48 % HB SC | 39% HB SC | D145 ** | 41% HB SC | 45% HB SC |
| **αD** | **D85** | 23% WB | | D94 | | | **D121** | 24% HB SC | | **D92** | 24% WB | |
| **Linker** | **G79** | | | **G88** | | 38% HB BB | G115 | | | G86 | | |
| **Linker** | **G81** | | 79% HB BB | **G90** | 58% HB | 24% HB | G117 | | | G88 | | |
| | K274 | | | **K283** | 31% HB SC | | K307 | | | K278 | | |
| **Hinge** | Y77 | | | Y86 | | | Y113 | | | **Y84** | | 34% WB |

**Table S7**. Percentage of hydrophobic ligand–protein interactions derived from Molecular Dynamics simulations for selected systems. The conserved residues within the kinase binding site are abbreviated as following: I–VIII – β-sheets I–VIII; **g.l.** – G-rich loop, **b.l.** – loop connecting αC-helix to IV; **GK** – gatekeeper; **linker** – loop connecting the hinge to αD-helix; **c.l.** – catalytic loop; DFG – DFG-motif; **RS3** – hydrophobic regulatory spine residue 3 (R-spine). Molecular dynamics timescale for individual replica is 5 μs. The displayed minimum contact strength is 20%.

| residue location | | MST1 | | | MST2 | | | MST3 | | | MST4 | |
|---|---|---|---|---|---|---|---|---|---|---|---|---|
| | | MR24 | MR26 | | MR24 | MR26 | | MR24 | MR26 | | MR24 | MR26 |
| **β-III** | **A30** | 45% | 34% | **A39** | 39% | 34% | **A63** | 37% | 42% | **A34** | 41% | 35% |
| **GK** | **M75** | 45% | 42% | **M84** | 37% | 40% | **M111** | 51% | 46% | **M82** | 50% | 47% |
| **β-VII** | L129 | | | **L138** | 29% | 34% | **L163** | 25% | 38% | **L134** | 24% | 23% |
| **β-V** | **I73** | 38% | 35% | I82 | | | **I109** | 47% | 37% | **I80** | 41% | 47% |
| **RS3** | **M50** | | | M59 | | | **L86** | 48% | 43% | **L57** | 46% | 50% |
| **αC** | I47 | | | **I56** | 21% | | I83 | | | I54 | | |

**Analytical data of compounds 9–36 and intermediates 38–70.**

$^{1}H$ NMR compound **38**:

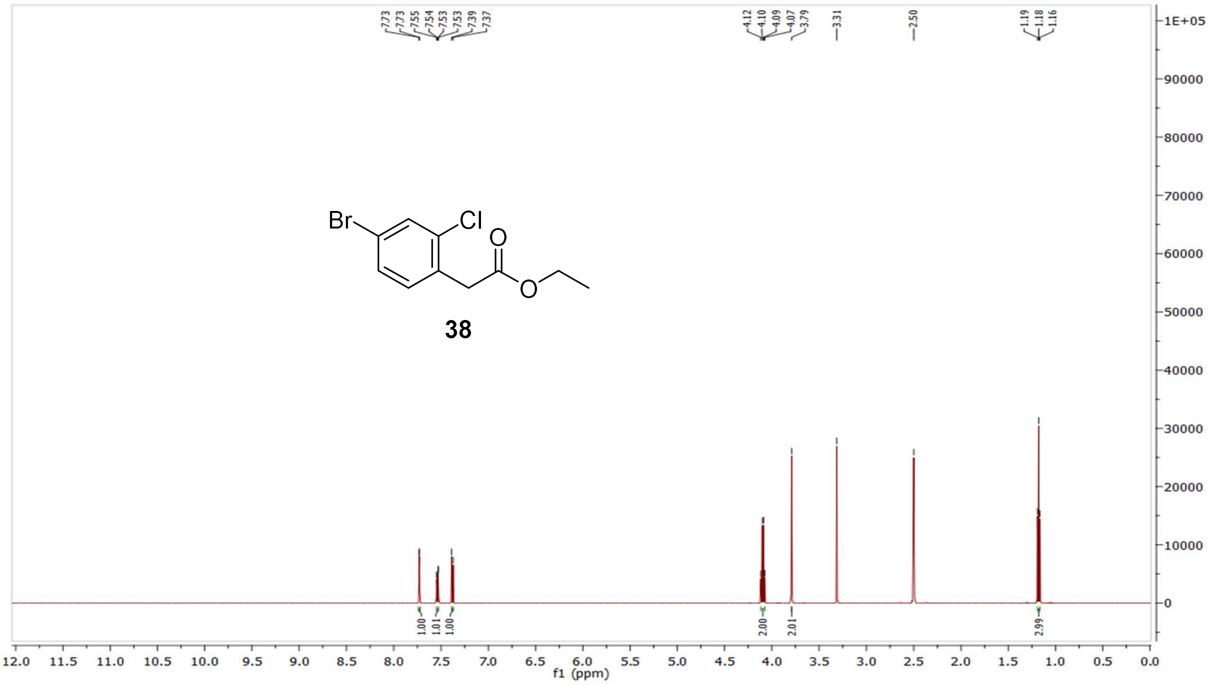

$^{13}C$ NMR compound **38**:

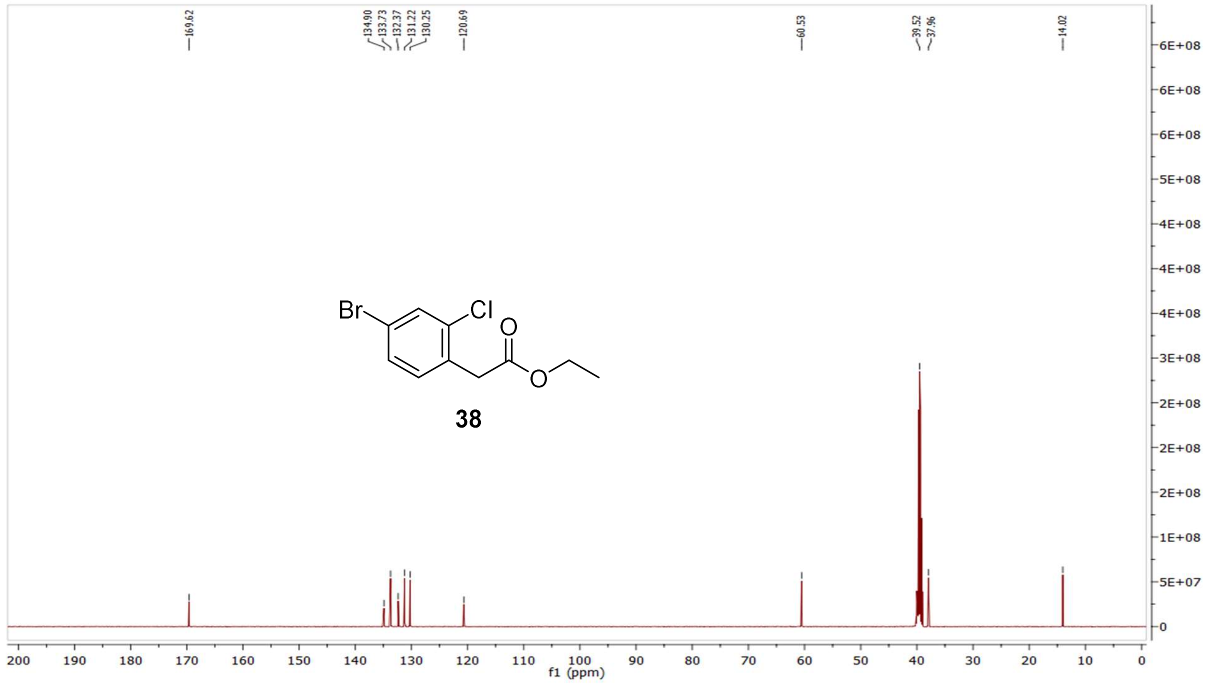

MS (ESI+) compound **38**:

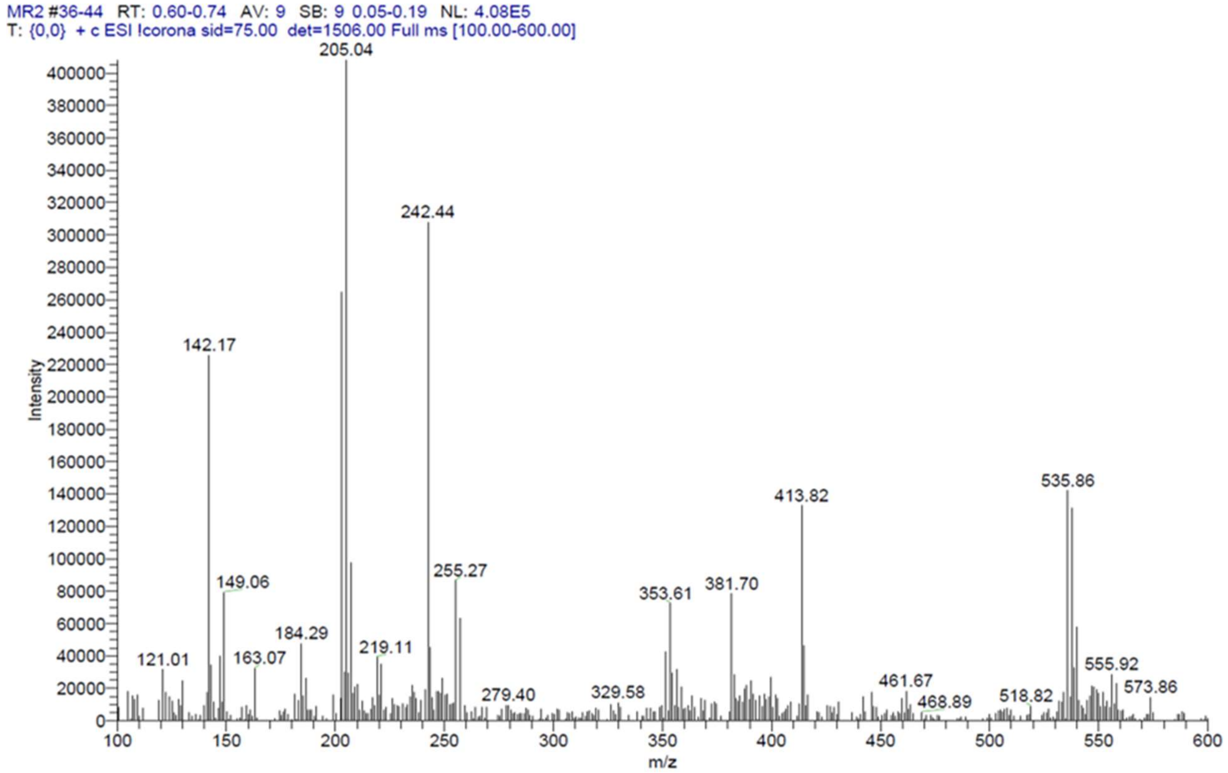


$^1$H NMR compound **39**:

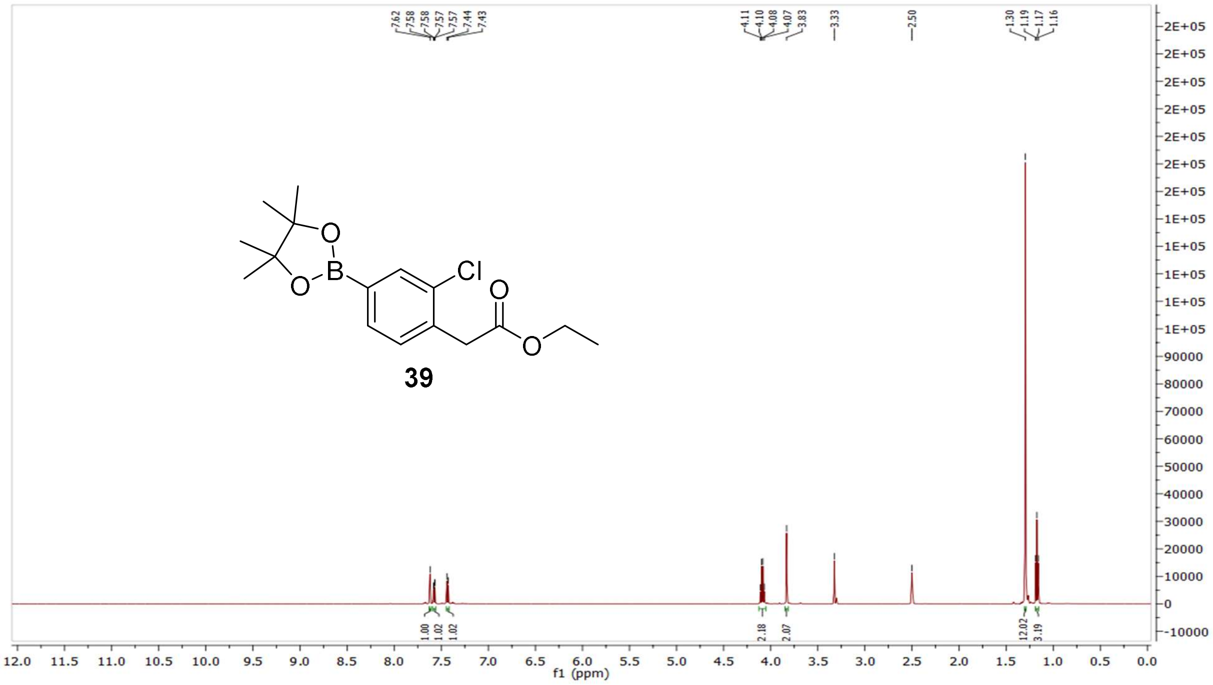

[13]C NMR compound **39**:

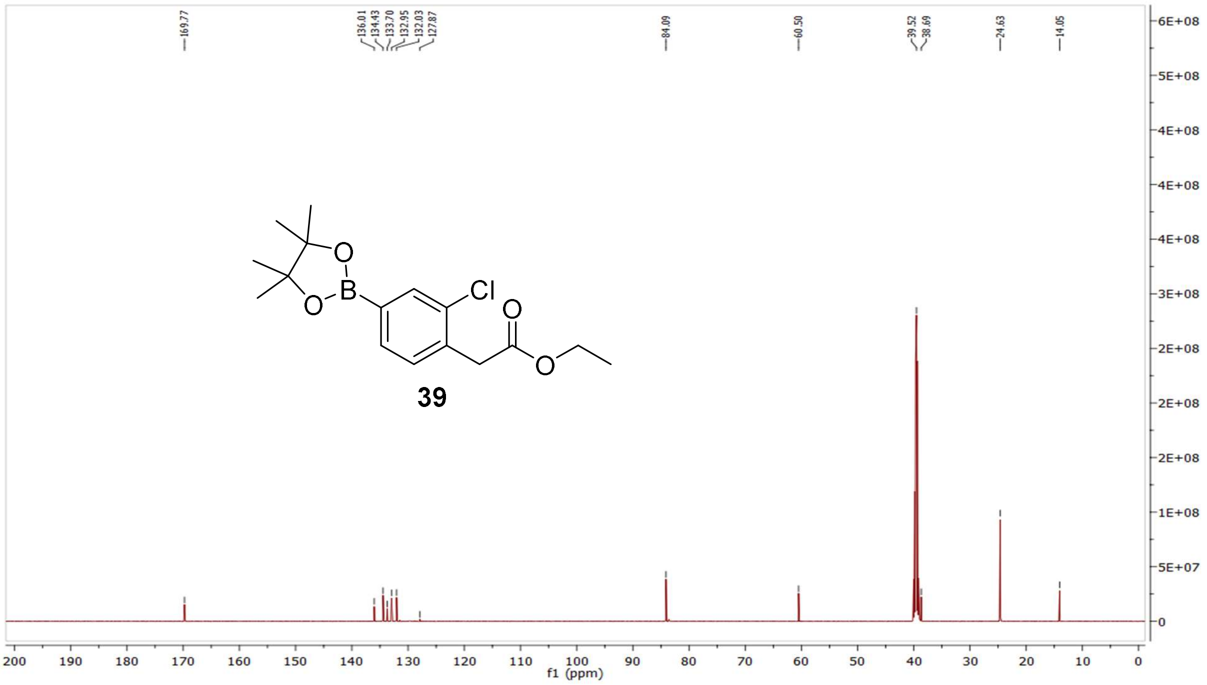


MS (ESI+) compound **39**:

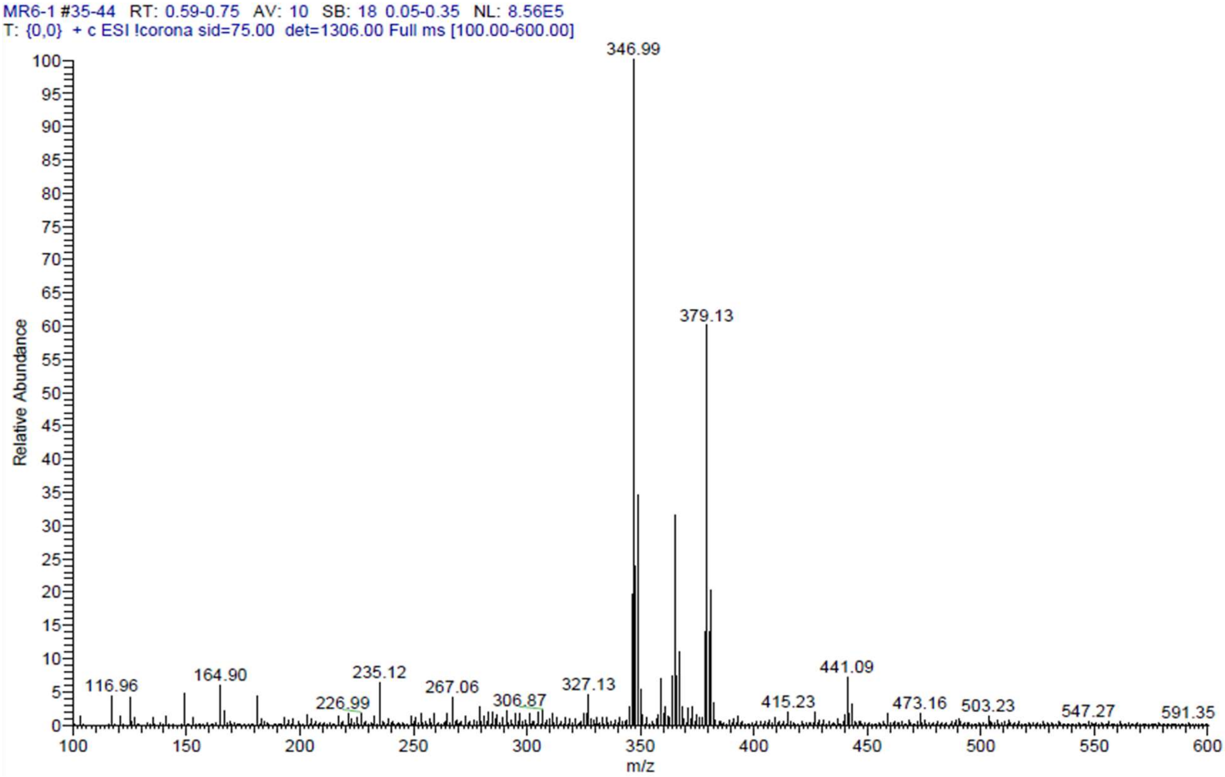

[1]H NMR compound **40**:

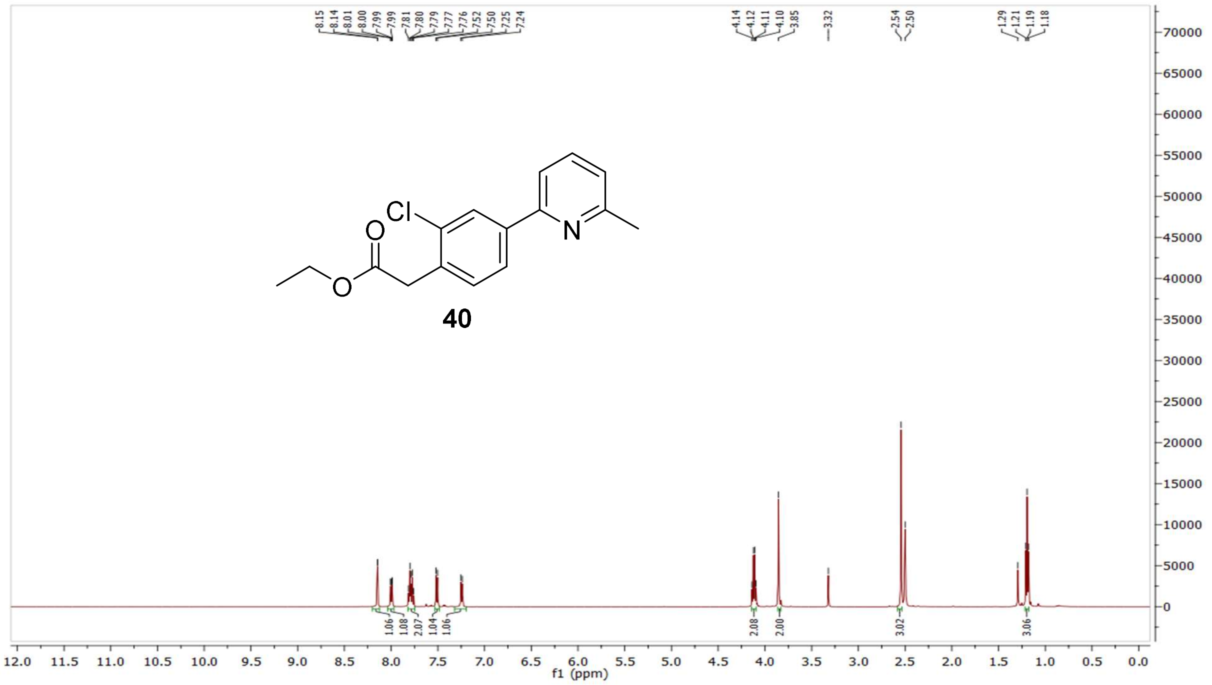

[13]C NMR compound **40**:

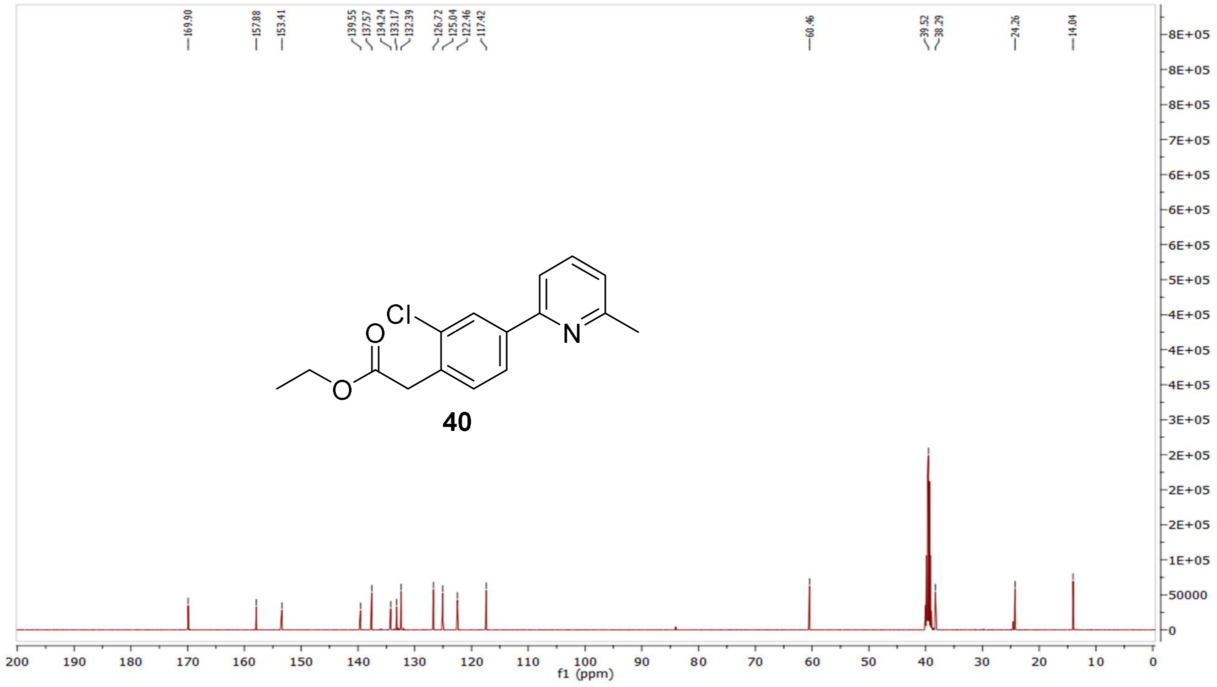

MS (ESI+) compound **40**:

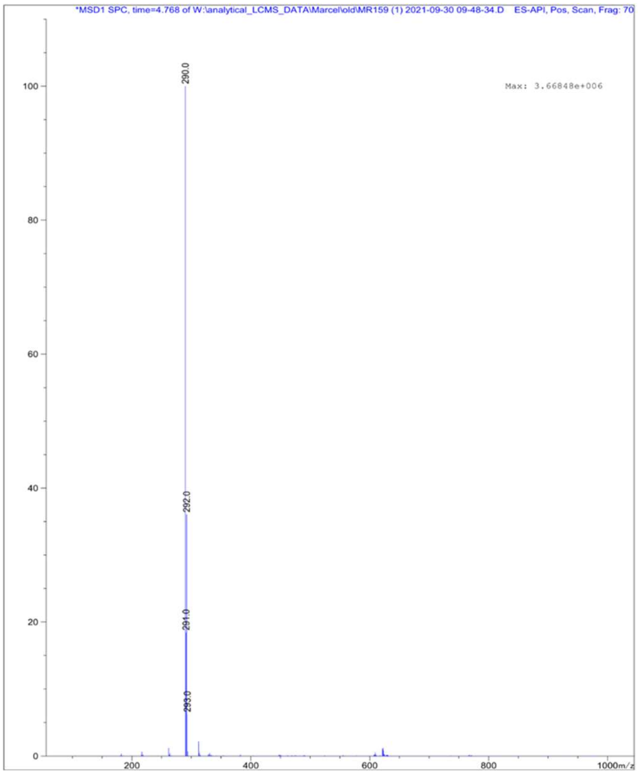


[1]H NMR compound **41**:

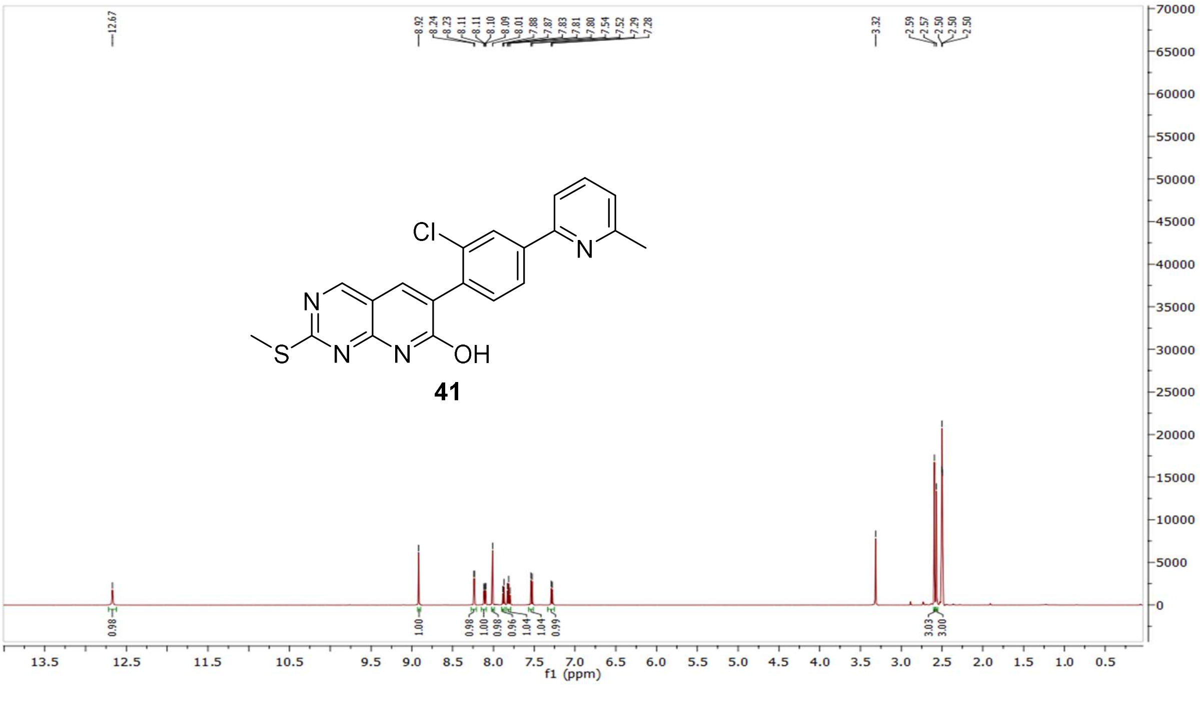

$^{13}C$ NMR compound **41:**

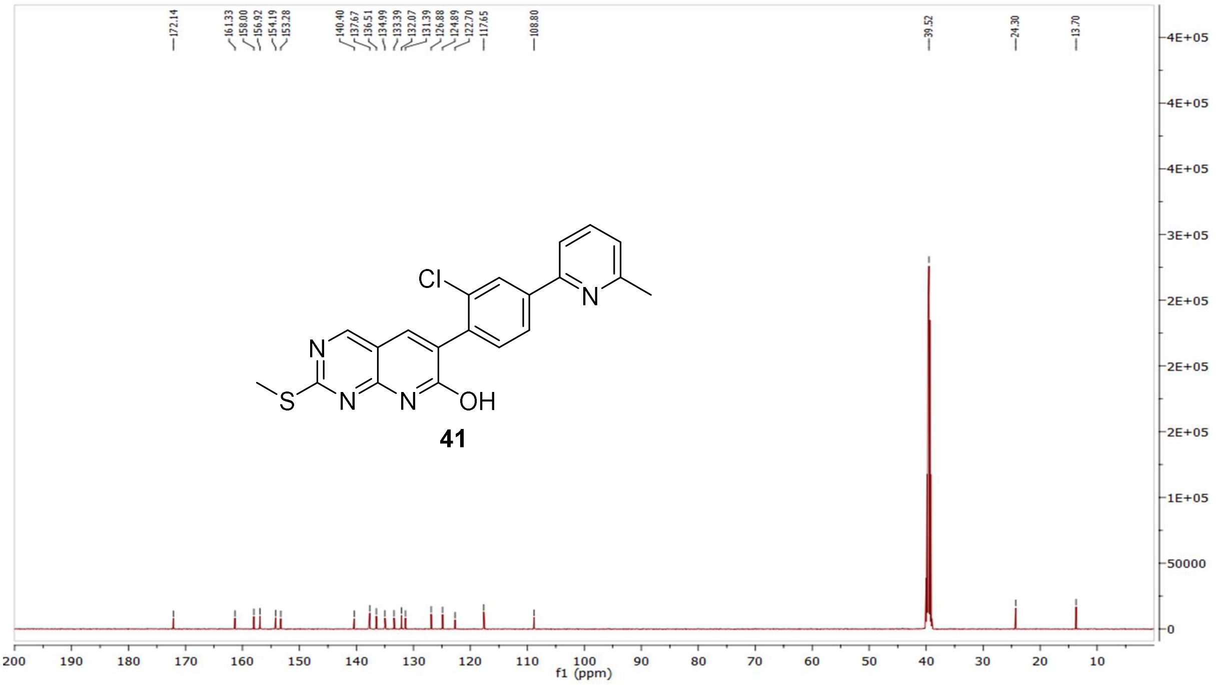

MS (ESI+) compound **41**:

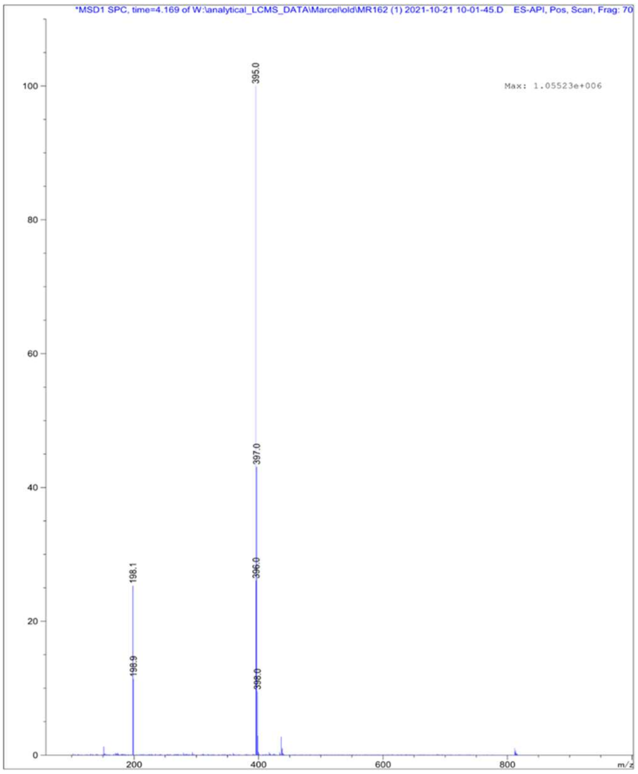

$^{1}H$ NMR compound **42**:

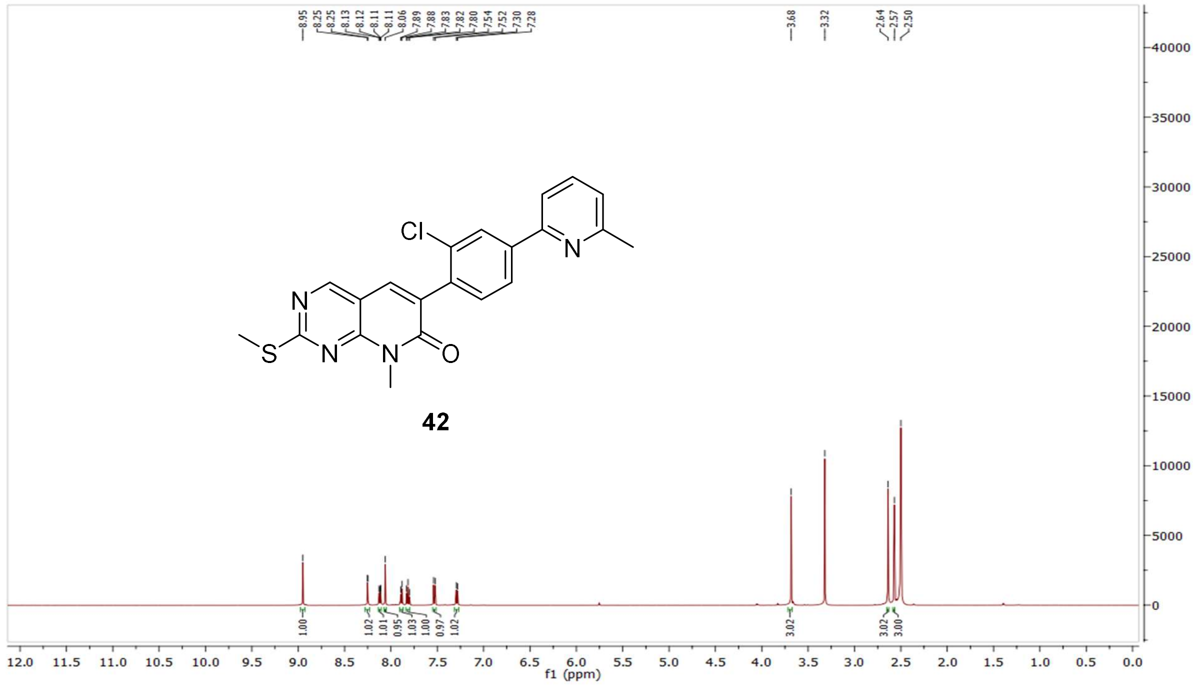

$^{13}C$ NMR compound **42:**

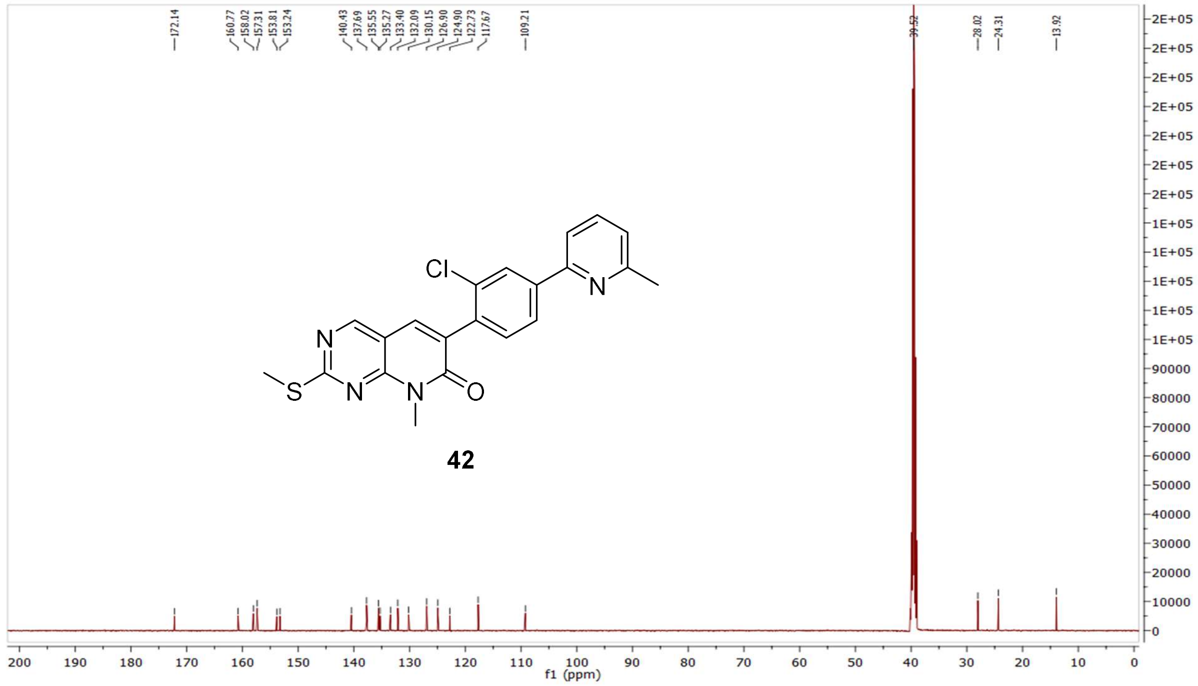

MS (ESI+) compound **42**:

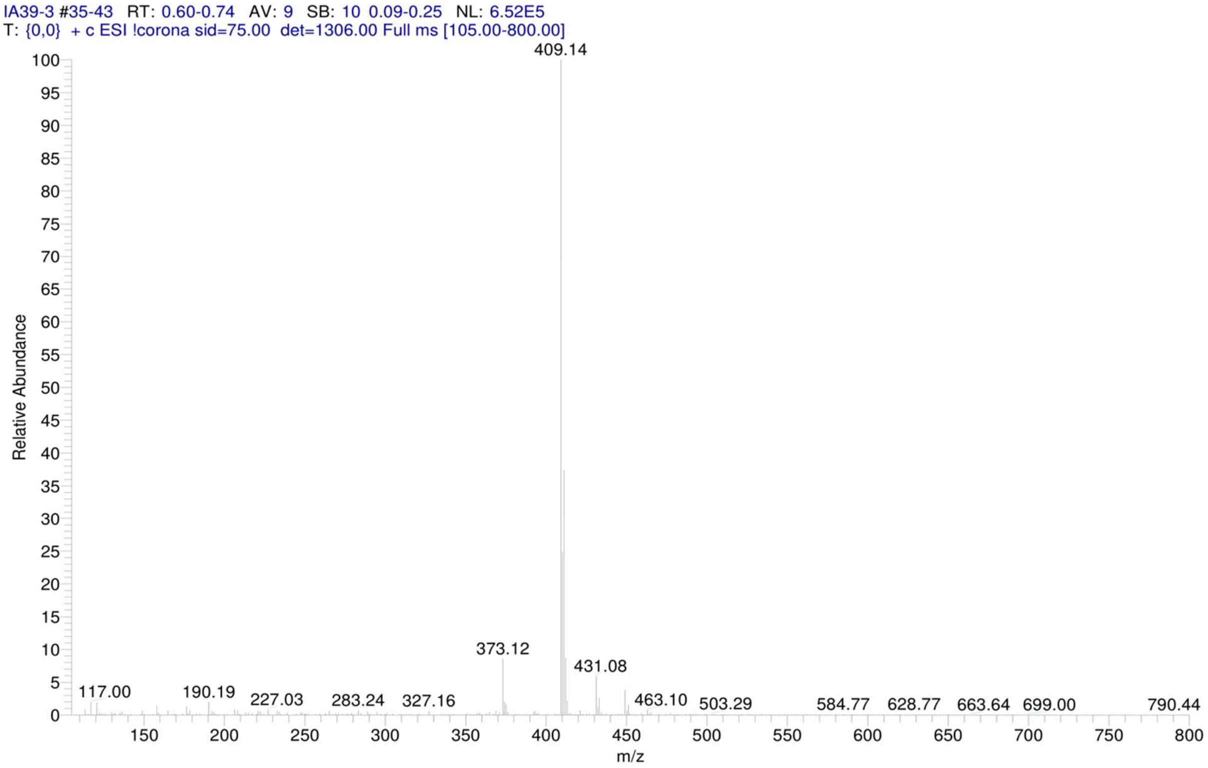


[1]H NMR compound **43**:

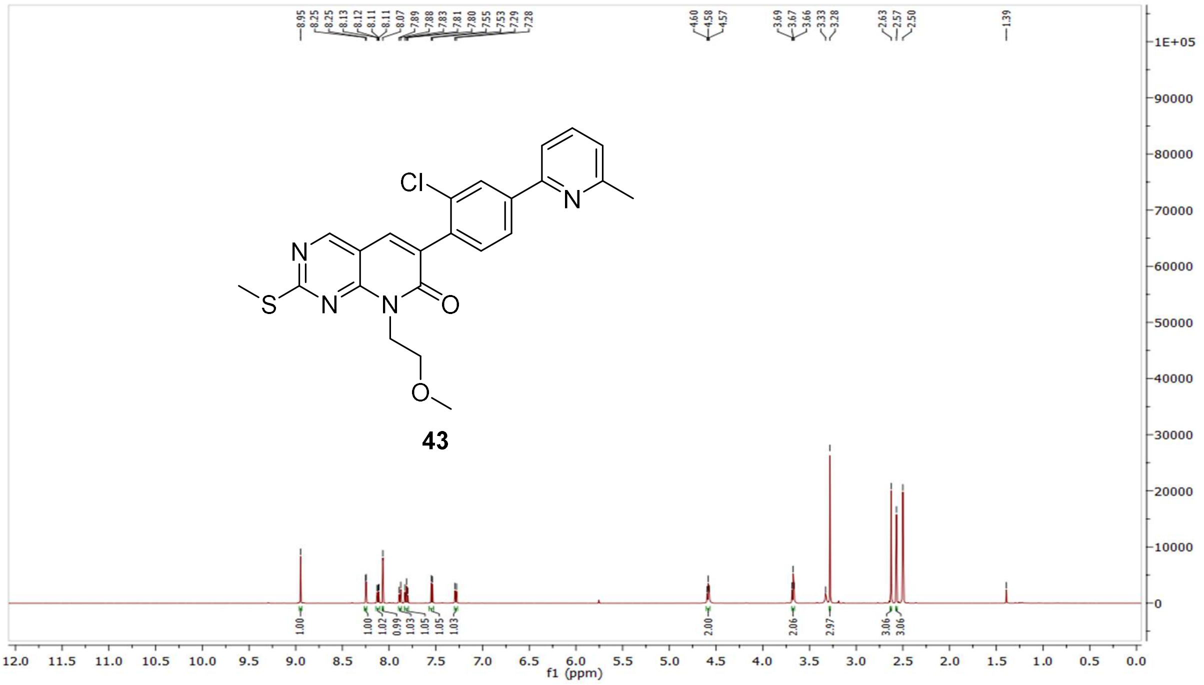

$^{13}$C NMR compound **43:**

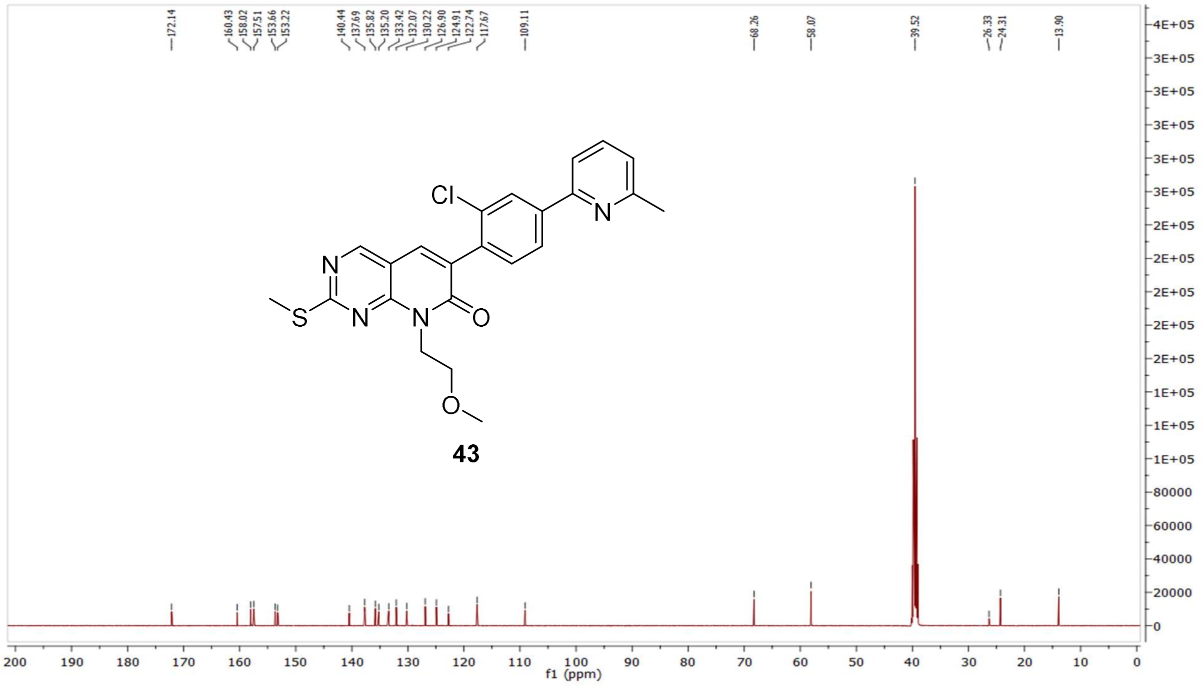


MS (ESI+) compound **43**:

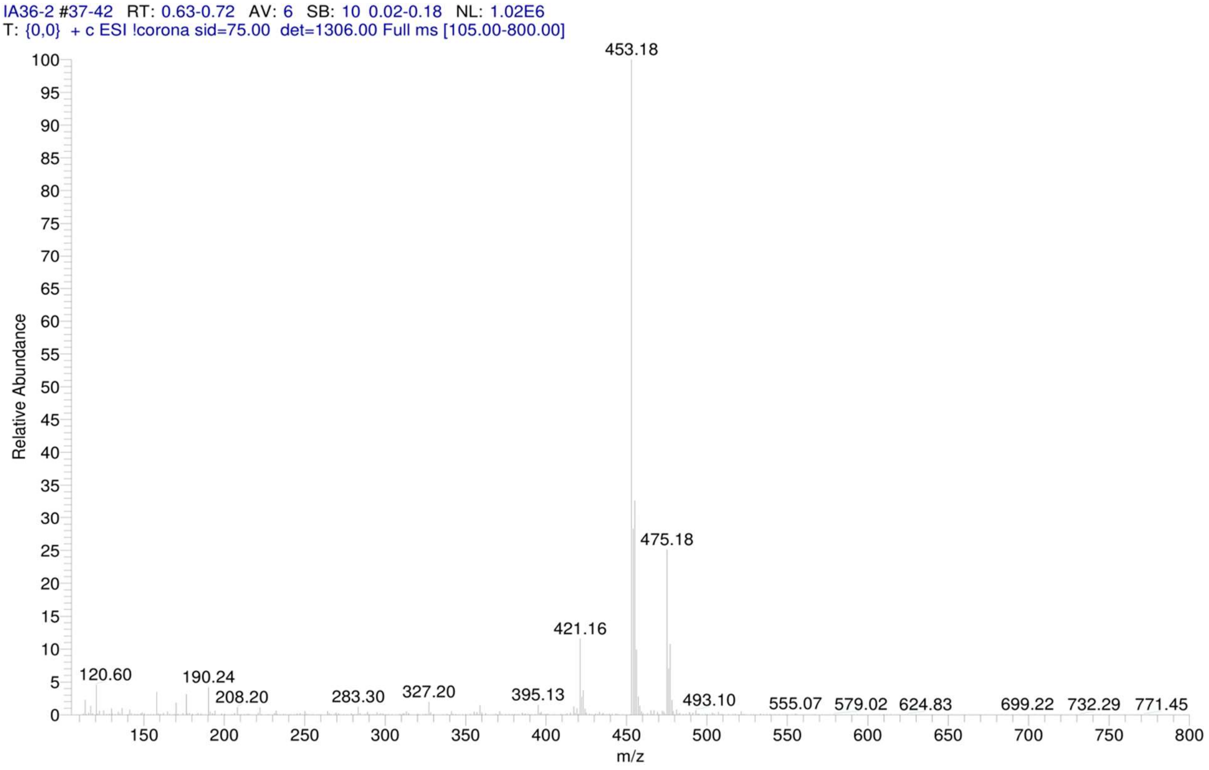

$^{1}$H NMR compound **44**:

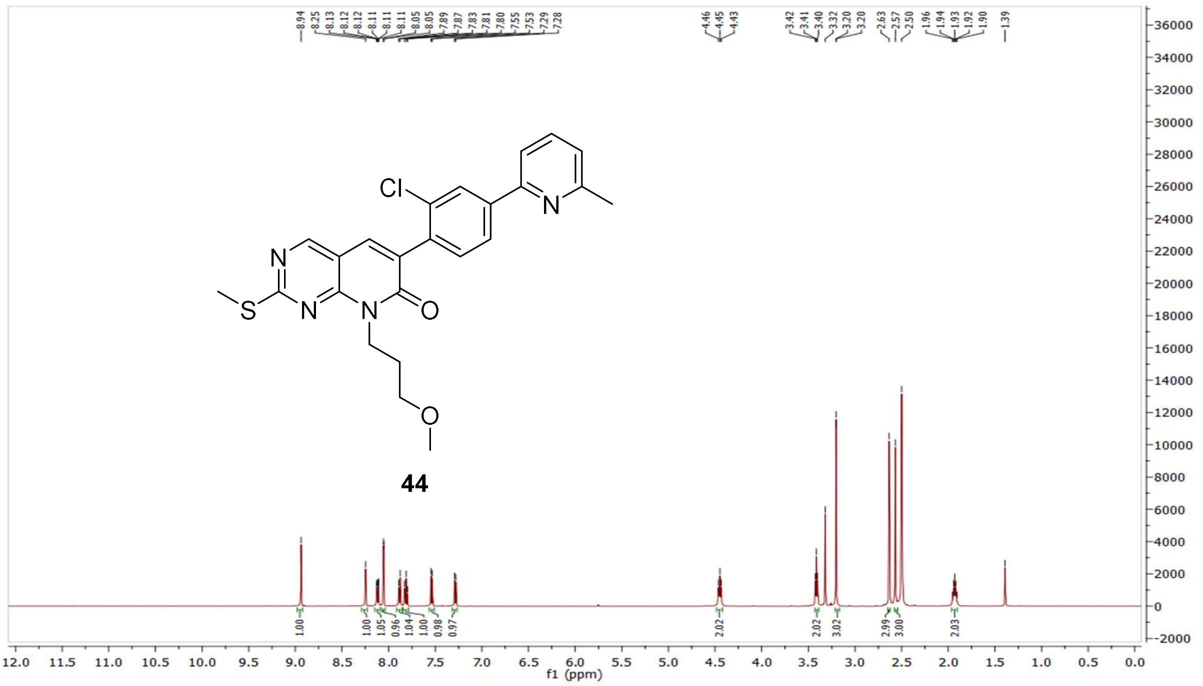

$^{13}$C NMR compound **44:**

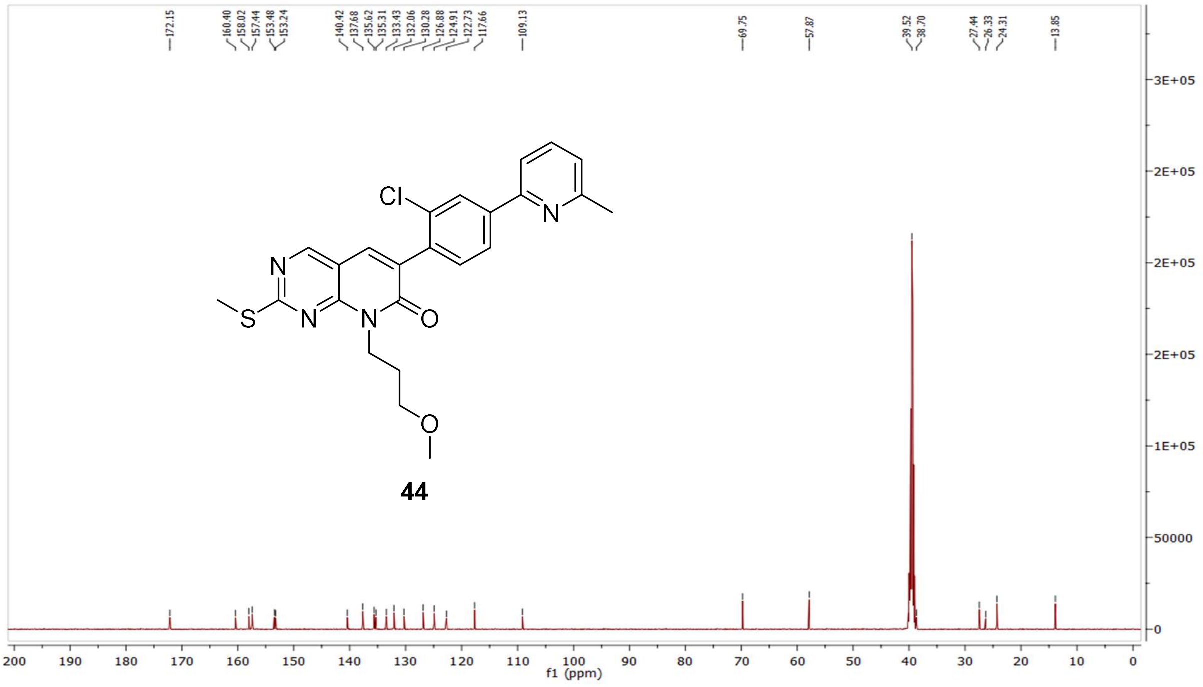

MS (ESI+) compound **44**:

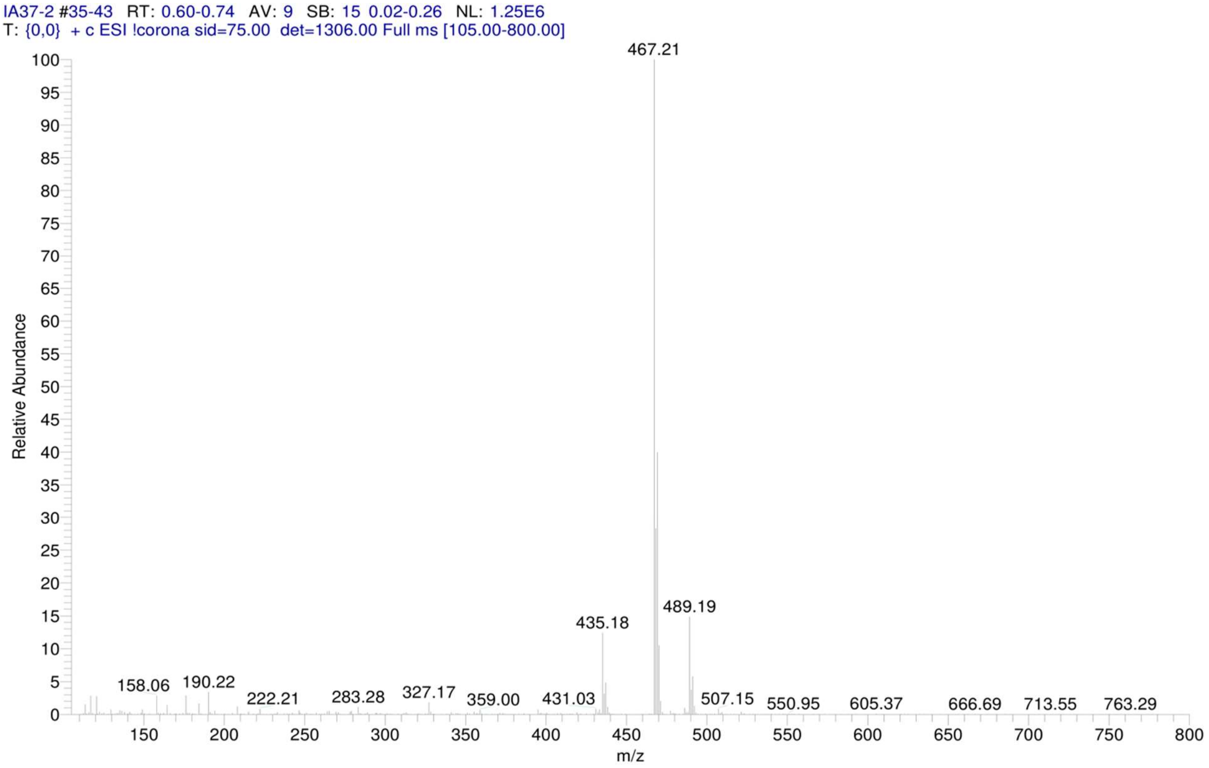


[1]H NMR compound **45**:

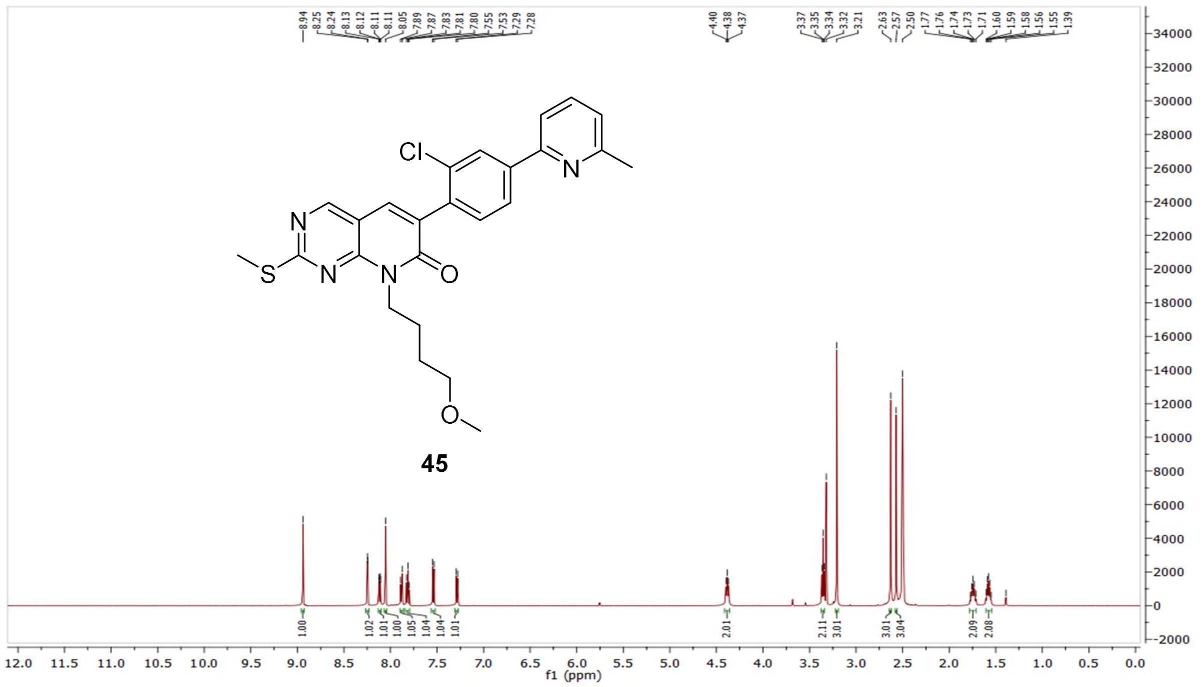

$^{13}$C NMR compound **45:**

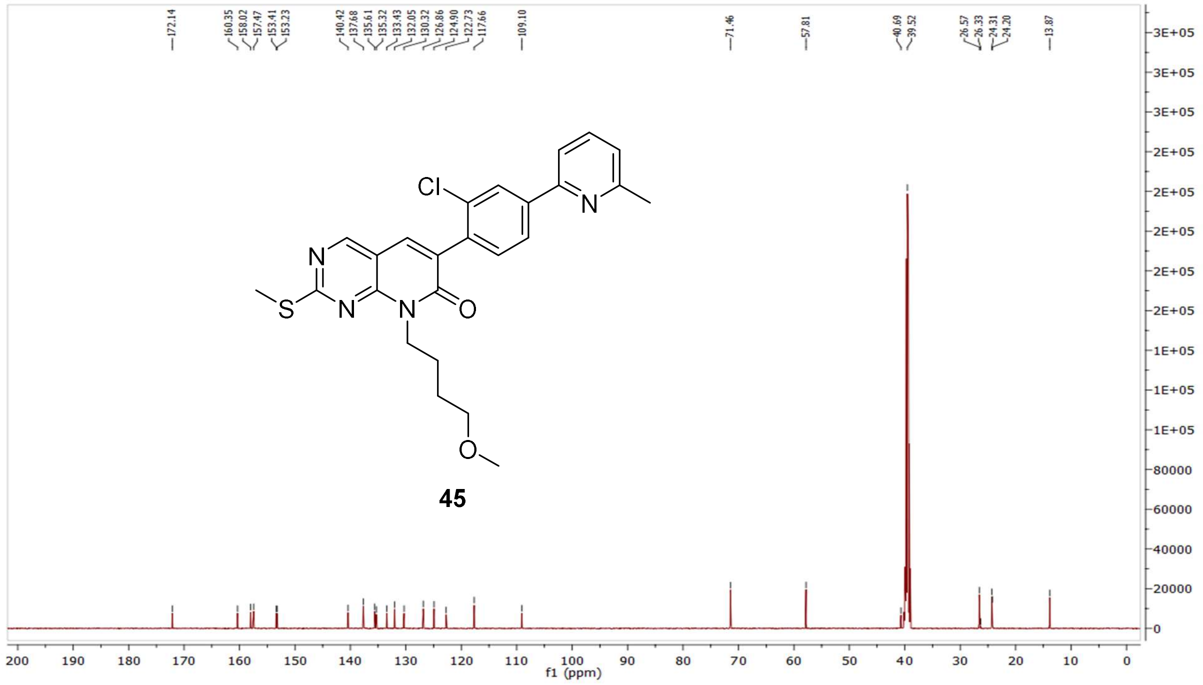


MS (ESI+) compound **45**:

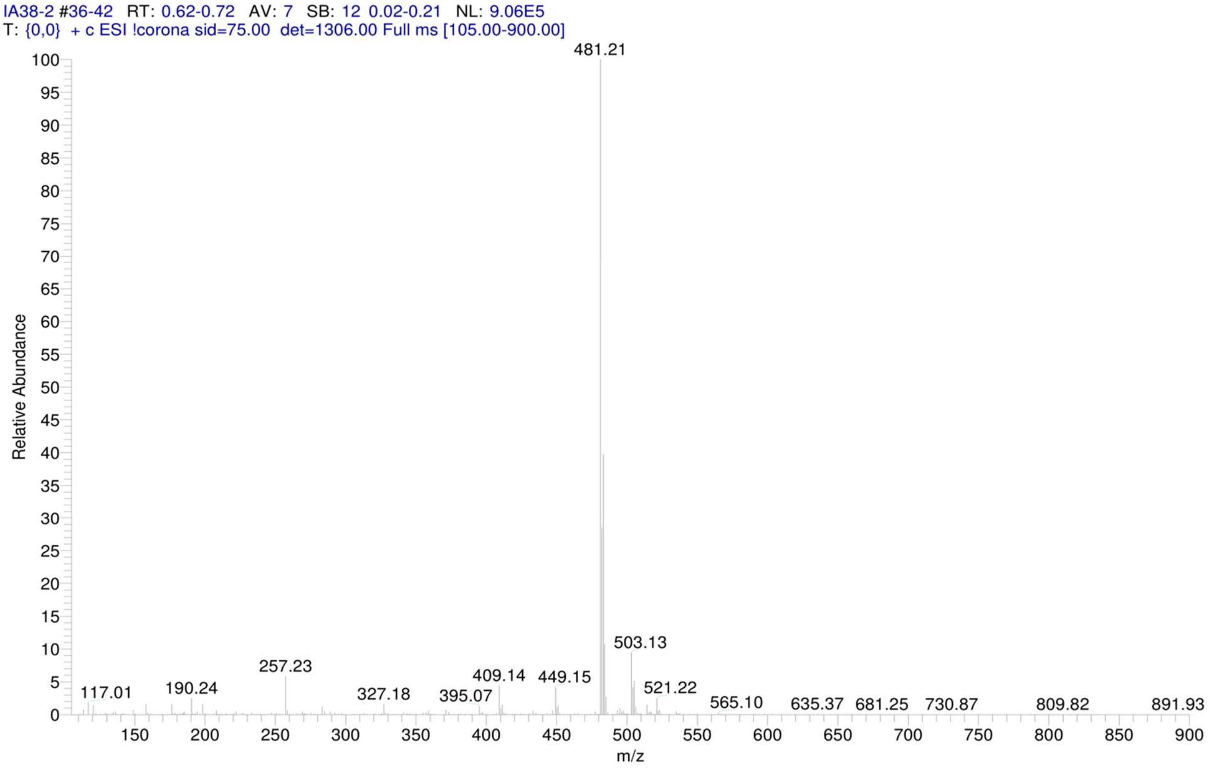

$^1$H NMR compound **46**:

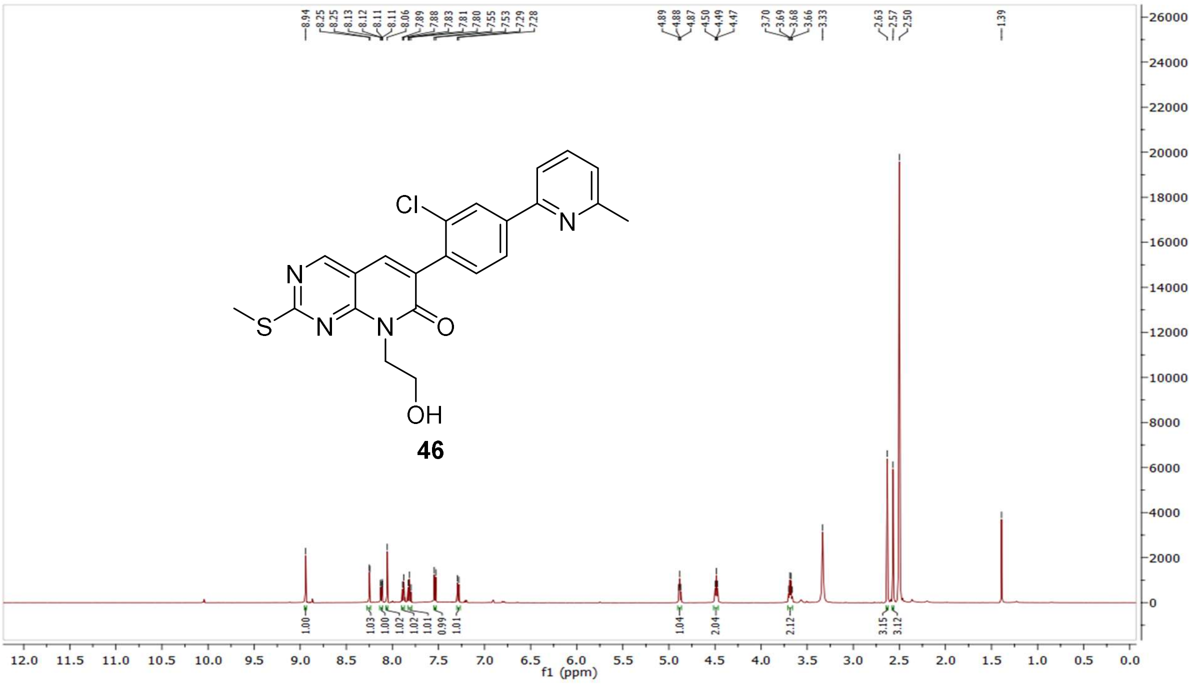

$^{13}$C NMR compound **46:**

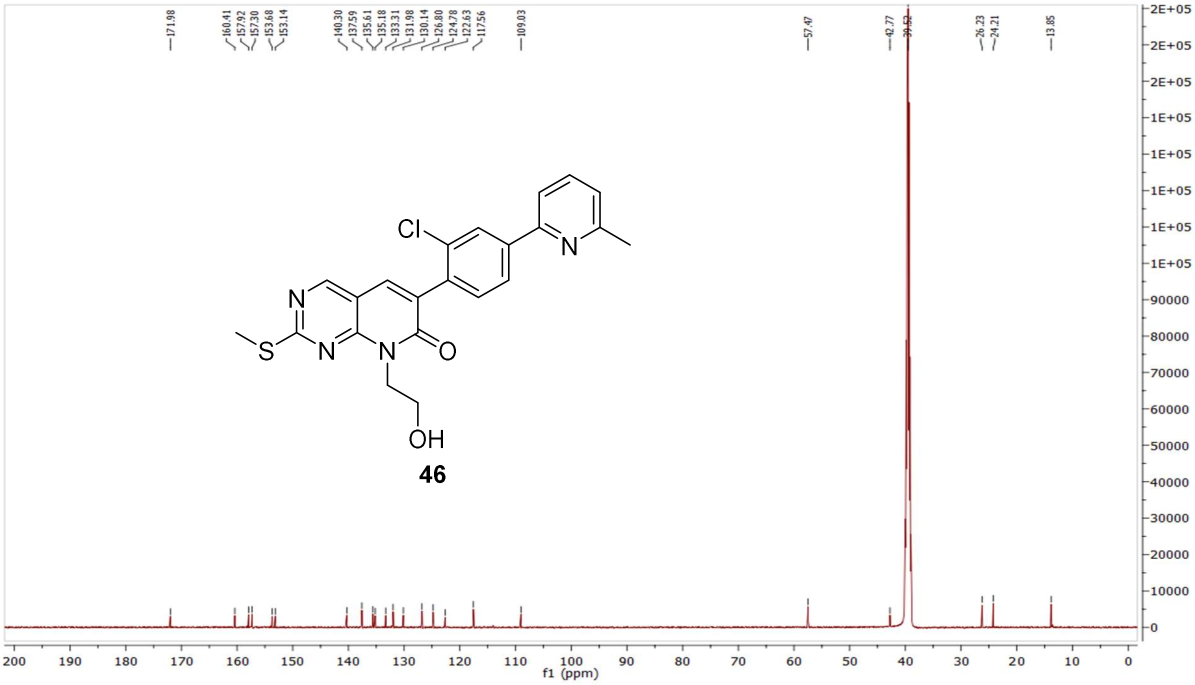

MS (ESI+) compound **46**:

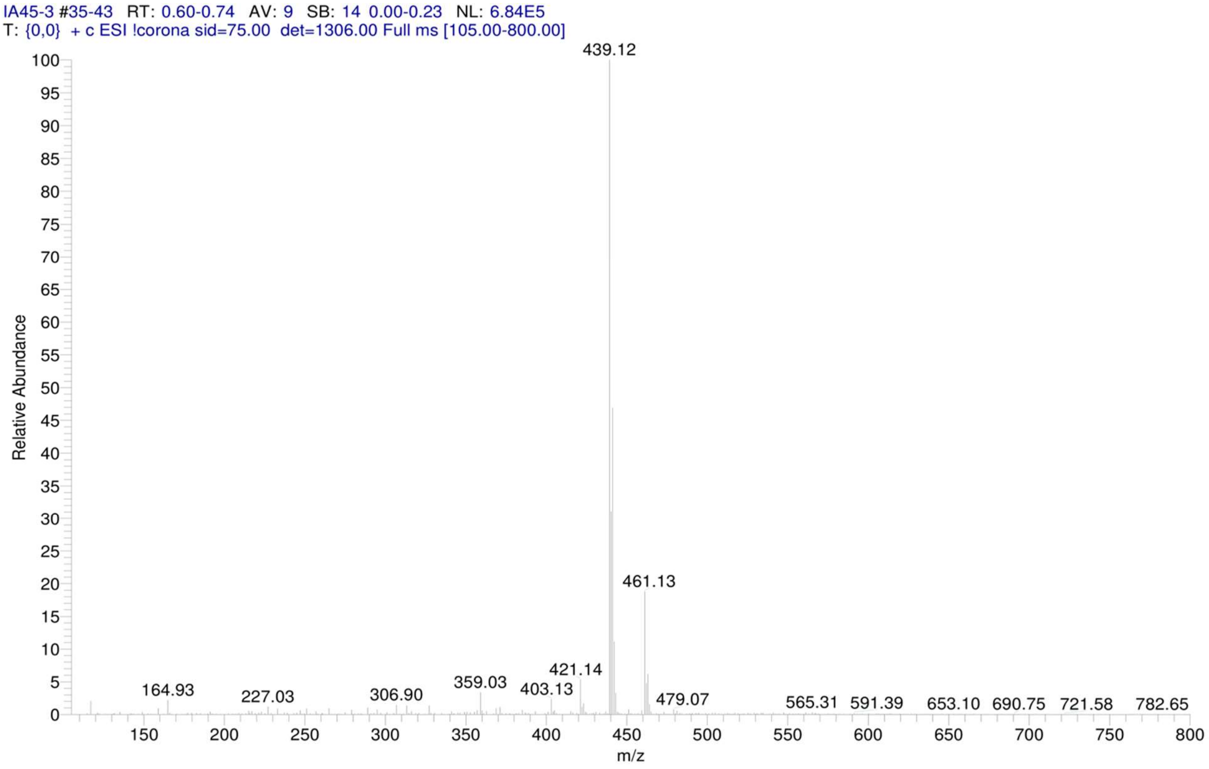


[1]H NMR compound **47**:

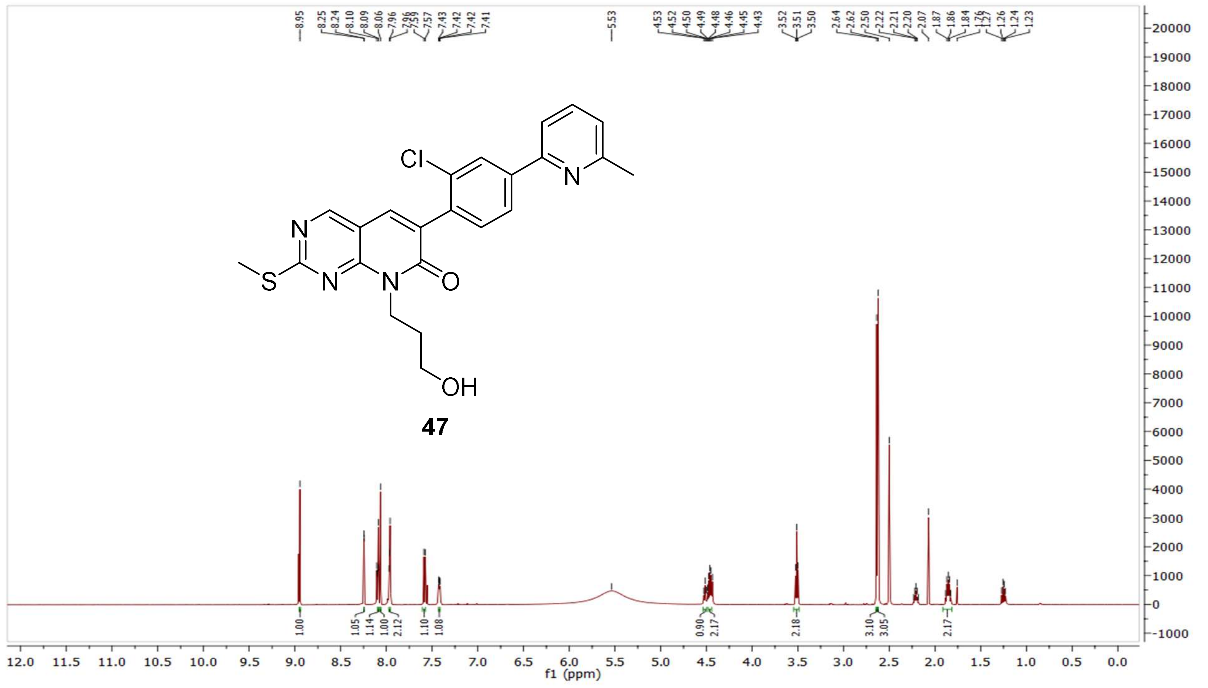

$^{13}$C NMR compound **47:**

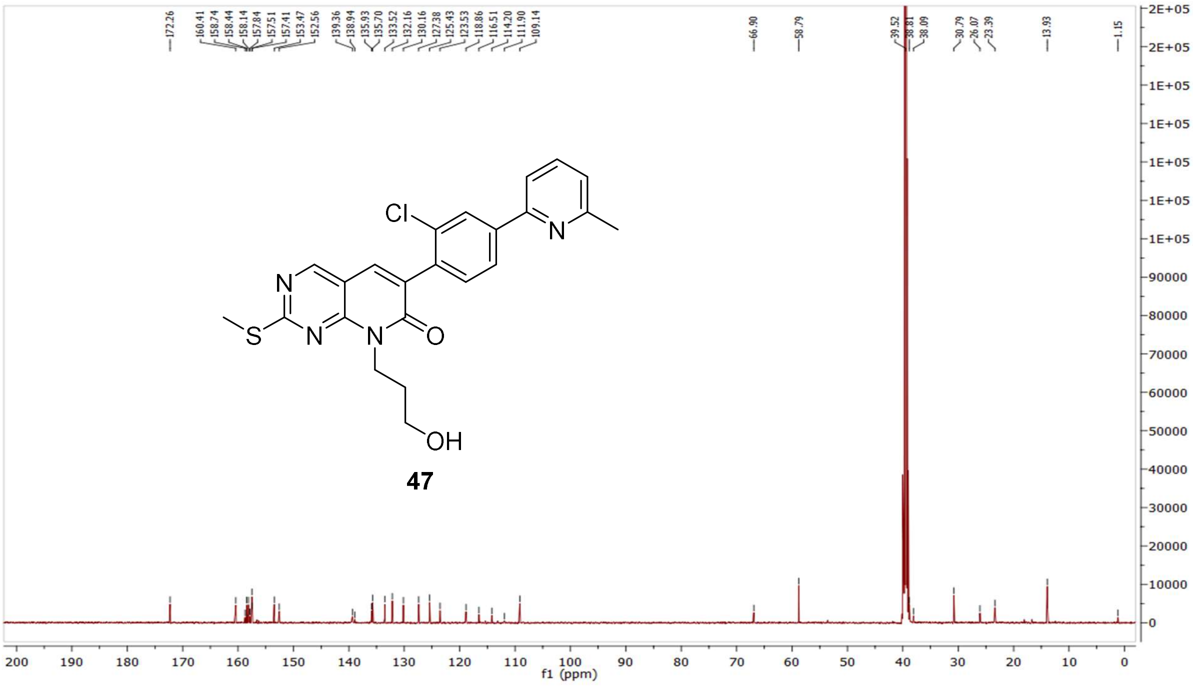


MS (ESI+) compound **47**:

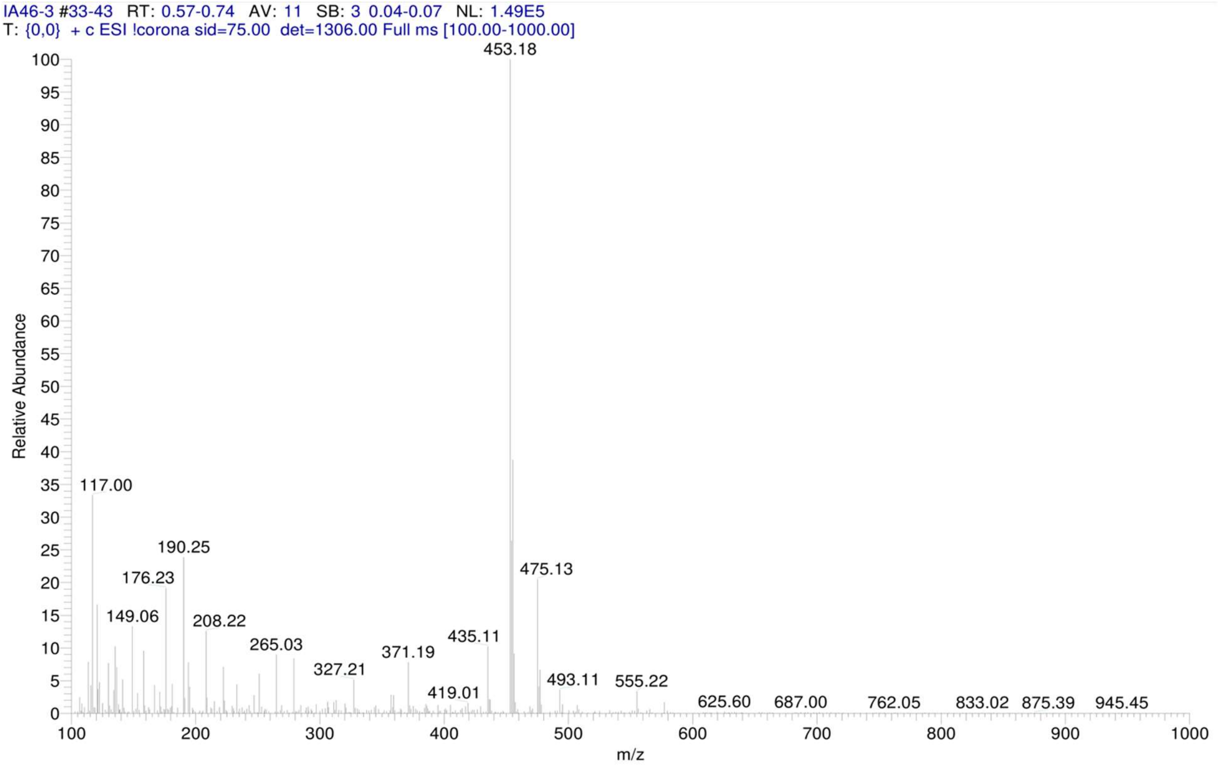

$^{1}H$ NMR compound **48**:

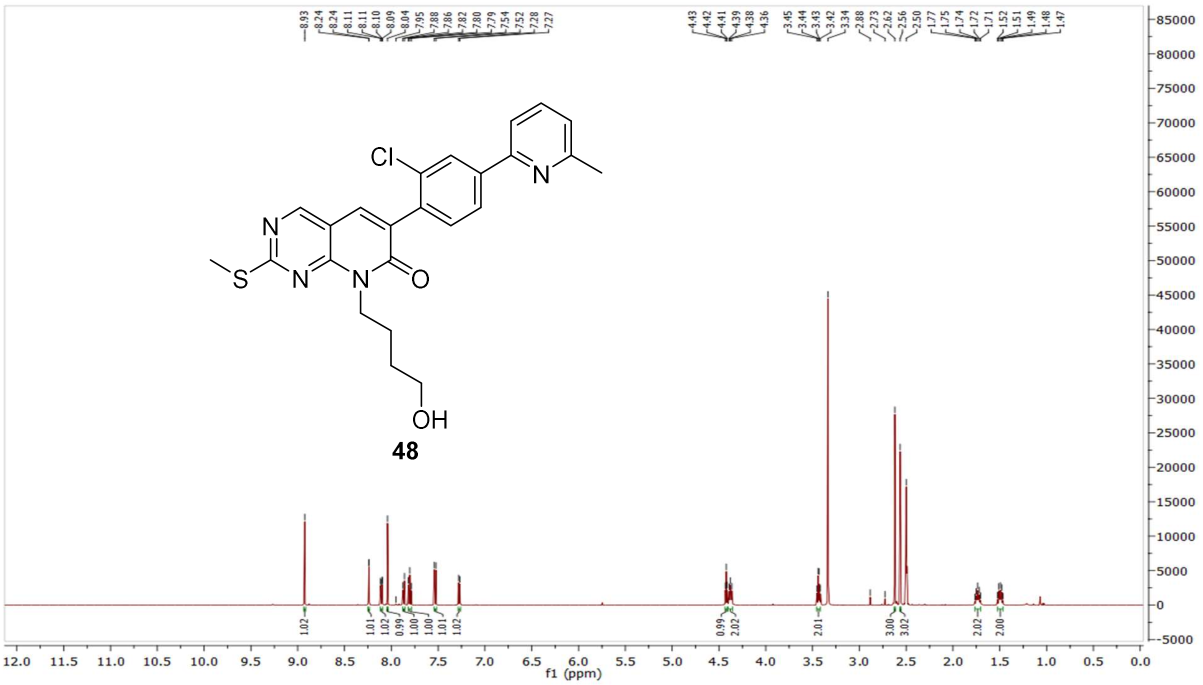


$^{13}C$ NMR compound **48:**

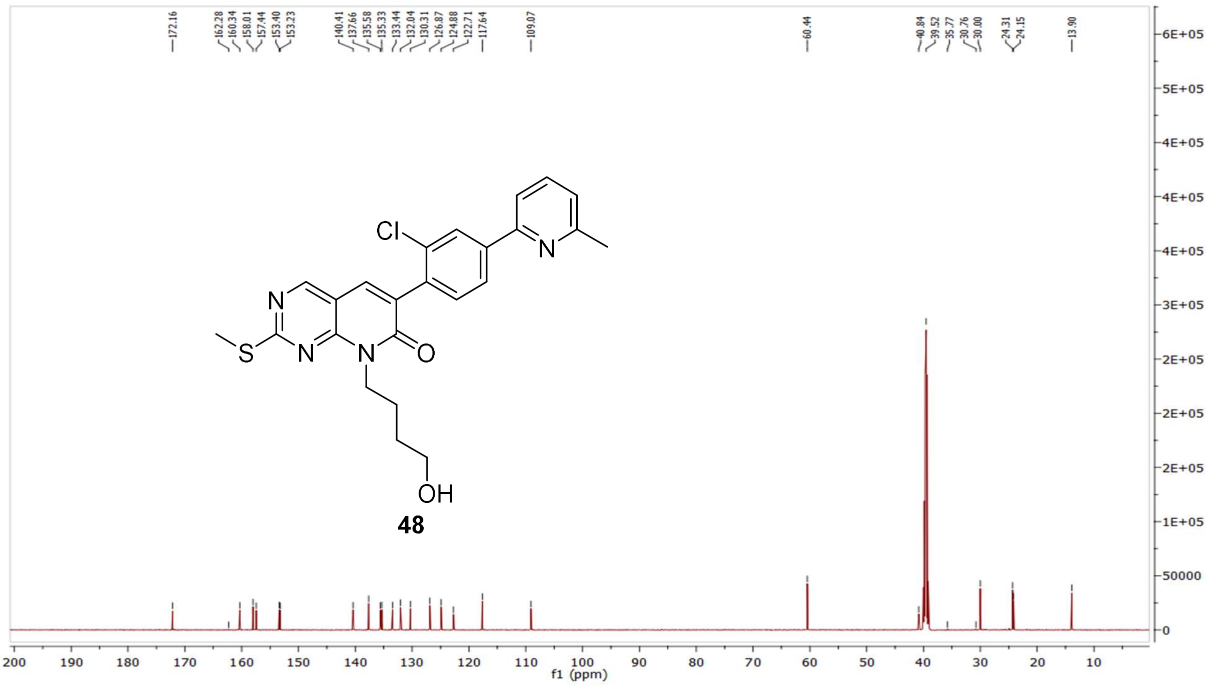

MS (ESI+) compound **48**:

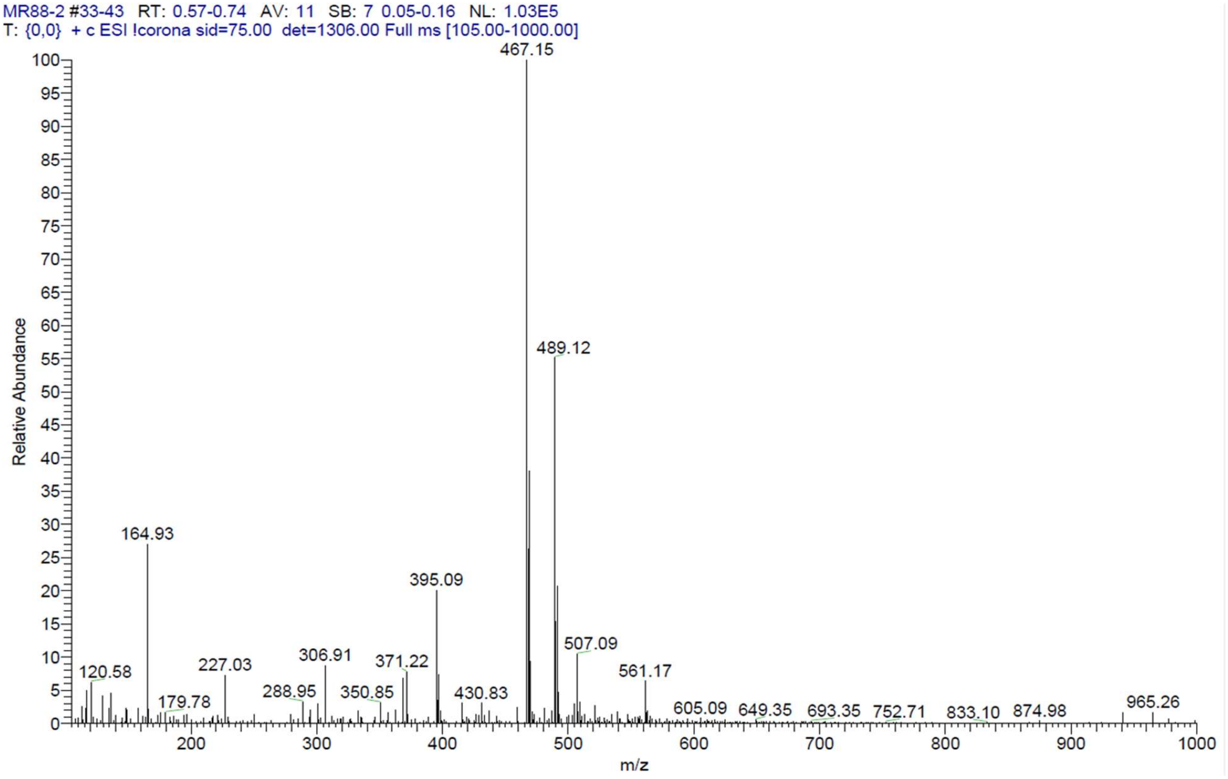


$^{1}$H NMR compound **49**:

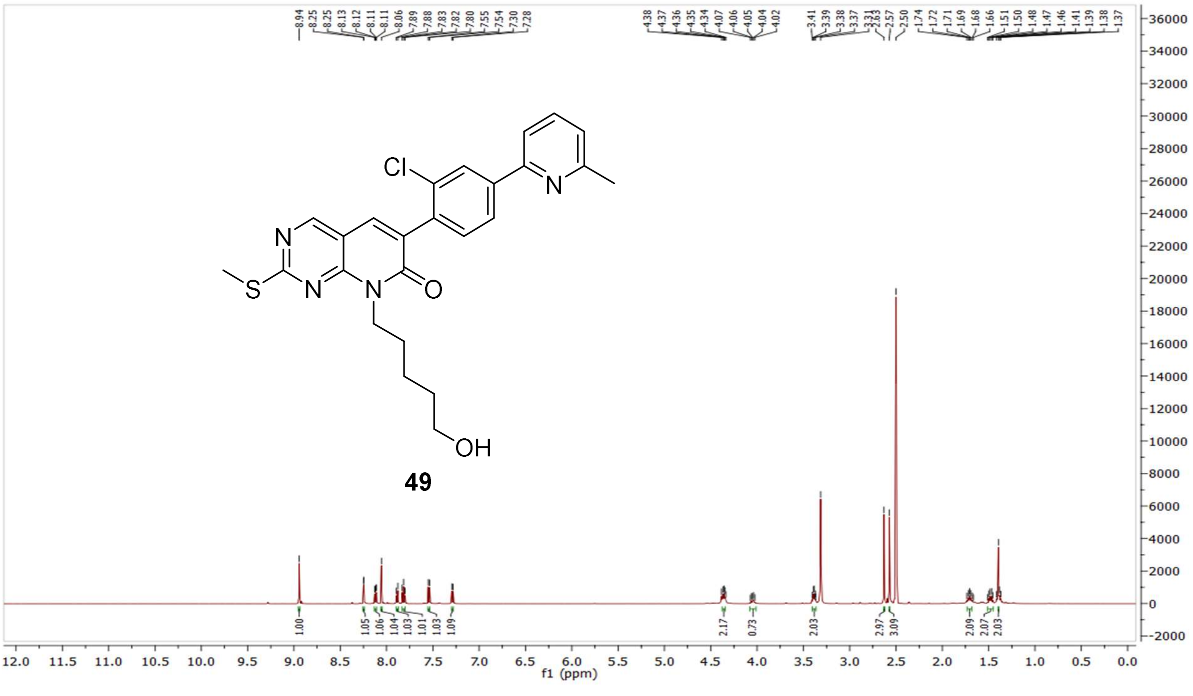

[13]C NMR compound **49:**

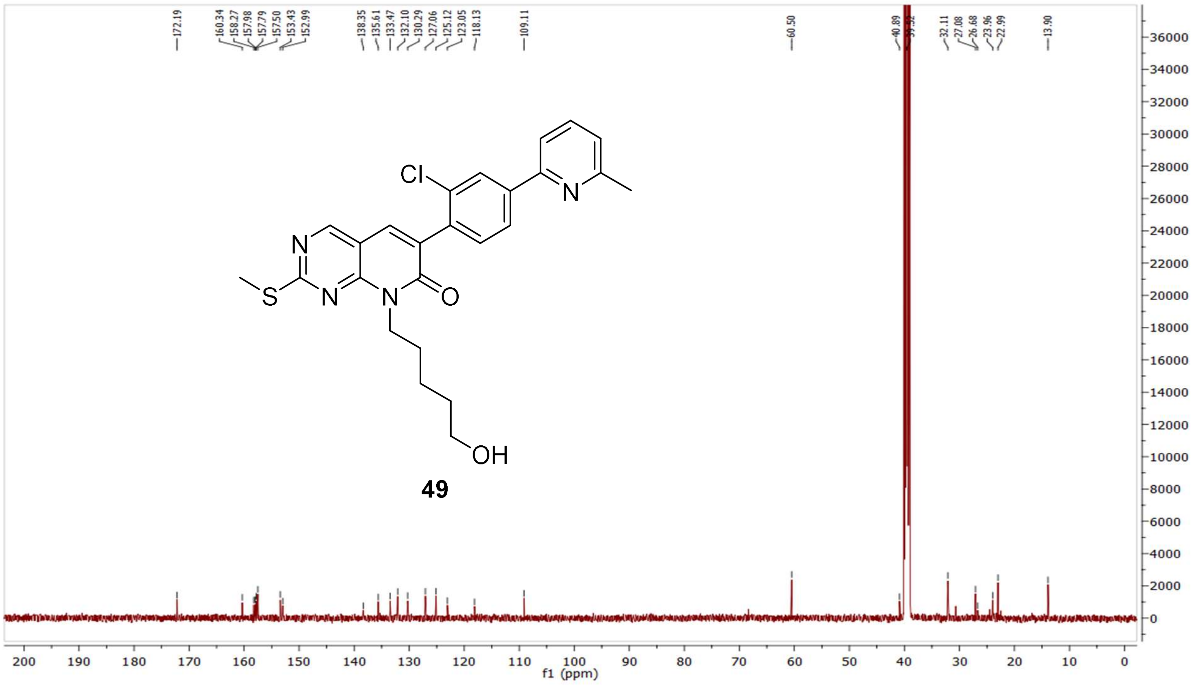


MS (ESI+) compound **49**:

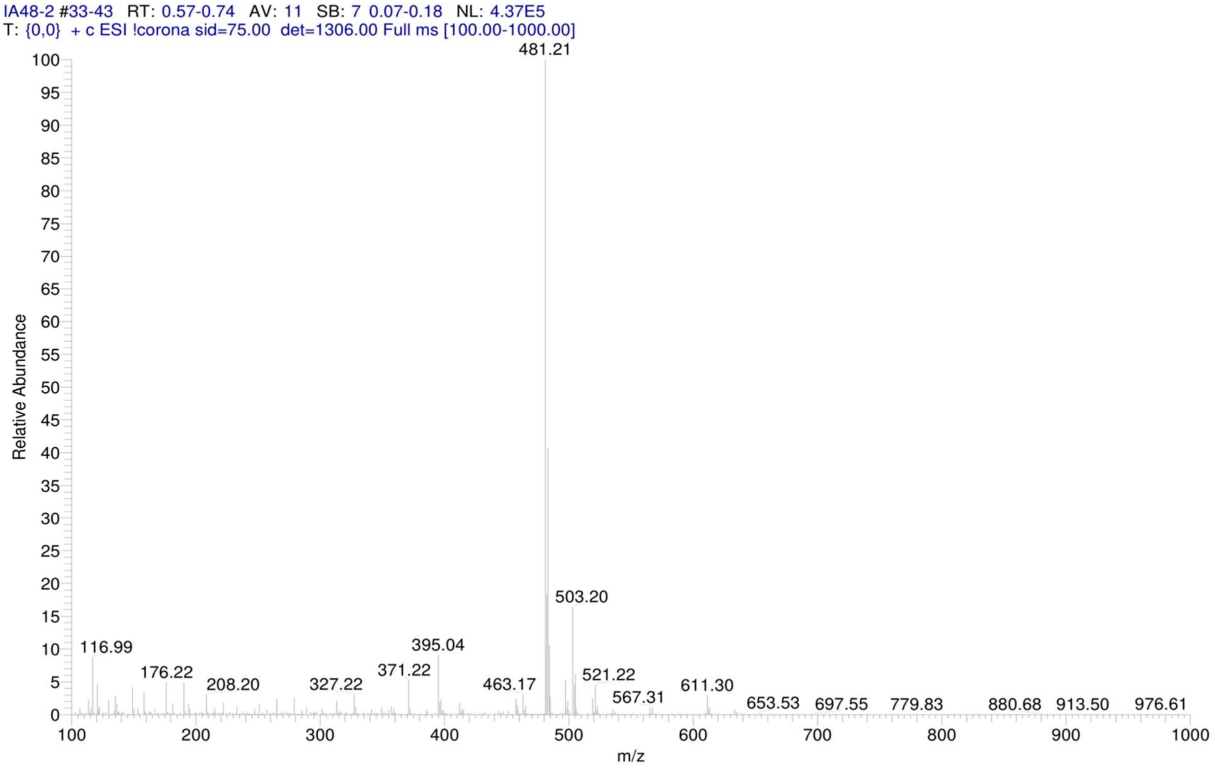

$^{1}$H NMR compound **9**:

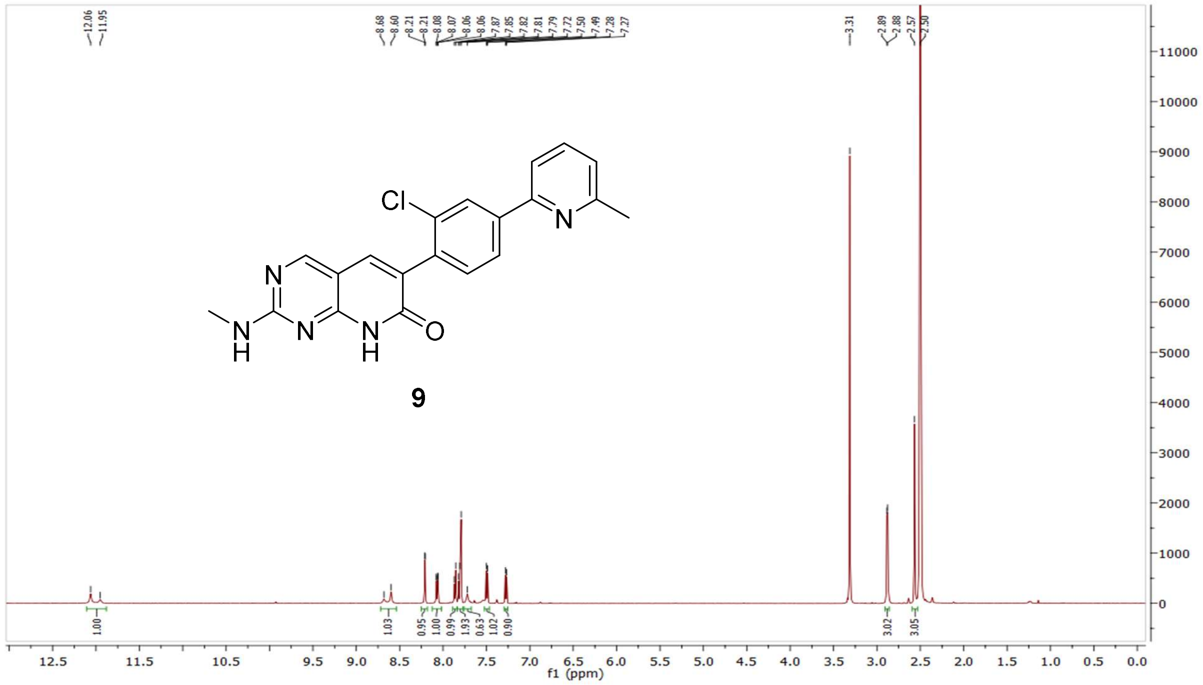


$^{13}$C NMR compound **9:**

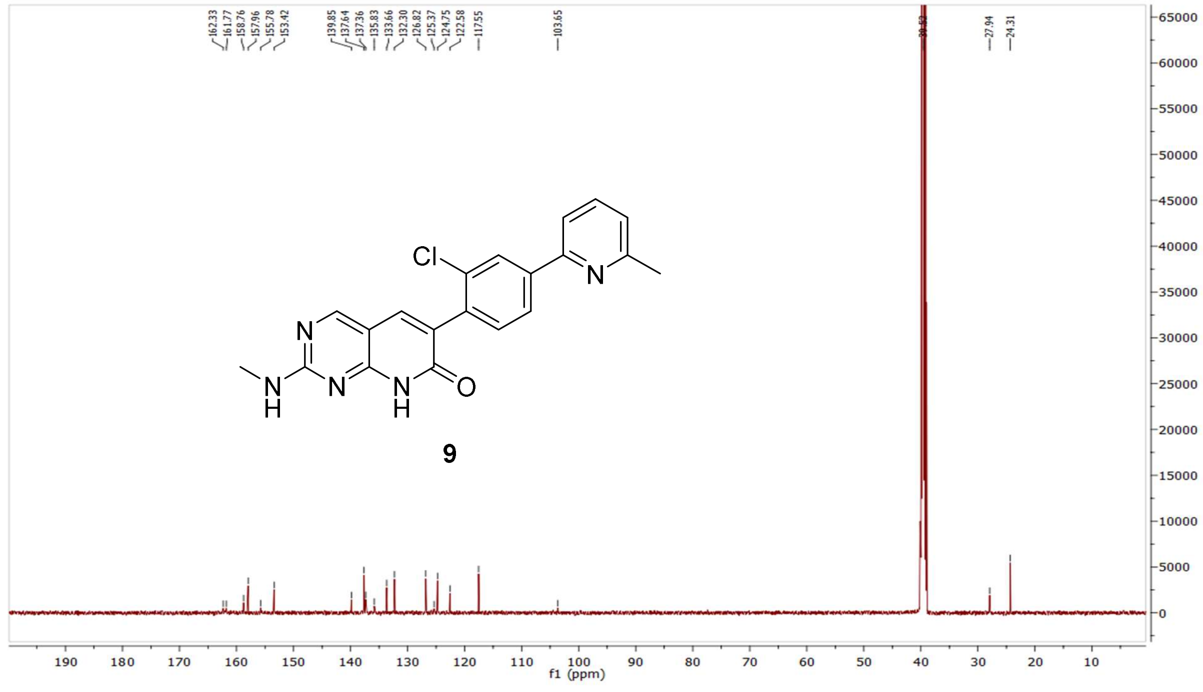

HRMS (FTMS +p MALDI) compound **9**:

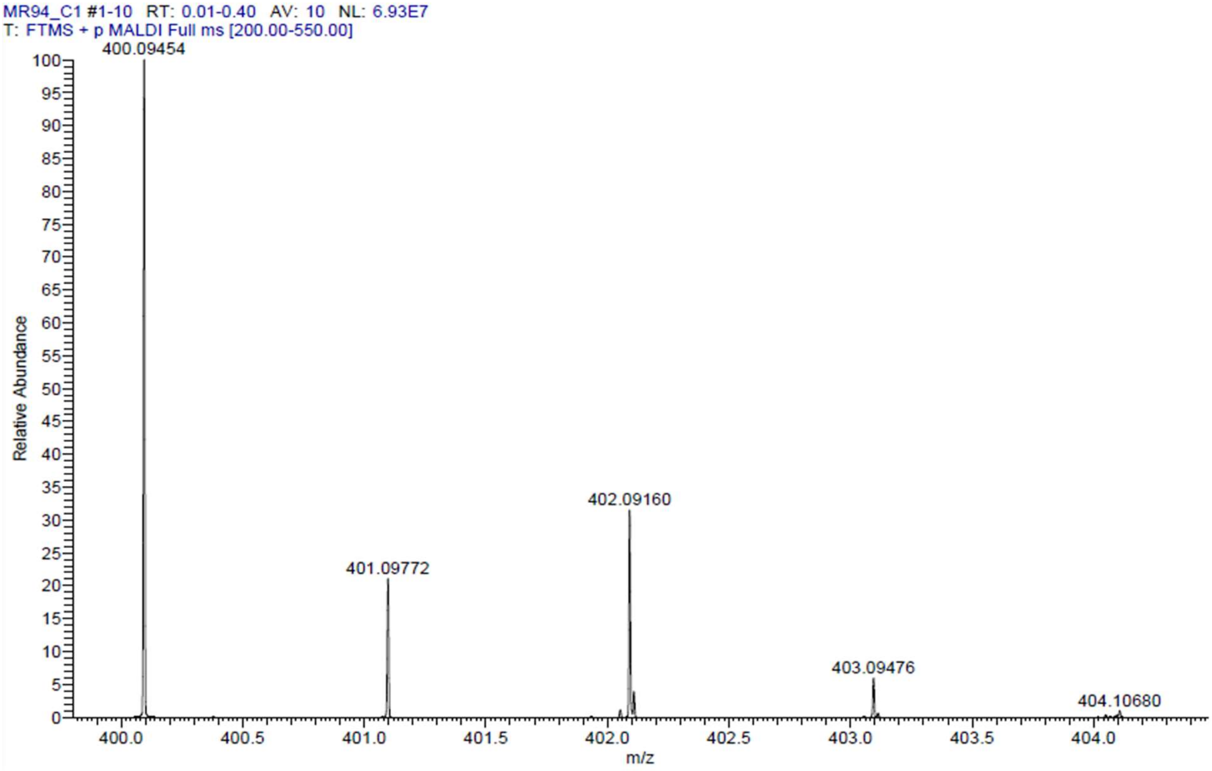

HPLC-chromatograms compound **9**:

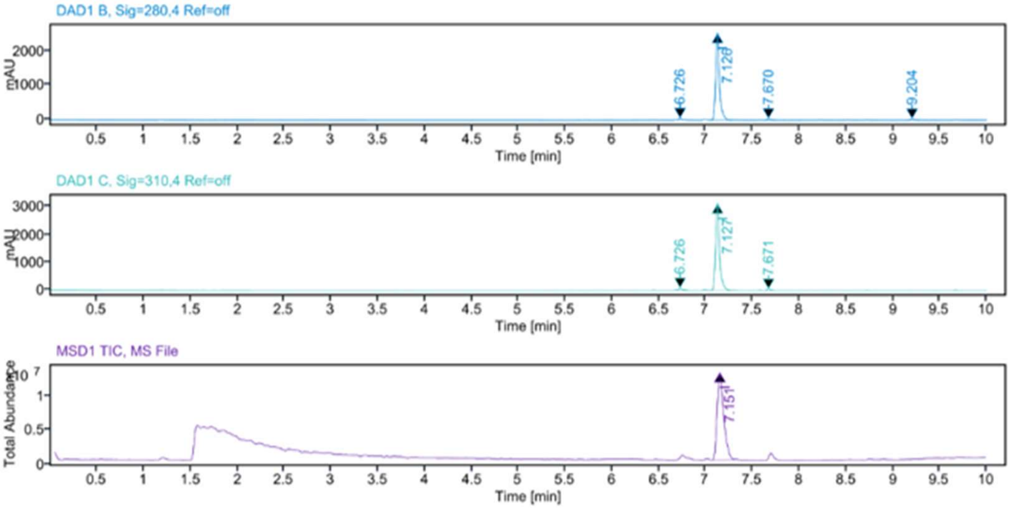


**Sample Purity**

| Signal Description | DAD1 B, Sig=280,4 Ref=off |
|---|---|

| Sample Name | Name | RT | Width | Area | Area% | Height |
|---|---|---|---|---|---|---|
| MR94 | | 6.726 | 0.028 | 117.8598 | 1.63 | 55.5862 |
| MR94 | | 7.126 | 0.038 | 6884.2095 | 95.07 | 2510.9246 |
| MR94 | | 7.670 | 0.036 | 120.2627 | 1.66 | 49.7059 |
| MR94 | | 9.204 | 0.043 | 118.5366 | 1.64 | 41.7601 |

| | |
|---|---|
| Max Area% | 95.074 |
| UV Signal Purity>95% | Pass |

| Signal Description | DAD1 C, Sig=310,4 Ref=off |
|---|---|

| Sample Name | Name | RT | Width | Area | Area% | Height |
|---|---|---|---|---|---|---|
| MR94 | | 6.726 | 0.029 | 173.9287 | 1.86 | 80.7885 |
| MR94 | | 7.127 | 0.040 | 9026.2363 | 96.64 | 3118.7725 |
| MR94 | | 7.671 | 0.036 | 139.6326 | 1.50 | 55.0632 |

| | |
|---|---|
| Max Area% | 96.643 |
| UV Signal Purity>95% | Pass |

**MSD Apex Spectra**

**RT:**

7.151

**Sample Name:**

MR94

**Signal Name:**

MSD1TIC

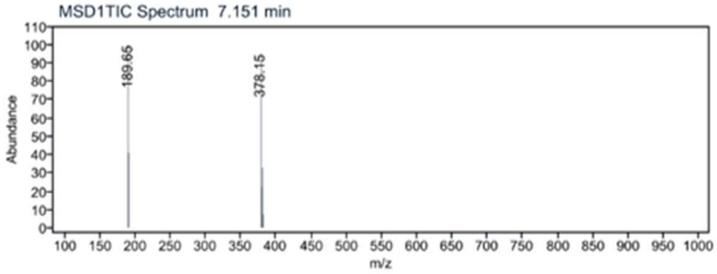

$^1$H NMR compound **10**:

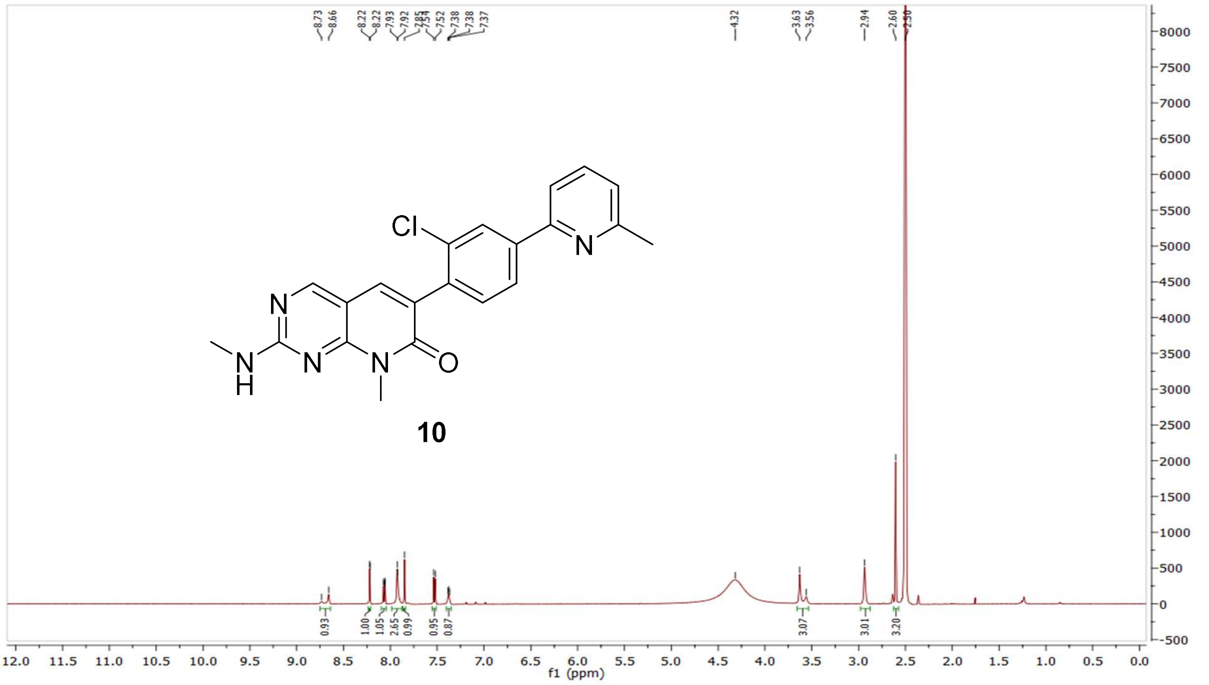


$^{13}$C NMR compound **10:**

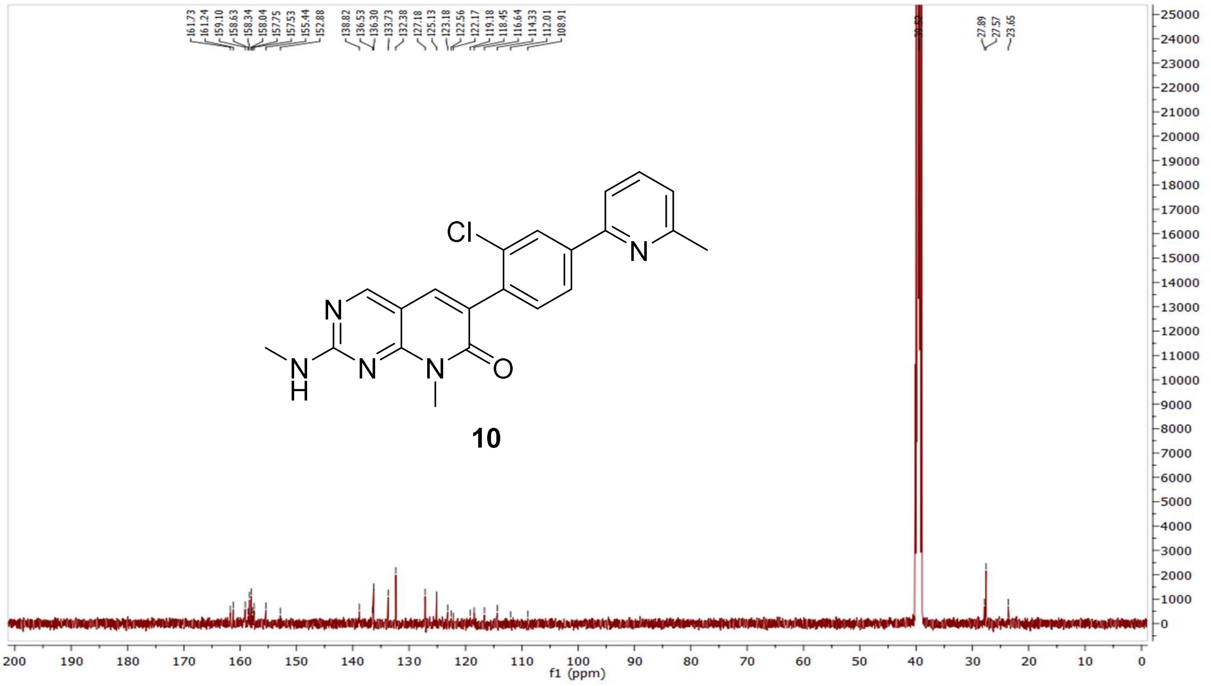

HRMS (FTMS +p MALDI) compound **10**:

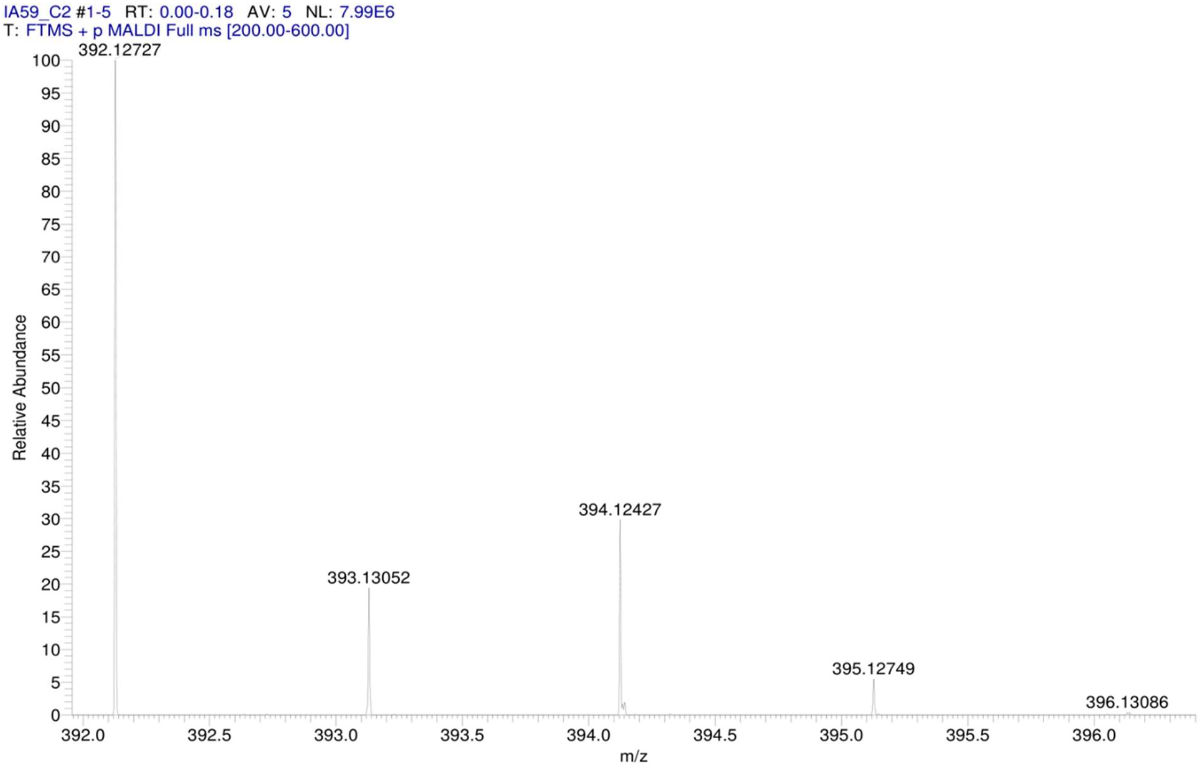


HPLC-chromatograms compound **10**:

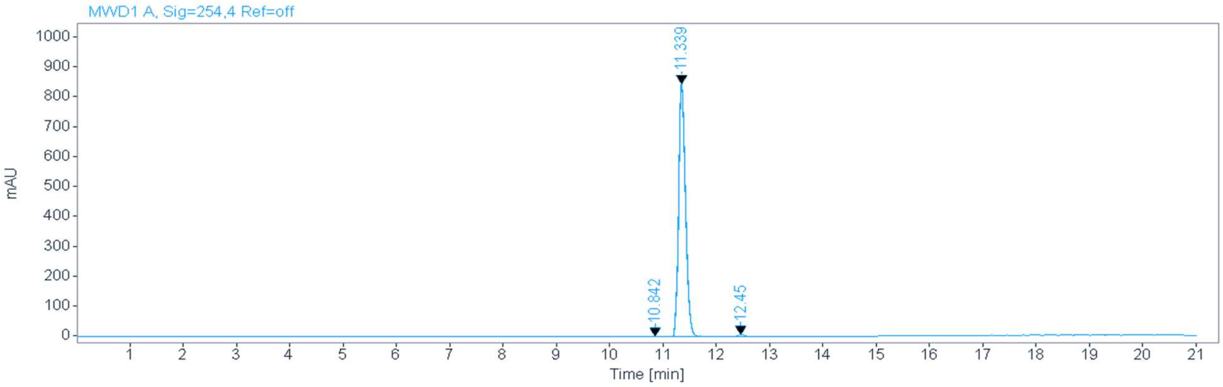


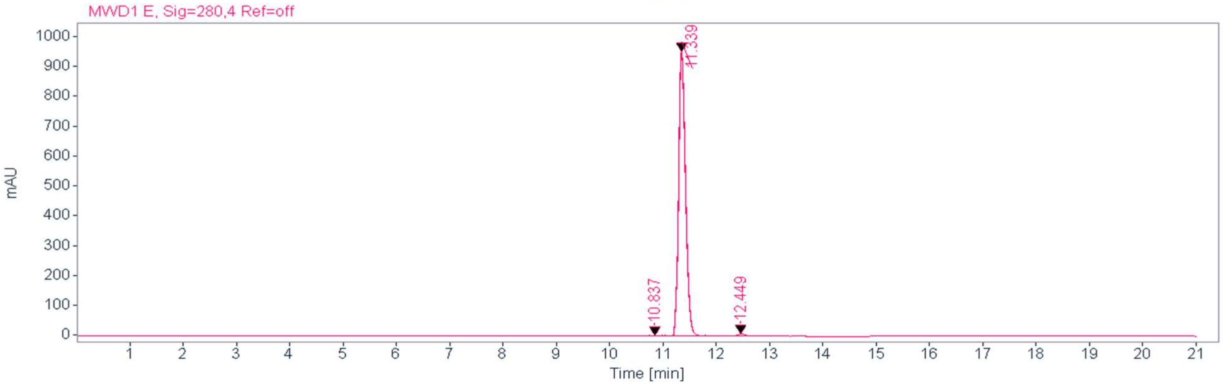

**Signal:** MWD1 A, Sig=254,4 Ref=off

| RT [min] | Type | Width [min] | Area | Height | Area% | Name |
|---|---|---|---|---|---|---|
| 10.842 | MM | 0.1375 | 10.4549 | 1.2672 | 0.1367 | |
| 11.339 | VV | 0.1377 | 578.6533 | 844.8054 | 99.0761 | |
| 12.450 | MM | 0.1529 | 60.2177 | 6.5655 | 0.7872 | |
| | | Sum | 7649.3259 | | | |

**Signal:** MWD1 E, Sig=280,4 Ref=off

| RT [min] | Type | Width [min] | Area | Height | Area% | Name |
|---|---|---|---|---|---|---|
| 10.837 | MM | 0.1666 | 16.4529 | 1.6464 | 0.1893 | |
| 11.339 | VV | 0.1382 | 8587.9229 | 952.6332 | 98.7932 | |
| 12.449 | MM | 0.1928 | 88.4481 | 7.6442 | 1.0175 | |
| | | Sum | 8692.8239 | | | |

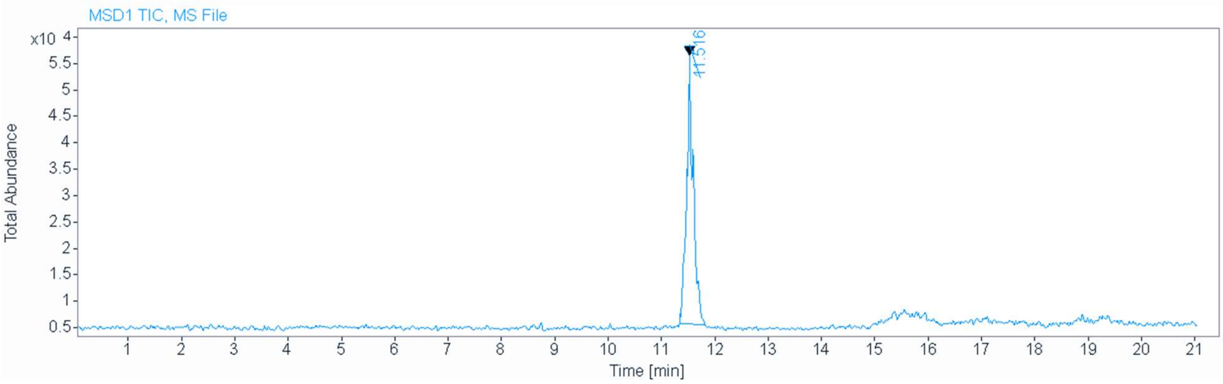

MS (ESI+) compound **10**:

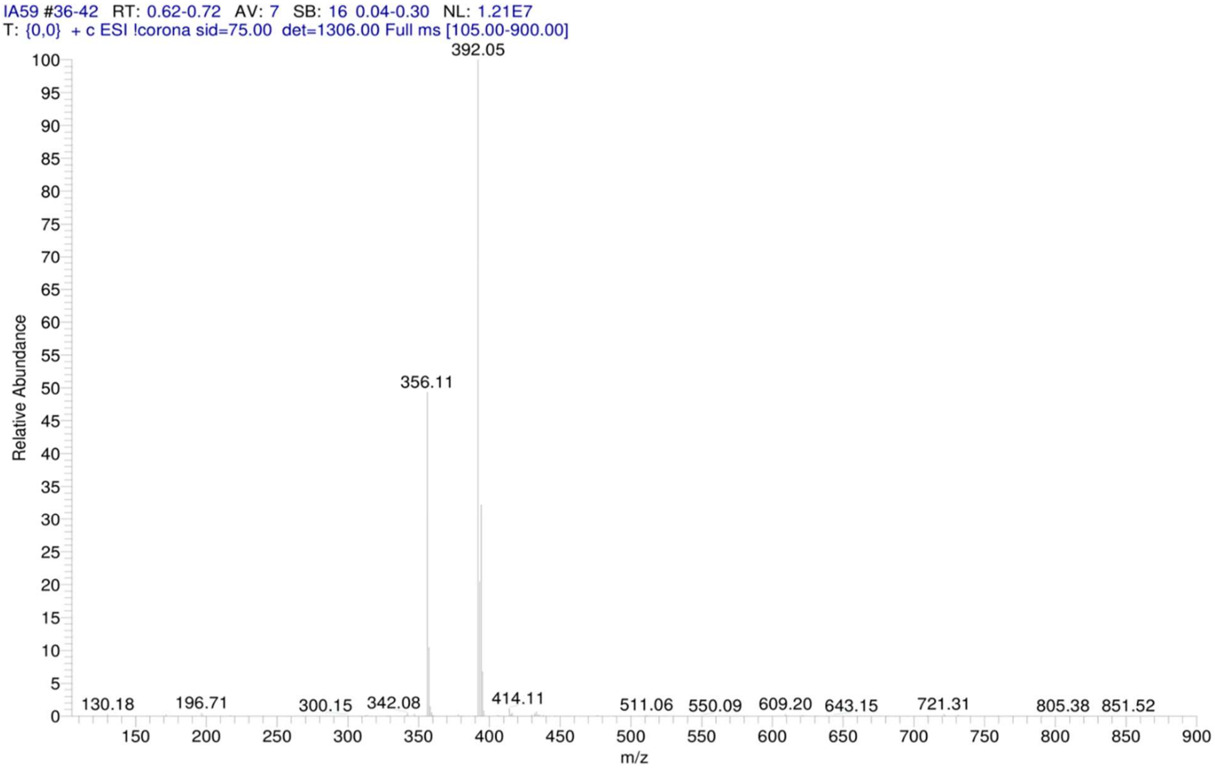


$^1$H NMR compound **11**:

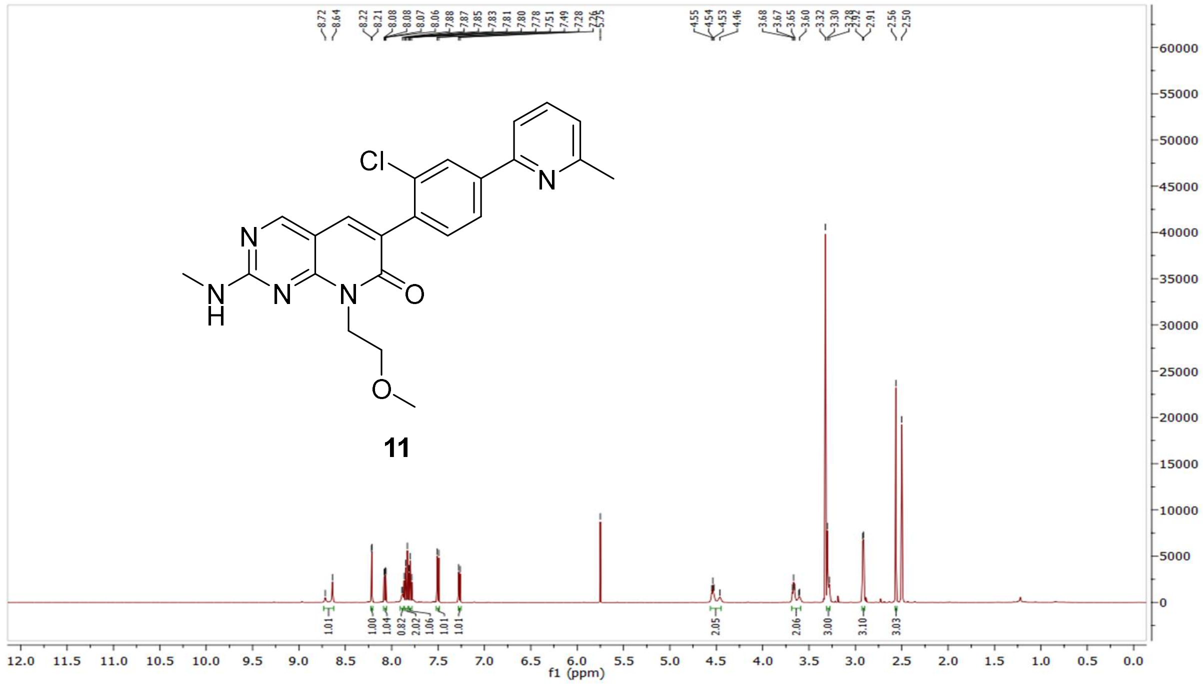

$^{13}C$ NMR compound **11:**

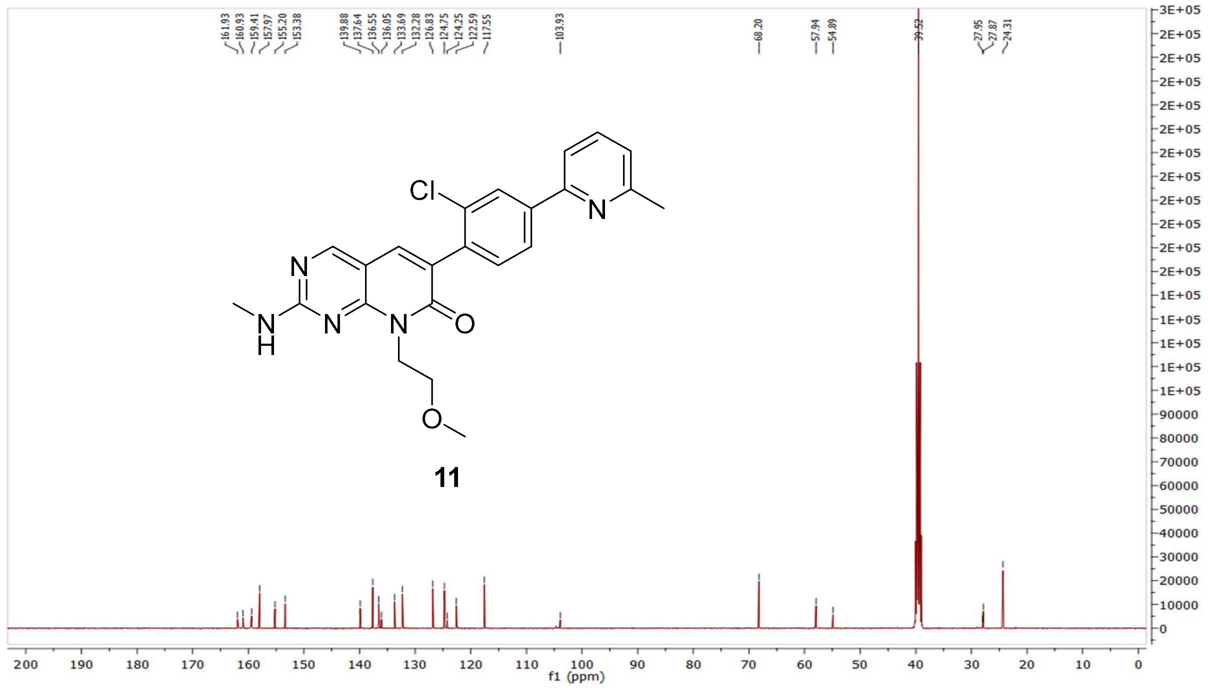

HRMS (FTMS +p MALDI) compound **11**:

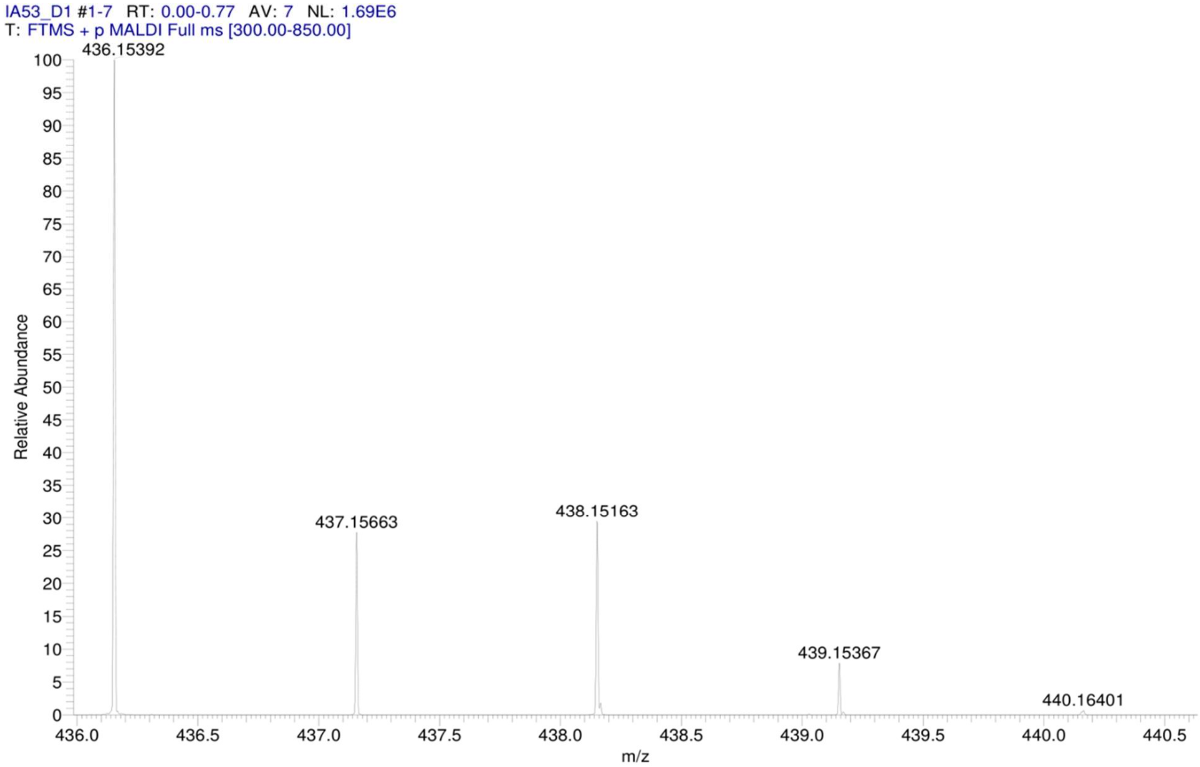

HPLC-chromatograms compound **11**:

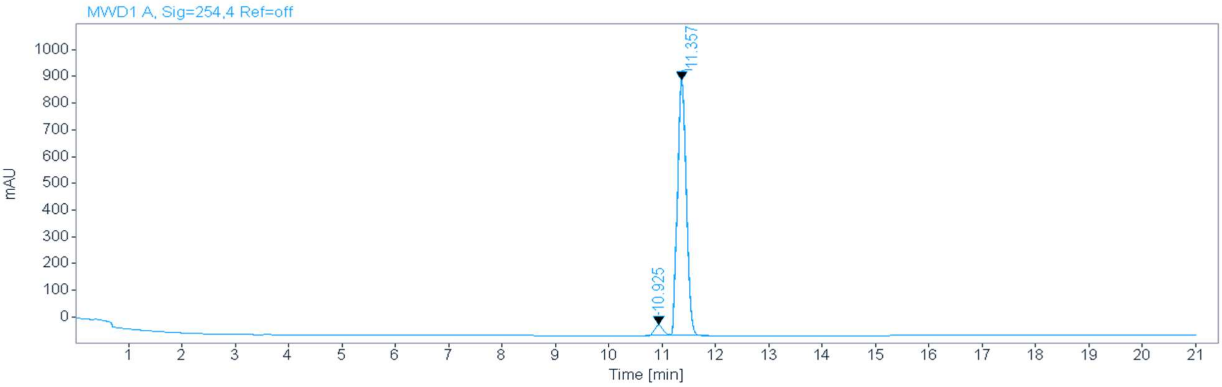


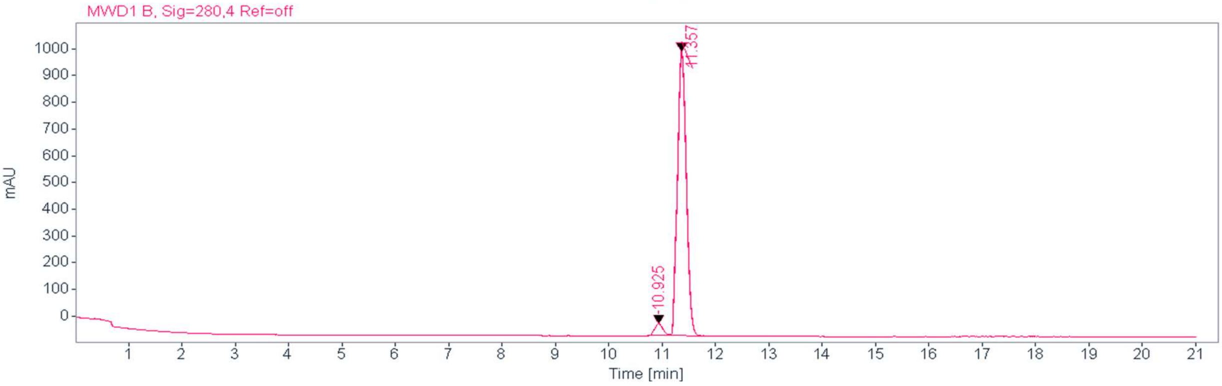


**Signal:** MWD1 A, Sig=254,4 Ref=off

| RT [min] | Type | Width [min] | Area | Height | Area% | Name |
|---|---|---|---|---|---|---|
| 10.925 | MM | 0.1839 | 450.3305 | 40.8190 | 3.8842 | |
| 11.357 | VV | 0.1780 | 11143.6914 | 955.7079 | 96.1158 | |
| | | Sum | 11594.0219 | | | |

**Signal:** MWD1 B, Sig=280,4 Ref=off

| RT [min] | Type | Width [min] | Area | Height | Area% | Name |
|---|---|---|---|---|---|---|
| 10.925 | MM | 0.1800 | 485.2560 | 44.9289 | 3.7629 | |
| 11.357 | VV | 0.1781 | 12410.6006 | 1063.9673 | 96.2371 | |
| | | Sum | 12895.8565 | | | |

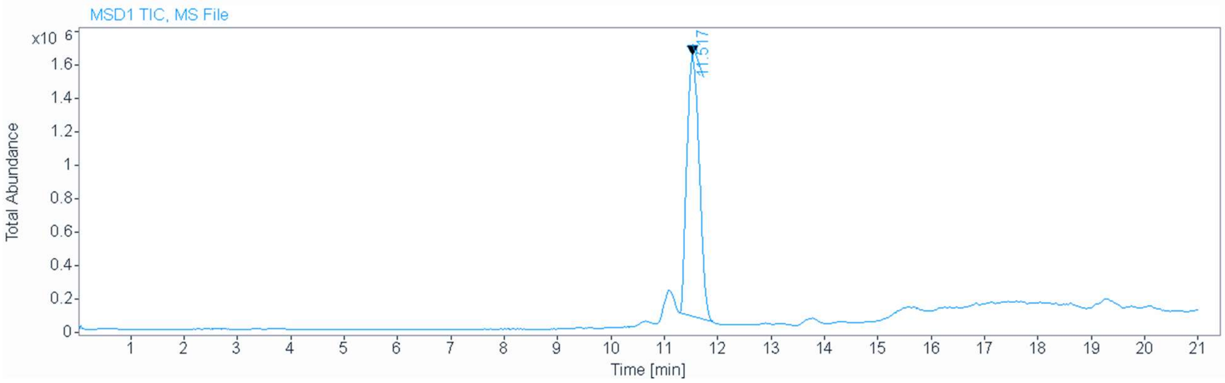


**Signal** MSD1 TIC, MS File

**Peak RT** 11.517

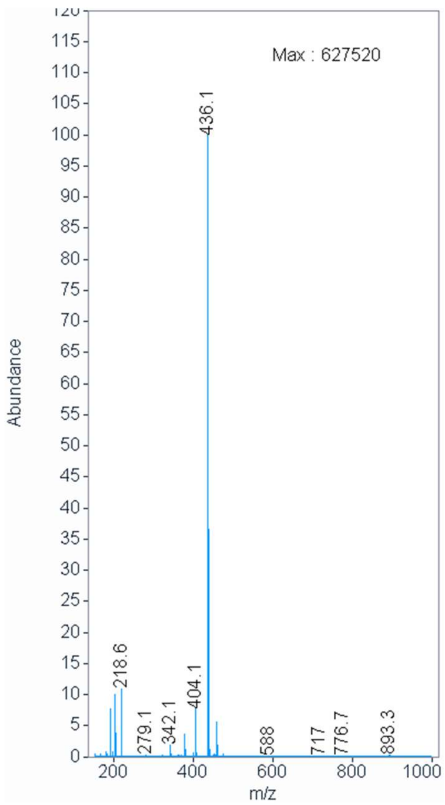

<sup></sup>

$^{1}$H NMR compound **12**:

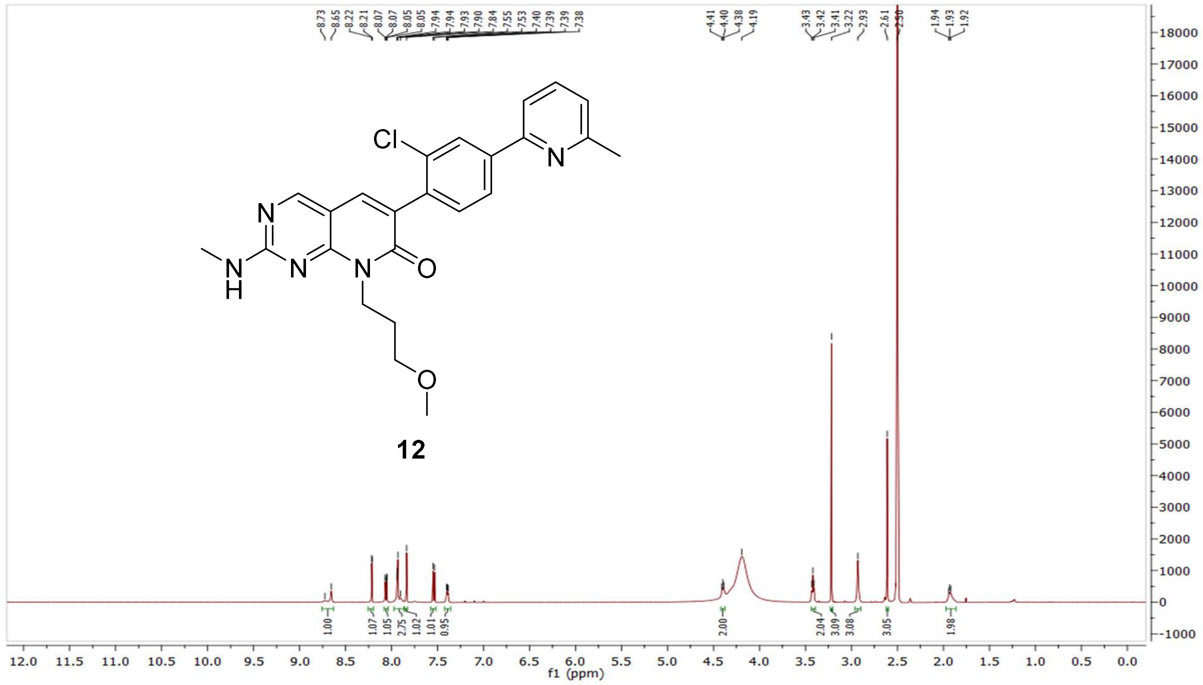

$^{13}$C NMR compound **12:**

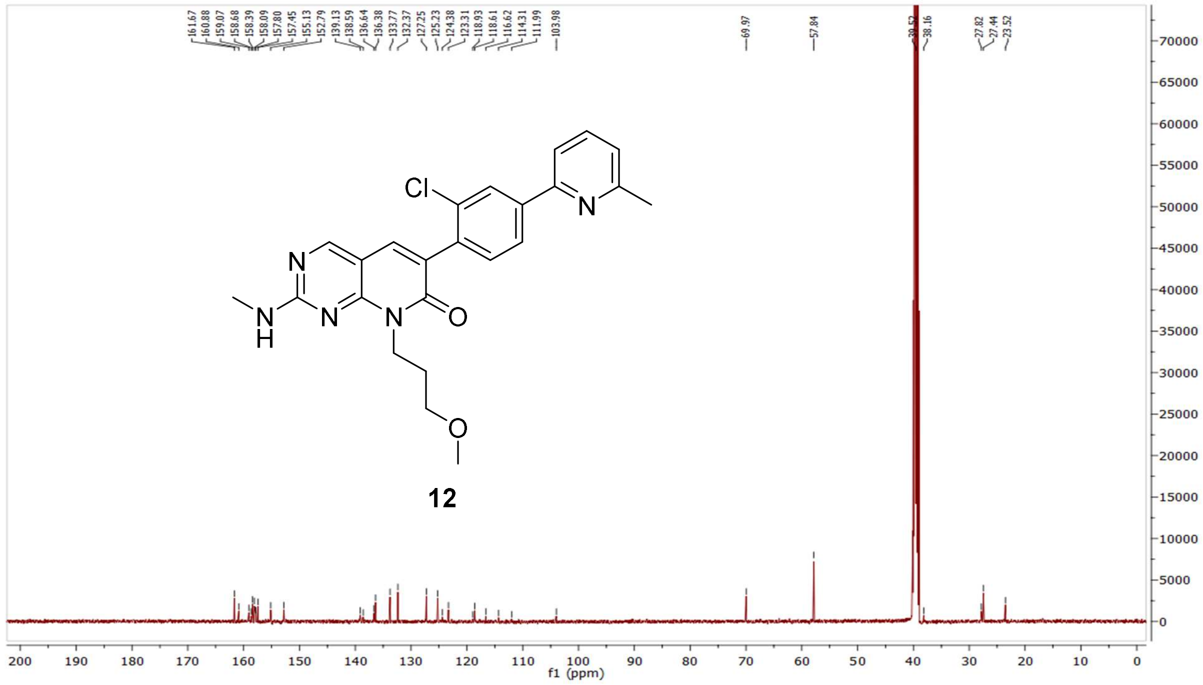

HRMS (FTMS +p MALDI) compound **12**:

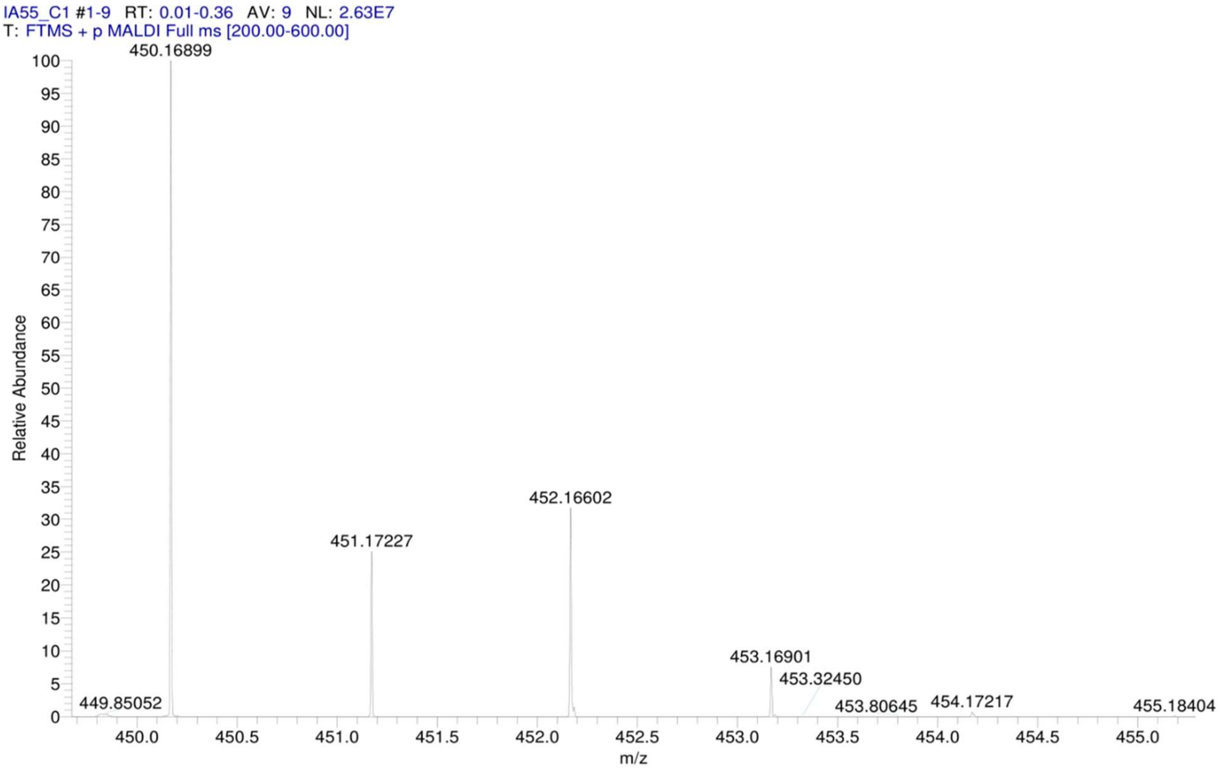


HPLC-chromatograms compound **12**:

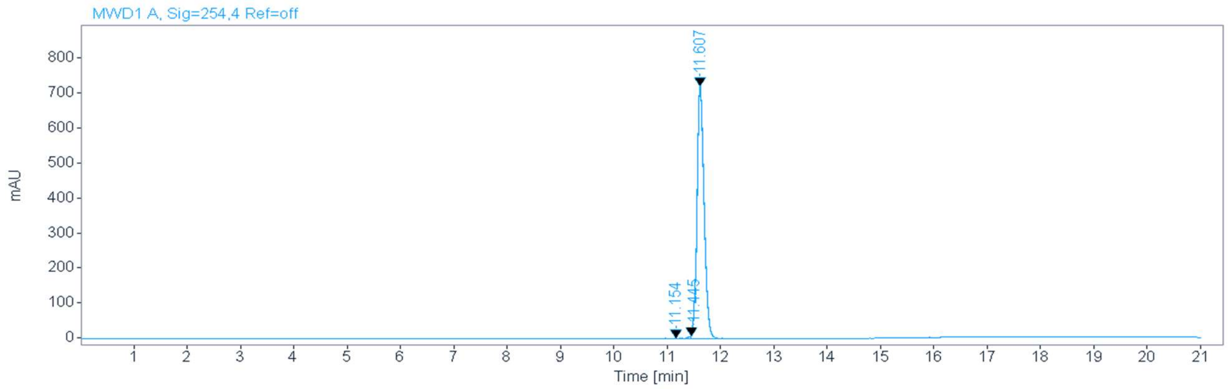


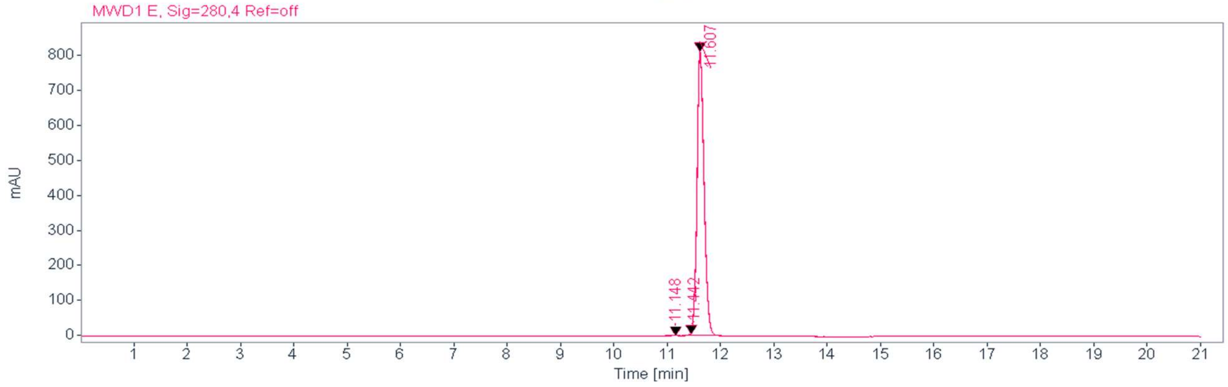

**Signal:** MWD1 A, Sig=254,4 Ref=off

| RT [min] | Type | Width [min] | Area | Height | Area% | Name |
|---|---|---|---|---|---|---|
| 11.154 | MM | 0.1880 | 13.5985 | 1.2053 | 0.2028 | |
| 11.445 | MM | 0.0746 | 32.9064 | 7.3566 | 0.4907 | |
| 11.607 | MM | 0.1538 | 6660.0605 | 721.8199 | 99.3066 | |
| | | Sum | 6706.5655 | | | |

**Signal:** MWD1 E, Sig=280,4 Ref=off

| RT [min] | Type | Width [min] | Area | Height | Area% | Name |
|---|---|---|---|---|---|---|
| 11.148 | MM | 0.1688 | 10.7099 | 1.0572 | 0.1427 | |
| 11.442 | MM | 0.0669 | 22.5312 | 5.6115 | 0.3001 | |
| 11.607 | MM | 0.1532 | 7473.8892 | 813.2397 | 99.5572 | |
| | | Sum | 7507.1302 | | | |

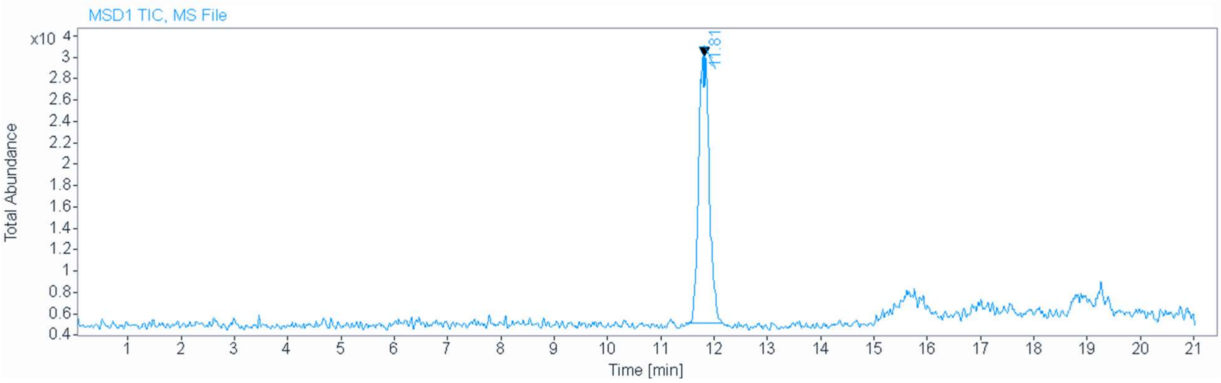

MS (ESI+) compound **12**:

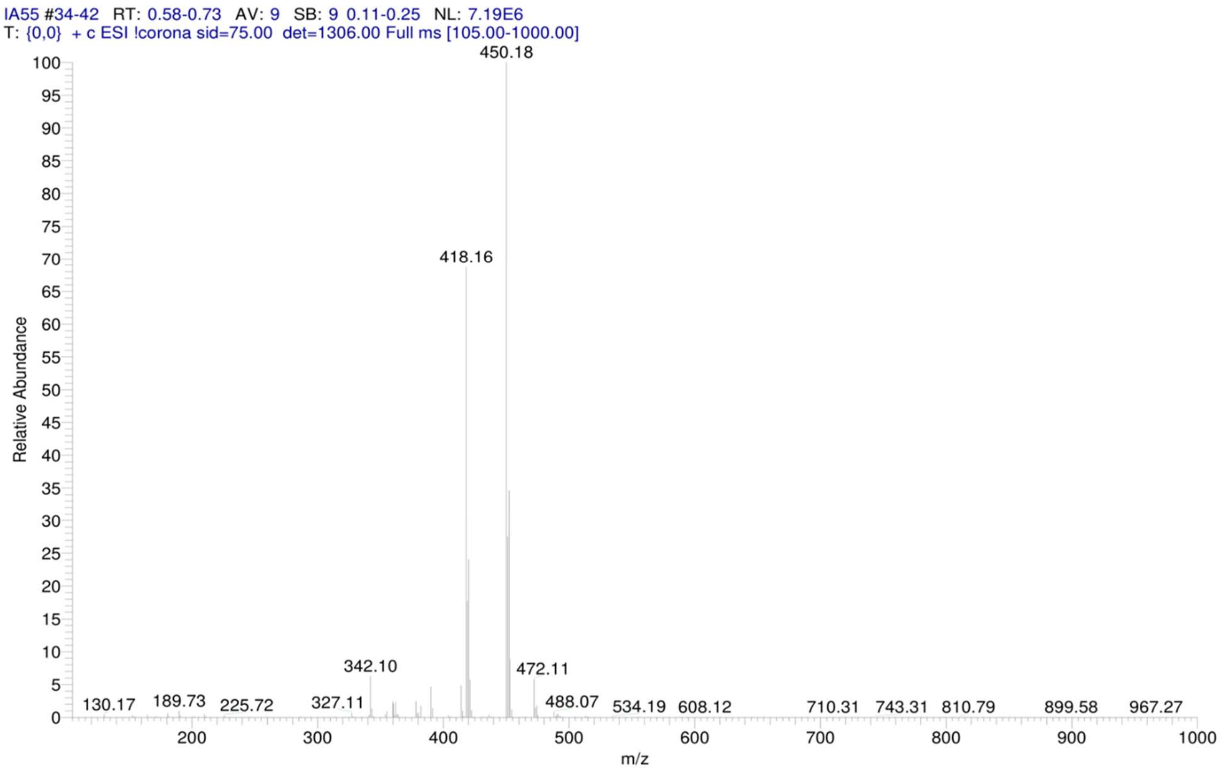


[1]H NMR compound **13**:

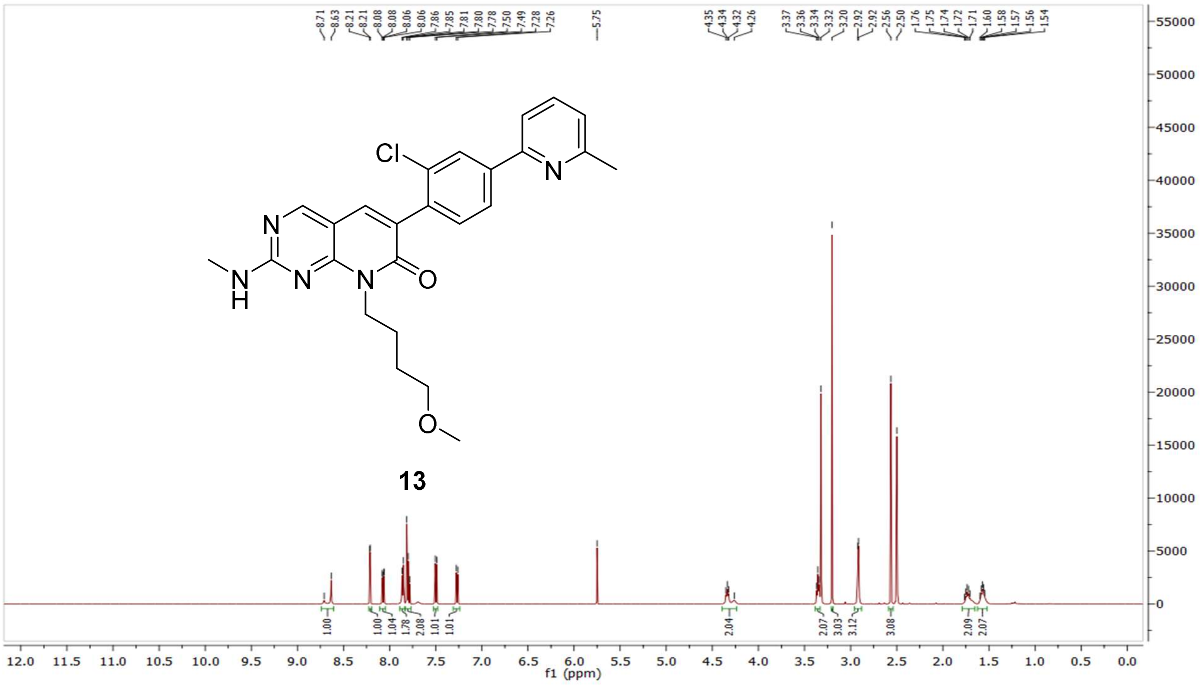

$^{13}$C NMR compound **13:**

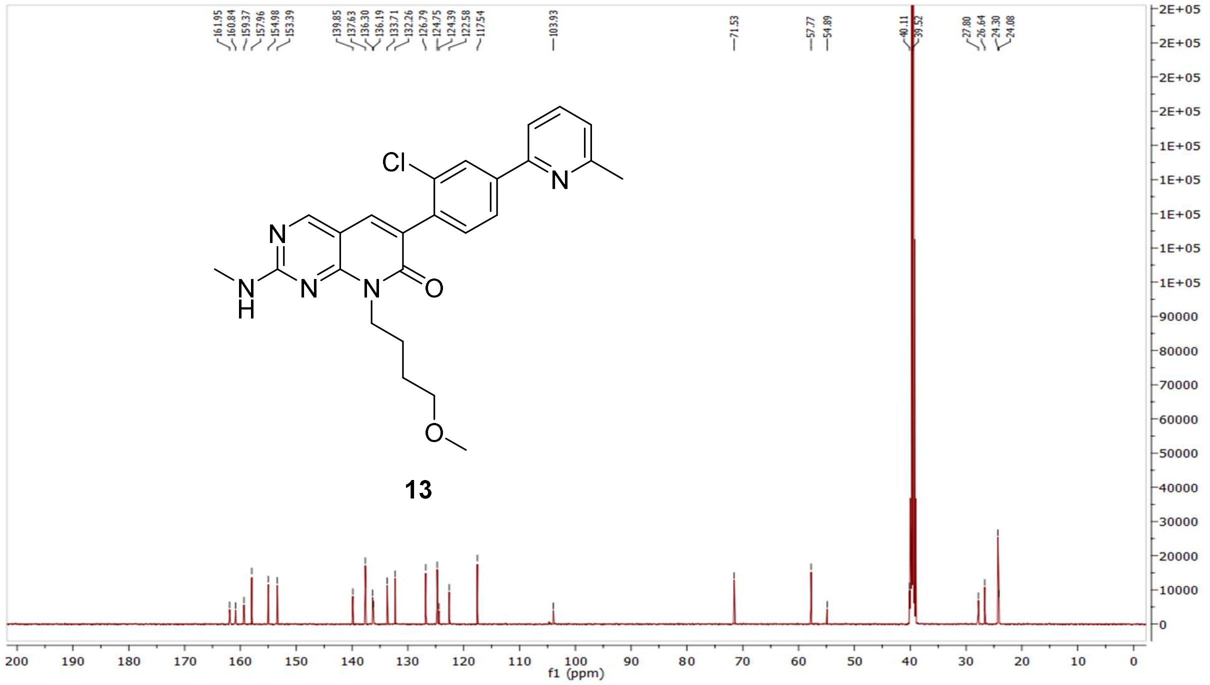


HRMS (FTMS +p MALDI) compound **13**:

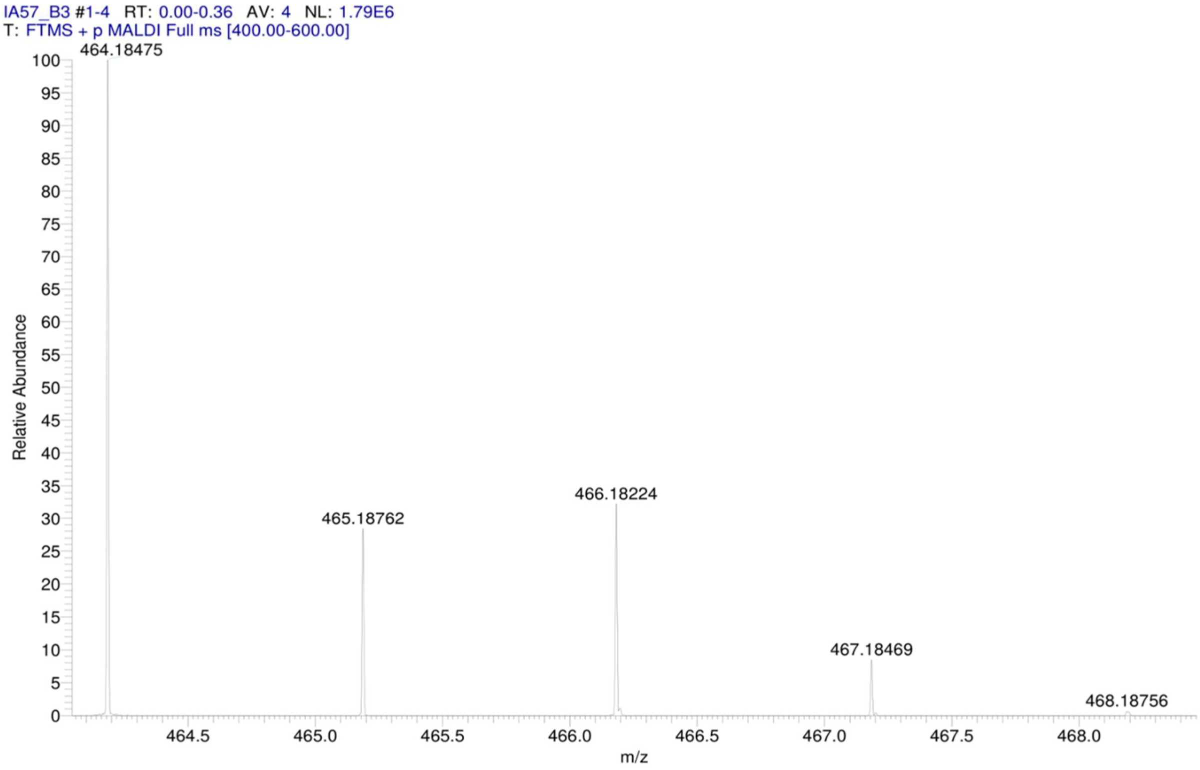

HPLC-chromatograms compound **13**:

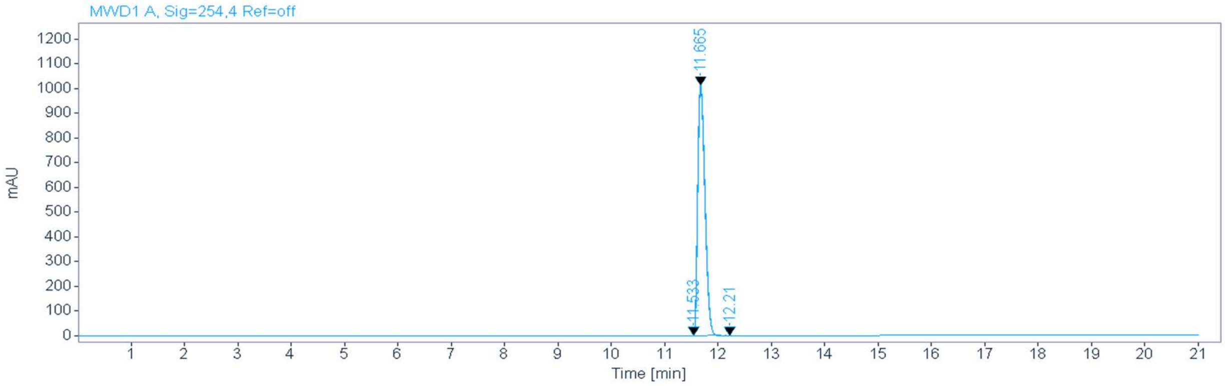


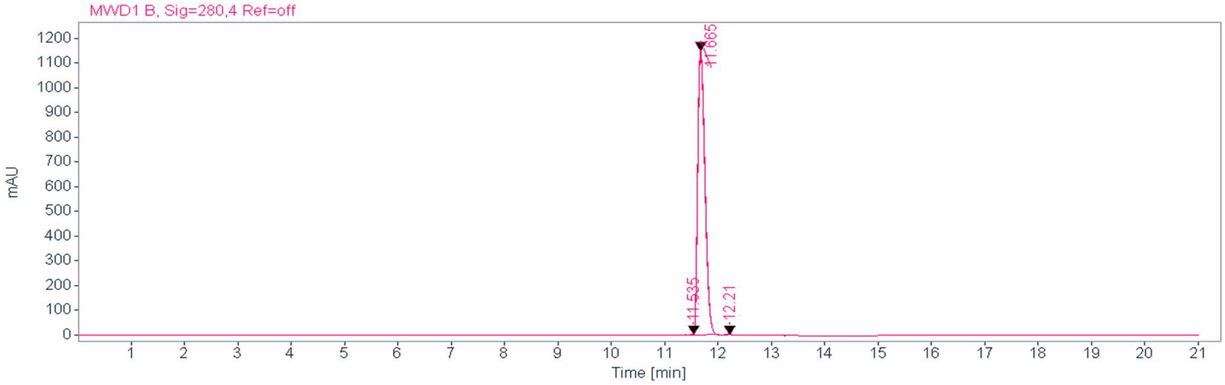


**Signal:** MWD1 A, Sig=254,4 Ref=off

| RT [min] | Type | Width [min] | Area | Height | Area% | Name |
|---|---|---|---|---|---|---|
| 11.533 | MM | 0.1061 | 16.0557 | 2.5214 | 0.1673 | |
| 11.665 | MM | 0.1570 | 9564.9688 | 1015.1769 | 99.6704 | |
| 12.210 | MM | 0.1409 | 15.5773 | 1.8423 | 0.1623 | |
| | | Sum | 9596.6018 | | | |

**Signal:** MWD1 B, Sig=280,4 Ref=off

| RT [min] | Type | Width [min] | Area | Height | Area% | Name |
|---|---|---|---|---|---|---|
| 11.535 | MM | 0.1000 | 20.4998 | 3.4169 | 0.1882 | |
| 11.665 | MM | 0.1570 | 10853.8027 | 1152.1707 | 99.6305 | |
| 12.210 | MM | 0.1535 | 19.7546 | 2.1444 | 0.1813 | |
| | | Sum | 10894.0571 | | | |

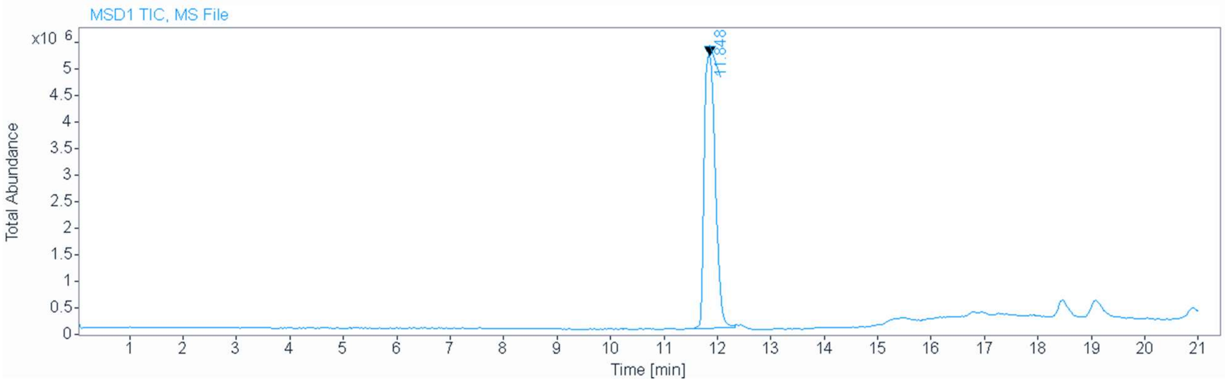


**Signal** MSD1 TIC, MS File

**Peak RT** 11.848

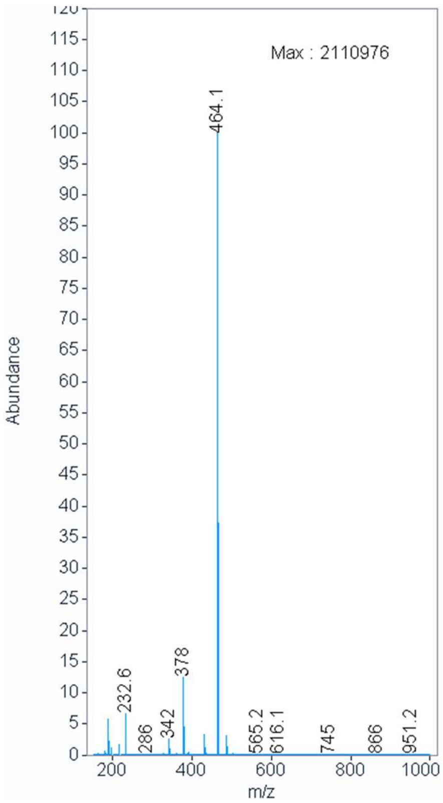

[1]H NMR compound **14**:

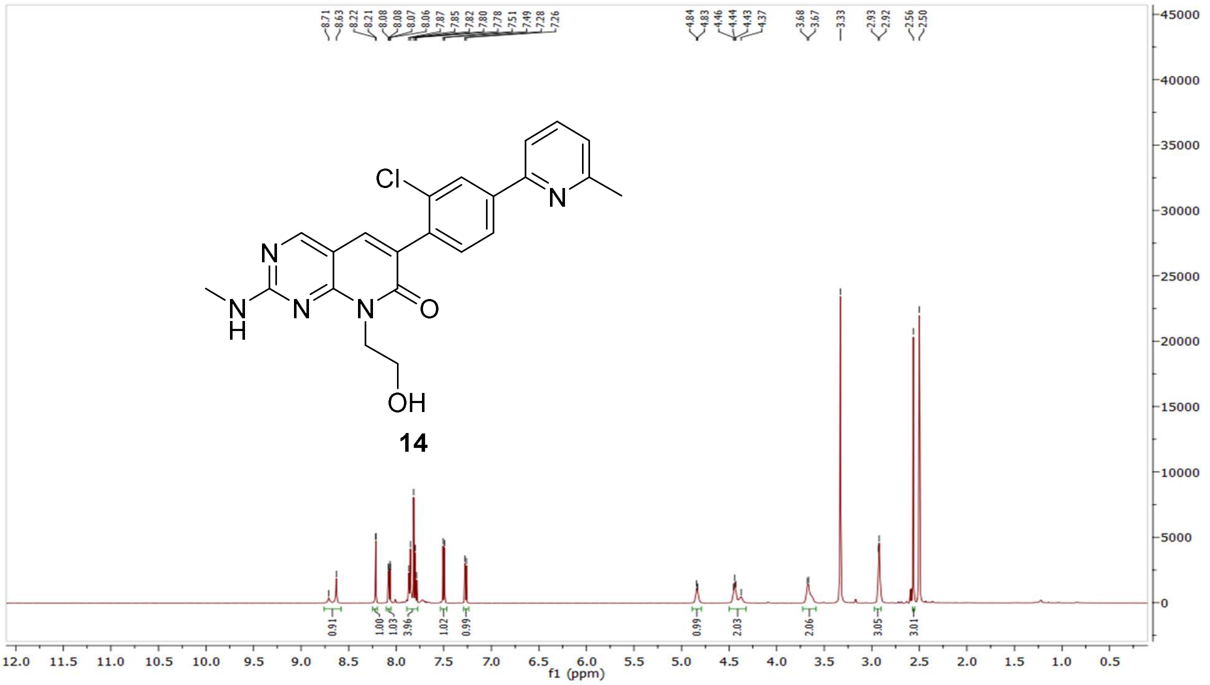

[13]C NMR compound **14:**

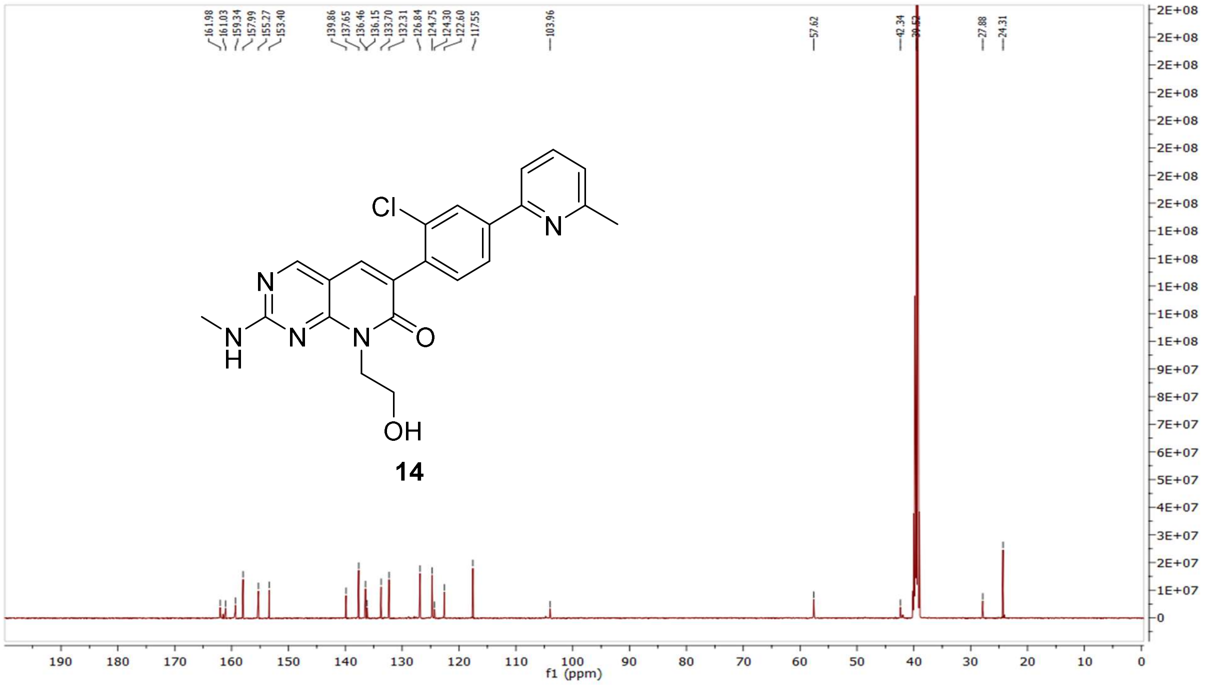

HRMS (FTMS +p MALDI) compound **14**:

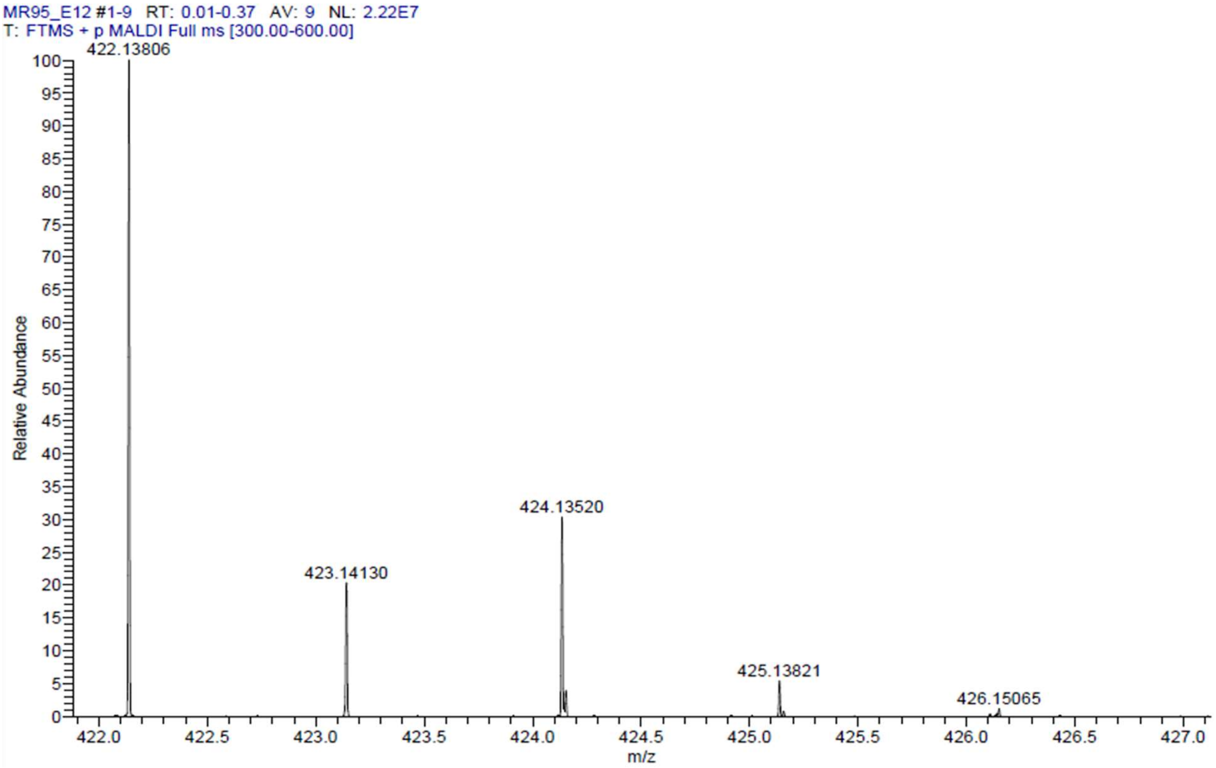

HPLC-chromatograms compound **14:**

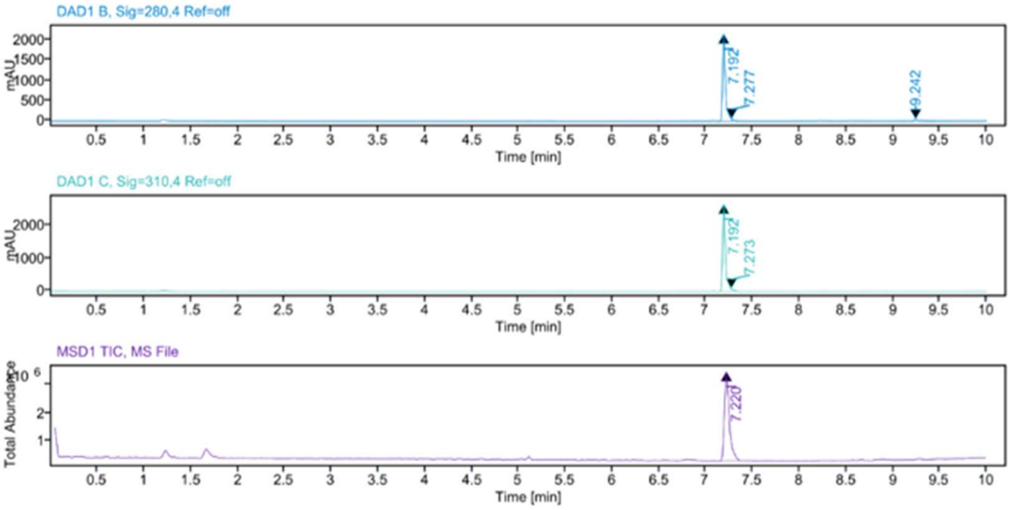


**Sample Purity**

| | |
|---|---|
| Signal Description | DAD1 B, Sig=280,4 Ref=off |

| Sample Name | Name | RT | Width | Area | Area% | Height |
|---|---|---|---|---|---|---|
| MR95 Fr.3 | | 7.192 | 0.029 | 4522.7153 | 95.72 | 2147.8464 |
| MR95 Fr.3 | | 7.277 | 0.020 | 79.6255 | 1.69 | 57.6529 |
| MR95 Fr.3 | | 9.242 | 0.037 | 122.7970 | 2.60 | 42.9522 |

| | |
|---|---|
| Max Area% | 95.716 |
| UV Signal Purity>95% | Pass |

| | |
|---|---|
| Signal Description | DAD1 C, Sig=310,4 Ref=off |

| Sample Name | Name | RT | Width | Area | Area% | Height |
|---|---|---|---|---|---|---|
| MR95 Fr.3 | | 7.192 | 0.029 | 5664.7227 | 97.98 | 2642.6379 |
| MR95 Fr.3 | | 7.273 | 0.022 | 116.7829 | 2.02 | 77.8921 |

| | |
|---|---|
| Max Area% | 97.980 |
| UV Signal Purity>95% | Pass |

**MSD Apex Spectra**

**RT:**

7.220

**Sample Name:**

MR95 Fr.3

**Signal Name:**

MSD1TIC

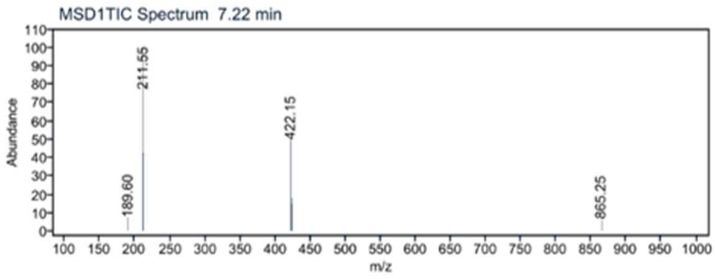

$^{1}$H NMR compound **15**:

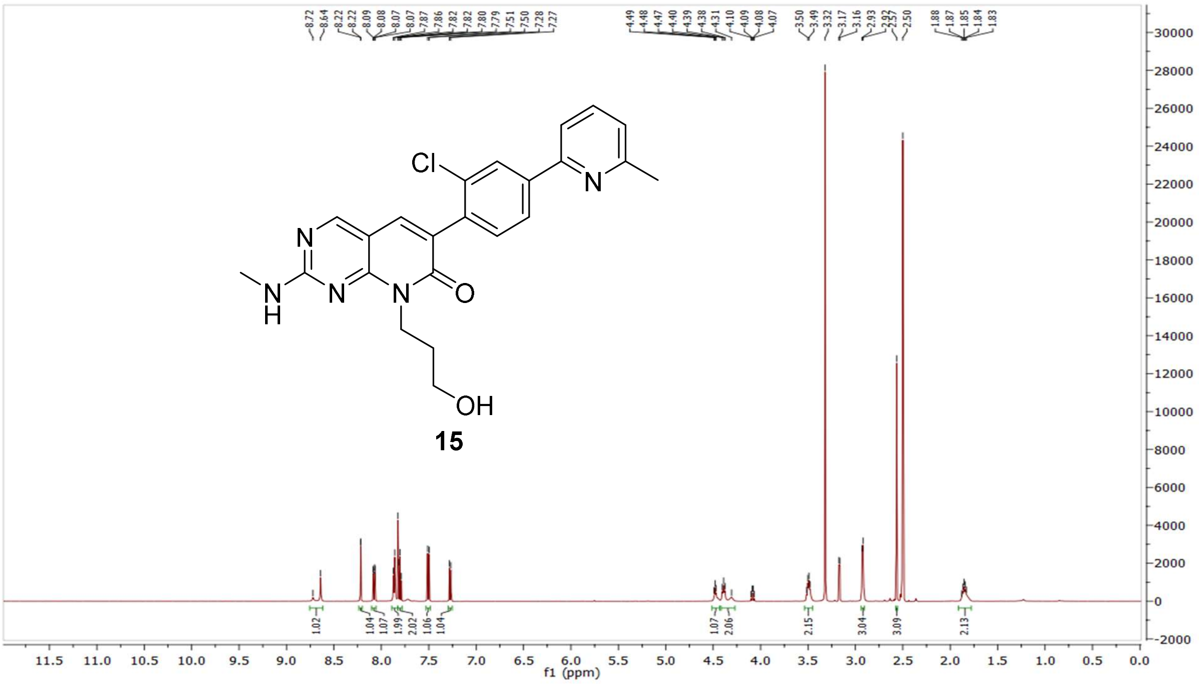


$^{13}$C NMR compound **15:**

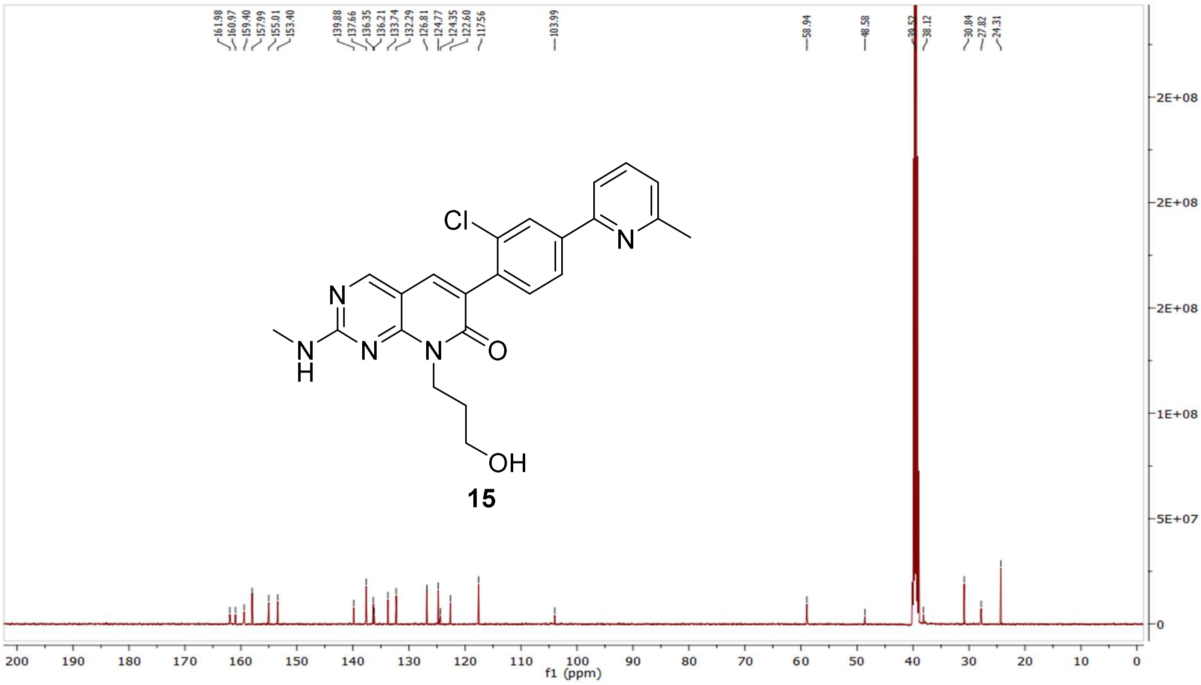

HRMS (FTMS +p MALDI) compound **15**:

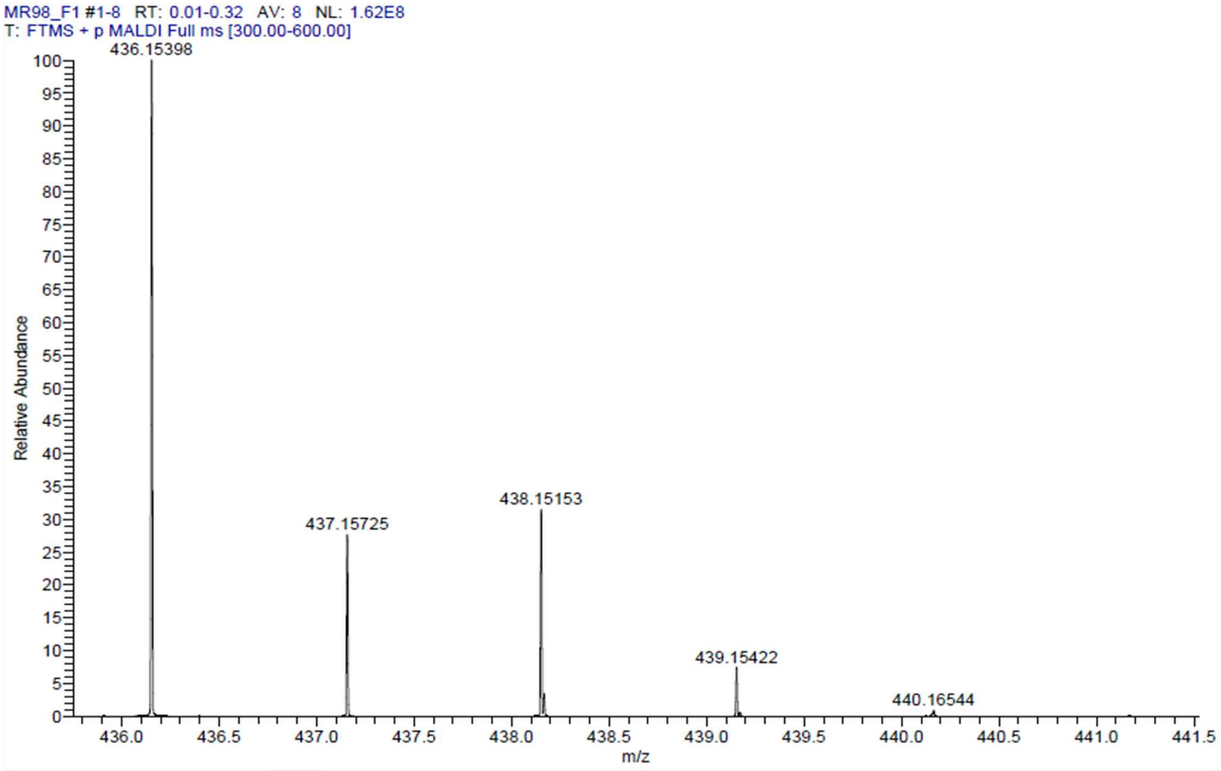

HPLC-chromatograms compound **15**:

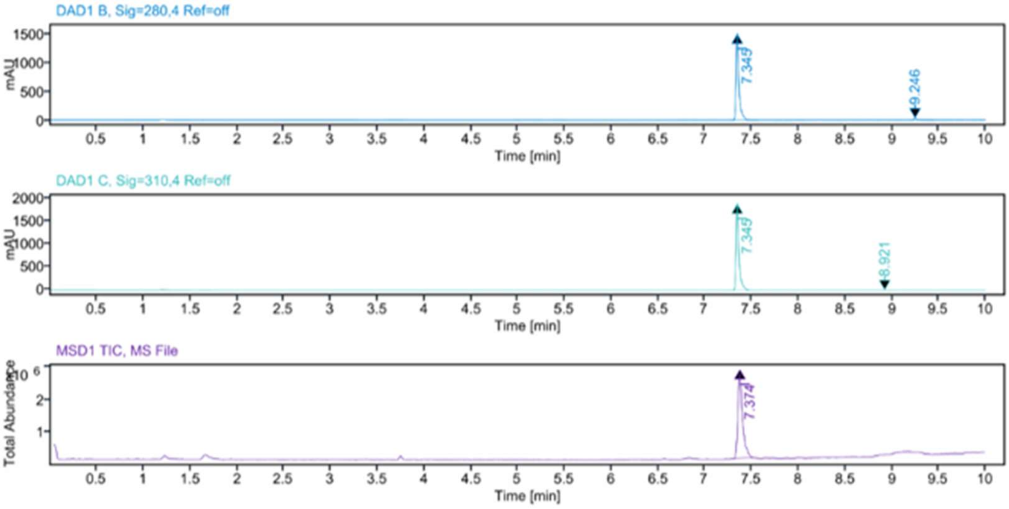


**Sample Purity**

Signal Description DAD1 B, Sig=280,4 Ref=off

| Sample Name | Name | RT | Width | Area | Area% | Height |
|---|---|---|---|---|---|---|
| MR98 Fr.1 | | 7.345 | 0.028 | 3252.5308 | 97.04 | 1520.6526 |
| MR98 Fr.1 | | 9.246 | 0.037 | 99.1988 | 2.96 | 34.0651 |

Max Area% 97.040

UV Signal Purity>95% Pass

Signal Description DAD1 C, Sig=310,4 Ref=off

| Sample Name | Name | RT | Width | Area | Area% | Height |
|---|---|---|---|---|---|---|
| MR98 Fr.1 | | 7.345 | 0.029 | 4090.5996 | 99.80 | 1901.1281 |
| MR98 Fr.1 | | 8.921 | 0.034 | 8.3521 | 0.20 | 3.1997 |

Max Area% 99.796

UV Signal Purity>95% Pass

**MSD Apex Spectra**

**RT:**

7.374

**Sample Name:**

MR98 Fr.1

**Signal Name:**

MSD1TIC

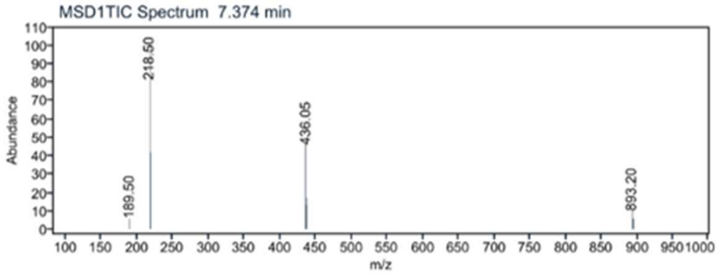

$^{1}H$ NMR compound **16**:

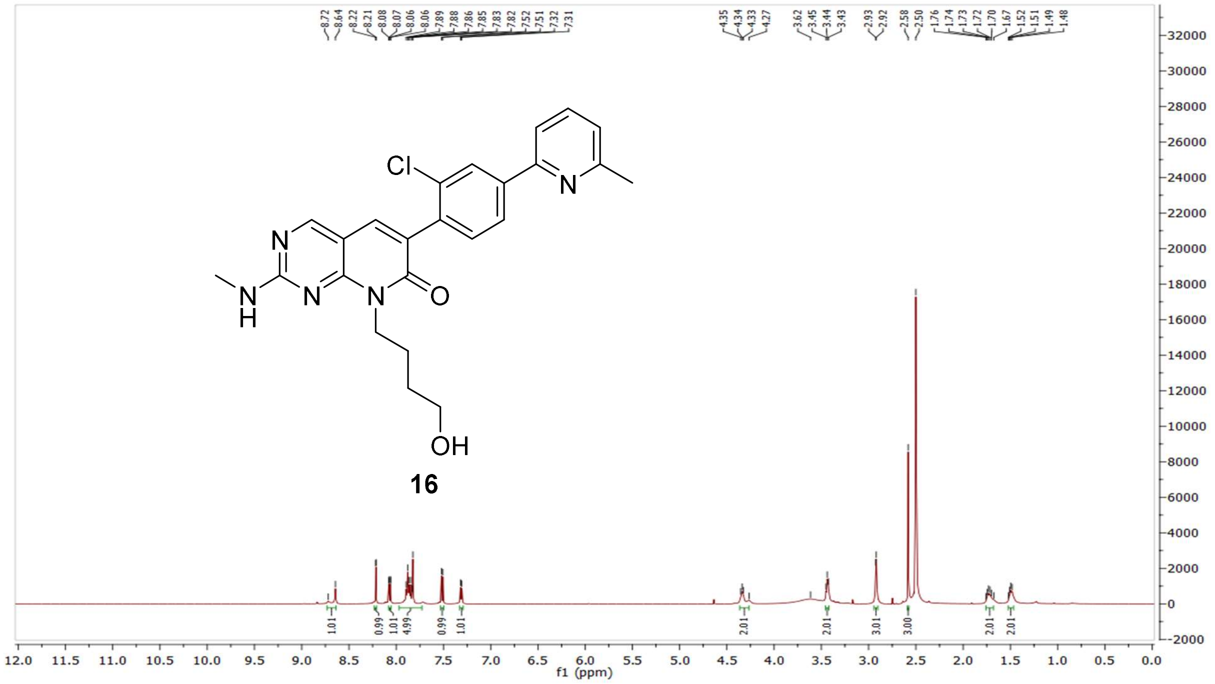


$^{13}C$ NMR compound **16:**

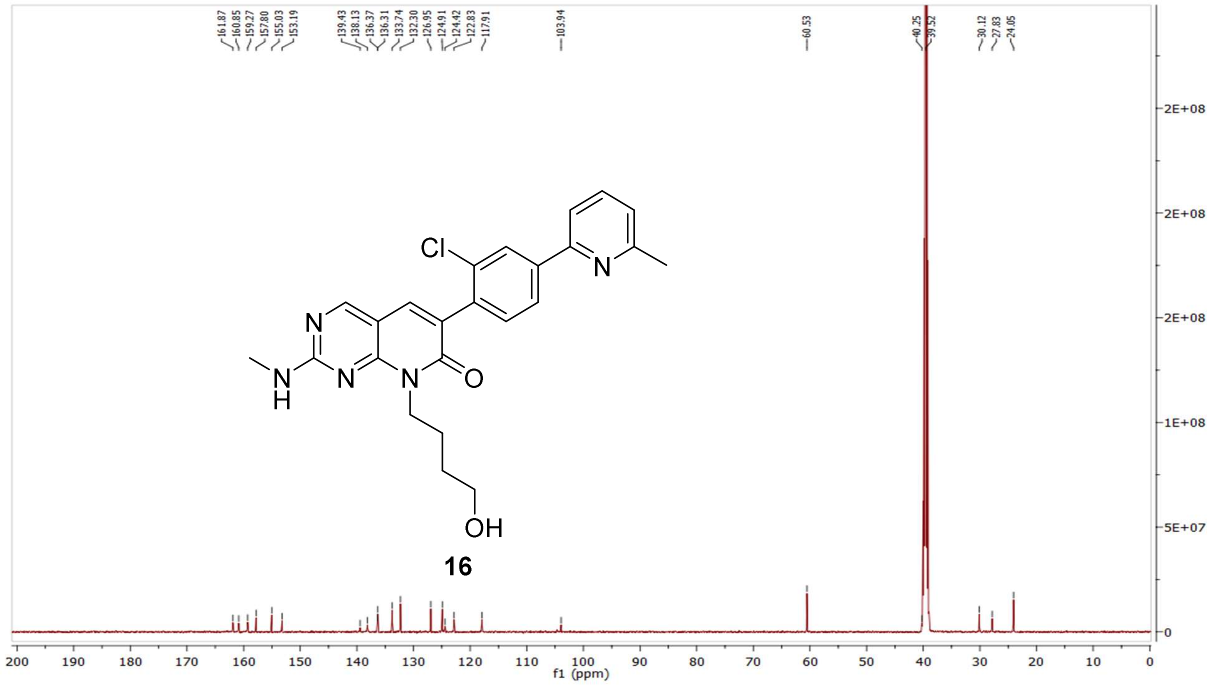

HRMS (FTMS +p MALDI) compound **16**:

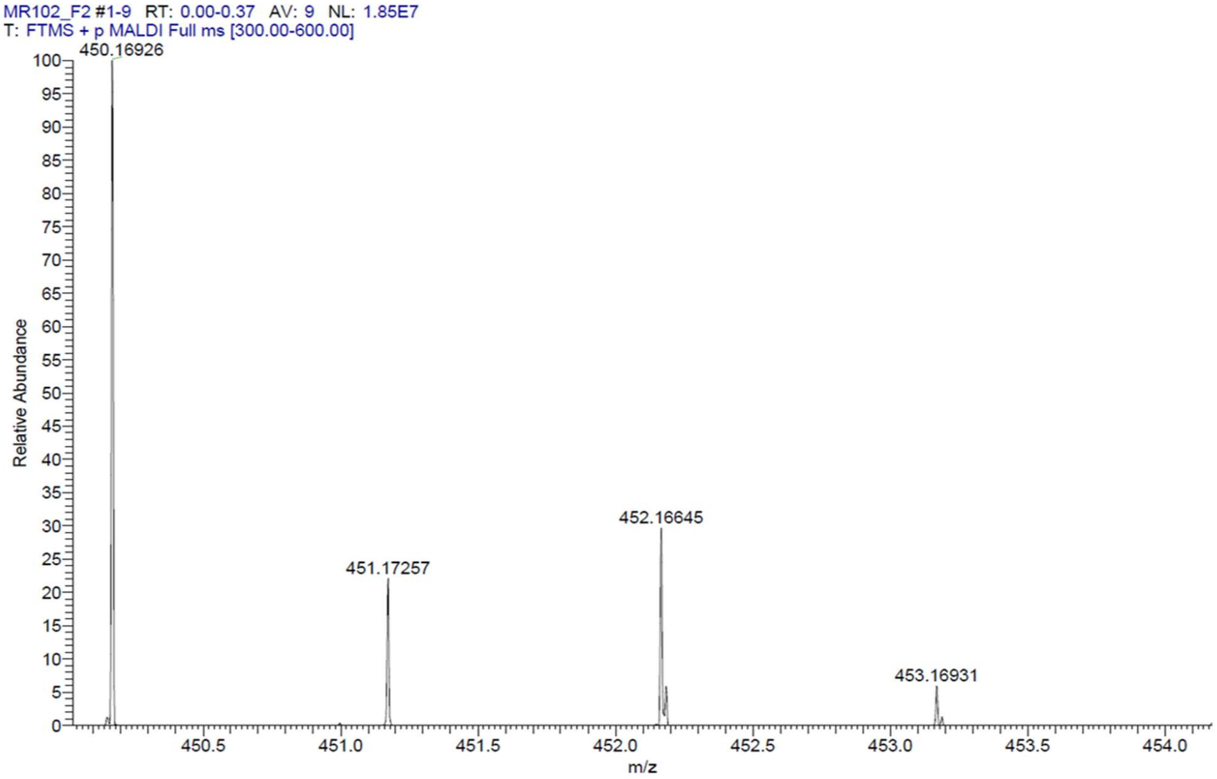

HPLC-chromatograms compound **16**:

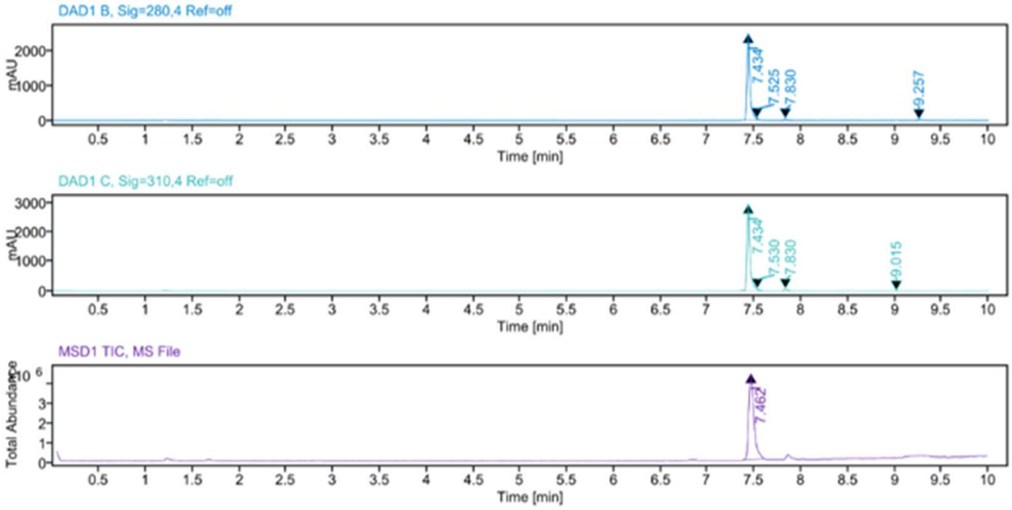


**Sample Purity**

Signal Description DAD1 B, Sig=280,4 Ref=off

| Sample Name | Name | RT | Width | Area | Area% | Height |
|---|---|---|---|---|---|---|
| MR102 Fr.1 | | 7.434 | 0.030 | 5540.8784 | 95.17 | 2532.4277 |
| MR102 Fr.1 | | 7.525 | 0.020 | 81.3289 | 1.40 | 64.4438 |
| MR102 Fr.1 | | 7.830 | 0.029 | 122.0781 | 2.10 | 59.3153 |
| MR102 Fr.1 | | 9.257 | 0.037 | 77.8087 | 1.34 | 33.7801 |

Max Area% 95.170

UV Signal Purity>95% Pass

Signal Description DAD1 C, Sig=310,4 Ref=off

| Sample Name | Name | RT | Width | Area | Area% | Height |
|---|---|---|---|---|---|---|
| MR102 Fr.1 | | 7.434 | 0.031 | 6833.9258 | 95.94 | 2988.2388 |
| MR102 Fr.1 | | 7.530 | 0.017 | 101.9787 | 1.43 | 79.8684 |
| MR102 Fr.1 | | 7.830 | 0.030 | 165.4347 | 2.32 | 72.7832 |
| MR102 Fr.1 | | 9.015 | 0.029 | 21.5153 | 0.30 | 11.6341 |

Max Area% 95.944

UV Signal Purity>95% Pass

**MSD Apex Spectra**

**RT:**

7.462

**Sample Name:**

MR102 Fr.1

**Signal Name:**

MSD1TIC

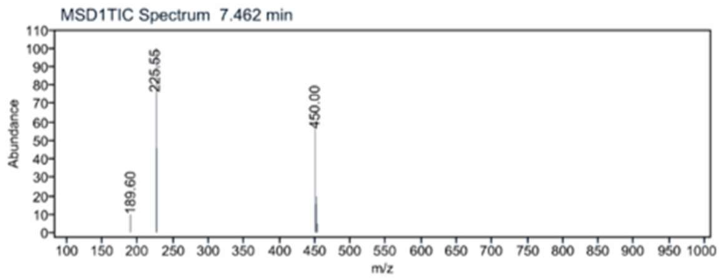


[1]H NMR compound **17**:

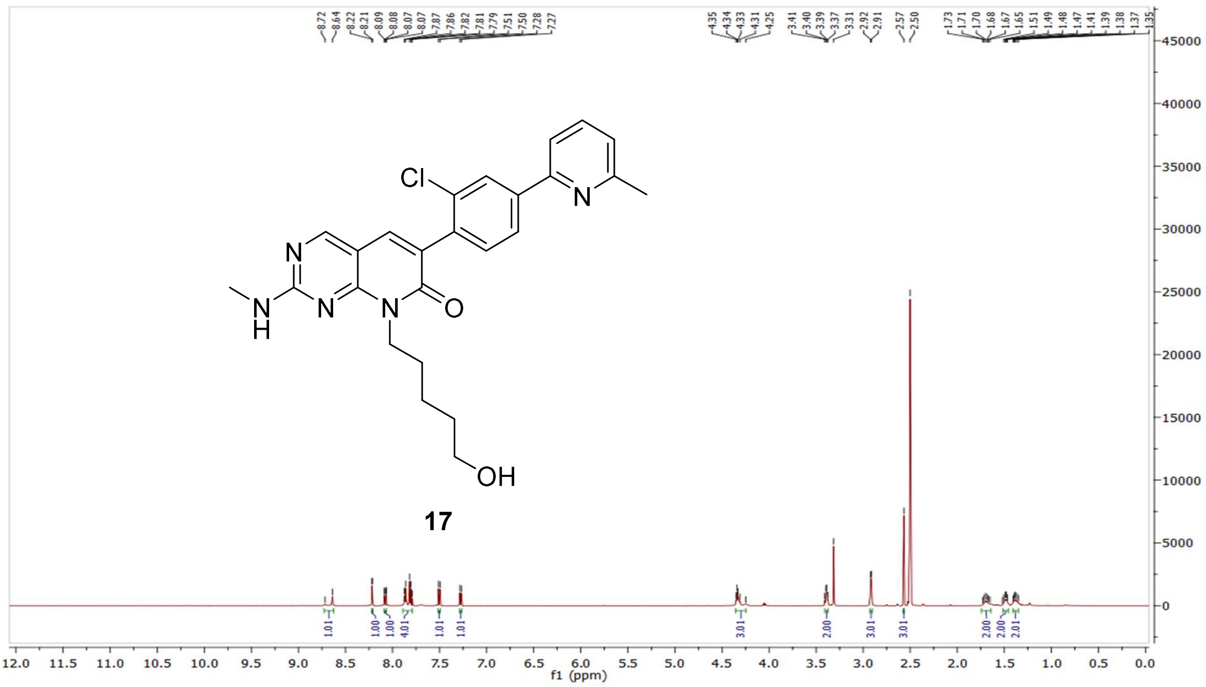

$^{13}C$ NMR compound **17:**

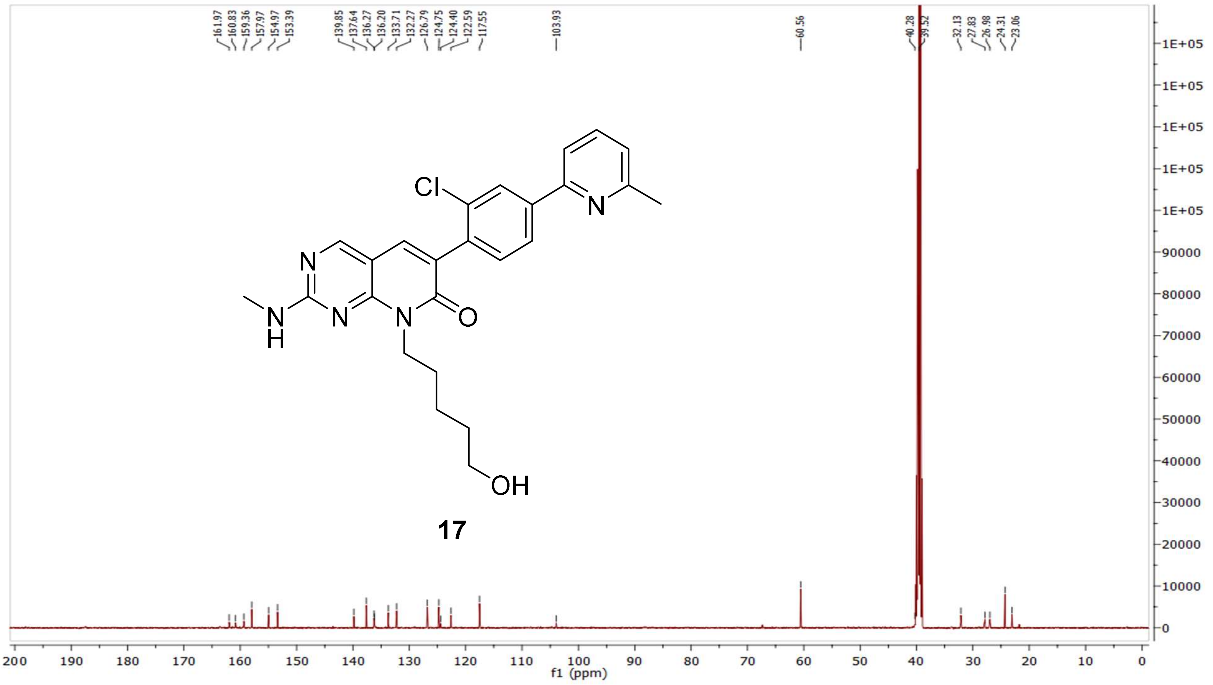


HRMS (FTMS +p MALDI) compound **17**:

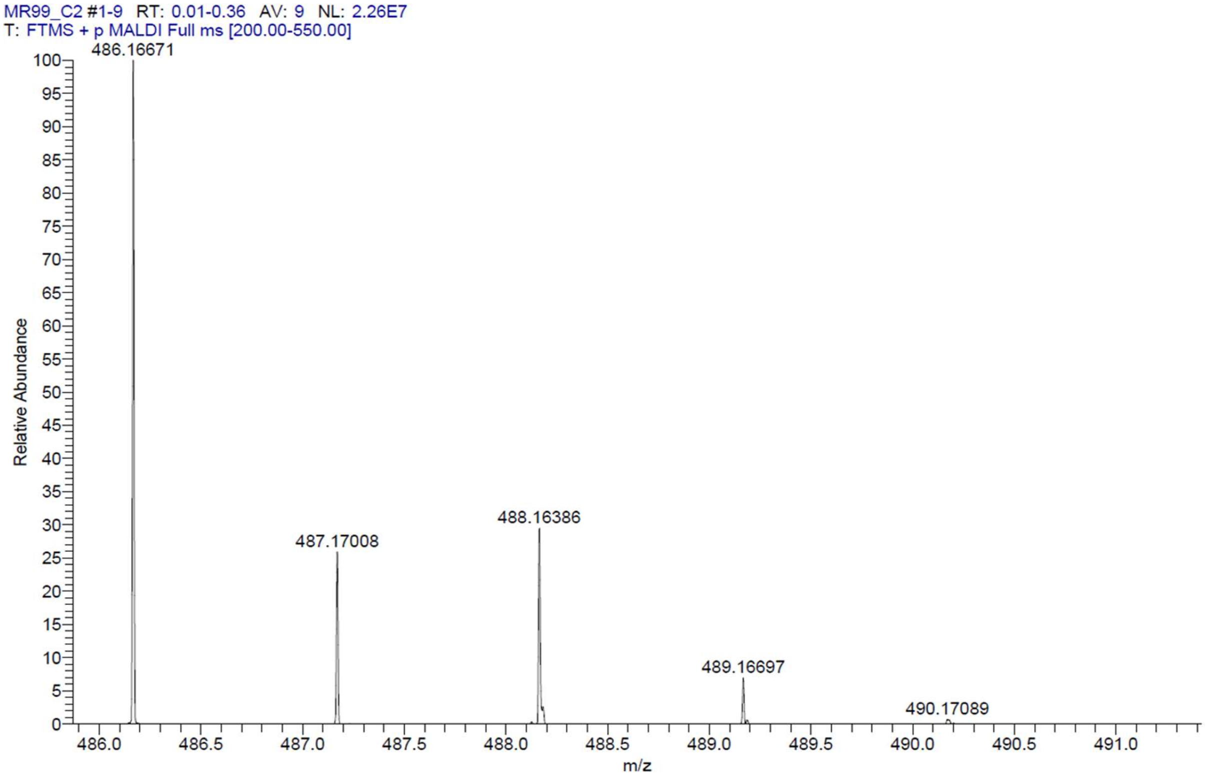

HPLC-chromatograms compound **17**:

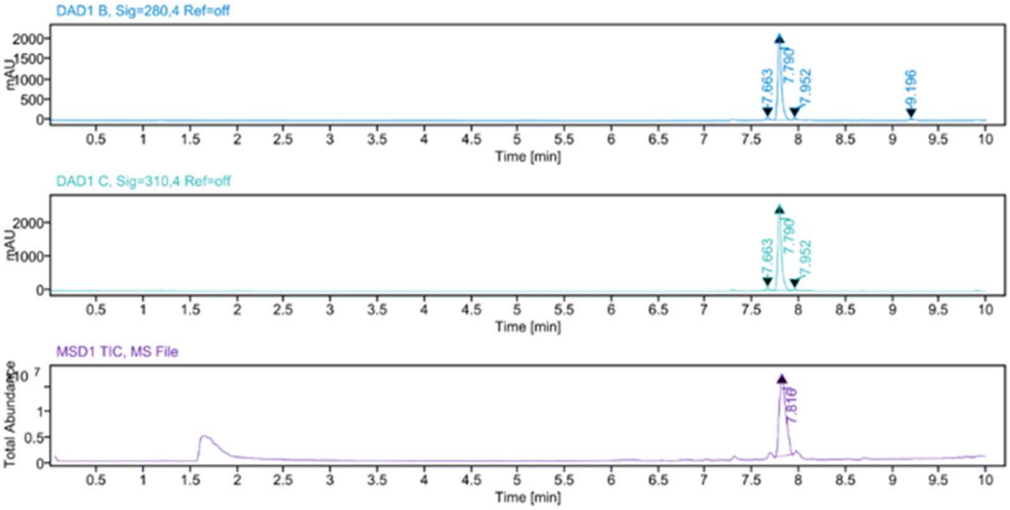


**Sample Purity**

Signal Description DAD1 B, Sig=280,4 Ref=off

| Sample Name | Name | RT | Width | Area | Area% | Height |
|---|---|---|---|---|---|---|
| MR99 Fr.1 | | 7.663 | 0.034 | 117.2607 | 1.93 | 55.9089 |
| MR99 Fr.1 | | 7.790 | 0.038 | 5778.2012 | 95.12 | 2142.6150 |
| MR99 Fr.1 | | 7.952 | 0.034 | 102.1053 | 1.68 | 49.8757 |
| MR99 Fr.1 | | 9.196 | 0.038 | 76.8694 | 1.27 | 34.0692 |

Max Area% 95.123

UV Signal Purity>95% Pass

Signal Description DAD1 C, Sig=310,4 Ref=off

| Sample Name | Name | RT | Width | Area | Area% | Height |
|---|---|---|---|---|---|---|
| MR99 Fr.1 | | 7.663 | 0.037 | 217.0914 | 2.96 | 91.4450 |
| MR99 Fr.1 | | 7.790 | 0.038 | 7033.5806 | 95.75 | 2588.5024 |
| MR99 Fr.1 | | 7.952 | 0.034 | 95.2009 | 1.30 | 45.8905 |

Max Area% 95.749

UV Signal Purity>95% Pass

**MSD Apex Spectra**

**RT:**

7.816

**Sample Name:**

MR99 Fr.1

**Signal Name:**

MSD1TIC

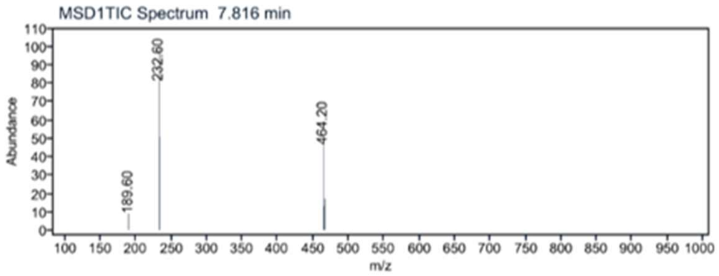

[1]H NMR compound **50**:

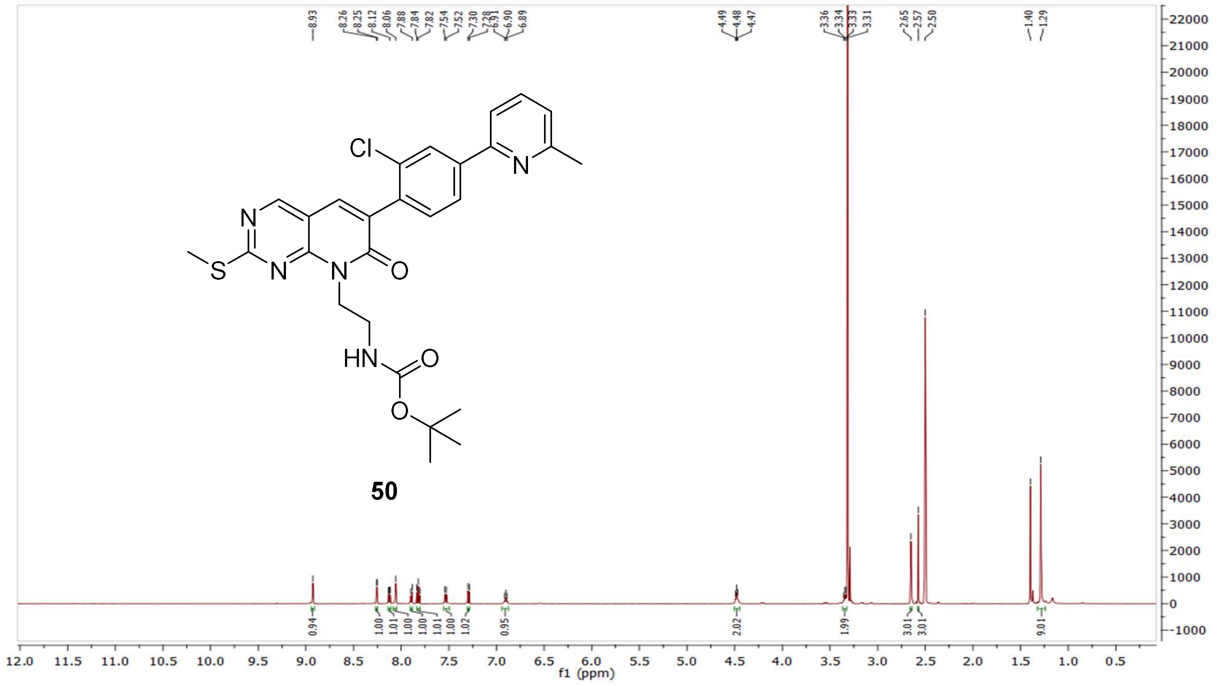


[13]C NMR compound **50:**

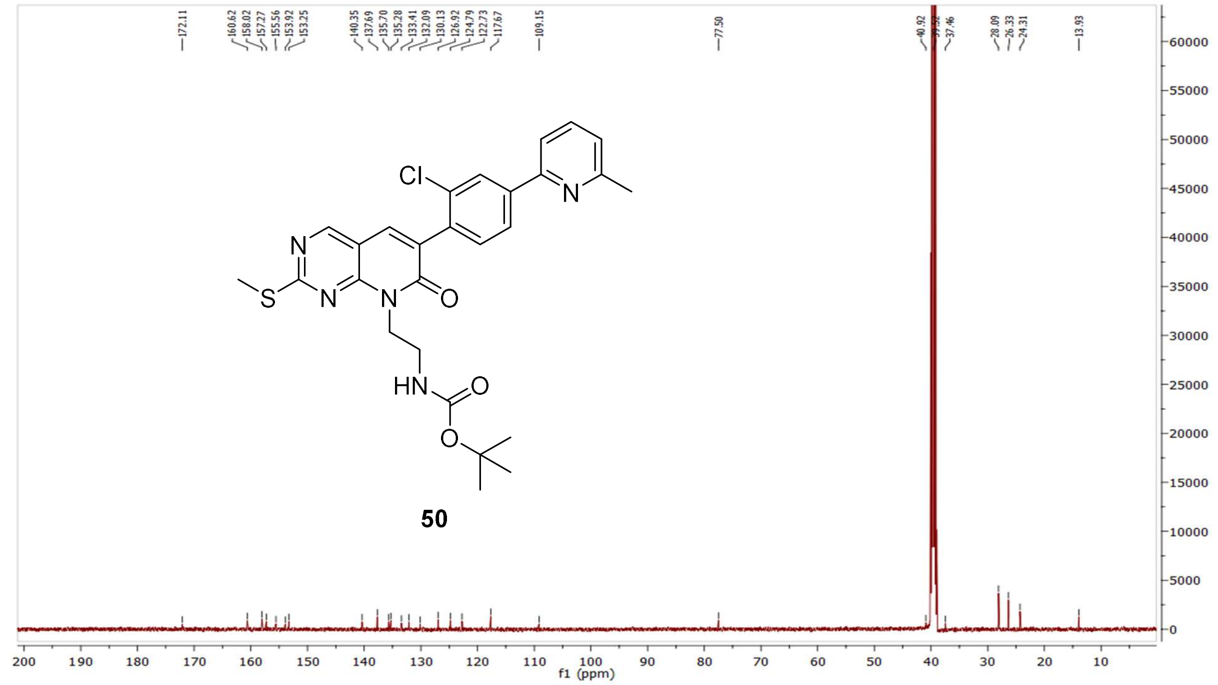

MS (ESI+) compound **50**:

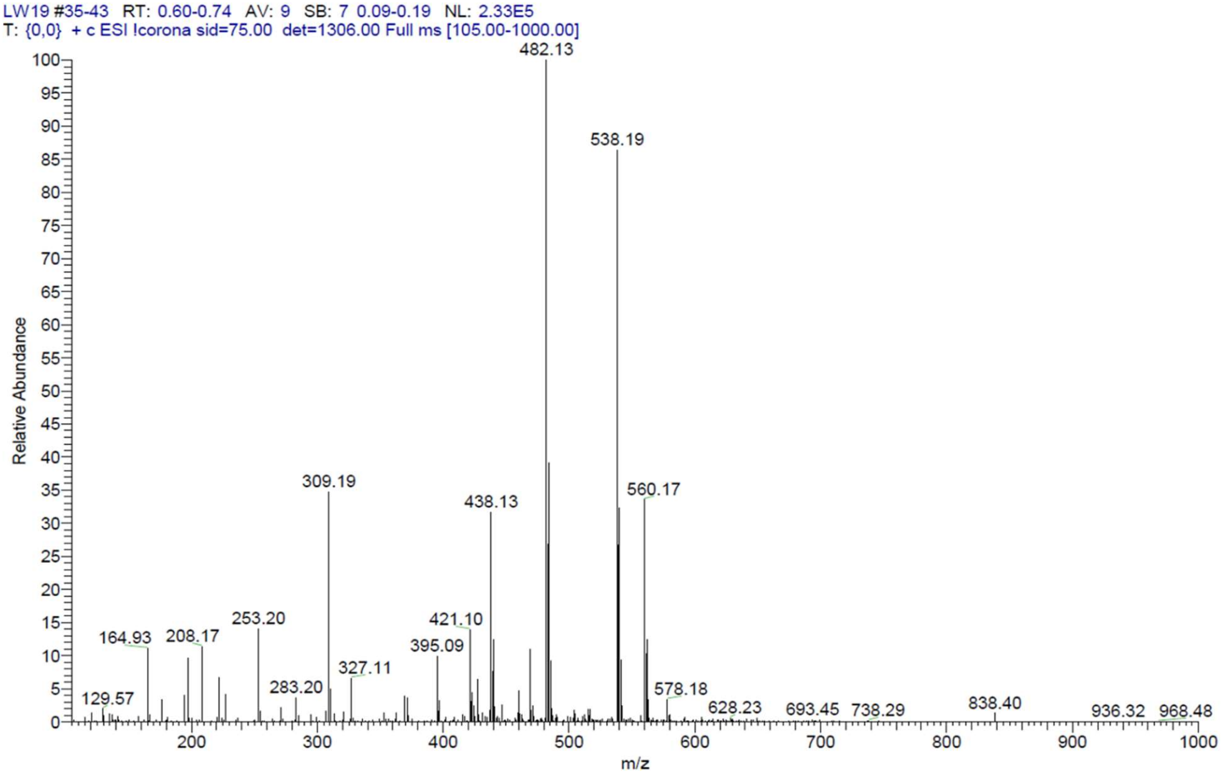


[1]H NMR compound **51**:

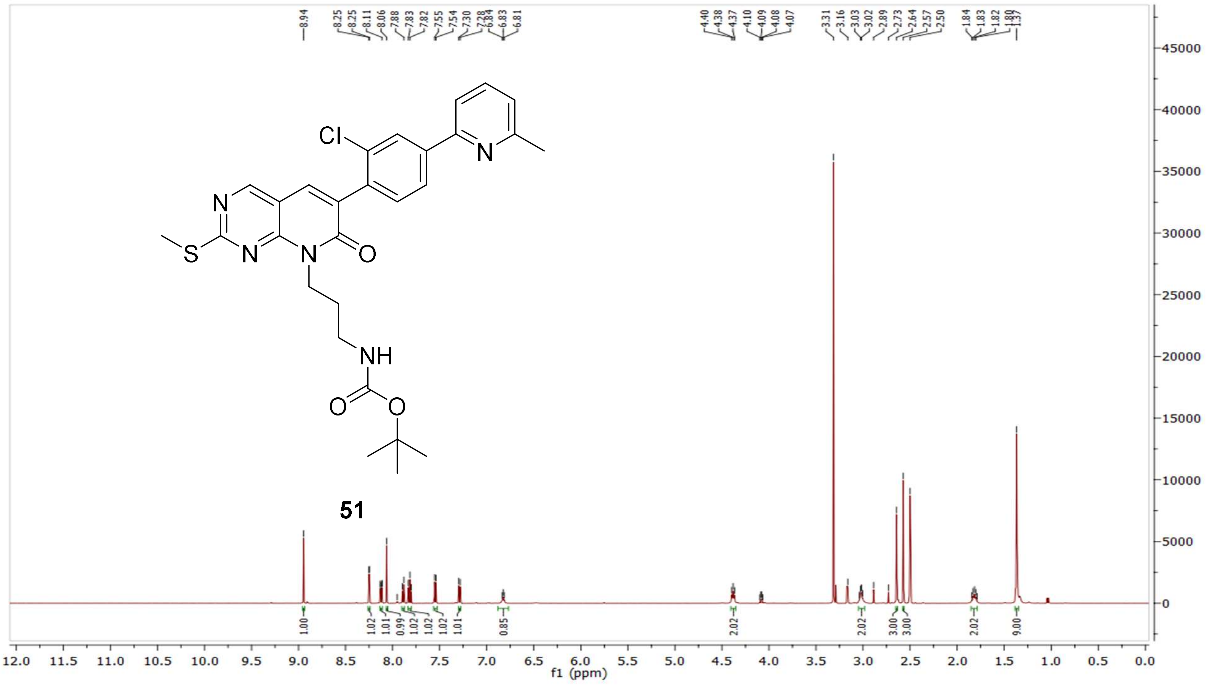

[13]C NMR compound **51:**

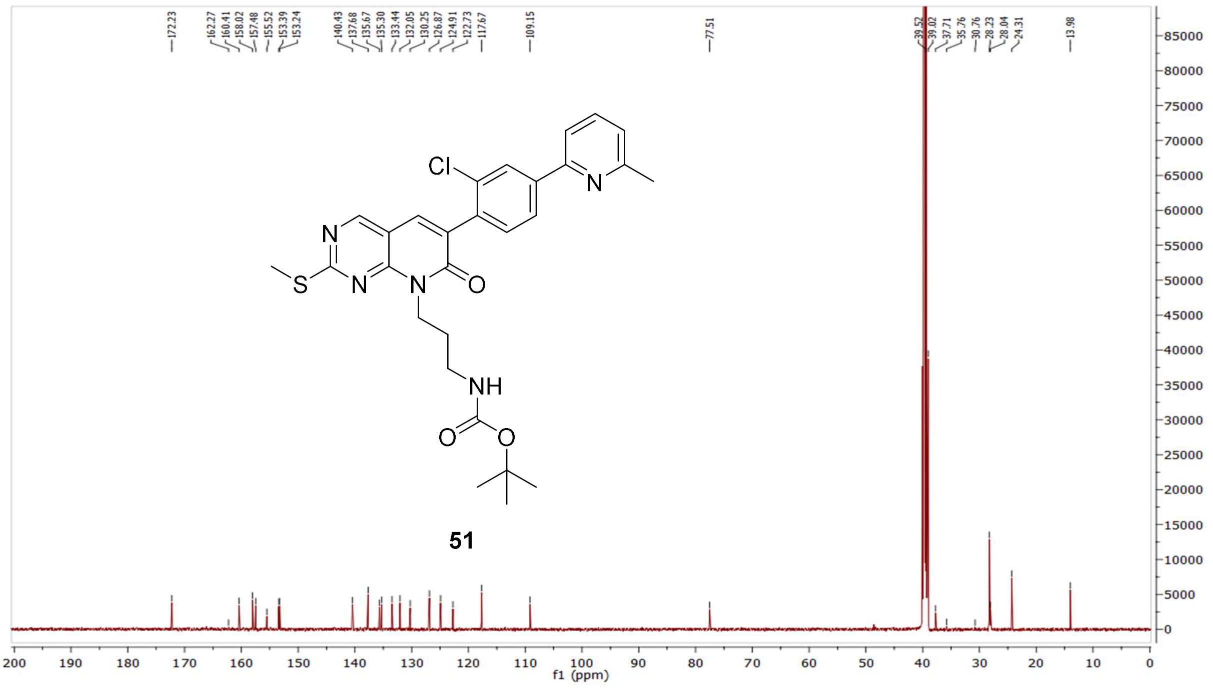


MS (ESI+) compound **51**:

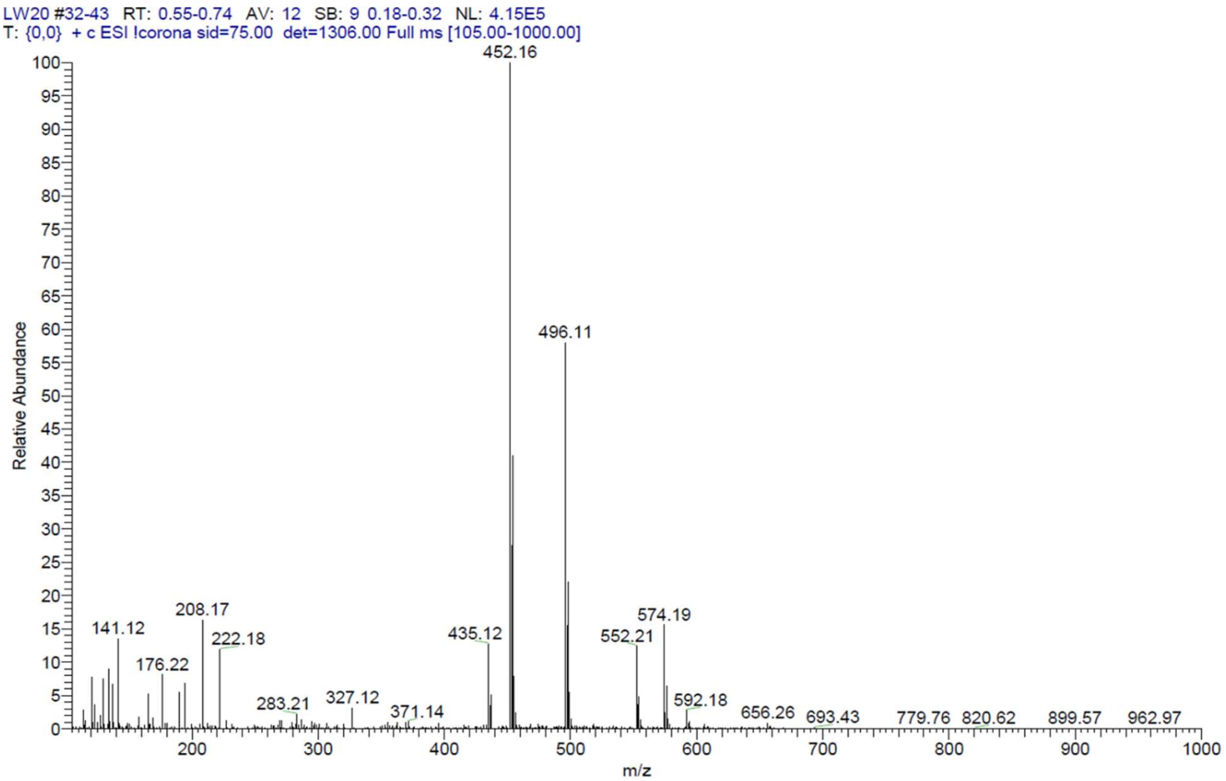

$^{1}$H NMR compound **52**:

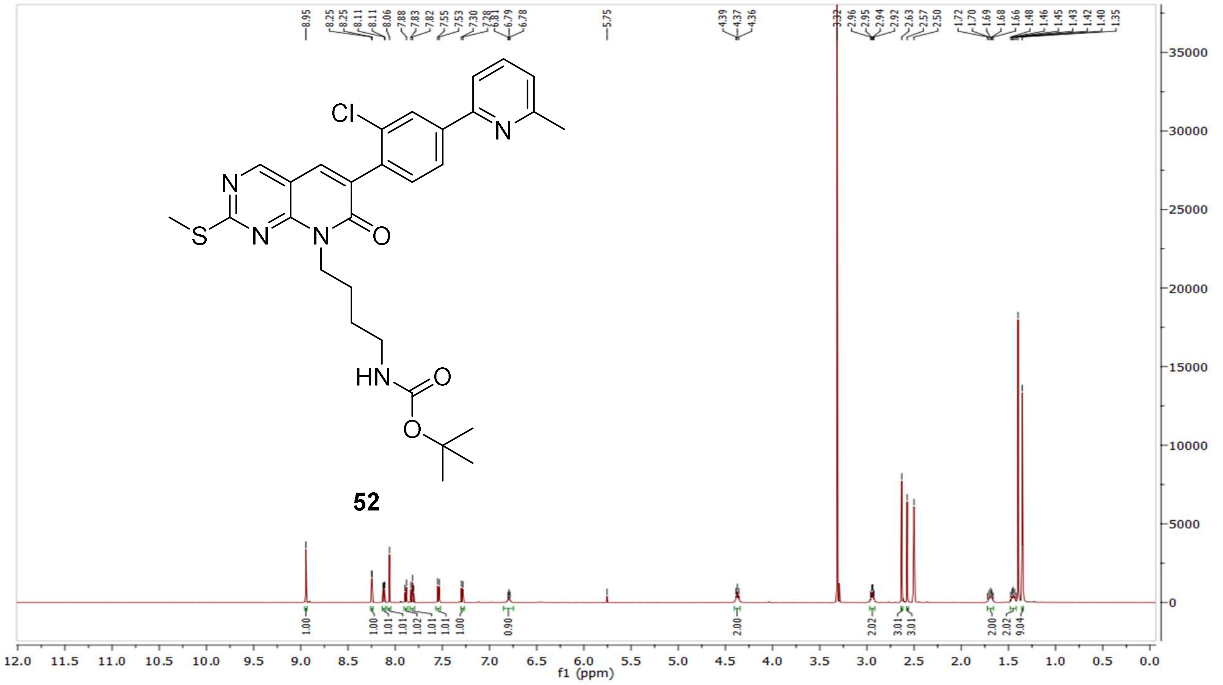

$^{13}$C NMR compound **52:**

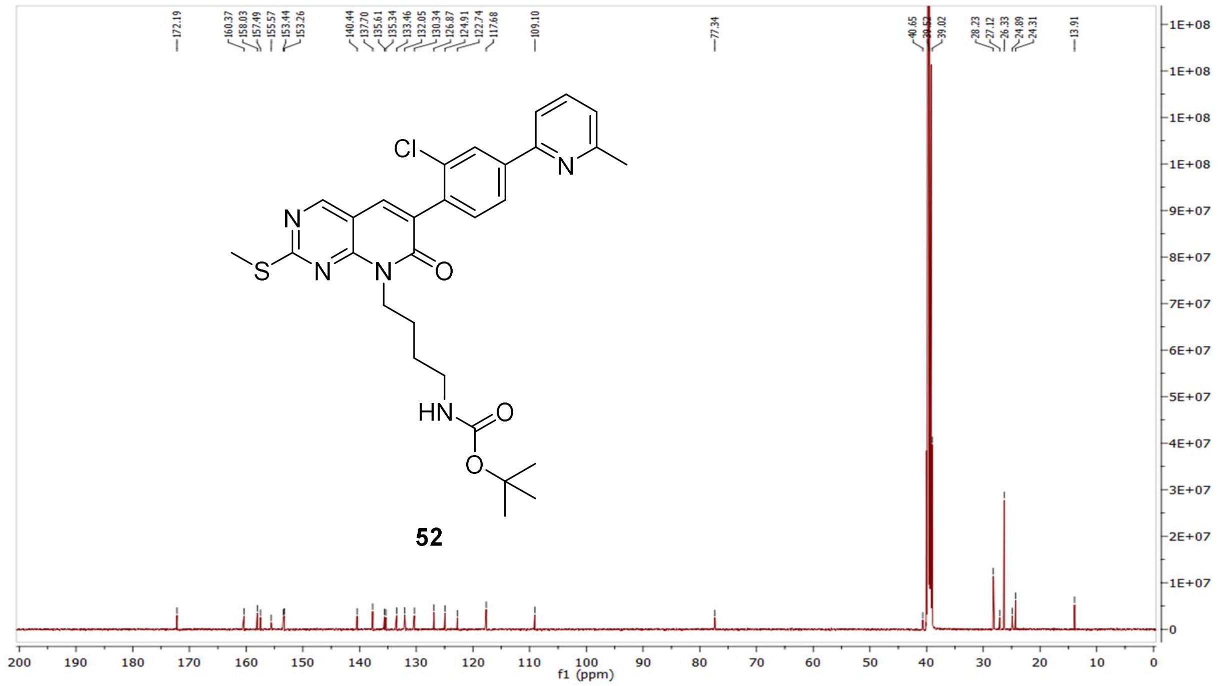

MS (ESI+) compound **52**:

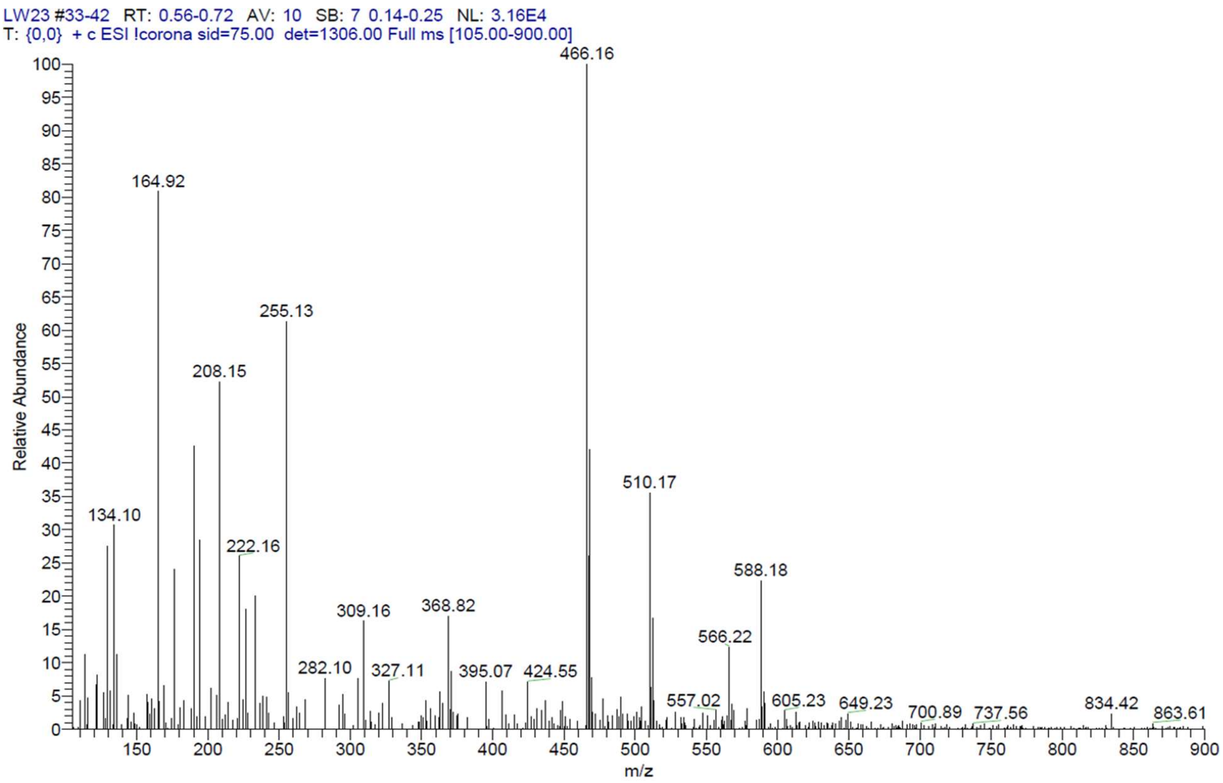


$^1$H NMR compound **53**:

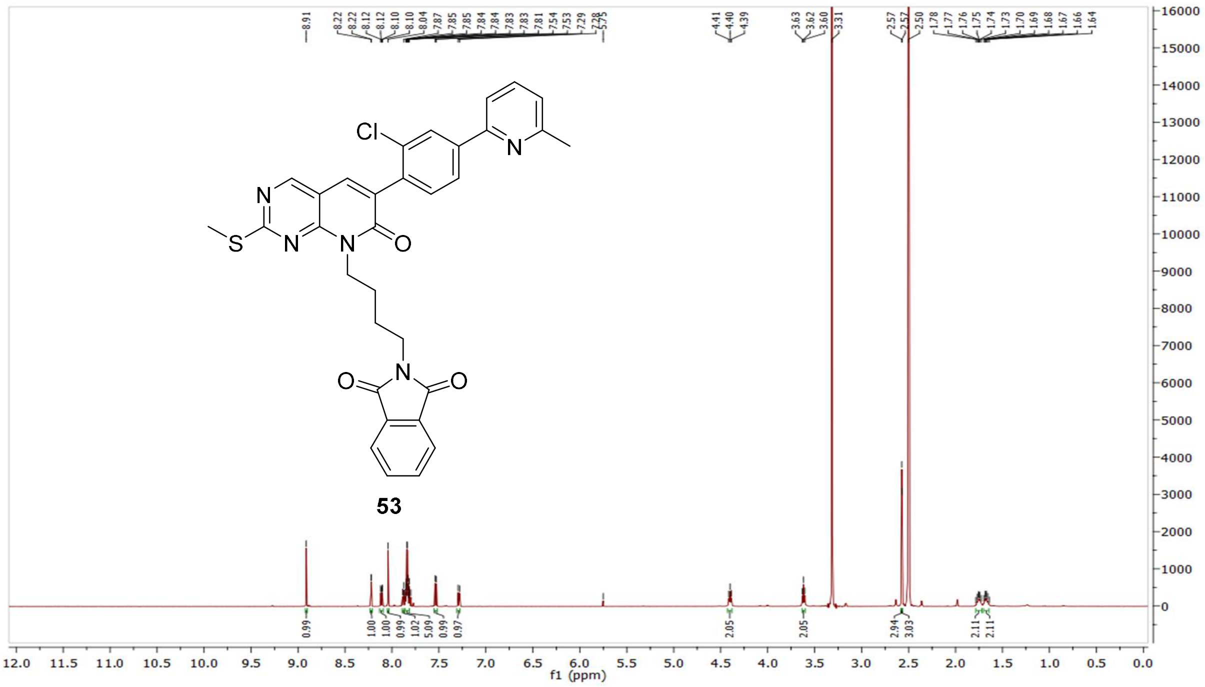

$^{13}$C NMR compound **53:**

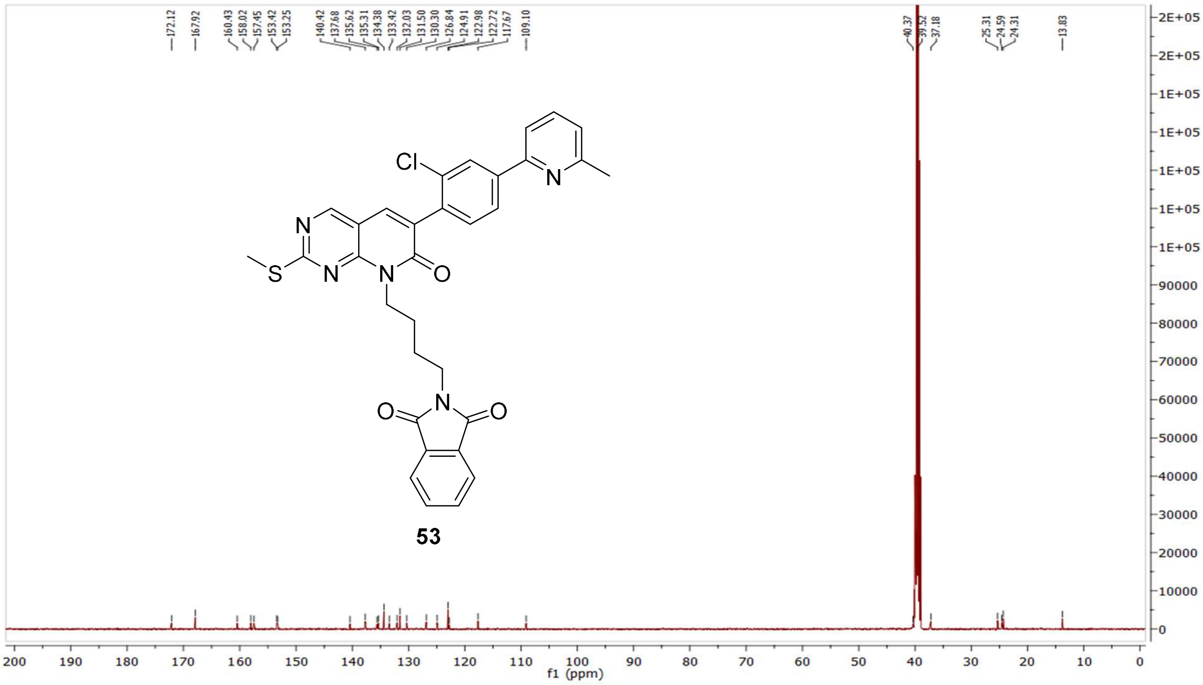


MS (ESI+) compound **53**:

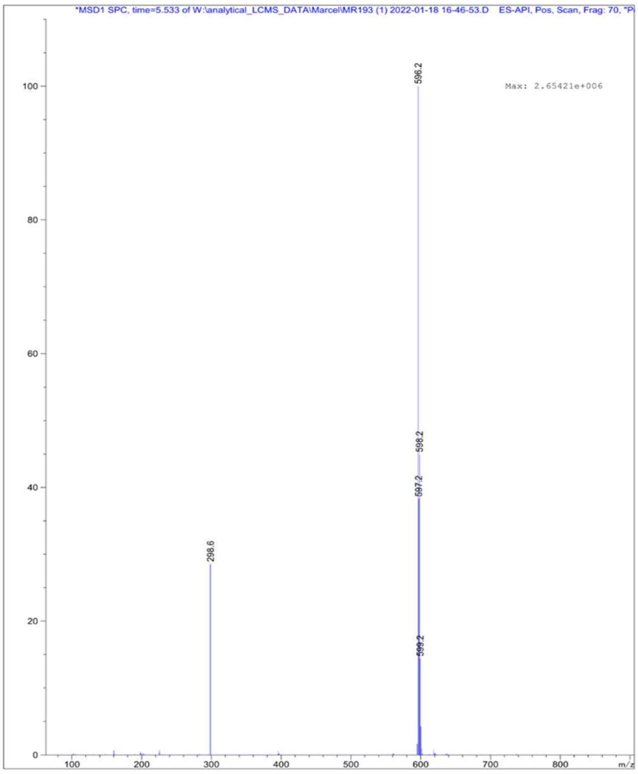

<sup></sup>

[1]H NMR compound **54**:

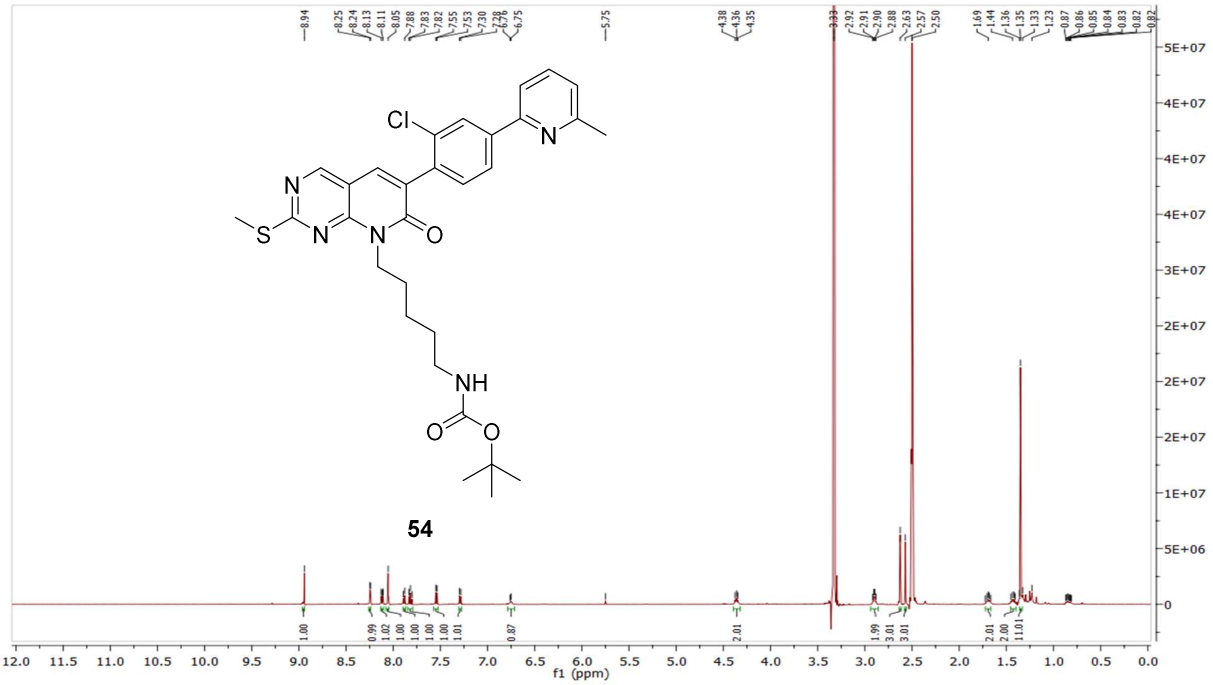

[13]C NMR compound **54:**

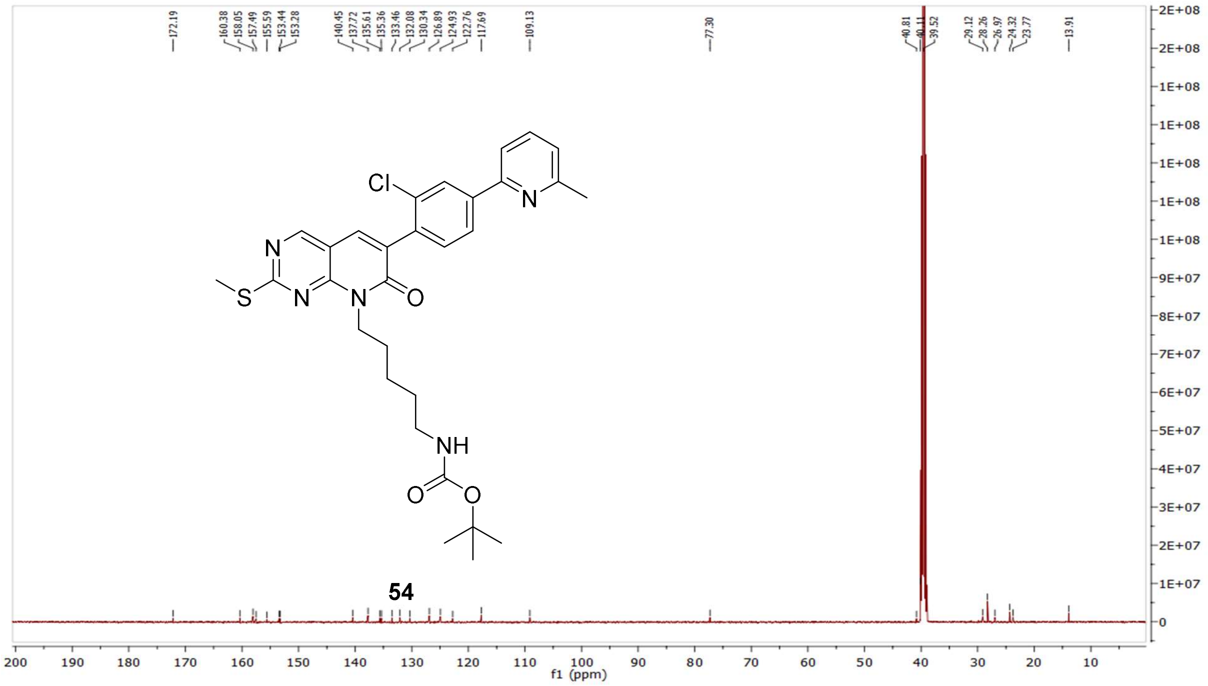

MS (ESI+) compound **54**:

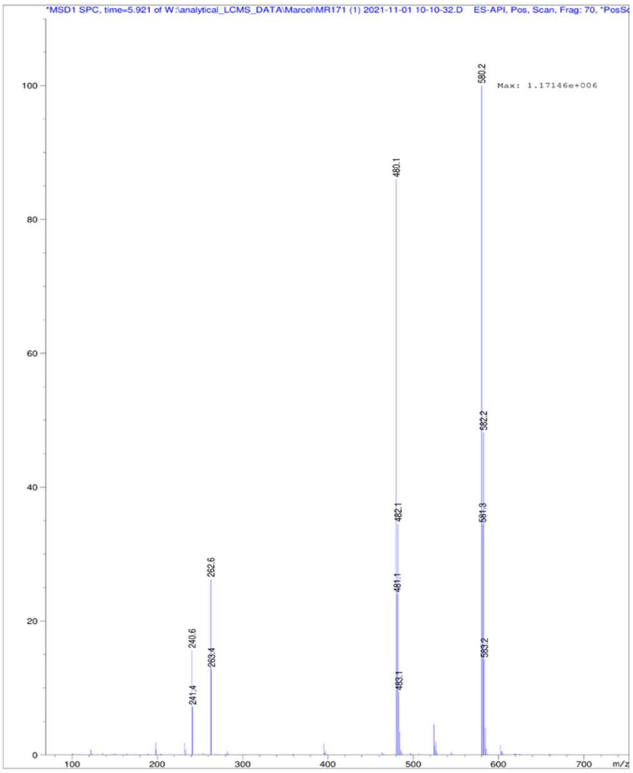


$^{1}$H NMR compound **55**:

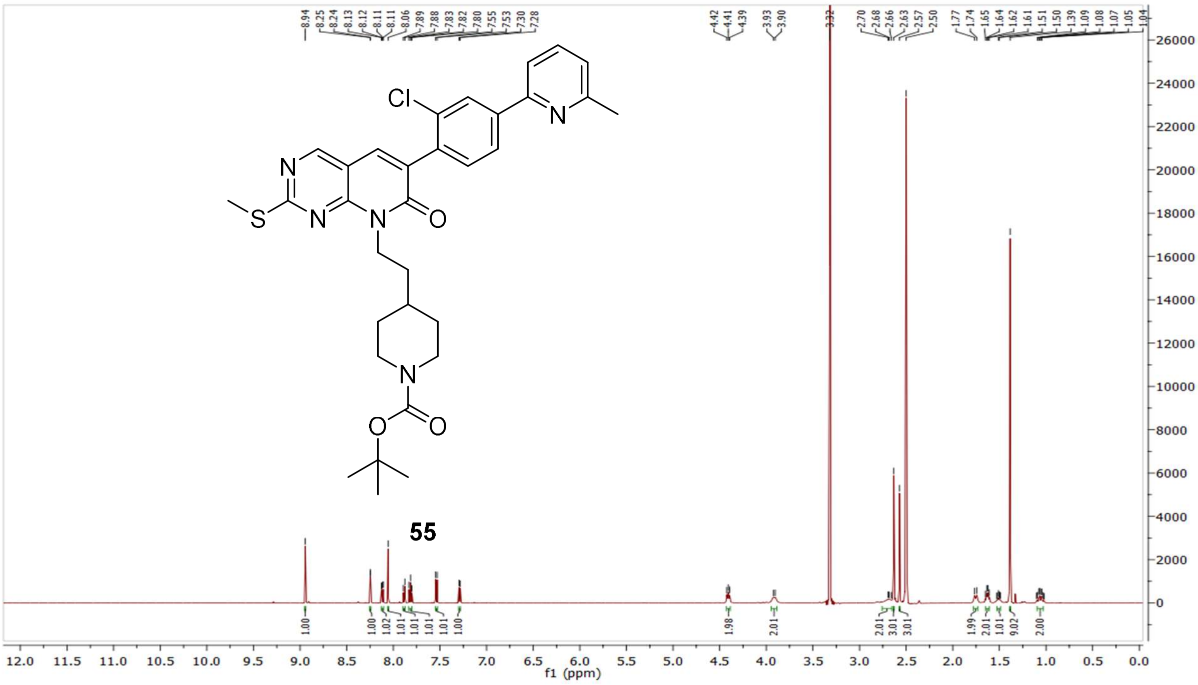

$^{13}$C NMR compound **55:**

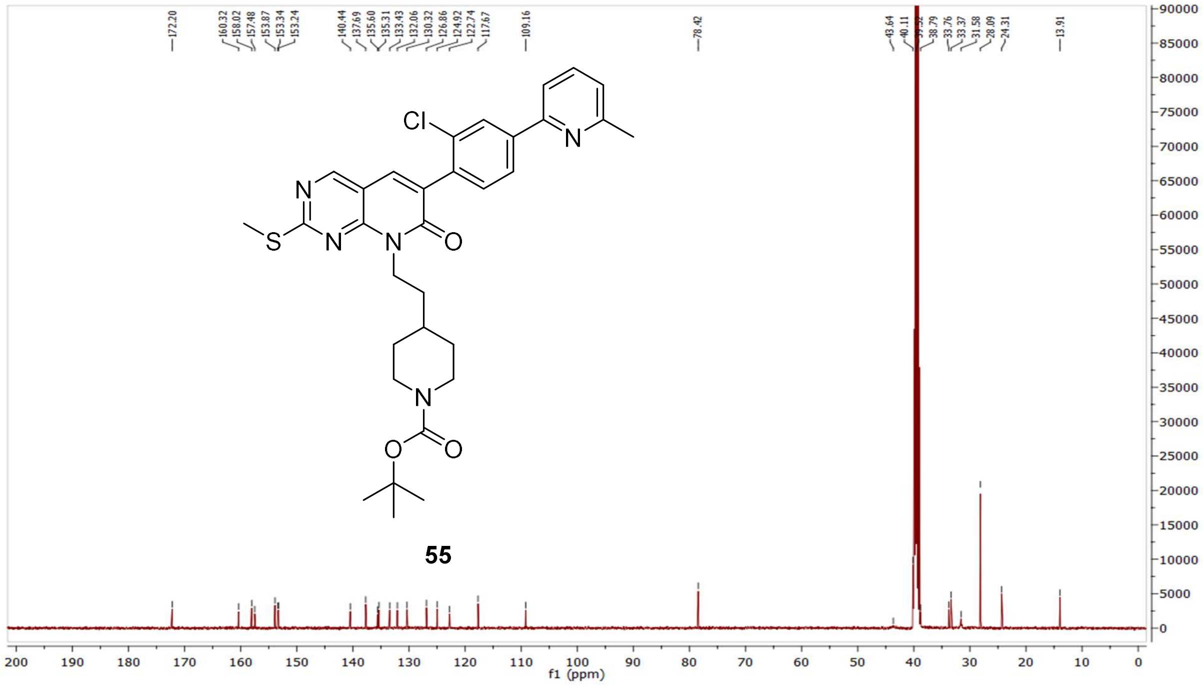

MS (ESI+) compound **55**:

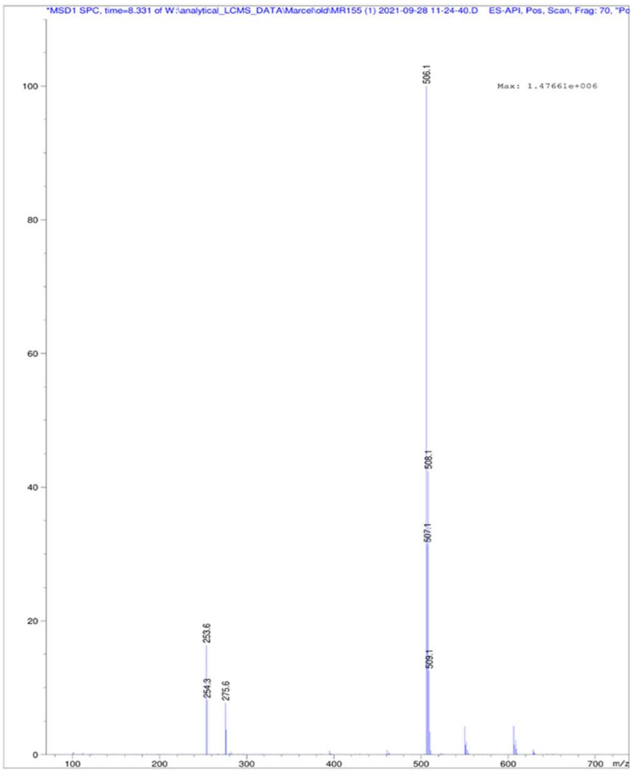

<sup></sup>

[1]H NMR compound **56**:

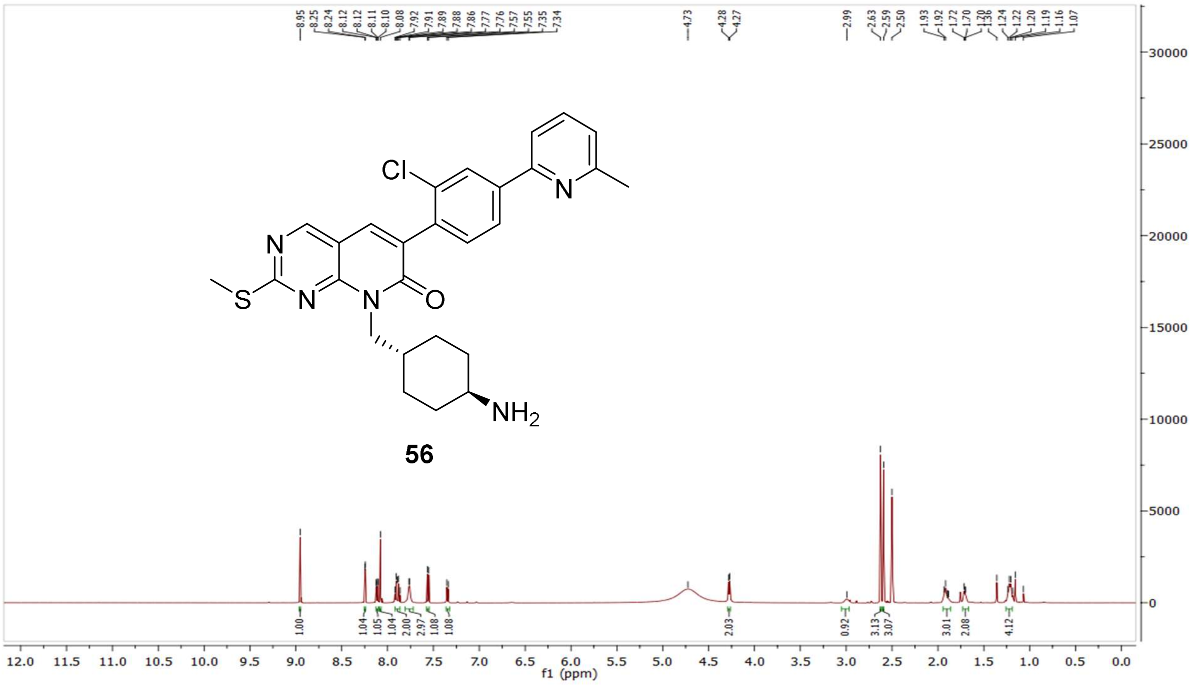


[13]C NMR compound **56:**

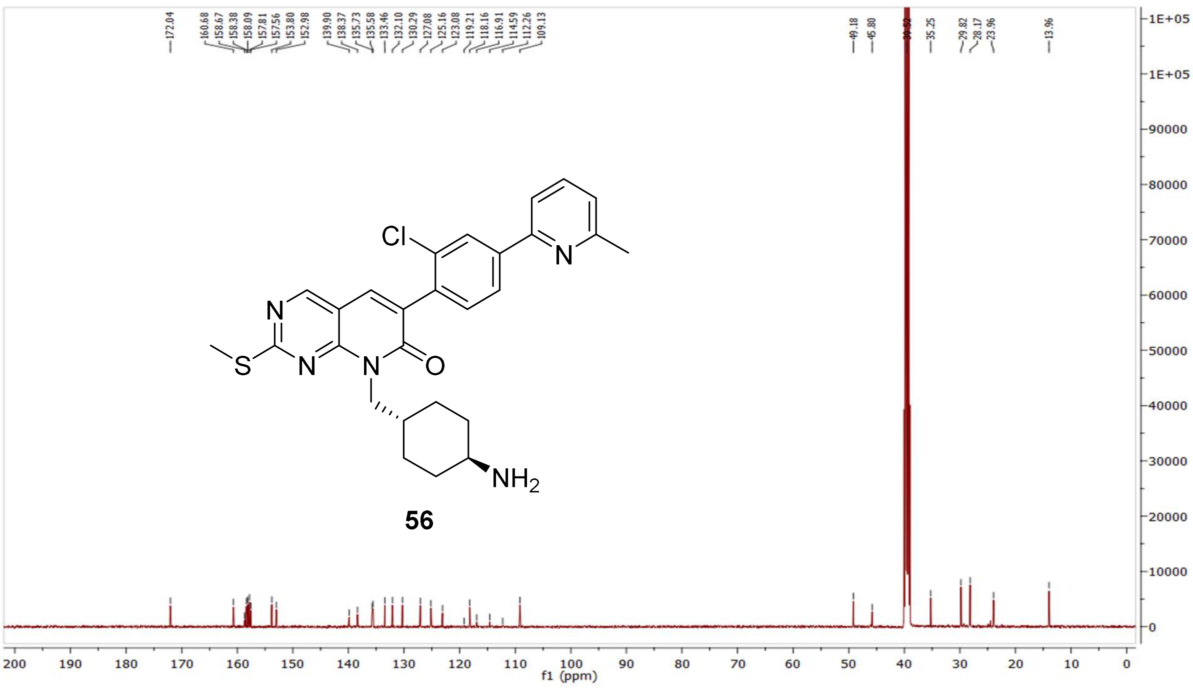

MS (ESI+) compound **56**:

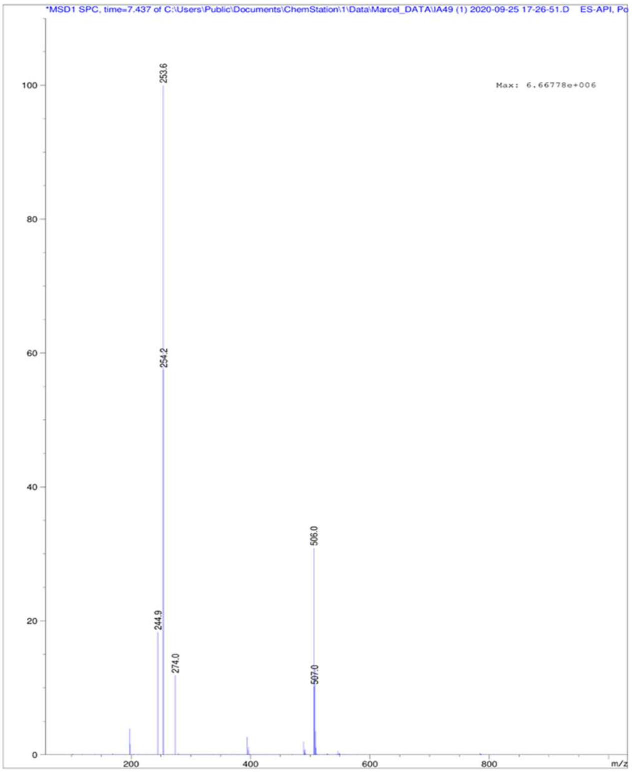

[1]H NMR compound **57**:

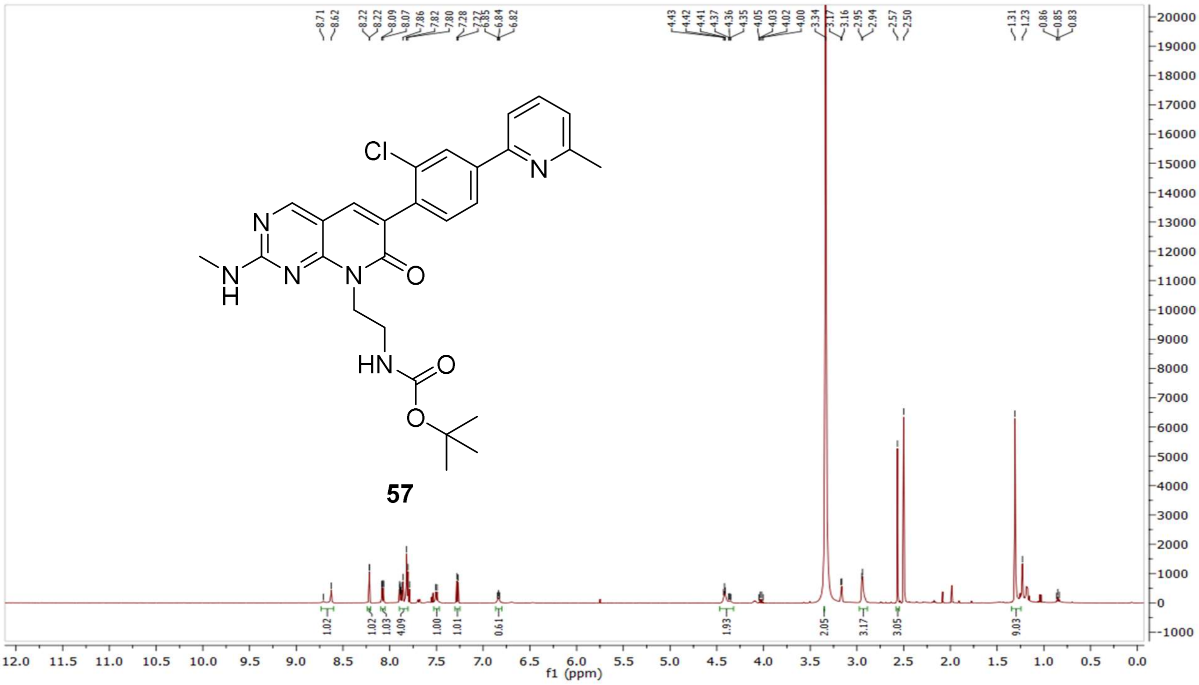

$^{13}$C NMR compound **57:**

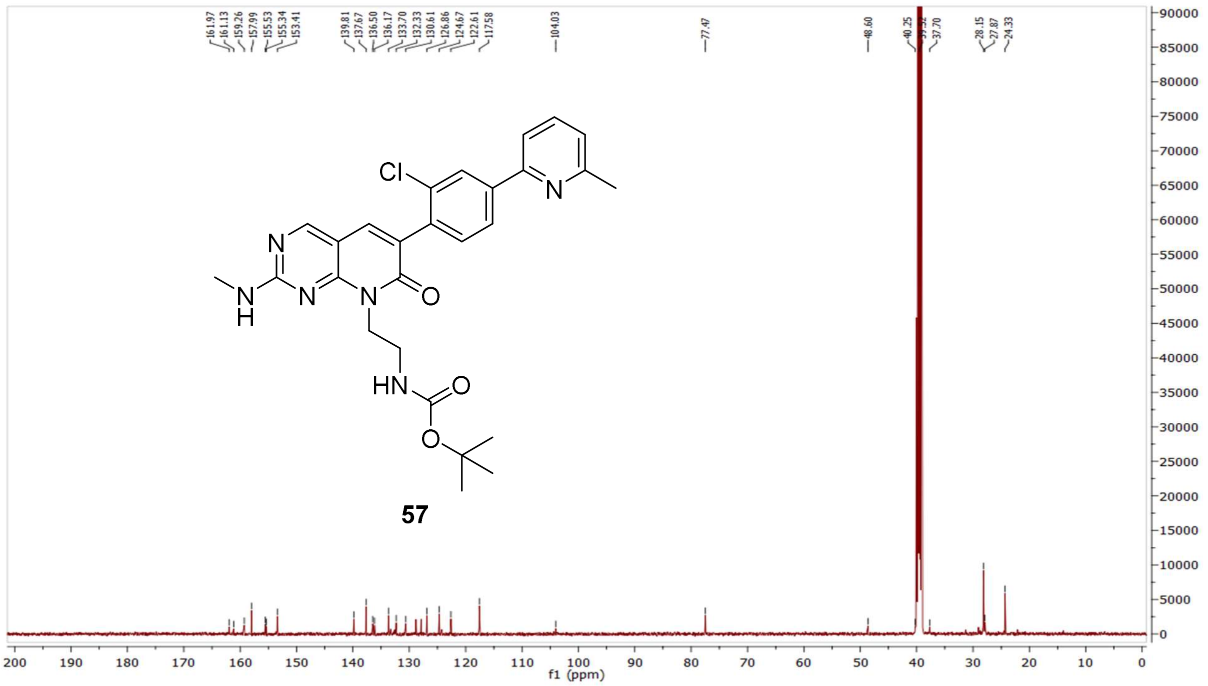

MS (ESI+) compound **57**:

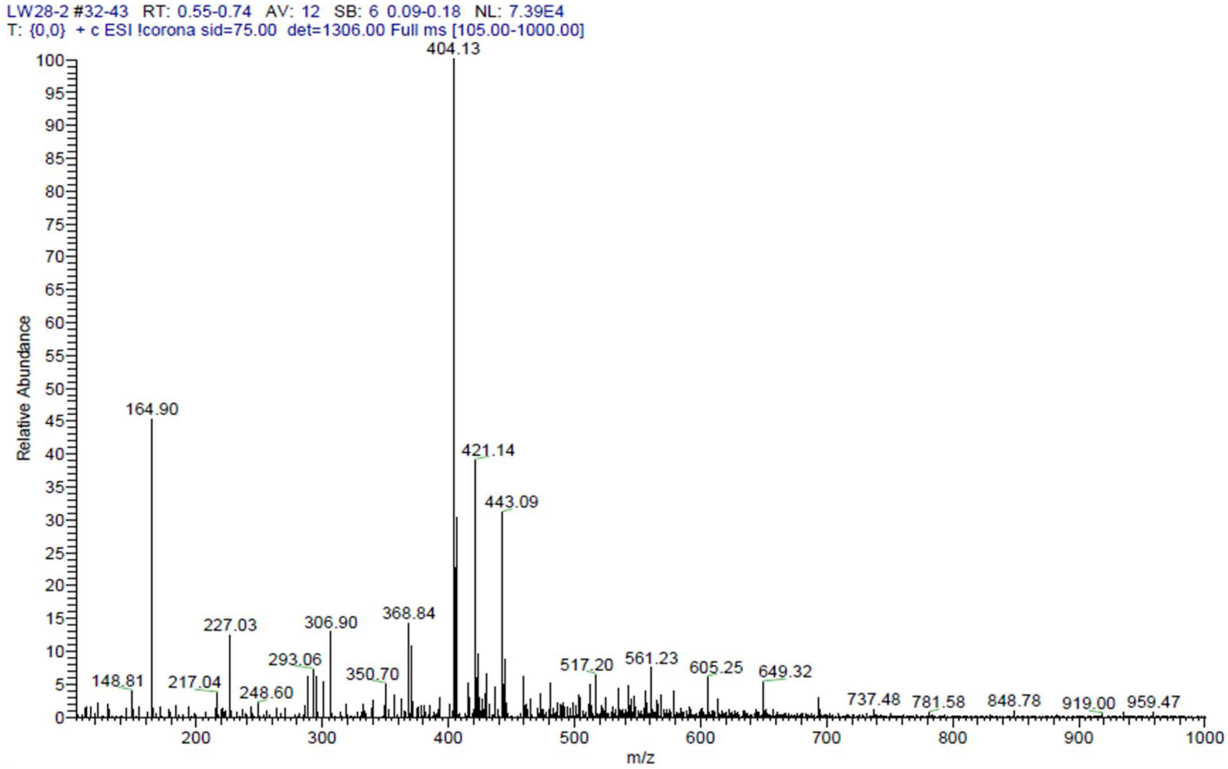

$^{1}$H NMR compound **58**:

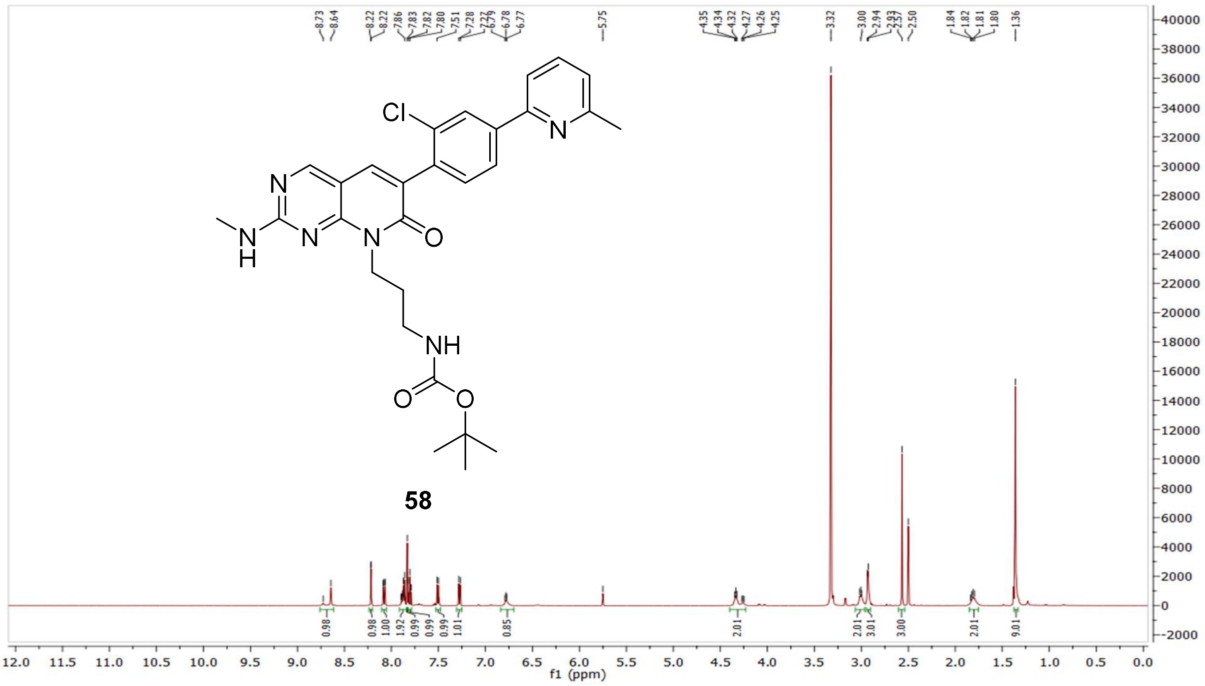

$^{13}$C NMR compound **58:**

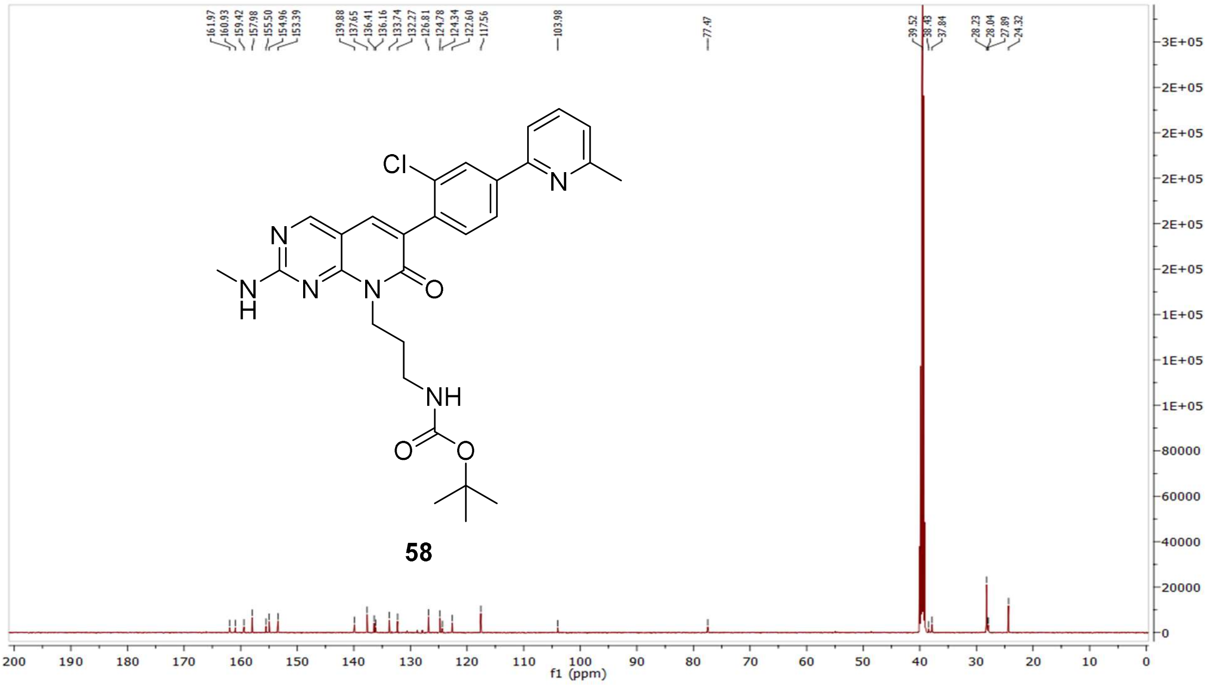

MS (ESI+) compound **58**:

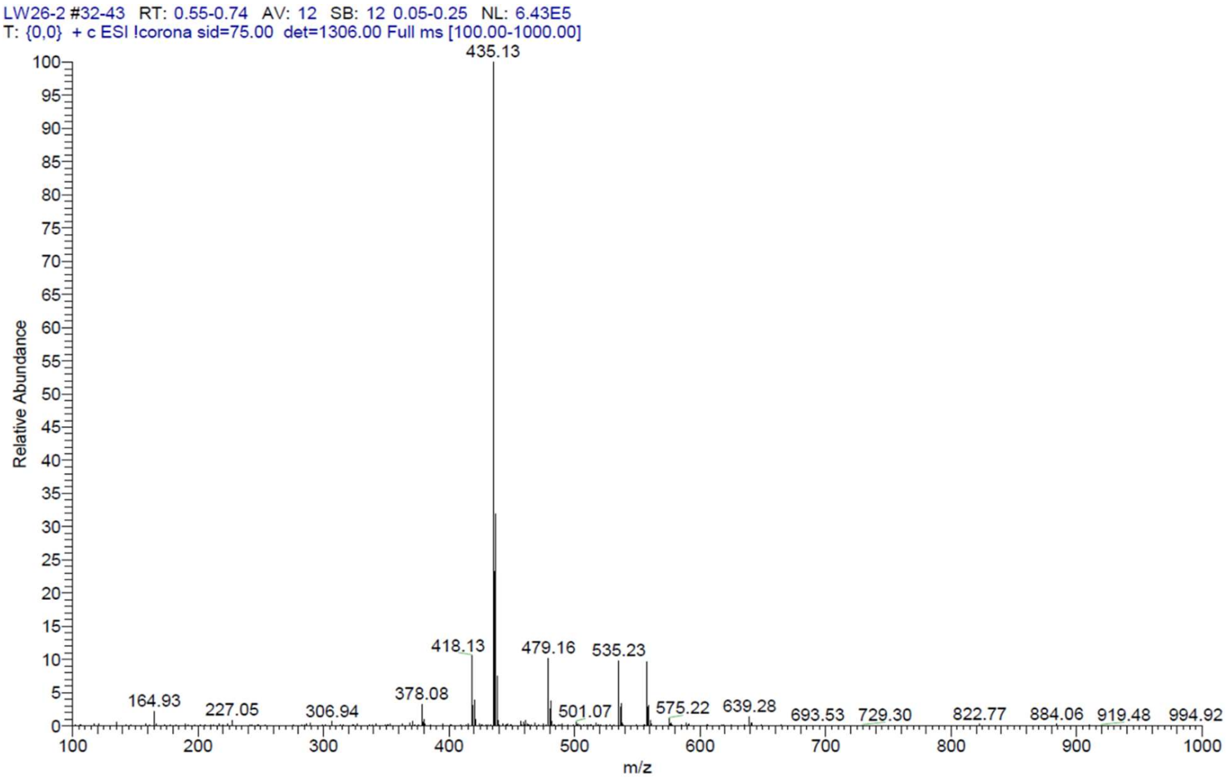


$^1$H NMR compound **59**:

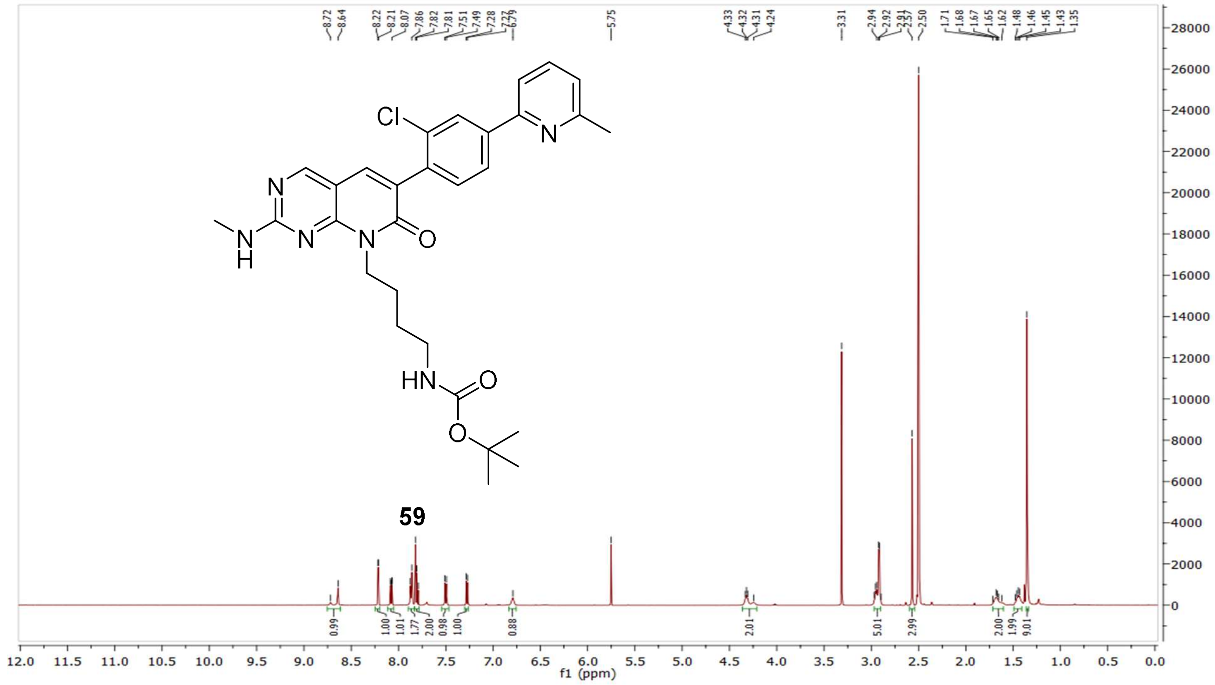

$^{13}C$ NMR compound **59:**

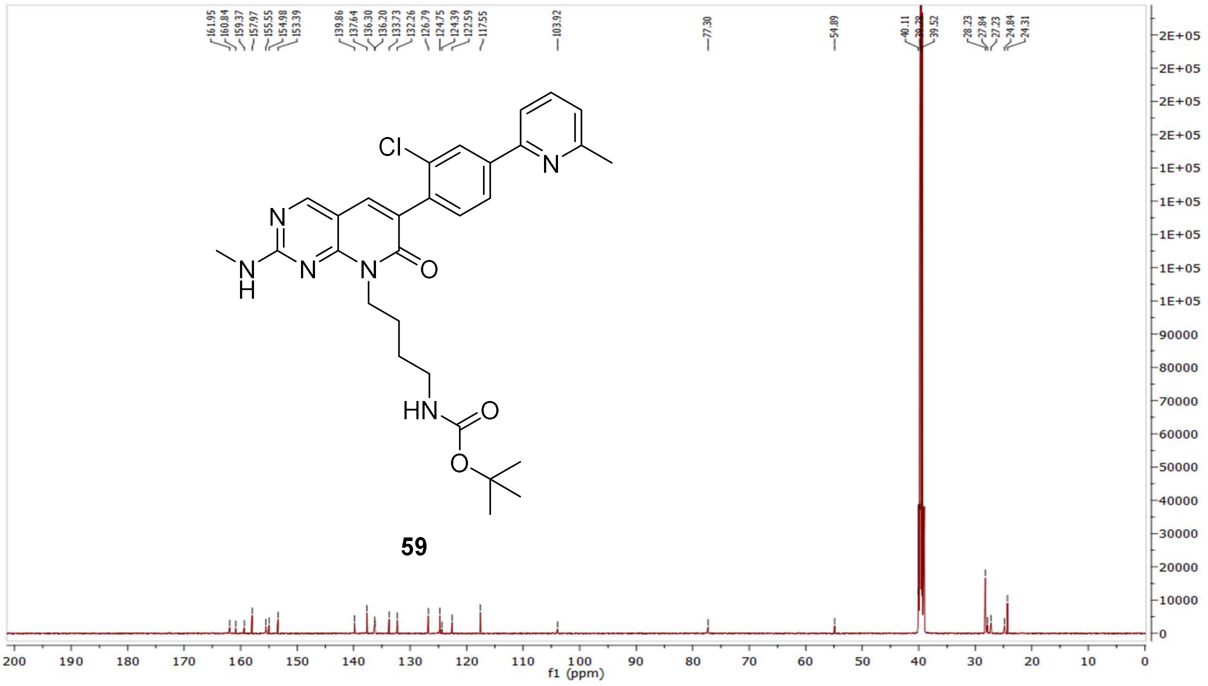

MS (ESI+) compound **59**:

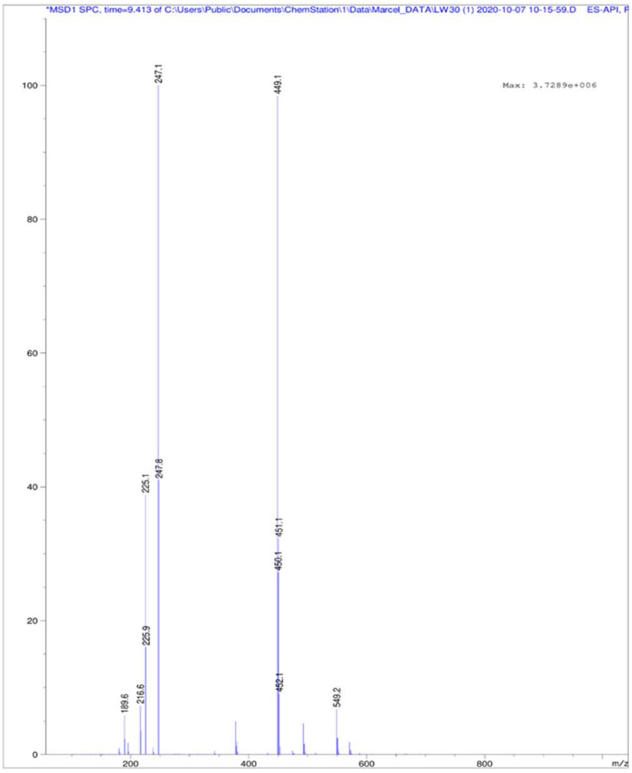

<sup></sup>[1]H NMR compound **60**:

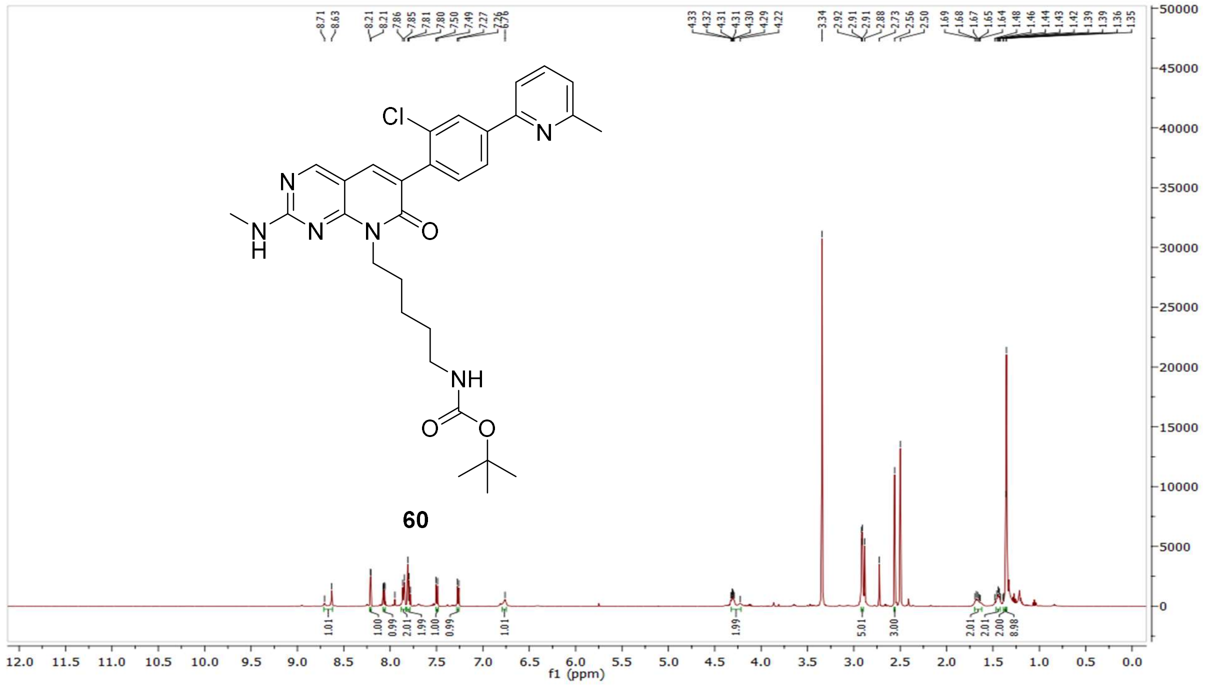

[13]C NMR compound **60:**

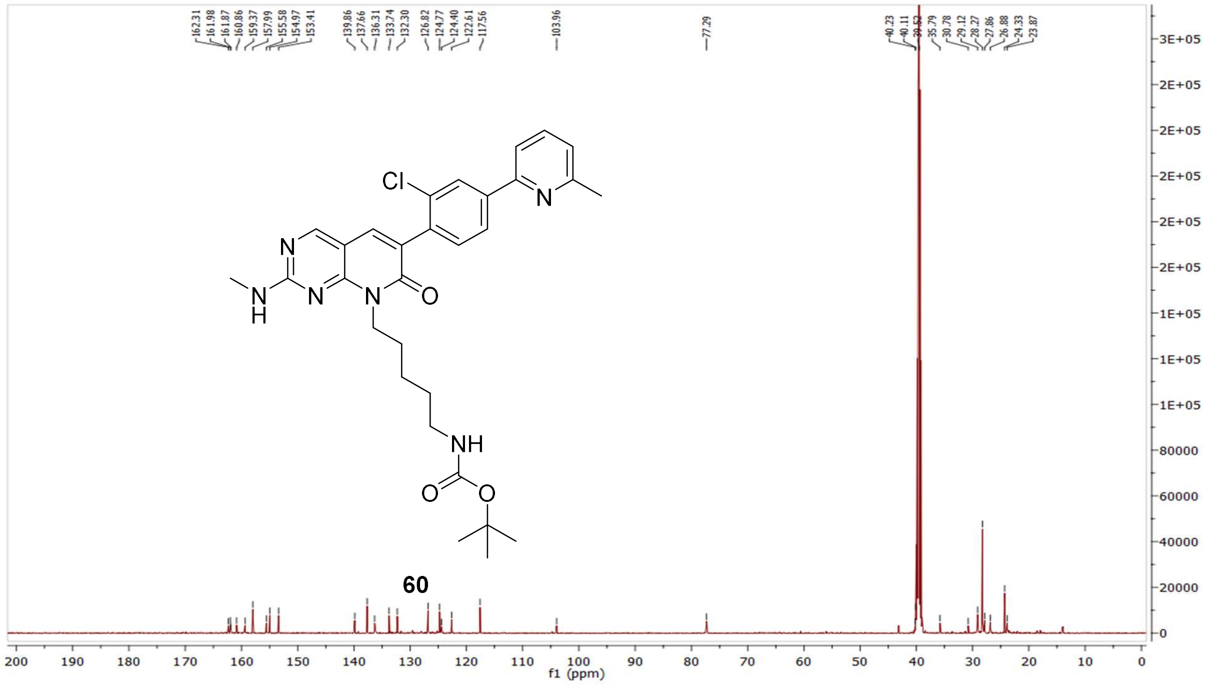

MS (ESI+) compound **60**:

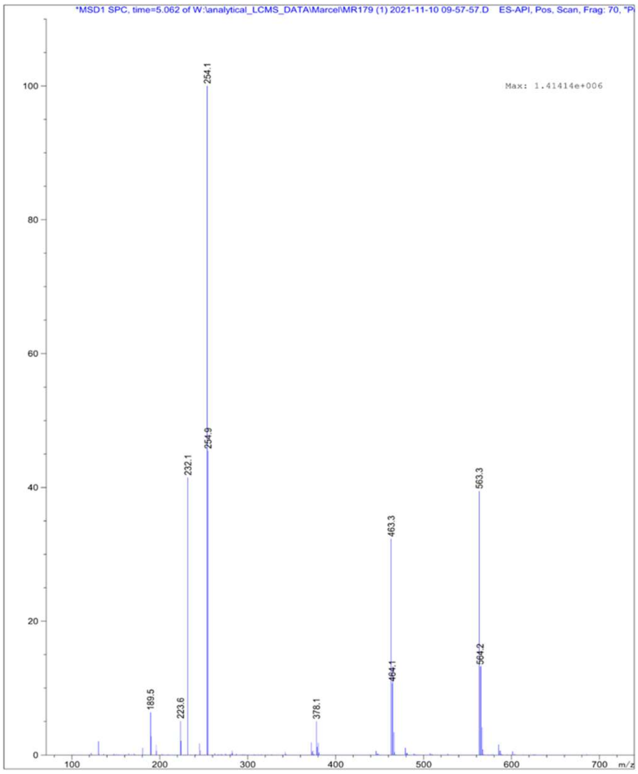


$^{1}$H NMR compound **61**:

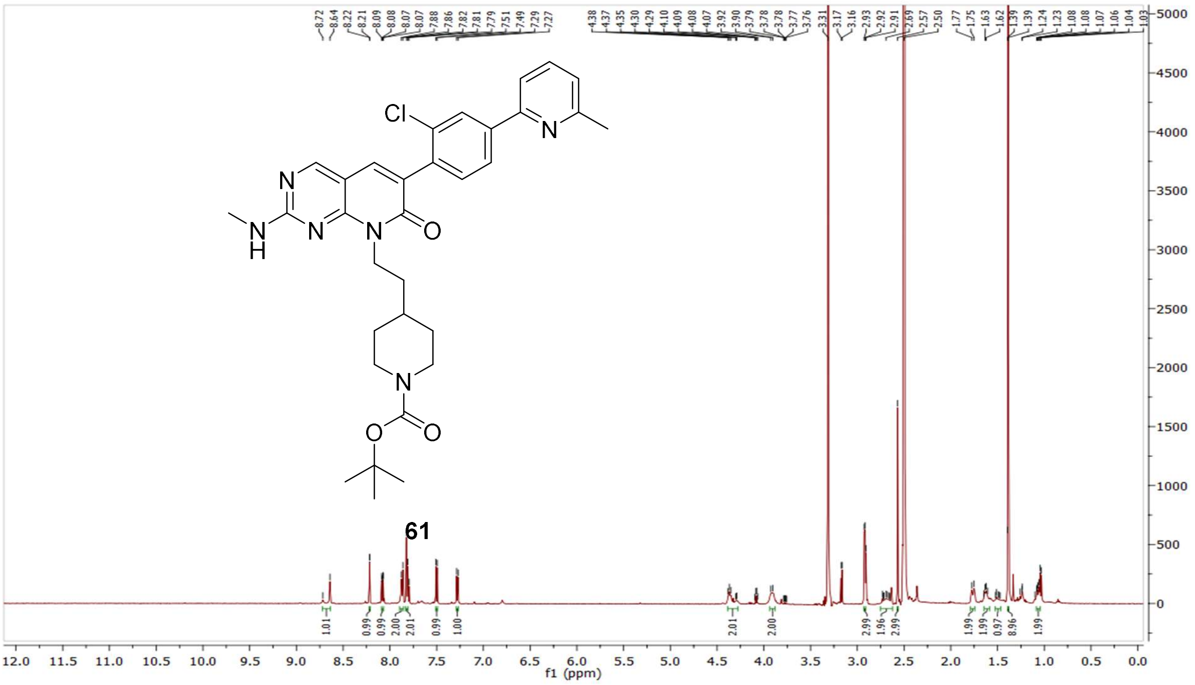

$^{13}$C NMR compound **61:**

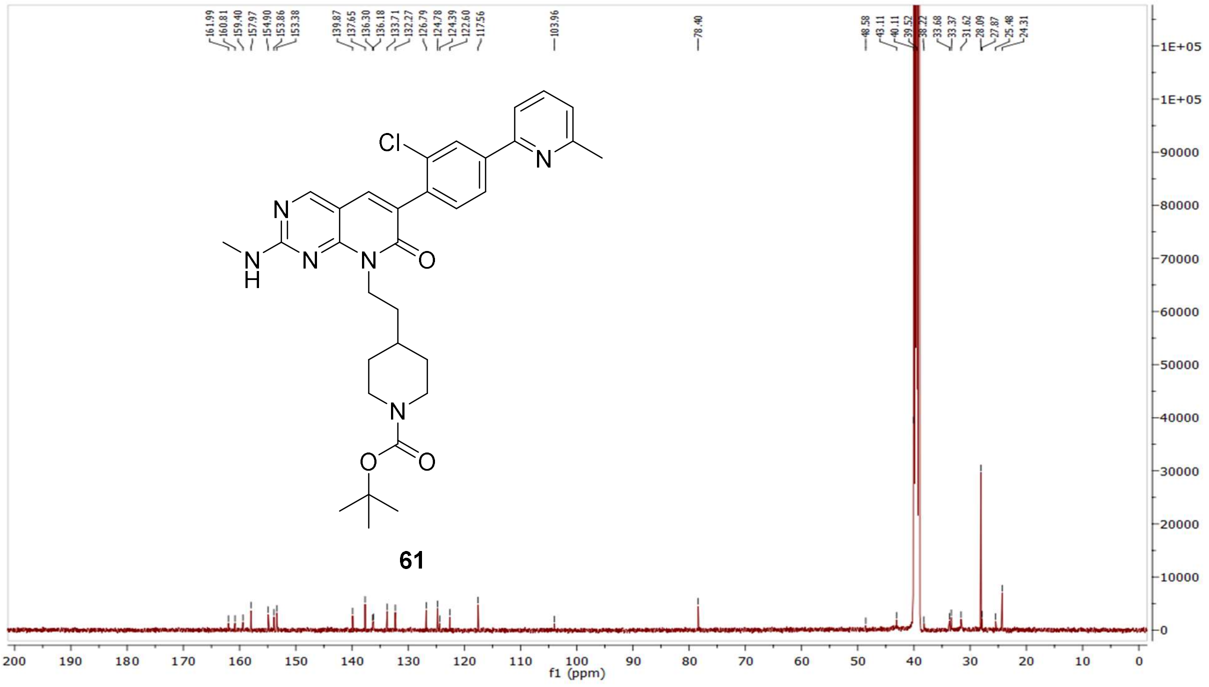


MS (ESI+) compound **61**:

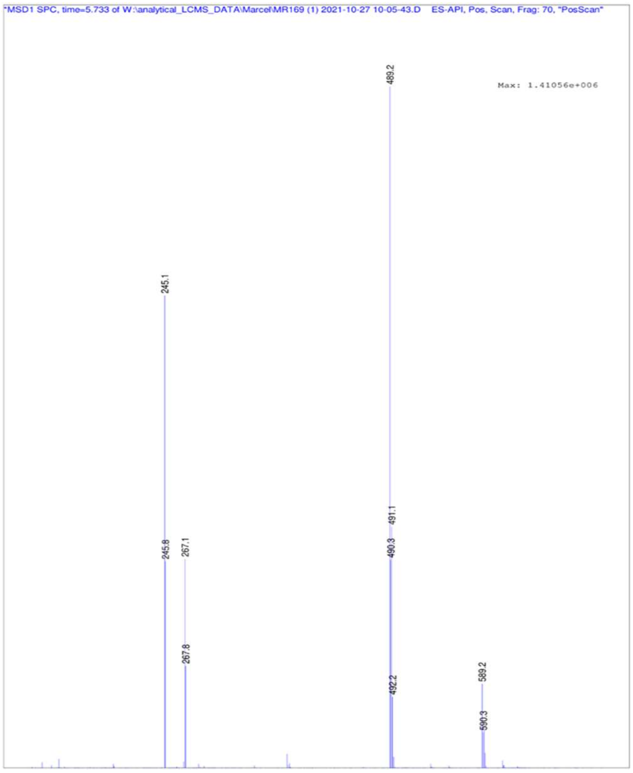

$^{1}H$ NMR compound **62**:

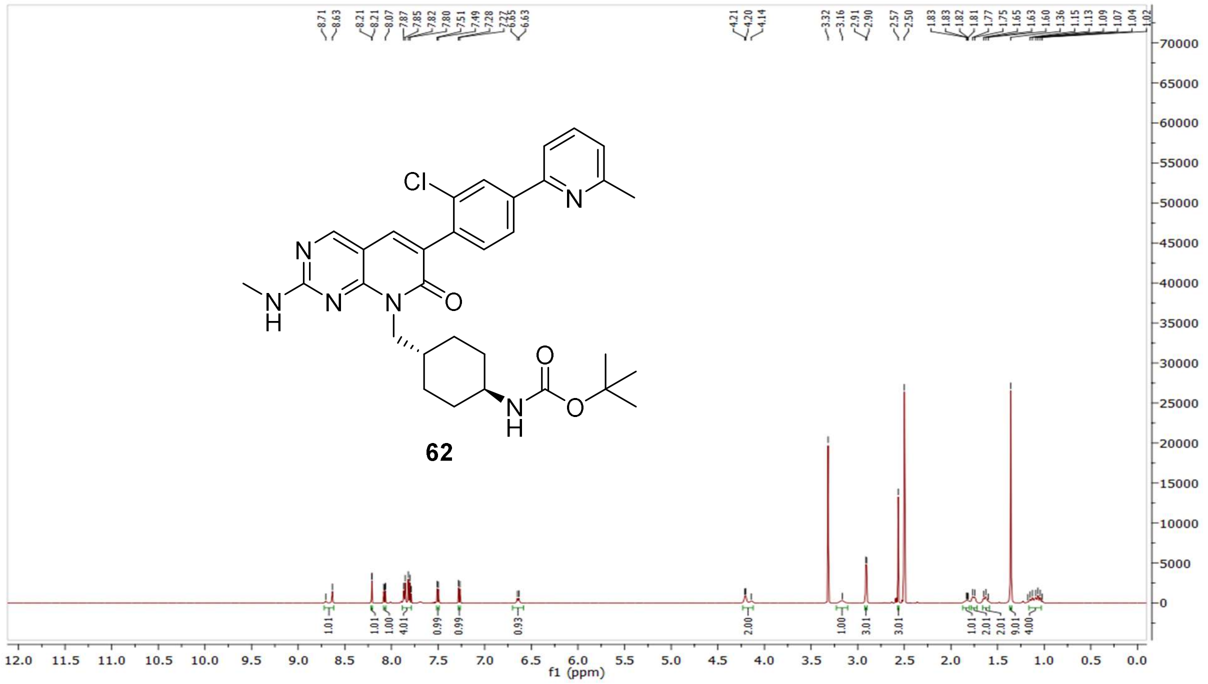

$^{13}C$ NMR compound **62:**

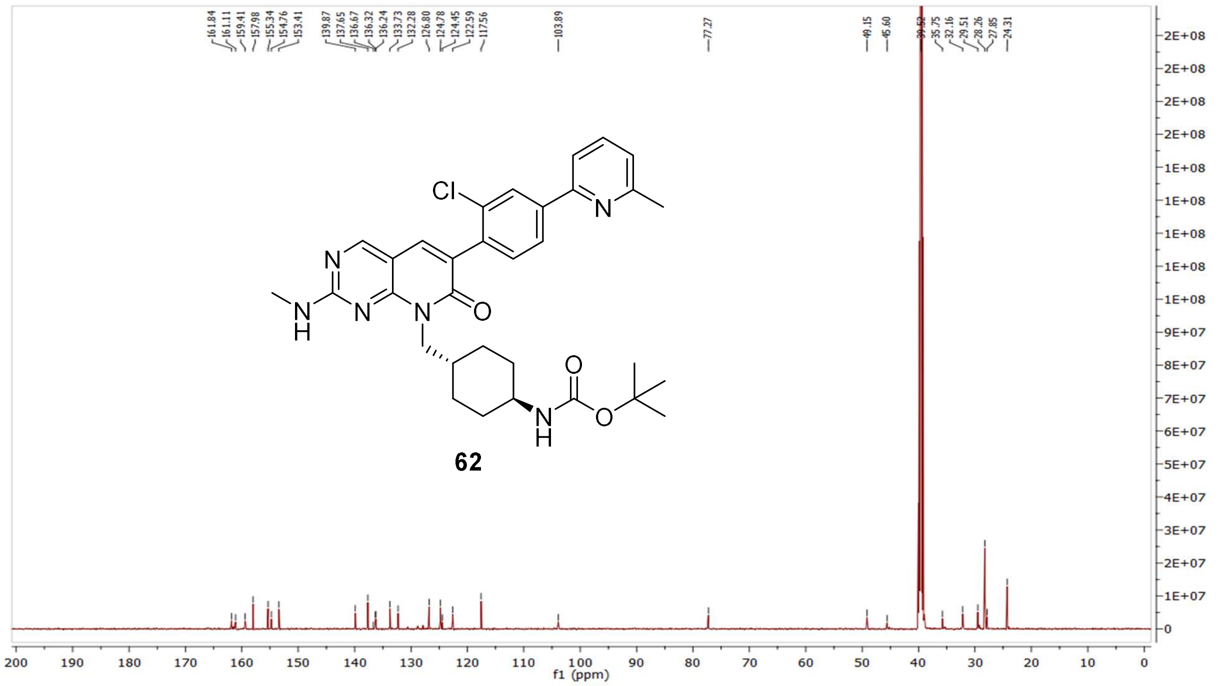

MS (ESI+) compound **62**:

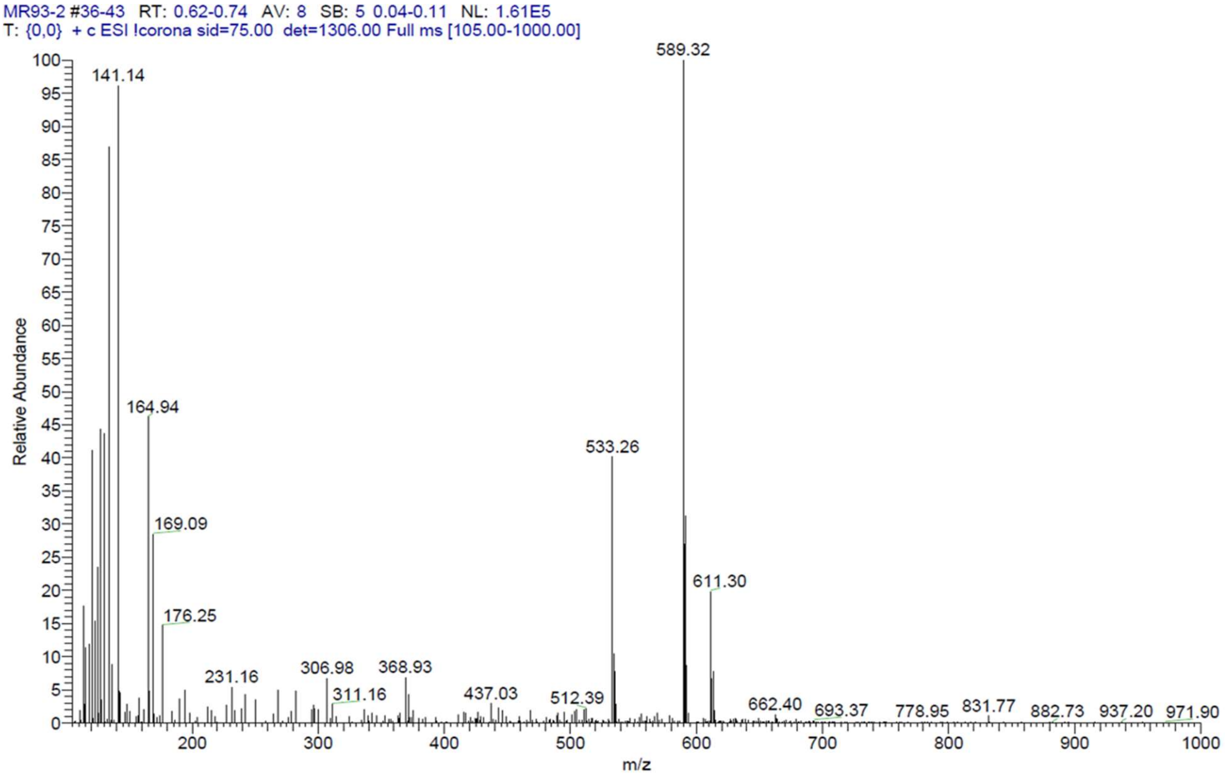


$^{1}H$ NMR compound **18**:

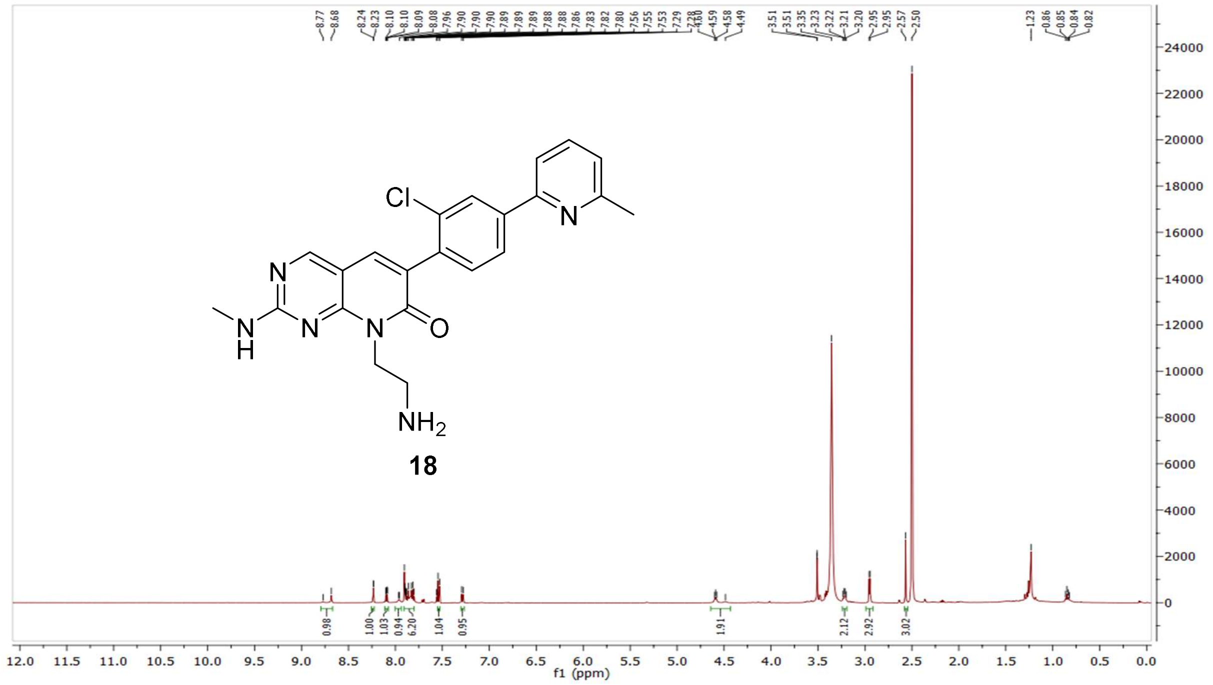

$^{13}C$ NMR compound **18:**

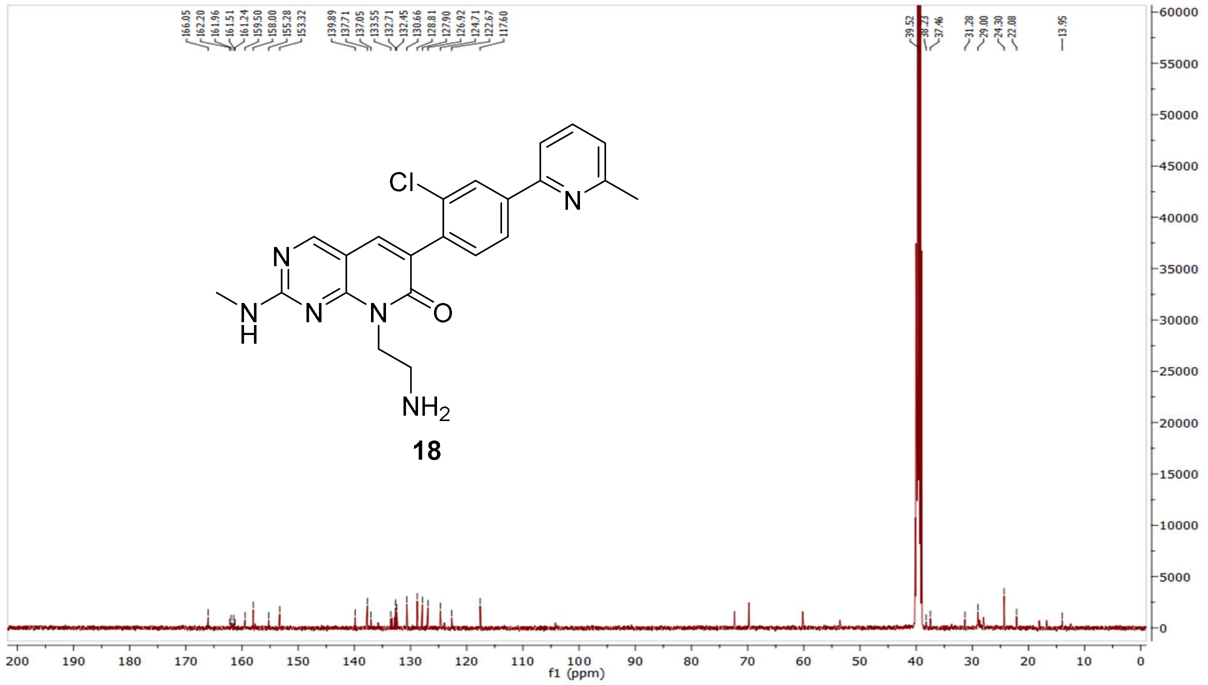


HRMS (FTMS +p MALDI) compound **18**:

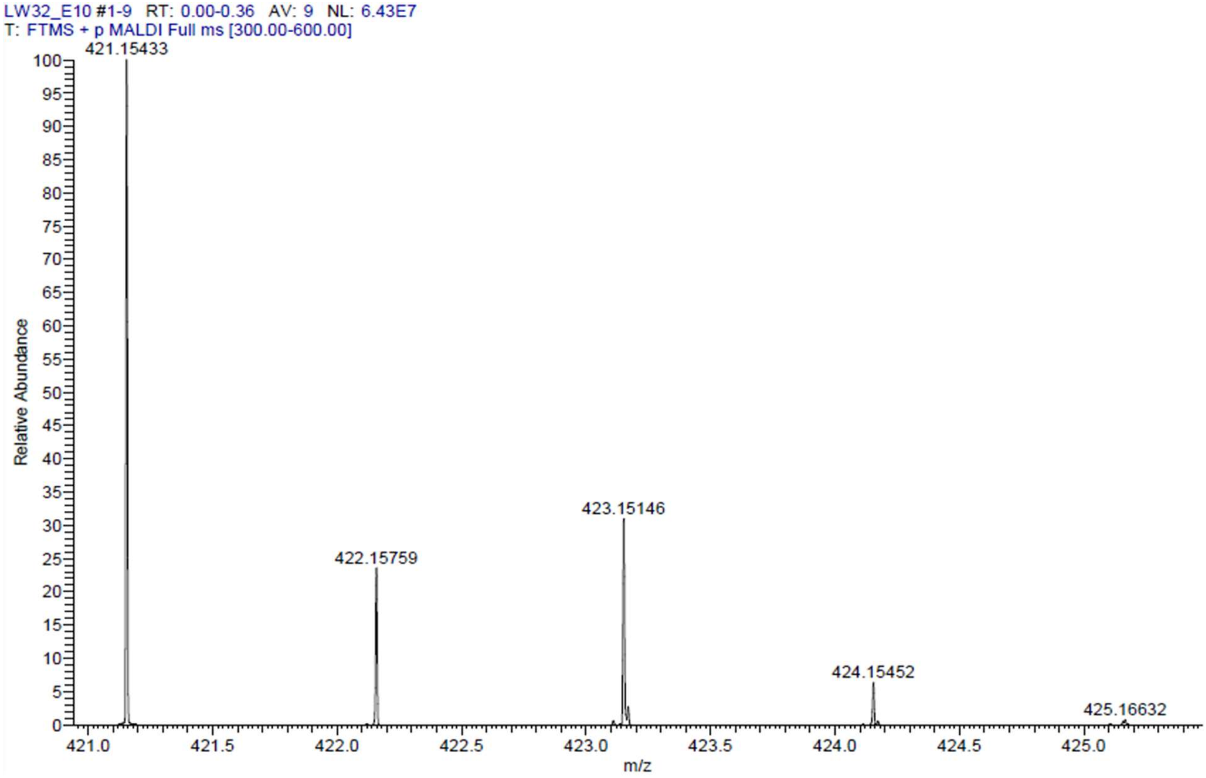

HPLC-chromatograms compound **18**:

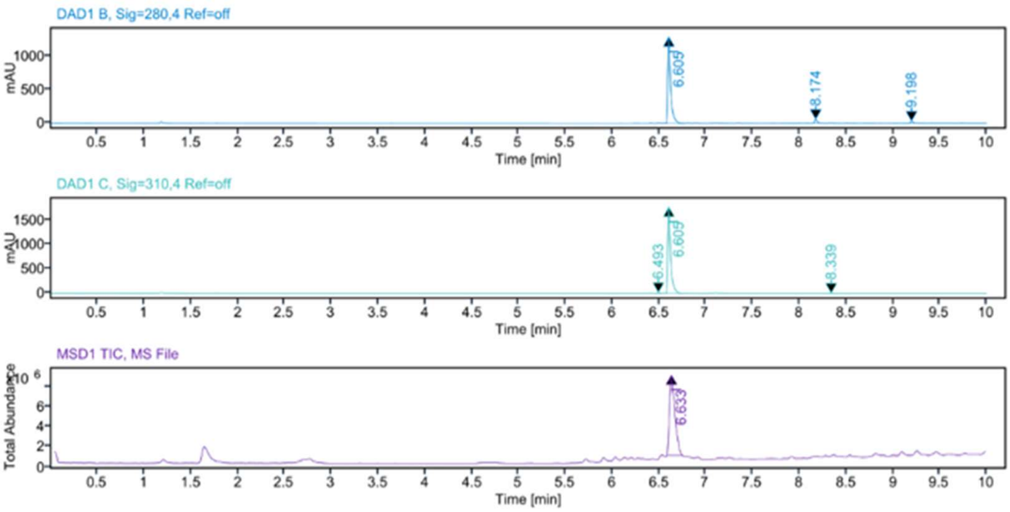


**Sample Purity**

Signal Description DAD1 B, Sig=280,4 Ref=off

| Sample Name | Name | RT | Width | Area | Area% | Height |
|---|---|---|---|---|---|---|
| LW32 | | 6.605 | 0.029 | 2721.5383 | 95.04 | 1292.3483 |
| LW32 | | 8.174 | 0.030 | 98.4081 | 3.44 | 51.7957 |
| LW32 | | 9.198 | 0.030 | 43.7584 | 1.53 | 24.0459 |

Max Area% 95.036

UV Signal Purity>95% Pass

Signal Description DAD1 C, Sig=310,4 Ref=off

| Sample Name | Name | RT | Width | Area | Area% | Height |
|---|---|---|---|---|---|---|
| LW32 | | 6.493 | 0.023 | 25.2417 | 0.66 | 17.1999 |
| LW32 | | 6.605 | 0.029 | 3786.9192 | 98.85 | 1784.2592 |
| LW32 | | 8.339 | 0.045 | 18.7843 | 0.49 | 5.5286 |

Max Area% 98.851

UV Signal Purity>95% Pass

**MSD Apex Spectra**

**RT:**

6.633

**Sample Name:**

LW32

**Signal Name:**

MSD1TIC

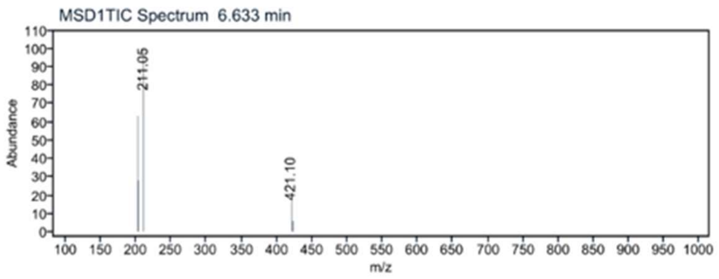

$^1$H NMR compound **19**:

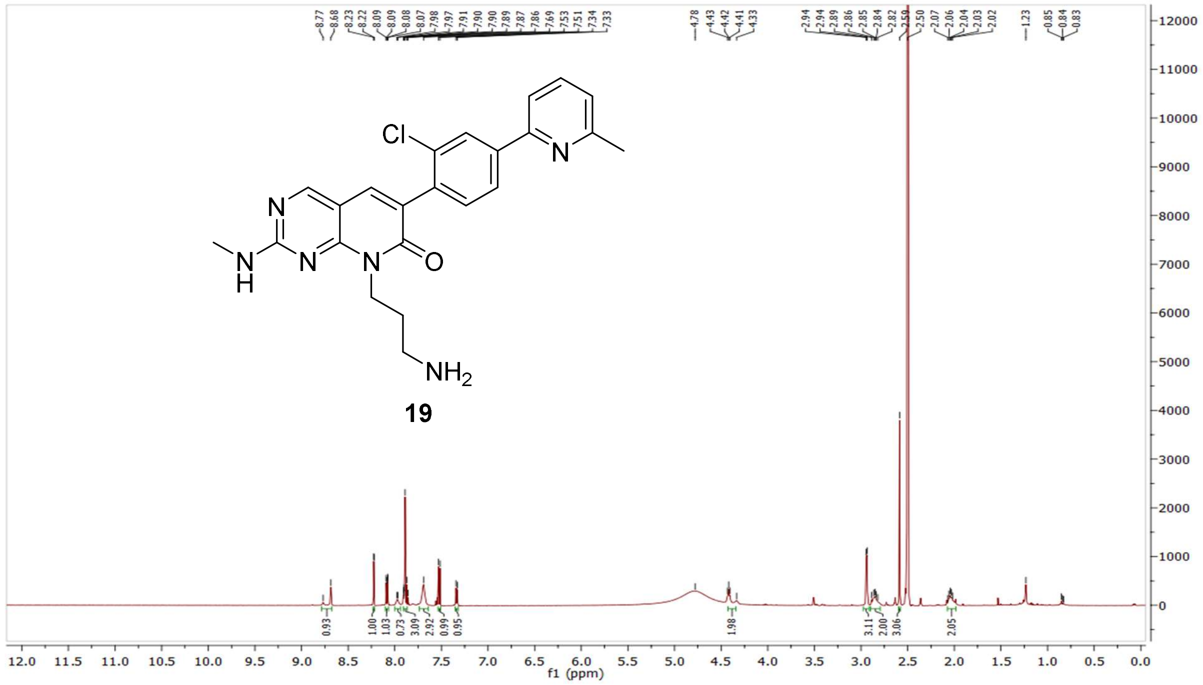

$^{13}$C NMR compound **19:**

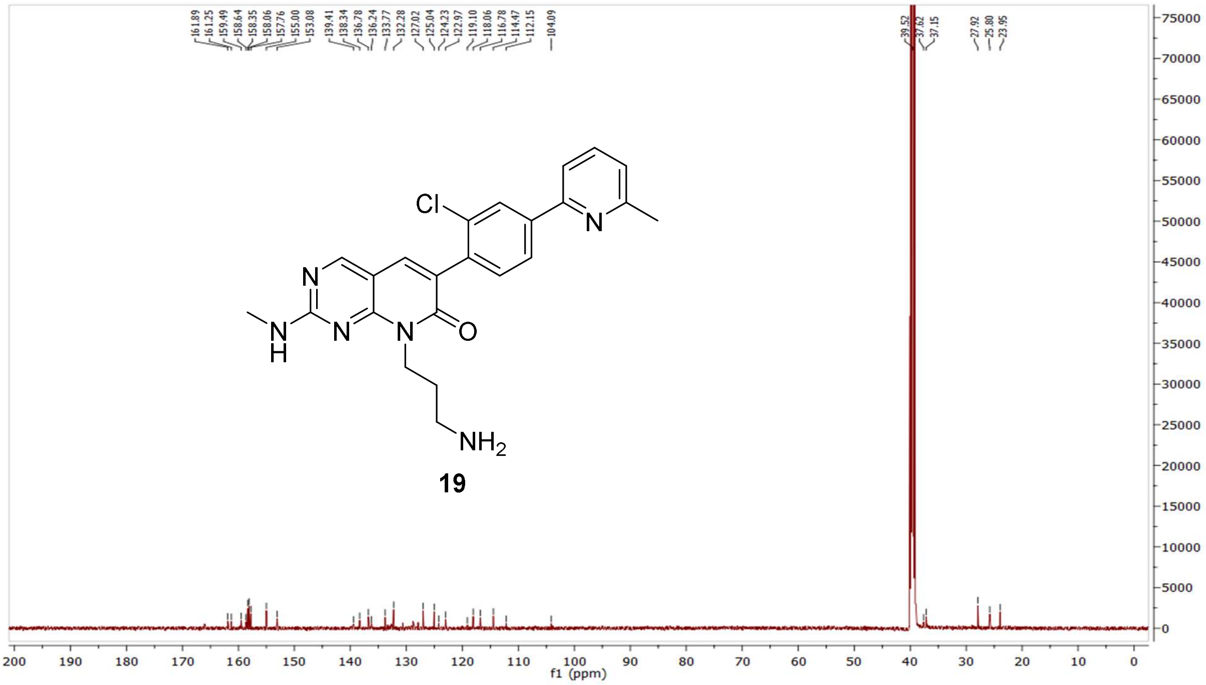

HRMS (FTMS +p MALDI) compound **19**:

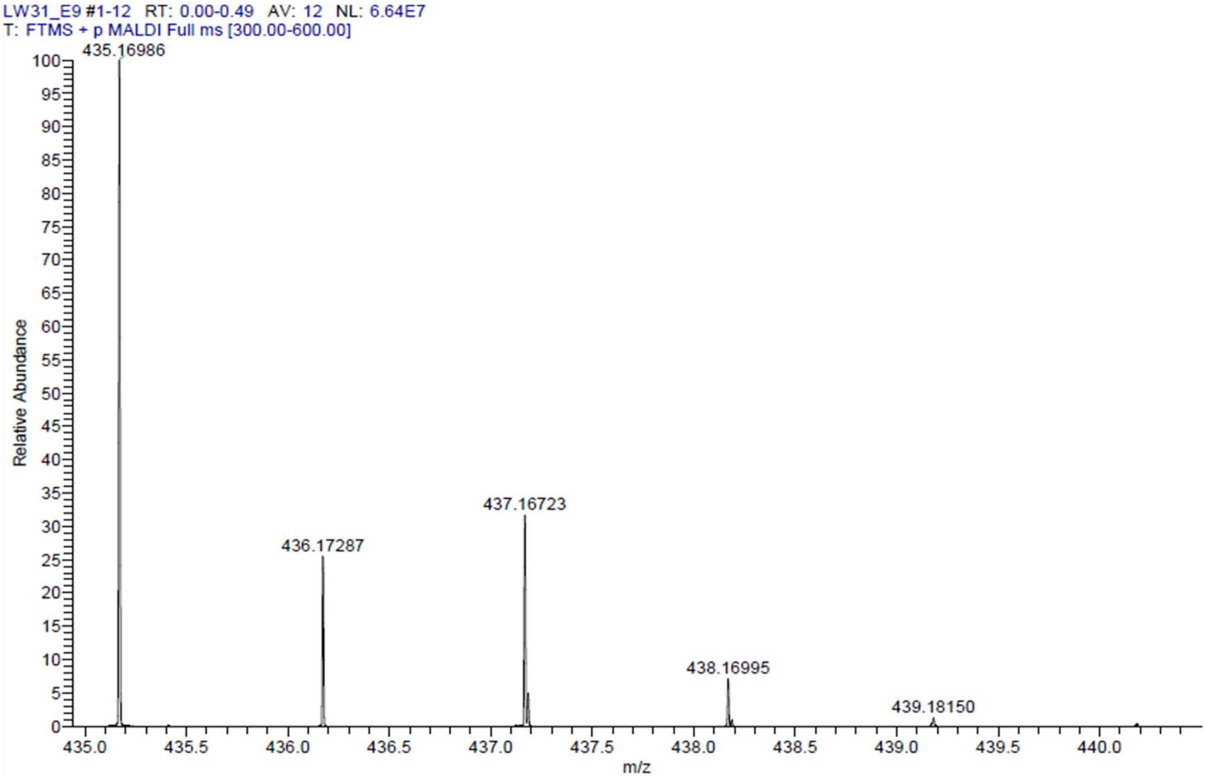

HPLC-chromatograms compound **19**:

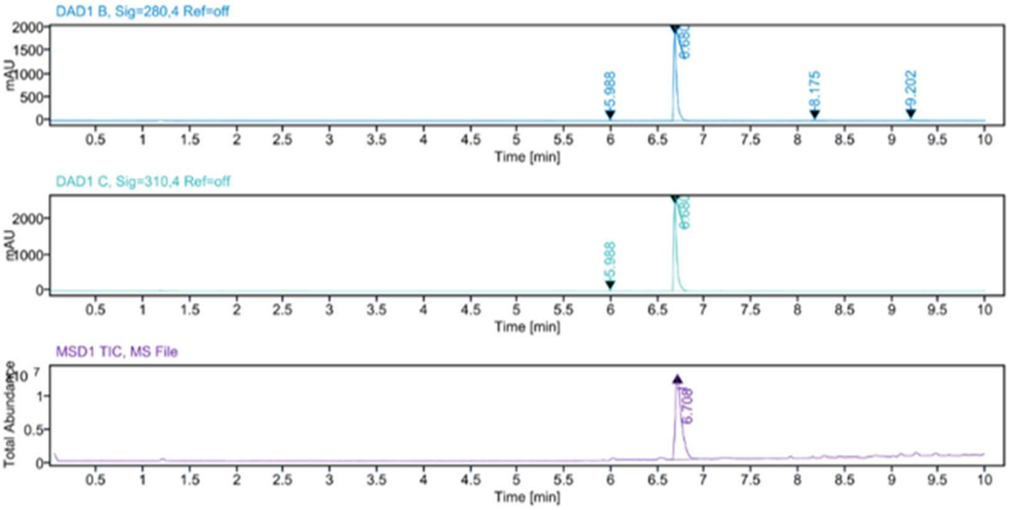


**Sample Purity**

Signal Description DAD1 B, Sig=280,4 Ref=off

| Sample Name | Name | RT | Width | Area | Area% | Height |
|---|---|---|---|---|---|---|
| LW31 | | 5.988 | 0.023 | 22.2106 | 0.51 | 12.7487 |
| LW31 | | 6.680 | 0.032 | 4184.0444 | 96.30 | 1829.5145 |
| LW31 | | 8.175 | 0.032 | 42.6073 | 0.98 | 17.8641 |
| LW31 | | 9.202 | 0.037 | 96.1437 | 2.21 | 33.9232 |

Max Area% 96.295
UV Signal Purity>95% Pass

Signal Description DAD1 C, Sig=310,4 Ref=off

| Sample Name | Name | RT | Width | Area | Area% | Height |
|---|---|---|---|---|---|---|
| LW31 | | 5.988 | 0.023 | 23.0060 | 0.41 | 13.1883 |
| LW31 | | 6.680 | 0.032 | 5545.7437 | 99.59 | 2405.1753 |

Max Area% 99.587
UV Signal Purity>95% Pass

**MSD Apex Spectra**

**RT:**

6.708

**Sample Name:**

LW31

**Signal Name:**

MSD1TIC

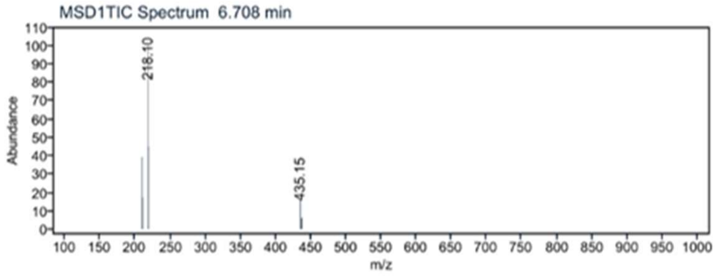

$^{1}$H NMR compound **20**:

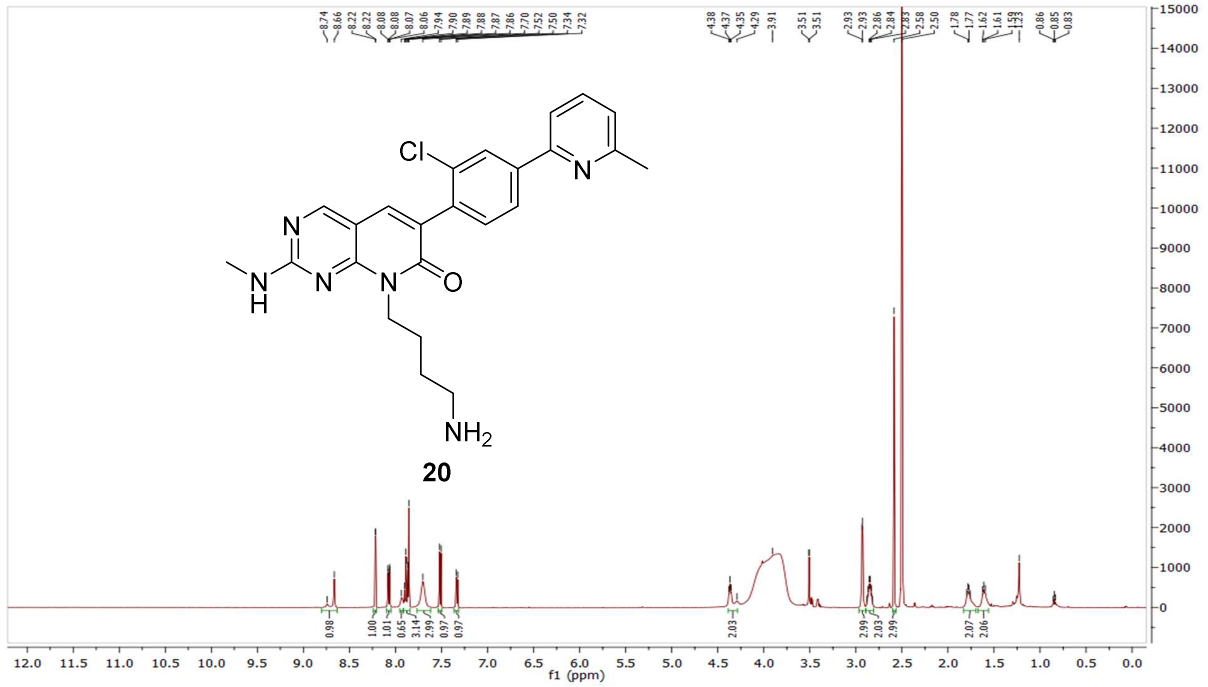


$^{13}$C NMR compound **20:**

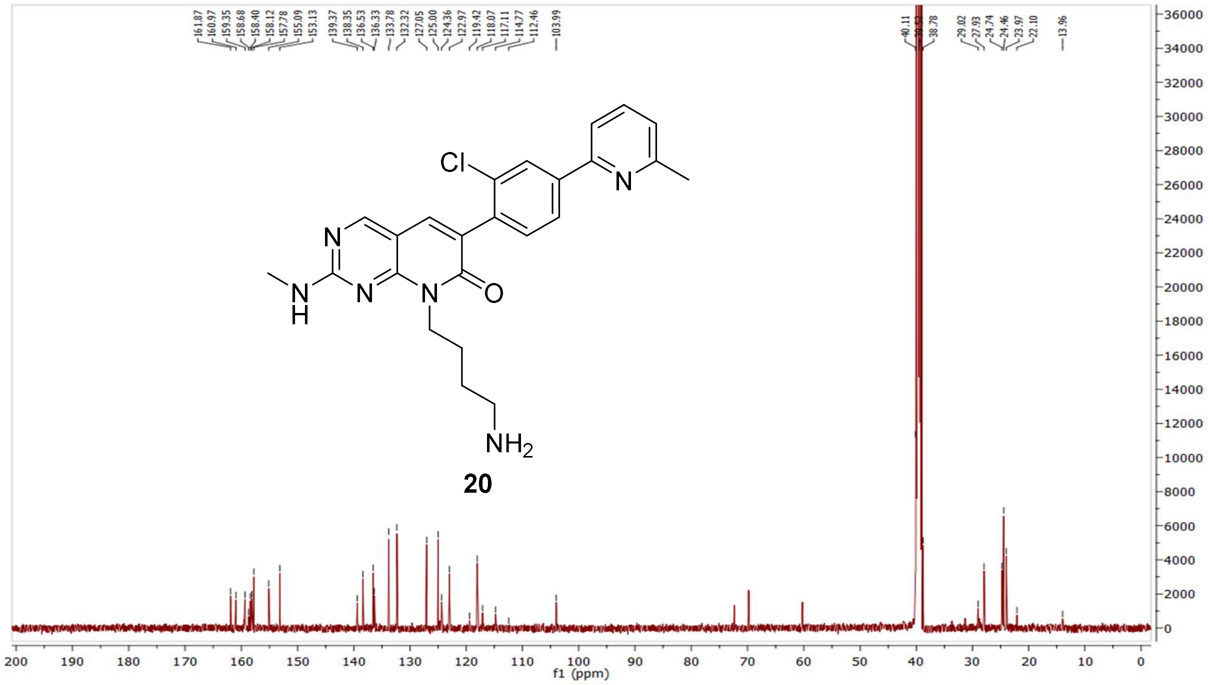

HRMS (FTMS +p MALDI) compound **20**:

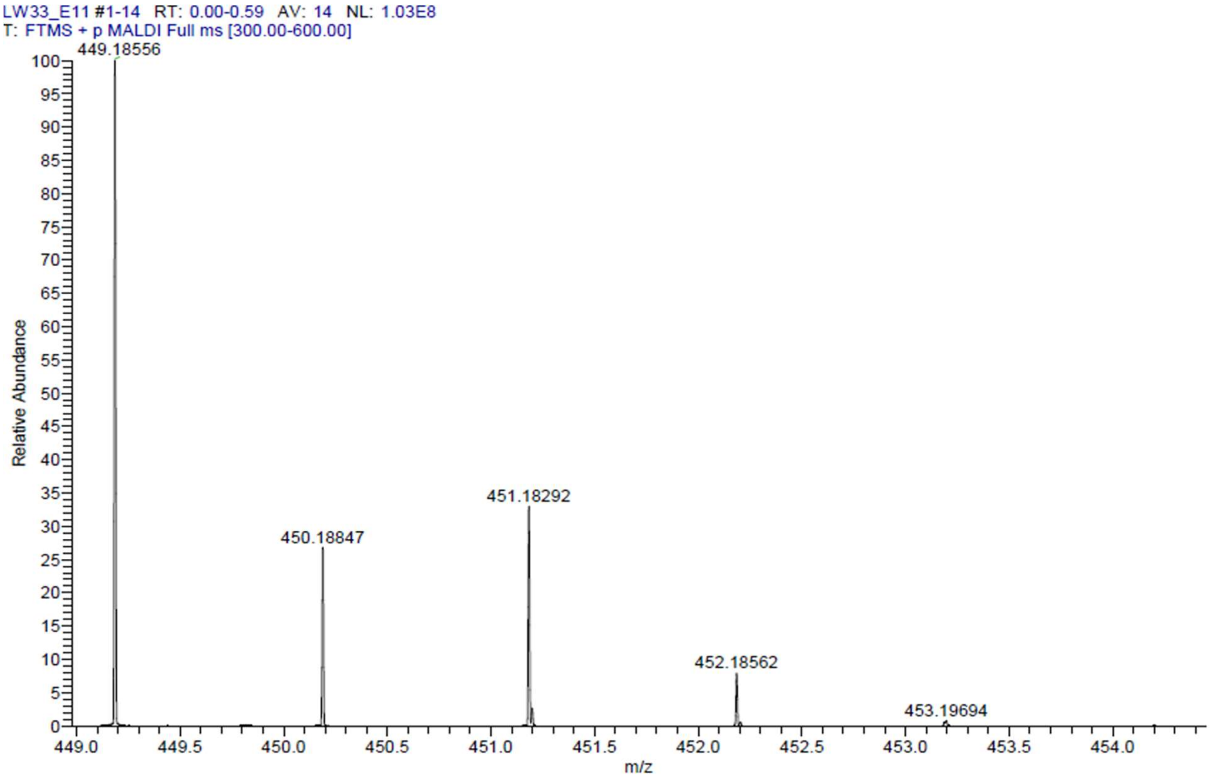

HPLC-chromatograms compound **20**:

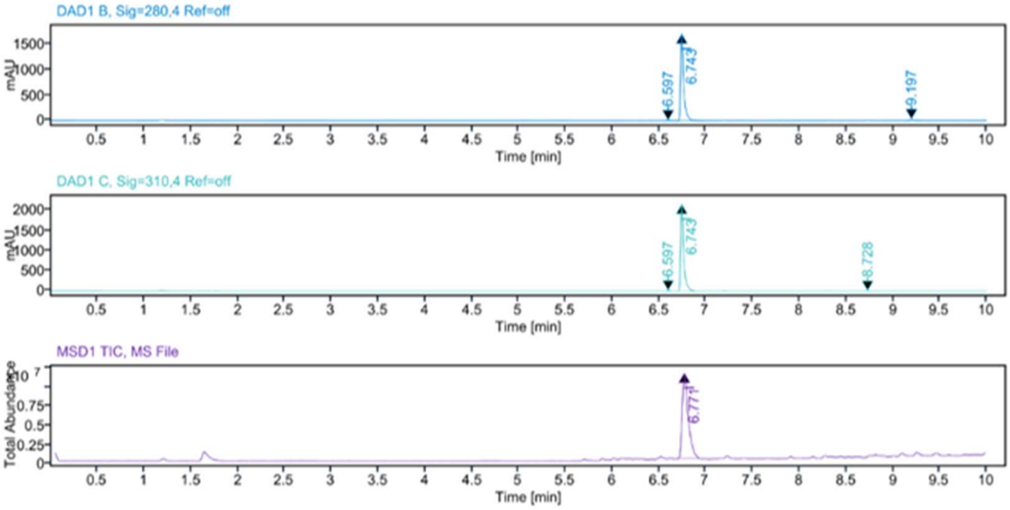


**Sample Purity**

Signal Description DAD1 B, Sig=280,4 Ref=off

| Sample Name | Name | RT | Width | Area | Area% | Height |
|---|---|---|---|---|---|---|
| LW33 | | 6.597 | 0.025 | 17.8330 | 0.45 | 9.7258 |
| LW33 | | 6.743 | 0.031 | 3865.4363 | 97.52 | 1704.5347 |
| LW33 | | 9.197 | 0.036 | 80.2835 | 2.03 | 30.7405 |

Max Area% 97.525

UV Signal Purity>95% Pass

Signal Description DAD1 C, Sig=310,4 Ref=off

| Sample Name | Name | RT | Width | Area | Area% | Height |
|---|---|---|---|---|---|---|
| LW33 | | 6.597 | 0.025 | 24.7654 | 0.50 | 13.4556 |
| LW33 | | 6.743 | 0.032 | 4909.3789 | 99.18 | 2164.3638 |
| LW33 | | 8.728 | 0.036 | 15.8413 | 0.32 | 5.3527 |

Max Area% 99.180

UV Signal Purity>95% Pass

**MSD Apex Spectra**

**RT:**

6.771

**Sample Name:**

LW33

**Signal Name:**

MSD1TIC

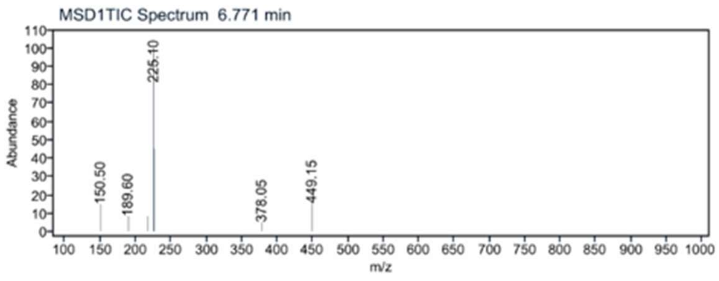

[1]H NMR compound **21**:

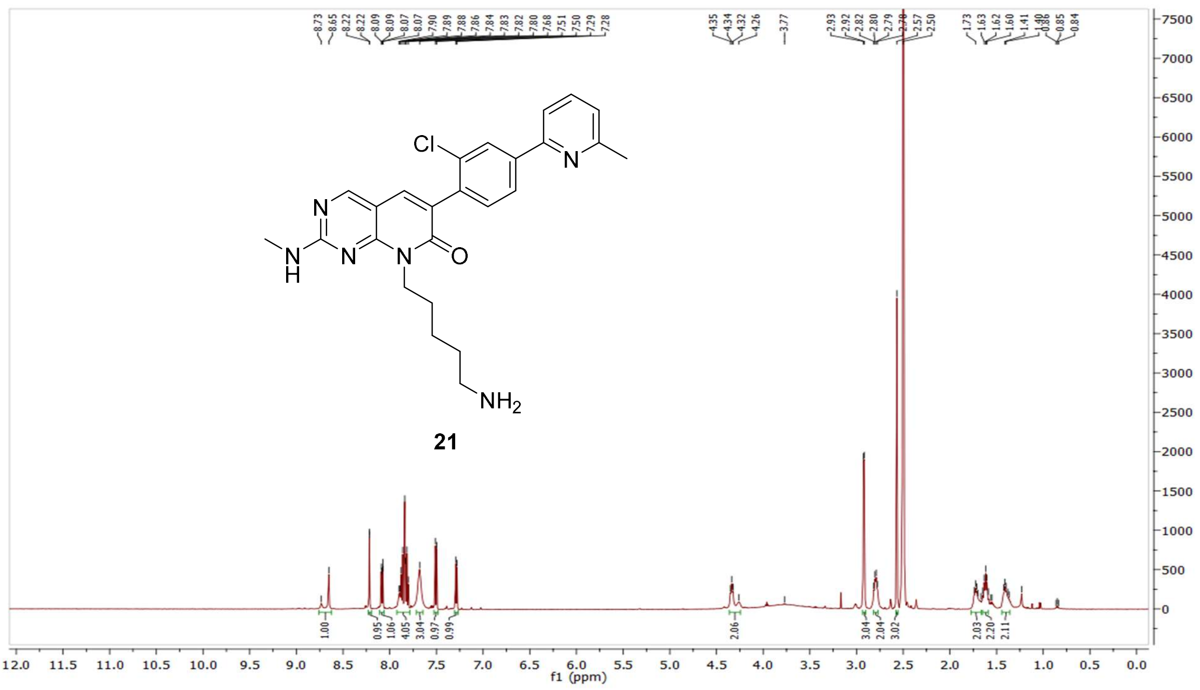

[13]C NMR compound **21:**

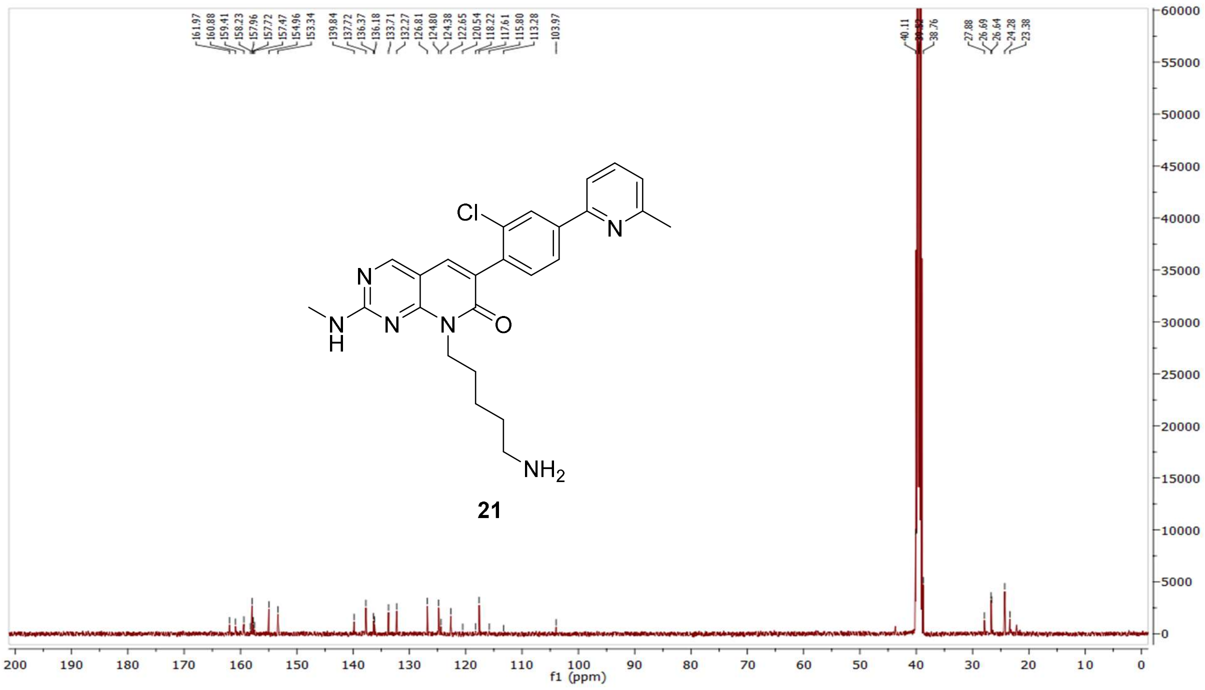

HRMS (FTMS +p MALDI) compound **21**:

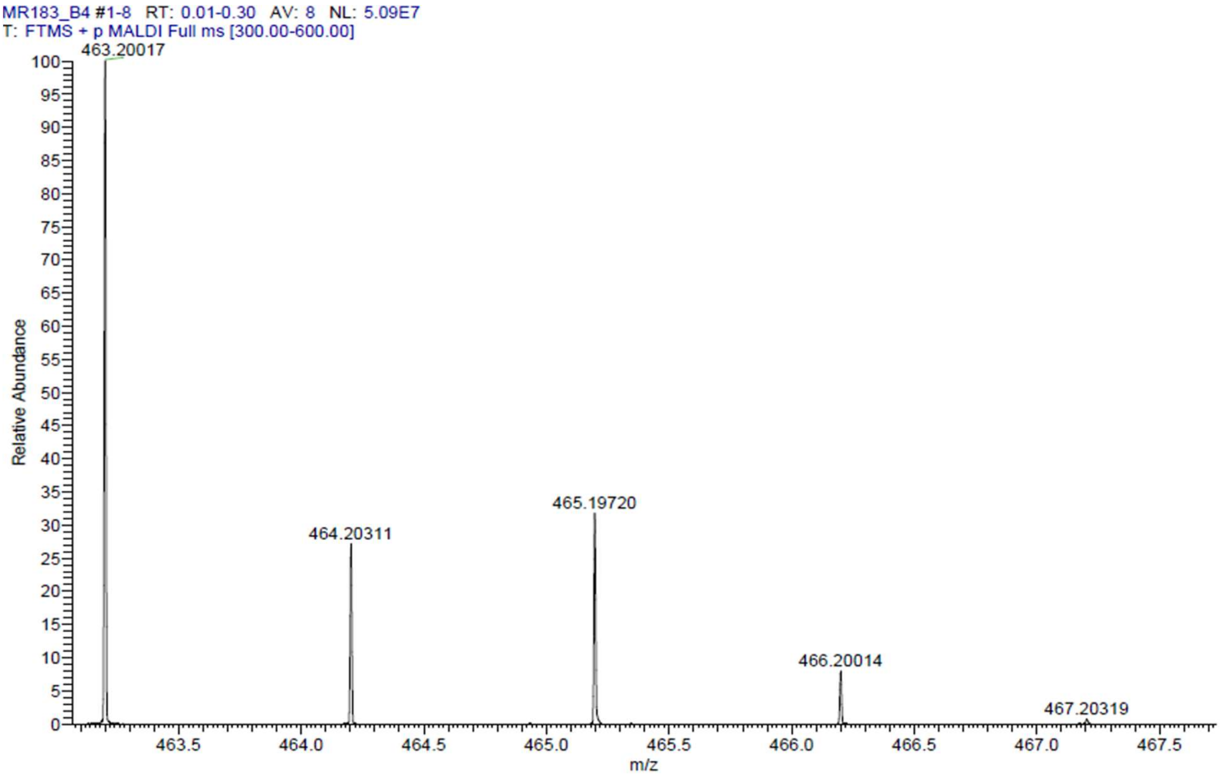

HPLC-chromatograms compound **21**:

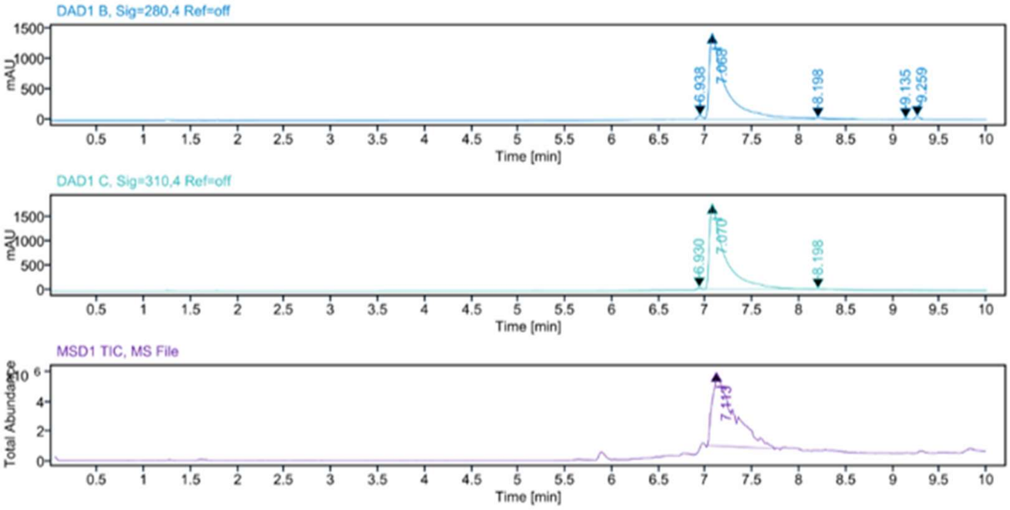


**Sample Purity**

Signal Description DAD1 B, Sig=280,4 Ref=off

| Sample Name | Name | RT | Width | Area | Area% | Height |
|---|---|---|---|---|---|---|
| MR183 Fr.10 | | 6.938 | 0.067 | 241.5397 | 1.34 | 60.6200 |
| MR183 Fr.10 | | 7.068 | 0.136 | 17131.51<br>95 | 95.22 | 1397.7190 |
| MR183 Fr.10 | | 8.198 | 0.189 | 355.4109 | 1.98 | 24.6875 |
| MR183 Fr.10 | | 9.135 | 0.061 | 72.8949 | 0.41 | 18.5080 |
| MR183 Fr.10 | | 9.259 | 0.059 | 189.7458 | 1.05 | 50.3698 |

Max Area% 95.222

UV Signal Purity>95% Pass

Signal Description DAD1 C, Sig=310,4 Ref=off

| Sample Name | Name | RT | Width | Area | Area% | Height |
|---|---|---|---|---|---|---|
| MR183 Fr.10 | | 6.930 | 0.058 | 130.5763 | 0.66 | 39.4717 |
| MR183 Fr.10 | | 7.070 | 0.133 | 19610.66<br>60 | 99.06 | 1731.5616 |
| MR183 Fr.10 | | 8.198 | 0.078 | 55.0989 | 0.28 | 8.7183 |

Max Area% 99.062

UV Signal Purity>95% Pass

**MSD Apex Spectra**

**RT:**

7.113

**Sample Name:**

MR183 Fr.10

**Signal Name:**

MSD1TIC

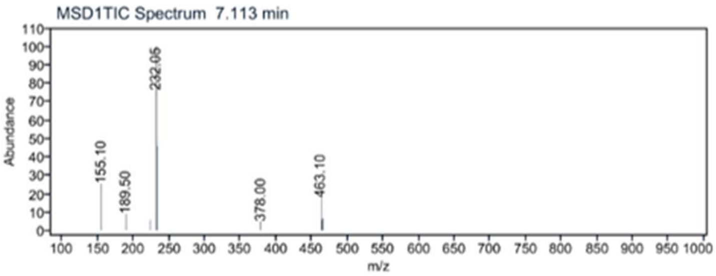


[1]H NMR compound **22**:

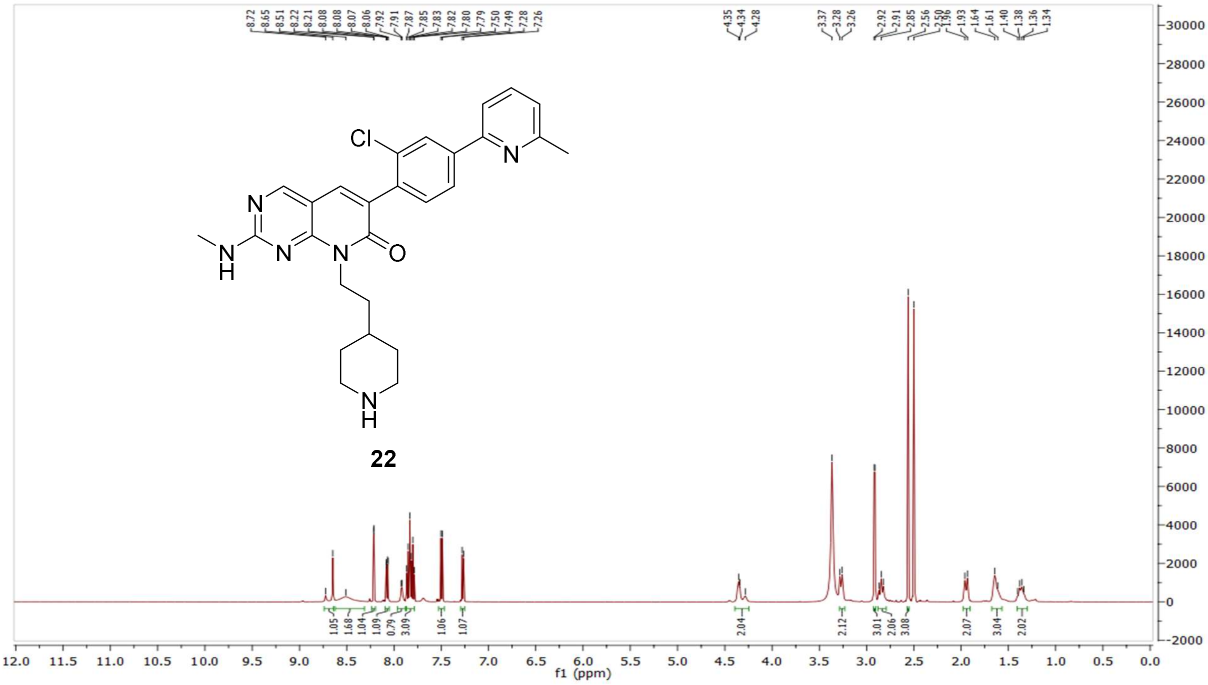

$^{13}$C NMR compound **22:**

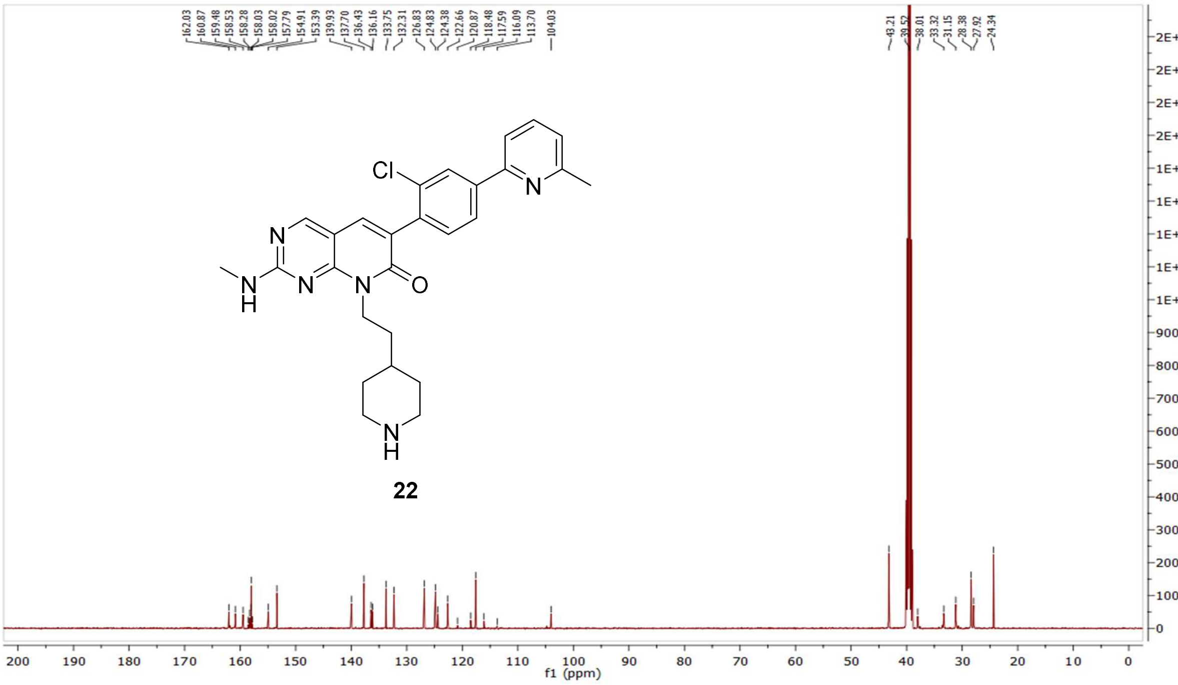

HRMS (FTMS +p MALDI) compound **22**:

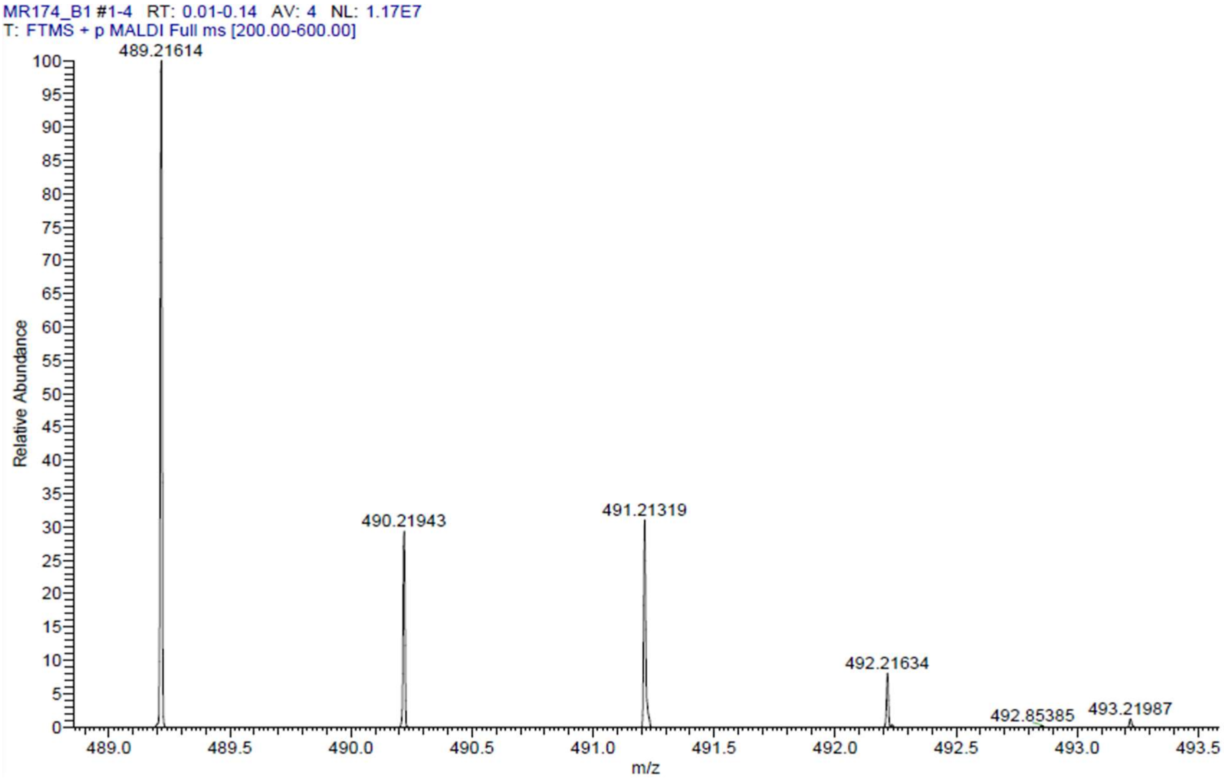

HPLC-chromatograms compound **22**:

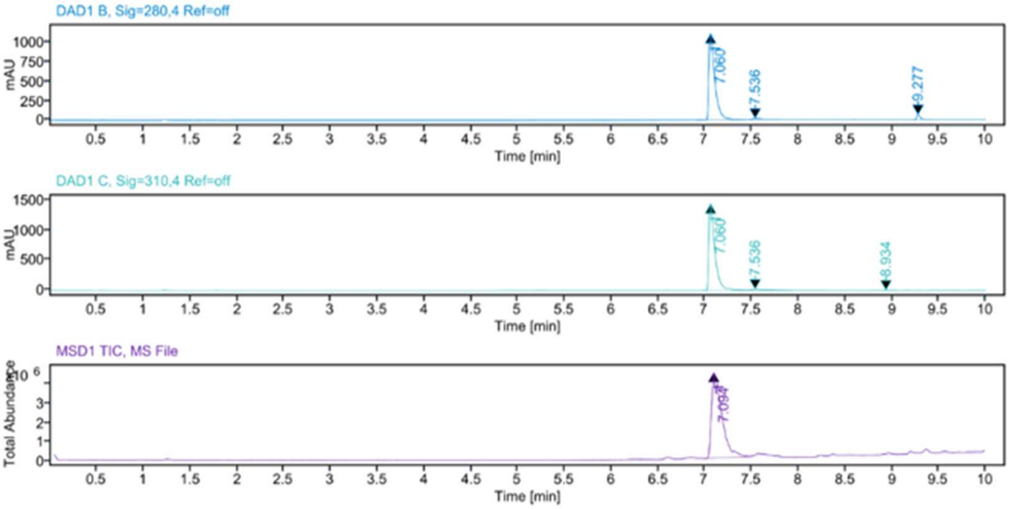


**Sample Purity**

| | |
|---|---|
| Signal Description | DAD1 B, Sig=280,4 Ref=off |

| Sample Name | Name | RT | Width | Area | Area% | Height |
|---|---|---|---|---|---|---|
| MR174 | | 7.060 | 0.063 | 4991.5537 | 95.17 | 1118.0280 |
| MR174 | | 7.536 | 0.123 | 90.6345 | 1.73 | 14.3690 |
| MR174 | | 9.277 | 0.042 | 162.9140 | 3.11 | 62.3949 |

| | |
|---|---|
| Max Area% | 95.166 |
| UV Signal Purity>95% | Pass |
| Signal Description | DAD1 C, Sig=310,4 Ref=off |

| Sample Name | Name | RT | Width | Area | Area% | Height |
|---|---|---|---|---|---|---|
| MR174 | | 7.060 | 0.063 | 6385.3379 | 96.96 | 1434.0374 |
| MR174 | | 7.536 | 0.167 | 193.4520 | 2.94 | 17.3180 |
| MR174 | | 8.934 | 0.047 | 6.8415 | 0.10 | 1.9418 |

| | |
|---|---|
| Max Area% | 96.959 |
| UV Signal Purity>95% | Pass |

**MSD Apex Spectra**

**RT:**

7.094

**Sample Name:**

MR174

**Signal Name:**

MSD1TIC

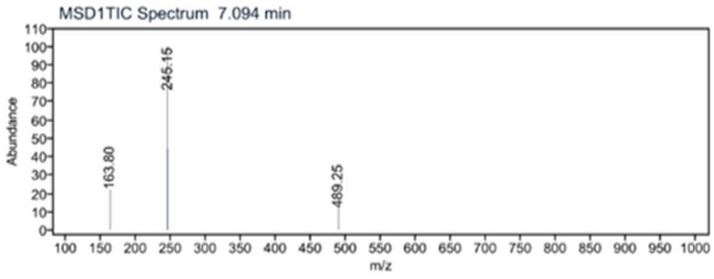

$^{1}$H NMR compound **23**:

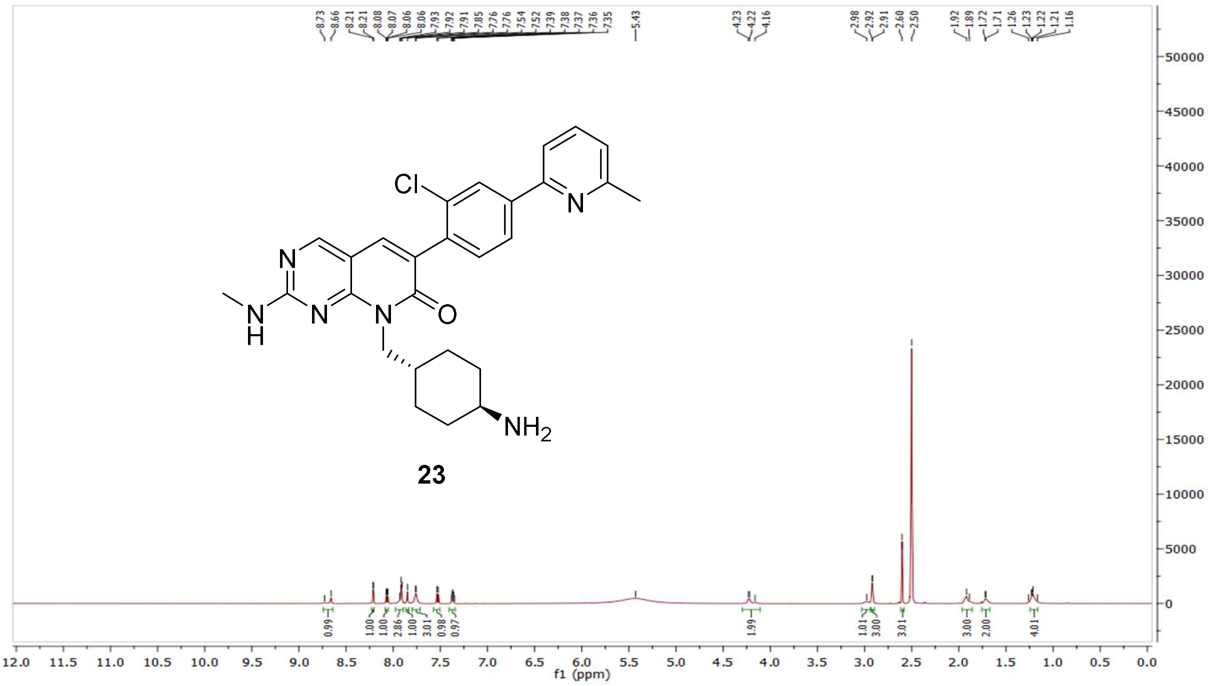

$^{13}$C NMR compound **23:**

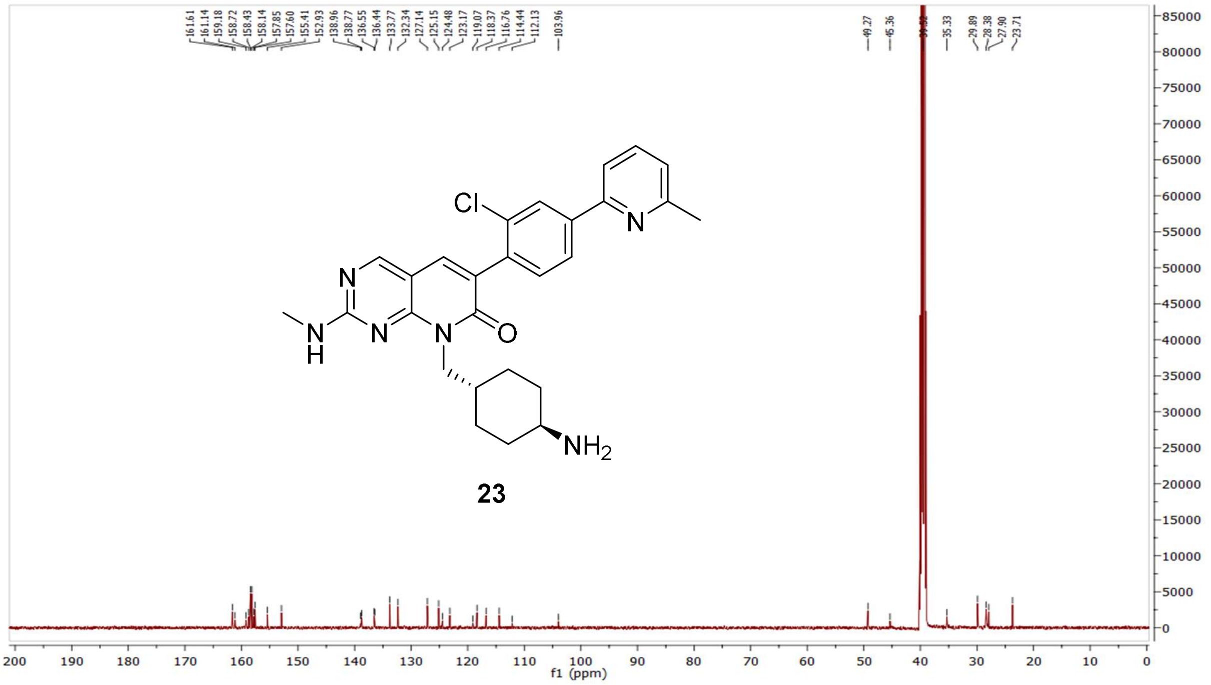

HRMS (FTMS +p MALDI) compound **23**:

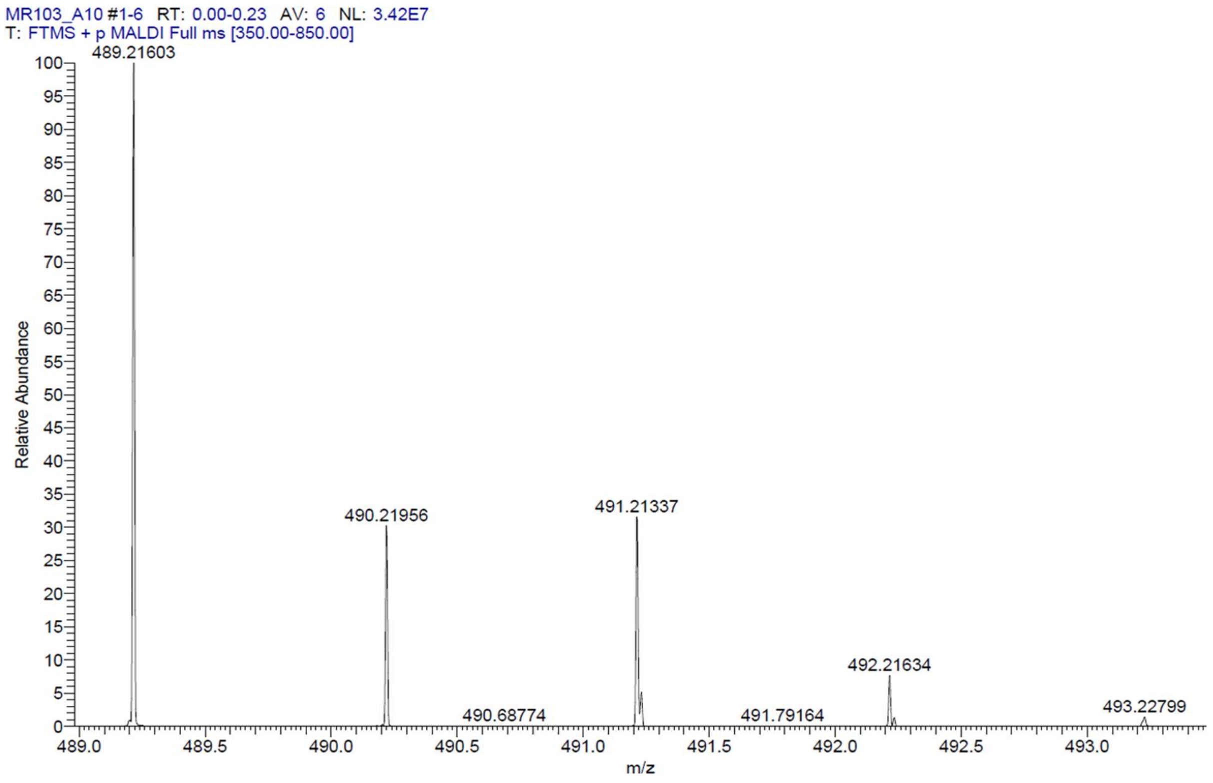

HPLC-chromatograms compound **23**:

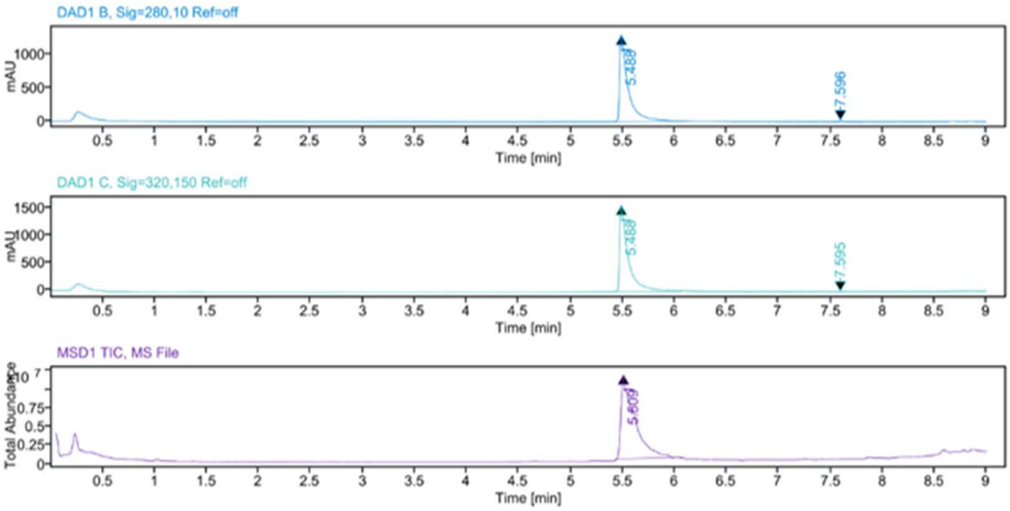


**Sample Purity**

Signal Description DAD1 B, Sig=280,10 Ref=off

| Sample Name | Name | RT | Width | Area | Area% | Height |
|---|---|---|---|---|---|---|
| MR103 Fr.1 | | 5.488 | 0.069 | 7566.7876 | 99.05 | 1312.4395 |
| MR103 Fr.1 | | 7.596 | 0.039 | 72.4722 | 0.95 | 24.2935 |

Max Area% 99.051

UV Signal Purity>95% Pass

Signal Description DAD1 C, Sig=320,150 Ref=off

| Sample Name | Name | RT | Width | Area | Area% | Height |
|---|---|---|---|---|---|---|
| MR103 Fr.1 | | 5.488 | 0.068 | 8980.7656 | 99.89 | 1573.0945 |
| MR103 Fr.1 | | 7.595 | 0.038 | 9.9523 | 0.11 | 4.2086 |

Max Area% 99.889

UV Signal Purity>95% Pass

**MSD Apex Spectra**

**RT:**

5.509

**Sample Name:**

MR103 Fr.1

**Signal Name:**

MSD1TIC

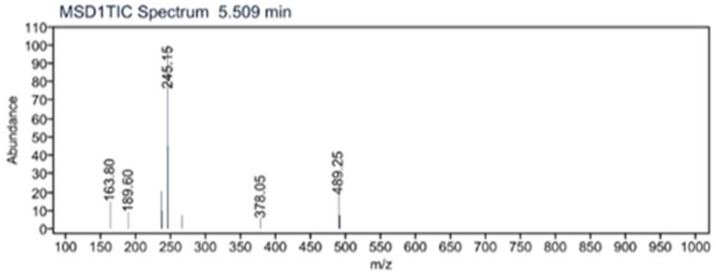

$^{1}$H NMR compound **24**:

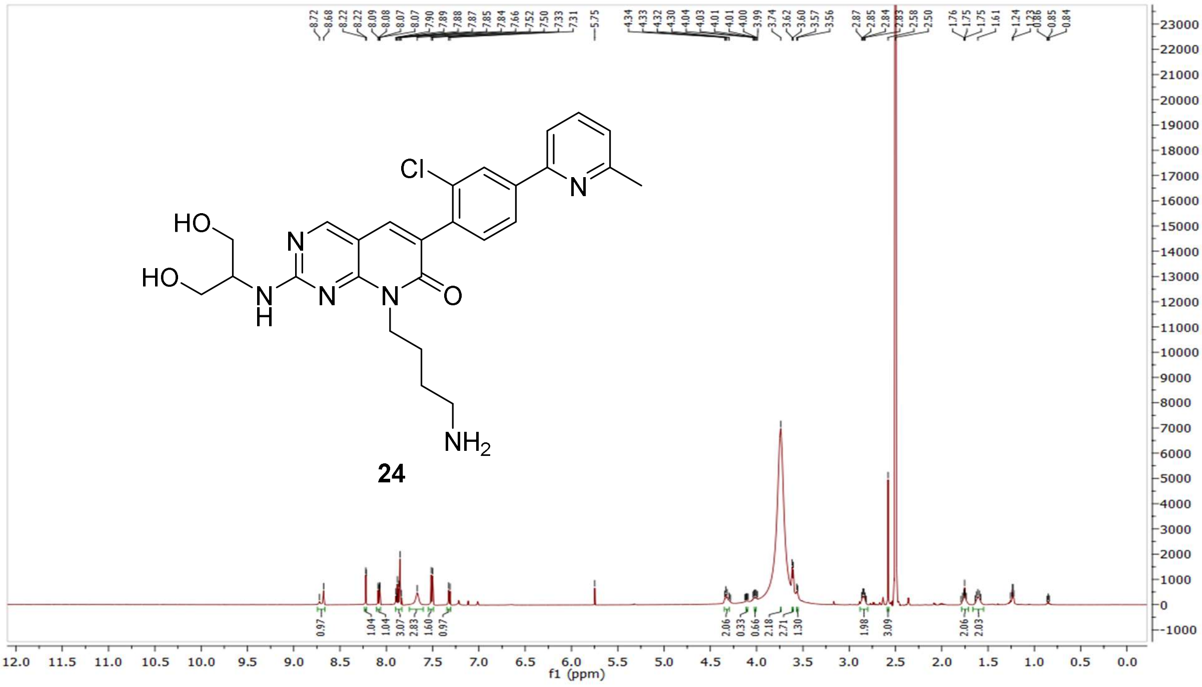


$^{13}$C NMR compound **24:**

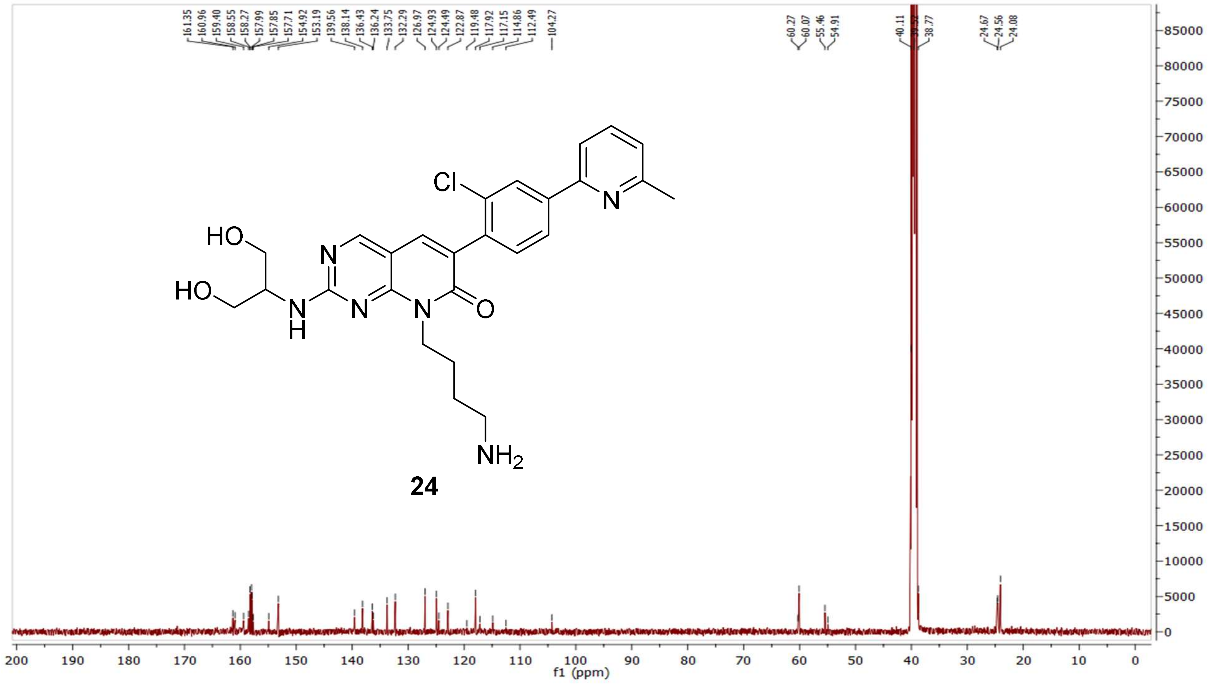

$^{1}H$-$^{1}H$ COSY NMR compound **24:**

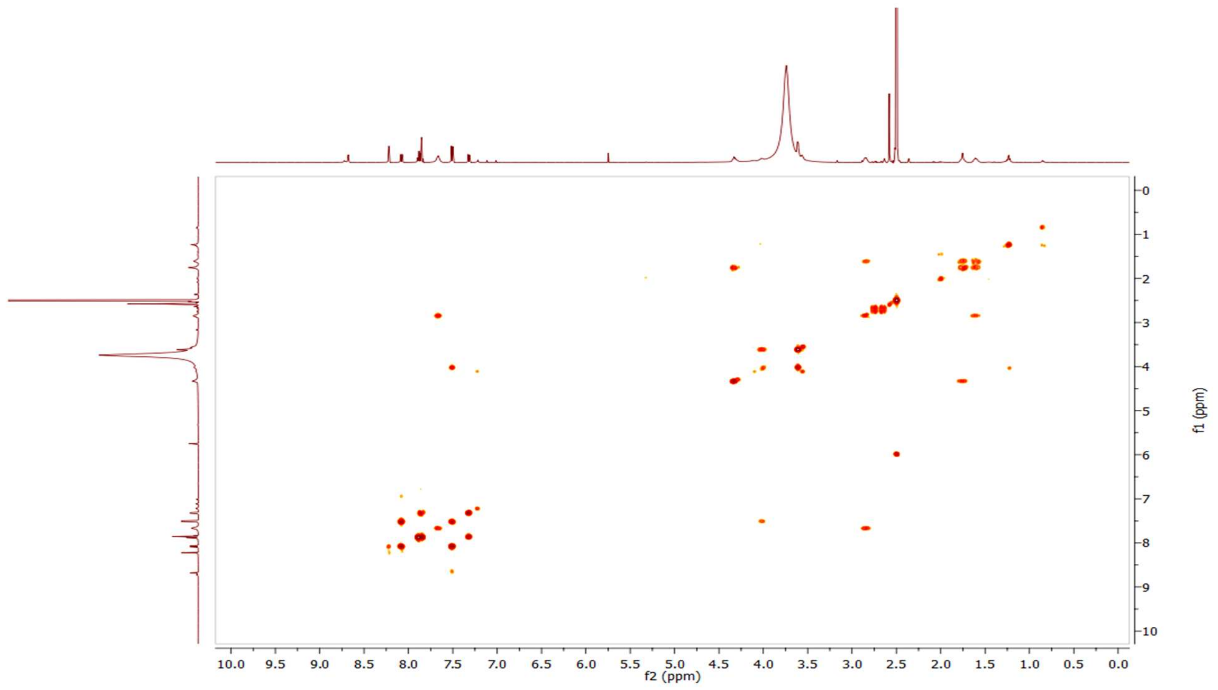


$^{1}H$-$^{13}C$ HSQC NMR compound **24:**

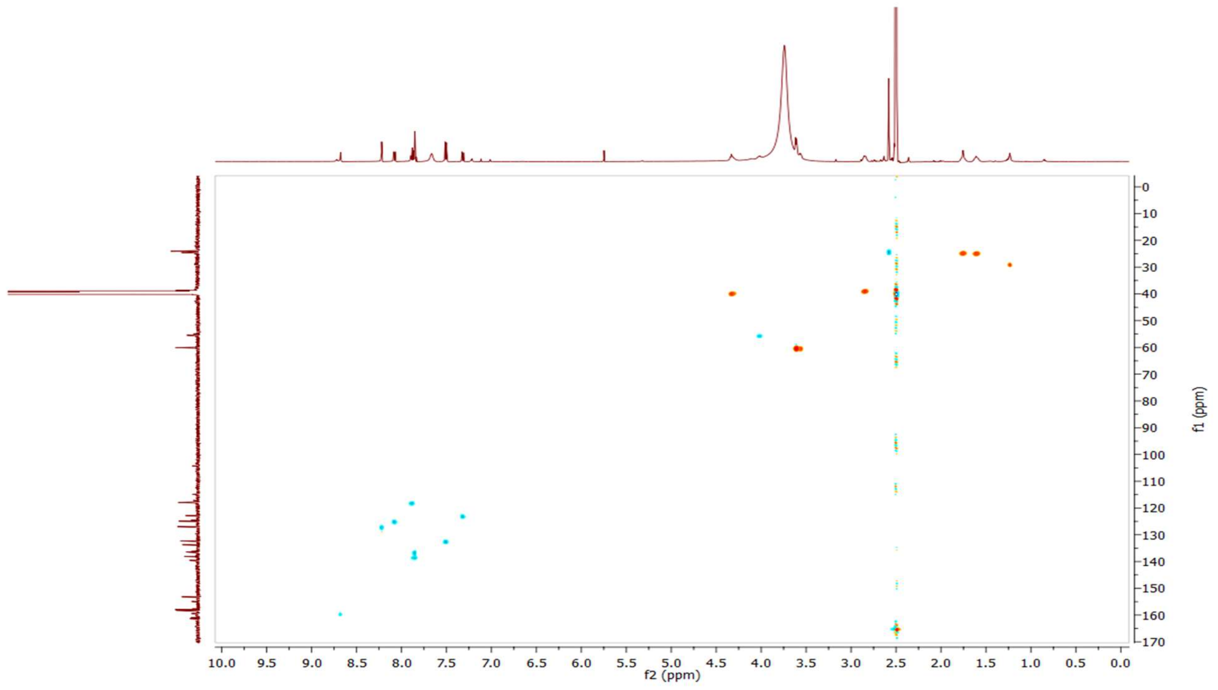

HRMS (FTMS +p MALDI) compound **24**:

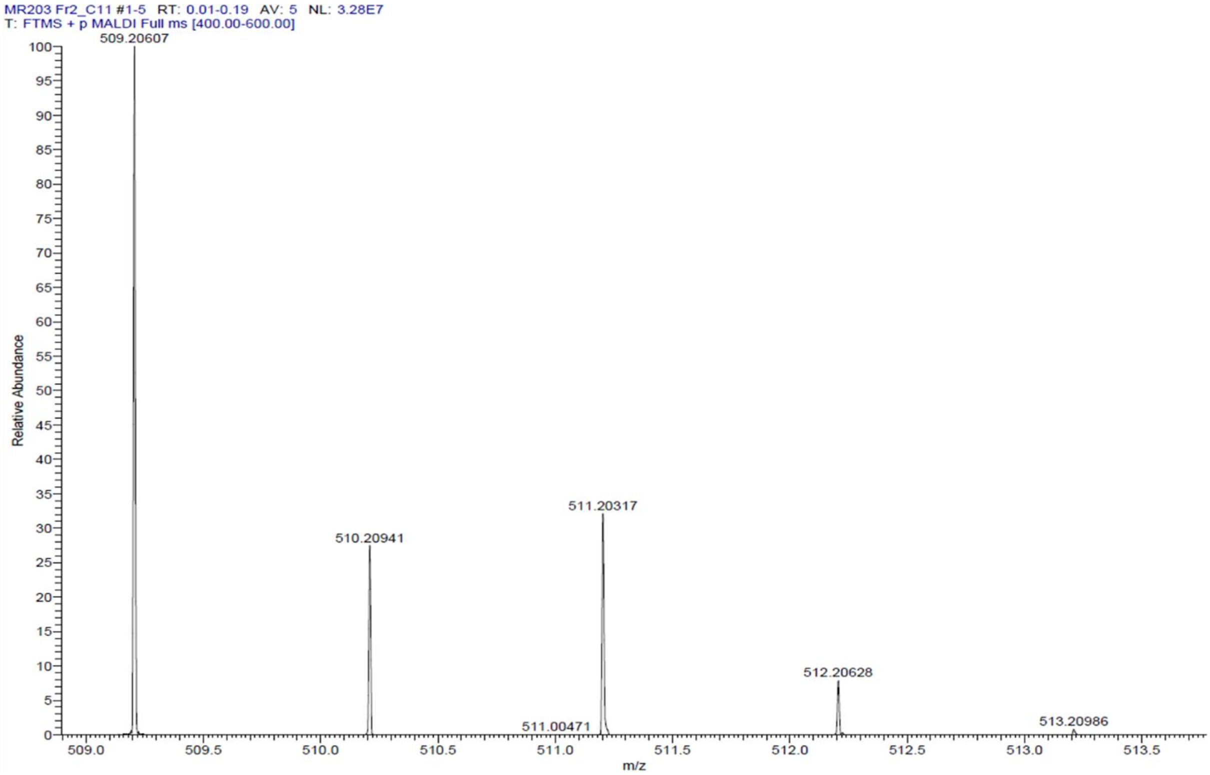

HPLC-chromatograms compound **24**:

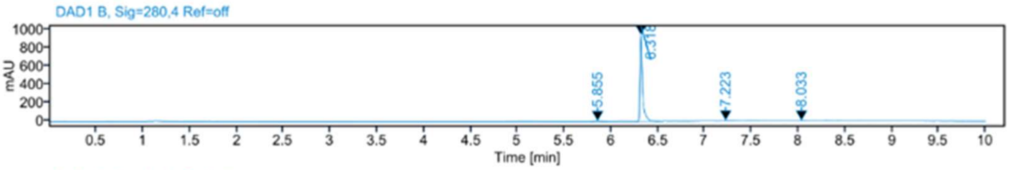


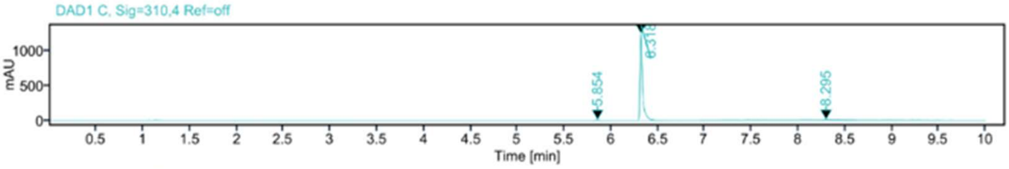


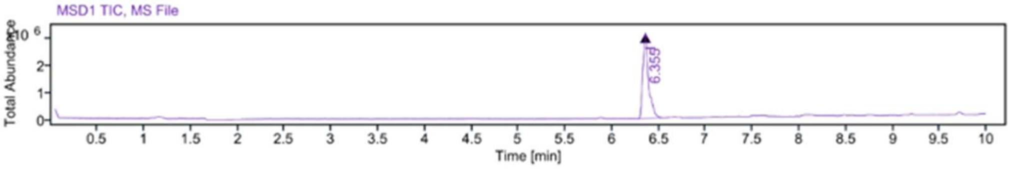


**Sample Purity**

Signal Description DAD1 B, Sig=280,4 Ref=off

| Sample Name | Name | RT | Width | Area | Area% | Height |
|---|---|---|---|---|---|---|
| MR203 Fr.2 | | 5.855 | 0.038 | 13.6211 | 0.74 | 4.6636 |
| MR203 Fr.2 | | 6.318 | 0.025 | 1774.8159 | 96.89 | 942.7355 |
| MR203 Fr.2 | | 7.223 | 0.121 | 29.9518 | 1.64 | 5.6283 |
| MR203 Fr.2 | | 8.033 | 0.043 | 13.4039 | 0.73 | 3.2504 |

Max Area% 96.890

UV Signal Purity>95% Pass

Signal Description DAD1 C, Sig=310,4 Ref=off

| Sample Name | Name | RT | Width | Area | Area% | Height |
|---|---|---|---|---|---|---|
| MR203 Fr.2 | | 5.854 | 0.039 | 16.5599 | 0.69 | 5.5920 |
| MR203 Fr.2 | | 6.318 | 0.025 | 2357.4634 | 97.63 | 1247.7914 |
| MR203 Fr.2 | | 8.295 | 0.079 | 40.6620 | 1.68 | 5.7415 |

Max Area% 97.630

UV Signal Purity>95% Pass

**MSD Apex Spectra**

**RT:**

6.355

**Sample Name:**

MR203 Fr.2

**Signal Name:**

MSD1TIC

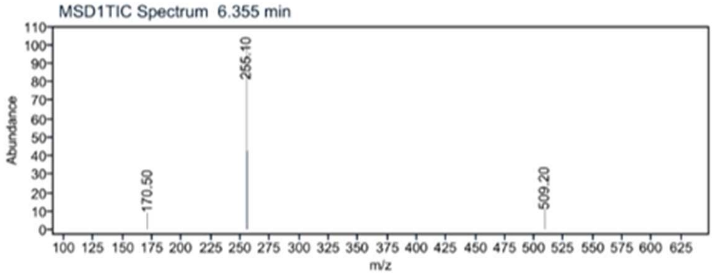

$^{1}$H NMR compound **25**:

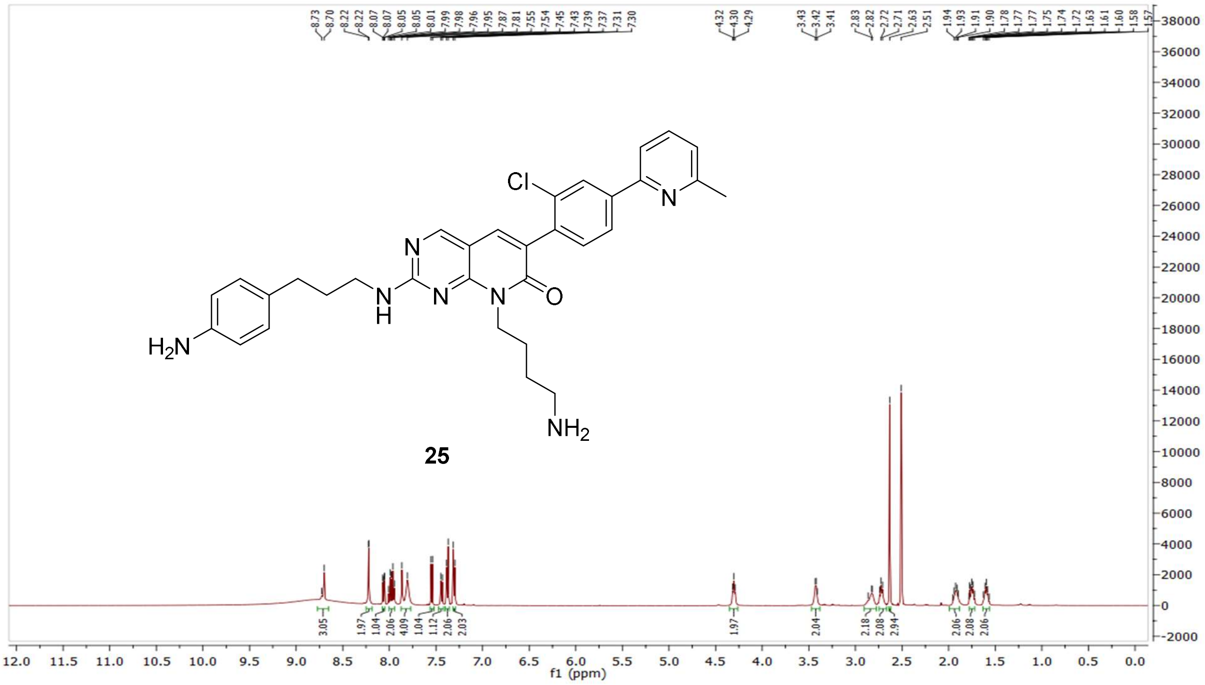

$^{13}$C NMR compound **25:**

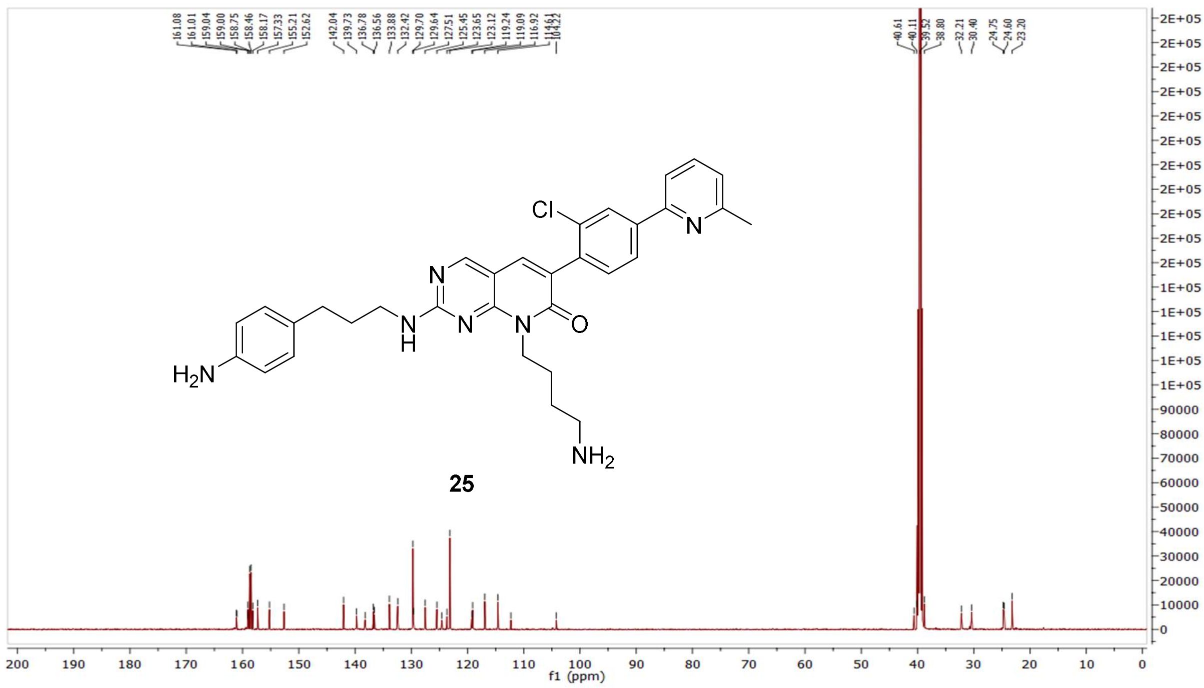

HRMS (FTMS +p MALDI) compound **25**:

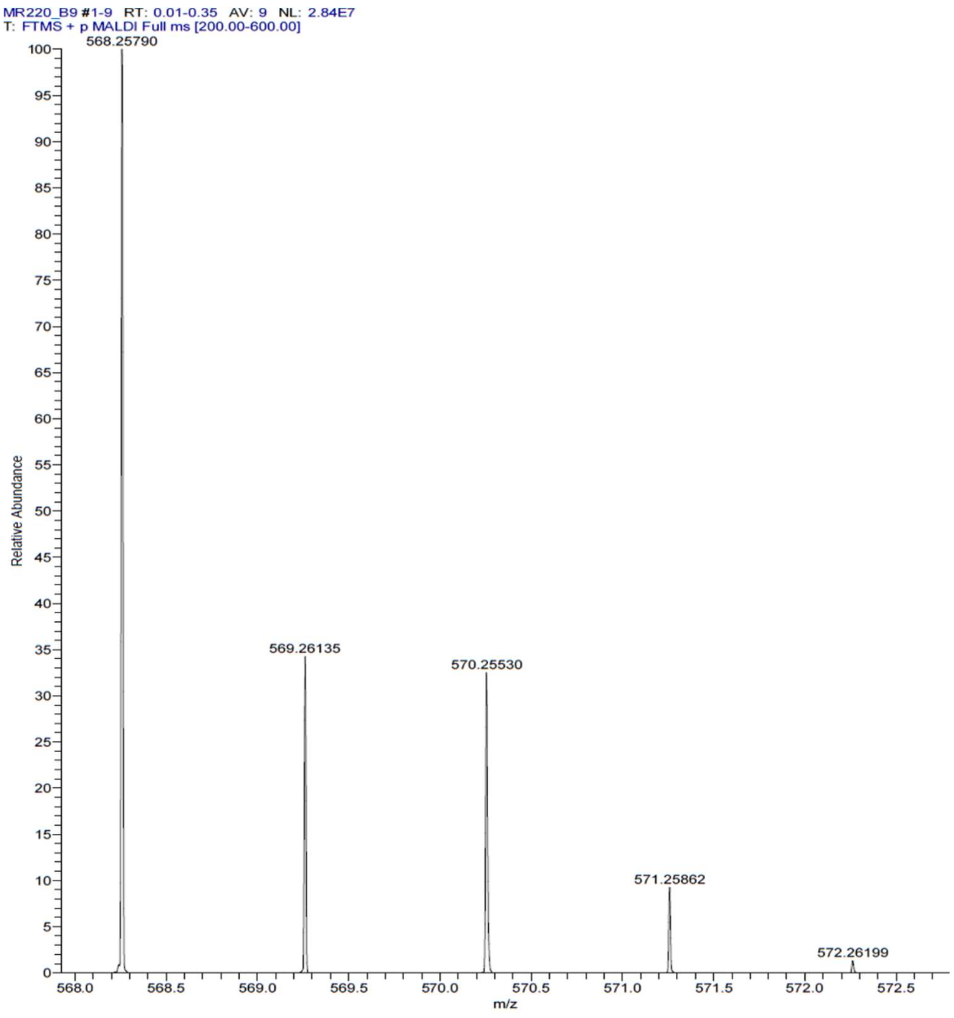

HPLC-chromatograms compound **25**:

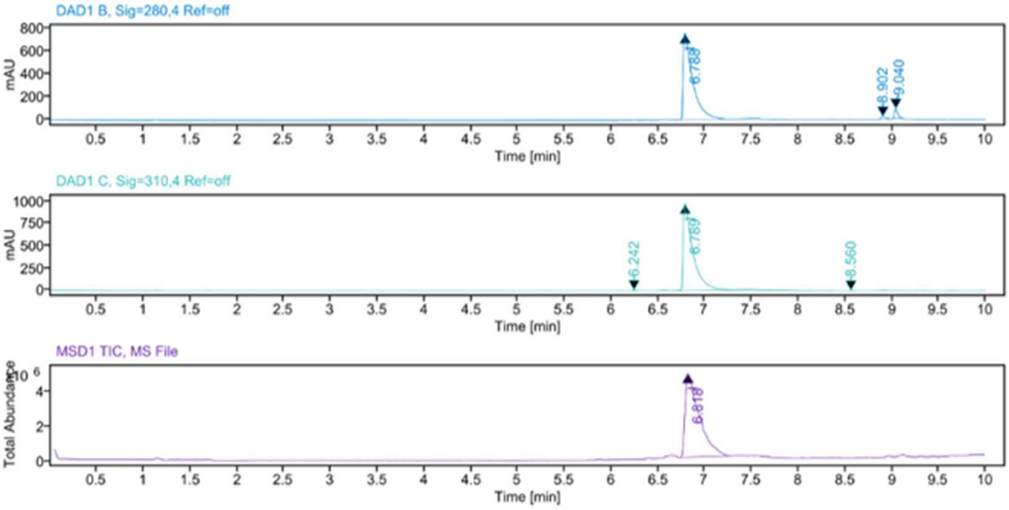


**Sample Purity**

Signal Description DAD1 B, Sig=280,4 Ref=off

| Sample Name | Name | RT | Width | Area | Area% | Height |
|---|---|---|---|---|---|---|
| MR220 | | 6.788 | 0.096 | 5365.5811 | 95.10 | 753.4427 |
| MR220 | | 8.902 | 0.050 | 65.7754 | 1.17 | 21.1441 |
| MR220 | | 9.040 | 0.037 | 210.6429 | 3.73 | 86.1849 |

Max Area% 95.101

UV Signal Purity>95% Pass

Signal Description DAD1 C, Sig=310,4 Ref=off

| Sample Name | Name | RT | Width | Area | Area% | Height |
|---|---|---|---|---|---|---|
| MR220 | | 6.242 | 0.034 | 13.6362 | 0.20 | 5.7600 |
| MR220 | | 6.789 | 0.095 | 6926.7007 | 99.71 | 973.9311 |
| MR220 | | 8.560 | 0.038 | 6.3518 | 0.09 | 2.3552 |

Max Area% 99.712

UV Signal Purity>95% Pass

**MSD Apex Spectra**

**RT:**

6.818

**Sample Name:**

MR220

**Signal Name:**

MSD1TIC

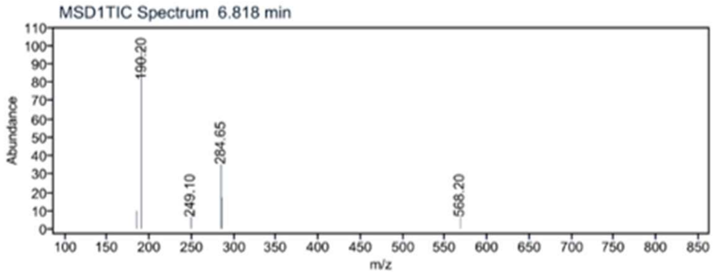

[1]H NMR compound **26**:

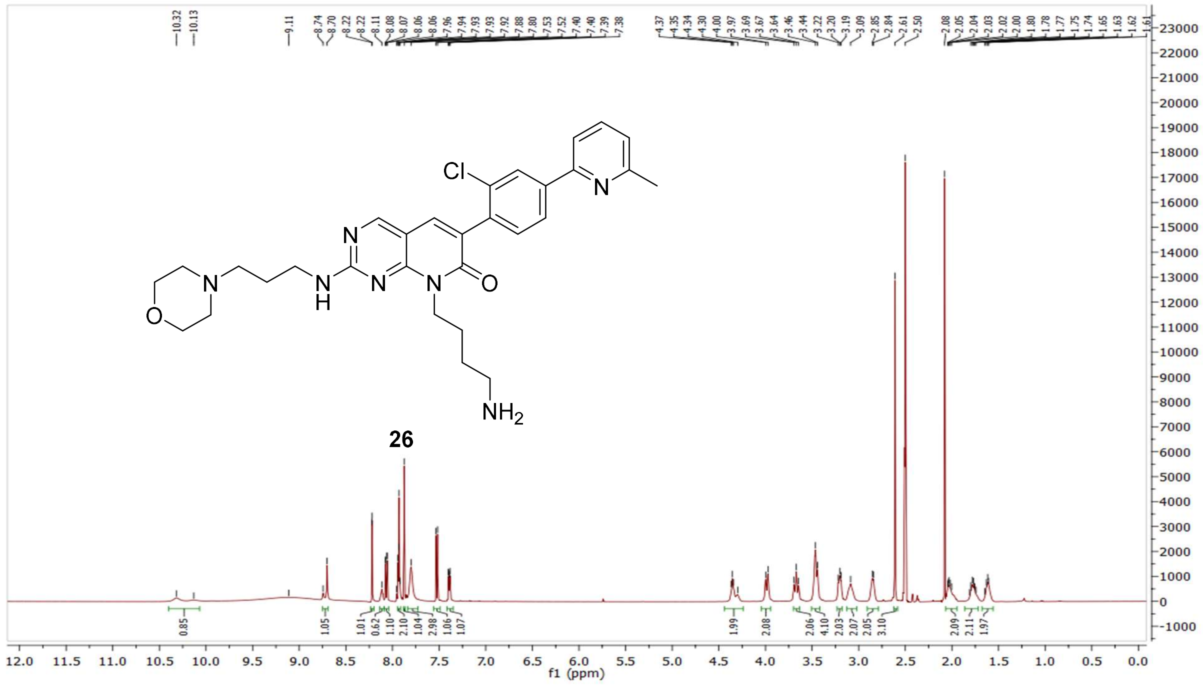


[13]C NMR compound **26:**

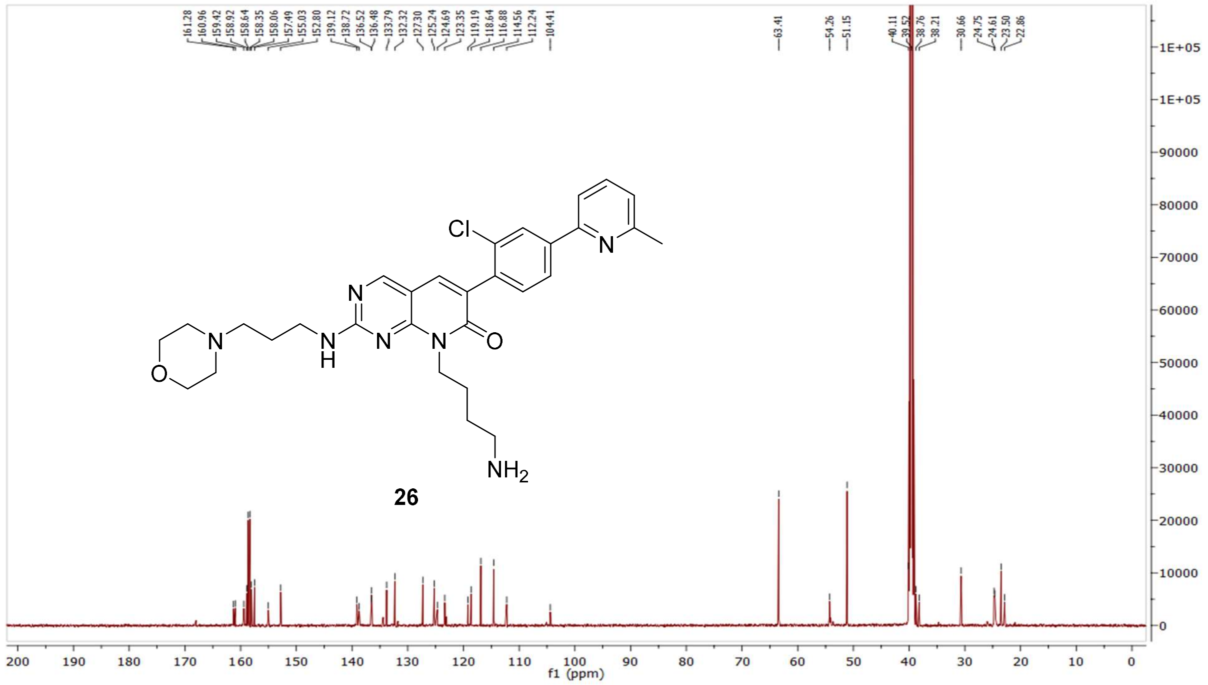

HRMS (FTMS +p MALDI) compound **26**:

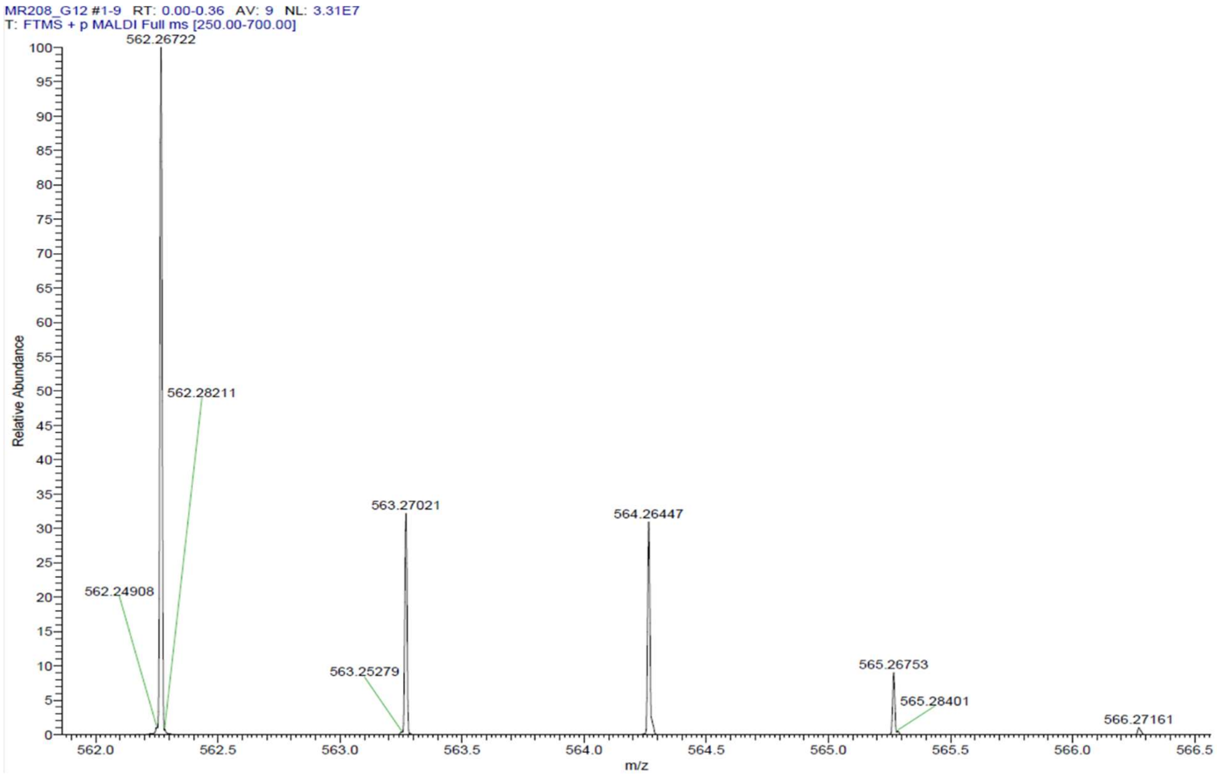

HPLC-chromatograms compound **26**:

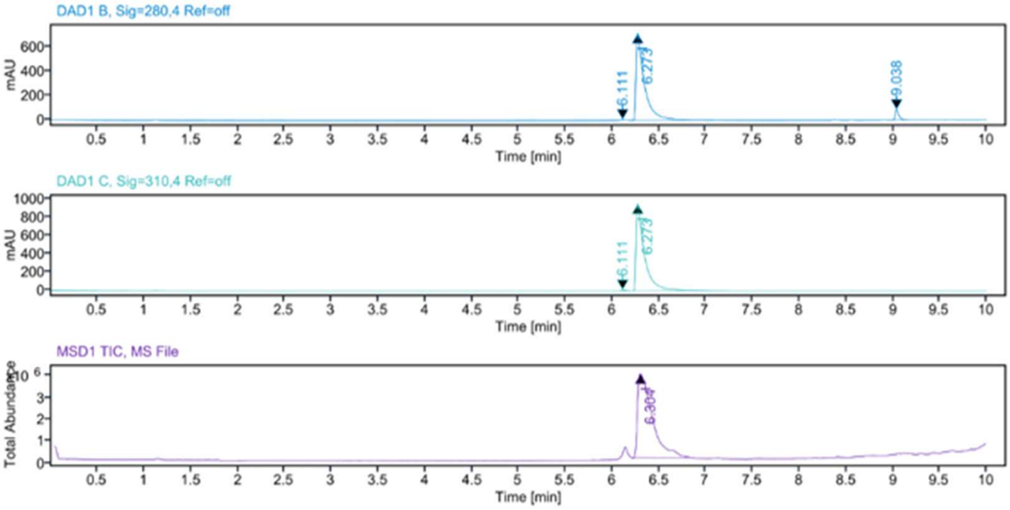


**Sample Purity**

Signal Description DAD1 B, Sig=280,4 Ref=off

| Sample Name | Name | RT | Width | Area | Area% | Height |
|---|---|---|---|---|---|---|
| MR208 | | 6.111 | 0.024 | 9.9063 | 0.21 | 6.6923 |
| MR208 | | 6.273 | 0.080 | 4480.9336 | 95.03 | 714.2936 |
| MR208 | | 9.038 | 0.037 | 224.4535 | 4.76 | 86.3237 |

Max Area% 95.030

UV Signal Purity>95% Pass

Signal Description DAD1 C, Sig=310,4 Ref=off

| Sample Name | Name | RT | Width | Area | Area% | Height |
|---|---|---|---|---|---|---|
| MR208 | | 6.111 | 0.030 | 28.2554 | 0.48 | 12.2316 |
| MR208 | | 6.273 | 0.080 | 5857.3203 | 99.52 | 956.0861 |

Max Area% 99.520

UV Signal Purity>95% Pass

**MSD Apex Spectra**

**RT:**

6.304

**Sample Name:**

MR208

**Signal Name:**

MSD1TIC

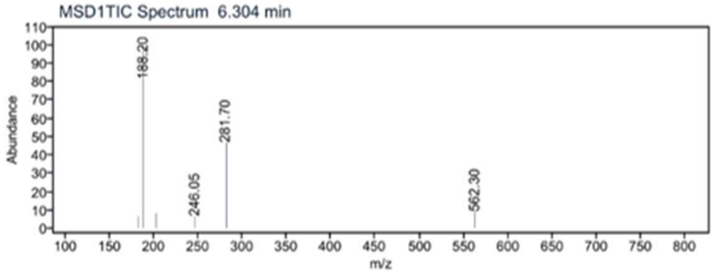

[1]H NMR compound **27**:

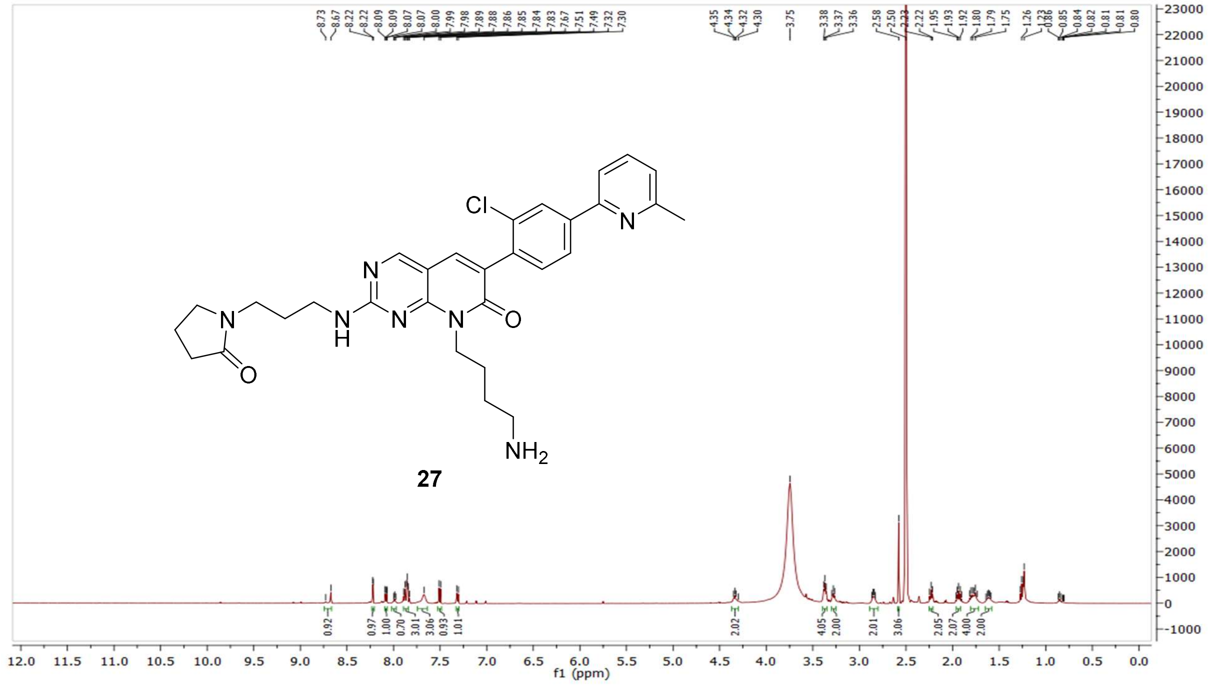

[13]C NMR compound **27:**

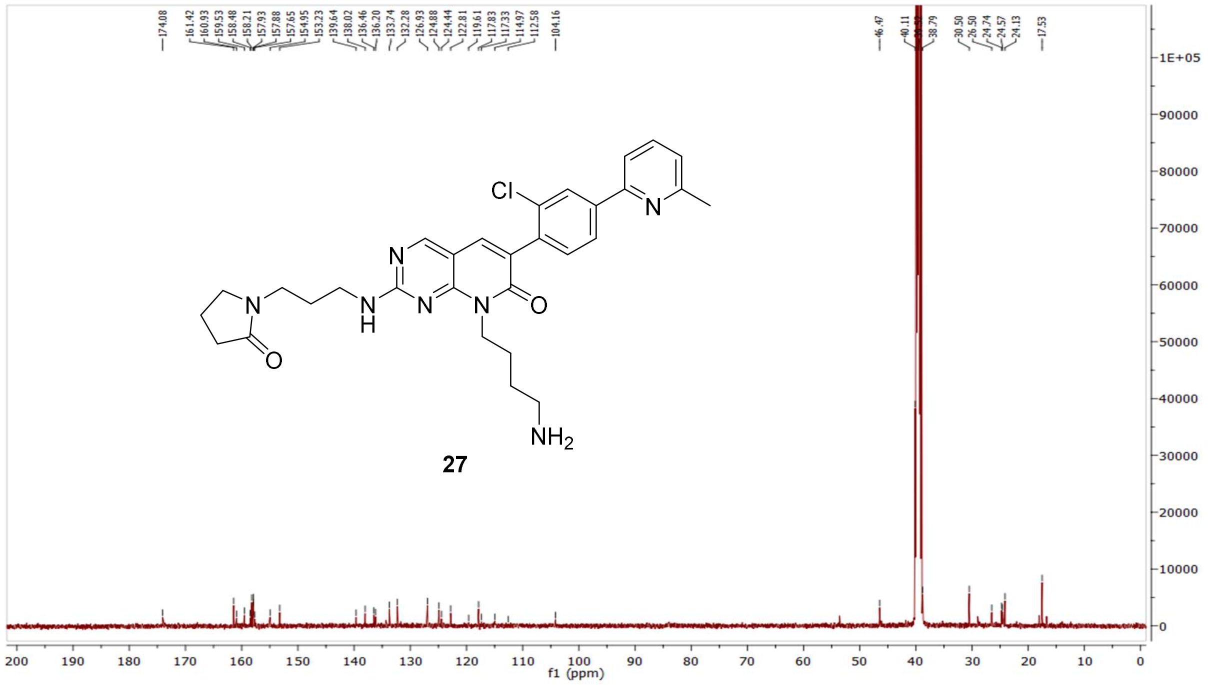

HRMS (FTMS +p MALDI) compound **27**:

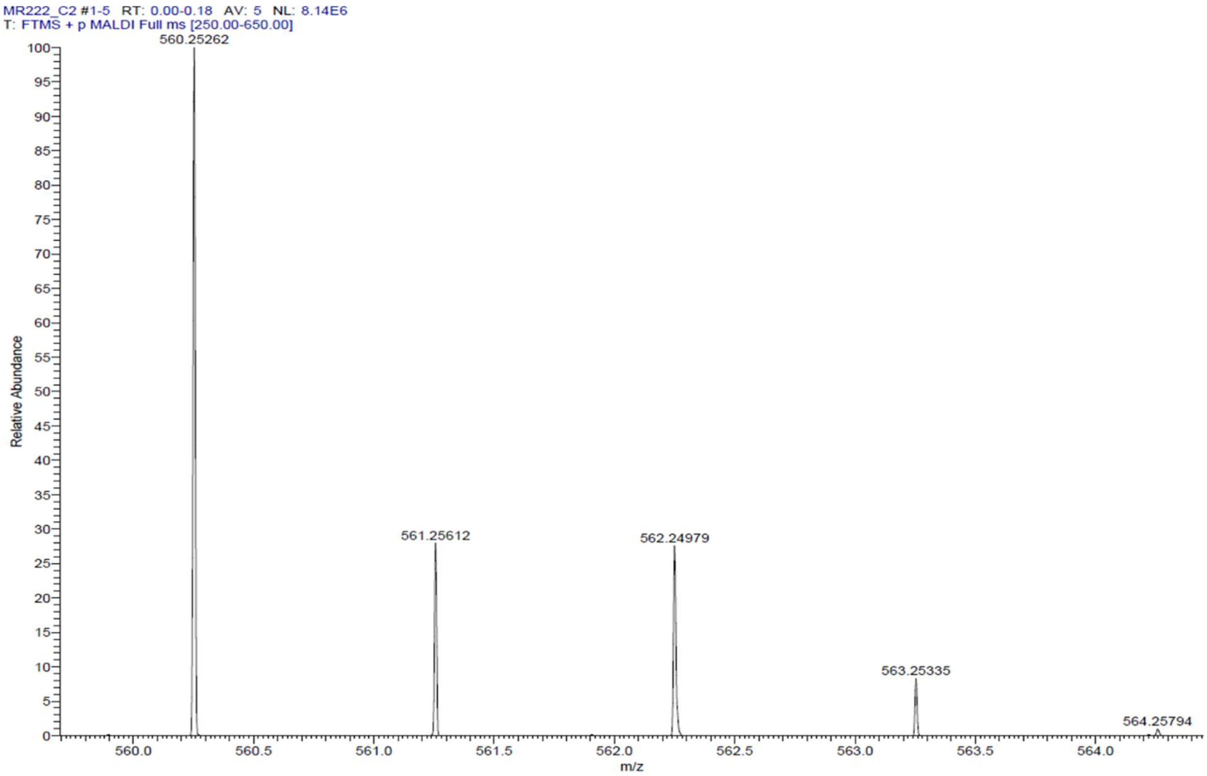

HPLC-chromatograms compound **27**:

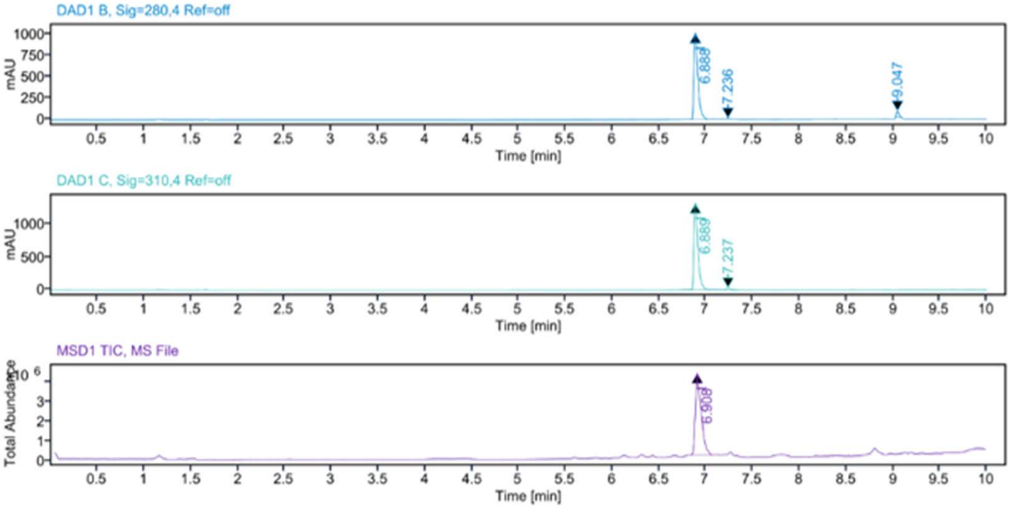


**Sample Purity**

Signal Description DAD1 B, Sig=280,4 Ref=off

| Sample Name | Name | RT | Width | Area | Area% | Height |
|---|---|---|---|---|---|---|
| MR222 F42 | | 6.888 | 0.046 | 3052.8311 | 95.16 | 1011.4472 |
| MR222 F42 | | 7.236 | 0.020 | 11.0566 | 0.34 | 9.3961 |
| MR222 F42 | | 9.047 | 0.031 | 144.1184 | 4.49 | 76.4450 |

Max Area% 95.163

UV Signal Purity>95% Pass

Signal Description DAD1 C, Sig=310,4 Ref=off

| Sample Name | Name | RT | Width | Area | Area% | Height |
|---|---|---|---|---|---|---|
| MR222 F42 | | 6.889 | 0.045 | 3980.4082 | 98.20 | 1316.7415 |
| MR222 F42 | | 7.237 | 0.032 | 72.8512 | 1.80 | 33.0787 |

Max Area% 98.203

UV Signal Purity>95% Pass

**MSD Apex Spectra**

**RT:**

6.908

**Sample Name:**

MR222 F42

**Signal Name:**

MSD1TIC

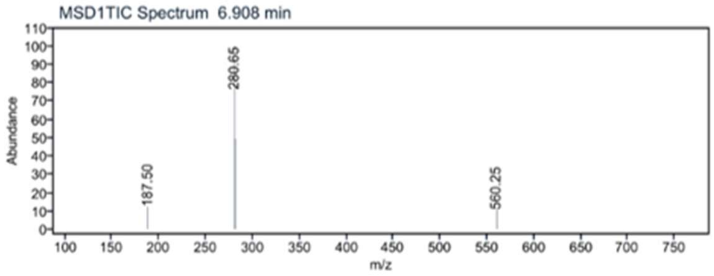

$^{1}H$ NMR compound **28**:

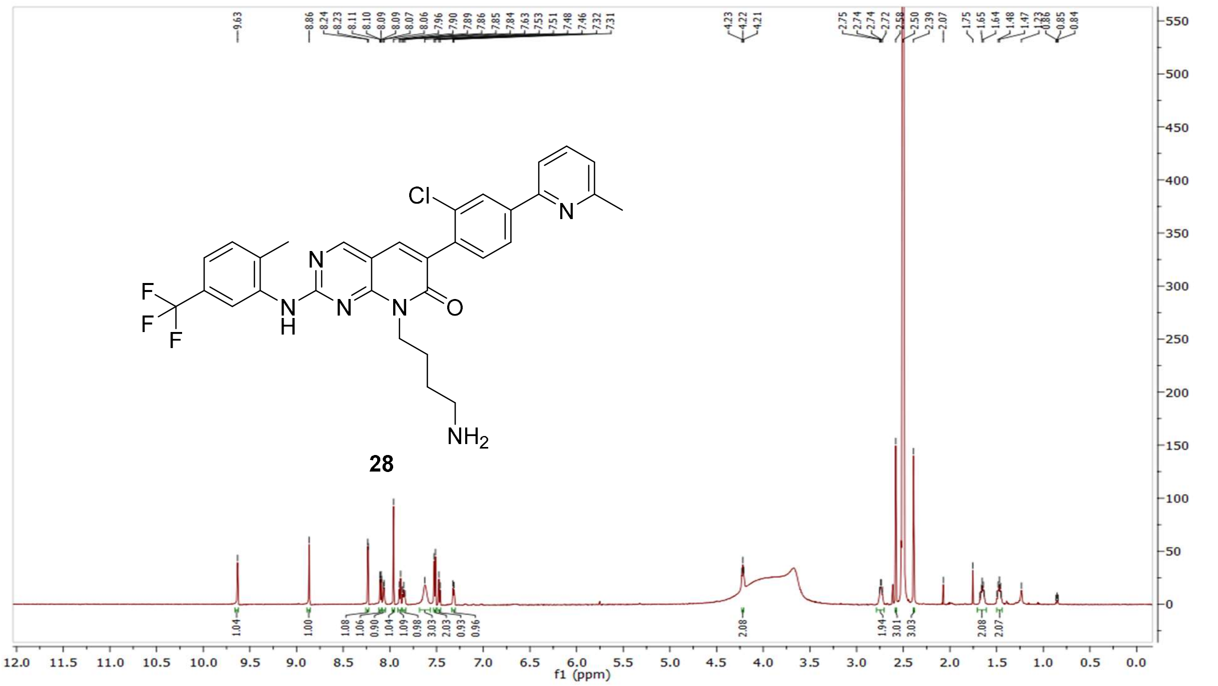


$^{13}C$ NMR compound **28:**

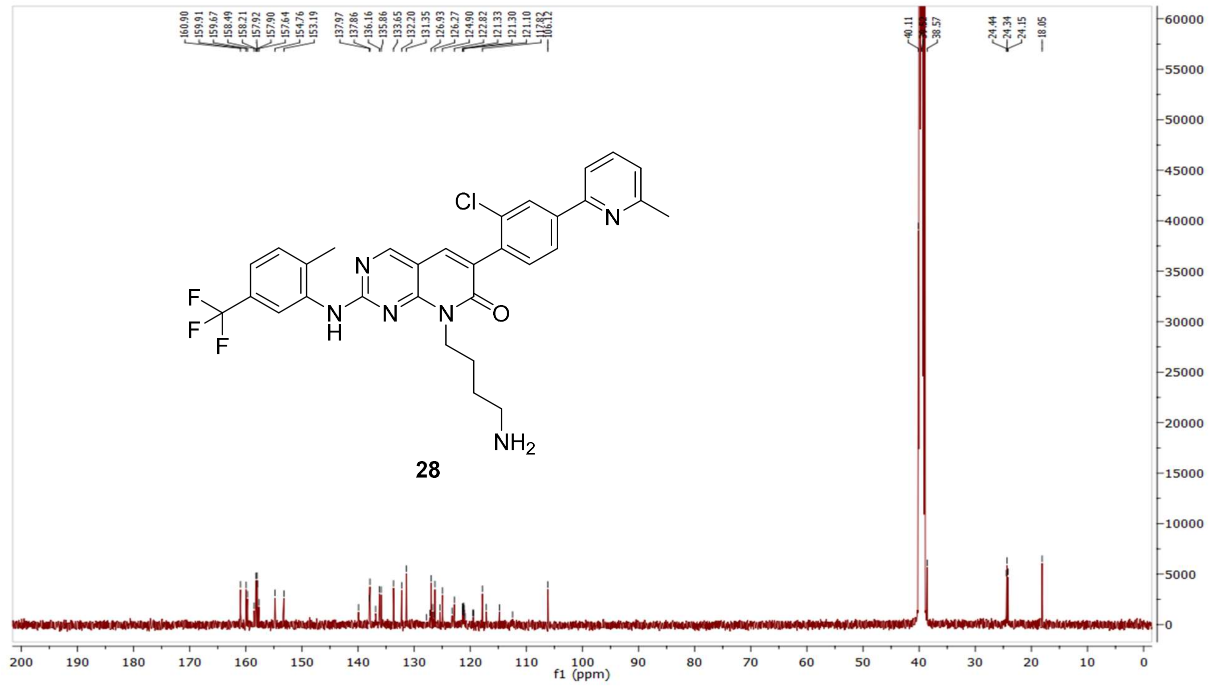

HRMS (FTMS +p MALDI) compound **28**:

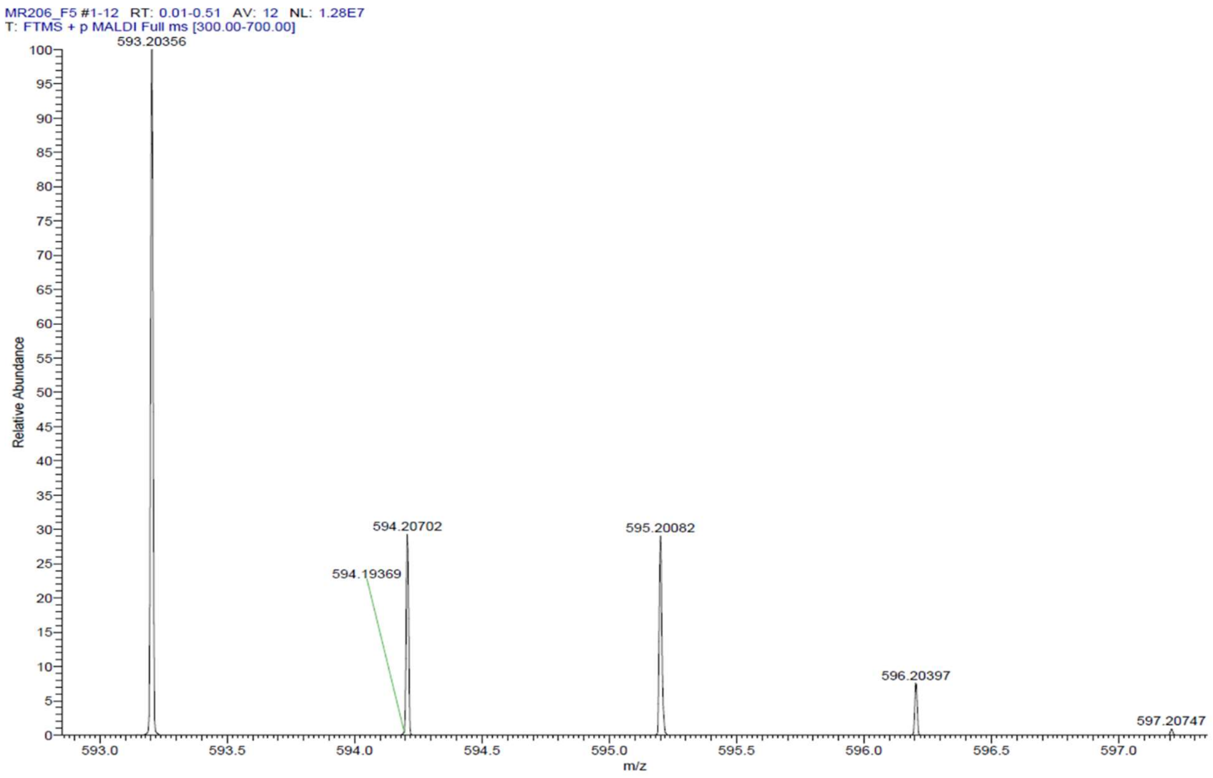

HPLC-chromatograms compound **28**:

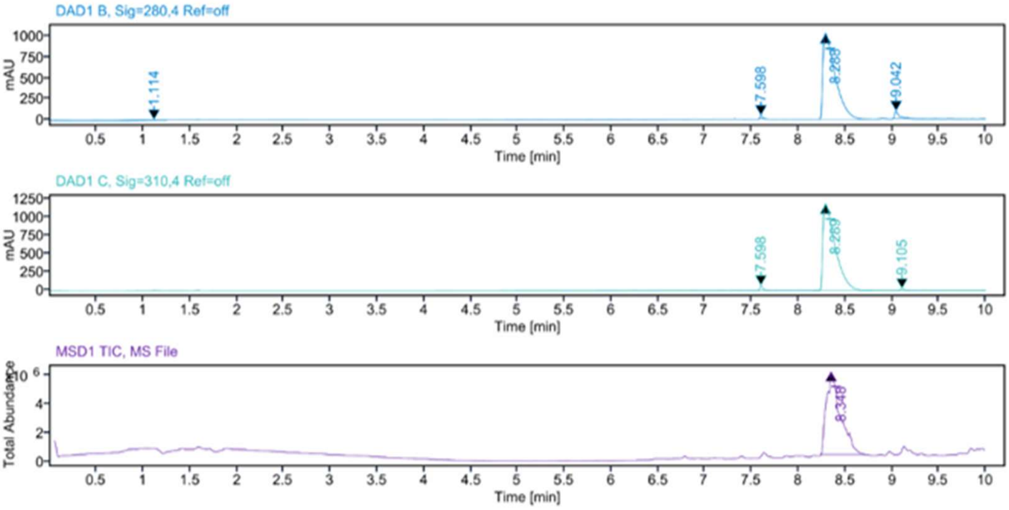


### Sample Purity

| Signal Description | DAD1 B, Sig=280,4 Ref=off |
|---|---|

| Sample Name | Name | RT | Width | Area | Area% | Height |
|---|---|---|---|---|---|---|
| MR206 Fr.1 | | 1.114 | 0.183 | 113.3712 | 1.17 | 8.6274 |
| MR206 Fr.1 | | 7.598 | 0.027 | 68.9130 | 0.71 | 40.8461 |
| MR206 Fr.1 | | 8.288 | 0.138 | 9243.1455 | 95.10 | 1010.7120 |
| MR206 Fr.1 | | 9.042 | 0.040 | 294.4494 | 3.03 | 87.5405 |

| | |
|---|---|
| Max Area% | 95.095 |
| UV Signal Purity>95% | Pass |
| Signal Description | DAD1 C, Sig=310,4 Ref=off |

| Sample Name | Name | RT | Width | Area | Area% | Height |
|---|---|---|---|---|---|---|
| MR206 Fr.1 | | 7.598 | 0.031 | 154.4546 | 1.41 | 69.3968 |
| MR206 Fr.1 | | 8.289 | 0.137 | 10770.55<br>86 | 98.04 | 1178.8027 |
| MR206 Fr.1 | | 9.105 | 0.037 | 60.7710 | 0.55 | 24.5136 |

| | |
|---|---|
| Max Area% | 98.041 |
| UV Signal Purity>95% | Pass |

### MSD Apex Spectra

**RT:**

8.348

**Sample Name:**

MR206 Fr.1

**Signal Name:**

MSD1TIC

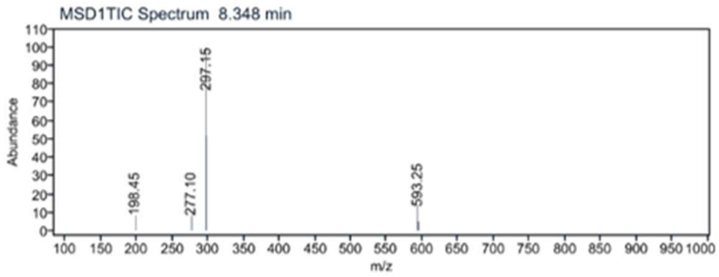

$^{1}$H NMR compound **63**:

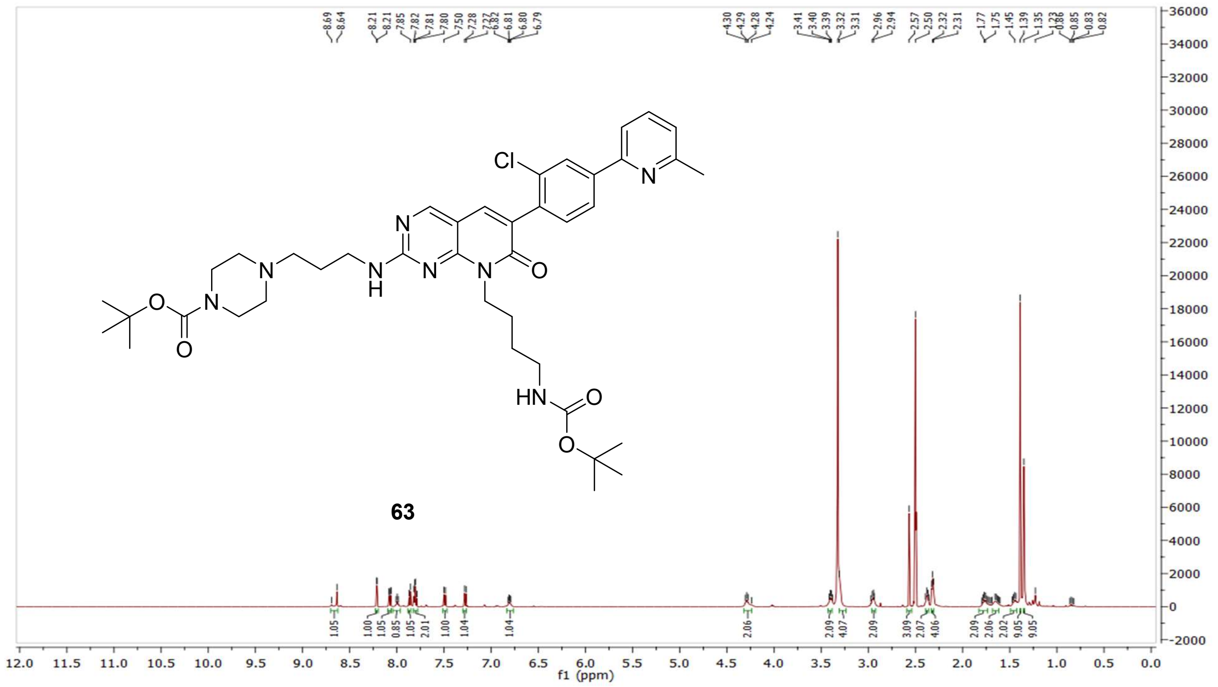

$^{13}$C NMR compound **63:**

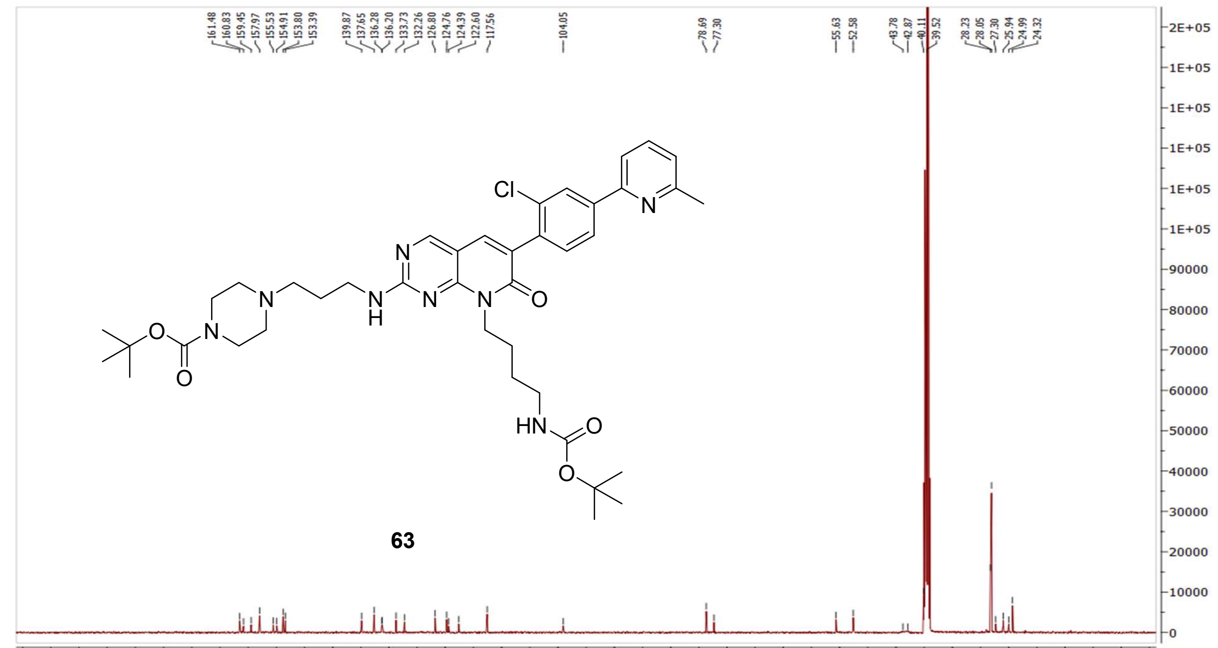

MS (ESI+) compound **63**:

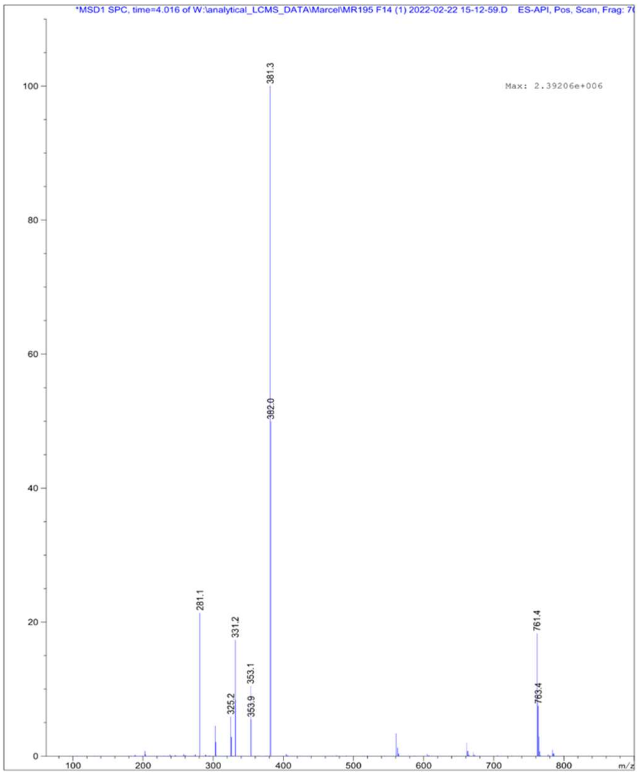


[1]H NMR compound **29**:

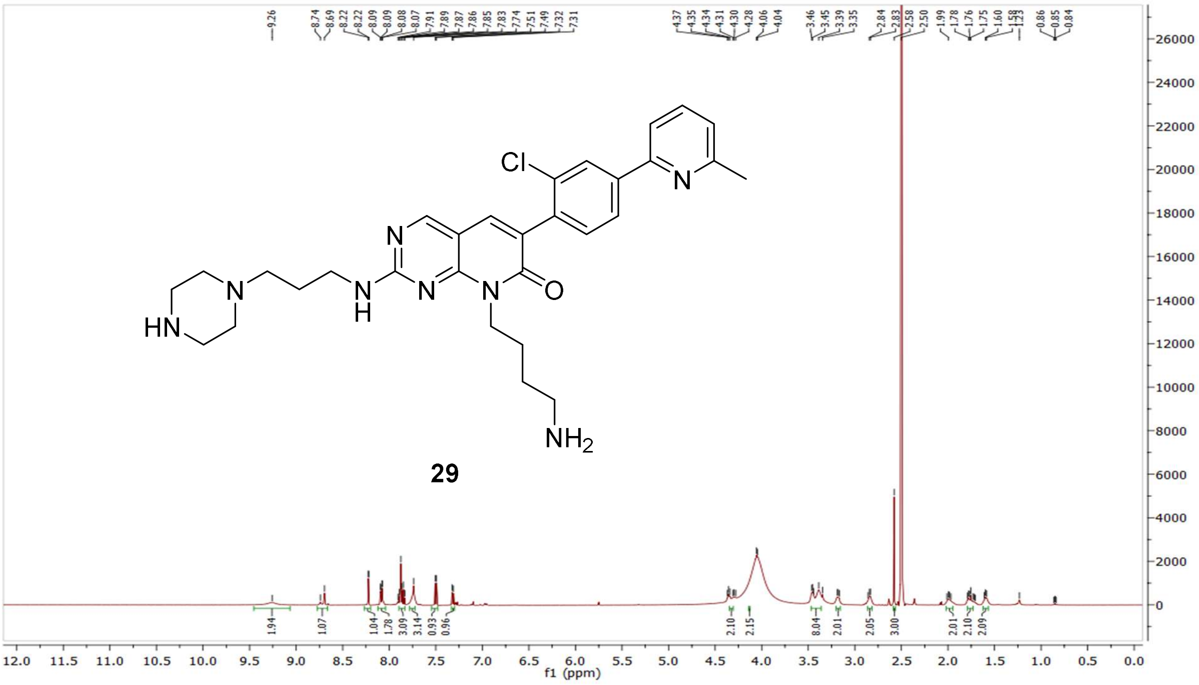

$^{13}$C NMR compound **29:**

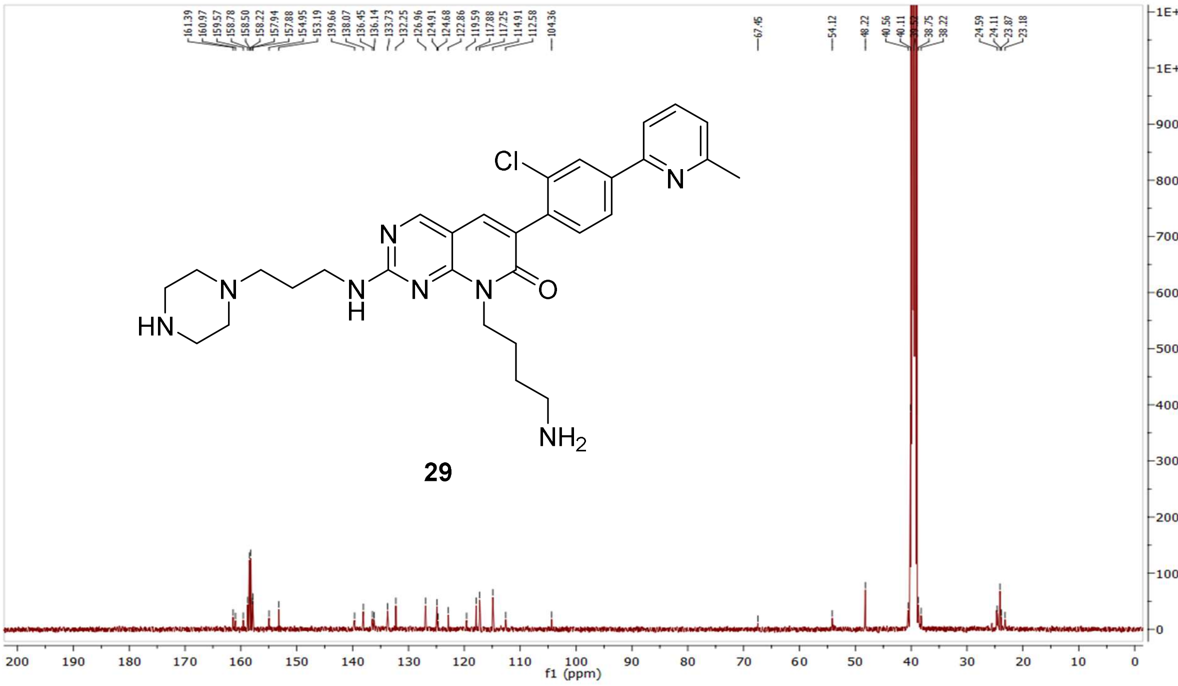


HRMS (FTMS +p MALDI) compound **29**:

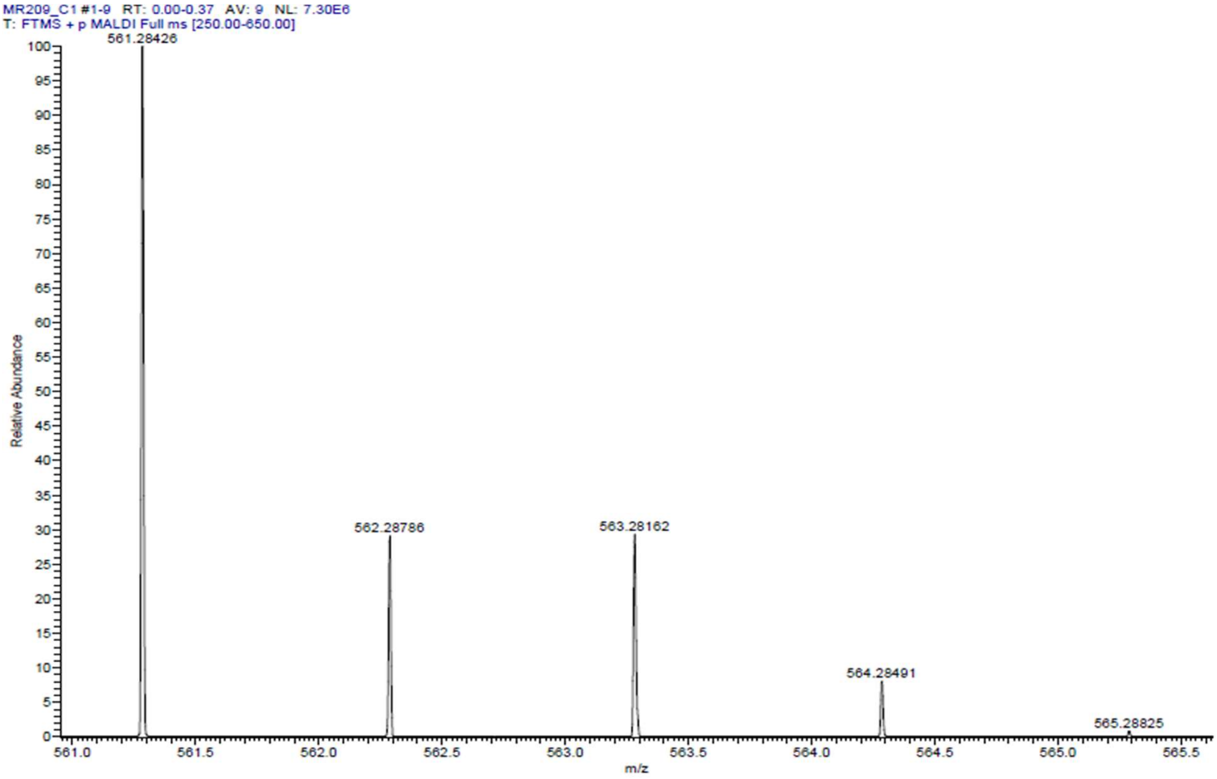

HPLC-chromatograms compound **29**:

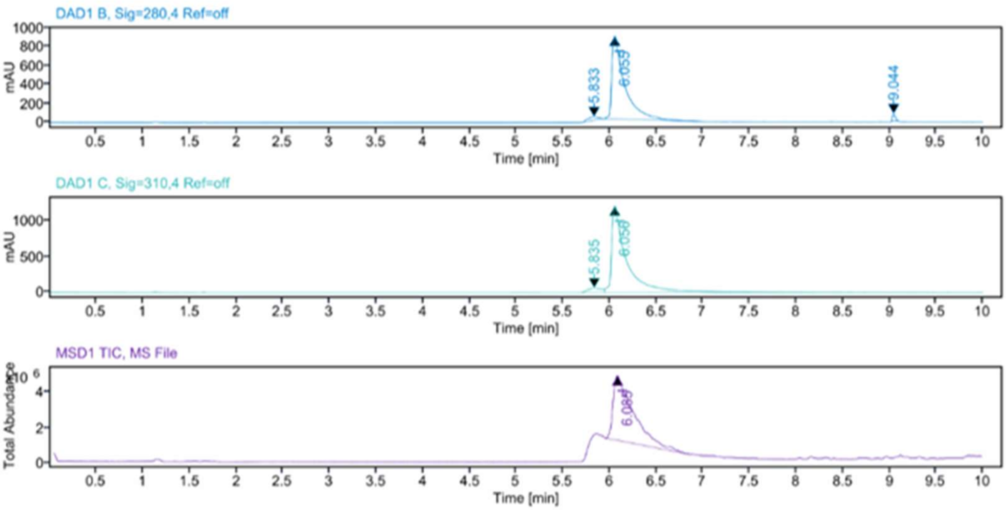


**Sample Purity**

Signal Description DAD1 B, Sig=280,4 Ref=off

| Sample Name | Name | RT | Width | Area | Area% | Height |
|---|---|---|---|---|---|---|
| MR209 | | 5.833 | 0.153 | 279.7994 | 3.18 | 35.4205 |
| MR209 | | 6.055 | 0.120 | 8357.6787 | 95.01 | 872.3410 |
| MR209 | | 9.044 | 0.035 | 159.1890 | 1.81 | 74.3656 |

Max Area% 95.010

UV Signal Purity>95% Pass

Signal Description DAD1 C, Sig=310,4 Ref=off

| Sample Name | Name | RT | Width | Area | Area% | Height |
|---|---|---|---|---|---|---|
| MR209 | | 5.835 | 0.197 | 640.5073 | 4.53 | 61.7644 |
| MR209 | | 6.056 | 0.125 | 13514.3330 | 95.47 | 1223.0841 |

Max Area% 95.475

UV Signal Purity>95% Pass

**MSD Apex Spectra**

**RT:**

6.085

**Sample Name:**

MR209

**Signal Name:**

MSD1TIC

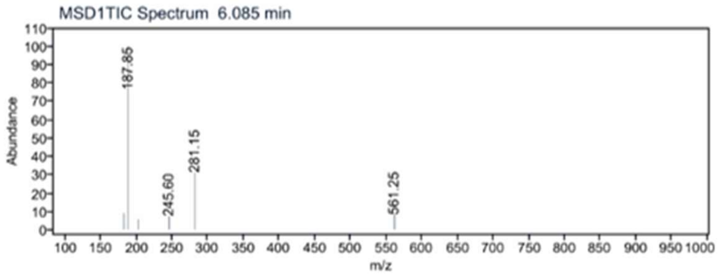

$^1$H NMR compound **64**:

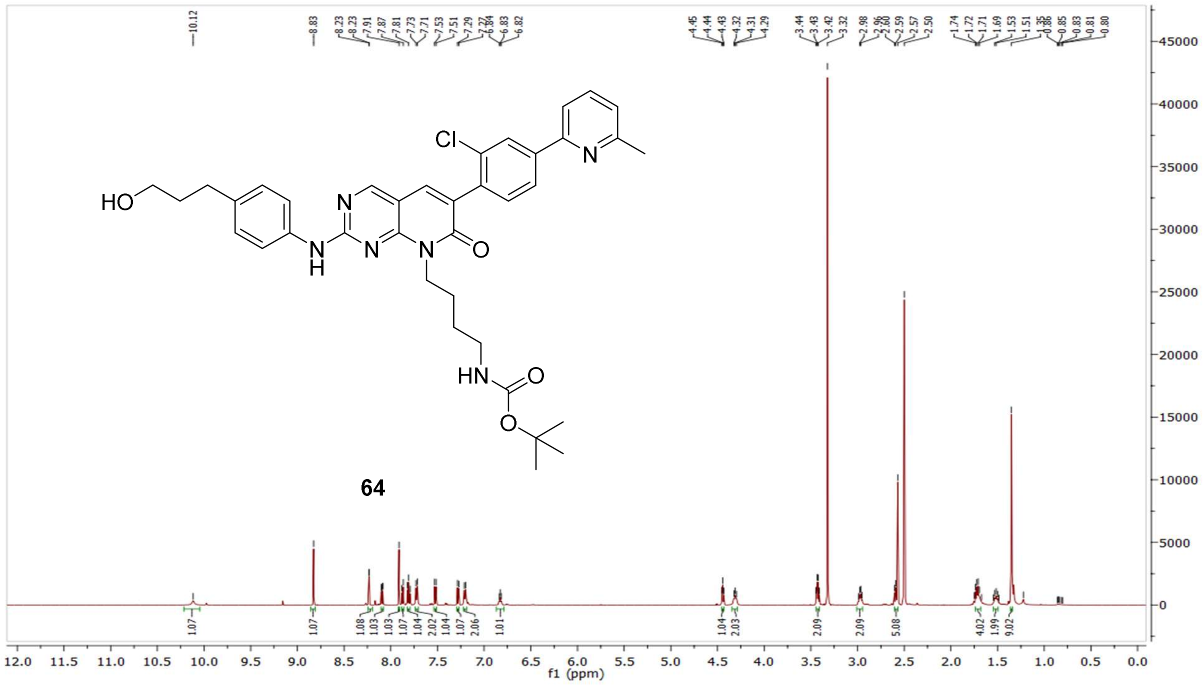

$^{13}$C NMR compound **64:**

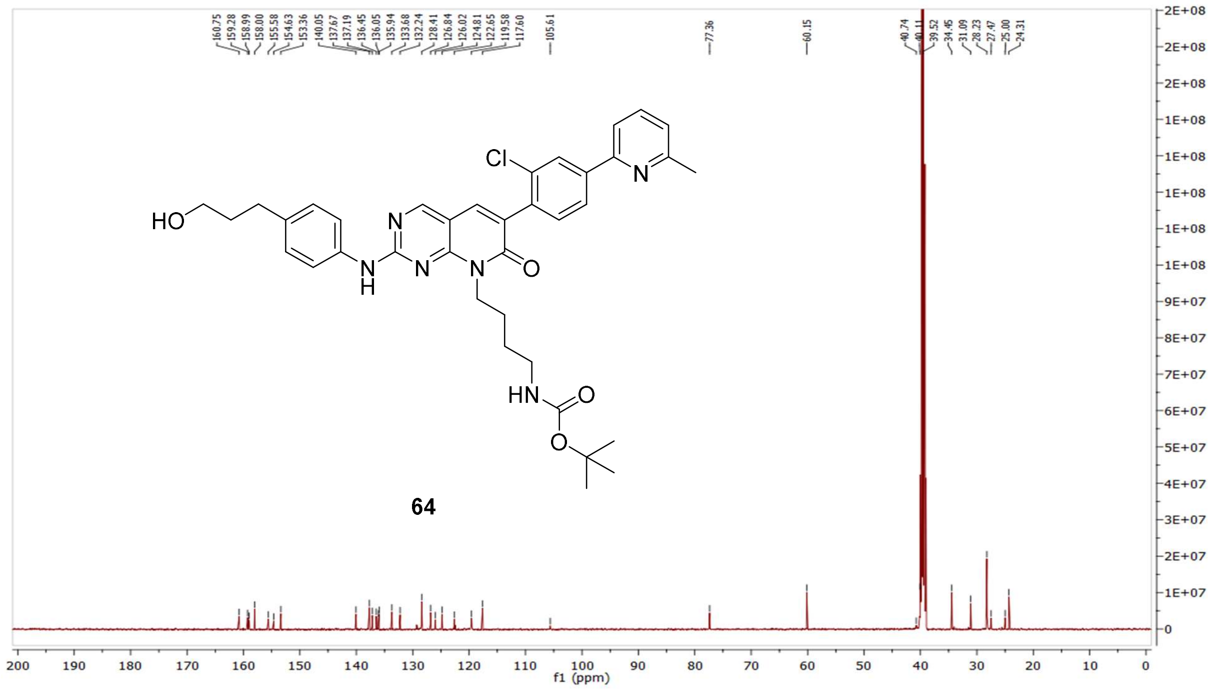

MS (ESI+) compound **64**:

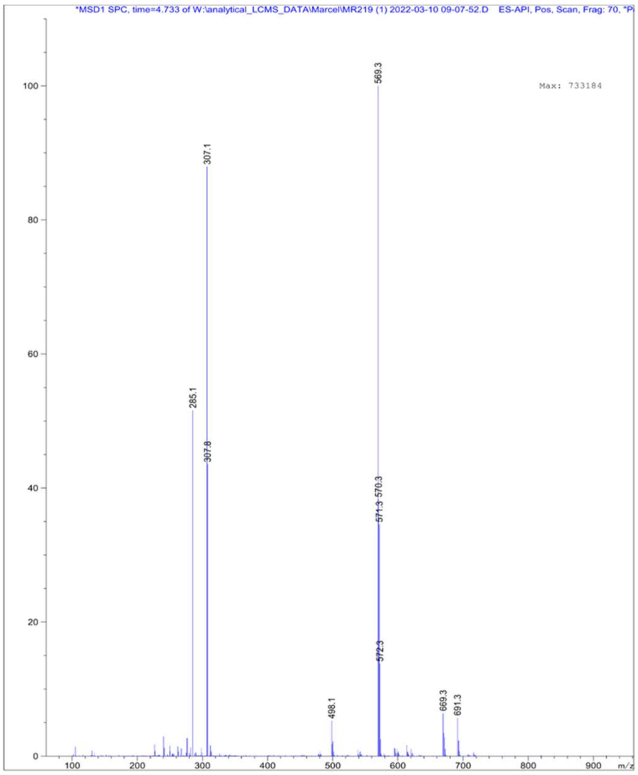


$^1$H NMR compound **65**:

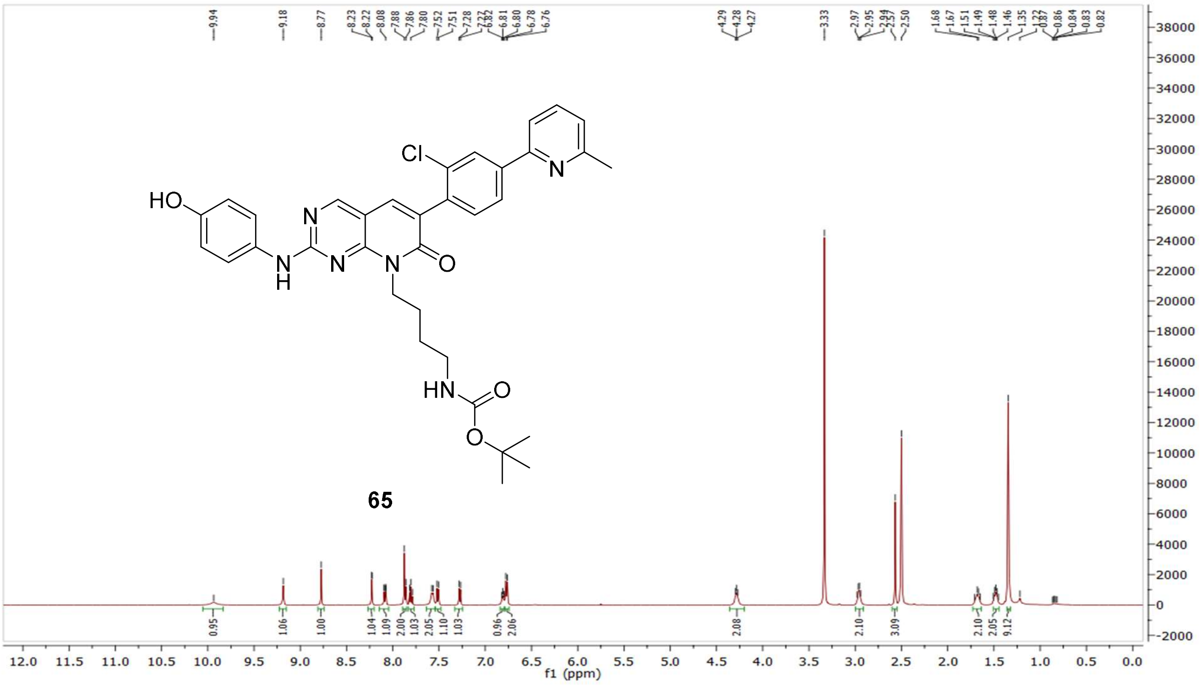

[13]C NMR compound **65:**

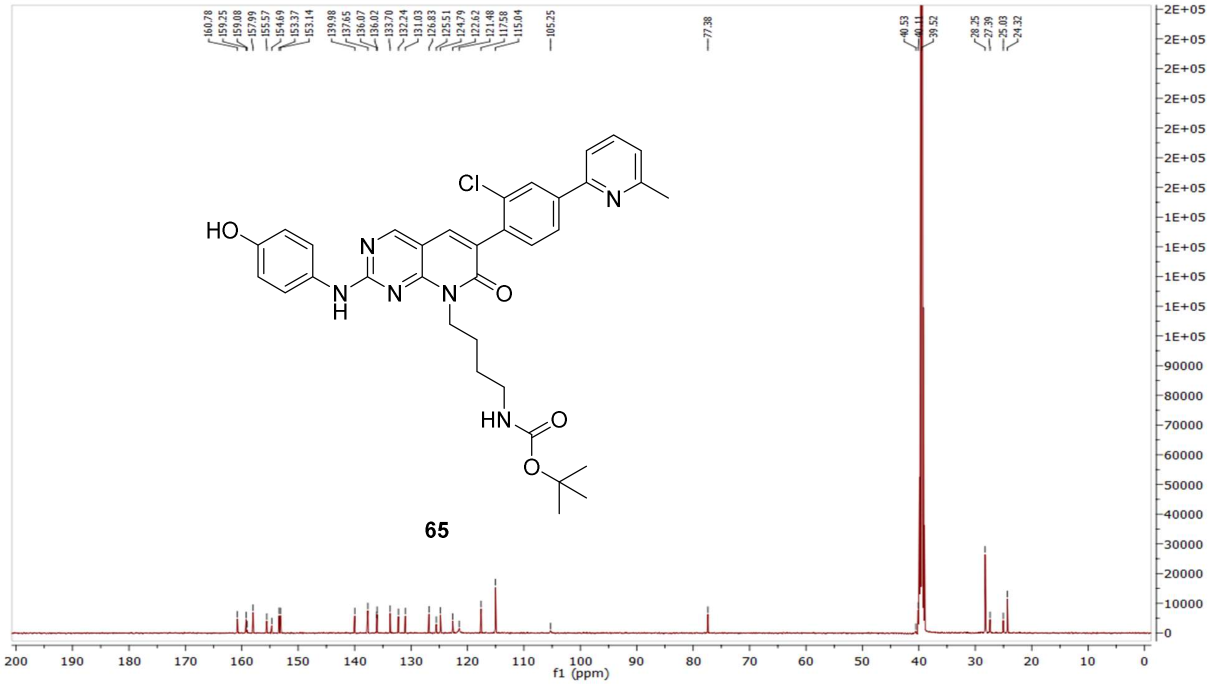

MS (ESI+) compound **65**:

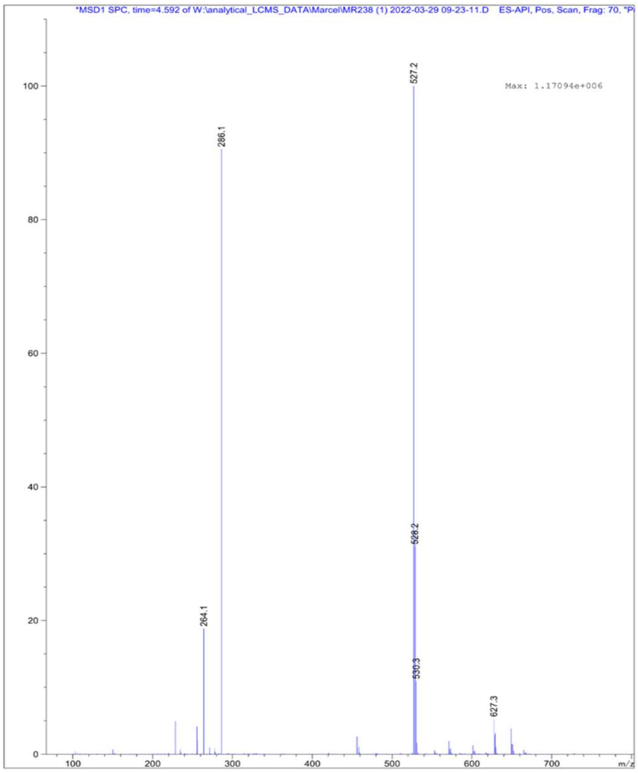

$^{1}$H NMR compound **66**:

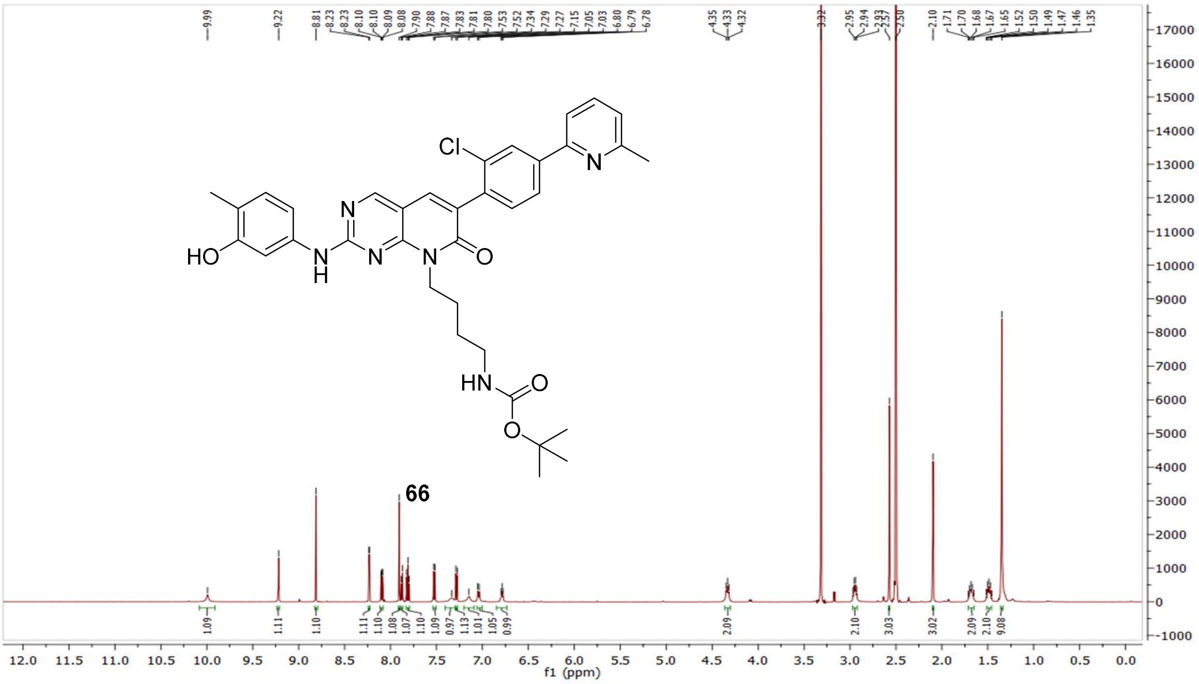


$^{13}$C NMR compound **66:**

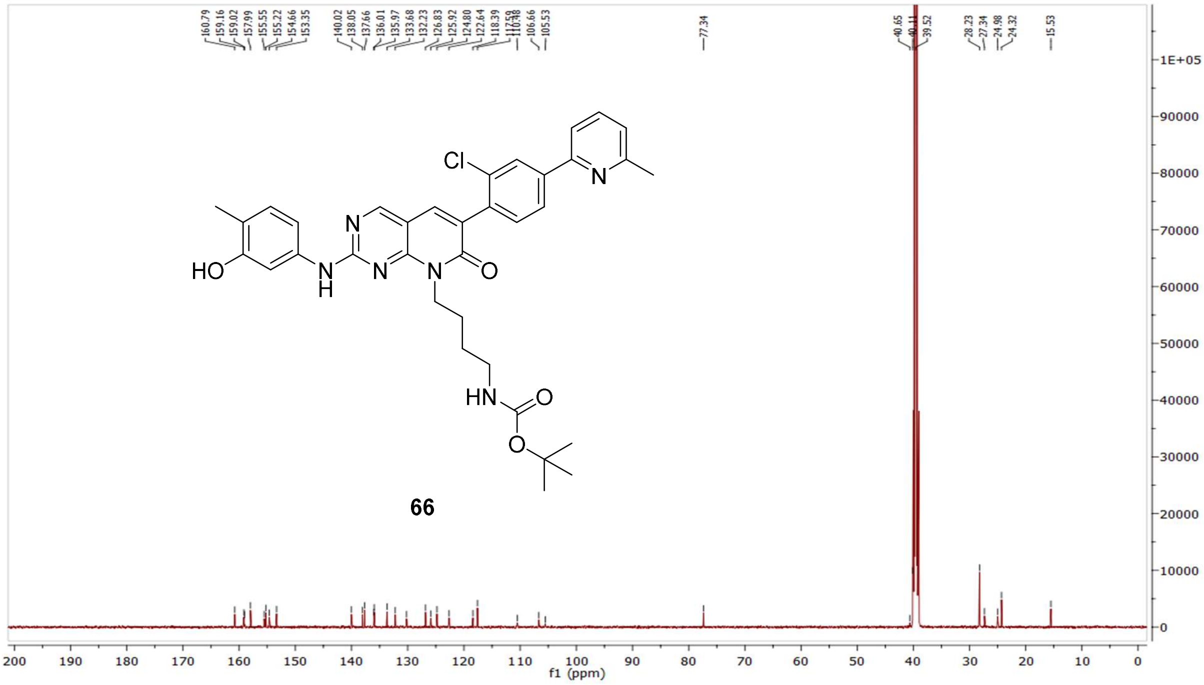

MS (ESI+) compound **66**:

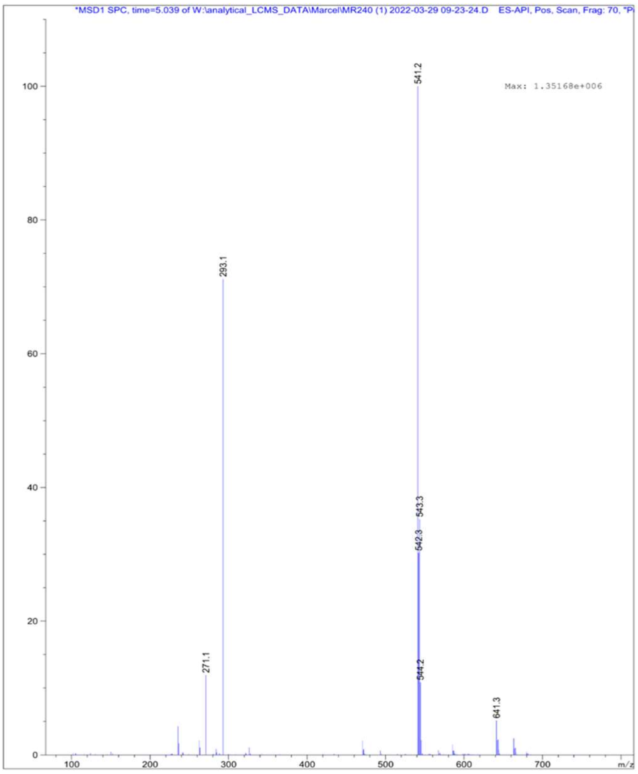

[1]H NMR compound **67**:

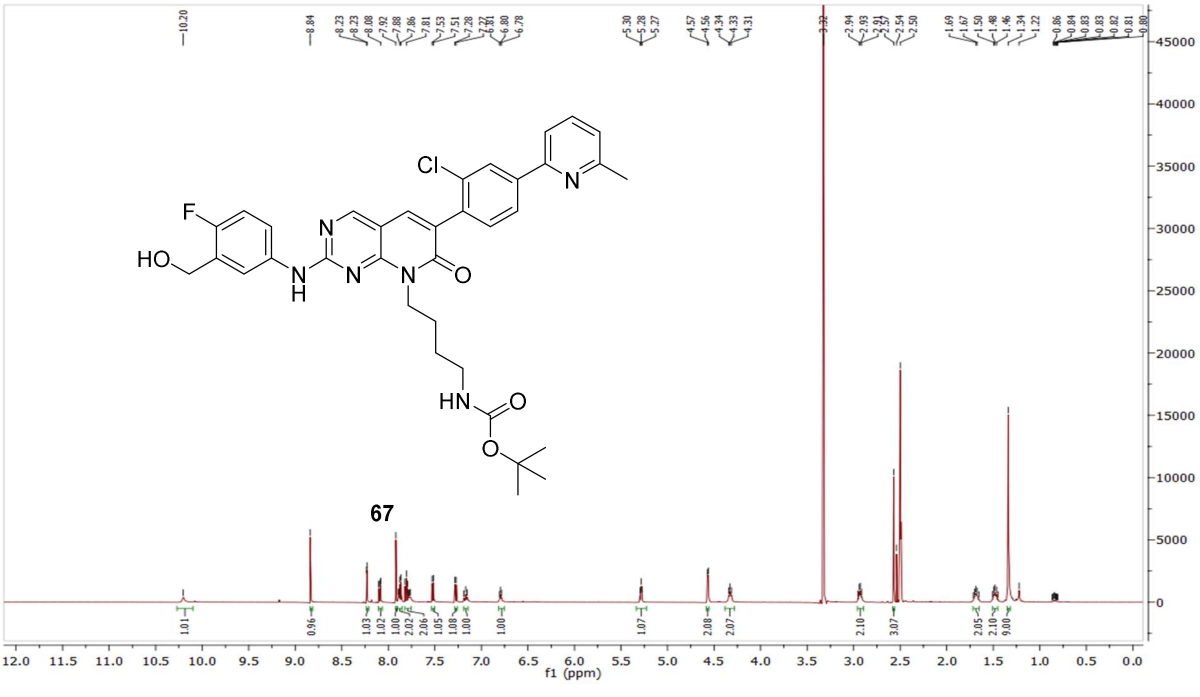

$^{13}C$ NMR compound **67:**

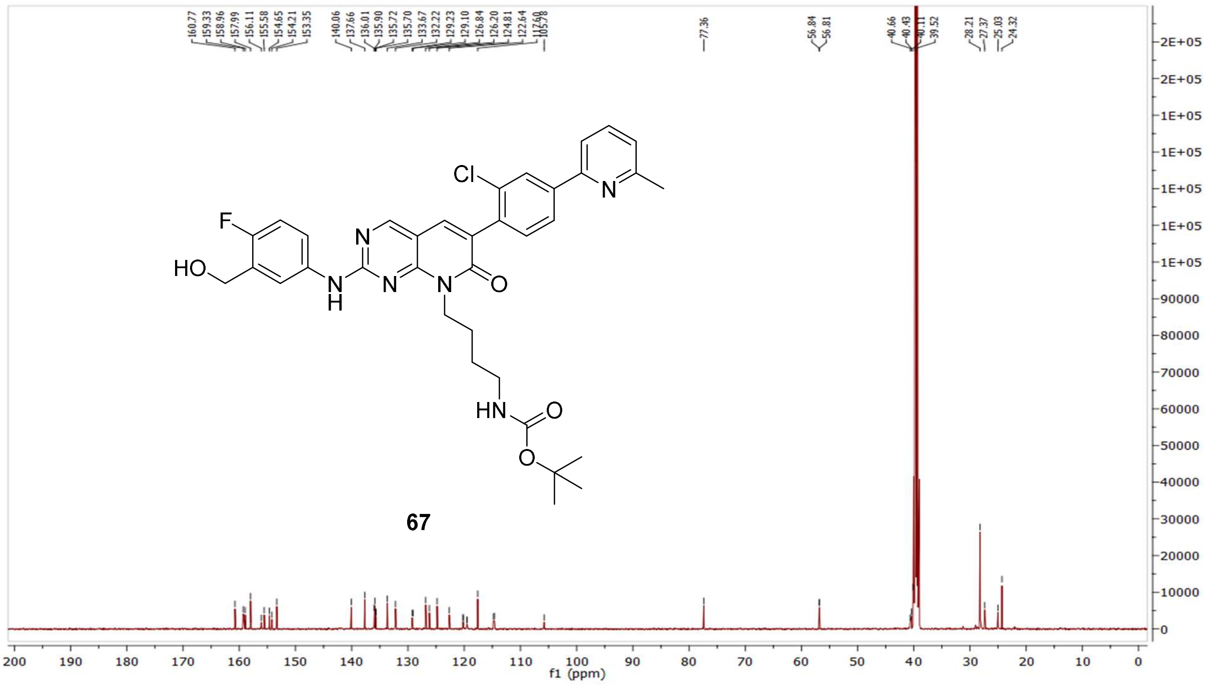


MS (ESI+) compound **67**:

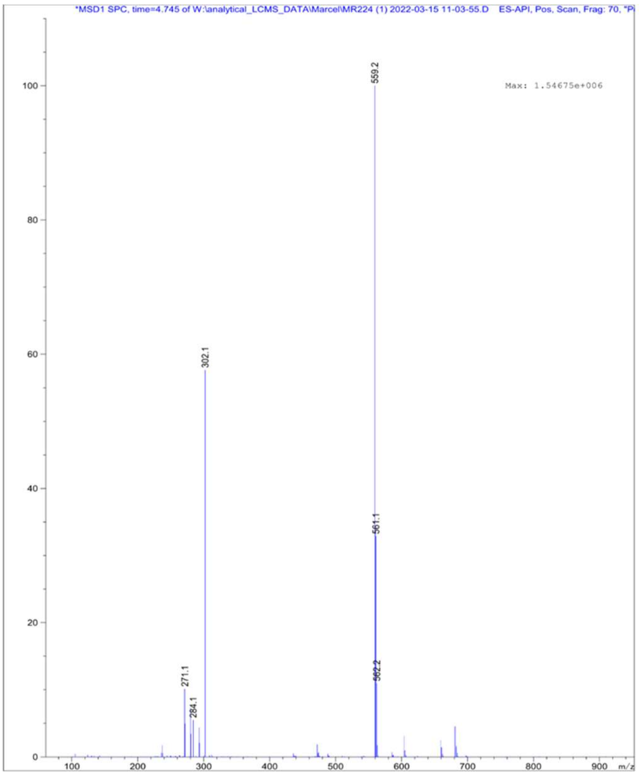

$^{1}$H NMR compound **68**:

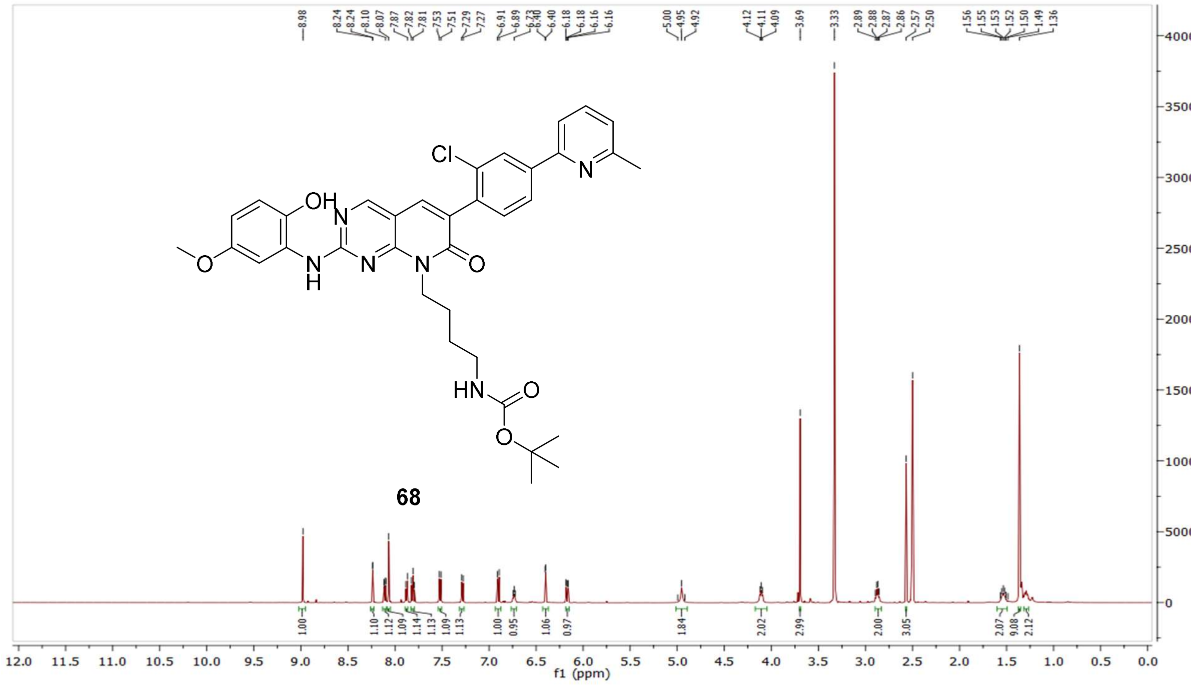

$^{13}$C NMR compound **68:**

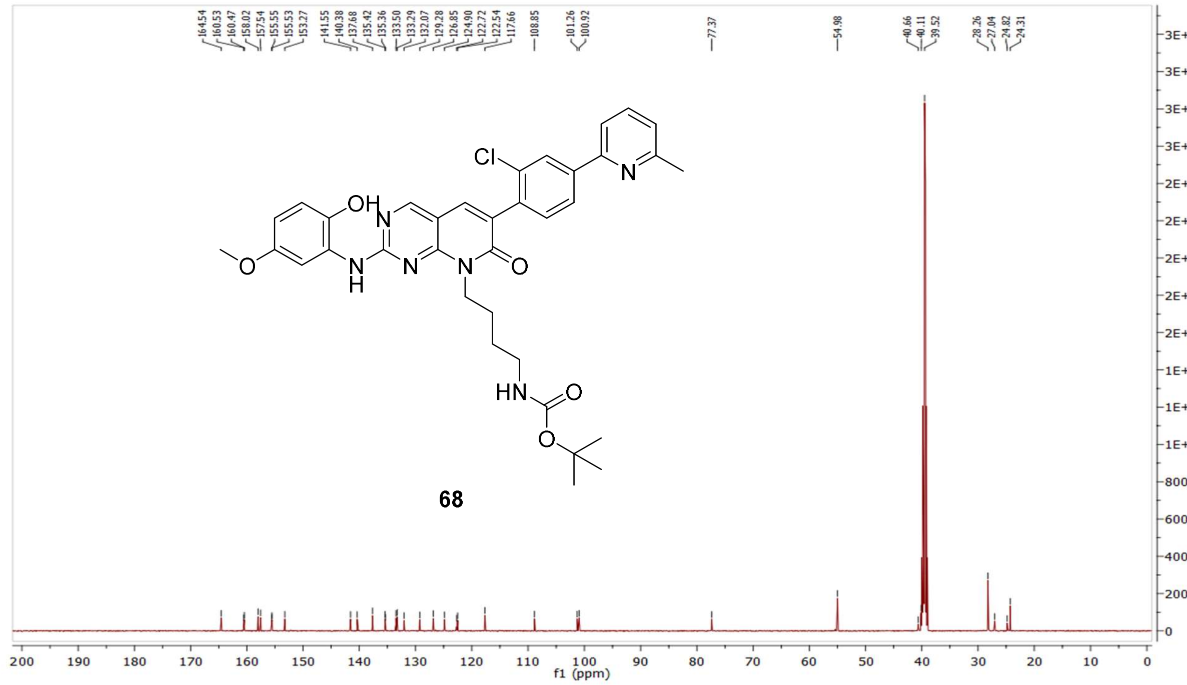

MS (ESI+) compound **68**:

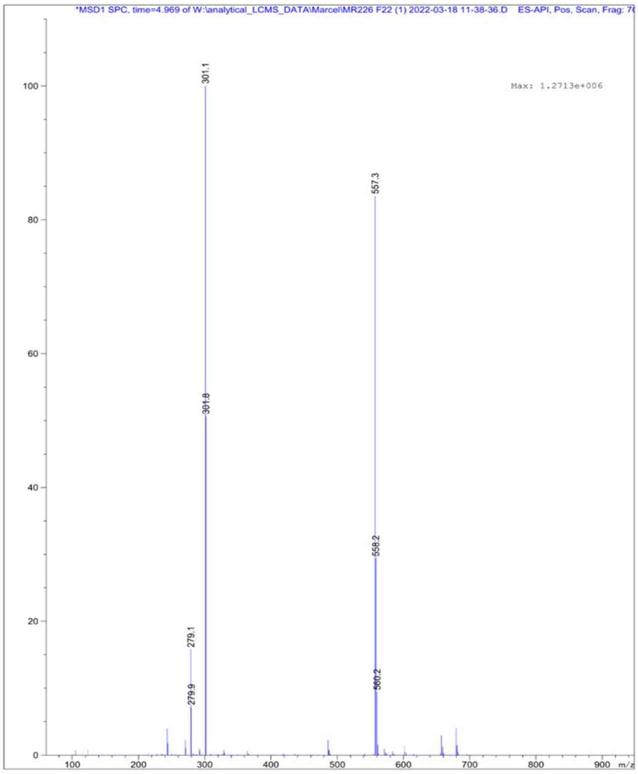

$^{1}$H NMR compound **69**:

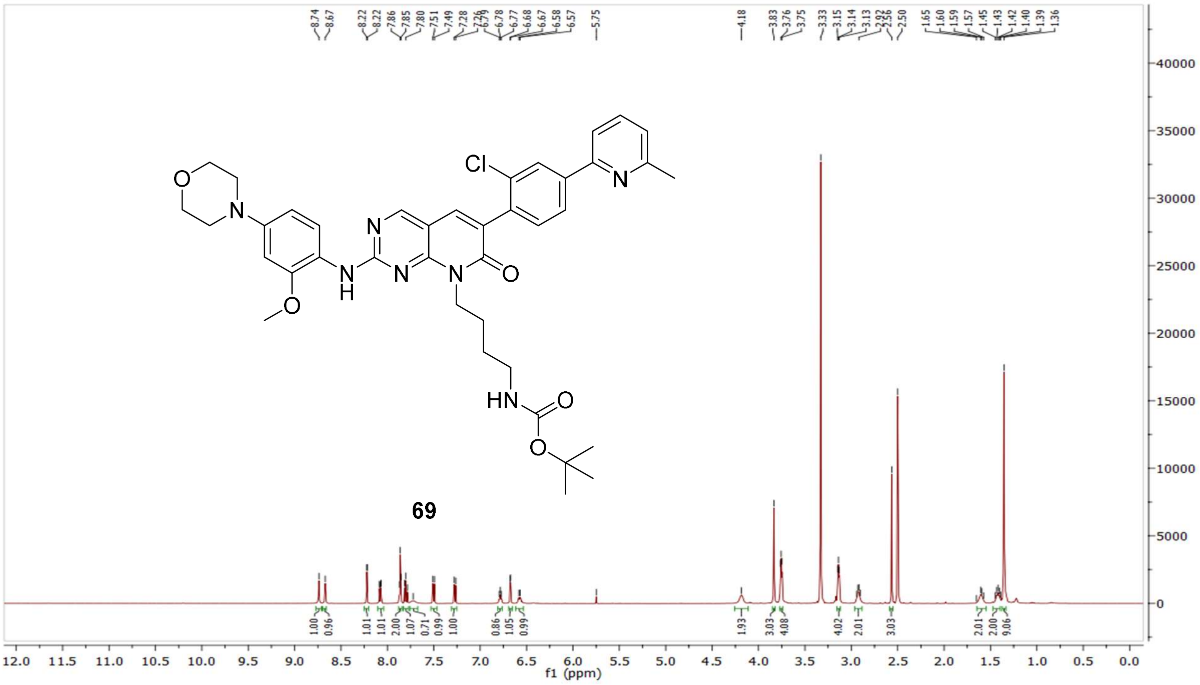

$^{13}$C NMR compound **69:**

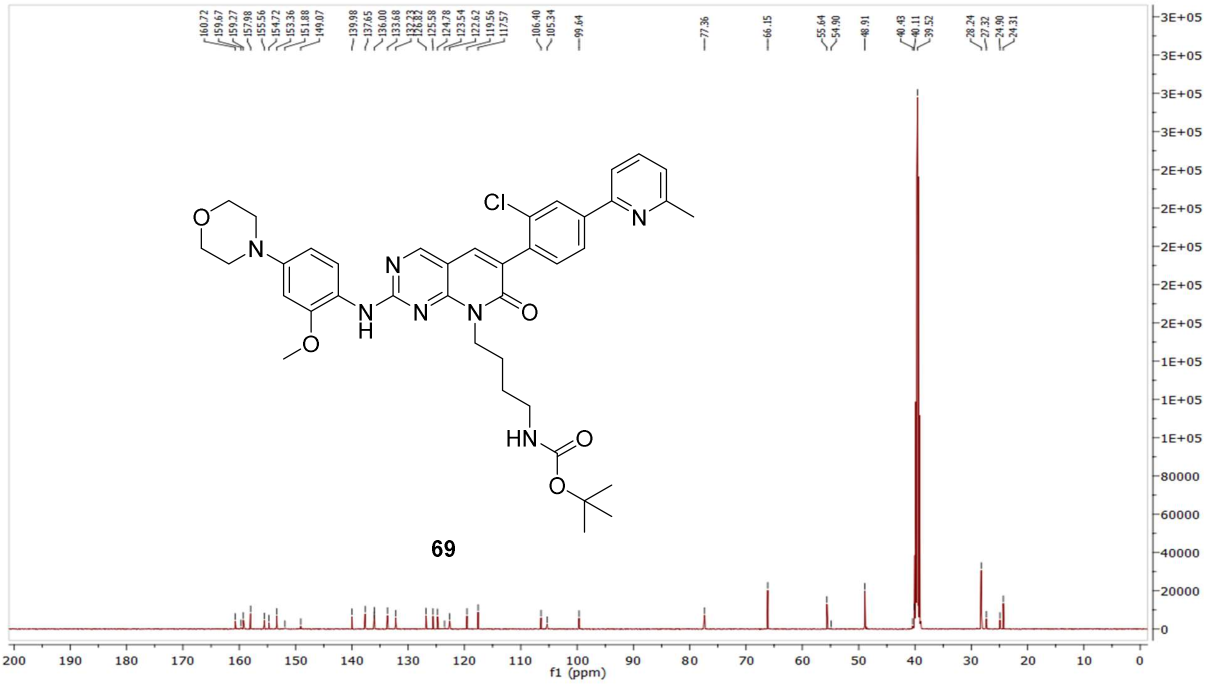

MS (ESI+) compound **69**:

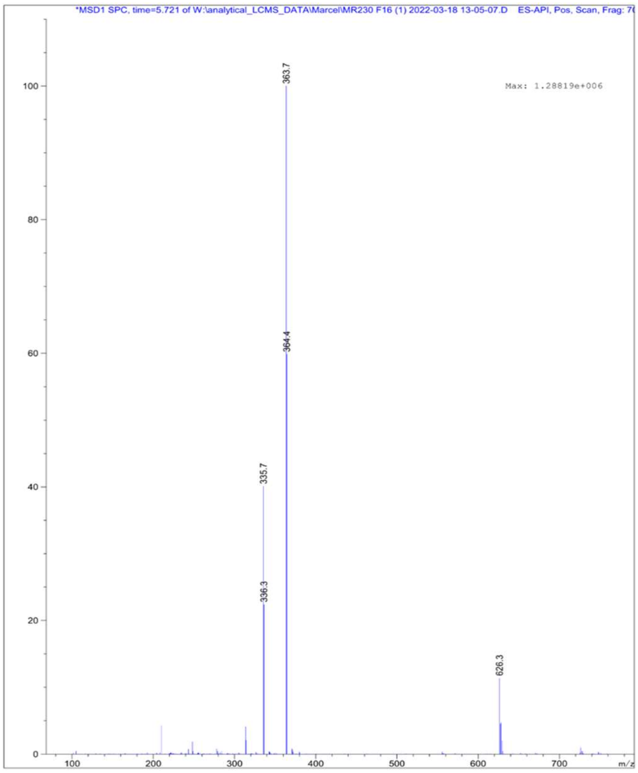

$^1$H NMR compound **70**:

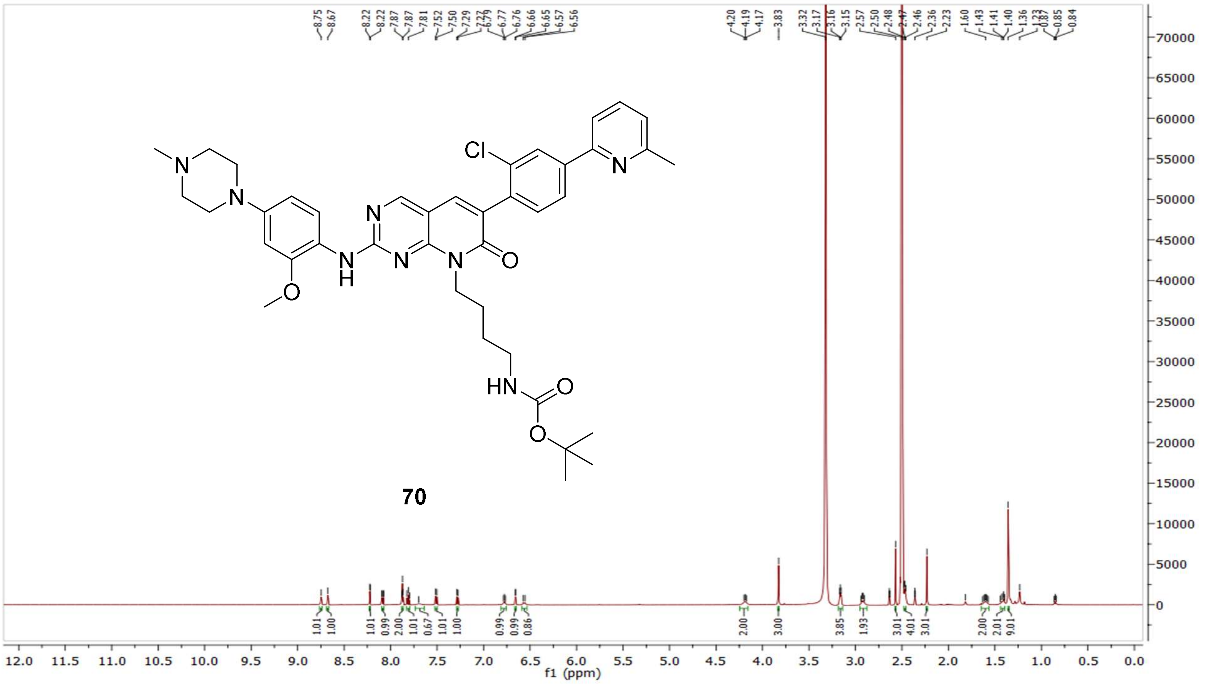

$^{13}$C NMR compound **70:**

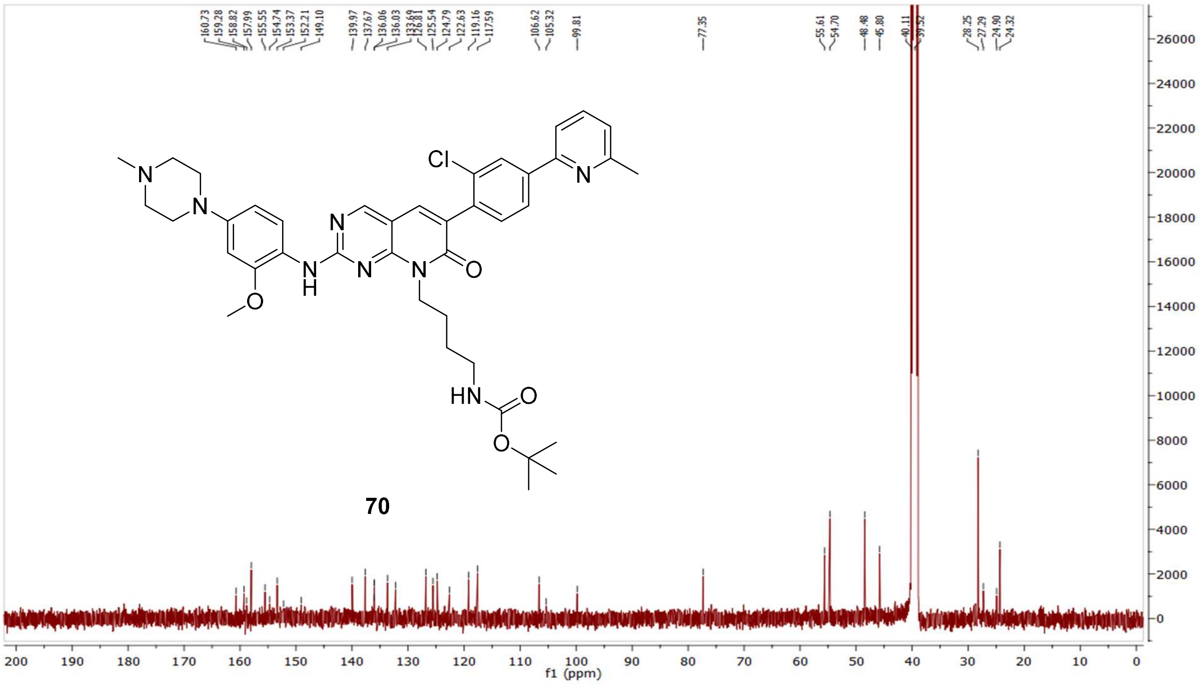

MS (ESI+) compound **70:**

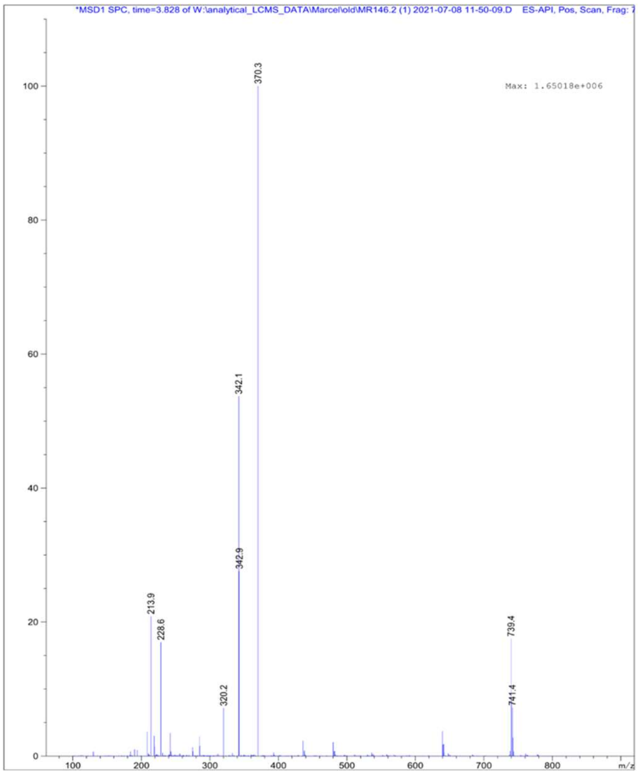


[1]H NMR compound **30**:

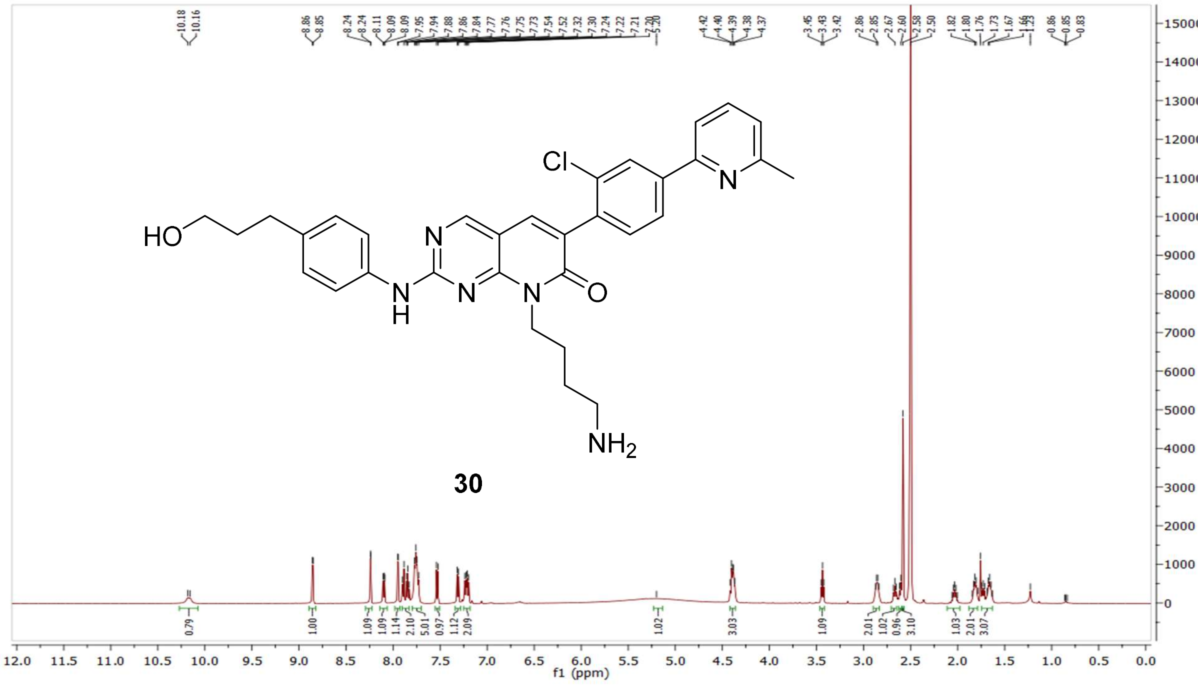

$^{13}$C NMR compound **30:**

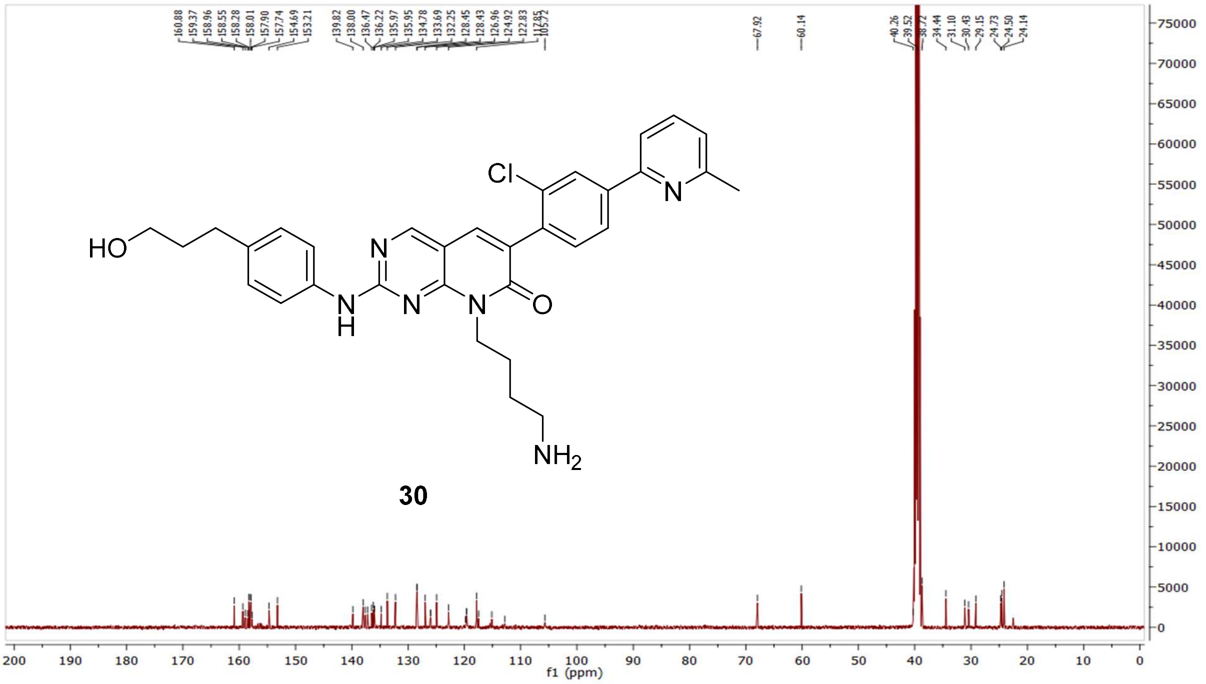


$^{1}$H-$^{1}$H COSY NMR compound **30:**

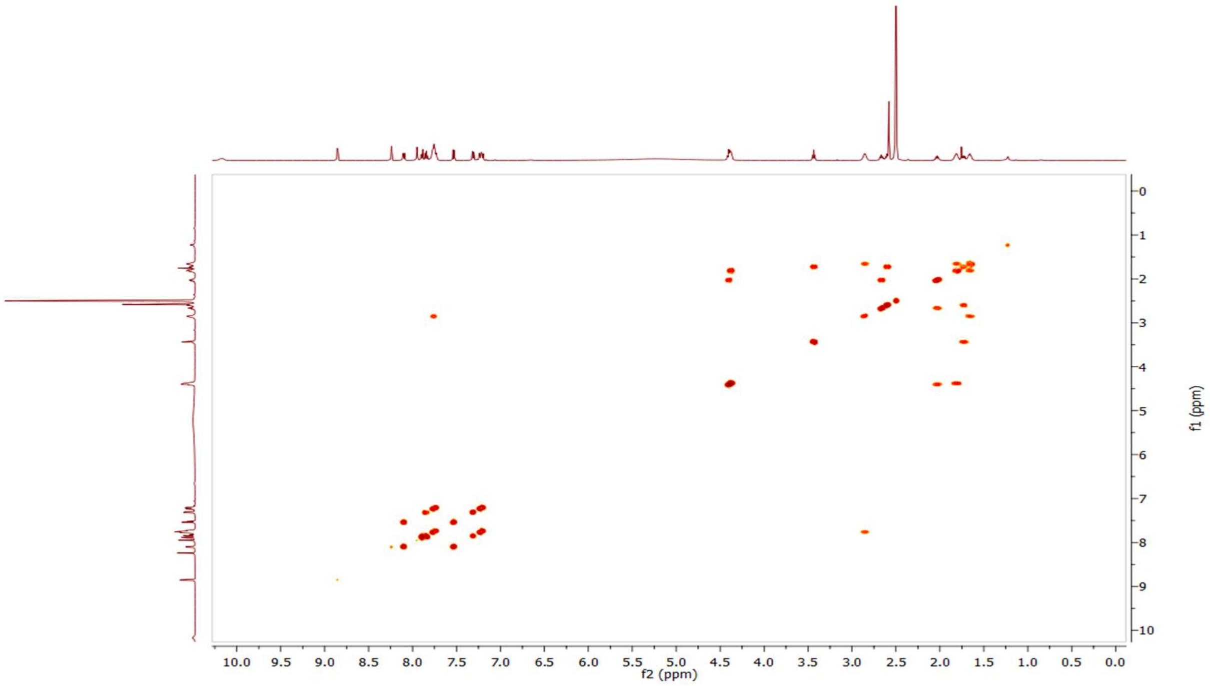

$^1H$-$^{13}C$ HSQC NMR compound **30:**

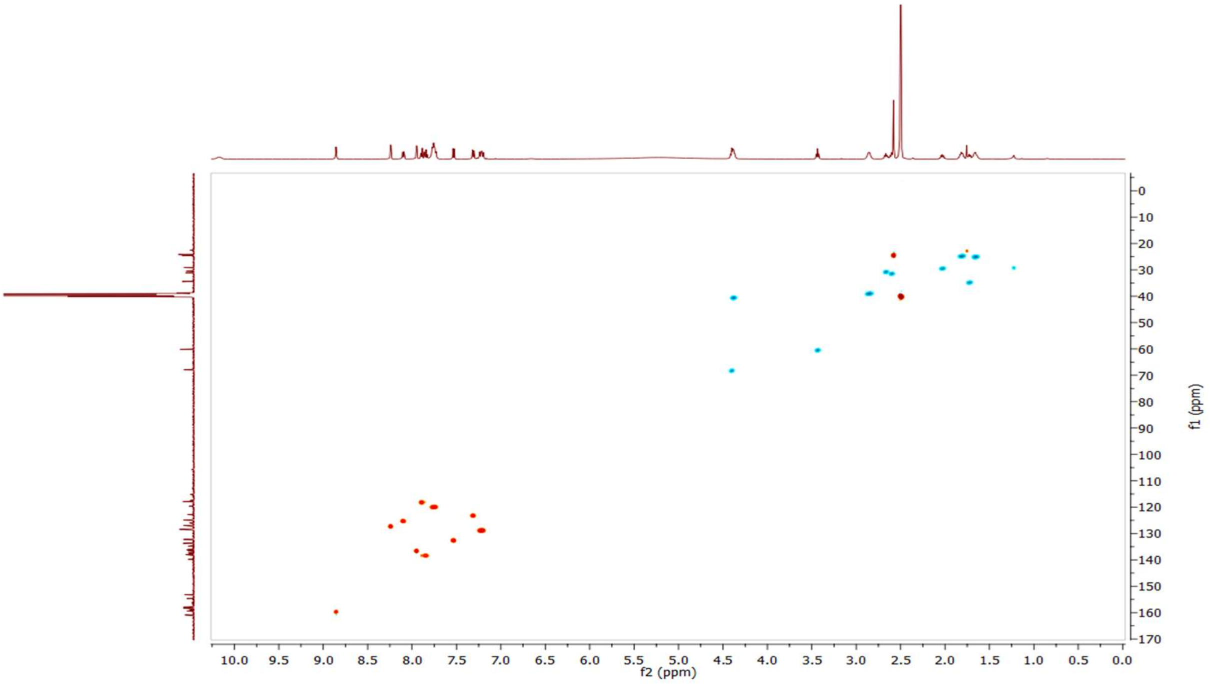


HRMS (FTMS +p MALDI) compound **30**:

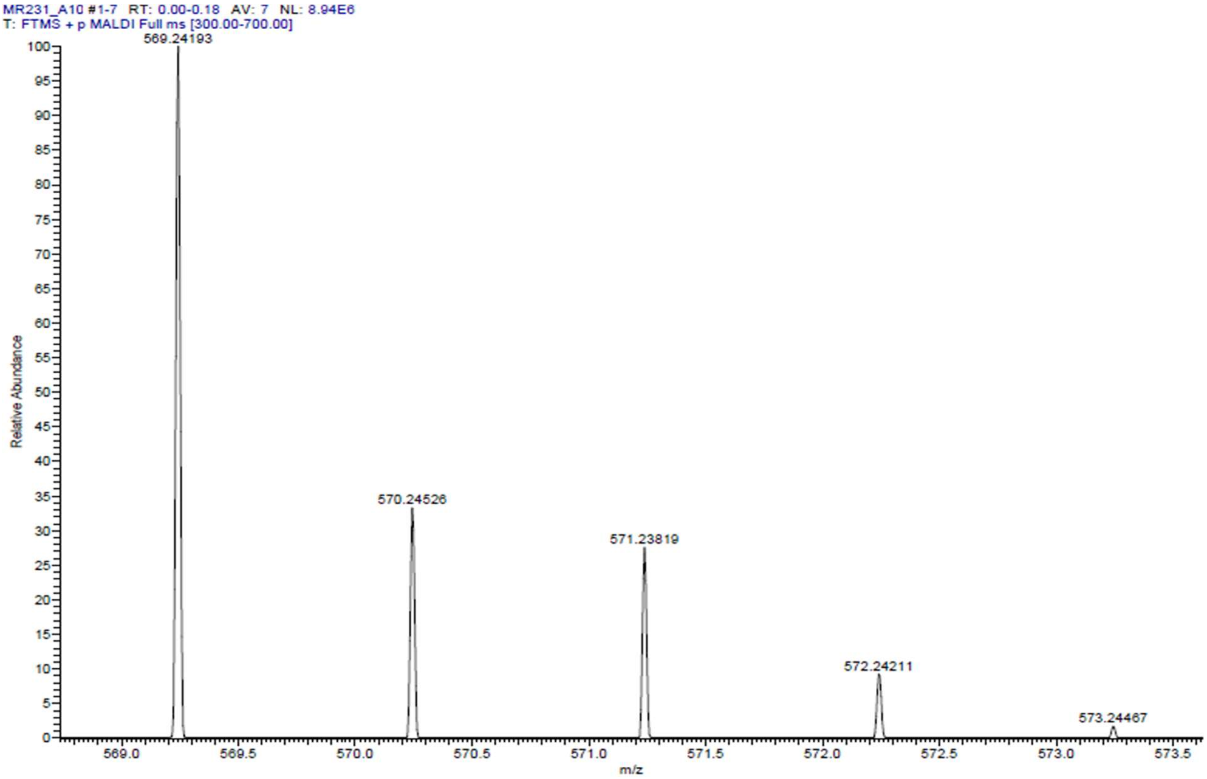

HPLC-chromatograms compound **30**:

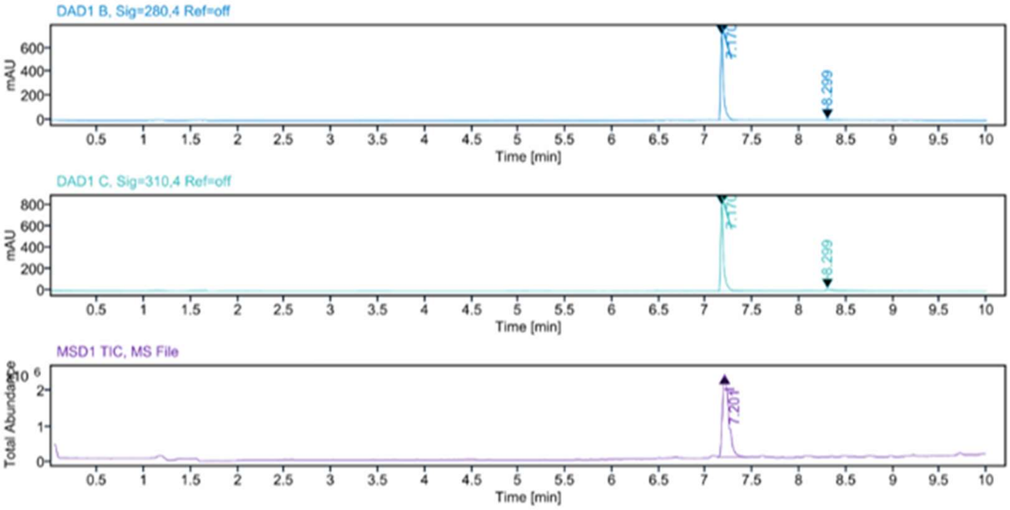


**Sample Purity**

Signal Description DAD1 B, Sig=280,4 Ref=off

| Sample Name | Name | RT | Width | Area | Area% | Height |
|---|---|---|---|---|---|---|
| MR231 Fr.2 | | 7.170 | 0.028 | 1542.3419 | 97.30 | 711.5419 |
| MR231 Fr.2 | | 8.299 | 0.052 | 42.7809 | 2.70 | 8.5474 |

Max Area% 97.301

UV Signal Purity>95% Pass

Signal Description DAD1 C, Sig=310,4 Ref=off

| Sample Name | Name | RT | Width | Area | Area% | Height |
|---|---|---|---|---|---|---|
| MR231 Fr.2 | | 7.170 | 0.028 | 1726.5173 | 95.32 | 801.6934 |
| MR231 Fr.2 | | 8.299 | 0.051 | 84.8554 | 4.68 | 18.4176 |

Max Area% 95.315

UV Signal Purity>95% Pass

**MSD Apex Spectra**

**RT:**

7.201

**Sample Name:**

MR231 Fr.2

**Signal Name:**

MSD1TIC

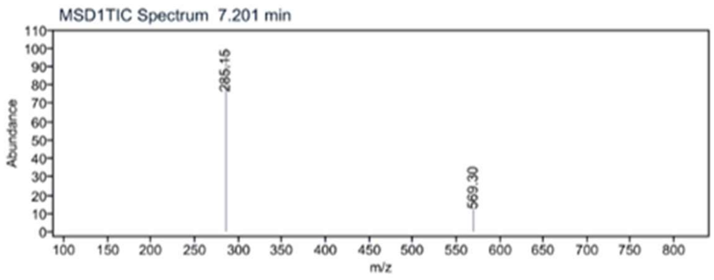

$^{1}H$ NMR compound **31**:

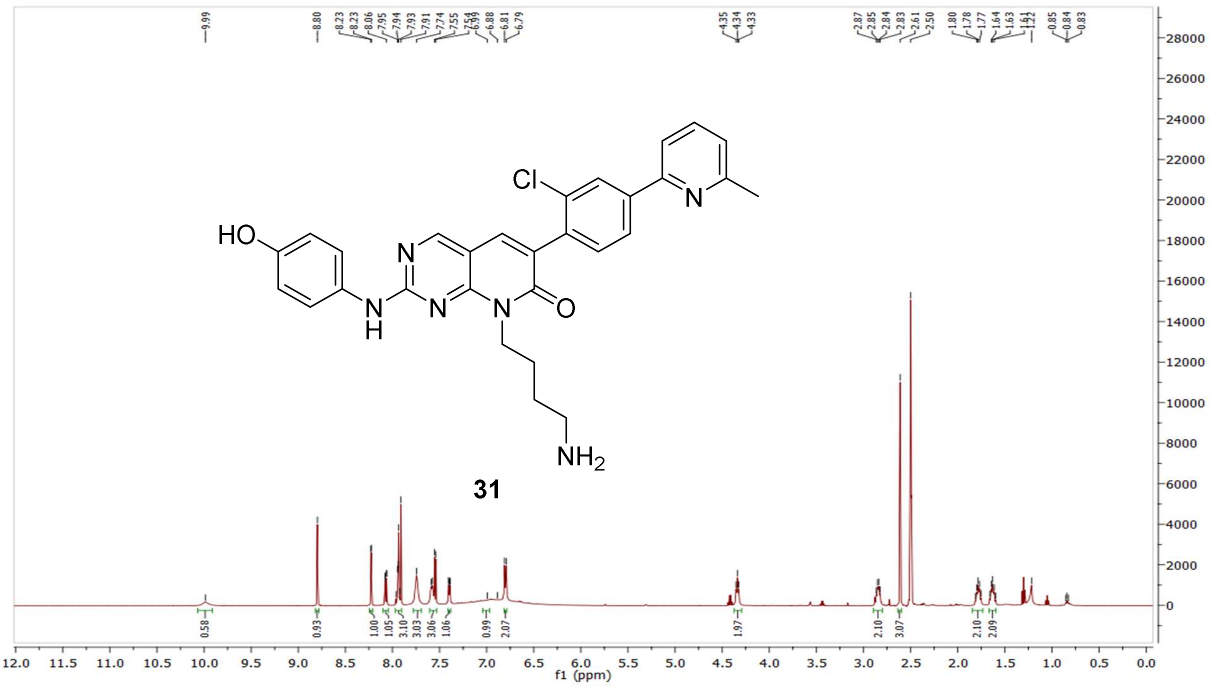

$^{13}C$ NMR compound **31:**

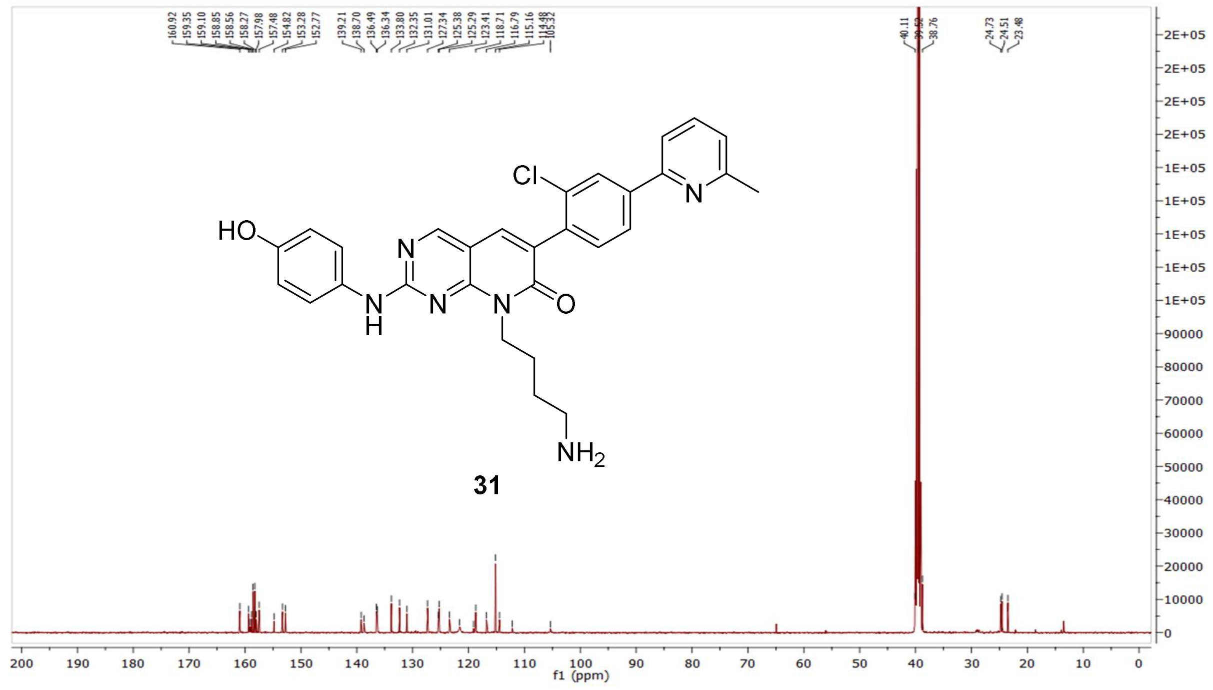

HRMS (FTMS +p MALDI) compound **31**:

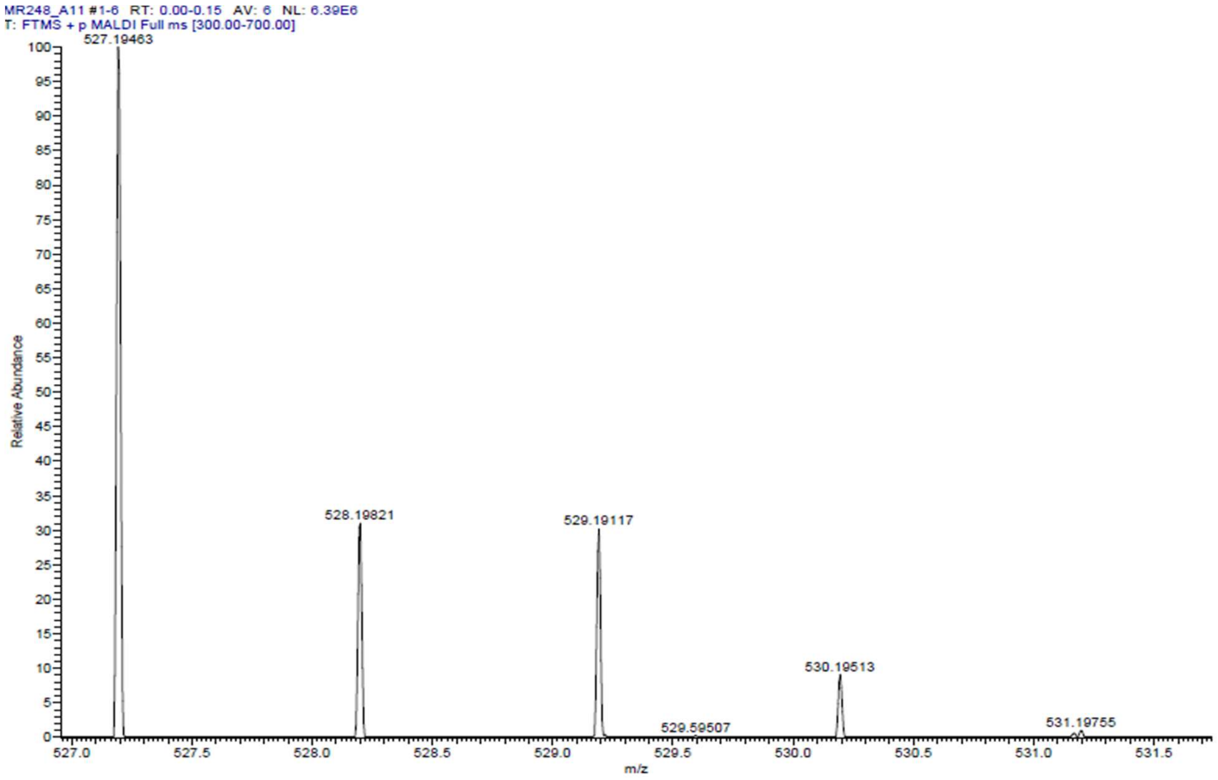

HPLC-chromatograms compound **31**:

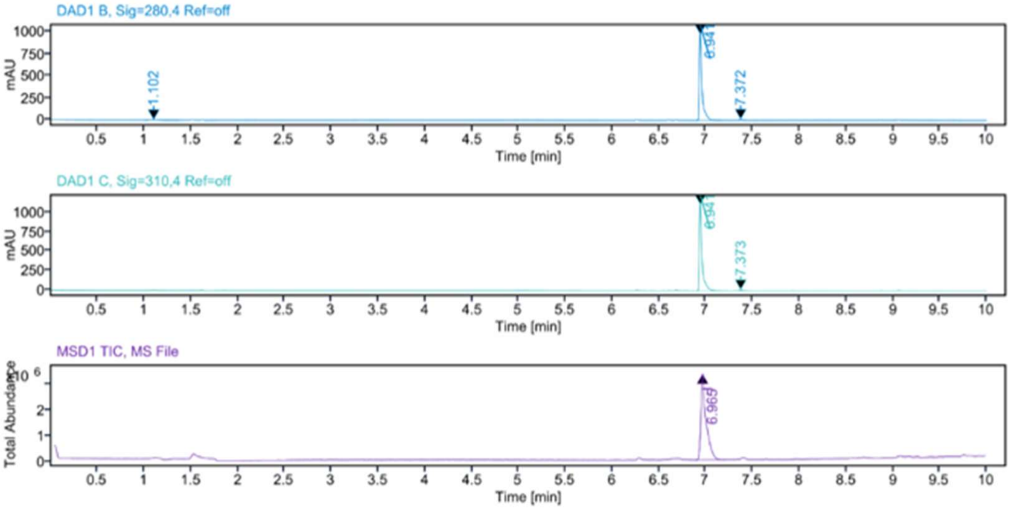


**Sample Purity**

Signal Description DAD1 B, Sig=280,4 Ref=off

| Sample Name | Name | RT | Width | Area | Area% | Height |
|---|---|---|---|---|---|---|
| MR248 | | 1.102 | 0.034 | 24.5789 | 1.17 | 6.5070 |
| MR248 | | 6.941 | 0.027 | 2063.8503 | 97.84 | 982.1979 |
| MR248 | | 7.372 | 0.023 | 21.0427 | 1.00 | 13.1992 |

Max Area% 97.837

UV Signal Purity>95% Pass

Signal Description DAD1 C, Sig=310,4 Ref=off

| Sample Name | Name | RT | Width | Area | Area% | Height |
|---|---|---|---|---|---|---|
| MR248 | | 6.941 | 0.027 | 2327.8308 | 98.99 | 1104.7898 |
| MR248 | | 7.373 | 0.023 | 23.8531 | 1.01 | 13.0145 |

Max Area% 98.986

UV Signal Purity>95% Pass

**MSD Apex Spectra**

**RT:**

6.965

**Sample Name:**

MR248

**Signal Name:**

MSD1TIC

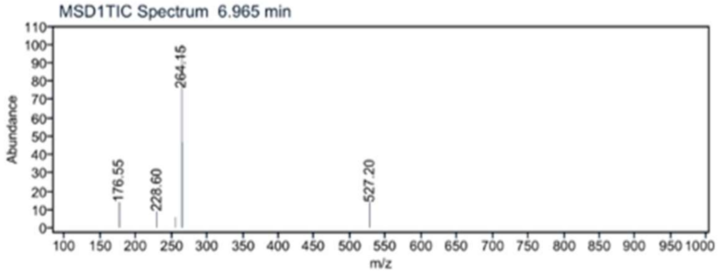

$^{1}$H NMR compound **32**:

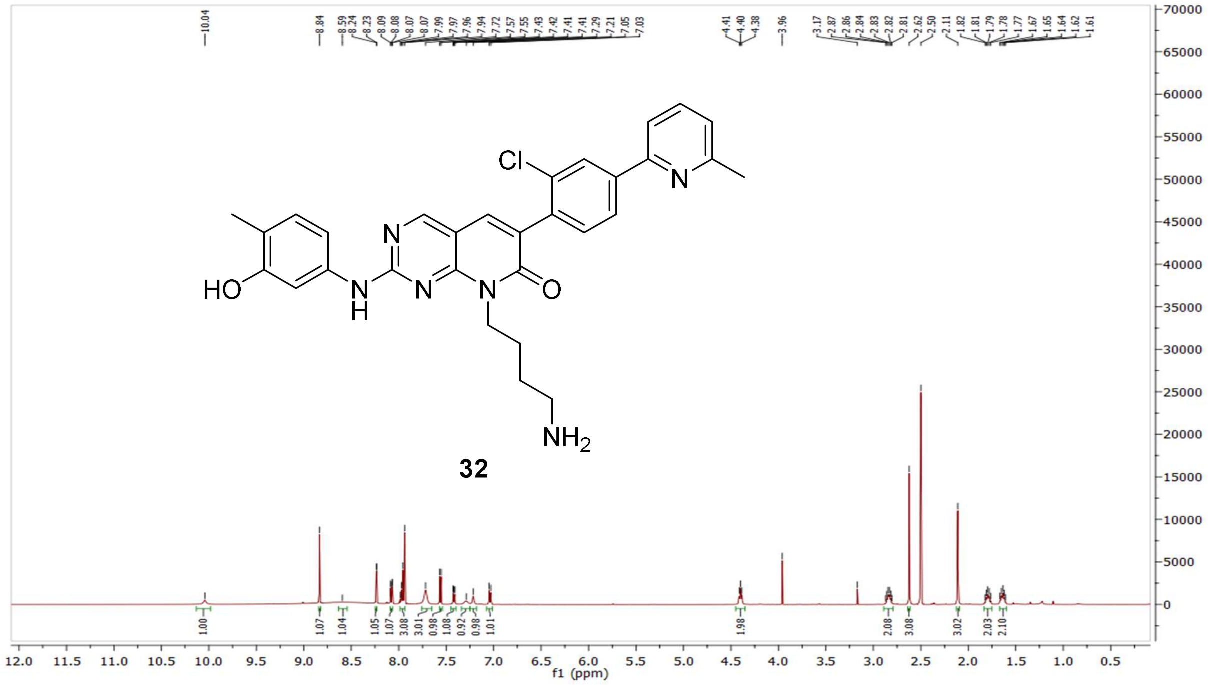

$^{13}$C NMR compound **32:**

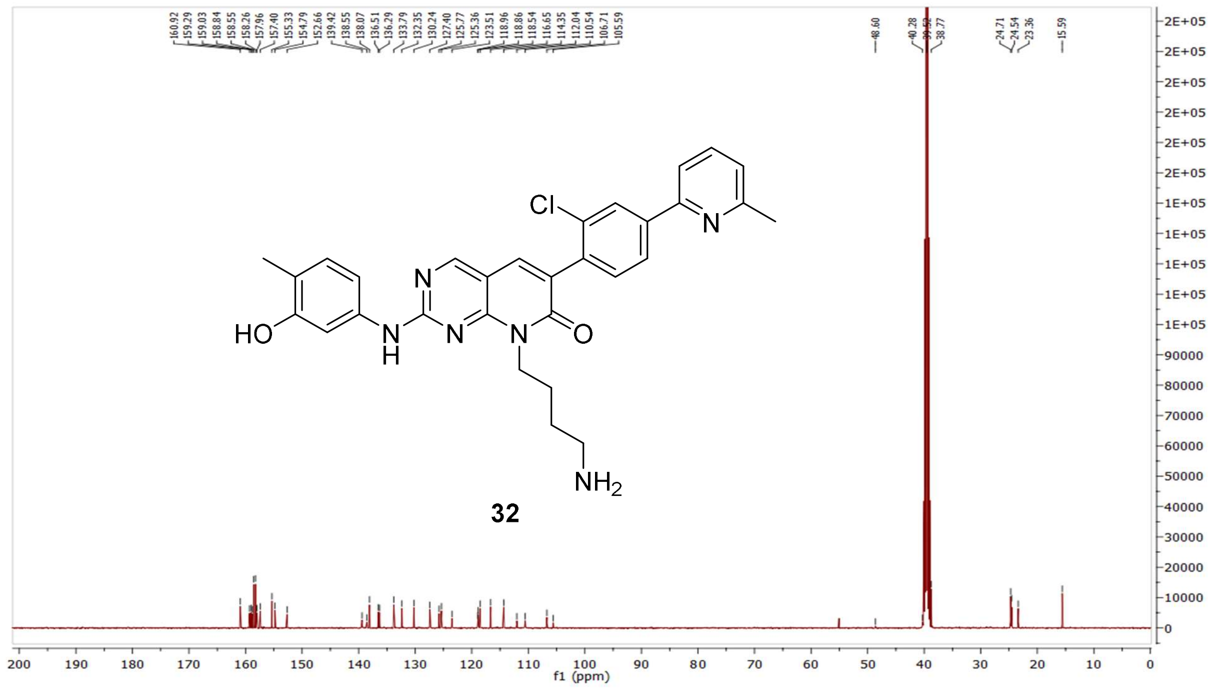

HRMS (FTMS +p MALDI) compound **32**:

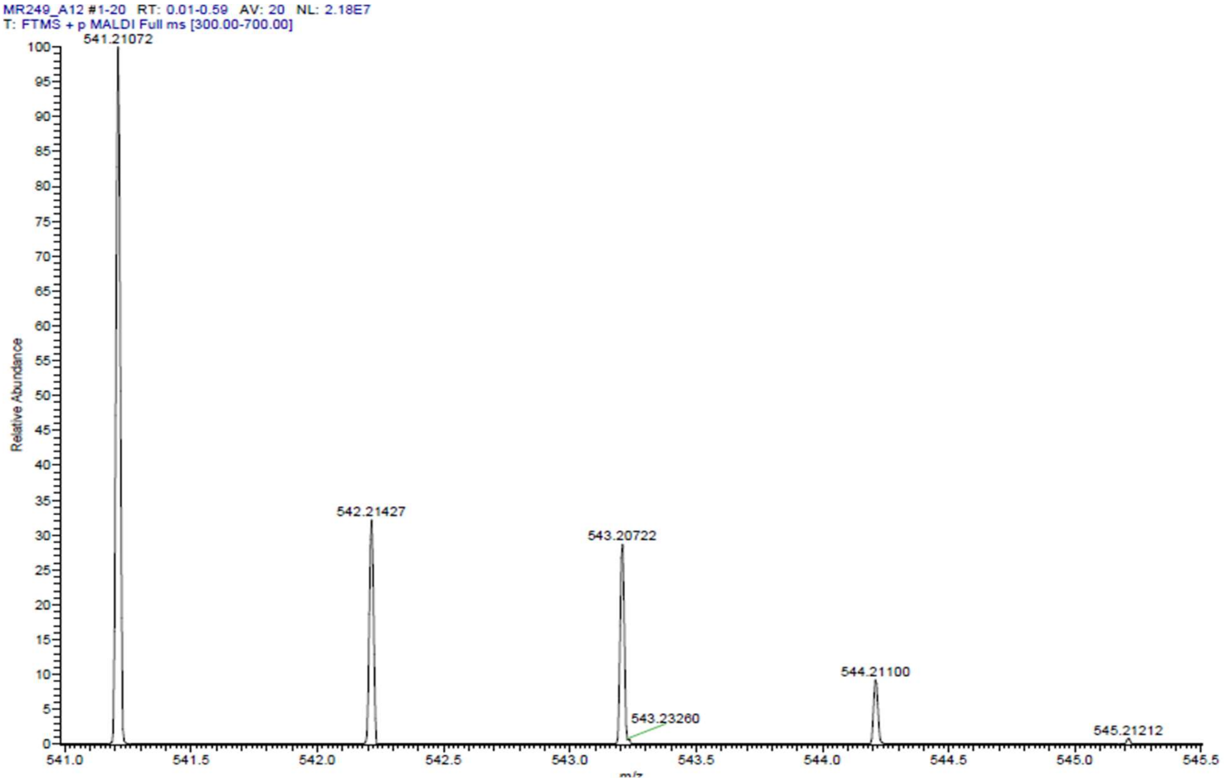

HPLC-chromatograms compound **32**:

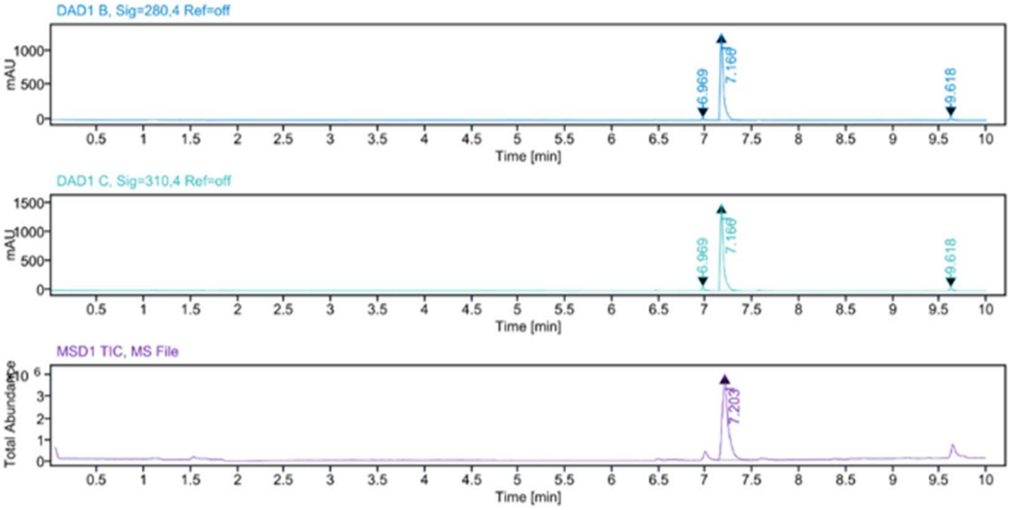


**Sample Purity**

Signal Description DAD1 B, Sig=280,4 Ref=off

| Sample Name | Name | RT | Width | Area | Area% | Height |
|---|---|---|---|---|---|---|
| MR249 | | 6.969 | 0.023 | 50.5189 | 1.60 | 32.6051 |
| MR249 | | 7.166 | 0.032 | 3008.7820 | 95.19 | 1263.8955 |
| MR249 | | 9.618 | 0.030 | 101.4152 | 3.21 | 47.3218 |

Max Area% 95.193

UV Signal Purity>95% Pass

Signal Description DAD1 C, Sig=310,4 Ref=off

| Sample Name | Name | RT | Width | Area | Area% | Height |
|---|---|---|---|---|---|---|
| MR249 | | 6.969 | 0.023 | 93.2061 | 2.47 | 60.4683 |
| MR249 | | 7.166 | 0.032 | 3583.1321 | 95.03 | 1501.9492 |
| MR249 | | 9.618 | 0.029 | 94.2872 | 2.50 | 49.9365 |

Max Area% 95.028

UV Signal Purity>95% Pass

**MSD Apex Spectra**

**RT:**

7.203

**Sample Name:**

MR249

**Signal Name:**

MSD1TIC

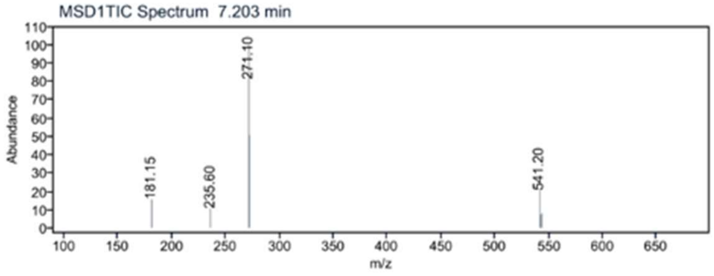

[1]H NMR compound **33**:

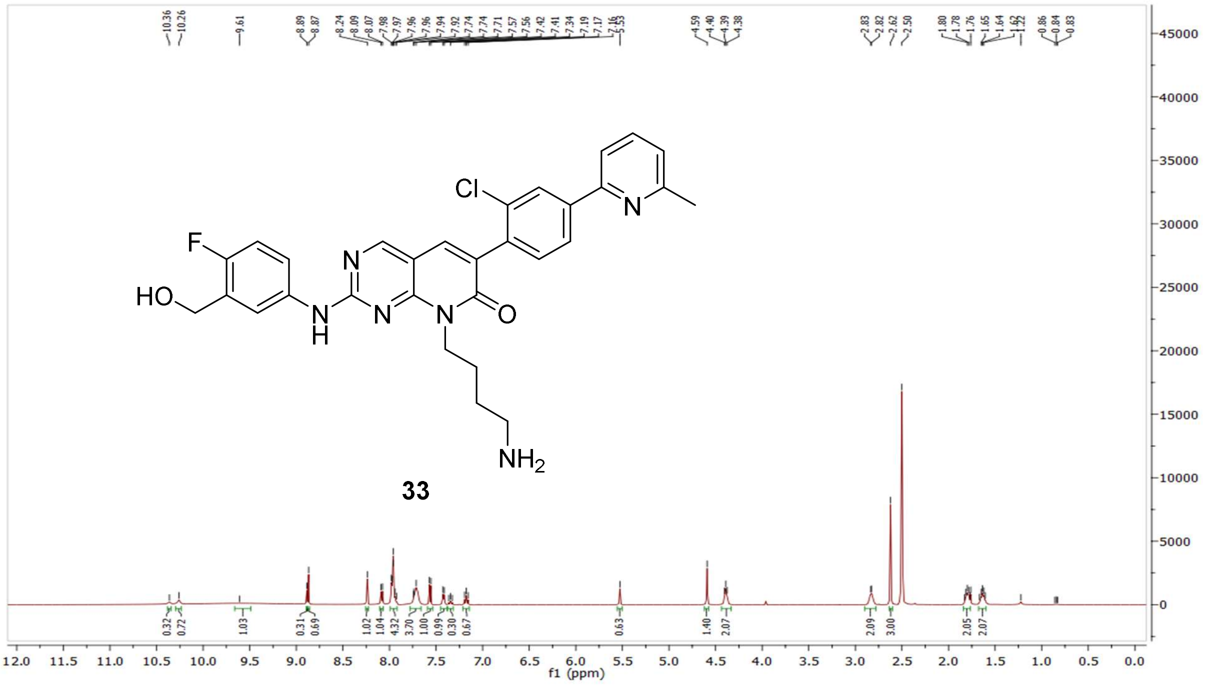

[13]C NMR compound **33:**

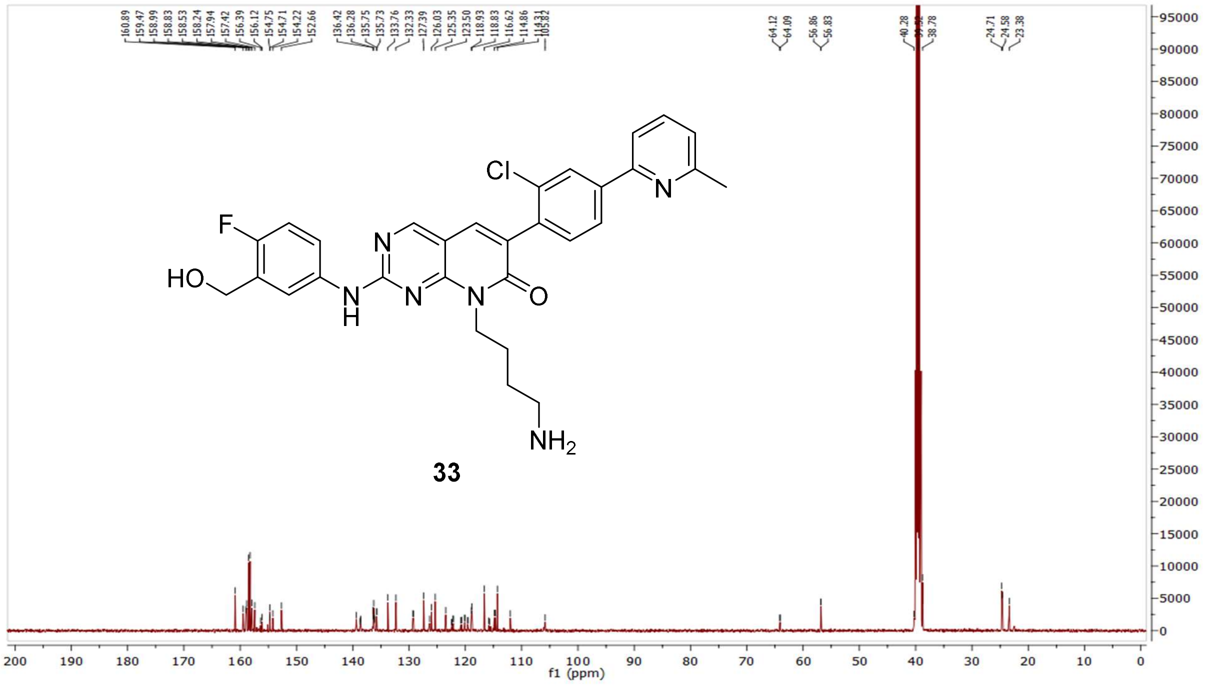

$^1$H-$^1$H COSY NMR compound **33:**

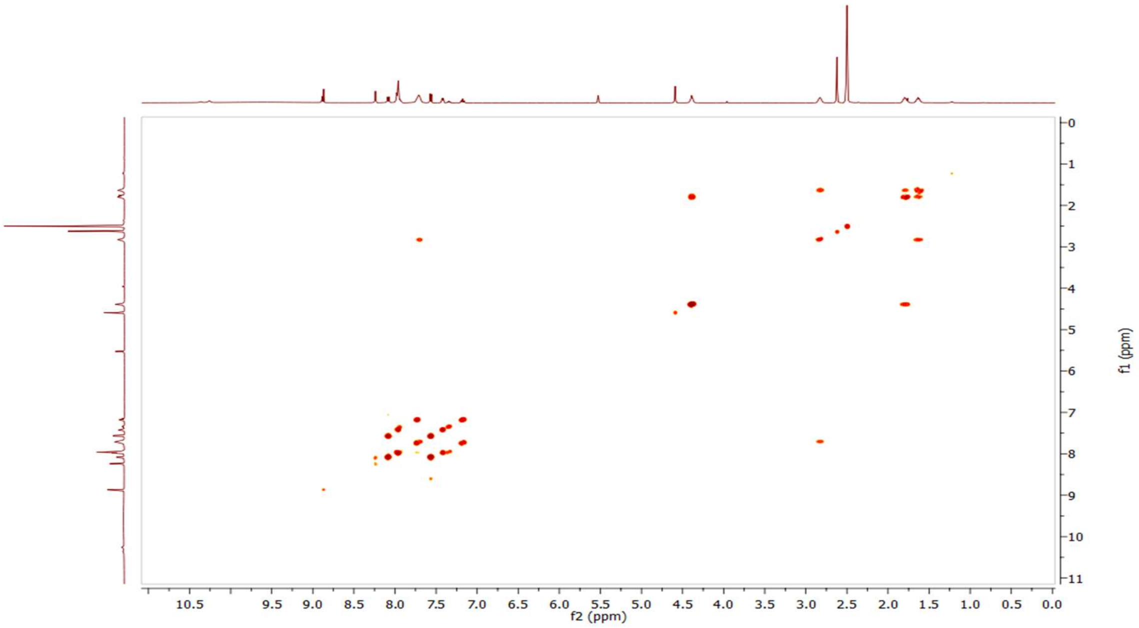

$^1$H-$^{13}$C HSQC NMR compound **33:**

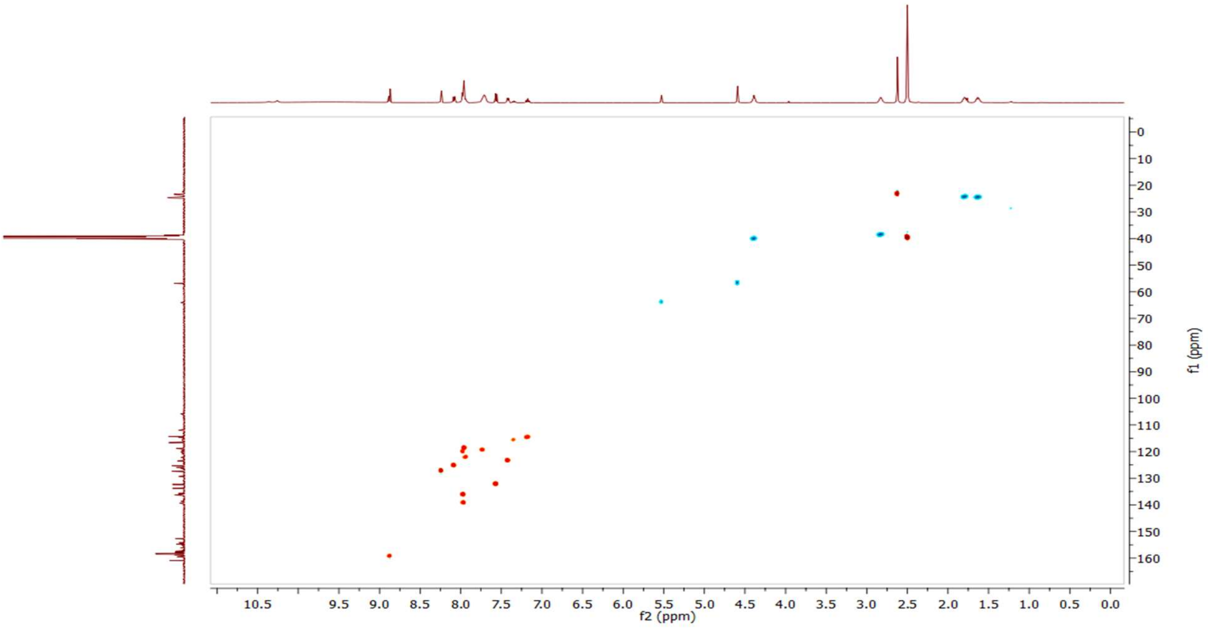

HRMS (FTMS +p MALDI) compound **33:**

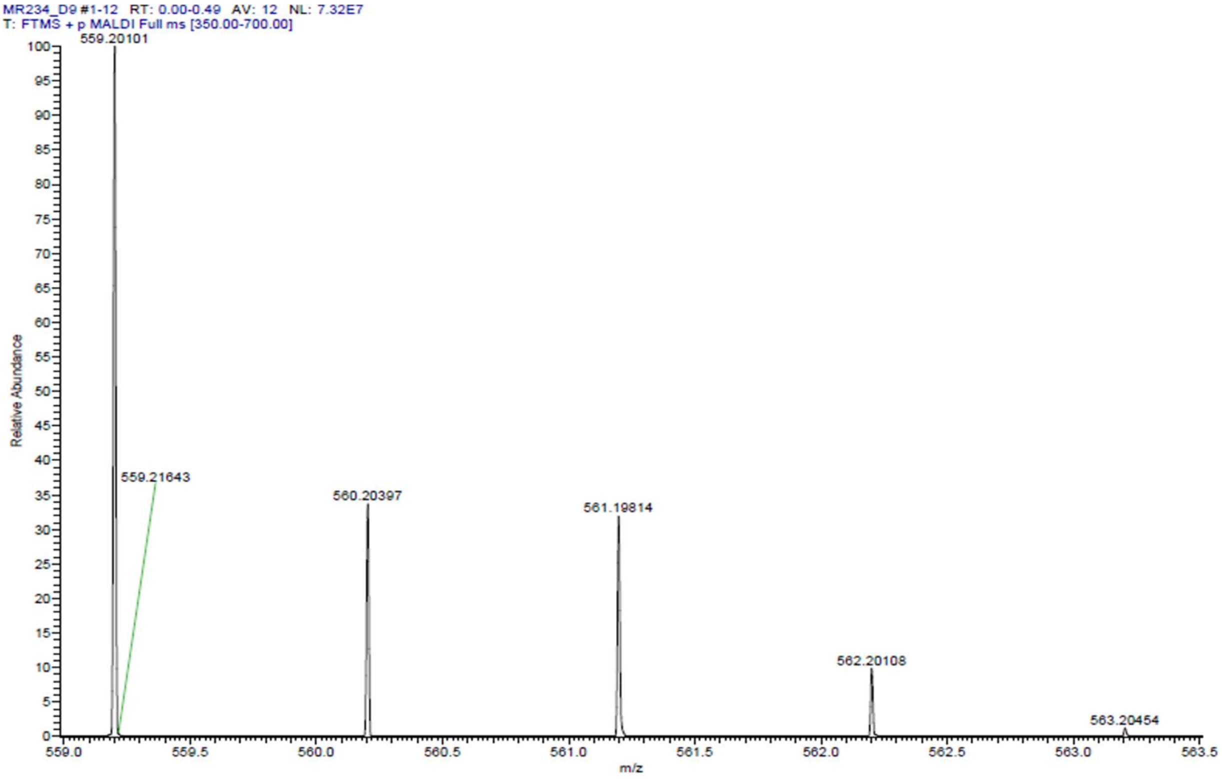

HPLC-chromatograms compound **33**:

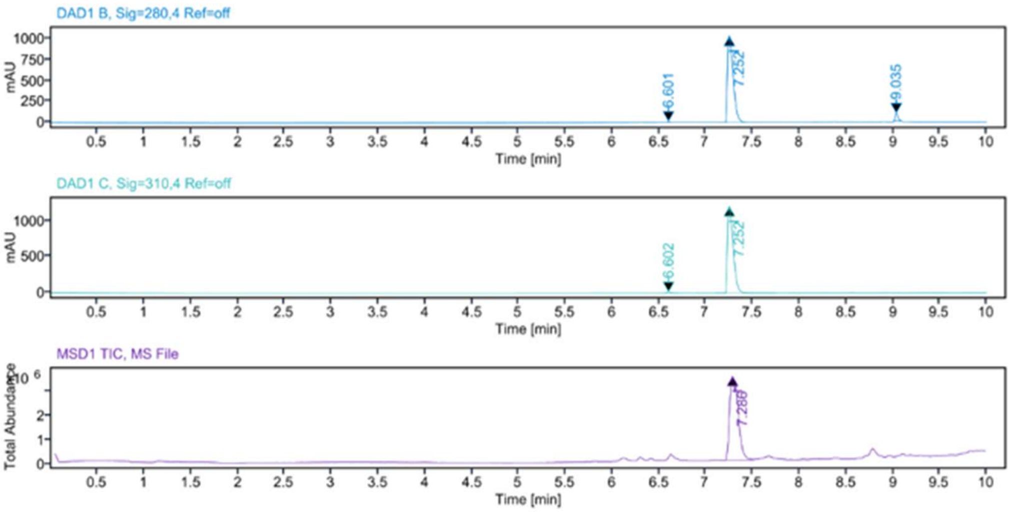


**Sample Purity**

Signal Description DAD1 B, Sig=280,4 Ref=off

| Sample Name | Name | RT | Width | Area | Area% | Height |
|---|---|---|---|---|---|---|
| MR234 F17 | | 6.601 | 0.024 | 12.4780 | 0.27 | 8.9440 |
| MR234 F17 | | 7.252 | 0.066 | 4341.9790 | 95.35 | 1030.4690 |
| MR234 F17 | | 9.035 | 0.033 | 199.1764 | 4.37 | 93.9332 |

Max Area% 95.352

UV Signal Purity>95% Pass

Signal Description DAD1 C, Sig=310,4 Ref=off

| Sample Name | Name | RT | Width | Area | Area% | Height |
|---|---|---|---|---|---|---|
| MR234 F17 | | 6.602 | 0.030 | 36.6391 | 0.72 | 16.4641 |
| MR234 F17 | | 7.252 | 0.066 | 5072.5186 | 99.28 | 1202.0598 |

Max Area% 99.283

UV Signal Purity>95% Pass

**MSD Apex Spectra**

**RT:**

7.286

**Sample Name:**

MR234 F17

**Signal Name:**

MSD1TIC

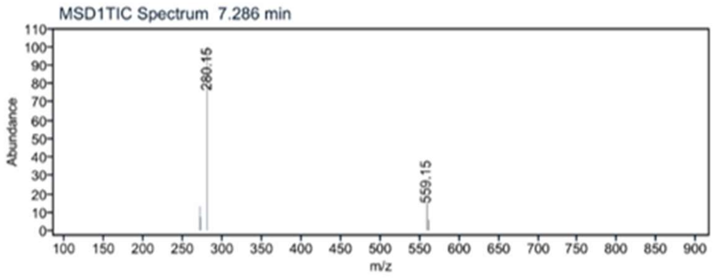

$^{1}H$ NMR compound **34**:

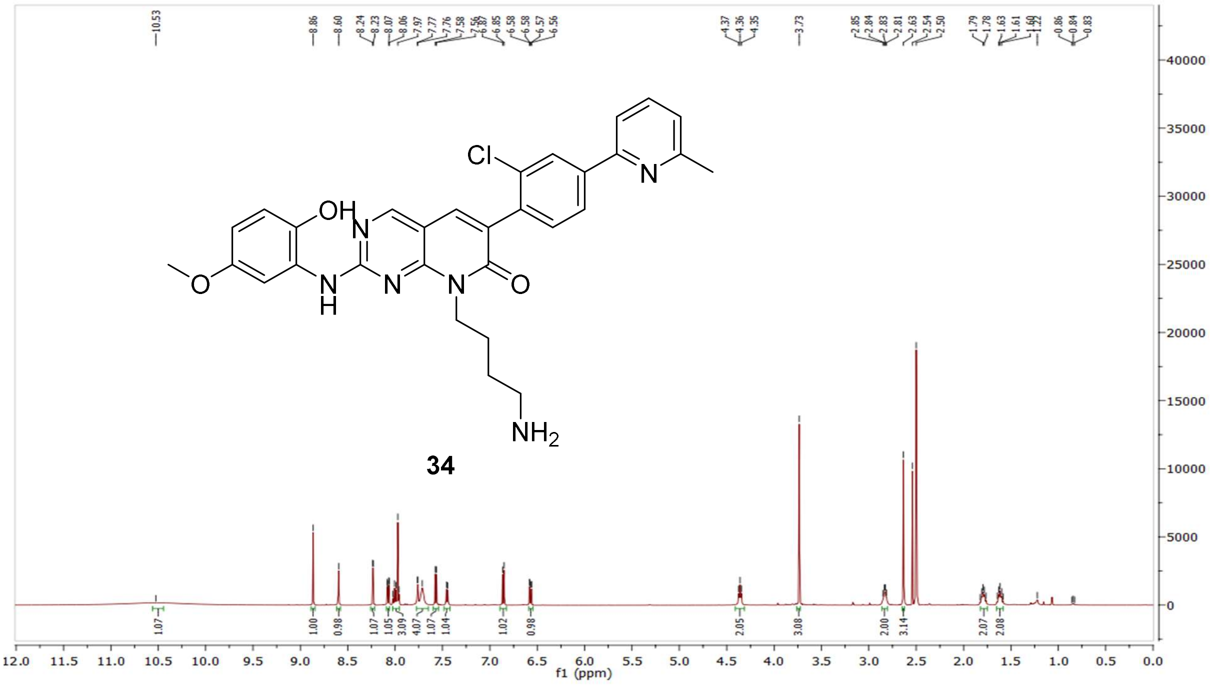


$^{13}C$ NMR compound **34:**

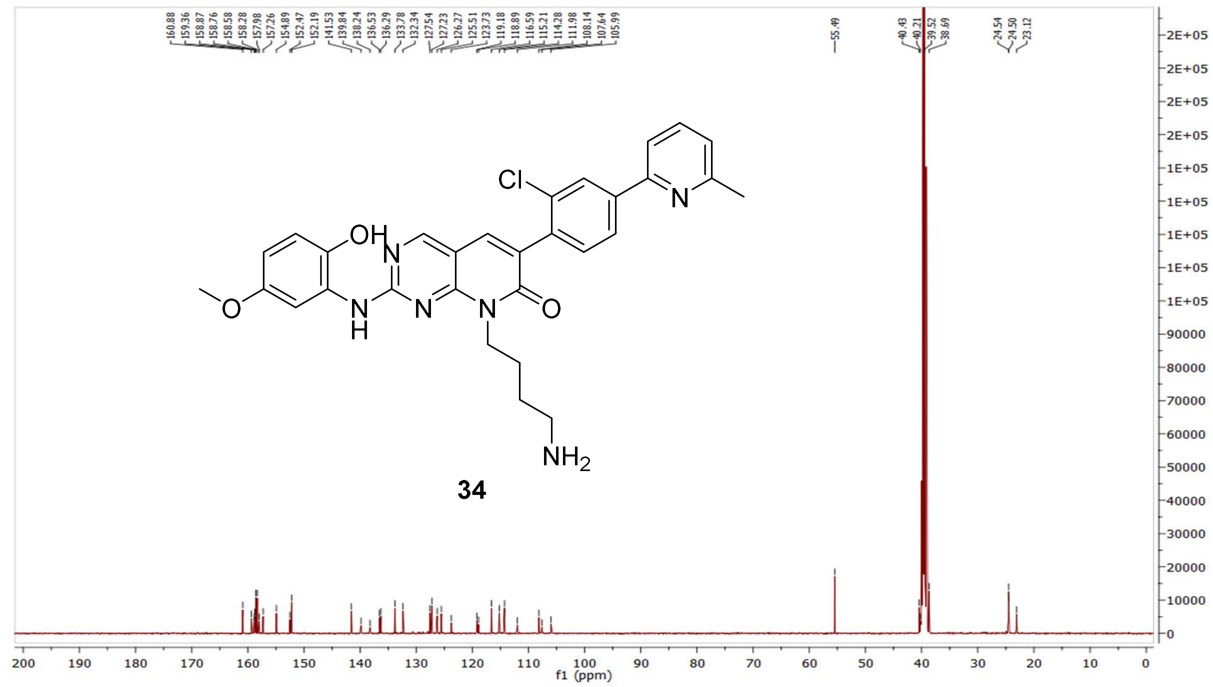

HRMS (FTMS +p MALDI) compound **34**:

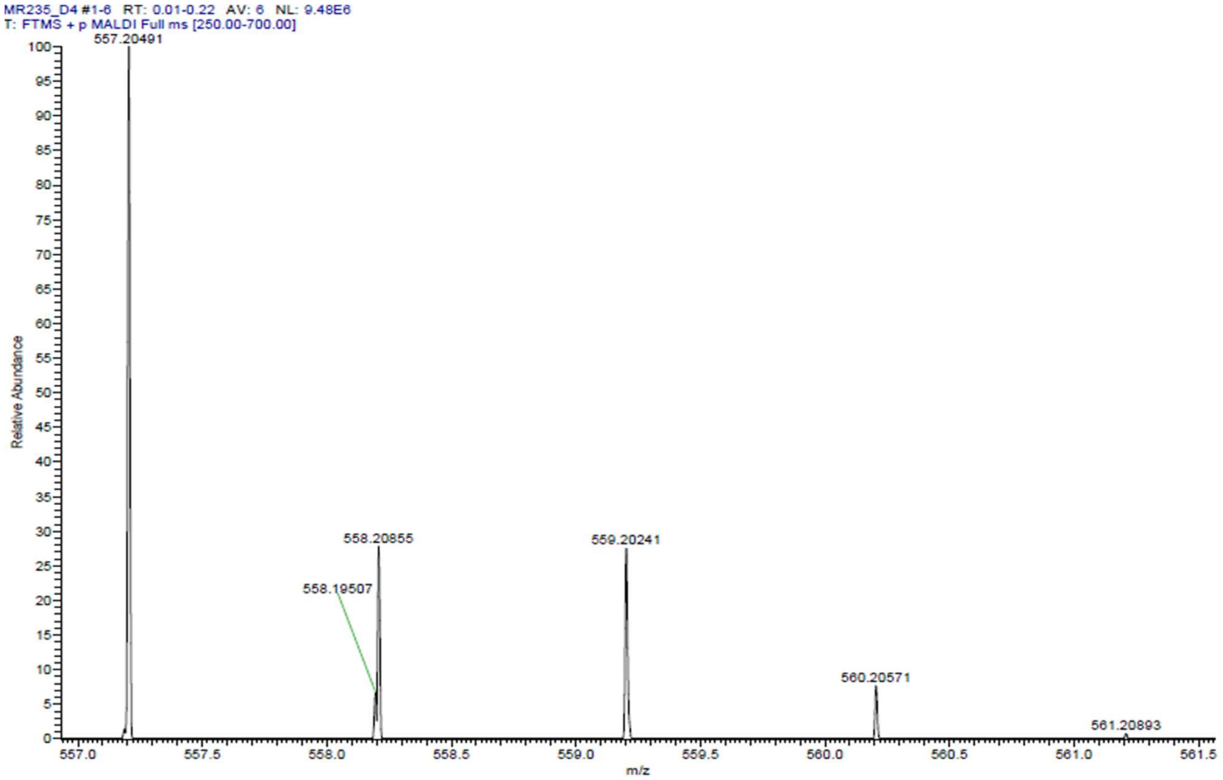

HPLC-chromatograms compound **34**:

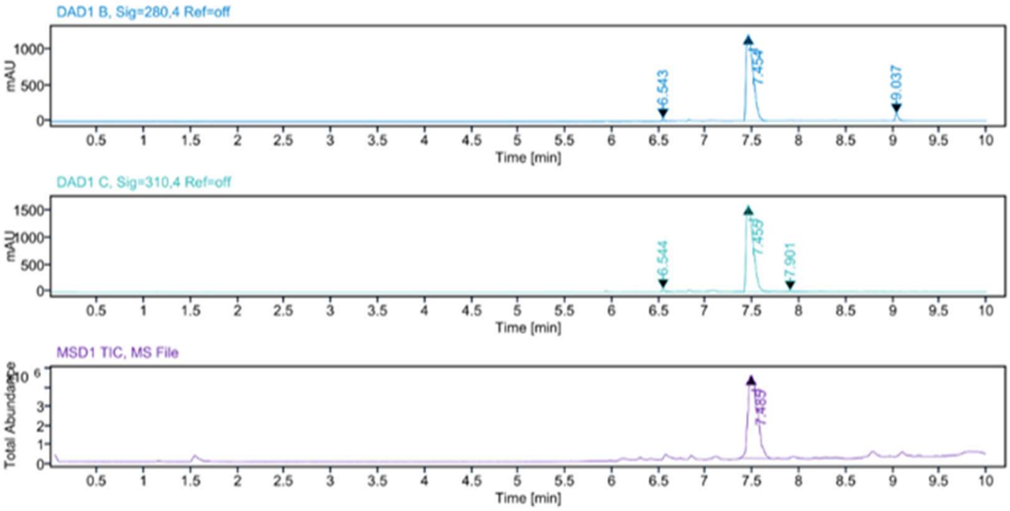


**Sample Purity**

Signal Description DAD1 B, Sig=280,4 Ref=off

| Sample Name | Name | RT | Width | Area | Area% | Height |
|---|---|---|---|---|---|---|
| MR235 | | 6.543 | 0.027 | 87.0370 | 1.32 | 35.0198 |
| MR235 | | 7.454 | 0.084 | 6262.4839 | 95.16 | 1200.6498 |
| MR235 | | 9.037 | 0.035 | 231.5378 | 3.52 | 96.2475 |

Max Area% 95.159

UV Signal Purity>95% Pass

Signal Description DAD1 C, Sig=310,4 Ref=off

| Sample Name | Name | RT | Width | Area | Area% | Height |
|---|---|---|---|---|---|---|
| MR235 | | 6.544 | 0.026 | 109.1202 | 1.28 | 47.7892 |
| MR235 | | 7.455 | 0.084 | 8360.3125 | 98.10 | 1603.0618 |
| MR235 | | 7.901 | 0.040 | 52.4847 | 0.62 | 14.6097 |

Max Area% 98.104

UV Signal Purity>95% Pass

**MSD Apex Spectra**

**RT:**

7.485

**Sample Name:**

MR235

**Signal Name:**

MSD1TIC

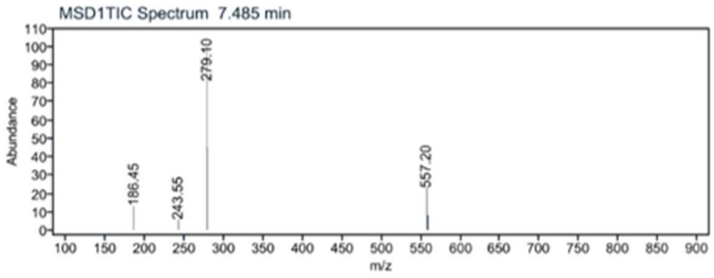

[1]H NMR compound **35**:

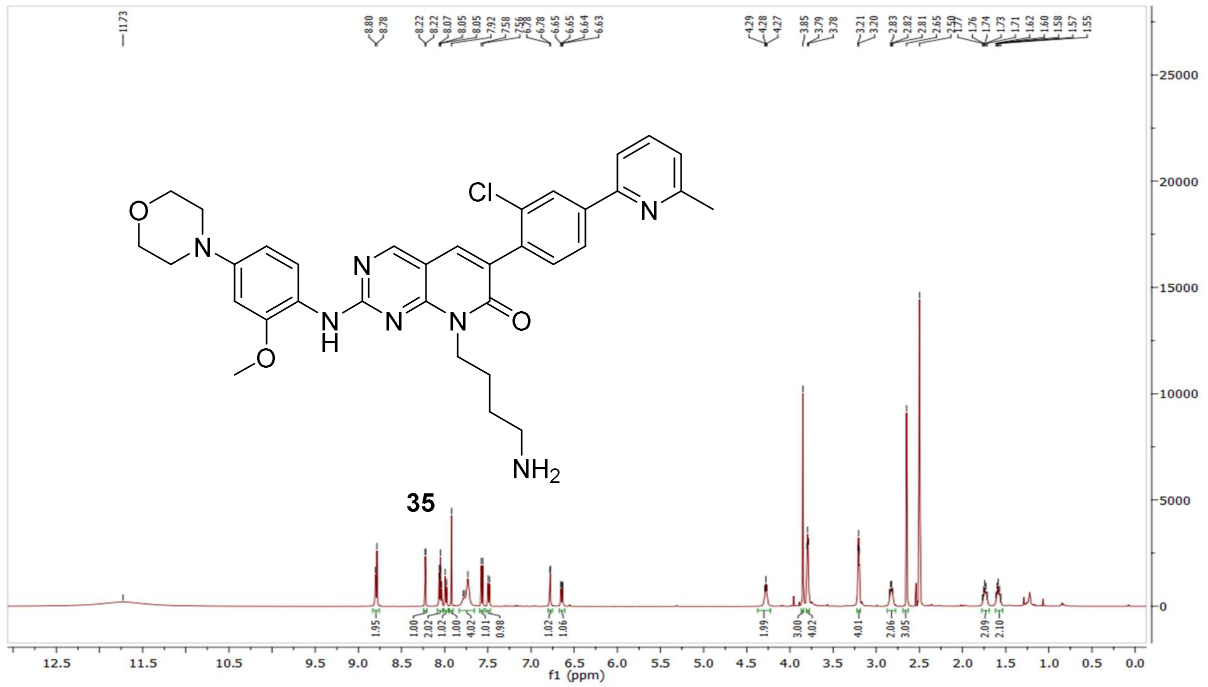

[13]C NMR compound **35:**

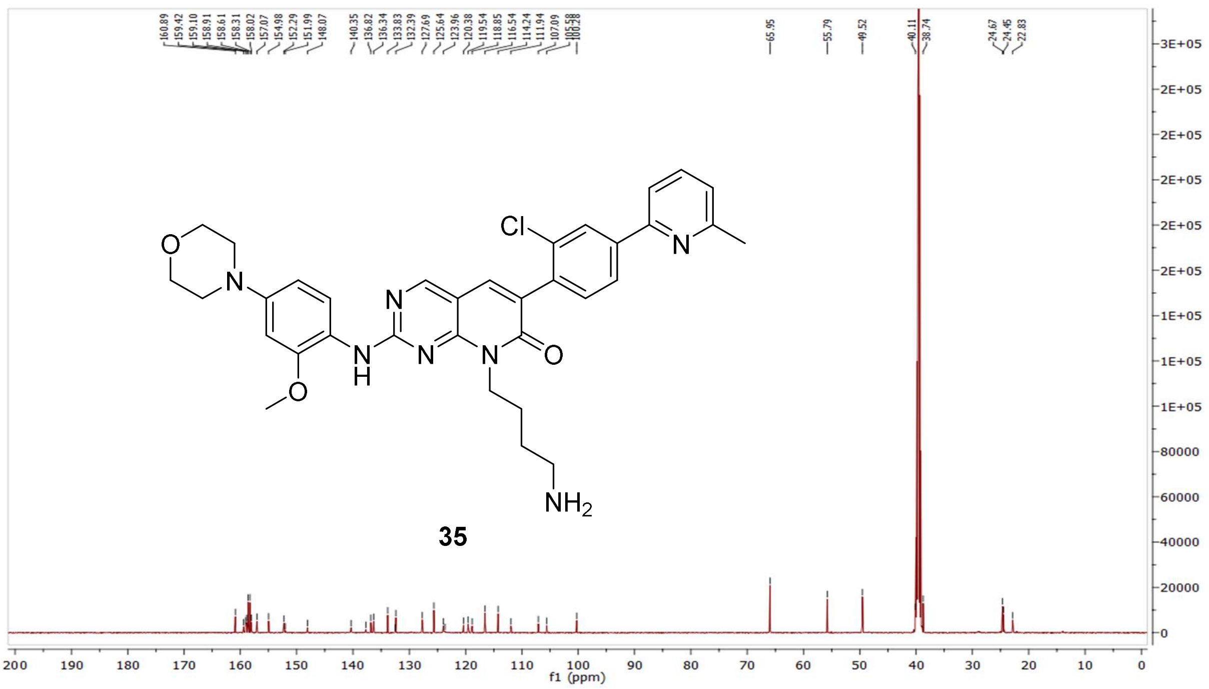

HRMS (FTMS +p MALDI) compound **35**:

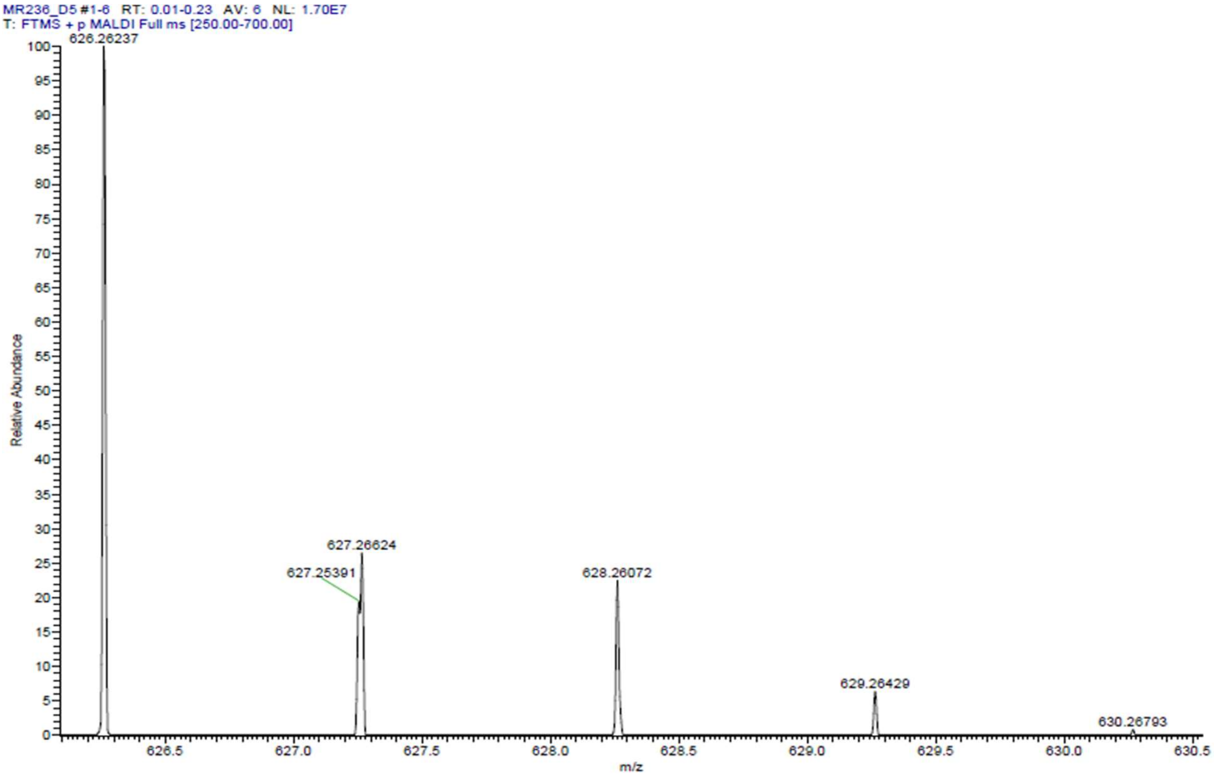

HPLC-chromatograms compound **35**:

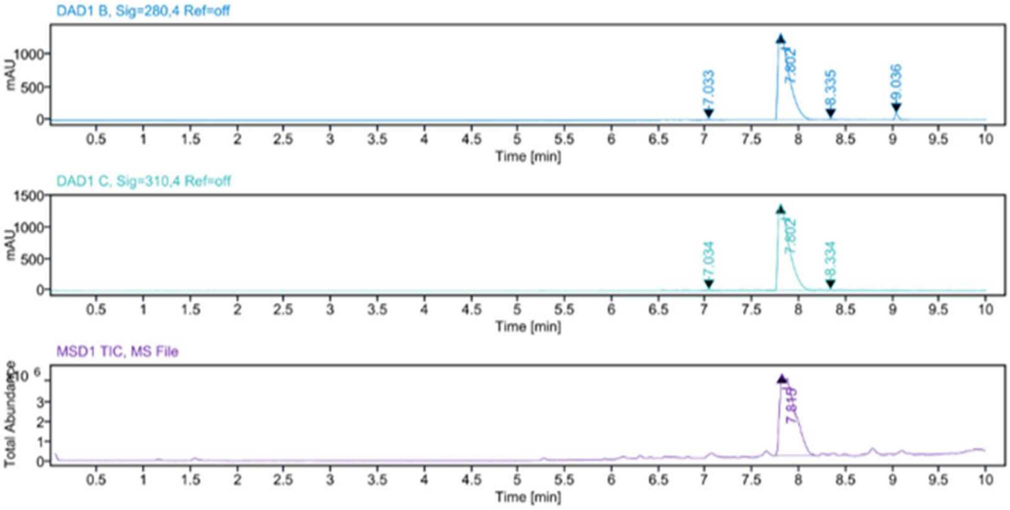


**Sample Purity**

| | |
|---|---|
| Signal Description | DAD1 B, Sig=280,4 Ref=off |

| Sample Name | Name | RT | Width | Area | Area% | Height |
|---|---|---|---|---|---|---|
| MR236 | | 7.033 | 0.069 | 77.9234 | 0.68 | 16.3943 |
| MR236 | | 7.802 | 0.127 | 11054.1250 | 96.52 | 1320.5002 |
| MR236 | | 8.335 | 0.062 | 42.6233 | 0.37 | 12.4577 |
| MR236 | | 9.036 | 0.037 | 278.0084 | 2.43 | 100.9268 |

| | |
|---|---|
| Max Area% | 96.520 |
| UV Signal Purity>95% | Pass |

| | |
|---|---|
| Signal Description | DAD1 C, Sig=310,4 Ref=off |

| Sample Name | Name | RT | Width | Area | Area% | Height |
|---|---|---|---|---|---|---|
| MR236 | | 7.034 | 0.062 | 63.9370 | 0.56 | 14.1509 |
| MR236 | | 7.802 | 0.126 | 11364.9502 | 99.23 | 1372.9155 |
| MR236 | | 8.334 | 0.037 | 24.3233 | 0.21 | 10.4064 |

| | |
|---|---|
| Max Area% | 99.229 |
| UV Signal Purity>95% | Pass |

**MSD Apex Spectra**

**RT:**

7.815

**Sample Name:**

MR236

**Signal Name:**

MSD1TIC

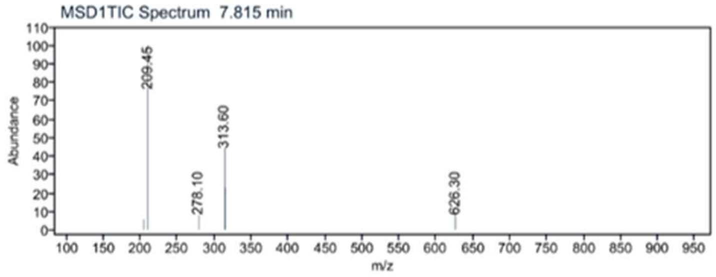


$^1$H NMR compound **36**:

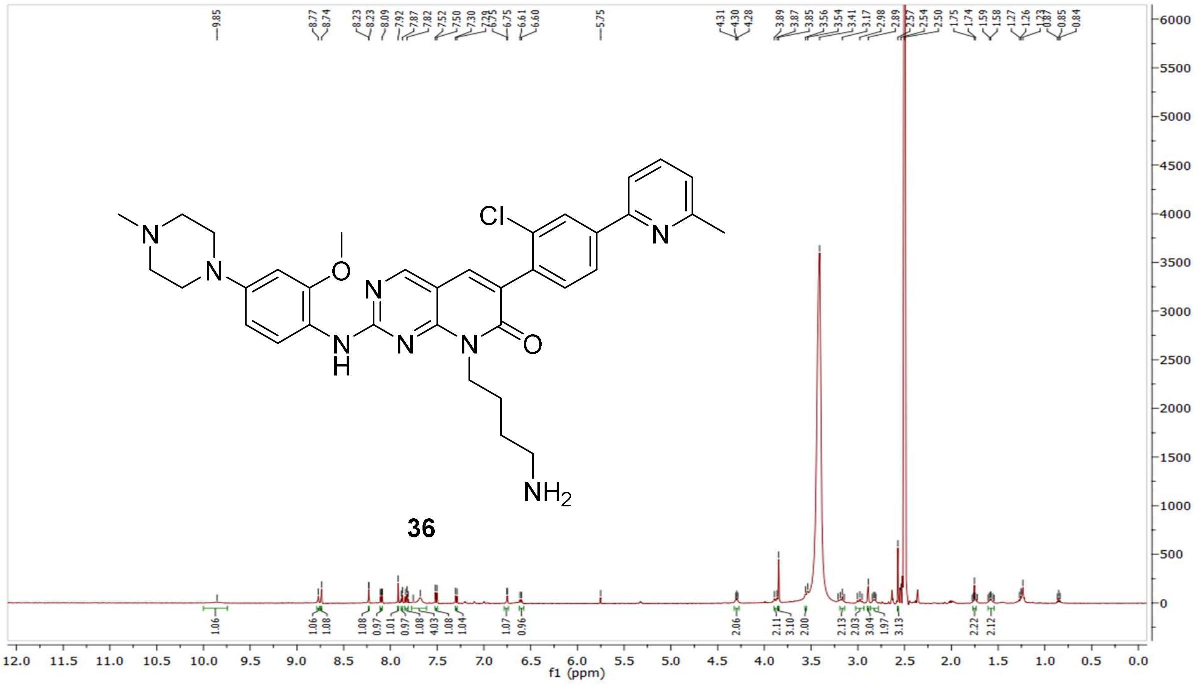

[13]C NMR compound **36:**

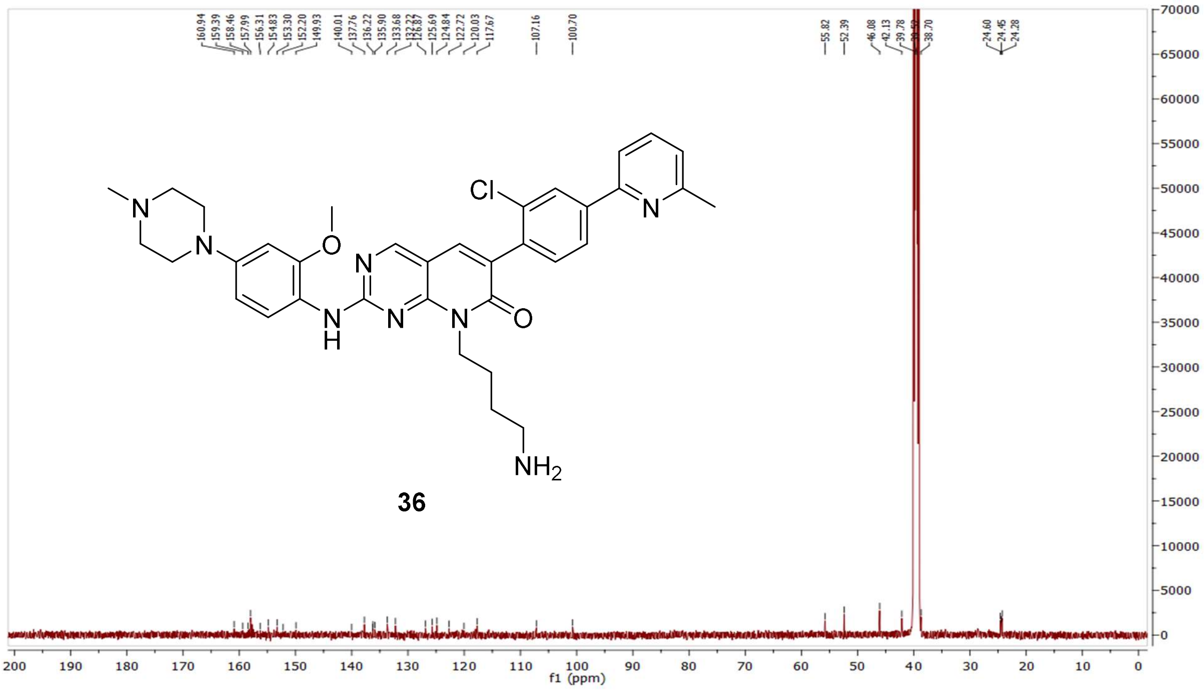


HRMS (FTMS +p MALDI) compound **36**:

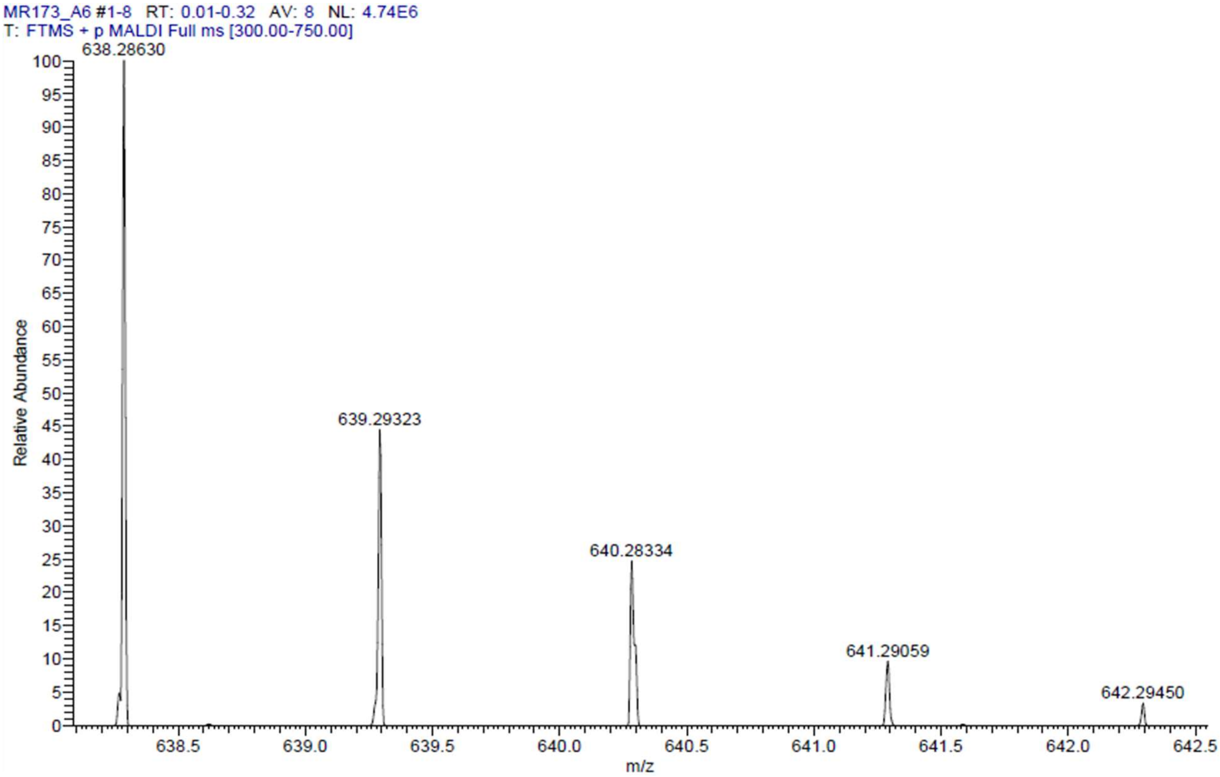

HPLC-chromatograms compound **36**:

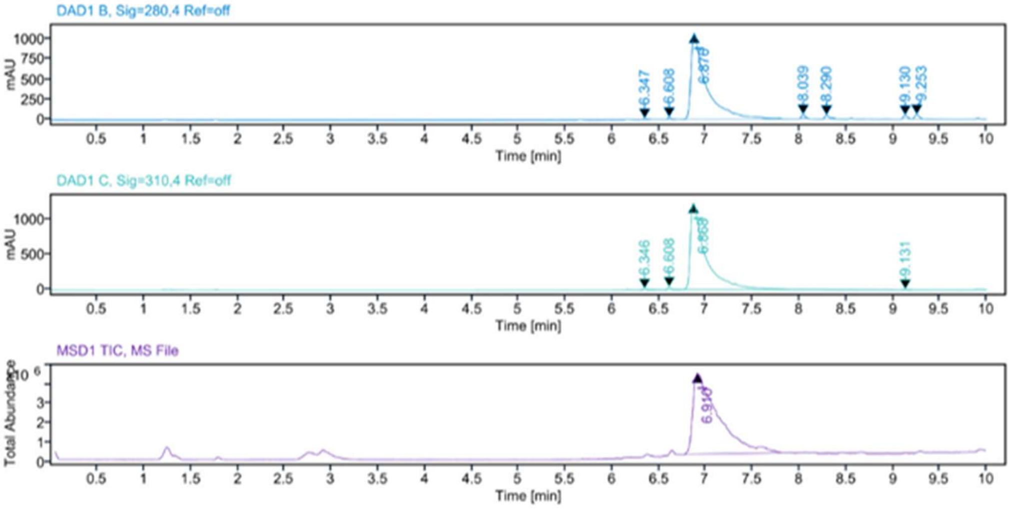


**Sample Purity**

Signal Description DAD1 B, Sig=280,4 Ref=off

| Sample Name | Name | RT | Width | Area | Area% | Height |
|---|---|---|---|---|---|---|
| MR173 F1 | | 6.347 | 0.055 | 37.8850 | 0.30 | 11.2140 |
| MR173 F1 | | 6.608 | 0.034 | 52.1530 | 0.41 | 23.9805 |
| MR173 F1 | | 6.876 | 0.131 | 11970.4082 | 95.00 | 1061.7455 |
| MR173 F1 | | 8.039 | 0.044 | 112.9122 | 0.90 | 43.5481 |
| MR173 F1 | | 8.290 | 0.045 | 147.4747 | 1.17 | 41.4767 |
| MR173 F1 | | 9.130 | 0.050 | 116.8831 | 0.93 | 38.8746 |
| MR173 F1 | | 9.253 | 0.053 | 162.2526 | 1.29 | 50.1435 |

Max Area% 95.003

UV Signal Purity>95% Pass

Signal Description DAD1 C, Sig=310,4 Ref=off

| Sample Name | Name | RT | Width | Area | Area% | Height |
|---|---|---|---|---|---|---|
| MR173 F1 | | 6.346 | 0.076 | 80.9464 | 0.59 | 16.8841 |
| MR173 F1 | | 6.608 | 0.037 | 100.1770 | 0.73 | 38.4242 |
| MR173 F1 | | 6.868 | 0.131 | 13601.4395 | 98.50 | 1236.4124 |
| MR173 F1 | | 9.131 | 0.065 | 25.7355 | 0.19 | 5.9878 |

Max Area% 98.502

UV Signal Purity>95% Pass

**MSD Apex Spectra**

**RT:**

6.910

**Sample Name:**

MR173 F1

**Signal Name:**

MSD1TIC

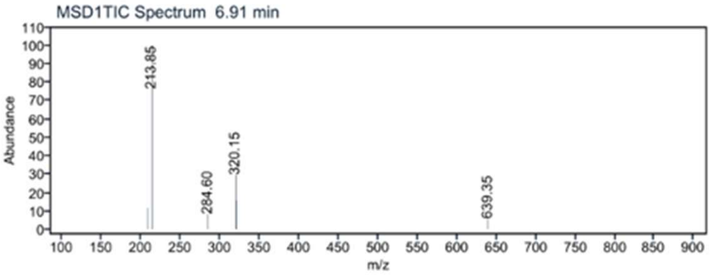
